# A phylogenetic approach reveals evolutionary aspects and novel genes of bradyzoite conversion in *Toxoplasma gondii*

**DOI:** 10.64898/2026.04.20.719551

**Authors:** Anupama C. A., Roopam Upadhyay, Swati Patankar

**Affiliations:** Department of Biosciences & Bioengineering, Indian Institute of Technology Bombay, Mumbai, India; International Finance Corporation, Bandra Kurla Complex, Mumbai, India

**Keywords:** *Toxoplasma gondii*, bradyzoite differentiation, phylogeny

## Abstract

*Toxoplasma gondii* is a widespread human pathogen that has multiple, clinically relevant stages in its complex life cycle, including fast-replicating tachyzoites and latent bradyzoites. Bradyzoite differentiation is triggered by stress responses that lead to changes in transcription, translation, and metabolism. Two aspects of this process are addressed in this report: first, whether proteins that play roles in bradyzoite differentiation are specific to *T. gondii* and other bradyzoite-forming parasites of the *Sarcocystidae* family, and second, whether new bradyzoite differentiation proteins can be identified in *T. gondii*. To answer these questions, a phylogenetic approach was used, comparing proteomes of select members of the *Sarcocystidae* family that form morphologically different bradyzoite cysts and members of the *Eimeriidae* family that do not form cysts. This approach resulted in 8 distinct clusters of *T. gondii* proteins that reflected different conservation patterns; for example, one cluster showed conservation among all organisms, while another showed conservation in bradyzoite cyst-forming organisms. Known *T. gondii* proteins involved in bradyzoite differentiation were found in all clusters, indicating that this process uses both highly conserved pathways as well as bradyzoite-specific pathways. Importantly, the cluster containing proteins that are conserved in bradyzoite-forming organisms contained several known regulators of bradyzoites, and will be a source for identifying novel *T. gondii* proteins that are involved in bradyzoite differentiation.

## Introduction

*Toxoplasma gondii* is an obligate intracellular parasite of the phylum Apicomplexa that infects warm-blooded animals, including economically important livestock (Jacobs, 1957). A third of the human population has been exposed to *T. gondii*, resulting in the zoonotic disease toxoplasmosis in susceptible individuals (Montoya & Liesenfeld, 2004).

*T. gondii*, like other apicomplexan parasites, has a complex life cycle involving multiple hosts and life cycle stages (Frenkel et al., 1970; Dubey & Frenkel, 1972; Dubey et al., 1998; Attias et al., 2020). Intermediary hosts like humans harbor the asexual stages, consisting of fast-replicating tachyzoites responsible for acute infections that can convert into latent cyst-forming bradyzoites that cause chronic infection (Dubey & Frenkel, 1972). Bradyzoite cysts persist in the brain and muscles, where they cause recrudescence of the disease in immunocompromised conditions (Jeffers et al., 2018). The thick cyst wall shields bradyzoites from drugs and host immune responses (Dubey et al., 1998; Van Der Ven et al., 1996; Villena et al., 1998), therefore, current therapeutic regimens, including pyrimethamine (Dunay et al., 2018) and sulphonamides, effectively target tachyzoites but fail to eliminate bradyzoite cysts. Therefore, understanding the molecular mechanisms governing tachyzoite-to-bradyzoite conversion is critical.

Bradyzoite differentiation in *T. gondii* is the outcome of stress responses (reviewed in Jeffers et al., 2018) to diverse environmental stressors, including the host immune response (Silva et al., 1998), nutritional starvation (Fox et al., 2004), and heat shock (Soete et al., 1993). In laboratory conditions, alkaline pH also induces bradyzoite differentiation (Bohne et al., 1995; Mayoral et al., 2020). Like other eukaryotes, *T. gondii* initiates an integrated stress response pathway, characterized by the phosphorylation of the α subunit of eukaryotic initiation factor-2 (eIF2) at serine 51 by specialized kinases, each responding to a specific stress. For example, eIF2α kinase A responds to endoplasmic reticulum (ER) stress (Augusto et al., 2018), eIF2α kinase B responds to alkaline and oxidative stress (Augusto et al., 2021), eIF2α kinase C responds to amino acid starvation (Konrad et al., 2014) and eIF2α kinase D responds to extra-cellular stress (Konrad et al., 2011). The phosphorylation of eIF2α leads to a global reduction of cap-dependent translation while selectively allowing the translation of stress-responsive transcripts like catalase and thioredoxin reductase for redox stress (Kalinin et al., 2023; Starck et al., 2016; Kwok et al., 2004), General Control Non-derepressible-5 (GCN-5) for alkaline stress (Naguleswaran et al., 2010), and calreticulin for ER stress (Joyce et al., 2013).

Regulators of bradyzoite conversion have been identified, including Bradyzoite Formation Deficient 1 (BFD-1), Bradyzoite Formation Deficient 2 (BFD-2) and eukaryotic initiation factor 1.2 (eIF1.2) (Licon et al., 2023; Waldman et al., 2020; F. Wang et al., 2024). These regulatory proteins further initiate transcription and translation of other bradyzoite-specific genes like bradyzoite antigen-1 (*bag-1*), lactate dehydrogenase-2 (*ldh-2*) and enolase-1 (*eno-1*) (Bohne et al., 1995b; Dzierszinski et al., 2001; Waldman et al., 2020; F. Wang et al., 2024).

Apart from the stress response pathway, cell cycle regulators (Naumov et al., 2022; Radke et al., 2018) and amylopectin metabolism enzymes (Lyu et al., 2021; Uboldi et al., 2015; J.-L. Wang et al., 2022; Yang et al., 2022) have also been implicated in bradyzoite formation and maintenance. Although many players in the pathway of differentiation are known in *T. gondii*, cell surface receptors and signal transduction pathways are still unclear. Further, while the stress response pathway is conserved among eukaryotes, it is unclear whether bradyzoite regulators evolved as specific effectors of differentiation in *T. gondii*. Here, we use an evolutionary approach to address these questions.

*T. gondii* is a member of the *Sarcocystidae* family, comprising of the genera *Toxoplasma*, *Hammondia*, *Neospora*, *Besnoitia*, *Cystoisospora*, and *Sarcocystis*. Members of this family establish chronic infections by forming tissue cysts, a conserved survival strategy across multiple genera. However, only the first four genera, classified into the subfamily *Toxoplasmatinae,* form bradyzoite cysts. In contrast, the *Cystoisopora* genus forms monozoic tissue cysts, containing a single parasite called a zoite (Lindsay et al., 2014; Shrestha et al., 2015), while *Sarcocystis* forms septate tissue cysts containing non-infectious stages called metrocytes along with infectious bradyzoites in the centre of the cysts (Fayer, 2004; Mehlhorn & Frenkel, 1980). These distinct cysts suggest divergent strategies for chronic infection, making comparative phylogenetic analysis a valuable approach for understanding the evolution of known bradyzoite-specific factors and identifying new ones in *T. gondii*.

We compared the *T. gondii* proteome with those of other *Sarcocystidae* members, and found the best hits using bit score, a sequence conservation metric, followed by clustering of the best hits using unsupervised clustering algorithms. This analysis partitioned the *T. gondii* proteome into distinct clusters: proteins that may have evolved based on phylogenetic relationships, conserved proteins indicative of housekeeping functions, and notably, a class of proteins conserved among bradyzoite cyst-forming organisms. Bradyzoite regulators were found in all these categories, suggesting that *T. gondii* uses ancient pathways, while evolving specialized proteins for bradyzoite differentiation. The *T. gondii* proteins that are conserved among bradyzoite cyst-forming organisms represent novel candidates that may be involved in stage differentiation; these encompass diverse functional categories, including cell surface receptors, signalling kinases, transcriptional regulators, translation factors, and nucleic acid-binding proteins. This work uncovers novel molecular mechanisms underlying tachyzoite-to-bradyzoite conversion in *T. gondii* and throws the field open for future studies on validation of their function.

## Materials and Methods

### Phylogenetic tree

Sequence IDs for the 18s rRNA sequences were taken for the selected organisms: *Toxoplasma gondii, Hammondia hammondi, Neospora caninum, Besnoitia besnoiti*, *Cystoisospora cuis*, *Sarcocystis neurona* and *Eimeria falciformis* (Oyarzún-Ruiz et al., 2023). The FASTA files of these sequences were downloaded from the NCBI database and sequences aligned using the Multiple Sequence Comparison by Log-expectation (MUSCLE) within MEGA 11 software(Tamura et al., 2021). The phylogenetic tree was made using the Maximum Likelihood (ML) method using the General Time Reversal (GTR) substitution model (Waddell & Steel, 1997) with bootstrap value of 500 to assess the tree robustness. Initial tree construction for the heuristic search was performed using Neighbour-Joining and BioNJ algorithms.

### Standalone BLAST of proteomes and selection of best alignments

The datasets containing the FASTA sequences of annotated proteins in *Toxoplasma gondii*, *Hammondia hammondi*, *Neospora caninum*, *Besnoitia besnoiti*, *Cystoisospora suis*, *Sarcocystis neurona*, *Eimeria falciformis* and *Cyclospora cayetanensis* were downloaded from ToxoDB (Version 68)(Alvarez-Jarreta et al., 2024). To identify the protein homologs between *T. gondii* and the selected organisms, a pairwise comparison was done using NCBI+ standalone pBLAST. Each *T. gondii* protein was aligned against the complete proteome of each selected organism. To capture all potential homologs, no threshold filters on the parameters like E-value, percentage identity, percentage alignment, or bit score were imposed.

### Ortholog-based validation of bit score derived homologs

To validate that the bit score derived homolog pairs represent orthologs, an orthology analysis was performed using OrthoVenn3 (Sun et al., 2023). The FASTA files consisting the protein sequences of selected organisms were uploaded to OrthoVenn3. The orthologous analysis used default settings of the OrthoMCL algorithm with an E-value threshold of 10^-5^ and an inflation value of 1.5. For each organism, the overlap between OrthoVenn3-derived ortholog pairs and bit score derived homolog pairs was calculated. Overlap was quantified as:

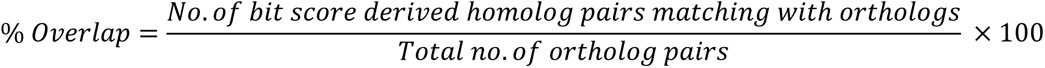

### Assigning a decile score to bit scores

The highest bit score for each *T. gondii* protein in each organism was obtained. If a protein showed no hit in a particular organism, it was assigned a bit score of 0; thus, these scores ranged from 0 to a maximum of approximately 24,000. To minimize skewness in the data distribution, bit scores were binned into categories (decile score) according to the decile range into which they fell. This created an 8206 x 7 matrix consisting of alignments of each *T. gondii* protein with the selected organisms and their corresponding decile score.

### Clustering

To identify conservation patterns among the selected organisms, a two-stage clustering approach was implemented. First, the optimal number of clusters (K) was determined by evaluating K in the range of 2 to 35 using multiple approaches, including the Elbow Method, Silhouette Score, and Gap Statistic. A K-means clustering was then performed (Jain & Dubes, 1988; Li & Wu, 2012) via the “kmeans( )” function in R with 500 bootstrap replicates. Among the bootstrap iterations, the clusters yielding the lowest Within-Cluster Sum of Squares (WCSS) value were selected to ensure maximum cluster compactness (Tibshirani et al., 2001). This approach identifies the minimum number of meaningful clusters beyond which additional clustering does not provide any significant insight.

In the second stage, the initial K-means clusters were further consolidated into higher-order meta-clusters using hierarchical clustering (Dasgupta & Long, 2005). Clustering was performed on the centroids of the K-means clusters using Euclidean distance and Ward’s minimum variance method (Ward.D2) (Ward Jr., 1963), implemented via the “hclust( )” function in R. The Dynamic Tree Cut algorithm was applied to identify the minimum number of clusters that could be derived while preserving meaningful structure in the data.

### COG Analysis

To assign proteins to broad functional categories, Cluster of Orthologous Groups (COG) analysis was performed using the Galaxy eggNOG-mapper (version 2.1.13) (Tatusov et al., 1997). Default parameters were used: eggNOG database version 5.0.2, BLOSUM62 scoring matrix, minimum query coverage of 0.001, and minimum E-value threshold of 0.001. Each gene was assigned to a single COG functional category (A–Z) or categorized as “nil” where no COG assignment was available.

The representation of individual COG categories within each cluster was assessed by comparing their frequency within the cluster to their background frequency across all the clusters. Enrichment analysis identified COG categories significantly overrepresented or underrepresented within each cluster. COG enrichment was analysed using a multivariate distribution analysis, calculated using an elastic net regression model with the R command “cv.glmnet( )”. Odds ratios were calculated to quantify the likelihood of observing proteins belonging to a certain COG category within a specific cluster relative to the background. Statistical significance was determined using Wald test with a p-value threshold of 0.05. The COG categories and their odds ratio for each cluster were plotted in a heat map using GraphPad Prism ver 8.4.3.

## Results and Discussion

This section describes the development and implementation of a comparative proteomics pipeline to identify conservation patterns of *T. gondii* proteins with proteins of the cyst-forming *Sarcocystidae* family, and two organisms from an *Eimeriidae* outgroup. The broad outcome was to make clusters that show different conservation patterns across these organisms. The *T. gondii* proteins in each cluster could then be further analysed for biological insights, especially regarding bradyzoite differentiation. Below, we describe each analytical step in detail, including the rationale for key methodological decisions and validation of our approach.

### Selection of organisms and pipeline development

*Toxoplasma* belongs to the family *Sarcocystidae*, consisting of tissue-cyst forming organisms with a wide host range. This family includes the genera *Sarcocystis, Neospora, Toxoplasma, Hammondia, Besnoitia, and Cystoisospora.* Among these, *Toxoplasma, Hammondia, Besnoitia, and Neospora* form tissue cysts containing bradyzoites, while *Cystoisospora* and *Sarcocystis* tissue cysts are morphologically distinct from bradyzoite cysts.

As the obligate parasitic nature of these organisms makes them challenging to culture, we selected representative species of each genus based on the availability of proteome datasets. These were *Hammondia hammondi*, *Toxoplasma gondii*, *Besnoitia besnoiti*, *Neospora caninum*, *Cystoisospora suis*, and *Sarcocystis neurona*. *Eimeria falciformis* and *Cyclospora cayetanensis* from the family *Eimeriidae* (subclass *Coccidiasina*) were included as negative controls, as these organisms do not form tissue cysts. Among available *Eimeriidae* proteomes, *E. falciformis* and *C. cayetanensis* were selected as extensively studied representatives.

To confirm evolutionary relationships between the selected organisms, a phylogenetic tree was constructed using 18S rRNA sequences (Figure 1a). As expected, bradyzoite cyst-forming organisms *T. gondii*, *H. hammondi*, and *N. caninum* formed a tight clade reflecting their close evolutionary relationships, while *B. besnoiti* occupied a separate branch. *C. suis* and *S. neurona* were positioned between the bradyzoite cyst-forming organisms and the outgroup formed by *E. falciformis* and *C. cayetanensis*.

**Figure 1:**
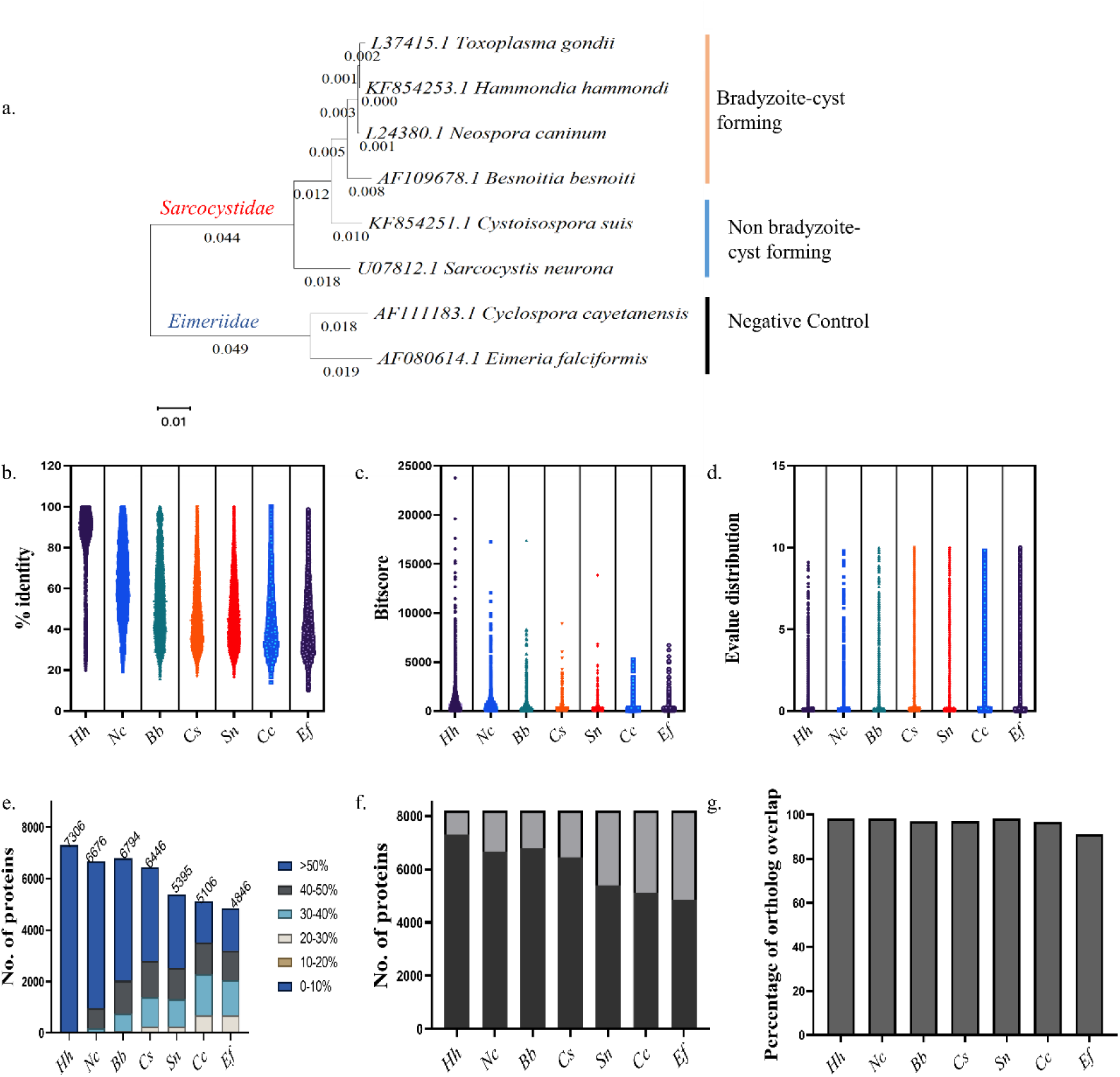
Phylogenetic relationships and parameter selection for conservation analysis. **(a)** Phylogenetic tree constructed from 18S rRNA sequences showing evolutionary relationships among *T. gondii* and selected organisms: *H. hammondi (Hh)*, *N. caninum (Nc)*, *B. besnoiti (Bb)*, *C. suis (Cs)*, *S. neurona (Sc)*, *C. cayetanensis (Cc)*, and *E. falciformis (Ef)*. **(b)** Percentage identity distributions across all alignments **(c)** Bit score distributions across all alignments for each organism. **(d)** E-value distributions across all alignments for each organism. **(e)** Percentage identity distributions across the bit-score selected homologs in each organism **(f)** Number of *T. gondii* proteins with detectable homologs in each organism (selected by highest bit score) and number with no detectable homologs. Black represents the number of proteins with homologs identified and grey represents the proteins with no homolog found. **(g)** Percentage overlap between bit score-derived homologs and OrthoVenn3-identified orthologs, demonstrating validation of the bit score approach.

The next step was to carry out a phylogenetic analysis by quantifying the conservation of each *T. gondii* protein across these organisms. For this, complete lists of annotated proteins for all selected organisms were retrieved from ToxoDB (version 68)(Alvarez-Jarreta et al., 2024) and aligned against the *T. gondii* reference proteome using the Basic Local Alignment Search Tool (BLAST). To capture all potential homologs or “hits”, regardless of divergence level, no threshold filters were imposed on E-value, percent identity, query coverage, or bit score. This approach yielded more hits than the 8,204 proteins in the *T. gondii* proteome, indicating that individual *T. gondii* proteins frequently matched with multiple sequences in the organisms being compared. For further analysis, a single hit for each *T. gondii* protein was selected from each organism. This one-to-one mapping requires that the highest-quality hit for each query is retained. Further, this hit should be associated with a quantitative measure of its evolutionary distance from the *T. gondii* homolog. Therefore, selecting an appropriate metric to assess alignment quality and identify true homologs among multiple hits was critical. Three commonly used parameters were evaluated: percent identity, expectation value (E-value), and bit score.

Percentage identity between two sequences gives a quantitative measure of homology. The distribution of percentage identity across all the alignment hits is given in Figure 1b. Percentage identity retained the evolutionary relationship between these organisms (median values ranged from 88.89% in *Hammondia* to 38.46% in *Eimeria*), it did not exclude the probability of finding chance alignments (Pearson, 2013). Moreover, percentage identity is particularly insufficient when working with closely related organisms as multiple hits are found for a single protein with high percentage identity (>70%); hence, percentage identity was not used further.

E-value represents the probability of obtaining an alignment by chance, with lower values indicating more significant matches. The formula for E-value includes the length of the query sequence, the size of the database, and the bit score. Bit score is a normalised value of raw alignment score calculated by match, mis-matches and gap penalties within the alignment based on the algorithm used. Statistically, an E-value < 10^-5^ or an alignment with a bit score > 50 is considered significant (Pearson, 2013; Svanberg Frisinger et al., 2021).

Since both these parameters can assess evolutionary distance reliably, a comparison of these parameters in all the hits of *T. gondii* proteins (8204) across all the selected organisms were done. The distribution of bit scores from the alignments across all organisms revealed that *Hammondia* exhibited a wider spread and generally higher bit scores compared to the other species. As the evolutionary distance from *Hammondia* to *Eimeria* increases, the bit score distribution becomes more compact, with maximum values decreasing from approximately 24,000 in *Hammondia* to around 6,000 in *Eimeria* reflecting their evolutionary distance (Figure 1 c). In contrast, E-value distributions were nearly identical across all species and failed to reflect evolutionary distances (Figure 1 d).

Many protein alignments between closely related species yielded E-values of zero, leading to difficulty in discrimination between top hits based on this metric alone. Importantly, bit score provided unique values for each alignment, enabling discrimination even when E-values were identical. For example, for a P-type ATPase (TGME49_201150), matched four sequences in *H. hammondi* with bit scores of 1,053, 1,008, 385, and 81. The top two alignments both yielded E-values of zero, while the remaining alignments had E-values of 1.09×10⁻¹⁰⁹ and 1.66×10⁻¹⁵. Based on this discriminatory power, bit score was selected as the metric for further analyses of conservation, with the alignment having the highest bit score for each *T. gondii* protein being retained. It is important to note that bit score has been used for phylogenetic comparisons in other reports (Shaukat et al., 2025; Svanberg Frisinger et al., 2021; Wheeler et al., 2016).

Using bit scores, the majority (>85%) of best hits across all organisms exhibited >30% sequence identity (Figure 1 e), a threshold commonly associated with detectable homology in protein comparisons. This distribution suggests that bit score selection preferentially retains alignments of sufficient similarity to infer functional relationships, while appropriately excluding low-complexity or domain-only matches. However, not all *T. gondii* proteins yielded detectable homologs in every organism. As expected, homolog detection rates correlated inversely with evolutionary distance (Figure 1 f). The proteins which did not match with any sequences in an organism were given a bit score value of zero. It was observed that *H. hammondi* showed the highest detectable homologs (∼89% of *T. gondii* proteins), while the more divergent *E. falciformis* and *C. cayetanensis* exhibited substantially lower number of homologs (59% and 62%, respectively). These patterns align with phylogenetic relationships, but raised the question of whether bit score-based selection reliably identifies functional orthologs.

To address this question, an orthology analysis was performed using OrthoVenn3 (Sun et al., 2023). The percentage overlap between bit score-selected homologs and OrthoVenn3-derived ortholog alignments was calculated. The overlap exceeded 90% for all organisms (Figure 1 g), validating that the bit score-based approach reliably identifies orthologs. This high degree of overlap confirmed the suitability of bit score for subsequent conservation analyses. It is important to note that OrthoVenn3 was also not able to distinguish evolutionary relationships (data not shown) and served merely as a confirmation for the homologs obtained using the highest bit score values.

Having selected the organisms, a pipeline for identifying conservation patterns among the selected organisms was developed (Figure 2). Using standalone BLAST, *T. gondii* proteins were aligned with all proteins from each of the organisms under study. If homologs were found, the protein with the highest bit score was selected and the bit score noted. If no homolog was found, a bit score of 0 was assigned. Next, bit scores of *T. gondii* proteins were converted into deciles, and these were used for clustering using K-means and hierarchical clustering. Three properties of each cluster were studied: conservation trends among the organisms under study, whether any known bradyzoite-formation proteins were present in the cluster, and COG analysis of proteins in the cluster.

**Figure 2:**
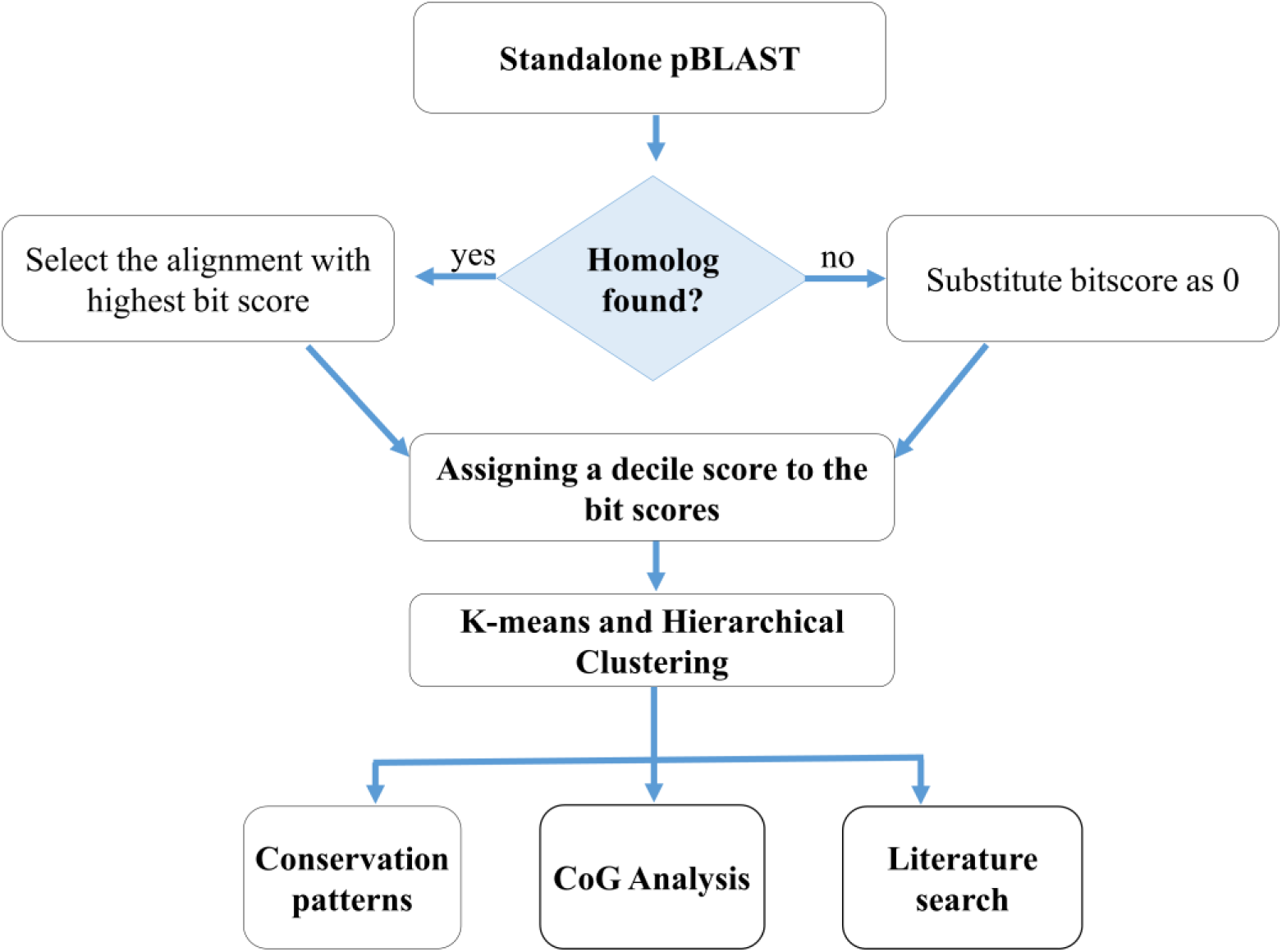
Pipeline for identifying candidate bradyzoite differentiation proteins. Flowchart depicting the comparative proteomics workflow used to identify *T. gondii* proteins potentially involved in bradyzoite differentiation through phylogenetic conservation analysis.

### Clustering of proteins with the highest bit score based on conservation patterns

The homologs of *T. gondii* proteins in the selected organisms showed bit score values ranging from 0 to ∼24,000. To minimise skewness in the data distribution, bit scores were binned into categories (decile score) according to the decile range they fell into, as shown in Table 1. Since the first two decile categories represented bit score values of zero, and the third category encompassed bit scores of 0-47, any alignment with a bit score less than 47 was assigned a decile score of 3. This adjustment resulted in a decile score scale ranging from 3 to 10, where 3 represents the least conserved proteins, and 10 represents the most highly conserved proteins. These data revealed that ∼30% of proteins yielded the lowest bit score (0-47); these had higher E-value and lower % identity compared to other alignments, confirming that they were not true homologs.

**Table 1:**
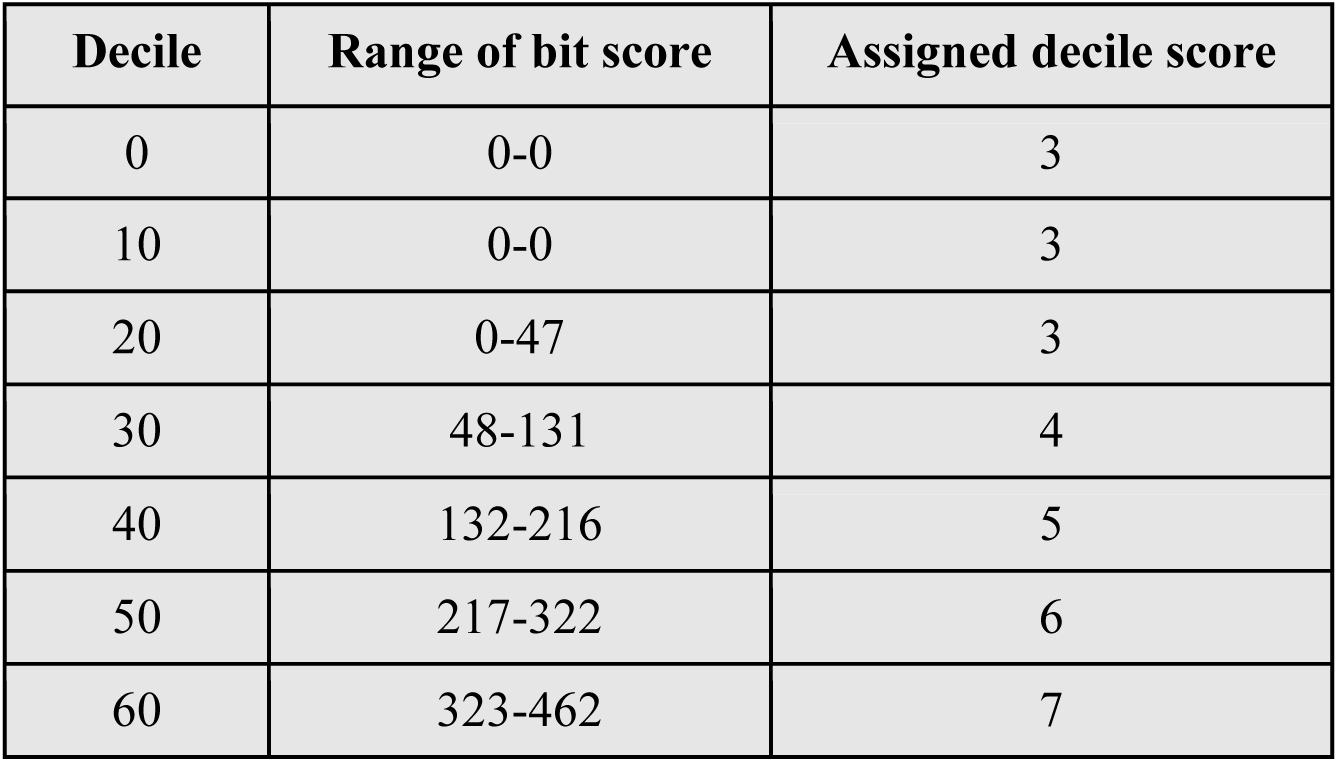
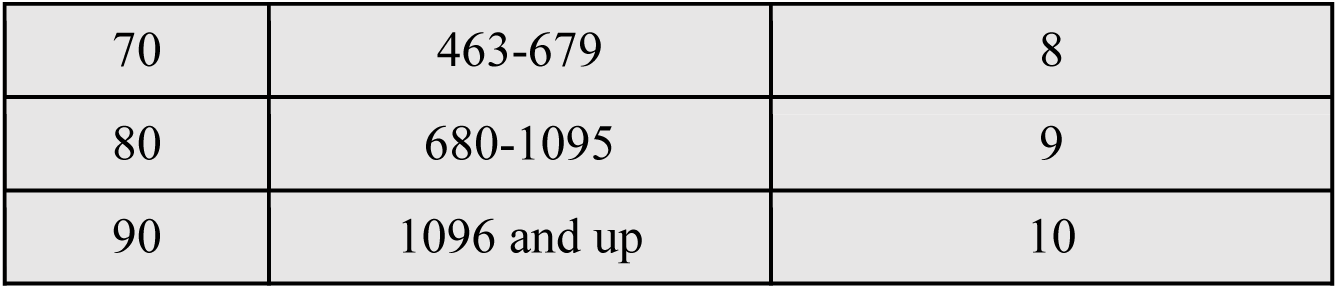
Bit score ranges and corresponding decile scores.

Having assigned deciles to the bit score values, each homolog of the 8,204 *T. gondii* proteins was assigned a decile score based on their bit scores (Figure 3 a). In *H. hammondi*, 924 proteins fell into the decile score of 3, while the majority (3179), clustered in the highly conserved decile score of 10. This pattern was reversed with increasing evolutionary distance: *E. falciformis* and *C. cayetanensis* exhibited more alignments in decile score 3 (3800 and 3922, respectively) and substantially fewer (106 and 69 respectively), in 10. This distribution pattern confirmed that substituting the raw bit score with decile scores preserved the underlying evolutionary relationship present in the dataset.

**Figure 3:**
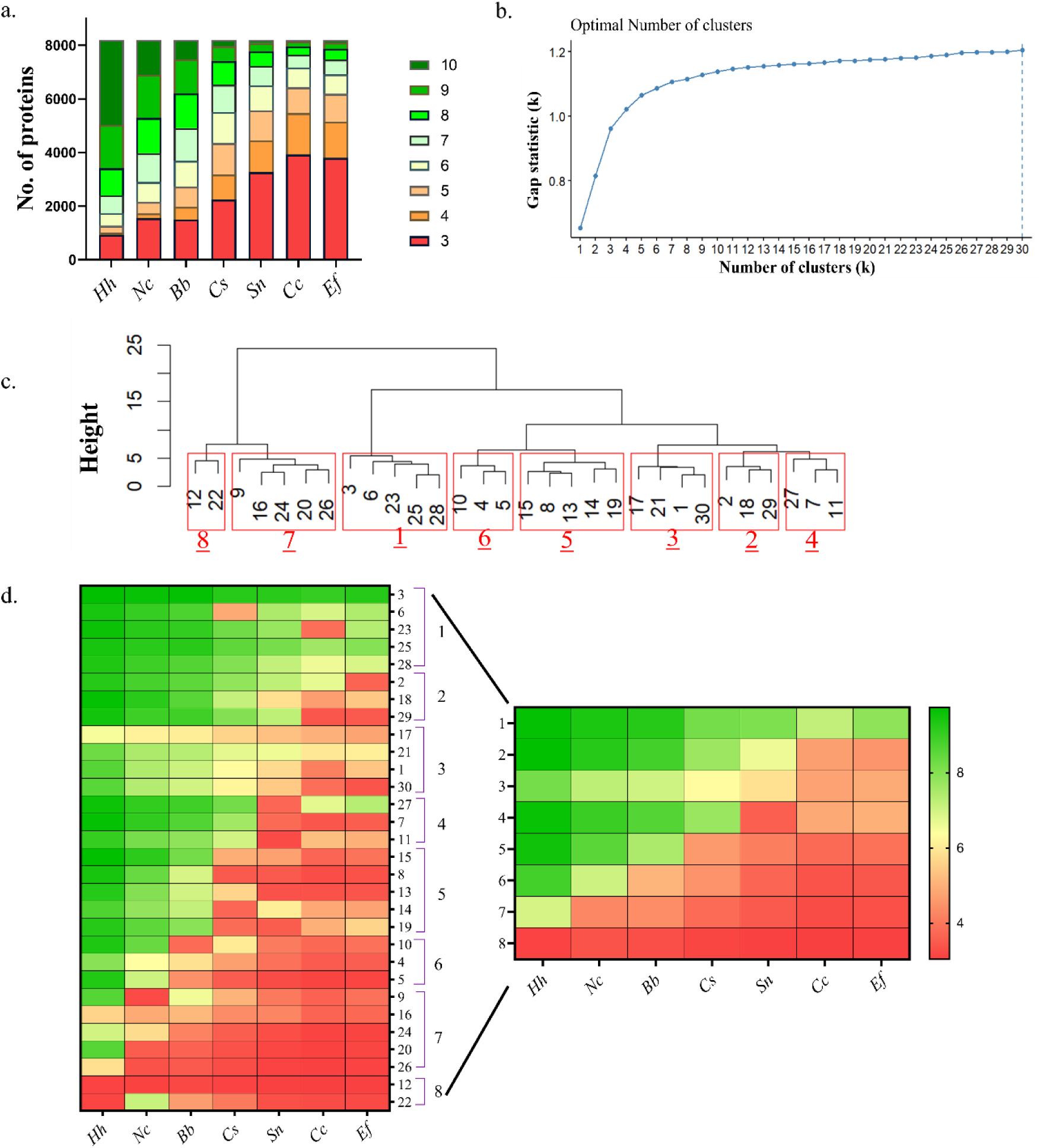
Clustering based on conservation pattern. (a) Distribution of protein alignments across decile score bins for each selected. The organisms are represented by two letter codes represent *H. hammondi (Hh)*, *N. caninum (Nc)*, *B. besnoiti (Bb)*, *C. suis (Cs)*, *S. neurona (Sc)*, *C. cayetanensis (Cc)*, and *E. falciformis (Ef)*. (b) Gap statistical analysis results indicating optimal cluster number (k = 30) based on a maximum of 30 clusters and 500 bootstrap iterations. (c) Dendrogram representing relationships among the 30 K-means clusters. Red boxes indicate K-means clusters grouped together to form 8 meta-clusters. (d) Heatmap showing conservation patterns within each of the 30 clusters obtained via K-means clustering compiled into 8 meta-clusters. Values represent mean decile scores for each organism within a cluster.

The mean decile score across all the 8204 *T. gondii* protein alignments was calculated for all the organisms. As expected, mean decile scores decreased progressively with phylogenetic distance, declining from *H. hammondi* to *C. cayetanensis* (Table 2). These numbers provide a reference for identifying clusters with conservation patterns driven by evolutionary distance. This phylogenetic gradient provides a reference framework for identifying proteins whose conservation patterns deviate from evolutionary expectations that may signal functional specialization, adaptive evolution, or lineage-specific divergence.

**Table 2:**
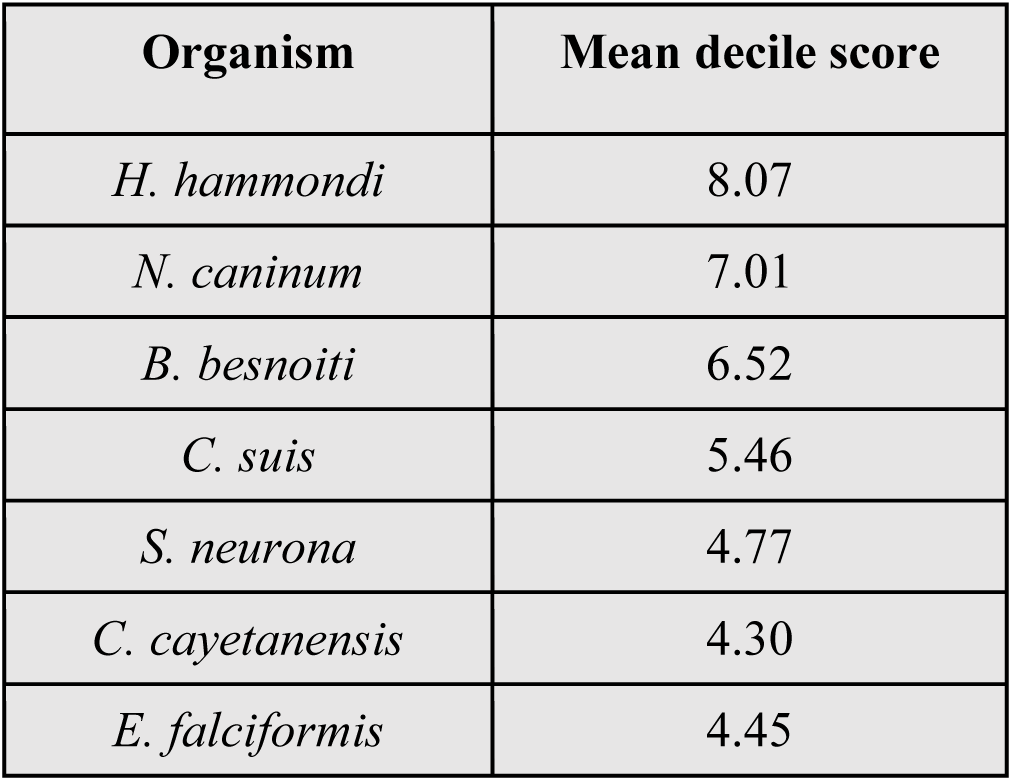
Mean decile score of all alignments reflecting evolutionary relationships.

| Organism | Mean decile score |
| --- | --- |
| *H. hammondi* | 8.07 |
| *N. caninum* | 7.01 |
| *B. besnoiti* | 6.52 |
| *C. suis* | 5.46 |
| *S. neurona* | 4.77 |
| *C. cayetanensis* | 4.30 |
| *E. falciformis* | 4.45 |

To find conservation patterns, two rounds of clustering were employed. To achieve this, in the first round, K-means clustering was applied to the decile score matrix. As shown in Figure 3 b, the Gap statistic value plateaued beyond 30 clusters, indicating that additional clusters would not yield meaningful new information. In contrast, the Silhouette and Elbow methods suggested *k* = 2 as optimal (Supplementary Figure 1 a and b). However, a two-cluster solution was deemed insufficient for resolving biologically meaningful conservation patterns, as it would merely separate highly conserved proteins from poorly conserved ones without distinguishing intermediate conservation levels or lineage-specific signatures. We therefore proceeded with *k* = 30.

To further group clusters with similar conservation profiles into meta-clusters, sub-clustering was performed. Silhouette and Elbow methods suggested only two meta-clusters (Supplementary Figure 2 a and b), again insufficient for biological resolution. Finally, a hierarchical clustering approach was implemented. The hierarchical clustering produced a dendrogram illustrating relationships among the 30 K-means clusters (Figure 3 c). A dynamic tree-cutting algorithm based on dendrogram topology yielded four meta-clusters (Supplementary Figure 2 c). However, these four suggested meta-clusters resulted in overly broad clusters that masked distinct conservation patterns, while higher number of clusters (K ≥ 10) led to over-fragmentation with limited additional biological interpretability. To balance statistical inference with biological interpretability, the dendrogram was cut at a height of 5.48 Euclidean distance units, yielding eight meta-clusters. The red boxes in Figure 3 c indicate the K-means clusters grouped to form each meta-cluster.

Conservation trends within the 30 clusters and the 8 meta-clusters were visualized using a heatmap of mean decile scores for each organism (Figure 3 d), confirming distinct conservation profiles across meta-clusters. The meta-clusters showed clear trends: one consisted of proteins that are conserved among all the organisms studied (cluster 1); another consisted of proteins that are unique to *T. gondii* (cluster 8). The remaining six clusters showed conservation between *T. gondii* proteins and different organisms: one contains proteins that are conserved in the cyst-forming *Sarcocystidae* family (cluster 2), and another contains proteins that are conserved in bradyzoite cyst-forming organisms (cluster 5).

### Conservation patterns of meta-clusters

To visualize conservation patterns for each cluster, heat maps along with the mean and standard deviation of the decile scores for each of the organisms were generated (Figure 4). Cluster 1, consisting of 1,101 proteins, exhibited high conservation (mean decile scores ranging from 9.67+/-0.64 to 7.87+/-1.10) across all selected organisms (Figure 4 a). This suggested that these proteins may perform housekeeping functions, regardless of family or cyst-forming capability. Interestingly, proteins with reduced conservation scores in *C. suis* and *C. cayetanensis* were found, and these mapped to distinct K-means subclusters (clusters 6 and 23, respectively; Figure 3d), despite their placement in the otherwise highly conserved meta-cluster 1. Gene ontology analysis of these proteins revealed divergence in distinct cellular functions: DNA recombination, energy metabolism, and protein catabolism (divergent in *C. suis*, K-means cluster 6); vesicle transport, cell motility, protein trafficking, and mRNA metabolism (divergent in *C. cayetanensis*, K-means cluster 23). These classes of proteins appear to have evolutionary trajectories that are lineage-specific.

**Figure 4:**
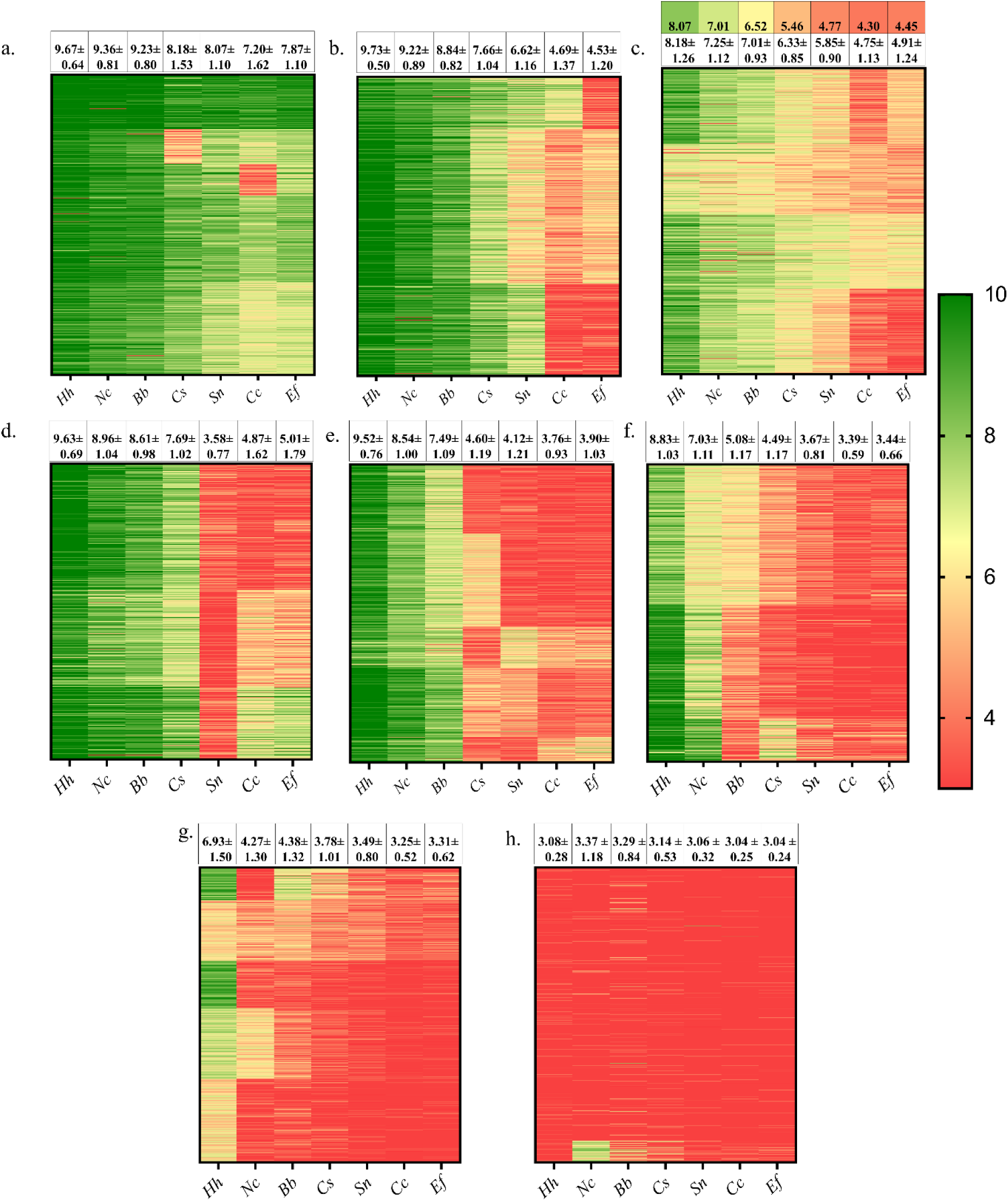
Patterns of conservation in each meta cluster. Each cluster is represented with a graph representing mean ± SD of decile_score for each organism and a corresponding heat map. Green = highly conserved; Yellow= moderately conserved and Red = Least conservation (a) Cluster 1 (b) Cluster 2 (c) Cluster 3; the color coded values represent the mean decile scores of the entire data set representing evolutionary relationship (d) Cluster 4 (e) Cluster 5 (f) Cluster 6 (g) Cluster 7 (h) Cluster 8. The organisms are represented by two letter codes represent *H. hammondi (Hh)*, *N. caninum (Nc)*, *B. besnoiti (Bb)*, *C. suis (Cs)*, *S. neurona (Sc)*, *C. cayetanensis (Cc)*, and *E. falciformis (Ef)*.

Cluster 2 (665 proteins) exhibited high conservation across all *Sarcocystidae* members (mean decile scores ranging from 9.73+/-0.50 to 6.62+/-1.16 from *Hammondia* to *Sarcocystis*) but markedly reduced conservation in the *Eimeriidae* outgroup consisting of *C. cayetanensis* and *E. falciformis* with mean decile scores 4.69+/-1.37 and 4.53+/-1.2, respectively (Figure 4 b). This trend indicated that these proteins are conserved within the *Sarcocystidae* family and may contribute to unique characteristics of this family, such as tissue cyst-forming abilities or their ability to infect diverse hosts.

Cluster 3 (1,157 proteins) showed a trend where the mean decile scores progressively decreased across the selected organisms. Since such patterns could reflect evolutionary distances, the mean decile score across the entire dataset (8204 *T. gondii* protein alignments) was calculated for all the organisms. As expected, mean decile scores decreased progressively with phylogenetic distance, declining from *H. hammondi* to *C. cayetanensis* (Figure 4 c, colour-coded values). Of the 8 clusters, cluster 3 exhibited a conservation profile that tracked the trend seen for mean decile scores of evolutionary distances (Figure 4 c).

Cluster 4 (593 proteins) showed a conservation profile with higher mean decile scores in all organisms within *Sarcocystidae* (9.63+/-0.69 to 7.69+/-1.02), but with notable divergence in *S. neurona*, where the mean decile score was 3.58+/-0.77 (Figure 4 d). The mean decile scores increased again in *Cyclospora* (4.87+/-1.62) and *Eimeria* (5.01+/-1.79). It is tempting to speculate this result reflects the exceptionally broad host range of the *Sarcocystis* genus, infecting numerous domestic and wild animal species (Dubey, 2022; Dubey et al., 1991).

Cluster 5 (1,338 proteins) showed conservation in the three bradyzoite cyst-forming species (*H. hammondi, N*. *caninum*, *B. besnoiti* with mean decile scores of 9.52+/-0.76, 8,54+/-1, and 7.49+/-1.09 respectively), with much lower conservation in *C. suis* and *S. neurona* (4.60+/-1.19 and 4.12+/-1.21 respectively) and the *Eimeriidae* outgroup (Figure 4 e). This cluster may contain proteins that play roles in bradyzoite differentiation and maintenance.

Cluster 6 (913 proteins) showed higher mean decile scores in *Hammondia* and *Neospora* (8.83+/-1.03 and 7.03+/-1.11, respectively) than other members of *Sarcocystidae viz. B. besnoiti* (5.08+/-1.17); *C. suis* (4.48+/-1.17) and *S. neurona* (3.67+/-0.81) as shown in Figure 4 f. Studies have found >95% synteny between *Hammondi*a and *Toxoplasma* (Walzer et al., 2013), and >80% between *Toxoplasma* and *Neospora* (Adomako-Ankomah et al., 2014) indicating close evolutionary relationships as confirmed in the phylogenetic tree (Figure 1a). Consistent with these relationships, for *Toxoplasma*, *Hammondia* and *Neospora*, cyst wall morphology and the requirement of bradyzoites for infection into the primary host are similar (Sokol-Borrelli et al., 2020). Proteins in cluster 6 may play roles in these biological functions.

Cluster 7 (1,460 proteins) exhibited high conservation exclusively in *H. hammondi* (mean decile score 6.93+/-1.5) and markedly reduced scores in all other organisms (mean decile scores 3-4) (Figure 4 g). This is consistent with the exceptionally close relationship between *Toxoplasma* and *Hammondia*. They share the same definitive host (felids), nearly identical life cycles, and morphological similarity (Walzer et al., 2013), indicating recent evolutionary divergence.

Cluster 8 (977 proteins) showed minimal conservation across all organisms (mean decile scores <4), indicating *T. gondii*-specific proteins or extensive divergence (Figure 4 h). *T. gondii* exhibits differences compared to other genera, including the ability to undergo robust bidirectional tachyzoite-bradyzoite interconversion in response to environmental changes (Sokol-Borrelli et al., 2020; Vizcarra et al., 2023). Cluster 8 proteins may thus contribute to *T. gondii*-specific adaptations of its persistence, host range, or metabolic specialisations. Interestingly, a subset of proteins exhibited elevated conservation in *N. caninum* (found in K-means cluster 22). These proteins were not enriched for any GO category and were annotated as hypothetical proteins. The details of *T. gondii* proteins found in each cluster are available in Supplementary Table 1.

### Each cluster displays enrichment of distinct functional categories

Functional enrichment analysis was performed to identify characteristic differences among clusters based on their predominant functional categories. Enriched and depleted functional categories with their corresponding odds ratios are visualized in Figure 5.

**Figure 5:**
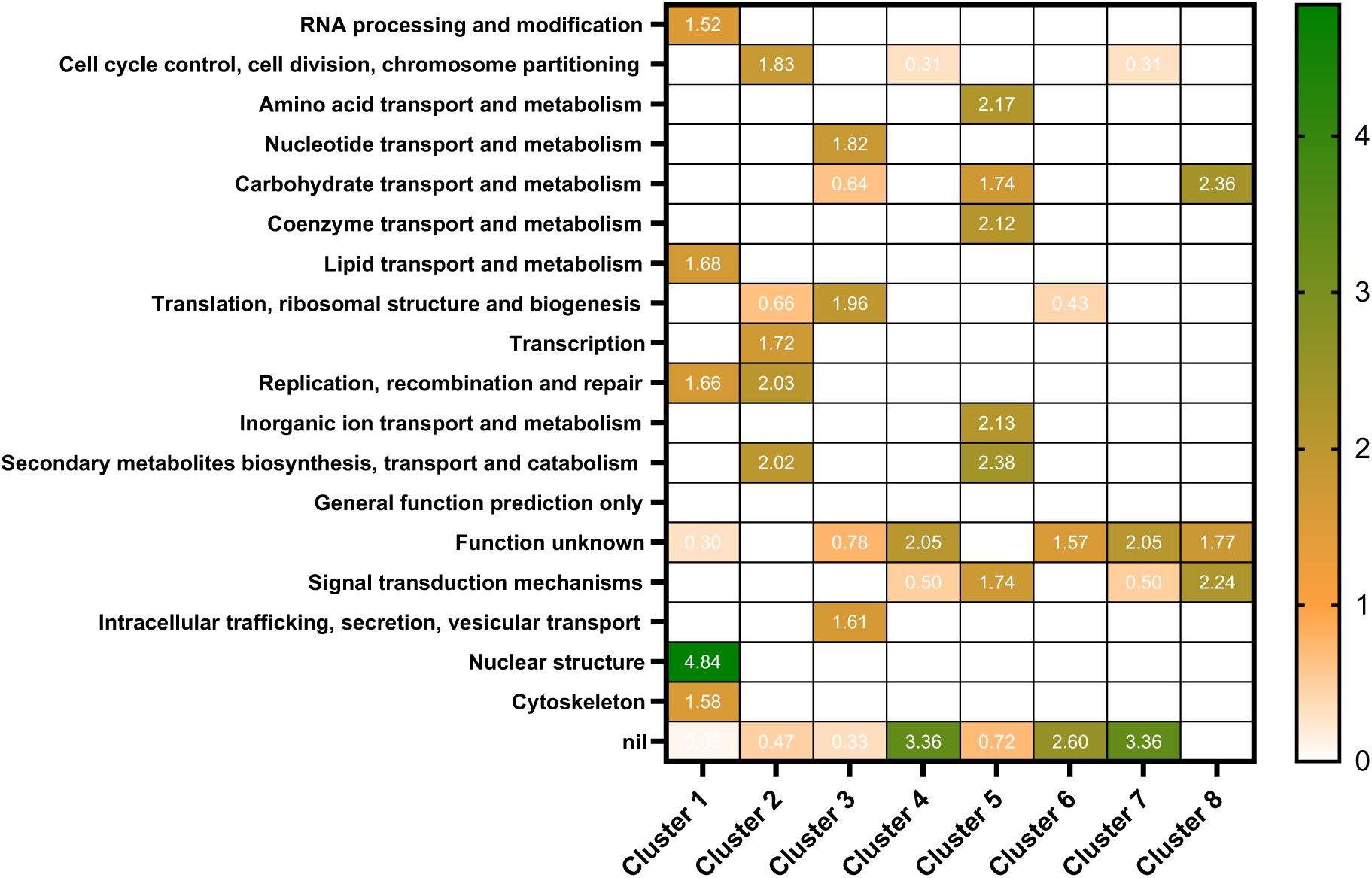
COG Functional annotations of each of the meta cluster. A heatmap showing the odds ratio of COG categories that are under or over represented in each of the cluster. The category “function unknown” represents proteins whose identifiers like KEGG pathway, pFAM domains are identified but exact functional annotation is difficult. “nil” represents proteins whose details are completely unknown and are unannotated by COG. Few COG categories that were not enriched were deleted and only the enriched are plotted.

Cluster 1 contains proteins that are conserved among all the organisms under study. This cluster showed significant enrichment of COG categories associated with housekeeping functions, including RNA processing and modification, lipid transport and metabolism, nuclear structure, and cytoskeleton organization. Notably, this cluster contained a low proportion of proteins with unknown function or lacking COG annotation (designated as “nil”). The scarcity of unannotated proteins supports the observation that this cluster predominantly represents essential cellular functions which are mostly conserved and well-studied across various eukaryotic systems.

Cluster 2 contains proteins that are highly conserved in the *Sarcocystidae* family. These proteins showed significant enrichment of the COG categories cell cycle control, transcription, DNA replication and repair, and secondary metabolite transport suggesting these regulatory functions underwent family-specific adaptations within the tissue cyst-forming *Sarcocystidae*. Translation and proteins classified as “function unknown” were significantly underrepresented.

Cluster 3 showed a conservation trend that tracked evolutionary relationships between the organisms under study. This cluster exhibited enrichment of proteins involved in nucleotide transport and metabolism, translation, and intracellular trafficking/vesicular transport. These categories were not enriched in any other cluster, suggesting these functions evolved with phylogenetic distance, accumulating sequence variation as lineages diverged.

Cluster 4 was divergent in *Sarcocystis neurona* and showed the highest proportion of proteins unannotated in COG (nil, odds ratio 3.36) or proteins with unknown function (odds ratio 2.05). Critical cellular processes including cell cycle regulation and signal transduction were significantly underrepresented. This pattern suggests two possibilities: (1) *S. neurona* has evolved lineage-specific proteins fulfilling these functions that lack clear orthologs in the *T. gondii* reference databases, or (2) annotation quality for *Sarcocystis* proteins is lower compared to other selected organisms due to limited functional characterization.

Cluster 5 showed an interesting trend of proteins being conserved in bradyzoite cyst-forming organisms. This cluster exhibited enrichment for signal transduction, secondary metabolite biosynthesis, inorganic ion transport and metabolism, coenzyme transport and metabolism, carbohydrate transport and metabolism, and amino acid transport and metabolism. This functional profile suggests that bradyzoite cyst-forming organisms may have evolved specific proteins involved in signal transduction cascades and diverse transport processes. This enrichment pattern correlates with bradyzoite biology: differentiation is triggered by environmental stress signals requiring complex signal transduction pathways, while cyst formation demands extensive protein trafficking, including secretion of cyst wall components. The specific enrichment of carbohydrate metabolism is particularly notable, as bradyzoites exhibit reduced TCA cycle activity and rely primarily on glycolysis for energy production (Cerutti et al., 2020; Shukla et al., 2018).

Clusters 6 and 7 showed high enrichment for proteins with unknown function and lacking COG annotation. These clusters contained proteins that are conserved in *T. gondii*, *H. hammondi*, and/or *N. caninum.* In cluster 6, lipid transport and metabolism were significantly underrepresented, suggesting that these processes have diverged between *T. gondii* versus *H. hammondi* and *N. caninum*. In cluster 7, signal transduction mechanisms and cell cycle control were underrepresented.

Cluster 8 (*Toxoplasma*-specific) showed no significantly underrepresented COG categories, indicating *T. gondii*-specific proteins span diverse functional classes. Nucleotide transport and metabolism and signal transduction were significantly enriched, suggesting *T. gondii* has evolved specialized adaptations in nucleotide homeostasis and signalling pathways.

### Proteins involved in bradyzoite differentiation fall into all clusters, except cluster 7 and 8

To assess the evolutionary origins of bradyzoite differentiation machinery, a manual curation of previously characterised proteins involved in bradyzoite conversion was performed. These proteins fell into two classes: first, a decrease in bradyzoite formation upon knockdown of the protein, indicating roles as positive regulators and second, an increase in bradyzoite formation upon knockdown, indicating roles as negative regulators. Each known regulator was mapped to its respective conservation cluster (Table 3) to determine whether *T. gondii* employs ancestral machinery broadly conserved across eukaryotes, family-specific conservation restricted to tissue cyst-forming *Sarcocystidae*, lineage-specific molecular players unique to bradyzoite cyst-forming organisms, or a combination of these strategies.

**Table 3:** Known molecular players of bradyzoite conversion and their corresponding clusters.

| Gene ID | Product description | Effect of KO on bradyzoite formation | Reference | Cluster No. |
| --- | --- | --- | --- | --- |
| TGME49_22<br>5490 | CDPK-2 | Affected amylopectin metabolism, unviable bradyzoites | Uboldi et al., 2015 | 1 |
| TGME49_24<br>6510<br>TGME49_20<br>0400 | PP2 A holoenzyme (regulatory subunit) | Affected amylopectin metabolism, unviable bradyzoites | Wang et al., 2022 | 1 |
| TGME49_31<br>5670 | PP2A holoenzyme (scaffolding) | Affected amylopectin metabolism, unviable bradyzoites | Wang et al., 2022 | 2 |
| TGME49_22<br>4220 | PP2A holoenzyme<br>(catalytic) | Affected amylopectin<br>metabolism, unviable<br>bradyzoites | Wang et al.,<br>2022 | 2 |
| TGME49_22<br>9630 | eIF2 Kinase A | Reduced cyst burden under ER<br>stress | Augusto et<br>al., 2018 | 2 |
| TGME49_22<br>4050 | AP2 domain<br>transcription factor<br>AP2X-4 | Increased transcription of<br>bradyzoite specific genes | Zhang et al.,<br>2022 | 2 |
| TGME49_26<br>8860 | Enolase 1 | No KO studies, marker for<br>bradyzoites | Dzierszynski<br>et al., 2001 | 2 |
| TGME49_22<br>3410 | eukaryotic<br>initiation factor<br>eIF4E1 | Induces bradyzoite formation | Holmes et al.,<br>2024 | 3 |
| TGME49_28<br>6090 | translation<br>initiation factor<br>SUI1, putative<br>eIF1.2 | Reduces cyst burden | Wang et al.,<br>2023 | 3 |
| TGME49_28<br>6470 | cAMP-dependent<br>protein kinase | Induces bradyzoite<br>differentiation | Sugi et al.,<br>2001 | 4 |
| TGME49_26<br>4660 | CST-1 | Fewer and fragile cysts | Zhang et al.,<br>2001 | 4 |
| TGME49_29<br>1040 | LDH-2 | Reduced cyst burden | Abdelbaset et<br>al., 2017 | 4 |
| TGME49_20<br>0385 | BFD-1 | KO stops differentiation | Waldman et<br>al., 2020 | 5 |
| TGME49_31<br>1100 | BFD-2 | Significantly reduces cyst-burden | Licon et<br>al., 2023 | 5 |
| TGME49_31<br>8470 | AP2 domain<br>transcription factor<br>AP2IV-4 | Repress expression of bradyzoite<br>genes in tachyzoites | Radke et al.,<br>2018 | 5 |
| TGME49_31<br>5760 | AP2 domain<br>transcription factor<br>AP2XI-4 | Reduces cyst-burden | Walker et al.,<br>2012 | 5 |
| TGME49_31<br>8610 | AP2 domain<br>transcription factor<br>AP2IV-3 | Reduces cyst-burden | Hong et al.,<br>2017 | 5 |
| TGME49_21<br>7700 | AP2 domain<br>transcription factor<br>AP2XII-2 | Increases cyst burden | Srivastava et<br>al., 2020 | 5 |
| TGME49_28<br>8950 | AP2 domain<br>transcription factor<br>AP2IX-2 | Reduces cyst-burden | Huang et<br>al.,2017 | 5 |
| TGME49_21<br>4960 | AP2 domain<br>transcription factor<br>AP2X-8 | Increases cyst burden | Li-Xiu sun et<br>al., 2025 | 5 |
| TGME49_29<br>3280 | Cyclin-5 | Reduces cyst formation | Naumov et<br>al,2022 | 6 |
| TGME49_31<br>1510 | eIF2 kinase IF2K-B | Reduced cysts in alkaline and<br>oxidative stress conditions | Augusto et<br>al., 2021 | 6 |
| TGME49_30<br>6620 | AP2 domain<br>transcription factor<br>AP2IX-9 | Repress expression of bradyzoite<br>genes in tachyzoites | Radke et al.,<br>2013 | 6 |
| TGME49_25<br>1740 | AP2 domain<br>transcription factor<br>AP2XII-9 | Increases cyst burden | Shi et al.,<br>2024 | 6 |
| TGME49_25<br>9020 | Bag 1 (Surface<br>antigen) | Reduces cyst burden | Zhang et<br>al.,1999 | 6 |

Cluster 1 contained three proteins involved in amylopectin metabolism: CDPK2 and two PP2A regulatory subunits; this cluster also contained the splicing factor, cdc 5. Amylopectin serves as the primary energy storage mechanism in bradyzoite cysts, and disruption of its metabolism results in morphologically abnormal, non-viable cysts (Uboldi et al., 2015; J.-L. Wang et al., 2022). However, both CDPK2 and PP2A also function in tachyzoite metabolism, with knockouts causing increased amylopectin accumulation in tachyzoites and reduced plaque formation (Uboldi et al., 2015; J.-L. Wang et al., 2022). Similarly, cdc 5 is involved in a plethora of functions like DNA replication and repair as well was protein folding (Kashyap et al., 2025). As cluster 1 contains proteins that are conserved in all the organisms under study, these proteins appear to have housekeeping functions with additional roles in bradyzoite cyst formation.

Cluster 2 contained five bradyzoite-associated proteins: PP2A catalytic and scaffolding subunits (Augusto et al., 2018), three eIF2α kinase paralogs (eIF2K-A, -C, -D)(Konrad et al., 2011, 2014; Narasimhan et al., 2008), AP2X-4 (J. Zhang et al., 2022), and enolase (Dzierszinski et al., 2001). Interestingly, PP2A regulatory subunits were found in cluster 1, while the catalytic and scaffolding subunits localized to cluster 2. This distribution suggests distinct evolutionary trajectories for different components of the same protein complex: regulatory subunits retain ancient functions conserved across apicomplexans, while catalytic/scaffolding subunits underwent family-specific adaptations. Enolase, while used as a bradyzoite marker in *T. gondii*, showed conservation in *Sarcocystidae* including genera forming alternative cyst types (*Cystoisospora*, *Sarcocystis*), indicating it functions in broader metabolic shifts during chronic infection, specifically the preference for glycolytic metabolism of differentiated stages (Denton et al., 1996; Shukla et al., 2018).

Cluster 3 that tracked the trend of evolutionary relationships between the organisms, contained two translation initiation factors, eIF4E1 and eIF1.2. These translation factors correlated with enrichment of translation in the COG categories found in this cluster. Both proteins maintain core translational functions across *Sarcocystidae* but have been adapted for bradyzoite-specific regulatory roles in *T. gondii*: eIF4E1 functions as a differentiation repressor maintaining the tachyzoite state (Holmes et al., 2024), while eIF1.2 regulates BFD1 and BFD2 translation (F. Wang et al., 2024).

Cluster 4 showed reduced conservation in *S. neurona*. This cluster contained four proteins: LDH2 (Abdelbaset et al., 2017), CST1(Tomita et al., 2013; Y. W. Zhang et al., 2001), cAMP-dependent kinase (Sugi et al., 2016). Notably, LDH2 (bradyzoite-specific lactate dehydrogenase in *T. gondii*) diverged in *Sarcocystis*, while LDH1 (tachyzoite-specific) was conserved in *Sarcocystis* (cluster 2). *Toxoplasma* bradyzoites rely primarily on glycolysis with reduced TCA cycle activity (Cerutti et al., 2020; Shukla et al., 2018); the LDH2 divergence may indicate fundamental differences in *Sarcocystis* cyst metabolism. The divergence of CST1 (cyst wall glycoprotein) aligns with the morphologically distinct septate cyst architecture of *Sarcocystis*, suggesting alternative cyst wall assembly mechanisms in this genus.

Cluster 5 (proteins conserved in bradyzoite-forming organisms) contained the highest number of characterized bradyzoite regulators and was strongly enriched for transcriptional and cell cycle control factors. This cluster included BFD1 and BFD2, key transcriptional and translational regulators controlling stage conversion (Waldman et al., 2020; Licon et al., 2023). BFD1 is a Myb-like transcription factor whose knockout completely abolishes cyst formation, while BFD2 post-transcriptionally regulates BFD1 expression, revealing regulatory hierarchy.

Notably, cluster 5 was the only cluster containing AP2 factors with demonstrated bradyzoite-enhancing activity; AP2 factors in other clusters function primarily in lytic cycle maintenance. Here we found AP2XI-4, AP2IV-3, and AP2IX-2, all essential for tachyzoite viability yet upregulated under bradyzoite-inducing conditions where they enhance transcription of bradyzoite-specific genes like BAG1 (Walker et al., 2013; Hong et al., 2017; Huang et al., 2017). Repressors in cluster 5 included AP2IV-4, AP2XII-2, and AP2X-8. Knockout of these factors causes cell cycle elongation and increased bradyzoite differentiation (Radke et al., 2018; Srivastava et al., 2020; Sun et al., 2025). However, AP2XII-2 and AP2X-8 are also essential for tachyzoite replication, indicating their primary function is maintaining rapid cell division, with bradyzoite suppression as a secondary consequence as disrupting the cell cycle favours differentiation.

The selective conservation of BFD1/BFD2 and bradyzoite-enhancing AP2 factors specifically in bradyzoite cyst-forming organisms validates cluster 5 as enriched for proteins that play roles in bradyzoite differentiation, representing evolutionary adaptations within the *Toxoplasmatinae* subfamily. The presence of these regulators within this cluster makes it the primary choice for finding new molecular players of stage conversion.

Clusters 6 contained proteins that are conserved in *T. gondii*, *H. hammondi*, and *N. caninum.* These proteins were eIF2K-B (Augusto et al., 2021), cyclin-5 (Naumov et al., 2022), AP2IX-9 (Radke et al., 2013), AP2XII-9 (Shi et al., 2024), and BAG1 (Bohne et al., 1995b; Y. W. Zhang et al., 1999). eIF2K-B is one of four eIF2α kinases in *T. gondii*, however, the other three were found in the *Sarcocystidae*-specific cluster 2. Cyclin-5 functions to prolong the cell cycle in bradyzoites (Naumov et al., 2022), while AP2IX-9 and AP2XII-9 primarily maintain lytic cycle progression but secondarily affect bradyzoite gene expression (Radke et al., 2013; Shi et al., n.d.). Most notably, BAG1, the most widely used bradyzoite marker protein also exhibited this specific pattern, indicating its diagnostic utility may be limited to *Toxoplasma*-like bradyzoite cysts rather than other tissue-cysts forming members of *Sarcocystidae*.

Cluster 7 (*Hammondia*-specific) and Cluster 8 (*Toxoplasma*-specific) contained no characterized bradyzoite regulators. This absence indicates that core bradyzoite differentiation machinery is evolutionarily ancient and conserved, with *Toxoplasma* or *Hammondia*-specific proteins playing minimal roles in fundamental stage conversion processes. It is also possible that, such proteins unique to *T. gondii* stage differentiation is not discovered yet.

Of 34 characterized proteins, 19 (∼56%) fell into broadly conserved clusters (1-4), while 15 (∼44%) showed lineage-specific conservation (clusters 5-6). A clear functional pattern emerged across clusters with proteins having dual roles in both tachyzoites and bradyzoite growth and maintenance (metabolic enzymes, translational factors, cyst wall proteins) localized predominantly to broadly conserved clusters (1, 2, 3, 4), while regulatory proteins (transcription factors, translational regulators, cell cycle controllers governing developmental fate decisions) concentrated in bradyzoite-cyst forming organisms-specific clusters (5, 6). This indicates, basic cellular processes like stress response, metabolism, protein synthesis, are of ancient origin and repurposed through differential regulation, while dedicated regulatory networks controlling developmental commitment (BFD1/BFD2, bradyzoite-enhancing AP2 factors) represent lineage-specific evolution enabling efficient, reversible stage conversion characteristic of chronic *Toxoplasma* infection.

## Summary

*Toxoplasma gondii* persists within hosts as cyst-forming bradyzoites, causing chronic toxoplasmosis. The stress-induced conversion from replicative tachyzoites to bradyzoites involves complex morphological and physiological changes, including reduced replication rates, organellar rearrangement, and metabolic shifts. While previous studies have focused on genetic manipulation of individual players and their impact on stage differentiation, genome-wide techniques to identify new proteins have largely focused on transcriptome analyses of tachyzoites and bradyzoites. This study utilises a novel strategy involving phylogenetic relationships among tissue cyst forming organisms to find conservation patterns of *T. gondii* proteins, and identify new proteins in bradyzoite differentiation.

Our approach relies on genome sequence availability and quality, which varies considerably across *Sarcocystidae*. While we employed ortholog-based comparisons to ensure functional relevance, we acknowledge that sequence assembly quality, particularly for such understudied organisms may affect cluster assignments. Despite these limitations, our comparative phylogenetic approach revealed many novel aspects of parasite biology. For example, this report reveals a fundamental evolutionary principle underlying bradyzoite differentiation in *T. gondii*: proteins involved in basic cellular mechanisms, which are common to both tachyzoite and bradyzoite biology are conserved in all the organisms chosen for the study, whereas proteins involved in specific regulatory network of stage conversion are lineage specific.

Beyond bradyzoite differentiation, the phylogenetic clustering framework has broader utility for investigating lineage-specific adaptations across *Sarcocystidae*. For instance, proteins conserved in *T. gondii* and *Hammondia* (cluster 6) may reveal determinants of their similar hosts and morphology, while proteins that diverge only in *S. neurona* (cluster 4) could explain its unique biology. This comparative approach is particularly valuable given the difficulty of experimentally manipulating most *Sarcocystidae* members, which require specific primary hosts for routine maintenance. Our study thus offers an alternative strategy for these organisms, enabling predictions about biological processes that would otherwise remain inaccessible.

Importantly, our approach identified eight clusters of proteins with distinct conservation patterns, of which cluster 5 contains proteins conserved exclusively in bradyzoite-cyst-forming lineages. Currently known bradyzoite regulators like BFD-1 and BFD-2 along with other regulatory AP2 transcription factors fell into this cluster. These results position the proteins found in cluster 5 as potential novel players in bradyzoite differentiation in *T. gondii*. Further analysis of cluster 5 using transcriptome datasets and proteomics data would shortlist *T. gondii* proteins that play roles in signal transduction pathways of bradyzoite differentiation, that are currently unknown. In conclusion, this report provides a useful dataset for researchers interested in bradyzoite differentiation to identify new proteins and validate existing candidates from the lens of evolutionary biology.

## Acknowledgements

We are grateful to the contributors of datasets and VEuPathDB for making publicly available the genomic and proteomic datasets that were essential to this study. We thank Aditya Sethi for initiating this project during his internship (current affiliation: University of Bristol, UK) and contributing to early conceptual development. We are grateful to P. V. Balaji, Kiran Kondabagil (IIT Bombay) and Dhanasekaran Shanmugam (National Chemical Laboratory, Pune) for their valuable feedback and insights.

## Funding

S. P. acknowledges funding from the Department of Biotechnology, Government of India (project number: BT/PR45380/COT/142/37/2022). A.C.A. acknowledges the Department of Biotechnology, Government of India, for her PhD fellowship.

## Author contributions

A.C.A., R.U., and S.P. designed and optimized the computational pipeline. R.U. provided technical expertise for pipeline optimization. A.C.A. performed the analyses. A.C.A and S.P. prepared the figures and wrote the manuscript. All authors reviewed and approved the final manuscript.

## Notes

### Competing Interest Statement

The authors have declared no competing interest.

### Summary of Updates

The corresponding author's name was revised to Swati Patankar

