## Supplementary figures for "A phylogenetic approach reveals evolutionary aspects and novel genes of bradyzoite conversion in *Toxoplasma gondii*"

**Supplementary file**

### Supplementary figures.

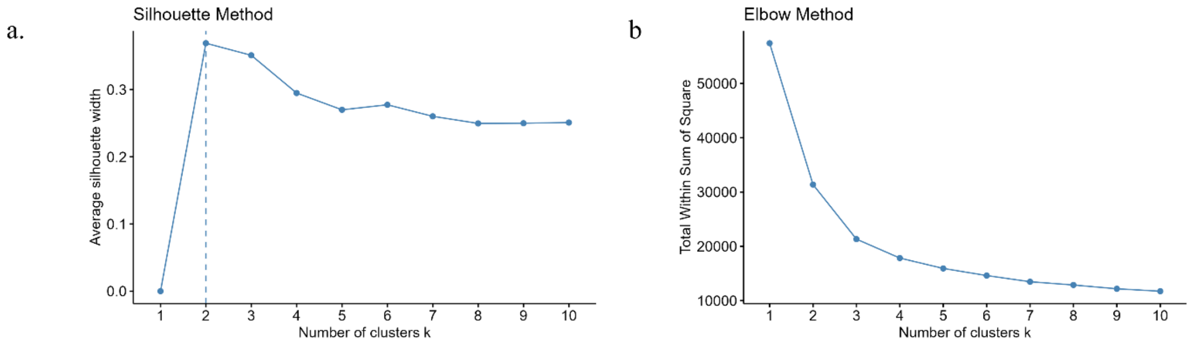


**Figure 1: Silhouette and Elbow method to identify the number of K. (a)** Silhouette method. **(b)** Elbow method. Both method suggested an optimum k value of 2.

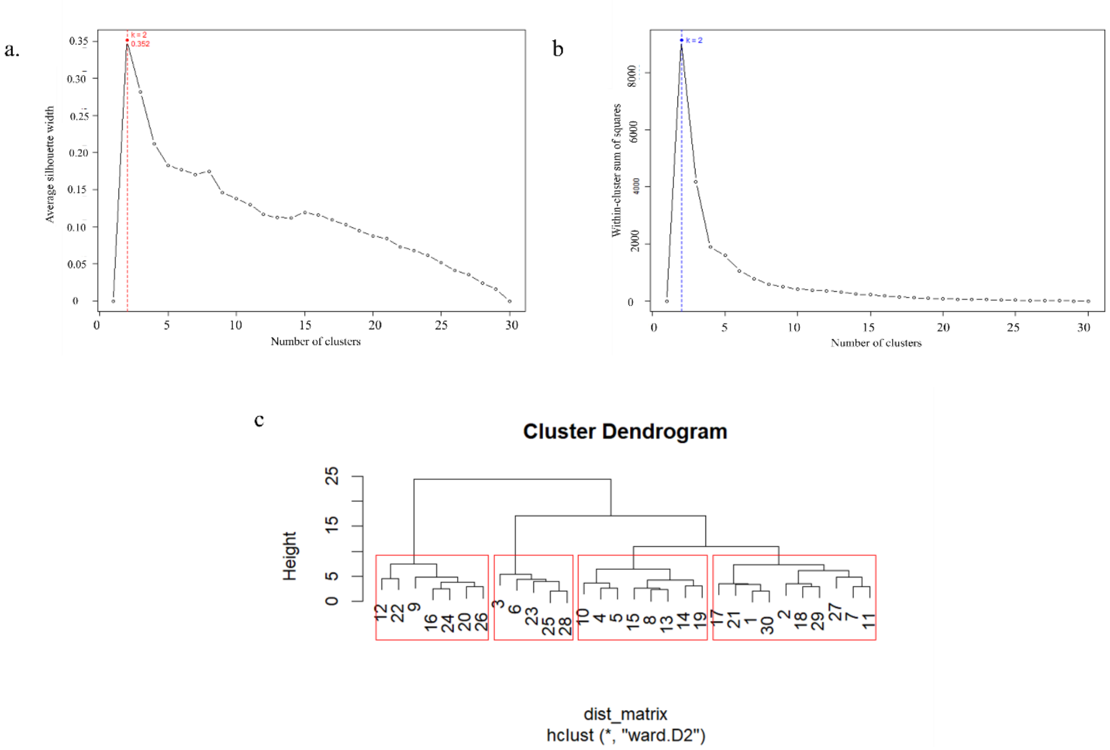


**Figure 2: Silhouette and Elbow method to identify the number of meta clusters. (a)** Silhouette method. **(b)** Elbow method. Both method suggested an optimum value of 2. **(c)** dynamic tree cut based on dendogram topology
