## Supplementary table 1 for "A phylogenetic approach reveals evolutionary aspects and novel genes of bradyzoite conversion in *Toxoplasma gondii*"

| Kmean_cluster | GeneID (all protein TG) | Product Description | Decile_bitHH | Decile_bitNcan | Decile_bitBB | Decilebit_CS | Decilebit_SN3 | Decile bit_Ef | Decile bit_CC | Meta_cluster |
|---|---|---|---|---|---|---|---|---|---|---|
| 28 | TGME49_200330.P453 | basal complex component BCC9 | 10 | 9 | 9 | 9 | 8 | 6 | 7 | 1 |
| 23 | TGME49_200350.P455 | subtilisin SUB3 | 10 | 10 | 9 | 6 | 8 | 8 | 3 | 1 |
| 28 | TGME49_200400.P458 | serine/threonine-protein phosphatase PP2A regulatory subunit PR48 | 10 | 9 | 9 | 9 | 8 | 7 | 7 | 1 |
| 28 | TGME49_201220.P467 | preconoidal ring protein PCR4 | 10 | 9 | 9 | 7 | 7 | 7 | 6 | 1 |
| 28 | TGME49_201250-t26_1-p1 | histone lysine methyltransferase, SET, putative | 10 | 10 | 10 | 9 | 8 | 7 | 7 | 1 |
| 3 | TGME49_201680-t26_1-p1 | eukaryotic initiation factor-3 subunit 10, putative | 10 | 10 | 10 | 9 | 10 | 9 | 8 | 1 |
| 6 | TGME49_201770.P472 | cullin 3, putative | 10 | 10 | 10 | 6 | 8 | 7 | 7 | 1 |
| 25 | TGME49_202120-t26_1-p1 | PH domain-containing protein | 10 | 10 | 10 | 7 | 9 | 8 | 8 | 1 |
| 28 | TGME49_202310-t26_1-p1 | O-sialoglycoprotein endopeptidase | 9 | 9 | 8 | 8 | 6 | 7 | 7 | 1 |
| 3 | TGME49_202370-t26_1-p1 | T-complex protein 1, epsilon subunit (TCP-1-epsilon), putative | 10 | 9 | 9 | 8 | 9 | 9 | 9 | 1 |
| 25 | TGME49_202380-t26_1-p1 | rab GDI alpha, putative | 9 | 9 | 9 | 9 | 7 | 9 | 8 | 1 |
| 6 | TGME49_202460.P491 | diacylglycerol kinase accessory domain (presumed) domain-containing protein | 10 | 9 | 9 | 4 | 8 | 7 | 6 | 1 |
| 23 | TGME49_202500.P492 | glideosome-associated protein with multiple-membrane spans GAPM1A | 8 | 8 | 8 | 8 | 8 | 7 | 3 | 1 |
| 25 | TGME49_202530-t26_1-p1 | aspartate-tRNA ligase | 10 | 9 | 9 | 9 | 9 | 8 | 8 | 1 |
| 25 | TGME49_202540.P493 | 3'5'-cyclic nucleotide phosphodiesterase PDE1 | 10 | 10 | 10 | 10 | 8 | 7 | 7 | 1 |
| 28 | TGME49_202572-t26_1-p1 | ribophorin i protein | 10 | 10 | 9 | 9 | 7 | 7 | 6 | 1 |
| 25 | TGME49_202630-t26_1-p1 | ATP-dependent metallopeptidase HflB subfamily protein | 10 | 10 | 10 | 7 | 8 | 9 | 9 | 1 |
| 6 | TGME49_202650-t26_1-p1 | hypothetical protein | 10 | 10 | 10 | 6 | 8 | 8 | 7 | 1 |
| 25 | TGME49_202680.P499 | cytochrome b-c1 complex subunit 2 | 10 | 9 | 9 | 9 | 9 | 9 | 8 | 1 |
| 28 | TGME49_202920.P505 | p-aminobenzoic acid synthase | 10 | 8 | 10 | 8 | 6 | 8 | 8 | 1 |
| 28 | TGME49_202980-t26_1-p1 | hypothetical protein | 10 | 10 | 10 | 10 | 6 | 6 | 6 | 1 |
| 25 | TGME49_203020-t26_1-p1 | ribosome biogenesis GTPase Der protein | 10 | 10 | 10 | 9 | 7 | 7 | 8 | 1 |
| 28 | TGME49_203110.P511 | citrate synthase, putative | 9 | 9 | 9 | 9 | 7 | 8 | 7 | 1 |
| 3 | TGME49_203135.P512 | dynein heavy chain family protein | 10 | 10 | 10 | 10 | 10 | 10 | 10 | 1 |
| 28 | TGME49_203160-t26_1-p1 | hypothetical protein | 10 | 9 | 9 | 8 | 8 | 7 | 7 | 1 |
| 28 | TGME49_203170.P513 | OB-fold nucleic acid binding domain-containing protein | 9 | 9 | 9 | 8 | 8 | 7 | 6 | 1 |
| 6 | TGME49_203220-t26_1-p1 | DEAD/DEAH box helicase domain-containing protein | 10 | 10 | 10 | 6 | 9 | 8 | 8 | 1 |
| 28 | TGME49_203320-t26_1-p1 | zinc finger in N-recognin protein | 10 | 10 | 10 | 8 | 6 | 7 | 6 | 1 |
| 25 | TGME49_203340.P518 | metallo-beta-lactamase domain-containing protein | 10 | 10 | 10 | 9 | 9 | 9 | 8 | 1 |
| 25 | TGME49_203380-t26_1-p1 | DnaJ domain-containing protein | 10 | 9 | 9 | 8 | 9 | 8 | 7 | 1 |
| 6 | TGME49_203910-t26_1-p1 | TBC domain-containing protein TBC10 | 10 | 10 | 9 | 5 | 8 | 6 | 6 | 1 |
| 23 | TGME49_203940-t26_1-p1 | ABC1 protein | 10 | 9 | 9 | 8 | 7 | 7 | 5 | 1 |
| 28 | TGME49_204020-t26_1-p1 | ribosomal protein RPL8 | 8 | 8 | 8 | 8 | 8 | 7 | 7 | 1 |
| 25 | TGME49_204130.P543 | perforin-like protein PLP1 | 10 | 10 | 9 | 9 | 9 | 8 | 8 | 1 |
| 23 | TGME49_204160.P544 | GYF domain-containing protein | 10 | 10 | 10 | 6 | 8 | 7 | 5 | 1 |
| 28 | TGME49_204280-t26_1-p1 | cell-cycle-associated protein kinase DYRK1 | 10 | 10 | 9 | 7 | 7 | 6 | 7 | 1 |
| 23 | TGME49_204310.P547 | hypothetical protein | 10 | 10 | 10 | 9 | 10 | 10 | 3 | 1 |
| 23 | TGME49_204360-t26_1-p1 | subtilisin SUB4 | 10 | 10 | 9 | 8 | 8 | 7 | 3 | 1 |
| 25 | TGME49_204400.P551 | ATPase synthase subunit alpha, putative | 9 | 10 | 9 | 8 | 9 | 9 | 7 | 1 |
| 28 | TGME49_204480-t26_1-p1 | thioredoxin domain-containing protein | 10 | 9 | 9 | 7 | 8 | 6 | 6 | 1 |
| 3 | TGME49_204560-t26_1-p1 | type I fatty acid synthase, putative | 10 | 10 | 10 | 10 | 10 | 9 | 9 | 1 |

| | | | | | | | | | | |
|---|---|---|---|---|---|---|---|---|---|---|
| 3 | TGME49_205010-t26_1-p1 | splicing factor 3B subunit 1, putative | 10 | 10 | 10 | 10 | 10 | 10 | 10 | 1 |
| 3 | TGME49_205220-t26_1-p1 | U5 snRNP-associated subunit, putative | 10 | 10 | 10 | 10 | 10 | 10 | 10 | 1 |
| 6 | TGME49_205240-t26_1-p1 | cleft lip and palate transmembrane protein 1 (clptm1) protein | 10 | 10 | 10 | 6 | 9 | 8 | 8 | 1 |
| 6 | TGME49_205290.P564 | glucan phosphatase Laforin | 9 | 9 | 9 | 5 | 7 | 7 | 7 | 1 |
| 23 | TGME49_205350.P565 | integral membrane protein, GNS1/SUR4 family protein, putative | 8 | 8 | 8 | 8 | 7 | 7 | 3 | 1 |
| 23 | TGME49_205360-t26_1-p1 | hypothetical protein | 10 | 10 | 10 | 8 | 7 | 7 | 3 | 1 |
| 28 | TGME49_205380.P567 | fructose-1,6-bisphosphatase 1 | 9 | 9 | 9 | 9 | 8 | 7 | 7 | 1 |
| 3 | TGME49_205440-t26_1-p1 | CCT chaperonin gamma subunit | 10 | 10 | 9 | 9 | 9 | 9 | 9 | 1 |
| 3 | TGME49_205470.P571 | translation elongation factor 2 family protein, putative | 10 | 10 | 10 | 10 | 10 | 10 | 10 | 1 |
| 28 | TGME49_205490.P572 | integral membrane protein, putative | 10 | 9 | 9 | 9 | 8 | 6 | 8 | 1 |
| 25 | TGME49_205510-t26_1-p1 | nucleolar protein 5, putative | 9 | 9 | 9 | 9 | 8 | 8 | 8 | 1 |
| 25 | TGME49_205658-t26_1-p1 | F5/8 type C domain-containing protein | 3 | 8 | 9 | 10 | 9 | 9 | 6 | 1 |
| 25 | TGME49_205710-t26_1-p1 | lysine-tRNA ligase | 10 | 10 | 10 | 9 | 9 | 8 | 8 | 1 |
| 28 | TGME49_205740-t26_1-p1 | ER membrane protein complex subunit 1, putative | 10 | 10 | 10 | 9 | 8 | 6 | 6 | 1 |
| 28 | TGME49_205750-t26_1-p1 | histone deacetylase complex subunit Sin3 | 10 | 9 | 10 | 8 | 7 | 6 | 8 | 1 |
| 25 | TGME49_206430.P584 | formin FRM1 | 10 | 10 | 9 | 7 | 9 | 8 | 8 | 1 |
| 25 | TGME49_206470-t26_1-p1 | pyruvate dehydrogenase complex subunit PDH-E3II | 9 | 9 | 9 | 9 | 8 | 8 | 7 | 1 |
| 23 | TGME49_206490.P586 | metacaspase MCA1 | 10 | 8 | 9 | 9 | 8 | 7 | 3 | 1 |
| 25 | TGME49_206510.P588 | toxolysin TLN4 | 10 | 10 | 9 | 9 | 8 | 7 | 8 | 1 |
| 3 | TGME49_206550.P590 | hypothetical protein | 10 | 10 | 10 | 10 | 10 | 9 | 10 | 1 |
| 28 | TGME49_206580-t26_1-p1 | formin FRM2 | 10 | 10 | 10 | 7 | 7 | 7 | 6 | 1 |
| 25 | TGME49_206590.P593 | calcium-dependent protein kinase CDPK2A | 10 | 10 | 9 | 9 | 9 | 9 | 8 | 1 |
| 28 | TGME49_206610.P594 | pyruvate dehydrogenase complex subunit PDH-E2 | 10 | 10 | 9 | 9 | 9 | 7 | 6 | 1 |
| 23 | TGME49_207060.P603 | ribonucleoside-diphosphate reductase small subunit | 9 | 9 | 9 | 8 | 8 | 8 | 3 | 1 |
| 28 | TGME49_207180-t26_1-p1 | indole-3-glycerol phosphate synthase domain-containing protein | 10 | 9 | 8 | 8 | 7 | 6 | 6 | 1 |
| 28 | TGME49_207390-t26_1-p1 | AP-4 complex subunit mu, putative | 9 | 9 | 9 | 8 | 8 | 8 | 6 | 1 |
| 25 | TGME49_207430.P616 | ATP-dependent RNA helicase DDX1, putative | 10 | 10 | 10 | 10 | 9 | 8 | 6 | 1 |
| 28 | TGME49_207440.P617 | ribosomal protein RPS4 | 8 | 8 | 8 | 8 | 7 | 7 | 7 | 1 |
| 25 | TGME49_207450.P618 | RNA-splicing ligase RtcB, putative | 9 | 9 | 9 | 6 | 9 | 9 | 9 | 1 |
| 3 | TGME49_207640-t26_1-p1 | isoleucyl-tRNA synthetase family protein | 10 | 10 | 10 | 9 | 9 | 9 | 9 | 1 |
| 25 | TGME49_207665.P623 | kinesin motor domain-containing protein | 9 | 9 | 9 | 7 | 7 | 8 | 8 | 1 |
| 25 | TGME49_207720-t26_1-p1 | Nrap domain-containing protein | 10 | 10 | 10 | 7 | 8 | 8 | 8 | 1 |
| 23 | TGME49_207760-t26_1-p1 | DnaJ C terminal region domain-containing protein | 9 | 9 | 8 | 6 | 8 | 8 | 3 | 1 |
| 25 | TGME49_207820.P627 | mitogen-activated protein kinase MAPK2 | 10 | 9 | 9 | 8 | 9 | 9 | 8 | 1 |
| 28 | TGME49_207900.P629 | transcription initiation factor TFIIIB | 10 | 9 | 9 | 8 | 8 | 8 | 6 | 1 |
| 28 | TGME49_207910-t26_1-p1 | calcium/proton exchanger CAX | 9 | 9 | 9 | 7 | 8 | 6 | 7 | 1 |
| 28 | TGME49_207990-t26_1-p1 | ribosomal RNA large subunit methyltransferase J protein | 10 | 10 | 10 | 8 | 7 | 7 | 7 | 1 |
| 28 | TGME49_208040-t26_1-p1 | aldo-keto reductase | 10 | 10 | 3 | 9 | 7 | 9 | 6 | 1 |
| 23 | TGME49_208420.P642 | signaling linker factor SLF | 10 | 10 | 10 | 8 | 9 | 8 | 3 | 1 |
| 28 | TGME49_208530-t26_1-p1 | nicotinate phosphoribosyltransferase | 10 | 9 | 9 | 8 | 7 | 6 | 8 | 1 |
| 6 | TGME49_208540-t26_1-p1 | DEAD/DEAH box helicase domain-containing protein | 10 | 10 | 10 | 6 | 7 | 8 | 6 | 1 |
| 25 | TGME49_208580.P650 | DNA ligase 1, putative | 10 | 10 | 10 | 8 | 7 | 9 | 8 | 1 |

| | | | | | | | | | | |
|---|---|---|---|---|---|---|---|---|---|---|
| 25 | TGME49_208590-t26_1-p1 | vacuolar ATP synthase subunit 54kD, putative | 9 | 9 | 9 | 8 | 8 | 8 | 8 | 1 |
| 23 | TGME49_208790-t26_1-p1 | MC family transporter, putative | 8 | 8 | 8 | 8 | 8 | 8 | 5 | 1 |
| 28 | TGME49_208820.P657 | 1-deoxy-D-xylulose-5-phosphate synthase | 10 | 10 | 10 | 9 | 6 | 7 | 6 | 1 |
| 28 | TGME49_209000.P661 | HECT-domain (ubiquitin-transferase) domain-containing protein | 10 | 10 | 10 | 8 | 6 | 7 | 7 | 1 |
| 3 | TGME49_209030.P664 | actin ACT1 | 9 | 9 | 9 | 9 | 9 | 9 | 9 | 1 |
| 28 | TGME49_209050.P665 | Tyrosine kinase-like (TKL) protein | 10 | 9 | 8 | 7 | 7 | 7 | 7 | 1 |
| 6 | TGME49_209130.P670 | regulator of chromosome condensation (RCC1) repeat-containing protein | 10 | 10 | 9 | 6 | 6 | 7 | 6 | 1 |
| 6 | TGME49_209160-t26_1-p1 | myristoyl CoA:protein N-myristoyltransferase | 9 | 9 | 9 | 6 | 8 | 8 | 7 | 1 |
| 6 | TGME49_209260.P673 | cytochrome c oxidase subunit COX5b | 8 | 8 | 8 | 5 | 7 | 7 | 7 | 1 |
| 25 | TGME49_209300.P677 | WD domain, G-beta repeat-containing protein | 10 | 9 | 10 | 9 | 9 | 8 | 6 | 1 |
| 23 | TGME49_209600.P684 | hypothetical protein | 9 | 8 | 8 | 8 | 8 | 7 | 5 | 1 |
| 28 | TGME49_209950.P701 | thioredoxin, putative | 9 | 9 | 8 | 7 | 8 | 7 | 6 | 1 |
| 3 | TGME49_209960.P702 | glycosyltransferase | 10 | 10 | 10 | 10 | 10 | 9 | 10 | 1 |
| 3 | TGME49_210360-t26_1-p1 | DEAD (Asp-Glu-Ala-Asp) box polypeptide 41 family protein | 10 | 10 | 10 | 9 | 9 | 9 | 9 | 1 |
| 28 | TGME49_210790-t26_1-p1 | dihydroorotate dehydrogenase | 10 | 9 | 9 | 8 | 8 | 7 | 7 | 1 |
| 6 | TGME49_210840-t26_1-p1 | arginyl-tRNA synthetase family protein | 10 | 9 | 9 | 5 | 8 | 9 | 9 | 1 |
| 28 | TGME49_210960-t26_1-p1 | replication factor C subunit 4, putative | 9 | 8 | 8 | 8 | 8 | 8 | 7 | 1 |
| 25 | TGME49_211090.P726 | aminotransferase, class V superfamily protein | 9 | 9 | 9 | 9 | 8 | 8 | 8 | 1 |
| 6 | TGME49_211230-t26_1-p1 | eukaryotic initiation factor-2B, alpha subunit, putative | 9 | 9 | 8 | 6 | 7 | 7 | 7 | 1 |
| 23 | TGME49_211270.P729 | sushi domain (scr repeat) domain-containing protein | 10 | 10 | 10 | 10 | 8 | 8 | 5 | 1 |
| 28 | TGME49_211480.P743 | GTP-binding protein engA, putative | 10 | 9 | 9 | 7 | 7 | 7 | 7 | 1 |
| 25 | TGME49_211680.P747 | protein disulfide isomerase | 9 | 9 | 9 | 9 | 8 | 8 | 8 | 1 |
| 3 | TGME49_211690-t26_1-p1 | AAA family protein | 10 | 10 | 10 | 9 | 10 | 10 | 9 | 1 |
| 6 | TGME49_211730-t26_1-p1 | histone lysine methyltransferase SET8 | 10 | 10 | 7 | 6 | 8 | 7 | 6 | 1 |
| 28 | TGME49_211910.P752 | kinesin-related protein 3A, putative | 10 | 9 | 9 | 9 | 6 | 8 | 8 | 1 |
| 25 | TGME49_212130-t26_1-p1 | phospholipase, patatin family protein | 10 | 10 | 10 | 8 | 9 | 8 | 7 | 1 |
| 23 | TGME49_212990.P775 | IMC sutures component ISC3 paralog | 10 | 10 | 9 | 8 | 7 | 6 | 3 | 1 |
| 28 | TGME49_213060.P779 | WD domain, G-beta repeat-containing protein | 10 | 10 | 9 | 8 | 7 | 7 | 7 | 1 |
| 28 | TGME49_213072.P782 | protein SLY1 | 10 | 9 | 9 | 7 | 7 | 8 | 7 | 1 |
| 28 | TGME49_213370-t26_1-p1 | formin FRM3 | 10 | 9 | 9 | 7 | 7 | 7 | 7 | 1 |
| 28 | TGME49_213520-t26_1-p1 | peptidase M20D, amidohydrolase | 9 | 9 | 8 | 8 | 7 | 7 | 7 | 1 |
| 28 | TGME49_213680-t26_1-p1 | MmgE/PrpD family protein | 9 | 9 | 9 | 8 | 8 | 7 | 7 | 1 |
| 28 | TGME49_213720.P803 | dynein regulatory complex subunit 7, putative | 9 | 9 | 9 | 10 | 6 | 6 | 7 | 1 |
| 6 | TGME49_213740-t26_1-p1 | methyltransferase, putative | 10 | 10 | 9 | 5 | 7 | 7 | 6 | 1 |
| 3 | TGME49_213770-t26_1-p1 | Superkiller viralicidic activity 2 family 2, putative | 10 | 10 | 10 | 8 | 10 | 10 | 9 | 1 |
| 25 | TGME49_213870-t26_1-p1 | UBA/TS-N domain-containing protein | 10 | 9 | 10 | 10 | 9 | 8 | 8 | 1 |
| 6 | TGME49_213900.P813 | regulator of chromosome condensation RCC1 | 10 | 9 | 8 | 3 | 7 | 7 | 6 | 1 |
| 28 | TGME49_213970-t26_1-p1 | ribosomal RNA small subunit methyltransferase B, putative | 9 | 8 | 9 | 8 | 6 | 8 | 8 | 1 |
| 23 | TGME49_214090.P815 | signal peptidase | 10 | 10 | 10 | 8 | 7 | 8 | 5 | 1 |
| 28 | TGME49_214200-t26_1-p1 | WD domain, G-beta repeat-containing protein | 10 | 10 | 10 | 8 | 8 | 6 | 7 | 1 |
| 25 | TGME49_214210-t26_1-p1 | rRNA pseudouridine synthase | 9 | 9 | 9 | 6 | 9 | 9 | 9 | 1 |
| 23 | TGME49_214230-t26_1-p1 | Dopey, N-terminal domain-containing protein | 10 | 10 | 9 | 6 | 8 | 7 | 5 | 1 |
| 28 | TGME49_214240-t26_1-p1 | bromodomain-containing protein | 10 | 10 | 10 | 8 | 8 | 7 | 7 | 1 |
| 3 | TGME49_214260-t26_1-p1 | alpha-glucan water dikinase 1, putative | 10 | 10 | 10 | 8 | 9 | 10 | 10 | 1 |

| | | | | | | | | | | |
|---|---|---|---|---|---|---|---|---|---|---|
| 25 | TGME49_214270.P821 | translation initiation factor IF-2, putative | 10 | 10 | 10 | 9 | 10 | 9 | 7 | 1 |
| 23 | TGME49_214300-t26_1-p1 | hypothetical protein | 10 | 10 | 10 | 9 | 7 | 7 | 5 | 1 |
| 25 | TGME49_214320.P822 | facilitative glucose transporter GT1 | 10 | 3 | 9 | 9 | 9 | 8 | 8 | 1 |
| 6 | TGME49_214490-t26_1-p1 | peptidase M16 inactive domain-containing protein | 10 | 10 | 10 | 6 | 7 | 9 | 7 | 1 |
| 23 | TGME49_214520.P827 | general transcription factor IIH polypeptide 4 GTF2H4 | 10 | 9 | 7 | 8 | 7 | 8 | 4 | 1 |
| 23 | TGME49_214560-t26_1-p1 | hypothetical protein | 10 | 10 | 10 | 6 | 7 | 7 | 3 | 1 |
| 28 | TGME49_214580.P829 | tetratricopeptide repeat-containing protein | 10 | 9 | 10 | 9 | 8 | 7 | 6 | 1 |
| 6 | TGME49_214820-t26_1-p1 | DNA repair protein DRE | 9 | 8 | 8 | 3 | 7 | 7 | 7 | 1 |
| 28 | TGME49_214850.P834 | 1-deoxy-D-xylulose 5-phosphate reductoisomerase, putative | 10 | 9 | 9 | 8 | 7 | 6 | 7 | 1 |
| 3 | TGME49_214970-t26_1-p1 | DNA replication licensing factor MCM2 | 10 | 10 | 10 | 10 | 10 | 10 | 7 | 1 |
| 23 | TGME49_215170.P847 | serine/threonine-protein phosphatase 2A catalytic subunit beta isoform, putative | 10 | 9 | 9 | 8 | 7 | 7 | 3 | 1 |
| 3 | TGME49_215260-t26_1-p1 | carbamoylphosphate synthetase | 10 | 10 | 10 | 10 | 9 | 10 | 9 | 1 |
| 6 | TGME49_215280.P850 | succinate dehydrogenase [ubiquinone] iron-sulfur subunit SDHB | 9 | 8 | 8 | 5 | 8 | 7 | 7 | 1 |
| 28 | TGME49_215500.P859 | TPR repeat-containing protein | 10 | 9 | 9 | 8 | 7 | 7 | 7 | 1 |
| 23 | TGME49_215590-t26_1-p1 | flavoprotein subunit of succinate dehydrogenase SDHA | 10 | 9 | 10 | 8 | 9 | 8 | 3 | 1 |
| 25 | TGME49_215740-t26_1-p1 | notchless, putative | 9 | 9 | 9 | 8 | 8 | 7 | 8 | 1 |
| 23 | TGME49_215940.P870 | acetyl-CoA transporter AT1 | 10 | 9 | 9 | 8 | 7 | 7 | 4 | 1 |
| 25 | TGME49_216000-t26_1-p1 | alveolin domain containing intermediate filament IMC3 | 9 | 9 | 9 | 8 | 8 | 8 | 8 | 1 |
| 28 | TGME49_216070.P875 | transmembrane protein TqsA-like, putative | 10 | 10 | 10 | 9 | 7 | 7 | 6 | 1 |
| 23 | TGME49_216080.P876 | apical complex lysine methyltransferase | 10 | 9 | 9 | 9 | 8 | 8 | 3 | 1 |
| 28 | TGME49_216170.P878 | cysteine desulfurase NFS2 | 10 | 9 | 8 | 7 | 7 | 7 | 6 | 1 |
| 25 | TGME49_216210-t26_1-p1 | hypothetical protein | 10 | 10 | 10 | 10 | 7 | 8 | 7 | 1 |
| 28 | TGME49_216430-t26_1-p1 | TBC domain-containing protein TBC7 | 10 | 10 | 8 | 8 | 8 | 7 | 7 | 1 |
| 25 | TGME49_216590-t26_1-p1 | HEAT repeat-containing protein | 10 | 10 | 10 | 9 | 9 | 9 | 6 | 1 |
| 23 | TGME49_216620.P890 | EF hand domain-containing protein | 10 | 10 | 10 | 10 | 10 | 10 | 3 | 1 |
| 28 | TGME49_216630-t26_1-p1 | trigger factor protein, putative | 10 | 10 | 9 | 9 | 9 | 6 | 6 | 1 |
| 25 | TGME49_216650.P891 | mature soluble cytoskeleton protein MSC1A | 9 | 9 | 8 | 8 | 8 | 8 | 8 | 1 |
| 28 | TGME49_216670-t26_1-p1 | FUSE-binding protein 2 / KH-type splicing regulatory protein | 10 | 10 | 9 | 9 | 7 | 6 | 6 | 1 |
| 23 | TGME49_216730-t26_1-p1 | DNA replication licensing factor MCM3, putative | 10 | 10 | 10 | 9 | 10 | 9 | 3 | 1 |
| 3 | TGME49_216790-t26_1-p1 | ATP-binding cassette sub-family E member 1, putative | 10 | 10 | 10 | 10 | 10 | 7 | 9 | 1 |
| 25 | TGME49_216820.P894 | transporter facilitator protein TFP1 | 10 | 9 | 9 | 9 | 9 | 9 | 8 | 1 |
| 3 | TGME49_216860-t26_1-p1 | DEAD (Asp-Glu-Ala-Asp) box polypeptide DDX39 | 9 | 9 | 9 | 9 | 9 | 9 | 9 | 1 |
| 3 | TGME49_216870-t26_1-p1 | DNA excision repair helicase | 10 | 7 | 10 | 10 | 10 | 9 | 8 | 1 |
| 25 | TGME49_216880.P895 | POC1 centriolar protein, putative | 8 | 8 | 8 | 8 | 8 | 8 | 8 | 1 |
| 28 | TGME49_216890-t26_1-p1 | hypothetical protein | 10 | 9 | 9 | 8 | 8 | 6 | 6 | 1 |
| 25 | TGME49_216950-t26_1-p1 | helicase | 10 | 10 | 10 | 9 | 8 | 8 | 7 | 1 |
| 28 | TGME49_216960-t26_1-p1 | elongation factor Tu GTP binding domain-containing protein | 10 | 9 | 9 | 8 | 6 | 7 | 7 | 1 |
| 6 | TGME49_216970-t26_1-p1 | coronin, putative | 10 | 9 | 9 | 6 | 8 | 8 | 8 | 1 |
| 28 | TGME49_217020-t26_1-p1 | ATPase, AFG1 family protein | 10 | 9 | 9 | 7 | 8 | 8 | 7 | 1 |

| | | | | | | | | | | |
|---|---|---|---|---|---|---|---|---|---|---|
| 23 | TGME49_217050-t26_1-p1 | ADA2-A transcriptional co-activator SAGA component | 10 | 10 | 9 | 9 | 8 | 7 | 5 | 1 |
| 25 | TGME49_217350.P900 | N6-adenosine-methyltransferase subunit METTL3 | 10 | 9 | 9 | 8 | 8 | 8 | 8 | 1 |
| 3 | TGME49_217460-t26_1-p1 | glutaminyl-tRNA synthetase (GlnRS) | 10 | 9 | 10 | 10 | 9 | 9 | 9 | 1 |
| 3 | TGME49_217600-t26_1-p1 | calcium-dependent protein kinase CDPK9 | 10 | 10 | 10 | 8 | 8 | 9 | 9 | 1 |
| 6 | TGME49_217820-t26_1-p1 | PCI domain-containing protein | 9 | 9 | 9 | 3 | 7 | 8 | 8 | 1 |
| 3 | TGME49_217910-t26_1-p1 | DNA polymerase (pol2) superfamily protein | 10 | 10 | 10 | 10 | 8 | 10 | 9 | 1 |
| 25 | TGME49_218200.P930 | UDP-sugar pyrophospharylase | 10 | 9 | 9 | 9 | 9 | 8 | 8 | 1 |
| 3 | TGME49_218210.P931 | Chromosome-associated kinesin KLP1, putative | 10 | 10 | 10 | 9 | 10 | 10 | 9 | 1 |
| 28 | TGME49_218220.P933 | CDK-related protein kinase CRK2 | 8 | 8 | 8 | 8 | 8 | 8 | 7 | 1 |
| 28 | TGME49_218280.P936 | mitochondrial import receptor subunit TOM40 | 9 | 9 | 8 | 8 | 8 | 7 | 7 | 1 |
| 25 | TGME49_218290.P937 | WD domain, G-beta repeat-containing protein | 10 | 9 | 10 | 10 | 7 | 8 | 7 | 1 |
| 28 | TGME49_218410-t26_1-p1 | ribosomal protein RPP0 | 8 | 8 | 8 | 8 | 7 | 7 | 7 | 1 |
| 25 | TGME49_218420-t26_1-p1 | pre-mRNA-splicing factor PRP46, putative | 10 | 9 | 9 | 7 | 9 | 8 | 8 | 1 |
| 28 | TGME49_218550-t26_1-p1 | PIK3R4 kinase-related protein | 10 | 10 | 9 | 9 | 7 | 7 | 7 | 1 |
| 25 | TGME49_218610-t26_1-p1 | ATPase (DUF699) protein | 10 | 10 | 10 | 9 | 9 | 9 | 7 | 1 |
| 25 | TGME49_218720.P946 | calcium-dependent protein kinase CDPK6 | 10 | 9 | 9 | 7 | 9 | 7 | 9 | 1 |
| 28 | TGME49_218840-t26_1-p1 | mutS domain protein | 10 | 10 | 10 | 9 | 7 | 6 | 6 | 1 |
| 25 | TGME49_218930.P956 | BTB/POZ domain-containing protein | 9 | 9 | 9 | 9 | 8 | 8 | 9 | 1 |
| 25 | TGME49_219070.P959 | cyclic nucleotide-binding domain-containing protein | 10 | 10 | 10 | 8 | 8 | 8 | 8 | 1 |
| 25 | TGME49_219150-t26_1-p1 | zinc finger, zz type domain-containing protein | 10 | 10 | 9 | 9 | 8 | 8 | 7 | 1 |
| 25 | TGME49_219200-t26_1-p1 | DEAD/DEAH box RNA helicase family protein | 10 | 10 | 9 | 9 | 9 | 8 | 6 | 1 |
| 28 | TGME49_219270-t26_1-p1 | glideosome-associated protein with multiple-membrane spans GAPM2A | 9 | 9 | 8 | 8 | 6 | 7 | 7 | 1 |
| 6 | TGME49_219280-t26_1-p1 | actin like protein ALP1 | 9 | 9 | 9 | 6 | 6 | 8 | 8 | 1 |
| 25 | TGME49_219310-t26_1-p1 | DnaK family protein | 10 | 10 | 10 | 9 | 9 | 8 | 8 | 1 |
| 25 | TGME49_219320-t26_1-p1 | gliding-associated protein GAP50 | 9 | 9 | 8 | 8 | 8 | 8 | 8 | 1 |
| 28 | TGME49_219510.P973 | GTP binding protein 7 isoform 2 family protein, putative | 9 | 9 | 8 | 8 | 7 | 7 | 6 | 1 |
| 25 | TGME49_219520-t26_1-p1 | histone arginine methyltransferase PRMT1 | 9 | 9 | 9 | 8 | 8 | 8 | 8 | 1 |
| 3 | TGME49_219540.P975 | cytosolic tRNA-Ala synthetase | 10 | 10 | 10 | 10 | 9 | 9 | 10 | 1 |
| 6 | TGME49_219550-t26_1-p1 | dihydrolipoyllysine-residue succinyltransferase component of oxoglutarate dehydrogenase | 9 | 9 | 8 | 4 | 7 | 8 | 6 | 1 |
| 6 | TGME49_219590-t26_1-p1 | RuvB family 1 protein | 9 | 9 | 9 | 6 | 8 | 8 | 8 | 1 |
| 23 | TGME49_219682-t26_1-p1 | pyruvate dehydrogenase kinase, putative | 10 | 9 | 9 | 8 | 7 | 7 | 5 | 1 |
| 3 | TGME49_219700-t26_1-p1 | DNA replication licensing factor MCM4, putative | 10 | 10 | 10 | 10 | 10 | 9 | 9 | 1 |
| 28 | TGME49_219790-t26_1-p1 | pre-mRNA processing factor PRP3 | 9 | 9 | 9 | 8 | 7 | 7 | 6 | 1 |
| 25 | TGME49_219800-t26_1-p1 | vacuolar ATP synthase subunit b, putative | 9 | 9 | 9 | 9 | 9 | 9 | 8 | 1 |
| 3 | TGME49_219850-t26_1-p1 | prolyl-tRNA synthetase (ProRS) | 10 | 10 | 10 | 7 | 9 | 9 | 9 | 1 |
| 3 | TGME49_219860-t26_1-p1 | DNA replication licensing factor MCM6, putative | 10 | 10 | 10 | 10 | 10 | 9 | 9 | 1 |
| 25 | TGME49_220100-t26_1-p1 | phosphoribosylpyrophosphate synthetase | 9 | 9 | 9 | 8 | 8 | 8 | 8 | 1 |
| 28 | TGME49_220270.P992 | alveolin domain containing intermediate filament IMC6 | 9 | 9 | 8 | 7 | 7 | 7 | 7 | 1 |
| 28 | TGME49_220600.P1003 | apicomplexan amino acid transporter ApiAT3-3 | 10 | 10 | 9 | 9 | 6 | 8 | 7 | 1 |
| 25 | TGME49_220860-t26_1-p1 | DEAD/DEAH box helicase | 10 | 9 | 9 | 9 | 8 | 8 | 7 | 1 |
| 3 | TGME49_220930.P1008 | IMC sutures component ISC3 | 10 | 10 | 9 | 9 | 8 | 9 | 9 | 1 |
| 28 | TGME49_221170-t26_1-p1 | CAAX metallo endopeptidase | 9 | 9 | 9 | 9 | 8 | 6 | 8 | 1 |
| 25 | TGME49_221250.P1015 | hypothetical protein | 10 | 10 | 10 | 9 | 7 | 8 | 8 | 1 |
| 3 | TGME49_221320-t26_1-p1 | acetyl-CoA carboxylase ACC1 | 10 | 10 | 10 | 10 | 9 | 9 | 9 | 1 |

| | | | | | | | | | | |
|---|---|---|---|---|---|---|---|---|---|---|
| 28 | TGME49_221330-t26_1-p1 | DNA gyrase subunit A | 10 | 10 | 10 | 7 | 7 | 7 | 7 | 1 |
| 3 | TGME49_221660-t26_1-p1 | DEAD/DEAH box helicase domain-containing protein | 10 | 10 | 10 | 10 | 10 | 9 | 8 | 1 |
| 3 | TGME49_221670-t26_1-p1 | transcriptional elongation factor FACT140 | 10 | 10 | 10 | 10 | 10 | 10 | 8 | 1 |
| 23 | TGME49_221690-t26_1-p1 | hypothetical protein | 10 | 10 | 10 | 9 | 7 | 6 | 4 | 1 |
| 6 | TGME49_221950.P1044 | splicing factor 3A subunit 3, putative | 10 | 9 | 9 | 3 | 9 | 8 | 6 | 1 |
| 28 | TGME49_222020.P1045 | phosphoglycerate kinase PGKII | 9 | 9 | 8 | 8 | 8 | 7 | 6 | 1 |
| 28 | TGME49_222140.P1049 | DEAD/DEAH box helicase domain-containing protein | 10 | 8 | 9 | 8 | 7 | 7 | 7 | 1 |
| 28 | TGME49_222160-t26_1-p1 | aldehyde dehydrogenase | 9 | 9 | 9 | 9 | 8 | 6 | 8 | 1 |
| 28 | TGME49_222210-t26_1-p1 | SPFH domain / Band 7 family protein | 9 | 8 | 8 | 8 | 8 | 7 | 7 | 1 |
| 3 | TGME49_222380.P1057 | importin-beta N-terminal domain-containing protein | 10 | 10 | 10 | 10 | 10 | 10 | 10 | 1 |
| 25 | TGME49_222430.P1060 | HECT-domain (ubiquitin-transferase) domain-containing protein | 10 | 10 | 10 | 7 | 7 | 9 | 7 | 1 |
| 25 | TGME49_222800-t26_1-p1 | glycogen synthase, putative | 10 | 10 | 10 | 8 | 10 | 7 | 9 | 1 |
| 3 | TGME49_222860.P1065 | eukaryotic translation initiation factor 3 subunit B, putative | 10 | 10 | 10 | 10 | 10 | 9 | 9 | 1 |
| 25 | TGME49_222960-t26_1-p1 | SCY kinase-related protein (incomplete catalytic triad) | 10 | 10 | 10 | 9 | 8 | 8 | 7 | 1 |
| 3 | TGME49_223390-t26_1-p1 | activating signal cointegrator 1 complex subunit 3 family 1 ASCC3L1, putative | 10 | 10 | 10 | 10 | 10 | 10 | 10 | 1 |
| 3 | TGME49_223480.P1081 | sushi domain (scr repeat) domain-containing protein | 10 | 10 | 10 | 10 | 9 | 10 | 9 | 1 |
| 28 | TGME49_223510.P1084 | serine hydrolase ASH3 | 9 | 8 | 8 | 8 | 7 | 7 | 6 | 1 |
| 28 | TGME49_223520-t26_1-p1 | EF hand domain-containing protein | 10 | 8 | 8 | 7 | 8 | 7 | 7 | 1 |
| 25 | TGME49_223540-t26_1-p1 | importin-beta N-terminal domain-containing protein | 10 | 10 | 10 | 10 | 8 | 9 | 8 | 1 |
| 3 | TGME49_223710-t26_1-p1 | tetratricopeptide repeat-containing protein | 10 | 10 | 10 | 10 | 8 | 9 | 9 | 1 |
| 28 | TGME49_223790-t26_1-p1 | hypothetical protein | 10 | 10 | 9 | 7 | 6 | 7 | 7 | 1 |
| 3 | TGME49_223840.P1093 | ATP-citrate lyase, putative | 10 | 10 | 10 | 10 | 10 | 10 | 10 | 1 |
| 25 | TGME49_223920-t26_1-p1 | rhoptry neck protein RON3 | 10 | 10 | 10 | 10 | 9 | 9 | 7 | 1 |
| 3 | TGME49_223970.P1101 | translation elongation and release factors (gtpases), putative | 10 | 10 | 10 | 9 | 8 | 9 | 9 | 1 |
| 28 | TGME49_224060-t26_1-p1 | thioredoxin, putative | 10 | 9 | 9 | 9 | 6 | 8 | 8 | 1 |
| 3 | TGME49_224190-t26_1-p1 | cation-transporting atpase family protein | 10 | 10 | 10 | 7 | 9 | 10 | 10 | 1 |
| 28 | TGME49_224280-t26_1-p1 | CPSF A subunit region protein | 10 | 10 | 10 | 9 | 8 | 7 | 6 | 1 |
| 3 | TGME49_224350.P1116 | aminopeptidase N2 | 10 | 10 | 10 | 10 | 10 | 9 | 10 | 1 |
| 28 | TGME49_224480.P1119 | cell-cycle-associated protein kinase CLK, putative | 10 | 9 | 8 | 8 | 8 | 8 | 7 | 1 |
| 28 | TGME49_224600-t26_1-p1 | GTP binding protein | 9 | 9 | 9 | 9 | 8 | 8 | 6 | 1 |
| 3 | TGME49_224840.P1131 | 3'5'-cyclic nucleotide phosphodiesterase PDE11 | 10 | 10 | 10 | 10 | 10 | 9 | 9 | 1 |
| 25 | TGME49_224850.P1133 | polyadenylate-binding protein PABC | 10 | 10 | 10 | 10 | 8 | 9 | 8 | 1 |
| 3 | TGME49_224950-t26_1-p1 | calcium-dependent protein kinase CDPK5 | 10 | 10 | 10 | 10 | 9 | 9 | 9 | 1 |
| 28 | TGME49_225000.P1140 | hypothetical protein | 10 | 10 | 9 | 7 | 6 | 7 | 7 | 1 |
| 28 | TGME49_225020.P1142 | CPH1-interacting protein CIP3 | 10 | 10 | 8 | 9 | 6 | 7 | 6 | 1 |
| 25 | TGME49_225050.P1144 | adenosylhomocysteinase, putative | 9 | 9 | 9 | 9 | 9 | 7 | 9 | 1 |
| 25 | TGME49_225125-t26_1-p1 | pre-rRNA processing protein | 10 | 10 | 10 | 10 | 9 | 6 | 8 | 1 |
| 3 | TGME49_225260.P1160 | DNA-directed RNA polymerase II RPB1 | 10 | 10 | 10 | 10 | 10 | 10 | 10 | 1 |
| 23 | TGME49_225310.P1163 | ARF1-directed GTPase-activating protein, putative | 9 | 8 | 9 | 8 | 7 | 6 | 3 | 1 |
| 28 | TGME49_225360-t26_1-p1 | hypothetical protein | 10 | 9 | 9 | 7 | 7 | 6 | 6 | 1 |

| | | | | | | | | | | |
|---|---|---|---|---|---|---|---|---|---|---|
| 3 | TGME49_225490.P1171 | calcium-dependent protein kinase CDPK2 | 10 | 10 | 10 | 10 | 9 | 9 | 9 | 1 |
| 23 | TGME49_225510.P1172 | RAP domain-containing protein | 10 | 9 | 9 | 9 | 8 | 6 | 5 | 1 |
| 28 | TGME49_225550-t26_1-p1 | phosphatidylserine decarboxylase | 9 | 9 | 9 | 8 | 8 | 6 | 6 | 1 |
| 28 | TGME49_225570.P1174 | hypothetical protein | 10 | 10 | 9 | 9 | 8 | 6 | 6 | 1 |
| 25 | TGME49_226020.P1192 | transporter, major facilitator family protein | 10 | 10 | 10 | 9 | 9 | 6 | 9 | 1 |
| 23 | TGME49_226030.P1193 | cAMP-dependent protein kinase subunit PKAc1 | 9 | 9 | 8 | 9 | 8 | 8 | 3 | 1 |
| 25 | TGME49_226080-t26_1-p1 | polyA polymerase | 10 | 9 | 9 | 9 | 9 | 9 | 8 | 1 |
| 3 | TGME49_226250.P1198 | DEAD (Asp-Glu-Ala-Asp) box polypeptide DDX3X | 10 | 10 | 10 | 9 | 9 | 9 | 9 | 1 |
| 25 | TGME49_226340.P1202 | hypothetical protein | 10 | 10 | 10 | 9 | 7 | 8 | 7 | 1 |
| 28 | TGME49_226400-t26_1-p1 | lipoic acid synthase LIPA | 9 | 8 | 8 | 8 | 8 | 7 | 7 | 1 |
| 28 | TGME49_226440-t26_1-p1 | SWI2/SNF2-containing protein RAD16 | 10 | 10 | 10 | 8 | 8 | 6 | 6 | 1 |
| 25 | TGME49_226480.P1210 | DNA topoisomerase family protein | 10 | 10 | 10 | 9 | 7 | 9 | 8 | 1 |
| 3 | TGME49_226510-t26_1-p1 | Sec23/Sec24 trunk domain-containing protein | 10 | 10 | 10 | 10 | 9 | 9 | 8 | 1 |
| 23 | TGME49_226660.P1220 | m6A methyltransferase component VIRMA | 10 | 10 | 10 | 9 | 7 | 6 | 4 | 1 |
| 3 | TGME49_226730-t26_1-p1 | aconitate hydratase ACN/IRP | 10 | 10 | 10 | 10 | 10 | 9 | 10 | 1 |
| 25 | TGME49_226830.P1225 | DnaK family protein | 10 | 10 | 10 | 9 | 9 | 8 | 8 | 1 |
| 28 | TGME49_226870.P1227 | tubulin gamma chain | 9 | 9 | 9 | 9 | 8 | 6 | 8 | 1 |
| 3 | TGME49_226960-t26_1-p1 | phosphofructokinase PFKII | 10 | 10 | 10 | 10 | 10 | 9 | 9 | 1 |
| 3 | TGME49_227290-t26_1-p1 | histone deacetylase HDAC3 | 9 | 9 | 9 | 9 | 9 | 9 | 9 | 1 |
| 25 | TGME49_227360-t26_1-p1 | ribosomal protein RPL3 | 9 | 9 | 9 | 9 | 9 | 9 | 8 | 1 |
| 25 | TGME49_227370.P1245 | hydrolase CocE/NonD family protein | 10 | 10 | 10 | 10 | 7 | 8 | 7 | 1 |
| 28 | TGME49_227420-t26_1-p1 | 4-hydroxy-3-methylbut-2-enyl diphosphate reductase | 9 | 8 | 8 | 8 | 8 | 7 | 7 | 1 |
| 23 | TGME49_227610.P1254 | hypothetical protein | 10 | 10 | 10 | 8 | 9 | 6 | 3 | 1 |
| 6 | TGME49_227830.P1261 | mitochondrial inner membrane translocase TIM44, putative | 9 | 9 | 8 | 6 | 8 | 7 | 6 | 1 |
| 25 | TGME49_227948.P1267 | toxolysin TLN2 | 3 | 10 | 10 | 10 | 9 | 9 | 8 | 1 |
| 25 | TGME49_227960-t26_1-p1 | PCI domain-containing protein | 9 | 9 | 9 | 9 | 9 | 8 | 7 | 1 |
| 25 | TGME49_228070.P1273 | zinc finger, C2H2 type domain-containing protein | 10 | 10 | 10 | 9 | 9 | 8 | 7 | 1 |
| 25 | TGME49_228080-t26_1-p1 | dihydrouridine synthase (dus) protein | 10 | 10 | 9 | 8 | 8 | 8 | 8 | 1 |
| 6 | TGME49_228100-t26_1-p1 | hypothetical protein | 10 | 10 | 10 | 3 | 7 | 7 | 6 | 1 |
| 3 | TGME49_228160.P1278 | acid phosphatase | 10 | 10 | 10 | 10 | 10 | 9 | 9 | 1 |
| 6 | TGME49_228210.P1283 | 26S proteasome regulatory subunit | 9 | 9 | 9 | 5 | 9 | 9 | 9 | 1 |
| 6 | TGME49_228250.P1287 | elongation factor Tu GTP binding domain-containing protein | 10 | 10 | 10 | 6 | 8 | 6 | 7 | 1 |
| 23 | TGME49_228400.P1293 | WD domain, G-beta repeat-containing protein | 10 | 9 | 9 | 9 | 8 | 7 | 4 | 1 |
| 25 | TGME49_228420.P1295 | cAMP-dependent protein kinase subunit PKAc2 | 9 | 7 | 8 | 8 | 8 | 8 | 8 | 1 |
| 25 | TGME49_228750.P1305 | calcium dependent protein kinase CDPK7 | 10 | 10 | 9 | 7 | 9 | 9 | 8 | 1 |
| 28 | TGME49_229000-t26_1-p1 | kelch repeat-containing protein | 10 | 9 | 9 | 8 | 8 | 7 | 7 | 1 |
| 25 | TGME49_229020.P1308 | CDK-related protein kinase CRK5 | 10 | 10 | 9 | 8 | 8 | 8 | 8 | 1 |
| 23 | TGME49_229170.P1313 | formate/nitrite transporter FNT3 | 9 | 8 | 8 | 7 | 7 | 7 | 3 | 1 |
| 3 | TGME49_229180-t26_1-p1 | HEAT repeat-containing protein | 10 | 10 | 10 | 10 | 10 | 9 | 9 | 1 |
| 28 | TGME49_229460.P1325 | SWI2/SNF2-containing protein | 10 | 10 | 8 | 8 | 7 | 7 | 7 | 1 |
| 28 | TGME49_229620.P1328 | hypothetical protein | 10 | 9 | 9 | 9 | 8 | 7 | 6 | 1 |
| 28 | TGME49_229900.P1342 | MACRO domain-containing protein | 9 | 9 | 9 | 8 | 8 | 7 | 6 | 1 |
| 28 | TGME49_229940.P1345 | cyclophilin, putative | 10 | 9 | 9 | 9 | 7 | 8 | 6 | 1 |
| 28 | TGME49_229950-t26_1-p1 | 26S proteasome regulatory subunit 6b, putative | 9 | 9 | 9 | 9 | 8 | 6 | 8 | 1 |
| 3 | TGME49_229990-t26_1-p1 | T-complex protein 1 subunit alpha, putative | 10 | 9 | 9 | 9 | 9 | 9 | 9 | 1 |
| 28 | TGME49_230010.P1346 | hypothetical protein | 10 | 9 | 10 | 8 | 7 | 7 | 8 | 1 |

| | | | | | | | | | | |
|---|---|---|---|---|---|---|---|---|---|---|
| 25 | TGME49_230210-t26_1-p1 | alveolin domain containing intermediate filament IMC10 | 10 | 9 | 9 | 8 | 8 | 8 | 8 | 1 |
| 6 | TGME49_230340.P1364 | apical annuli protein AAP4 | 10 | 9 | 9 | 3 | 8 | 6 | 6 | 1 |
| 25 | TGME49_230350.P1365 | rhoptry neck protein RON11 | 10 | 10 | 10 | 8 | 9 | 9 | 8 | 1 |
| 3 | TGME49_230420.P1367 | sarco/endoplasmic reticulum Ca2+-ATPase | 10 | 10 | 10 | 7 | 10 | 10 | 10 | 1 |
| 25 | TGME49_230450-t26_1-p1 | bifunctional GMP synthase/glutamine amidotransferase protein | 10 | 9 | 9 | 9 | 9 | 8 | 8 | 1 |
| 25 | TGME49_230460-t26_1-p1 | Dpy-30 motif protein | 10 | 9 | 10 | 10 | 9 | 6 | 8 | 1 |
| 28 | TGME49_230690.P1382 | phosphatidate phosphatase LIPIN | 10 | 10 | 9 | 8 | 7 | 7 | 8 | 1 |
| 23 | TGME49_230920.P1390 | AP-2 complex subunit mu | 10 | 9 | 9 | 9 | 8 | 6 | 5 | 1 |
| 3 | TGME49_230960.P1392 | splicing factor 3b, subunit 3, putative | 10 | 10 | 10 | 10 | 10 | 10 | 9 | 1 |
| 28 | TGME49_231000.P1396 | START domain-containing protein | 10 | 10 | 10 | 9 | 6 | 7 | 7 | 1 |
| 28 | TGME49_231060.P1400 | subtilisin SUB9 | 9 | 9 | 9 | 8 | 7 | 7 | 7 | 1 |
| 28 | TGME49_231070.P1401 | basal complex component BCC2 | 10 | 9 | 9 | 7 | 6 | 7 | 7 | 1 |
| 25 | TGME49_231170.P1407 | structural maintenance of chromosomes protein SMC4, putative | 10 | 10 | 10 | 9 | 9 | 9 | 6 | 1 |
| 6 | TGME49_231190-t26_1-p1 | arsenite-activated ATPase family protein | 9 | 8 | 8 | 5 | 7 | 8 | 8 | 1 |
| 3 | TGME49_231210.P1409 | sarcalumenin/eps15 family protein | 10 | 10 | 9 | 8 | 9 | 9 | 9 | 1 |
| 3 | TGME49_231350.P1412 | glucosamine-fructose-6-phosphate aminotransferase | 10 | 10 | 10 | 9 | 9 | 9 | 9 | 1 |
| 25 | TGME49_231430-t26_1-p1 | oligosaccharyl transferase stt3 protein, putative | 10 | 10 | 10 | 8 | 9 | 9 | 7 | 1 |
| 25 | TGME49_231480.P1420 | GCN1, putative | 10 | 10 | 10 | 10 | 8 | 9 | 7 | 1 |
| 25 | TGME49_231600.P1421 | HEAT repeat-containing protein | 10 | 10 | 10 | 10 | 8 | 9 | 8 | 1 |
| 25 | TGME49_231630-t26_1-p1 | alveolin domain containing intermediate filament IMC4 | 9 | 9 | 8 | 8 | 8 | 8 | 8 | 1 |
| 25 | TGME49_231640-t26_1-p1 | alveolin domain containing intermediate filament IMC1 | 10 | 9 | 9 | 9 | 9 | 8 | 8 | 1 |
| 28 | TGME49_231850-t26_1-p1 | protein phosphatase PPM13, putative | 9 | 8 | 8 | 8 | 8 | 7 | 6 | 1 |
| 6 | TGME49_231870.P1427 | peroxisome biogenesis protein PEX5 | 10 | 9 | 9 | 6 | 7 | 8 | 8 | 1 |
| 28 | TGME49_231890.P1429 | beta-ketoacyl-acyl carrier protein synthase III, putative | 10 | 9 | 8 | 8 | 7 | 7 | 7 | 1 |
| 28 | TGME49_231900-t26_1-p1 | acyl-CoA dehydrogenase domain-containing protein | 9 | 6 | 9 | 8 | 7 | 8 | 6 | 1 |
| 3 | TGME49_231970.P1434 | pre-mRNA processing splicing factor PRP8 | 10 | 10 | 10 | 10 | 10 | 10 | 10 | 1 |
| 28 | TGME49_232040.P1441 | cell division control protein CDC50.2 | 10 | 9 | 9 | 8 | 8 | 7 | 7 | 1 |
| 25 | TGME49_232050.P1442 | DnaJ domain-containing protein | 10 | 10 | 9 | 8 | 8 | 8 | 7 | 1 |
| 3 | TGME49_232080.P1444 | VPS13 domain-containing protein | 10 | 10 | 10 | 10 | 8 | 9 | 9 | 1 |
| 28 | TGME49_232090-t26_1-p1 | 3-hydroxyacyl-CoA dehydrogenase, NAD binding domain-containing protein | 8 | 8 | 8 | 8 | 7 | 7 | 7 | 1 |
| 3 | TGME49_232170.P1450 | hypothetical protein | 10 | 10 | 10 | 10 | 9 | 8 | 9 | 1 |
| 25 | TGME49_232250-t26_1-p1 | catalase | 9 | 9 | 9 | 9 | 8 | 9 | 9 | 1 |
| 28 | TGME49_232340.P1459 | protein phosphatase PPM2A | 9 | 9 | 9 | 9 | 7 | 6 | 6 | 1 |
| 28 | TGME49_232380.P1462 | WD domain, G-beta repeat-containing protein | 10 | 10 | 10 | 7 | 9 | 7 | 6 | 1 |
| 28 | TGME49_232440-t26_1-p1 | Brf1p family coiled coil protein | 9 | 9 | 9 | 9 | 8 | 7 | 7 | 1 |
| 3 | TGME49_232450.P1467 | SWI2/SNF2-containing protein RAD54 | 10 | 10 | 10 | 8 | 10 | 9 | 9 | 1 |
| 25 | TGME49_232590.P1473 | glutamate-cysteine ligase, catalytic subunit domain-containing protein | 10 | 10 | 10 | 8 | 6 | 8 | 8 | 1 |
| 23 | TGME49_232710-t26_1-p1 | ribosomal protein RPS3A | 8 | 8 | 8 | 8 | 8 | 7 | 3 | 1 |
| 28 | TGME49_232730.P1481 | acyl-CoA:diacylglycerol acyltransferase 1-related enzyme | 9 | 7 | 9 | 9 | 8 | 7 | 7 | 1 |

| | | | | | | | | | | |
|---|---|---|---|---|---|---|---|---|---|---|
| 3 | TGME49_232830.P1484 | V-type proton ATPase subunit a1 | 10 | 10 | 10 | 10 | 9 | 9 | 9 | 1 |
| 25 | TGME49_233000.P1488 | KOW motif domain-containing protein | 10 | 10 | 10 | 9 | 9 | 8 | 8 | 1 |
| 6 | TGME49_233010.P1489 | mitogen-activated protein kinase ERK7 | 9 | 9 | 9 | 3 | 8 | 8 | 8 | 1 |
| 25 | TGME49_233040.P1491 | 3'5'-cyclic nucleotide phosphodiesterase PDE15 | 10 | 10 | 10 | 10 | 10 | 8 | 7 | 1 |
| 23 | TGME49_233090.P1493 | XPG N-terminal domain-containing protein | 10 | 9 | 8 | 8 | 7 | 7 | 4 | 1 |
| 25 | TGME49_233110-t26_1-p1 | inosine-5'-monophosphate dehydrogenase | 9 | 9 | 9 | 9 | 9 | 8 | 9 | 1 |
| 28 | TGME49_233190.P1498 | SKIP/SNW domain-containing protein | 10 | 9 | 9 | 9 | 7 | 7 | 8 | 1 |
| 6 | TGME49_233260.P1502 | GTP-binding protein, putative | 9 | 9 | 9 | 6 | 8 | 8 | 7 | 1 |
| 25 | TGME49_233310.P1504 | peptidase D, putative | 10 | 9 | 9 | 9 | 8 | 8 | 7 | 1 |
| 28 | TGME49_233330-t26_1-p1 | radial spokehead family protein | 9 | 9 | 9 | 9 | 8 | 7 | 7 | 1 |
| 3 | TGME49_233520.P1509 | ATP-dependent RNA helicase | 10 | 10 | 10 | 10 | 10 | 10 | 10 | 1 |
| 28 | TGME49_233680.P1511 | nuclear movement family protein | 9 | 8 | 8 | 8 | 7 | 7 | 7 | 1 |
| 25 | TGME49_233770.P1515 | calcium-translocating P-type ATPase, PMCA-type protein | 10 | 10 | 10 | 9 | 10 | 7 | 9 | 1 |
| 28 | TGME49_233820.P1518 | DNA polymerase delta small subunit, putative | 10 | 9 | 9 | 8 | 7 | 6 | 7 | 1 |
| 25 | TGME49_234190-t26_1-p1 | serine hydroxymethyltransferase 2, putative | 10 | 9 | 9 | 9 | 7 | 8 | 8 | 1 |
| 3 | TGME49_234280-t26_1-p1 | AMP deaminase | 10 | 10 | 10 | 9 | 10 | 9 | 8 | 1 |
| 3 | TGME49_234360-t26_1-p1 | DNA topoisomerase I, putative | 10 | 10 | 10 | 10 | 9 | 9 | 9 | 1 |
| 25 | TGME49_234420-t26_1-p1 | ATPase, AAA family protein | 10 | 9 | 9 | 9 | 9 | 9 | 8 | 1 |
| 25 | TGME49_234505-t26_1-p1 | phenylalanyl-tRNA synthetase alpha chain A, putative | 9 | 9 | 9 | 7 | 8 | 8 | 8 | 1 |
| 23 | TGME49_234520-t26_1-p1 | U2 snRNP auxilliary factor, large subunit, splicing factor subfamily protein | 10 | 8 | 9 | 9 | 6 | 8 | 3 | 1 |
| 25 | TGME49_234570.P1541 | sterol carrier protein-2 HAD-2SCP-2 | 10 | 9 | 9 | 9 | 8 | 8 | 8 | 1 |
| 28 | TGME49_234900-t26_1-p1 | PHD-finger domain-containing protein | 10 | 10 | 8 | 8 | 7 | 7 | 6 | 1 |
| 28 | TGME49_234970.P1546 | tyrosine kinase-like protein TKL2 | 10 | 9 | 9 | 8 | 6 | 8 | 6 | 1 |
| 3 | TGME49_235020.P1548 | COPI protein, putative | 10 | 10 | 10 | 9 | 10 | 9 | 9 | 1 |
| 25 | TGME49_235130-t26_1-p1 | rhoptry apical surface protein RASP1 | 9 | 9 | 9 | 8 | 8 | 8 | 7 | 1 |
| 6 | TGME49_235160-t26_1-p1 | hypothetical protein | 10 | 10 | 10 | 5 | 10 | 7 | 5 | 1 |
| 28 | TGME49_235170-t26_1-p1 | ATPase, AAA family protein | 10 | 10 | 9 | 9 | 8 | 7 | 6 | 1 |
| 25 | TGME49_235370.P1555 | protein kinase, putative | 10 | 10 | 9 | 9 | 9 | 9 | 8 | 1 |
| 28 | TGME49_235398-t26_1-p1 | hypothetical protein | 9 | 8 | 9 | 8 | 7 | 6 | 6 | 1 |
| 3 | TGME49_235470.P1562 | myosin A | 10 | 10 | 10 | 10 | 10 | 10 | 10 | 1 |
| 23 | TGME49_235610-t26_1-p1 | ATPase, AAA family protein | 10 | 10 | 9 | 8 | 8 | 8 | 3 | 1 |
| 28 | TGME49_235620.P1566 | deoxyhypusine synthase | 9 | 9 | 9 | 8 | 6 | 7 | 7 | 1 |
| 6 | TGME49_235750.P1575 | ULK kinase | 9 | 9 | 8 | 6 | 8 | 6 | 8 | 1 |
| 25 | TGME49_235860.P1576 | subtilisin SUB11 | 10 | 10 | 10 | 10 | 8 | 8 | 9 | 1 |
| 3 | TGME49_235920.P1578 | dynein, axonemal, heavy chain 2 family protein | 10 | 10 | 10 | 10 | 10 | 10 | 10 | 1 |
| 25 | TGME49_235930.P1579 | domain K- type RNA binding proteins family protein | 10 | 9 | 9 | 9 | 8 | 8 | 8 | 1 |
| 25 | TGME49_235970-t26_1-p1 | eukaryotic initiation factor-2 gamma, putative | 9 | 9 | 9 | 9 | 9 | 8 | 9 | 1 |
| 6 | TGME49_236080-t26_1-p1 | replication factor-a protein 1 (rpa1) subfamily protein | 9 | 9 | 9 | 6 | 8 | 8 | 6 | 1 |
| 25 | TGME49_236210-t26_1-p1 | cytochrome b-c1 complex subunit 1 | 9 | 9 | 9 | 9 | 9 | 9 | 8 | 1 |
| 6 | TGME49_236620-t26_1-p1 | protein kinase (incomplete catalytic triad) | 10 | 9 | 9 | 6 | 8 | 8 | 6 | 1 |
| 3 | TGME49_236650.P1601 | DEAD (Asp-Glu-Ala-Asp) box polypeptide 17 | 10 | 9 | 9 | 9 | 8 | 9 | 9 | 1 |
| 25 | TGME49_236960.P1610 | transporter facilitator protein TFP2 | 9 | 7 | 9 | 9 | 8 | 8 | 8 | 1 |
| 3 | TGME49_236970.P1611 | SWI2/SNF2-containing PHD finger protein | 10 | 10 | 10 | 10 | 10 | 10 | 9 | 1 |
| 28 | TGME49_237110-t26_1-p1 | replication factor C subunit 2, putative | 9 | 8 | 8 | 8 | 7 | 7 | 7 | 1 |
| 28 | TGME49_237170.P1618 | hypothetical protein | 9 | 9 | 9 | 9 | 6 | 7 | 6 | 1 |

| | | | | | | | | | | |
|---|---|---|---|---|---|---|---|---|---|---|
| 3 | TGME49_237220-t26_1-p1 | DNA replication licensing factor Mcm7, putative | 10 | 10 | 10 | 10 | 10 | 9 | 9 | 1 |
| 25 | TGME49_237510-t26_1-p1 | hypothetical protein | 10 | 10 | 10 | 8 | 8 | 8 | 7 | 1 |
| 3 | TGME49_238050.P1641 | nuclease and tudor domain-containing protein | 10 | 10 | 10 | 10 | 9 | 9 | 9 | 1 |
| 28 | TGME49_238190-t26_1-p1 | DNA-directed RNA polymerase II RPB3 | 9 | 9 | 8 | 8 | 8 | 7 | 6 | 1 |
| 6 | TGME49_238230-t26_1-p1 | Ser/Thr phosphatase family protein | 9 | 8 | 8 | 4 | 8 | 7 | 7 | 1 |
| 28 | TGME49_238240.P1645 | bystin protein | 9 | 8 | 8 | 8 | 7 | 7 | 7 | 1 |
| 28 | TGME49_238390.P1649 | guanylate cyclase organizer UGO | 10 | 10 | 9 | 8 | 8 | 6 | 6 | 1 |
| 28 | TGME49_238400-t26_1-p1 | endonuclease/exonuclease/phosphatase domain-containing protein | 10 | 10 | 10 | 7 | 6 | 7 | 7 | 1 |
| 28 | TGME49_238940.P1657 | GDP mannose 4,6-dehydratase, putative | 9 | 8 | 8 | 8 | 8 | 7 | 7 | 1 |
| 23 | TGME49_238950.P1658 | stearoyl-CoA desaturase | 10 | 10 | 10 | 9 | 8 | 8 | 3 | 1 |
| 25 | TGME49_238995.P1661 | ion channel protein | 10 | 10 | 10 | 9 | 9 | 8 | 7 | 1 |
| 3 | TGME49_239020-t26_1-p1 | ABC transporter transmembrane region domain-containing protein | 10 | 10 | 10 | 8 | 10 | 9 | 9 | 1 |
| 6 | TGME49_239430.P1675 | EF hand domain-containing protein | 9 | 9 | 9 | 5 | 6 | 7 | 7 | 1 |
| 28 | TGME49_239440-t26_1-p1 | protein kinase (incomplete catalytic triad) | 10 | 10 | 8 | 7 | 7 | 6 | 6 | 1 |
| 25 | TGME49_239490.P1676 | 2-oxoisovalerate dehydrogenase subunit alpha | 9 | 9 | 9 | 9 | 9 | 8 | 8 | 1 |
| 28 | TGME49_239500-t26_1-p1 | proteasome subunit alpha type-4, putative | 8 | 8 | 8 | 8 | 8 | 7 | 7 | 1 |
| 23 | TGME49_239560-t26_1-p1 | myosin E | 10 | 10 | 10 | 7 | 9 | 8 | 3 | 1 |
| 25 | TGME49_239620-t26_1-p1 | 5'-nucleotidase, C-terminal domain-containing protein | 10 | 10 | 10 | 10 | 6 | 8 | 8 | 1 |
| 28 | TGME49_239680-t26_1-p1 | hypothetical protein | 9 | 9 | 9 | 8 | 7 | 6 | 6 | 1 |
| 28 | TGME49_239752-t26_1-p1 | hypothetical protein | 10 | 10 | 10 | 9 | 7 | 8 | 6 | 1 |
| 6 | TGME49_239820.P1689 | D-3-phosphoglycerate dehydrogenase | 10 | 9 | 9 | 6 | 8 | 8 | 8 | 1 |
| 28 | TGME49_239930-t26_1-p1 | beta-catenin family protein 1, putative | 9 | 9 | 9 | 8 | 9 | 8 | 6 | 1 |
| 28 | TGME49_240210.P1697 | phosphate transporter family protein, putative | 10 | 9 | 7 | 7 | 7 | 7 | 7 | 1 |
| 23 | TGME49_240240-t26_1-p1 | subtilisin SUB5 | 10 | 9 | 9 | 8 | 7 | 7 | 5 | 1 |
| 23 | TGME49_240430-t26_1-p1 | glyoxalase family protein | 10 | 10 | 9 | 9 | 7 | 6 | 3 | 1 |
| 3 | TGME49_240600-t26_1-p1 | chaperonin cpn60, putative | 10 | 10 | 10 | 10 | 9 | 9 | 9 | 1 |
| 25 | TGME49_240640-t26_1-p1 | casein kinase I | 8 | 8 | 8 | 8 | 8 | 8 | 8 | 1 |
| 3 | TGME49_240650-t26_1-p1 | coatomer protein complex, subunit alpha, putative | 10 | 10 | 10 | 10 | 10 | 10 | 9 | 1 |
| 25 | TGME49_240690.P1712 | S-adenosyl methionine synthetase | 9 | 9 | 9 | 9 | 8 | 7 | 8 | 1 |
| 25 | TGME49_240710.P1713 | RNA recognition motif-containing protein | 10 | 10 | 10 | 7 | 9 | 8 | 8 | 1 |
| 28 | TGME49_240740.P1714 | Sec1 family protein | 10 | 9 | 9 | 7 | 8 | 8 | 7 | 1 |
| 23 | TGME49_240770-t26_1-p1 | cytochrome b5 family heme/steroid binding domain-containing protein | 9 | 8 | 8 | 7 | 7 | 7 | 3 | 1 |
| 28 | TGME49_240810.P1716 | apicomplexan amino acid transporter ApiAT6-1 | 9 | 9 | 9 | 8 | 7 | 7 | 7 | 1 |
| 25 | TGME49_240870-t26_1-p1 | beta adaptin protein, putative | 10 | 10 | 10 | 7 | 9 | 9 | 8 | 1 |
| 3 | TGME49_240890-t26_1-p1 | 6-phosphofructokinase | 10 | 10 | 10 | 10 | 10 | 10 | 10 | 1 |
| 3 | TGME49_240910.P1720 | double C2-like domain-containing protein DOC2.1 | 10 | 10 | 10 | 7 | 7 | 10 | 10 | 1 |
| 6 | TGME49_240970-t26_1-p1 | hypothetical protein | 10 | 9 | 9 | 6 | 8 | 7 | 7 | 1 |
| 6 | TGME49_241150-t26_1-p1 | N2227 domain-containing protein | 9 | 9 | 9 | 5 | 8 | 7 | 6 | 1 |
| 28 | TGME49_241830-t26_1-p1 | peptidyl-prolyl cis-trans isomerase | 9 | 9 | 8 | 8 | 7 | 6 | 6 | 1 |
| 23 | TGME49_241880-t26_1-p1 | 3'5'-cyclic nucleotide phosphodiesterase PDE9 | 10 | 10 | 10 | 9 | 5 | 8 | 3 | 1 |
| 28 | TGME49_242070.P1730 | cAMP-dependent protein kinase regulatory subunit 1 | 9 | 9 | 9 | 8 | 8 | 7 | 7 | 1 |
| 28 | TGME49_242290-t26_1-p1 | proteasome subunit alpha type-6, putative | 8 | 8 | 8 | 8 | 8 | 7 | 7 | 1 |
| 25 | TGME49_242320.P1734 | preconoidal ring protein PCR5 | 10 | 10 | 10 | 8 | 7 | 9 | 7 | 1 |
| 6 | TGME49_242330.P1735 | ribosomal protein RPS5 | 7 | 7 | 7 | 3 | 7 | 7 | 7 | 1 |

| | | | | | | | | | | |
|---|---|---|---|---|---|---|---|---|---|---|
| 25 | TGME49_242400.P1738 | calcium-dependent protein kinase CDPK2B | 10 | 9 | 9 | 9 | 9 | 9 | 6 | 1 |
| 25 | TGME49_242625.P1739 | ATPase family associated with various cellular activities (AAA) subfamily protein | 10 | 10 | 10 | 10 | 9 | 9 | 6 | 1 |
| 3 | TGME49_242660-t26_1-p1 | vacuolar sorting protein 35, putative | 10 | 10 | 10 | 9 | 10 | 9 | 8 | 1 |
| 25 | TGME49_242720.P1743 | aspartyl protease ASP5 | 10 | 9 | 9 | 9 | 8 | 8 | 8 | 1 |
| 25 | TGME49_242830-t26_1-p1 | XRN 5'-3' exonuclease N-terminus protein | 10 | 10 | 10 | 8 | 9 | 6 | 9 | 1 |
| 23 | TGME49_242890-t26_1-p1 | hypothetical protein | 10 | 10 | 10 | 9 | 6 | 8 | 5 | 1 |
| 3 | TGME49_243250-t26_1-p1 | myosin H | 10 | 10 | 10 | 10 | 10 | 9 | 10 | 1 |
| 25 | TGME49_243440-t26_1-p1 | histone lysine acetyltransferase GCN5-B | 10 | 9 | 10 | 8 | 9 | 8 | 8 | 1 |
| 28 | TGME49_243470-t26_1-p1 | hypothetical protein | 10 | 10 | 10 | 8 | 7 | 6 | 6 | 1 |
| 28 | TGME49_243490-t26_1-p1 | mitochondrial chaperone BCS1, putative | 9 | 9 | 9 | 7 | 9 | 8 | 6 | 1 |
| 28 | TGME49_243530-t26_1-p1 | pentatricopeptide repeat domain-containing protein | 10 | 10 | 10 | 9 | 6 | 7 | 8 | 1 |
| 28 | TGME49_243540-t26_1-p1 | U4/U6 small nuclear ribonucleoprotein PRP4, putative | 10 | 9 | 9 | 9 | 7 | 7 | 7 | 1 |
| 28 | TGME49_243710-t26_1-p1 | t-complex protein beta subunit | 10 | 9 | 9 | 9 | 6 | 9 | 7 | 1 |
| 25 | TGME49_243800-t26_1-p1 | acyl-CoA synthetase ACS4, putative | 10 | 10 | 10 | 9 | 9 | 8 | 8 | 1 |
| 3 | TGME49_243920-t26_1-p1 | DNA replication licensing factor MCM5, putative | 10 | 10 | 10 | 9 | 10 | 9 | 8 | 1 |
| 23 | TGME49_244000.P1777 | DEAD/DEAH box helicase domain-containing protein | 10 | 10 | 9 | 9 | 9 | 9 | 5 | 1 |
| 25 | TGME49_244100.P1781 | U4/U6 small nuclear ribonucleoprotein PRP31, putative | 10 | 9 | 9 | 8 | 8 | 8 | 7 | 1 |
| 3 | TGME49_244200-t26_1-p1 | 2-oxoglutarate dehydrogenase e1 component, mitochondrial precursor, putative | 10 | 10 | 10 | 10 | 10 | 10 | 9 | 1 |
| 28 | TGME49_244210-t26_1-p1 | RNA polymerase-associated protein RTF1 | 10 | 9 | 9 | 7 | 7 | 6 | 6 | 1 |
| 6 | TGME49_244290-t26_1-p1 | AP-4 complex subunit epsilon, putative | 10 | 10 | 10 | 5 | 8 | 8 | 7 | 1 |
| 25 | TGME49_244310-t26_1-p1 | GTP-binding protein | 9 | 9 | 9 | 7 | 9 | 7 | 8 | 1 |
| 3 | TGME49_244380-t26_1-p1 | cactin | 10 | 10 | 9 | 9 | 9 | 9 | 9 | 1 |
| 23 | TGME49_244500-t26_1-p1 | Tubulin-tyrosine ligase family protein | 10 | 10 | 10 | 9 | 8 | 8 | 3 | 1 |
| 3 | TGME49_244560.P1794 | heat shock protein 90, putative | 10 | 10 | 10 | 10 | 9 | 9 | 9 | 1 |
| 6 | TGME49_244650.P1800 | eukaryotic initiation factor-5, putative | 9 | 9 | 9 | 3 | 8 | 6 | 8 | 1 |
| 3 | TGME49_244880-t26_1-p1 | DNA-directed RNA polymerase I RPA1 | 10 | 10 | 10 | 10 | 10 | 10 | 9 | 1 |
| 28 | TGME49_245470-t26_1-p1 | mitotic checkpoint protein, BUB3 family protein | 9 | 8 | 8 | 8 | 8 | 6 | 7 | 1 |
| 28 | TGME49_245500-t26_1-p1 | dipeptidyl peptidase iv (dpp iv) n-terminal region domain-containing protein | 10 | 10 | 10 | 10 | 7 | 6 | 6 | 1 |
| 25 | TGME49_245510-t26_1-p1 | phospholipid-translocating P-type ATPase, flippase subfamily protein | 10 | 10 | 10 | 9 | 7 | 8 | 8 | 1 |
| 25 | TGME49_245670-t26_1-p1 | pyruvate dehydrogenase complex subunit PDH-E1Alpha | 10 | 9 | 9 | 9 | 8 | 8 | 8 | 1 |
| 28 | TGME49_245710.P1816 | membrane occupation and recognition nexus protein MORN3 | 8 | 8 | 8 | 8 | 8 | 7 | 7 | 1 |
| 25 | TGME49_246060.P1821 | DNA-dependent RNA polymerase | 10 | 10 | 10 | 10 | 6 | 8 | 8 | 1 |
| 28 | TGME49_246220.P1830 | LITAF-like zinc finger protein, putative | 10 | 9 | 8 | 8 | 7 | 7 | 6 | 1 |
| 28 | TGME49_246330-t26_1-p1 | CRAL/TRIO domain-containing protein | 9 | 9 | 8 | 7 | 7 | 7 | 7 | 1 |
| 25 | TGME49_246340-t26_1-p1 | DnaJ domain-containing protein | 10 | 9 | 9 | 8 | 9 | 9 | 8 | 1 |
| 23 | TGME49_246500.P1832 | splicing factor 3A subunit 1, putative | 10 | 9 | 9 | 8 | 7 | 7 | 5 | 1 |
| 25 | TGME49_246510-t26_1-p1 | protein phosphatase 2a regulatory b subunit (b56 family) protein | 10 | 9 | 9 | 9 | 9 | 8 | 8 | 1 |
| 6 | TGME49_246540.P1834 | cytochrome c1, heme protein | 9 | 8 | 9 | 3 | 9 | 8 | 8 | 1 |
| 25 | TGME49_246550-t26_1-p1 | aspartyl protease ASP3 | 10 | 9 | 9 | 9 | 9 | 8 | 8 | 1 |

| | | | | | | | | | | |
|---|---|---|---|---|---|---|---|---|---|---|
| 3 | TGME49_246600-t26_1-p1 | ABC1 family protein | 10 | 10 | 10 | 8 | 9 | 9 | 9 | 1 |
| 25 | TGME49_246690-t26_1-p1 | alpha amylase, catalytic domain-containing protein | 10 | 10 | 9 | 7 | 8 | 9 | 8 | 1 |
| 6 | TGME49_246800-t26_1-p1 | acylaminoacyl-peptidase, putative | 10 | 9 | 9 | 6 | 8 | 7 | 7 | 1 |
| 28 | TGME49_247250-t26_1-p1 | RbAp46 | 10 | 8 | 8 | 8 | 7 | 7 | 6 | 1 |
| 28 | TGME49_247370.P1851 | polycystin cation channel protein | 10 | 10 | 10 | 7 | 8 | 6 | 7 | 1 |
| 25 | TGME49_247470-t26_1-p1 | nucleolar protein, putative | 9 | 9 | 9 | 8 | 9 | 8 | 8 | 1 |
| 25 | TGME49_247500-t26_1-p1 | acyl-CoA dehydrogenase, middle domain-containing protein | 10 | 9 | 10 | 10 | 8 | 9 | 8 | 1 |
| 28 | TGME49_247510.P1856 | fructose-1,6-bisphosphatase 2 | 9 | 9 | 8 | 8 | 8 | 6 | 8 | 1 |
| 28 | TGME49_247540.P1858 | ATP-binding cassette G family transporter ABCG107 | 10 | 10 | 9 | 8 | 6 | 7 | 7 | 1 |
| 3 | TGME49_247550.P1859 | heat shock protein HSP60 | 10 | 10 | 9 | 9 | 9 | 9 | 9 | 1 |
| 25 | TGME49_247690-t26_1-p1 | phospholipid-translocating P-type ATPase, flippase subfamily protein | 10 | 10 | 10 | 8 | 8 | 9 | 8 | 1 |
| 28 | TGME49_247710-t26_1-p1 | casein kinase I | 10 | 9 | 9 | 8 | 8 | 7 | 7 | 1 |
| 25 | TGME49_247760.P1865 | acyl-CoA synthetase ACS5, putative | 10 | 10 | 10 | 9 | 9 | 8 | 8 | 1 |
| 28 | TGME49_247850.P1868 | 28 kDa A-kinase anchor, putative | 10 | 10 | 9 | 9 | 7 | 7 | 6 | 1 |
| 23 | TGME49_248120.P1874 | hypothetical protein | 10 | 9 | 9 | 9 | 8 | 7 | 4 | 1 |
| 28 | TGME49_248400.P1887 | glyoxalase I, putative | 9 | 8 | 8 | 8 | 7 | 7 | 7 | 1 |
| 25 | TGME49_248470.P1891 | subtilisin SUB6 | 10 | 8 | 10 | 9 | 9 | 8 | 8 | 1 |
| 28 | TGME49_248630.P1898 | actin-related protein ARP1 | 9 | 9 | 9 | 7 | 8 | 8 | 6 | 1 |
| 3 | TGME49_248640.P1899 | ND6 protein | 10 | 10 | 10 | 10 | 10 | 10 | 8 | 1 |
| 3 | TGME49_248670-t26_1-p1 | V-type H(+)-translocating pyrophosphatase VP1 | 10 | 10 | 10 | 10 | 10 | 10 | 9 | 1 |
| 28 | TGME49_248830.P1907 | phosphoinositide phospholipase PIPLC | 10 | 10 | 10 | 8 | 7 | 8 | 6 | 1 |
| 23 | TGME49_249020.P1916 | kinesin motor domain-containing protein | 10 | 10 | 9 | 8 | 6 | 7 | 3 | 1 |
| 6 | TGME49_249170-t26_1-p1 | Ras family protein | 10 | 10 | 9 | 6 | 7 | 8 | 6 | 1 |
| 23 | TGME49_249180.P1919 | bifunctional dihydrofolate reductase-thymidylate synthase | 10 | 9 | 9 | 7 | 8 | 7 | 5 | 1 |
| 6 | TGME49_249260.P1923 | CDK-related protein kinase CRK3 | 10 | 10 | 7 | 5 | 6 | 6 | 7 | 1 |
| 25 | TGME49_249270.P1924 | protein disulfide isomerase-related protein (provisional), putative | 9 | 9 | 9 | 7 | 8 | 8 | 8 | 1 |
| 6 | TGME49_249380.P1930 | palmitoyltransferase DHHC13 | 9 | 8 | 8 | 5 | 7 | 7 | 6 | 1 |
| 3 | TGME49_249390.P1931 | glutamate/leucine/phenylalanine/valine dehydrogenase family protein | 10 | 10 | 10 | 7 | 10 | 8 | 10 | 1 |
| 28 | TGME49_249480.P1933 | co-chaperone UNC | 10 | 10 | 10 | 10 | 6 | 8 | 7 | 1 |
| 3 | TGME49_249530.P1935 | exportin 1, putative | 10 | 10 | 10 | 10 | 10 | 10 | 10 | 1 |
| 28 | TGME49_249590-t26_1-p1 | proteasome subunit alpha type-5, putative | 8 | 8 | 8 | 8 | 8 | 7 | 7 | 1 |
| 6 | TGME49_249620-t26_1-p1 | histone deacetylase HDAC2 | 10 | 9 | 9 | 4 | 7 | 8 | 8 | 1 |
| 25 | TGME49_249670-t26_1-p1 | cathepsin B | 9 | 9 | 9 | 8 | 7 | 8 | 8 | 1 |
| 25 | TGME49_249730.P1941 | ND9 protein | 10 | 10 | 10 | 9 | 8 | 9 | 7 | 1 |
| 3 | TGME49_249810.P1947 | activating signal cointegrator 1 complex subunit 3, putative | 10 | 10 | 10 | 8 | 8 | 10 | 10 | 1 |
| 6 | TGME49_249820-t26_1-p1 | ATP-binding cassette sub-family B member 5 | 10 | 9 | 9 | 3 | 7 | 7 | 7 | 1 |
| 3 | TGME49_249840.P1948 | dynein heavy chain 2, putative | 10 | 10 | 10 | 10 | 10 | 10 | 10 | 1 |
| 25 | TGME49_249850-t26_1-p1 | GAP40 protein | 9 | 9 | 9 | 8 | 8 | 8 | 8 | 1 |
| 28 | TGME49_250690-t26_1-p1 | zinc finger (CCCH type) motif-containing protein | 10 | 10 | 7 | 8 | 8 | 8 | 7 | 1 |
| 6 | TGME49_250760-t26_1-p1 | pentatricopeptide repeat domain-containing protein | 10 | 10 | 9 | 3 | 8 | 8 | 7 | 1 |
| 25 | TGME49_250770.P1963 | eukaryotic initiation factor-4A, putative | 9 | 9 | 9 | 9 | 9 | 8 | 8 | 1 |

| | | | | | | | | | | |
|---|---|---|---|---|---|---|---|---|---|---|
| 23 | TGME49_250790-t26_1-p1 | hypothetical protein | 10 | 9 | 9 | 8 | 7 | 7 | 4 | 1 |
| 28 | TGME49_251500.P1976 | eukaryotic initiation factor-3, subunit 3, putative | 9 | 9 | 9 | 9 | 8 | 7 | 7 | 1 |
| 25 | TGME49_251620-t26_1-p1 | flap structure-specific endonuclease 1, putative | 9 | 9 | 9 | 8 | 8 | 8 | 7 | 1 |
| 28 | TGME49_251690-t26_1-p1 | seryl-tRNA synthetase, cytoplasmic, putative | 9 | 9 | 9 | 9 | 8 | 8 | 6 | 1 |
| 3 | TGME49_251780-t26_1-p1 | heat shock protein | 10 | 10 | 10 | 10 | 10 | 9 | 9 | 1 |
| 3 | TGME49_251850.P1983 | serine/threonine protein phosphatase PP7 | 10 | 10 | 10 | 10 | 10 | 10 | 10 | 1 |
| 25 | TGME49_251880-t26_1-p1 | tRNA synthetases class I family protein | 9 | 9 | 9 | 9 | 8 | 8 | 8 | 1 |
| 28 | TGME49_251930-t26_1-p1 | enoyl-acyl carrier reductase ENR | 9 | 8 | 8 | 8 | 8 | 7 | 7 | 1 |
| 25 | TGME49_252220-t26_1-p1 | tetratricopeptide repeat domain containing protein | 10 | 9 | 9 | 9 | 9 | 9 | 8 | 1 |
| 25 | TGME49_252290.P1990 | importin alpha, putative | 10 | 9 | 9 | 9 | 9 | 9 | 8 | 1 |
| 28 | TGME49_252420.P1997 | histone arginine methyltransferase PRMT3 | 10 | 9 | 8 | 8 | 6 | 7 | 6 | 1 |
| 28 | TGME49_252430.P1998 | START domain-containing protein | 10 | 9 | 9 | 7 | 7 | 8 | 7 | 1 |
| 3 | TGME49_252640-t26_1-p1 | P-type ATPase PMA1 | 10 | 10 | 10 | 10 | 10 | 9 | 9 | 1 |
| 28 | TGME49_252850.P2003 | initiation factor, subunit 2 family protein | 9 | 9 | 9 | 8 | 7 | 7 | 6 | 1 |
| 28 | TGME49_253030-t26_1-p1 | glycosyl hydrolase, family 31 protein | 10 | 10 | 10 | 10 | 6 | 8 | 6 | 1 |
| 3 | TGME49_253290.P2015 | valyl-tRNA synthetase | 10 | 10 | 10 | 9 | 10 | 10 | 10 | 1 |
| 25 | TGME49_253430.P2022 | asparagine synthetase, putative | 10 | 9 | 9 | 9 | 9 | 8 | 8 | 1 |
| 25 | TGME49_253440.P2023 | cell-cycle-associated protein kinase SRPK, putative | 10 | 10 | 10 | 10 | 8 | 9 | 7 | 1 |
| 28 | TGME49_253550.P2027 | WD repeat-containing protein 92, putative | 9 | 8 | 8 | 8 | 8 | 7 | 6 | 1 |
| 28 | TGME49_253700-t26_1-p1 | transporter, major facilitator family protein | 10 | 10 | 10 | 7 | 7 | 6 | 6 | 1 |
| 23 | TGME49_253730-t26_1-p1 | importin-beta N-terminal domain-containing protein | 10 | 10 | 10 | 10 | 10 | 8 | 5 | 1 |
| 25 | TGME49_254020.P2043 | dynein intermediate chain, putative | 10 | 10 | 9 | 9 | 8 | 8 | 8 | 1 |
| 23 | TGME49_254300-t26_1-p1 | hypothetical protein | 10 | 10 | 10 | 10 | 8 | 7 | 4 | 1 |
| 3 | TGME49_254370-t26_1-p1 | guanylyl cyclase | 10 | 10 | 10 | 10 | 6 | 10 | 10 | 1 |
| 28 | TGME49_254390-t26_1-p1 | CRAL/TRIO domain-containing protein | 10 | 9 | 9 | 9 | 8 | 7 | 7 | 1 |
| 6 | TGME49_254490-t26_1-p1 | Sel1 repeat-containing protein | 10 | 10 | 9 | 5 | 8 | 8 | 7 | 1 |
| 23 | TGME49_254570-t26_1-p1 | hypothetical protein | 10 | 10 | 10 | 8 | 9 | 7 | 4 | 1 |
| 25 | TGME49_254630.P2064 | CDK-related protein kinase CRK6 | 10 | 10 | 10 | 8 | 8 | 8 | 7 | 1 |
| 28 | TGME49_254650.P2066 | pre-mRNA-splicing factor Cwc2, putative | 9 | 9 | 9 | 8 | 8 | 7 | 7 | 1 |
| 28 | TGME49_254690.P2068 | serine hydrolase ASH2 | 9 | 9 | 8 | 8 | 8 | 7 | 7 | 1 |
| 28 | TGME49_254710-t26_1-p1 | serine esterase (DUF676) protein | 10 | 10 | 10 | 9 | 6 | 7 | 6 | 1 |
| 23 | TGME49_254750.P2071 | c3orf15 protein | 9 | 8 | 9 | 9 | 8 | 7 | 5 | 1 |
| 28 | TGME49_254910.P2080 | hypothetical protein | 10 | 10 | 9 | 8 | 7 | 7 | 6 | 1 |
| 6 | TGME49_254940-t26_1-p1 | MIF4G domain-containing protein | 10 | 10 | 10 | 5 | 7 | 7 | 7 | 1 |
| 23 | TGME49_255190.P2086 | myosin C, myosin B | 10 | 10 | 10 | 10 | 10 | 10 | 3 | 1 |
| 28 | TGME49_255250.P2093 | tRNA (cytosine(34)-C(5))-methyltransferase, putative | 10 | 10 | 10 | 9 | 6 | 6 | 8 | 1 |
| 6 | TGME49_255260-t26_1-p1 | apical membrane antigen AMA1 | 9 | 9 | 9 | 6 | 7 | 6 | 7 | 1 |
| 25 | TGME49_255350.P2096 | ATPase, putative | 9 | 8 | 9 | 9 | 8 | 9 | 8 | 1 |
| 28 | TGME49_255510-t26_1-p1 | ankyrin repeat-containing protein | 10 | 10 | 9 | 9 | 7 | 8 | 6 | 1 |
| 25 | TGME49_255880.P2119 | hypothetical protein | 10 | 10 | 10 | 7 | 8 | 8 | 7 | 1 |
| 28 | TGME49_255970.P2125 | histone lysine methyltransferase SET/SUV39 | 10 | 9 | 9 | 8 | 8 | 7 | 7 | 1 |
| 23 | TGME49_256020.P2128 | EFHC1, putative | 10 | 9 | 9 | 7 | 8 | 7 | 3 | 1 |
| 3 | TGME49_256760-t26_1-p1 | pyruvate kinase PyK1 | 9 | 9 | 9 | 9 | 9 | 9 | 9 | 1 |
| 25 | TGME49_256770-t26_1-p1 | eukaryotic translation initiation factor 4A, isoform 3, putative | 9 | 9 | 9 | 9 | 8 | 9 | 9 | 1 |
| 28 | TGME49_256800.P2138 | hypothetical protein | 10 | 10 | 9 | 7 | 7 | 6 | 6 | 1 |
| 3 | TGME49_256970-t26_1-p1 | vacuolar ATP synthase subunit A, putative | 10 | 10 | 10 | 10 | 10 | 9 | 9 | 1 |

| | | | | | | | | | | |
|---|---|---|---|---|---|---|---|---|---|---|
| 6 | TGME49_256980.P2148 | glycerol-3-phosphate-acyltransferase | 10 | 10 | 9 | 6 | 8 | 8 | 7 | 1 |
| 3 | TGME49_256990-t26_1-p1 | glycyl-tRNA synthetase | 10 | 10 | 10 | 10 | 10 | 9 | 9 | 1 |
| 28 | TGME49_257050.P2151 | ketopantoate hydroxymethyltransferase - ketopantoate reductase | 10 | 9 | 9 | 9 | 7 | 6 | 6 | 1 |
| 23 | TGME49_257080.P2154 | 3'5'-cyclic nucleotide phosphodiesterase PDE13 | 10 | 10 | 10 | 9 | 8 | 8 | 3 | 1 |
| 28 | TGME49_257390.P2171 | ribosome biogenesis GTP-binding protein YsxC protein | 10 | 10 | 8 | 8 | 7 | 6 | 6 | 1 |
| 25 | TGME49_257470.P2175 | myosin J | 10 | 10 | 10 | 10 | 7 | 7 | 8 | 1 |
| 28 | TGME49_257480-t26_1-p1 | NADP-dependent succinate-semialdehyde dehydrogenase | 10 | 9 | 9 | 9 | 8 | 7 | 7 | 1 |
| 28 | TGME49_257560.P2179 | WD domain, G-beta repeat-containing protein | 10 | 10 | 9 | 9 | 7 | 7 | 7 | 1 |
| 28 | TGME49_258010.P2198 | calcium signaling protein kinase RAD53, putative | 10 | 9 | 8 | 8 | 7 | 7 | 7 | 1 |
| 3 | TGME49_258030.P2200 | DNA polymerase | 10 | 10 | 10 | 9 | 8 | 10 | 10 | 1 |
| 3 | TGME49_258210.P2213 | DNA-directed RNA polymerase II RPB2 | 10 | 10 | 10 | 10 | 10 | 9 | 10 | 1 |
| 28 | TGME49_258390.P2219 | DnaJ protein, putative | 9 | 8 | 8 | 8 | 8 | 7 | 7 | 1 |
| 6 | TGME49_258400-t26_1-p1 | LCCL-related protein CCp5A | 10 | 10 | 10 | 3 | 6 | 7 | 7 | 1 |
| 25 | TGME49_258690.P2235 | 5-aminolevulinic acid synthase domain-containing protein | 10 | 9 | 9 | 8 | 8 | 8 | 8 | 1 |
| 28 | TGME49_258700.P2236 | transporter, major facilitator family protein | 10 | 9 | 9 | 8 | 6 | 8 | 8 | 1 |
| 28 | TGME49_258740.P2238 | eukaryotic initiation factor-2A, putative | 10 | 9 | 9 | 8 | 7 | 7 | 7 | 1 |
| 6 | TGME49_258770.P2239 | polypeptide n-acetylgalactosaminyltransferase 12, putative | 10 | 10 | 9 | 5 | 8 | 8 | 8 | 1 |
| 6 | TGME49_258832.P2243 | Vps52 / Sac2 family protein | 9 | 10 | 8 | 6 | 7 | 7 | 8 | 1 |
| 23 | TGME49_258950.P2251 | lectin family protein | 10 | 9 | 9 | 9 | 8 | 6 | 4 | 1 |
| 28 | TGME49_258980.P2253 | CCR4-NOT transcription complex subunit 9, putative | 9 | 8 | 8 | 8 | 6 | 7 | 7 | 1 |
| 25 | TGME49_259010.P2256 | vacuolar ATP synthase subunit d, putative | 9 | 9 | 9 | 8 | 8 | 8 | 7 | 1 |
| 28 | TGME49_259180.P2268 | cystathionine beta-synthase | 9 | 9 | 8 | 9 | 8 | 7 | 7 | 1 |
| 25 | TGME49_259200.P2270 | Na+/H+ exchanger NHE1 | 10 | 10 | 10 | 7 | 9 | 9 | 8 | 1 |
| 25 | TGME49_259260-t26_1-p1 | membrane protein FtsH1 | 10 | 10 | 10 | 9 | 7 | 9 | 8 | 1 |
| 25 | TGME49_259530.P2277 | GalNac | 10 | 10 | 9 | 9 | 8 | 8 | 7 | 1 |
| 28 | TGME49_259600.P2280 | hypothetical protein | 10 | 9 | 9 | 8 | 7 | 7 | 6 | 1 |
| 28 | TGME49_259720.P2288 | membrane inositol phospholipid binding protein | 10 | 9 | 9 | 7 | 6 | 7 | 7 | 1 |
| 3 | TGME49_260170.P2308 | elongation factor EF-G, putative | 10 | 10 | 10 | 8 | 9 | 9 | 9 | 1 |
| 3 | TGME49_260200.P2309 | helicase associated domain (ha2) protein | 10 | 10 | 10 | 10 | 9 | 10 | 10 | 1 |
| 25 | TGME49_260210-t26_1-p1 | DnaJ domain-containing protein | 10 | 9 | 9 | 8 | 9 | 8 | 8 | 1 |
| 6 | TGME49_260220.P2310 | folate/biopterin transporter subfamily protein | 10 | 9 | 9 | 5 | 6 | 8 | 8 | 1 |
| 28 | TGME49_260240.P2312 | ccr4-associated factor family protein | 10 | 9 | 8 | 7 | 6 | 7 | 7 | 1 |
| 3 | TGME49_260470.P2324 | ferlin 2 | 10 | 10 | 10 | 7 | 9 | 10 | 10 | 1 |
| 28 | TGME49_260500-t26_1-p1 | COPI associated protein, putative | 8 | 8 | 8 | 8 | 8 | 7 | 7 | 1 |
| 3 | TGME49_260630.P2334 | DnaJ family Sec63 protein | 10 | 10 | 10 | 9 | 9 | 9 | 9 | 1 |
| 6 | TGME49_260640.P2335 | autophagy-related protein ATG9 | 10 | 10 | 10 | 5 | 7 | 8 | 5 | 1 |
| 6 | TGME49_260790.P2340 | RAP domain-containing protein | 9 | 9 | 9 | 4 | 8 | 8 | 5 | 1 |
| 25 | TGME49_261000.P2346 | DNA mismatch repair protein MSH2, putative | 10 | 10 | 10 | 8 | 8 | 9 | 8 | 1 |
| 6 | TGME49_261010.P2347 | tat-binding family protein, putative | 9 | 9 | 9 | 6 | 8 | 9 | 9 | 1 |
| 3 | TGME49_261050.P2350 | exosome complex exonuclease RRP44, putative | 10 | 9 | 10 | 9 | 9 | 9 | 9 | 1 |
| 28 | TGME49_261060.P2351 | diphthamide biosynthesis protein 2-related domain-containing protein | 9 | 9 | 9 | 9 | 7 | 8 | 6 | 1 |
| 3 | TGME49_261080.P2353 | cysteine repeat modular protein CRMPA | 10 | 10 | 10 | 10 | 10 | 10 | 10 | 1 |
| 28 | TGME49_261210.P2355 | 26s proteasome subunit p55, putative | 9 | 9 | 9 | 7 | 8 | 8 | 7 | 1 |

| | | | | | | | | | | |
|---|---|---|---|---|---|---|---|---|---|---|
| 25 | TGME49_261230.P2357 | ankyrin repeat-containing protein | 10 | 10 | 10 | 7 | 9 | 8 | 8 | 1 |
| 25 | TGME49_261460-t26_1-p1 | transcriptional elongation factor FACT80 | 10 | 9 | 9 | 9 | 9 | 8 | 8 | 1 |
| 6 | TGME49_261480.P2366 | phosphatidyl serine synthase | 9 | 9 | 9 | 6 | 7 | 8 | 6 | 1 |
| 23 | TGME49_261500.P2368 | hydrolase | 9 | 9 | 8 | 8 | 8 | 7 | 3 | 1 |
| 6 | TGME49_261520.P2370 | nucleolar GTP-binding protein 1, putative | 10 | 10 | 10 | 3 | 9 | 8 | 8 | 1 |
| 28 | TGME49_261530-t26_1-p1 | aspartyl protease ASP7 | 10 | 9 | 7 | 9 | 6 | 7 | 7 | 1 |
| 28 | TGME49_261570-t26_1-p1 | ribosomal protein RPL7A | 8 | 8 | 8 | 8 | 8 | 7 | 7 | 1 |
| 25 | TGME49_261600.P2372 | aminopeptidase P | 10 | 10 | 10 | 9 | 9 | 8 | 6 | 1 |
| 25 | TGME49_261850.P2384 | plastid replication-repair enzyme PREX | 10 | 10 | 10 | 9 | 8 | 8 | 8 | 1 |
| 3 | TGME49_261950-t26_1-p1 | ATP synthase beta subunit ATP-B | 10 | 9 | 9 | 9 | 9 | 9 | 9 | 1 |
| 28 | TGME49_262070-t26_1-p1 | ribosomal rna assembly protein mis3, putative | 9 | 9 | 8 | 8 | 6 | 8 | 7 | 1 |
| 25 | TGME49_262150.P2397 | kelch protein K13 | 10 | 10 | 9 | 7 | 9 | 9 | 8 | 1 |
| 28 | TGME49_262430.P2403 | 4-hydroxy-3-methylbut-2-en-1-yl diphosphate synthase | 10 | 10 | 9 | 8 | 7 | 7 | 7 | 1 |
| 25 | TGME49_262880.P2430 | hypothetical protein | 10 | 10 | 10 | 10 | 8 | 8 | 8 | 1 |
| 28 | TGME49_262920.P2432 | serine protease DegP | 10 | 9 | 8 | 7 | 7 | 7 | 7 | 1 |
| 28 | TGME49_263000.P2437 | BEACH domain-containing protein BDCP | 10 | 10 | 10 | 10 | 7 | 7 | 6 | 1 |
| 25 | TGME49_263060.P2441 | Proteasome/cyclosome repeat-containing protein | 10 | 10 | 10 | 7 | 9 | 8 | 7 | 1 |
| 25 | TGME49_263070-t26_1-p1 | CMGC kinase, CK2 family | 9 | 9 | 9 | 8 | 8 | 7 | 8 | 1 |
| 28 | TGME49_263090.P2444 | 14-3-3 protein | 8 | 8 | 8 | 8 | 8 | 7 | 7 | 1 |
| 25 | TGME49_263100.P2445 | hypothetical protein | 10 | 10 | 10 | 10 | 9 | 8 | 7 | 1 |
| 6 | TGME49_263130-t26_1-p1 | citrate synthase, putative | 10 | 8 | 8 | 6 | 7 | 7 | 7 | 1 |
| 28 | TGME49_263140.P2448 | SWI2/SNF2-containing protein RAD26 | 10 | 10 | 10 | 9 | 8 | 7 | 7 | 1 |
| 28 | TGME49_263190.P2452 | adenylosuccinate lyase, putative | 9 | 9 | 9 | 8 | 8 | 7 | 6 | 1 |
| 6 | TGME49_263410-t26_1-p1 | scavenger receptor cysteine-rich domain-containing protein | 10 | 10 | 10 | 3 | 7 | 8 | 7 | 1 |
| 23 | TGME49_263420.P2467 | ubiquitin-specific protease USP4 | 10 | 10 | 10 | 10 | 8 | 8 | 3 | 1 |
| 28 | TGME49_263480.P2471 | sodium/hydrogen exchanger 3 protein | 10 | 9 | 9 | 9 | 7 | 6 | 6 | 1 |
| 3 | TGME49_263650.P2484 | pre-mRNA-processing protein 8, putative | 10 | 10 | 10 | 10 | 9 | 10 | 10 | 1 |
| 28 | TGME49_263710-t26_1-p1 | acyl-CoA:cholesterol acyltransferase alpha ACAT1-alpha | 9 | 9 | 9 | 7 | 8 | 7 | 8 | 1 |
| 6 | TGME49_263730.P2489 | FAD-dependent glycerol-3-phosphate dehydrogenase | 10 | 9 | 9 | 6 | 8 | 7 | 7 | 1 |
| 6 | TGME49_263820.P2495 | DEAD/DEAH box helicase domain-containing protein | 10 | 10 | 9 | 5 | 7 | 7 | 6 | 1 |
| 3 | TGME49_263870.P2498 | glutamate-tRNA ligase | 10 | 10 | 10 | 10 | 9 | 9 | 9 | 1 |
| 28 | TGME49_264000.P2501 | aldehyde dehydrogenase | 10 | 9 | 9 | 9 | 8 | 8 | 6 | 1 |
| 25 | TGME49_264060-t26_1-p1 | PCI domain-containing protein | 9 | 9 | 9 | 8 | 9 | 8 | 8 | 1 |
| 23 | TGME49_264070-t26_1-p1 | F5/8 type C domain-containing protein | 10 | 9 | 10 | 8 | 9 | 8 | 3 | 1 |
| 25 | TGME49_264178.P2510 | flagellar associated protein | 10 | 10 | 10 | 10 | 9 | 9 | 7 | 1 |
| 25 | TGME49_264450-t26_1-p1 | DNA topoisomerase III beta-1, putative | 10 | 10 | 10 | 10 | 9 | 9 | 6 | 1 |
| 6 | TGME49_264690.P2520 | cyclin 4, putative | 9 | 8 | 8 | 6 | 8 | 7 | 7 | 1 |
| 6 | TGME49_264760-t26_1-p1 | Oxysterol-binding protein | 10 | 10 | 10 | 6 | 8 | 8 | 8 | 1 |
| 28 | TGME49_264820-t26_1-p1 | RbAp48 | 10 | 10 | 9 | 8 | 8 | 7 | 7 | 1 |
| 23 | TGME49_264870.P2526 | Sodium:neurotransmitter symporter family protein | 10 | 10 | 9 | 9 | 6 | 7 | 4 | 1 |
| 28 | TGME49_264880-t26_1-p1 | NEDD8-activating enzyme E1 catalytic subunit | 9 | 9 | 9 | 8 | 8 | 7 | 6 | 1 |
| 25 | TGME49_265110-t26_1-p1 | ribosome biogenesis protein, putative | 10 | 10 | 10 | 10 | 8 | 7 | 8 | 1 |
| 3 | TGME49_265205.P2536 | WD domain, G-beta repeat-containing protein | 10 | 10 | 10 | 10 | 10 | 10 | 10 | 1 |
| 28 | TGME49_265310-t26_1-p1 | heat shock protein 40, putative | 8 | 8 | 8 | 8 | 8 | 7 | 7 | 1 |

| | | | | | | | | | | |
|---|---|---|---|---|---|---|---|---|---|---|
| 28 | TGME49_265330-t26_1-p1 | cell-cycle-associated protein kinase GSK, putative | 9 | 9 | 9 | 9 | 7 | 8 | 6 | 1 |
| 25 | TGME49_265410.P2538 | pre-mRNA-processing factor 17, putative | 10 | 9 | 9 | 9 | 9 | 7 | 8 | 1 |
| 28 | TGME49_265450-t26_1-p1 | hexokinase | 9 | 9 | 9 | 7 | 8 | 7 | 7 | 1 |
| 28 | TGME49_266060-t26_1-p1 | ribosomal protein RPSA | 8 | 8 | 8 | 8 | 8 | 7 | 7 | 1 |
| 6 | TGME49_266280-t26_1-p1 | AP-4 complex subunit beta, putative | 10 | 10 | 10 | 5 | 9 | 9 | 8 | 1 |
| 25 | TGME49_266400.P2567 | hypothetical protein | 10 | 10 | 9 | 9 | 7 | 8 | 8 | 1 |
| 28 | TGME49_266630.P2573 | conoid protein hub CPH1 | 10 | 9 | 9 | 7 | 8 | 6 | 6 | 1 |
| 25 | TGME49_266640.P2574 | acetyl-CoA synthetase ACS | 10 | 10 | 10 | 9 | 9 | 8 | 7 | 1 |
| 28 | TGME49_266730-t26_1-p1 | leucyl-tRNA synthetase (LeuRS2) | 10 | 9 | 10 | 9 | 8 | 6 | 6 | 1 |
| 28 | TGME49_266740.P2579 | RNA recognition motif-containing protein | 10 | 9 | 9 | 9 | 7 | 8 | 7 | 1 |
| 28 | TGME49_266800.P2582 | vacuolar iron transporter VIT | 8 | 8 | 8 | 8 | 8 | 7 | 7 | 1 |
| 3 | TGME49_266960.P2590 | beta tubulin | 9 | 9 | 9 | 9 | 9 | 9 | 9 | 1 |
| 3 | TGME49_266990-t26_1-p1 | beta-COP | 10 | 10 | 10 | 9 | 9 | 9 | 9 | 1 |
| 28 | TGME49_267040-t26_1-p1 | hypothetical protein | 9 | 9 | 9 | 7 | 9 | 8 | 6 | 1 |
| 6 | TGME49_267080-t26_1-p1 | 26S protease regulatory subunit 4, putative | 9 | 9 | 9 | 6 | 9 | 9 | 7 | 1 |
| 6 | TGME49_267270-t26_1-p1 | divalent metal transporter, putative | 10 | 9 | 9 | 6 | 7 | 7 | 6 | 1 |
| 3 | TGME49_267330.P2598 | fumarate hydratase | 10 | 9 | 9 | 9 | 9 | 9 | 9 | 1 |
| 6 | TGME49_267410-t26_1-p1 | scavenger receptor protein SR1 precursor | 10 | 9 | 9 | 3 | 7 | 8 | 6 | 1 |
| 28 | TGME49_267430-t26_1-p1 | DnaJ domain-containing protein | 9 | 9 | 8 | 8 | 8 | 6 | 6 | 1 |
| 28 | TGME49_267450-t26_1-p1 | alpha-tubulin suppressor protein | 10 | 10 | 10 | 9 | 9 | 7 | 6 | 1 |
| 25 | TGME49_267480-t26_1-p1 | tRNA (guanine(26)-N(2))-dimethyltransferase, putative | 10 | 10 | 9 | 9 | 7 | 8 | 8 | 1 |
| 23 | TGME49_267490-t26_1-p1 | cathepsin CPC3, putative | 10 | 6 | 7 | 9 | 7 | 8 | 3 | 1 |
| 6 | TGME49_267600.P2608 | FHA domain-containing protein | 9 | 9 | 3 | 3 | 6 | 8 | 8 | 1 |
| 25 | TGME49_267620.P2609 | IMC sutures component ISC6 | 9 | 9 | 9 | 8 | 9 | 8 | 8 | 1 |
| 3 | TGME49_267680-t26_1-p1 | microneme protein MIC12 | 10 | 10 | 10 | 10 | 10 | 10 | 10 | 1 |
| 3 | TGME49_267800-t26_1-p1 | dynamin-related protein DRPA | 10 | 9 | 10 | 9 | 9 | 9 | 9 | 1 |
| 28 | TGME49_268200.P2625 | RNA recognition motif-containing protein | 10 | 9 | 9 | 8 | 8 | 7 | 6 | 1 |
| 28 | TGME49_268210.P2626 | store potentiating/activating regulatory kinase SPARK | 9 | 9 | 8 | 8 | 6 | 7 | 7 | 1 |
| 25 | TGME49_268370.P2634 | non-specific serine/threonine protein kinase | 10 | 10 | 10 | 10 | 9 | 8 | 6 | 1 |
| 25 | TGME49_268590.P2641 | rhomboid protease ROM4 | 10 | 9 | 9 | 9 | 9 | 9 | 8 | 1 |
| 28 | TGME49_268630-t26_1-p1 | YagE family protein | 9 | 8 | 8 | 8 | 8 | 8 | 7 | 1 |
| 25 | TGME49_268640-t26_1-p1 | BING4CT (NUC141) domain-containing protein | 10 | 8 | 9 | 9 | 8 | 8 | 7 | 1 |
| 28 | TGME49_268710.P2645 | elongation factor Tu GTP binding domain-containing protein | 10 | 10 | 10 | 10 | 6 | 8 | 6 | 1 |
| 25 | TGME49_268830.P2651 | ATP synthase subunit delta | 9 | 9 | 9 | 8 | 8 | 8 | 8 | 1 |
| 3 | TGME49_268870.P2655 | tetratricopeptide repeat-containing protein | 10 | 10 | 10 | 10 | 9 | 9 | 9 | 1 |
| 28 | TGME49_268890-t26_1-p1 | citrate synthase I | 9 | 9 | 9 | 7 | 7 | 7 | 7 | 1 |
| 28 | TGME49_268950-t26_1-p1 | hypothetical protein | 9 | 9 | 9 | 7 | 8 | 7 | 6 | 1 |
| 28 | TGME49_269000.P2662 | ABC transporter family protein | 10 | 9 | 9 | 7 | 7 | 6 | 8 | 1 |
| 25 | TGME49_269110-t26_1-p1 | ornithine aminotransferase, mitochondrial precursor, putative | 9 | 9 | 9 | 9 | 8 | 8 | 8 | 1 |
| 28 | TGME49_269160-t26_1-p1 | outer dynein arm docking complex protein ODA1, putative | 9 | 9 | 9 | 9 | 8 | 7 | 7 | 1 |
| 6 | TGME49_269180.P2669 | MIF4G domain-containing protein | 10 | 10 | 10 | 5 | 9 | 8 | 6 | 1 |
| 6 | TGME49_269190.P2670 | glyceraldehyde-3-phosphate dehydrogenase GAPDH2 | 10 | 10 | 10 | 3 | 9 | 8 | 8 | 1 |
| 3 | TGME49_269200.P2671 | crooked neck family 1 protein isoform 2, putative | 10 | 10 | 10 | 10 | 10 | 9 | 9 | 1 |
| 28 | TGME49_269250.P2674 | Mov34/MPN/PAD-1 family protein | 9 | 8 | 8 | 8 | 8 | 7 | 7 | 1 |

| | | | | | | | | | | |
|---|---|---|---|---|---|---|---|---|---|---|
| 28 | TGME49_269340-t26_1-p1 | hypothetical protein | 9 | 9 | 8 | 8 | 7 | 6 | 6 | 1 |
| 25 | TGME49_269700.P2696 | NLI interacting factor family phosphatase | 10 | 10 | 10 | 8 | 9 | 8 | 8 | 1 |
| 28 | TGME49_269730.P2699 | myosin-light-chain kinase | 10 | 10 | 8 | 8 | 6 | 7 | 6 | 1 |
| 25 | TGME49_269740.P2700 | R3H domain-containing protein | 10 | 10 | 10 | 9 | 6 | 9 | 7 | 1 |
| 25 | TGME49_269770.P2702 | WD domain, G-beta repeat-containing protein | 10 | 10 | 10 | 9 | 9 | 8 | 8 | 1 |
| 25 | TGME49_269800.P2703 | glutamine-dependent NAD(+) synthetase protein, putative | 10 | 10 | 10 | 7 | 9 | 7 | 8 | 1 |
| 25 | TGME49_269840-t26_1-p1 | proteasome regulatory subunit | 8 | 8 | 8 | 8 | 8 | 8 | 8 | 1 |
| 25 | TGME49_269885.P2707 | rhoptry metalloprotease toxolysin TLN1 | 10 | 9 | 9 | 8 | 7 | 8 | 8 | 1 |
| 28 | TGME49_269920-t26_1-p1 | phosphatidylserine decarboxylase | 10 | 9 | 8 | 8 | 7 | 6 | 6 | 1 |
| 28 | TGME49_269930.P2709 | calcium-binding egf domain-containing protein CBDP3 | 10 | 10 | 9 | 8 | 7 | 8 | 7 | 1 |
| 3 | TGME49_269980-t26_1-p1 | preprotein translocase Sec61, putative | 9 | 9 | 9 | 9 | 9 | 9 | 9 | 1 |
| 28 | TGME49_269990.P2712 | hypothetical protein | 10 | 10 | 10 | 8 | 6 | 7 | 7 | 1 |
| 25 | TGME49_270100.P2719 | oxidoreductase, aldo/keto reductase family protein | 10 | 10 | 10 | 10 | 9 | 9 | 6 | 1 |
| 25 | TGME49_270510-t26_1-p1 | asparaginyl-tRNA synthetase (NOB+tRNA synthase) | 10 | 9 | 9 | 7 | 9 | 9 | 9 | 1 |
| 28 | TGME49_270550.P2772 | gamma-glutamyl phosphate reductase, putative | 9 | 9 | 9 | 8 | 8 | 7 | 7 | 1 |
| 3 | TGME49_270690.P2777 | dynamin-related protein DRPC | 10 | 10 | 10 | 10 | 9 | 10 | 9 | 1 |
| 3 | TGME49_270840.P2786 | poly(ADP-ribose) polymerase catalytic domain-containing protein | 10 | 10 | 10 | 8 | 9 | 9 | 9 | 1 |
| 25 | TGME49_270850-t26_1-p1 | glycosyl hydrolases family 17 protein | 10 | 9 | 9 | 9 | 8 | 8 | 8 | 1 |
| 3 | TGME49_270865.P2787 | adenylate cyclase beta | 10 | 10 | 10 | 10 | 10 | 10 | 10 | 1 |
| 28 | TGME49_270910.P2789 | glycerol 3-phosphate acyltransferase ATS1 | 9 | 9 | 8 | 8 | 7 | 6 | 6 | 1 |
| 25 | TGME49_270930.P2791 | THO complex subunit 2, putative | 10 | 10 | 10 | 10 | 7 | 8 | 7 | 1 |
| 25 | TGME49_271060.P2798 | vacuolar protein sorting-associated protein 45 | 10 | 10 | 9 | 9 | 9 | 8 | 8 | 1 |
| 28 | TGME49_271120.P2802 | endonuclease/exonuclease/phosphatase family protein | 10 | 9 | 9 | 9 | 8 | 7 | 6 | 1 |
| 25 | TGME49_271210-t26_1-p1 | 4-alpha-glucanotransferase | 10 | 10 | 10 | 9 | 9 | 8 | 7 | 1 |
| 28 | TGME49_271240.P2805 | hypothetical protein | 10 | 10 | 9 | 7 | 7 | 6 | 6 | 1 |
| 28 | TGME49_271440.P2815 | nuclear protein localization protein 4, putative | 9 | 9 | 9 | 8 | 8 | 8 | 6 | 1 |
| 25 | TGME49_271740.P2821 | zinc finger protein, putative | 10 | 10 | 10 | 10 | 8 | 8 | 6 | 1 |
| 3 | TGME49_271780.P2823 | Filamin/ABP280 repeat-containing protein | 10 | 10 | 10 | 10 | 9 | 9 | 8 | 1 |
| 28 | TGME49_271920.P2830 | hypothetical protein | 10 | 9 | 9 | 9 | 7 | 7 | 6 | 1 |
| 28 | TGME49_271970-t26_1-p1 | glideosome-associated protein with multiple-membrane spans GAPM3 | 8 | 8 | 8 | 8 | 8 | 7 | 7 | 1 |
| 23 | TGME49_272240-t26_1-p1 | Toxoplasma gondii family D protein | 10 | 10 | 9 | 9 | 9 | 8 | 5 | 1 |
| 25 | TGME49_272290.P2850 | pyruvate dehydrogenase complex subunit PD-HE1Beta | 9 | 9 | 9 | 8 | 8 | 8 | 8 | 1 |
| 28 | TGME49_272420.P2862 | phosphatidylcholine-sterol acyltransferase | 10 | 9 | 9 | 8 | 8 | 7 | 7 | 1 |
| 28 | TGME49_272500.P2868 | sugar transporter ST2 | 10 | 9 | 9 | 8 | 7 | 7 | 7 | 1 |
| 23 | TGME49_272720.P2882 | methyltransferase domain-containing protein | 10 | 10 | 9 | 9 | 9 | 6 | 4 | 1 |
| 3 | TGME49_272740.P2884 | DNA-directed RNA polymerase III RPC2 | 10 | 10 | 10 | 10 | 10 | 10 | 10 | 1 |
| 25 | TGME49_272900.P2887 | DNA repair protein RAD51 | 9 | 9 | 8 | 8 | 8 | 8 | 8 | 1 |
| 3 | TGME49_272910-t26_1-p1 | T-complex protein 1 delta subunit | 9 | 9 | 9 | 9 | 9 | 9 | 9 | 1 |
| 3 | TGME49_273090-t26_1-p1 | cell division protein CDC48CY | 10 | 10 | 10 | 10 | 10 | 10 | 10 | 1 |
| 25 | TGME49_273140.P2893 | radical SAM methylthiotransferase, MiaB/RimO family protein | 10 | 9 | 9 | 8 | 7 | 8 | 8 | 1 |
| 23 | TGME49_273370.P2899 | coatomer gamma 2-subunit protein, putative | 10 | 10 | 10 | 10 | 10 | 8 | 5 | 1 |

| | | | | | | | | | | |
|---|---|---|---|---|---|---|---|---|---|---|
| 25 | TGME49_273380.P2900 | ion channel protein | 10 | 10 | 10 | 8 | 9 | 8 | 8 | 1 |
| 28 | TGME49_273445.P2903 | iron-sulfur cluster assembly protein SufD | 10 | 10 | 8 | 8 | 7 | 7 | 7 | 1 |
| 25 | TGME49_273460.P2904 | eukaryotic translation initiation factor 3 subunit 6 interacting protein | 10 | 10 | 10 | 9 | 9 | 8 | 8 | 1 |
| 23 | TGME49_273490.P2905 | glutamine synthetase, type I, putative | 9 | 9 | 9 | 9 | 4 | 8 | 3 | 1 |
| 25 | TGME49_273520.P2907 | eukaryotic translation initiation factor 3 subunit E, putative | 10 | 9 | 9 | 7 | 9 | 8 | 8 | 1 |
| 28 | TGME49_273540-t26_1-p1 | phosphatidylserine synthase, putative | 9 | 9 | 8 | 8 | 8 | 6 | 7 | 1 |
| 25 | TGME49_273560.P2909 | kinesin heavy chain, putative | 10 | 10 | 10 | 8 | 9 | 6 | 8 | 1 |
| 28 | TGME49_273730.P2917 | phospholipase, patatin family protein | 10 | 9 | 9 | 9 | 6 | 8 | 7 | 1 |
| 25 | TGME49_273740-t26_1-p1 | acetyl-CoA acyltransferase B, putative | 9 | 8 | 9 | 8 | 7 | 8 | 8 | 1 |
| 3 | TGME49_273760-t26_1-p1 | heat shock protein HSP70 | 10 | 10 | 10 | 10 | 10 | 10 | 10 | 1 |
| 23 | TGME49_273800.P2921 | WD domain, G-beta repeat-containing protein | 10 | 10 | 10 | 8 | 7 | 6 | 3 | 1 |
| 3 | TGME49_273870.P2924 | SWI2/SNF2 ISWI-like (AT hook) | 10 | 10 | 10 | 9 | 10 | 10 | 9 | 1 |
| 23 | TGME49_273950.P2927 | replication factor C subunit 5, putative | 9 | 8 | 8 | 8 | 8 | 7 | 3 | 1 |
| 28 | TGME49_274060-t26_1-p1 | 2-oxoglutarate/malate translocase OMT | 8 | 8 | 8 | 8 | 8 | 7 | 7 | 1 |
| 28 | TGME49_274190.P2946 | eukaryotic initiation factor 2B epsilon subunit, putative | 10 | 10 | 10 | 9 | 7 | 7 | 7 | 1 |
| 25 | TGME49_275320.P2948 | penicillin amidase | 10 | 10 | 10 | 10 | 8 | 7 | 8 | 1 |
| 6 | TGME49_275350.P2949 | TBC domain-containing protein TBC1 | 10 | 10 | 9 | 4 | 7 | 7 | 6 | 1 |
| 23 | TGME49_275430.P2952 | hypothetical protein | 10 | 10 | 10 | 9 | 8 | 9 | 3 | 1 |
| 25 | TGME49_275480.P2954 | pre-mRNA-splicing factor CEF1, putative | 10 | 10 | 10 | 7 | 9 | 8 | 8 | 1 |
| 25 | TGME49_275630-t26_1-p1 | HECT-domain (ubiquitin-transferase) domain-containing protein | 10 | 10 | 10 | 9 | 9 | 8 | 8 | 1 |
| 25 | TGME49_275650-t26_1-p1 | hypothetical protein | 10 | 10 | 10 | 9 | 7 | 8 | 8 | 1 |
| 3 | TGME49_275690.P2959 | chaperone protein ClpB3 | 10 | 3 | 10 | 10 | 10 | 10 | 10 | 1 |
| 6 | TGME49_276130.P2965 | cathepsin CPC2 | 10 | 9 | 9 | 3 | 6 | 9 | 6 | 1 |
| 23 | TGME49_276170-t26_1-p1 | phosphatidylinositol 3- and 4-kinase | 10 | 10 | 9 | 9 | 9 | 8 | 3 | 1 |
| 28 | TGME49_276180-t26_1-p1 | histone acetyltransferase TAF1/250 | 10 | 10 | 10 | 10 | 6 | 9 | 6 | 1 |
| 3 | TGME49_277000-t26_1-p1 | transport protein Sec24, putative | 10 | 9 | 10 | 10 | 9 | 8 | 9 | 1 |
| 25 | TGME49_277500-t26_1-p1 | 26S proteasome regulatory subunit 7, putative | 9 | 9 | 9 | 8 | 8 | 9 | 9 | 1 |
| 6 | TGME49_277510.P2981 | cytoplasmic dynein intermediate chain | 10 | 10 | 9 | 6 | 7 | 8 | 7 | 1 |
| 25 | TGME49_277720-t26_1-p1 | GDA1/CD39 (nucleoside phosphatase) family protein | 10 | 9 | 9 | 9 | 7 | 8 | 8 | 1 |
| 3 | TGME49_277760-t26_1-p1 | adenylosuccinate lyase, putative | 10 | 10 | 10 | 9 | 9 | 9 | 9 | 1 |
| 25 | TGME49_277850-t26_1-p1 | trypsin domain-containing protein | 10 | 9 | 9 | 9 | 9 | 9 | 8 | 1 |
| 6 | TGME49_277860.P2987 | hypothetical protein | 9 | 8 | 8 | 5 | 6 | 8 | 8 | 1 |
| 3 | TGME49_278110-t26_1-p1 | 1,3-beta-glucan synthase component protein | 10 | 10 | 10 | 10 | 9 | 10 | 9 | 1 |
| 3 | TGME49_278440-t26_1-p1 | SWI2/SNF2 Brahma-like putative | 10 | 10 | 10 | 7 | 10 | 10 | 9 | 1 |
| 6 | TGME49_278500-t26_1-p1 | ribosomal RNA large subunit methyltransferase J protein | 10 | 10 | 10 | 5 | 8 | 8 | 7 | 1 |
| 25 | TGME49_278550-t26_1-p1 | elongation factor Tu GTP binding domain-containing protein | 10 | 10 | 10 | 9 | 9 | 6 | 8 | 1 |
| 28 | TGME49_278630-t26_1-p1 | tetratricopeptide repeat-containing protein | 10 | 9 | 9 | 9 | 8 | 6 | 6 | 1 |
| 3 | TGME49_278660.P3007 | P-type ATPase4 | 10 | 10 | 10 | 10 | 10 | 9 | 10 | 1 |
| 23 | TGME49_278830.P3016 | glucose-6-phosphate 1-dehydrogenase | 10 | 3 | 10 | 8 | 9 | 9 | 3 | 1 |
| 3 | TGME49_278870-t26_1-p1 | myosin F | 10 | 10 | 10 | 10 | 9 | 10 | 8 | 1 |
| 23 | TGME49_278975.P3026 | metacaspase MCA2 | 10 | 10 | 9 | 7 | 7 | 7 | 4 | 1 |
| 23 | TGME49_278990.P3027 | phosphate transporter MPT | 9 | 8 | 8 | 8 | 8 | 8 | 3 | 1 |
| 28 | TGME49_279330-t26_1-p1 | DEAD/DEAH box helicase family protein | 9 | 9 | 9 | 9 | 9 | 7 | 6 | 1 |

| | | | | | | | | | | |
|---|---|---|---|---|---|---|---|---|---|---|
| 23 | TGME49_279390.P3031 | proliferation-associated protein 2G4, putative | 9 | 9 | 9 | 8 | 8 | 7 | 5 | 1 |
| 25 | TGME49_279420-t26_1-p1 | hypothetical protein | 10 | 10 | 9 | 9 | 8 | 8 | 8 | 1 |
| 25 | TGME49_279450-t26_1-p1 | adenylosuccinate synthetase, putative | 9 | 9 | 9 | 8 | 8 | 8 | 8 | 1 |
| 28 | TGME49_280400.P3034 | Dpy-19-like C-mannosyltransferase | 10 | 10 | 10 | 9 | 6 | 8 | 7 | 1 |
| 28 | TGME49_280410.P3035 | 3'5'-cyclic nucleotide phosphodiesterase PDE7 | 10 | 10 | 10 | 8 | 7 | 7 | 7 | 1 |
| 25 | TGME49_280490-t26_1-p1 | U-box domain-containing protein | 10 | 10 | 10 | 10 | 8 | 7 | 8 | 1 |
| 6 | TGME49_280500-t26_1-p1 | inorganic anion transporter, sulfate permease (SulP) family protein | 10 | 10 | 9 | 3 | 9 | 6 | 7 | 1 |
| 25 | TGME49_280600-t26_1-p1 | histidyl-tRNA synthetase (HisRS), putative | 10 | 10 | 10 | 10 | 7 | 8 | 8 | 1 |
| 28 | TGME49_280610.P3047 | signal recognition particle receptor alpha subunit | 10 | 9 | 9 | 9 | 6 | 8 | 8 | 1 |
| 3 | TGME49_280700.P3050 | glutamate decarboxylase | 10 | 10 | 10 | 10 | 9 | 10 | 10 | 1 |
| 25 | TGME49_280800-t26_1-p1 | SWI2/SNF2 SRCAP/Ino80 | 10 | 10 | 10 | 9 | 8 | 8 | 8 | 1 |
| 6 | TGME49_281450.P3061 | cell-cycle-associated protein kinase, putative | 9 | 9 | 8 | 5 | 8 | 7 | 5 | 1 |
| 28 | TGME49_281980.P3075 | phosphatidate cytidylyltransferase | 10 | 9 | 9 | 8 | 6 | 7 | 7 | 1 |
| 25 | TGME49_282200.P3081 | chaperone protein ClpB4, putative | 10 | 10 | 10 | 10 | 7 | 8 | 8 | 1 |
| 3 | TGME49_282230-t26_1-p1 | sulfate adenylyltransferase/adenylylsulfate kinase | 10 | 10 | 9 | 9 | 9 | 9 | 9 | 1 |
| 28 | TGME49_283480.P3084 | cell-cycle-associated protein kinase DYRK2, putative | 9 | 10 | 9 | 8 | 8 | 7 | 7 | 1 |
| 25 | TGME49_283490.P3085 | Alpha-amylase AMY3, putative | 10 | 8 | 8 | 9 | 9 | 9 | 8 | 1 |
| 25 | TGME49_283780.P3101 | glucose-6-phosphate isomerase GPI | 10 | 9 | 9 | 9 | 9 | 8 | 8 | 1 |
| 28 | TGME49_283850-t26_1-p1 | peptidyl-prolyl cis-trans isomerase | 9 | 9 | 9 | 7 | 8 | 7 | 8 | 1 |
| 6 | TGME49_283900.P3104 | origin recognition complex subunit 1, putative | 10 | 9 | 9 | 5 | 7 | 8 | 7 | 1 |
| 28 | TGME49_284780.P3116 | hypothetical protein | 10 | 10 | 10 | 10 | 9 | 6 | 6 | 1 |
| 3 | TGME49_285200.P3121 | cleavage and polyadenylation specificity factor subunit 3 | 10 | 10 | 10 | 9 | 9 | 9 | 9 | 1 |
| 3 | TGME49_285470.P3132 | patched family protein | 10 | 10 | 10 | 10 | 9 | 9 | 9 | 1 |
| 23 | TGME49_285660.P3135 | DEAD/DEAH box helicase domain-containing protein | 10 | 10 | 10 | 10 | 9 | 8 | 3 | 1 |
| 28 | TGME49_285950-t26_1-p1 | RNA methyltransferase, putative | 10 | 9 | 9 | 8 | 7 | 7 | 6 | 1 |
| 3 | TGME49_285970-t26_1-p1 | 30S ribosomal protein S5, putative | 10 | 10 | 10 | 10 | 9 | 9 | 8 | 1 |
| 3 | TGME49_285980.P3143 | phosphoglucomutase PGM1 | 10 | 10 | 10 | 9 | 8 | 9 | 9 | 1 |
| 28 | TGME49_285990-t26_1-p1 | Filamin/ABP280 repeat-containing protein | 10 | 10 | 9 | 9 | 6 | 7 | 7 | 1 |
| 3 | TGME49_286080-t26_1-p1 | elongation factor 2 family protein | 10 | 10 | 10 | 9 | 9 | 10 | 8 | 1 |
| 6 | TGME49_286120-t26_1-p1 | prolyl endopeptidase | 10 | 10 | 10 | 3 | 8 | 9 | 9 | 1 |
| 28 | TGME49_286210.P3151 | protein phosphatase 4, catalytic subunit | 8 | 8 | 8 | 7 | 6 | 8 | 8 | 1 |
| 25 | TGME49_286240.P3153 | kelch repeat protein, putative | 10 | 9 | 9 | 7 | 7 | 8 | 8 | 1 |
| 28 | TGME49_286250-t26_1-p1 | hypothetical protein | 9 | 9 | 9 | 9 | 7 | 7 | 7 | 1 |
| 25 | TGME49_286410-t26_1-p1 | hypothetical protein | 10 | 10 | 10 | 10 | 7 | 8 | 7 | 1 |
| 28 | TGME49_286490-t26_1-p1 | G protein-coupled receptor 89 isoform 5, putative | 10 | 8 | 9 | 9 | 6 | 7 | 7 | 1 |
| 6 | TGME49_286690-t26_1-p1 | DNA/RNA-binding protein KIN17 domain-containing protein | 9 | 9 | 9 | 6 | 8 | 7 | 8 | 1 |
| 28 | TGME49_286750.P3163 | MA3 domain-containing protein | 9 | 9 | 9 | 9 | 8 | 7 | 7 | 1 |
| 3 | TGME49_286932-t26_1-p1 | ATPase family associated with various cellular activities (AAA) domain-containing protein | 10 | 10 | 10 | 8 | 9 | 10 | 10 | 1 |
| 23 | TGME49_287160.P3166 | kinesin-like protein KIF24, putative | 10 | 10 | 9 | 8 | 8 | 8 | 3 | 1 |
| 3 | TGME49_287500-t26_1-p1 | T complex chaperonin, putative | 10 | 9 | 9 | 9 | 9 | 9 | 9 | 1 |
| 6 | TGME49_288060-t26_1-p1 | hypothetical protein | 9 | 8 | 8 | 3 | 7 | 7 | 6 | 1 |
| 28 | TGME49_288210.P3179 | PUL domain-containing protein | 10 | 9 | 9 | 9 | 7 | 7 | 7 | 1 |
| 3 | TGME49_288380.P3182 | heat shock protein HSP90 | 10 | 10 | 10 | 9 | 10 | 10 | 10 | 1 |
| 6 | TGME49_288400-t26_1-p1 | LETM1 family protein | 10 | 9 | 9 | 6 | 8 | 8 | 7 | 1 |

| | | | | | | | | | | |
|---|---|---|---|---|---|---|---|---|---|---|
| 25 | TGME49_288460-t26_1-p1 | hypothetical protein | 10 | 10 | 10 | 9 | 10 | 8 | 7 | 1 |
| 25 | TGME49_288500-t26_1-p1 | malate:quinone oxidoreductase | 10 | 9 | 9 | 9 | 9 | 9 | 8 | 1 |
| 25 | TGME49_288530-t26_1-p1 | NOL1/NOP2/sun family protein | 10 | 10 | 9 | 9 | 9 | 8 | 8 | 1 |
| 28 | TGME49_288600.P3183 | ATG18/WIPI-like protein PROP1 | 10 | 7 | 9 | 9 | 9 | 8 | 6 | 1 |
| 23 | TGME49_288700.P3186 | structural maintenance of chromosomes protein SMC1 | 10 | 10 | 10 | 8 | 9 | 7 | 5 | 1 |
| 28 | TGME49_288740-t26_1-p1 | L-asparaginase, putative | 10 | 9 | 9 | 8 | 6 | 7 | 7 | 1 |
| 28 | TGME49_288800.P3190 | endonuclease/exonuclease/phosphatase family protein | 10 | 9 | 9 | 8 | 6 | 6 | 7 | 1 |
| 25 | TGME49_288830-t26_1-p1 | NADH dehydrogenase (NDH2-II) | 10 | 9 | 9 | 8 | 8 | 8 | 7 | 1 |
| 25 | TGME49_288860.P3194 | RuvB family 2 protein | 9 | 9 | 9 | 7 | 9 | 8 | 9 | 1 |
| 28 | TGME49_289010-t26_1-p1 | RNA recognition motif-containing protein | 10 | 10 | 9 | 9 | 8 | 7 | 7 | 1 |
| 28 | TGME49_289070-t26_1-p1 | E1-E2 ATPase subfamily protein | 10 | 10 | 10 | 8 | 6 | 7 | 7 | 1 |
| 25 | TGME49_289080-t26_1-p1 | AP-3 complex subunit beta, putative | 10 | 10 | 10 | 9 | 7 | 8 | 8 | 1 |
| 6 | TGME49_289100.P3198 | protein hook | 10 | 10 | 9 | 6 | 9 | 7 | 6 | 1 |
| 3 | TGME49_289170-t26_1-p1 | adenylate cyclase alpha 1 | 10 | 10 | 10 | 9 | 10 | 9 | 9 | 1 |
| 28 | TGME49_289190-t26_1-p1 | tetratricopeptide repeat-containing protein | 10 | 9 | 9 | 8 | 7 | 7 | 7 | 1 |
| 23 | TGME49_289200-t26_1-p1 | RAP domain-containing protein | 10 | 9 | 9 | 8 | 8 | 6 | 5 | 1 |
| 25 | TGME49_289300-t26_1-p1 | methionyl-tRNA synthetase | 10 | 10 | 10 | 8 | 9 | 8 | 8 | 1 |
| 25 | TGME49_289310-t26_1-p1 | cullin family protein | 10 | 10 | 10 | 8 | 7 | 9 | 9 | 1 |
| 25 | TGME49_289320-t26_1-p1 | casein kinase I, putative | 9 | 9 | 9 | 9 | 8 | 8 | 7 | 1 |
| 25 | TGME49_289330-t26_1-p1 | ubiquitin carboxyl-terminal hydrolase family 2 protein | 10 | 10 | 10 | 8 | 6 | 9 | 9 | 1 |
| 25 | TGME49_289350-t26_1-p1 | ATP-binding cassette G family transporter ABCG84 | 10 | 10 | 9 | 9 | 9 | 8 | 8 | 1 |
| 6 | TGME49_289550-t26_1-p1 | WD domain, G-beta repeat-containing protein | 9 | 9 | 9 | 6 | 8 | 7 | 6 | 1 |
| 25 | TGME49_289620-t26_1-p1 | cathepsin CPC1 | 10 | 10 | 9 | 9 | 9 | 8 | 8 | 1 |
| 3 | TGME49_289650.P3206 | PEP-carboxykinase I | 10 | 10 | 10 | 10 | 9 | 9 | 9 | 1 |
| 25 | TGME49_289690.P3209 | glyceraldehyde-3-phosphate dehydrogenase GAPDH1 | 9 | 9 | 8 | 8 | 8 | 8 | 8 | 1 |
| 23 | TGME49_289730.P3212 | vacuolar protein sorting-associated protein 18 | 10 | 10 | 10 | 9 | 8 | 7 | 5 | 1 |
| 25 | TGME49_289770-t26_1-p1 | AP-1 complex subunit mu-1, putative | 9 | 9 | 9 | 9 | 8 | 8 | 8 | 1 |
| 6 | TGME49_289780-t26_1-p1 | ATP-dependent hsl protease ATP-binding subunit hslU, putative | 10 | 9 | 9 | 5 | 7 | 8 | 8 | 1 |
| 25 | TGME49_289830-t26_1-p1 | eukaryotic initiation factor-3, delta subunit, putative | 9 | 9 | 9 | 8 | 8 | 8 | 7 | 1 |
| 6 | TGME49_289940.P3218 | uroporphyrinogen decarboxylase | 10 | 9 | 9 | 6 | 7 | 7 | 7 | 1 |
| 3 | TGME49_290160.P3223 | sortilin-like receptor SORTLR | 10 | 10 | 10 | 9 | 9 | 10 | 10 | 1 |
| 3 | TGME49_290170-t26_1-p1 | protein phosphatase PPKL | 10 | 10 | 10 | 10 | 9 | 10 | 8 | 1 |
| 6 | TGME49_290200-t26_1-p1 | NAD/NADP octopine/nopaline dehydrogenase, alpha-helical domain-containing protein | 9 | 9 | 9 | 3 | 8 | 7 | 7 | 1 |
| 28 | TGME49_290240-t26_1-p1 | TCF25 domain-containing protein | 10 | 9 | 9 | 8 | 7 | 7 | 7 | 1 |
| 25 | TGME49_290270-t26_1-p1 | SPRY domain-containing protein | 9 | 9 | 9 | 9 | 8 | 7 | 8 | 1 |
| 3 | TGME49_290290-t26_1-p1 | ubiquitin-activating enzyme E1 family protein | 10 | 10 | 10 | 10 | 9 | 8 | 9 | 1 |
| 6 | TGME49_290580-t26_1-p1 | ATP-binding cassette G family transporter ABCG89 | 10 | 10 | 9 | 6 | 8 | 8 | 7 | 1 |
| 25 | TGME49_290600-t26_1-p1 | succinyl-CoA-synthetase alpha SCSA | 8 | 8 | 8 | 8 | 8 | 8 | 8 | 1 |
| 25 | TGME49_290640-t26_1-p1 | DNA mismatch repair protein MSH6-1, putative | 10 | 10 | 10 | 10 | 8 | 9 | 6 | 1 |
| 28 | TGME49_290660.P3228 | RNA recognition motif-containing protein | 9 | 9 | 8 | 8 | 8 | 7 | 7 | 1 |
| 28 | TGME49_290670.P3229 | leucyl aminopeptidase LAP | 10 | 9 | 9 | 9 | 8 | 7 | 7 | 1 |
| 28 | TGME49_290720.P3232 | V-type proton ATPase subunit a2, putative | 10 | 10 | 10 | 7 | 7 | 9 | 6 | 1 |
| 6 | TGME49_290840.P3234 | serine protease | 10 | 9 | 9 | 6 | 8 | 8 | 7 | 1 |

| | | | | | | | | | | |
|---|---|---|---|---|---|---|---|---|---|---|
| 25 | TGME49_290850.P3235 | WD-40 repeat protein | 9 | 9 | 9 | 8 | 9 | 7 | 9 | 1 |
| 25 | TGME49_290940-t26_1-p1 | EMP/nonaspanin domain family protein | 10 | 9 | 10 | 9 | 8 | 9 | 8 | 1 |
| 3 | TGME49_290950-t26_1-p1 | clathrin heavy chain | 10 | 10 | 10 | 10 | 10 | 10 | 10 | 1 |
| 3 | TGME49_290960-t26_1-p1 | pyruvate phosphate dikinase, pep/pyruvate binding domain-containing protein | 10 | 10 | 10 | 8 | 8 | 9 | 9 | 1 |
| 25 | TGME49_290980.P3242 | serine palmitoyltransferase SPT1 | 10 | 10 | 9 | 9 | 9 | 8 | 8 | 1 |
| 25 | TGME49_291020.P3244 | myosin L | 10 | 10 | 10 | 7 | 8 | 9 | 7 | 1 |
| 23 | TGME49_291140-t26_1-p1 | CCR4-Not complex component, Not1 protein | 10 | 10 | 10 | 9 | 9 | 8 | 4 | 1 |
| 3 | TGME49_291180.P3247 | vacuolar protein sorting-associated protein 13, putative | 10 | 10 | 10 | 10 | 9 | 10 | 9 | 1 |
| 23 | TGME49_291680-t26_1-p1 | protein transport protein SEC23, putative | 10 | 10 | 10 | 10 | 10 | 10 | 3 | 1 |
| 23 | TGME49_291820.P3250 | RNA helicase (UPF2 interacting domain) protein | 10 | 10 | 10 | 9 | 10 | 8 | 5 | 1 |
| 25 | TGME49_292010-t26_1-p1 | transcription initiation factor IIB | 9 | 9 | 9 | 8 | 7 | 8 | 8 | 1 |
| 3 | TGME49_292020.P3257 | cysteine repeat modular protein CRMPB | 10 | 10 | 10 | 10 | 10 | 10 | 10 | 1 |
| 3 | TGME49_292080-t26_1-p1 | leucyl-tRNA synthetase | 10 | 10 | 10 | 10 | 9 | 10 | 10 | 1 |
| 28 | TGME49_292120-t26_1-p1 | membrane occupation and recognition nexus protein MORN2 | 9 | 8 | 8 | 8 | 8 | 7 | 7 | 1 |
| 28 | TGME49_292140-t26_1-p1 | NIMA-related protein kinase NIMA1 | 10 | 10 | 9 | 8 | 8 | 7 | 7 | 1 |
| 25 | TGME49_292170-t26_1-p1 | histone lysine methyltransferase, SET, putative | 10 | 10 | 10 | 10 | 8 | 8 | 9 | 1 |
| 28 | TGME49_292220-t26_1-p1 | PCI domain-containing protein | 9 | 9 | 9 | 7 | 7 | 7 | 7 | 1 |
| 25 | TGME49_292920-t26_1-p1 | heat shock protein 75, putative | 10 | 10 | 10 | 9 | 9 | 9 | 8 | 1 |
| 25 | TGME49_293000.P3270 | 3'5'-cyclic nucleotide phosphodiesterase PDE2 | 10 | 10 | 10 | 8 | 8 | 8 | 7 | 1 |
| 25 | TGME49_293180.P3275 | NADP-specific glutamate dehydrogenase | 9 | 9 | 9 | 8 | 9 | 8 | 8 | 1 |
| 6 | TGME49_293190.P3276 | endonuclease/exonuclease/phosphatase family protein | 10 | 9 | 8 | 5 | 7 | 7 | 7 | 1 |
| 25 | TGME49_293368-t26_1-p1 | TIP49 C-terminus family protein | 9 | 3 | 9 | 9 | 9 | 8 | 9 | 1 |
| 23 | TGME49_293570.P3309 | translocation protein sec62, putative | 9 | 8 | 8 | 8 | 8 | 6 | 4 | 1 |
| 25 | TGME49_293590.P3311 | 3-oxoacyl-acyl-carrier protein synthase I/II, putative | 9 | 9 | 9 | 9 | 9 | 8 | 8 | 1 |
| 28 | TGME49_293730.P3322 | palmitoyltransferase DHHC14 | 10 | 10 | 9 | 8 | 7 | 7 | 7 | 1 |
| 28 | TGME49_293810.P3330 | carboxyvinyl-carboxyphosphonate phosphorylmutase | 8 | 8 | 8 | 8 | 7 | 7 | 7 | 1 |
| 25 | TGME49_293820.P3331 | calpain family cysteine protease domain-containing protein | 10 | 10 | 10 | 8 | 8 | 9 | 7 | 1 |
| 28 | TGME49_294270-t26_1-p1 | histone arginine methyltransferase PRMT4/CARM1 | 10 | 9 | 9 | 8 | 6 | 6 | 7 | 1 |
| 25 | TGME49_294280-t26_1-p1 | E3 ubiquitin-protein ligase, putative | 9 | 10 | 10 | 9 | 8 | 8 | 7 | 1 |
| 25 | TGME49_294330.P3351 | apical membrane antigen AMA4 | 10 | 10 | 10 | 10 | 9 | 7 | 8 | 1 |
| 3 | TGME49_294350.P3353 | DEAD/DEAH box helicase domain-containing protein | 10 | 10 | 10 | 9 | 10 | 10 | 8 | 1 |
| 3 | TGME49_294360.P3354 | ubiquitin specific protease 39 isoform 2, putative | 10 | 9 | 9 | 9 | 9 | 9 | 9 | 1 |
| 28 | TGME49_294430-t26_1-p1 | protein SDA1, putative | 9 | 9 | 9 | 8 | 8 | 8 | 6 | 1 |
| 3 | TGME49_294550-t26_1-p1 | dynein heavy chain | 10 | 10 | 10 | 10 | 10 | 10 | 10 | 1 |
| 23 | TGME49_294630.P3363 | PH domain-containing protein, putative | 10 | 9 | 9 | 8 | 8 | 7 | 5 | 1 |
| 3 | TGME49_294640-t26_1-p1 | ribonucleoside-diphosphate reductase large chain | 10 | 10 | 10 | 10 | 10 | 10 | 10 | 1 |
| 25 | TGME49_294770-t26_1-p1 | Armadillo/beta-catenin family repeat-containing protein | 10 | 10 | 10 | 9 | 9 | 8 | 8 | 1 |
| 3 | TGME49_294820.P3369 | type I fatty acid synthase, putative | 10 | 10 | 10 | 10 | 10 | 10 | 10 | 1 |
| 23 | TGME49_294898-t26_1-p1 | tetratricopeptide repeat-containing protein | 10 | 10 | 10 | 9 | 8 | 8 | 3 | 1 |
| 23 | TGME49_295000.P3377 | vacuolar protein sorting-associated protein VPS33 | 10 | 10 | 10 | 9 | 8 | 7 | 5 | 1 |

| | | | | | | | | | | |
|---|---|---|---|---|---|---|---|---|---|---|
| 3 | TGME49_295040-t26_1-p1 | HEAT repeat-containing protein | 10 | 10 | 10 | 10 | 7 | 9 | 9 | 1 |
| 23 | TGME49_295320-t26_1-p1 | embryo sac development arrest EDA7, putative | 10 | 10 | 9 | 9 | 9 | 8 | 5 | 1 |
| 25 | TGME49_295710-t26_1-p1 | HECT-domain (ubiquitin-transferase) domain-containing protein | 10 | 10 | 10 | 8 | 9 | 7 | 8 | 1 |
| 28 | TGME49_296010.P3396 | phosphatidylinositol 3- and 4-kinase | 10 | 9 | 3 | 8 | 8 | 8 | 8 | 1 |
| 28 | TGME49_297060.P3398 | phosphoglycerate mutase PGMII | 8 | 8 | 8 | 8 | 7 | 7 | 7 | 1 |
| 28 | TGME49_297070.P3399 | carbonic anhydrase-related protein CARP | 9 | 9 | 9 | 9 | 6 | 8 | 8 | 1 |
| 28 | TGME49_297110-t26_1-p1 | kinesin motor domain-containing protein | 10 | 10 | 10 | 9 | 7 | 7 | 6 | 1 |
| 25 | TGME49_297220-t26_1-p1 | acyl-CoA synthetase ACS1 | 10 | 10 | 10 | 9 | 8 | 8 | 8 | 1 |
| 28 | TGME49_297320-t26_1-p1 | hypothetical protein | 10 | 9 | 9 | 9 | 6 | 7 | 7 | 1 |
| 23 | TGME49_297420-t26_1-p1 | beta-tubulin cofactor D, putative | 10 | 10 | 10 | 8 | 7 | 7 | 4 | 1 |
| 6 | TGME49_297500-t26_1-p1 | T-complex protein 1 eta subunit | 10 | 9 | 9 | 6 | 9 | 8 | 9 | 1 |
| 3 | TGME49_297530-t26_1-p1 | DNA-directed RNA polymerase I RPA2 | 10 | 10 | 10 | 10 | 9 | 9 | 10 | 1 |
| 3 | TGME49_297720-t26_1-p1 | trehalose-phosphatase | 10 | 10 | 10 | 9 | 9 | 10 | 9 | 1 |
| 23 | TGME49_297800.P3419 | structural maintenance of chromosomes protein SMC2, putative | 10 | 10 | 10 | 7 | 10 | 10 | 3 | 1 |
| 28 | TGME49_297820.P3421 | sperm associated antigen 6, putative | 9 | 9 | 9 | 9 | 9 | 7 | 6 | 1 |
| 6 | TGME49_297840-t26_1-p1 | DNA primase, large subunit | 10 | 9 | 9 | 6 | 8 | 8 | 7 | 1 |
| 25 | TGME49_297970.P3426 | M18 aspartyl aminopeptidase | 9 | 9 | 9 | 8 | 8 | 8 | 8 | 1 |
| 25 | TGME49_298020-t26_1-p1 | DEAD-family helicase | 7 | 9 | 10 | 9 | 9 | 9 | 6 | 1 |
| 23 | TGME49_299070-t26_1-p1 | pyruvate kinase PyKII | 10 | 9 | 9 | 8 | 6 | 7 | 3 | 1 |
| 3 | TGME49_299810.P3445 | cysteine-tRNA synthetase (CysRS) | 10 | 9 | 10 | 9 | 9 | 9 | 9 | 1 |
| 3 | TGME49_300100.P3453 | rhoptry neck protein RON2 | 10 | 10 | 10 | 10 | 9 | 9 | 9 | 1 |
| 28 | TGME49_300130.P3456 | apical membrane antigen AMA2 | 9 | 9 | 9 | 8 | 7 | 7 | 7 | 1 |
| 28 | TGME49_300140.P3457 | elongation factor 1-gamma, putative | 9 | 9 | 9 | 8 | 8 | 7 | 7 | 1 |
| 3 | TGME49_300260.P3462 | threonyl-tRNA synthetase family protein | 10 | 10 | 10 | 10 | 9 | 9 | 9 | 1 |
| 6 | TGME49_300310.P3465 | 26S proteasome regulatory subunit, S6a family AAA ATpase | 9 | 9 | 9 | 6 | 9 | 8 | 7 | 1 |
| 28 | TGME49_300350-t26_1-p1 | cysteine desulfurase/selenocysteine lyase family PLP dependent transferase superfamily protein | 10 | 9 | 9 | 9 | 7 | 8 | 7 | 1 |
| 25 | TGME49_301010-t26_1-p1 | serine/threonine protein phosphatase 6 catalytic subunit PP6C | 10 | 9 | 8 | 8 | 8 | 8 | 8 | 1 |
| 28 | TGME49_301120-t26_1-p1 | acetyl-CoA acetyltransferase | 9 | 9 | 8 | 8 | 6 | 7 | 6 | 1 |
| 28 | TGME49_301360.P3475 | phosphate transporter, putative | 10 | 10 | 9 | 9 | 8 | 7 | 6 | 1 |
| 6 | TGME49_301390-t26_1-p1 | ribosome biogenesis protein BOP1, putative | 10 | 10 | 10 | 3 | 9 | 7 | 6 | 1 |
| 3 | TGME49_301440.P3480 | calcium-dependent protein kinase CDPK1 | 9 | 9 | 9 | 9 | 9 | 9 | 9 | 1 |
| 28 | TGME49_301450-t26_1-p1 | FG-GAP repeat-containing protein | 10 | 10 | 9 | 8 | 7 | 7 | 7 | 1 |
| 28 | TGME49_304460-t26_1-p1 | zinc finger, C3HC4 type (RING finger) domain-containing protein | 10 | 10 | 9 | 8 | 8 | 6 | 7 | 1 |
| 23 | TGME49_304650-t26_1-p1 | histidine acid phosphatase superfamily protein | 10 | 10 | 9 | 7 | 8 | 8 | 5 | 1 |
| 28 | TGME49_304680.P3486 | DNA damage-inducible protein 1 | 9 | 9 | 9 | 8 | 8 | 6 | 6 | 1 |
| 3 | TGME49_304710.P3487 | eukaryotic peptide chain release factor, putative | 9 | 9 | 9 | 9 | 9 | 9 | 9 | 1 |
| 6 | TGME49_304970.P3493 | CDK-related protein kinase CRK1 | 10 | 9 | 3 | 4 | 8 | 8 | 8 | 1 |
| 25 | TGME49_304990-t26_1-p1 | guanylate-binding protein, N-terminal domain-containing protein | 10 | 10 | 10 | 8 | 10 | 6 | 7 | 1 |
| 28 | TGME49_305020-t26_1-p1 | CS domain-containing protein | 10 | 9 | 10 | 9 | 8 | 7 | 6 | 1 |
| 25 | TGME49_305040-t26_1-p1 | HEAT repeat-containing protein | 10 | 10 | 10 | 10 | 7 | 8 | 7 | 1 |
| 3 | TGME49_305240-t26_1-p1 | XPA binding protein 2 family protein | 10 | 10 | 10 | 10 | 9 | 9 | 8 | 1 |
| 23 | TGME49_305320-t26_1-p1 | mRNA capping enzyme | 9 | 9 | 9 | 9 | 8 | 7 | 5 | 1 |
| 3 | TGME49_305340.P3502 | ATPase MORC | 10 | 10 | 10 | 8 | 8 | 10 | 9 | 1 |

| | | | | | | | | | | |
|---|---|---|---|---|---|---|---|---|---|---|
| 28 | TGME49_305460-t26_1-p1 | methionine aminopeptidase 2, putative | 9 | 9 | 9 | 8 | 6 | 8 | 8 | 1 |
| 3 | TGME49_305480-t26_1-p1 | elongator complex protein ELP3 | 10 | 10 | 10 | 8 | 9 | 9 | 9 | 1 |
| 25 | TGME49_305590-t26_1-p1 | ABC transporter transmembrane region domain-containing protein | 10 | 10 | 10 | 10 | 8 | 9 | 8 | 1 |
| 25 | TGME49_305640-t26_1-p1 | metallo-beta-lactamase domain-containing protein | 7 | 9 | 9 | 8 | 8 | 7 | 8 | 1 |
| 28 | TGME49_305750-t26_1-p1 | nucleolar gtp-binding protein 2, putative | 9 | 9 | 9 | 7 | 8 | 8 | 7 | 1 |
| 25 | TGME49_305780-t26_1-p1 | 5'-3' exoribonuclease, putative | 10 | 10 | 10 | 8 | 8 | 8 | 9 | 1 |
| 25 | TGME49_305860-t26_1-p1 | calcium-dependent protein kinase CDPK3 | 10 | 9 | 9 | 7 | 8 | 9 | 9 | 1 |
| 28 | TGME49_305940-t26_1-p1 | peptidyl-prolyl cis-trans isomerase, cyclophilin-type domain-containing protein | 10 | 8 | 9 | 9 | 7 | 8 | 7 | 1 |
| 25 | TGME49_305980-t26_1-p1 | pyruvate dehydrogenase complex subunit PDH-E3I | 10 | 9 | 9 | 9 | 9 | 8 | 7 | 1 |
| 6 | TGME49_306020.P3513 | VPS13 domain-containing protein | 10 | 10 | 10 | 6 | 7 | 9 | 7 | 1 |
| 3 | TGME49_306060-t26_1-p1 | rhoptry neck protein RON8 | 10 | 10 | 10 | 10 | 10 | 9 | 8 | 1 |
| 25 | TGME49_306080-t26_1-p1 | ATP-dependent DNA helicase, RecQ family protein | 10 | 9 | 10 | 9 | 9 | 8 | 6 | 1 |
| 6 | TGME49_306220-t26_1-p1 | pre-mRNA-processing factor 40, putative | 10 | 9 | 9 | 6 | 7 | 8 | 7 | 1 |
| 3 | TGME49_306290-t26_1-p1 | DNA-directed RNA polymerase III RPC1 | 10 | 10 | 10 | 10 | 9 | 10 | 10 | 1 |
| 28 | TGME49_306310.P3519 | structural maintenance of chromosomes protein SMC3, putative | 10 | 10 | 10 | 8 | 7 | 6 | 6 | 1 |
| 6 | TGME49_306350.P3520 | variable surface lipoprotein | 10 | 9 | 9 | 5 | 6 | 7 | 7 | 1 |
| 25 | TGME49_306660.P3527 | RNA pseudouridine synthase superfamily protein | 10 | 10 | 10 | 10 | 8 | 9 | 6 | 1 |
| 28 | TGME49_306960-t26_1-p1 | phenylalanine--tRNA ligase, beta subunit protein | 10 | 10 | 9 | 7 | 7 | 7 | 8 | 1 |
| 23 | TGME49_307010.P3534 | histone lysine demethylase JMJC1/KDM5D/JARID1D | 10 | 10 | 9 | 8 | 6 | 8 | 5 | 1 |
| 28 | TGME49_307770.P3538 | pantothenate kinase 1 | 10 | 9 | 8 | 7 | 8 | 7 | 7 | 1 |
| 28 | TGME49_307800-t26_1-p1 | GDA1/CD39 (nucleoside phosphatase) family protein | 10 | 10 | 9 | 9 | 6 | 6 | 7 | 1 |
| 25 | TGME49_307850-t26_1-p1 | 6-phosphogluconate dehydrogenase | 9 | 9 | 9 | 9 | 8 | 8 | 7 | 1 |
| 28 | TGME49_308040-t26_1-p1 | ZPR1 zinc finger domain-containing protein | 9 | 9 | 9 | 9 | 7 | 7 | 7 | 1 |
| 25 | TGME49_308580-t26_1-p1 | Lon protease family protein | 10 | 10 | 10 | 10 | 6 | 9 | 9 | 1 |
| 6 | TGME49_308810-t26_1-p1 | rhoptry neck protein RON9 | 10 | 9 | 9 | 3 | 7 | 8 | 8 | 1 |
| 25 | TGME49_308890.P3547 | transcription elongation factor SPT6 | 10 | 10 | 10 | 10 | 8 | 9 | 8 | 1 |
| 28 | TGME49_308920.P3548 | splicing factor U2AF protein | 10 | 9 | 9 | 9 | 7 | 7 | 7 | 1 |
| 6 | TGME49_309110.P3559 | tRNA-specific 2-thiouridylase MNMA | 10 | 9 | 9 | 5 | 7 | 6 | 6 | 1 |
| 25 | TGME49_309120.P3560 | ribosomal protein RPL4 | 9 | 9 | 9 | 8 | 8 | 8 | 8 | 1 |
| 6 | TGME49_309140-t26_1-p1 | transducin beta-like protein TBL1 | 10 | 10 | 9 | 3 | 6 | 8 | 6 | 1 |
| 6 | TGME49_309190.P3563 | ARO-interacting protein AIP | 10 | 9 | 9 | 6 | 8 | 7 | 6 | 1 |
| 23 | TGME49_309250-t26_1-p1 | hypothetical protein | 10 | 10 | 9 | 8 | 7 | 7 | 5 | 1 |
| 23 | TGME49_309370-t26_1-p1 | AP-3 complex subunit delta, putative | 10 | 10 | 10 | 8 | 6 | 7 | 3 | 1 |
| 3 | TGME49_309420.P3569 | ferlin 1 | 10 | 10 | 10 | 9 | 10 | 10 | 9 | 1 |
| 28 | TGME49_309570-t26_1-p1 | DEAD/DEAH box helicase domain-containing protein | 10 | 9 | 9 | 8 | 7 | 7 | 7 | 1 |
| 6 | TGME49_309580-t26_1-p1 | transporter, major facilitator family protein | 10 | 9 | 9 | 6 | 7 | 7 | 7 | 1 |
| 6 | TGME49_309730-t26_1-p1 | thioredoxin reductase | 10 | 9 | 9 | 5 | 9 | 7 | 7 | 1 |
| 25 | TGME49_309752.P3574 | succinate-Coenzyme A ligase, beta subunit, putative | 9 | 9 | 9 | 9 | 9 | 8 | 6 | 1 |
| 3 | TGME49_309980-t26_1-p1 | dynein heavy chain family protein | 10 | 10 | 10 | 10 | 10 | 10 | 10 | 1 |
| 28 | TGME49_310000.P3588 | membrane protein, putative | 10 | 10 | 9 | 8 | 8 | 7 | 6 | 1 |
| 28 | TGME49_310118.P3595 | ribosomal protein S29, putative | 10 | 9 | 9 | 9 | 8 | 8 | 6 | 1 |

| | | | | | | | | | | |
|---|---|---|---|---|---|---|---|---|---|---|
| 23 | TGME49_310130.P3596 | gamma-tubulin complex component, putative | 10 | 10 | 10 | 9 | 7 | 7 | 4 | 1 |
| 28 | TGME49_310210-t26_1-p1 | hypothetical protein | 10 | 10 | 9 | 9 | 7 | 6 | 6 | 1 |
| 3 | TGME49_310430-t26_1-p1 | Hsp90 domain-containing protein | 10 | 10 | 10 | 9 | 10 | 9 | 9 | 1 |
| 25 | TGME49_310440-t26_1-p1 | membrane occupation and recognition nexus protein MORN1 | 9 | 9 | 9 | 8 | 8 | 8 | 8 | 1 |
| 3 | TGME49_310670.P3619 | glycogen phosphorylase GP | 10 | 10 | 10 | 10 | 10 | 10 | 10 | 1 |
| 25 | TGME49_310700-t26_1-p1 | serine/threonine phosphatase PP1 | 8 | 8 | 8 | 8 | 8 | 8 | 8 | 1 |
| 25 | TGME49_310820-t26_1-p1 | SLU7 splicing factor, putative | 9 | 9 | 9 | 8 | 9 | 8 | 7 | 1 |
| 28 | TGME49_310860-t26_1-p1 | U5 snRNP-specific protein | 9 | 8 | 8 | 7 | 8 | 7 | 7 | 1 |
| 28 | TGME49_311060-t26_1-p1 | metal-dependent phosphohydrolase HD domain-containing protein | 10 | 10 | 9 | 8 | 8 | 7 | 6 | 1 |
| 25 | TGME49_311090-t26_1-p1 | ubiquitin carboxyl-terminal hydrolase | 10 | 9 | 9 | 9 | 8 | 8 | 7 | 1 |
| 25 | TGME49_311240-t26_1-p1 | DnaJ family chaperone J1 | 9 | 9 | 9 | 9 | 8 | 7 | 8 | 1 |
| 28 | TGME49_311300.P3640 | cAMP-dependent protein kinase regulatory subunit 2, putative | 9 | 9 | 8 | 8 | 8 | 7 | 6 | 1 |
| 23 | TGME49_311360.P3643 | protein kinase G AGC kinase family member PKG | 10 | 10 | 10 | 10 | 10 | 10 | 3 | 1 |
| 28 | TGME49_311430.P3648 | fibrillarin | 8 | 8 | 8 | 8 | 8 | 7 | 7 | 1 |
| 25 | TGME49_311500-t26_1-p1 | ThiF family protein | 10 | 10 | 9 | 7 | 7 | 8 | 8 | 1 |
| 3 | TGME49_311720-t26_1-p1 | chaperonin protein BiP | 10 | 10 | 10 | 10 | 10 | 10 | 9 | 1 |
| 28 | TGME49_311800-t26_1-p1 | endonuclease/exonuclease/phosphatase family protein | 10 | 7 | 9 | 9 | 6 | 8 | 8 | 1 |
| 28 | TGME49_311870-t26_1-p1 | WD domain, G-beta repeat-containing protein | 9 | 9 | 9 | 7 | 8 | 7 | 7 | 1 |
| 6 | TGME49_311880-t26_1-p1 | hypothetical protein | 10 | 9 | 9 | 5 | 6 | 8 | 7 | 1 |
| 3 | TGME49_312100.P3662 | inner membrane complex protein IMC42 | 10 | 10 | 10 | 10 | 10 | 9 | 9 | 1 |
| 25 | TGME49_312110.P3663 | apicoplast-associated thioredoxin family protein Atrx1 | 10 | 9 | 9 | 8 | 8 | 8 | 7 | 1 |
| 23 | TGME49_312190-t26_1-p1 | hypothetical protein | 10 | 10 | 10 | 6 | 7 | 8 | 4 | 1 |
| 25 | TGME49_312200.P3665 | serine/threonine protein phosphatase PP5 | 10 | 9 | 9 | 8 | 9 | 8 | 8 | 1 |
| 3 | TGME49_312280-t26_1-p1 | pre-mRNA-splicing factor ATP-dependent RNA helicase, putative | 10 | 10 | 10 | 10 | 10 | 10 | 10 | 1 |
| 28 | TGME49_312310-t26_1-p1 | ATPase, AAA family protein | 10 | 9 | 9 | 7 | 8 | 6 | 6 | 1 |
| 23 | TGME49_312500-t26_1-p1 | hypothetical protein | 10 | 10 | 10 | 9 | 8 | 7 | 5 | 1 |
| 28 | TGME49_312570.P3675 | CMGC kinase, MAPK family (ERK) MAPK-1 | 10 | 10 | 9 | 9 | 7 | 8 | 7 | 1 |
| 25 | TGME49_312622-t26_1-p1 | DUF803 domain-containing protein | 10 | 10 | 9 | 9 | 8 | 8 | 8 | 1 |
| 3 | TGME49_312630.P3676 | glideosome-associated connector GAC | 10 | 10 | 10 | 10 | 10 | 10 | 10 | 1 |
| 6 | TGME49_312860.P3682 | PA domain-containing protein | 10 | 9 | 9 | 6 | 7 | 8 | 8 | 1 |
| 25 | TGME49_312930.P3683 | cystathione gamma lyase | 9 | 9 | 9 | 9 | 8 | 8 | 7 | 1 |
| 25 | TGME49_312990-t26_1-p1 | DEAD/DEAH box ATP-dependent RNA helicase | 10 | 10 | 9 | 9 | 9 | 6 | 9 | 1 |
| 25 | TGME49_313010-t26_1-p1 | DEAD (Asp-Glu-Ala-Asp) box polypeptide DDX6 | 9 | 9 | 9 | 9 | 9 | 8 | 8 | 1 |
| 3 | TGME49_313020.P3687 | STAS domain-containing protein | 10 | 10 | 10 | 9 | 9 | 9 | 9 | 1 |
| 28 | TGME49_313060-t26_1-p1 | eukaryotic translation initiation factor 2B, putative | 10 | 9 | 9 | 7 | 7 | 8 | 7 | 1 |
| 25 | TGME49_313100.P3690 | signal recognition particle SRP54 protein | 10 | 9 | 9 | 9 | 9 | 7 | 9 | 1 |
| 23 | TGME49_313120-t26_1-p1 | DNA-directed RNA polymerase, alpha subunit | 10 | 10 | 9 | 9 | 8 | 6 | 4 | 1 |
| 25 | TGME49_313140.P3691 | isocitrate dehydrogenase 2 | 3 | 9 | 9 | 9 | 9 | 6 | 9 | 1 |
| 25 | TGME49_313180.P3693 | pre-mRNA processing factor 4 kinase | 10 | 9 | 9 | 9 | 9 | 9 | 8 | 1 |
| 6 | TGME49_313230-t26_1-p1 | eukaryotic initiation factor-2, alpha subunit | 9 | 8 | 8 | 6 | 8 | 7 | 7 | 1 |
| 25 | TGME49_313240-t26_1-p1 | ethylene-responsive RNA helicase, putative | 10 | 10 | 9 | 9 | 8 | 8 | 7 | 1 |
| 3 | TGME49_313410.P3704 | proteasome 26S regulatory subunit | 10 | 10 | 10 | 9 | 9 | 9 | 9 | 1 |
| 28 | TGME49_313570-t26_1-p1 | regulator of chromosome condensation (RCC1) repeat-containing protein | 10 | 9 | 9 | 8 | 7 | 8 | 6 | 1 |

| | | | | | | | | | | |
|---|---|---|---|---|---|---|---|---|---|---|
| 28 | TGME49_313590.P3714 | CLEC16A domain-containing protein | 10 | 9 | 8 | 8 | 7 | 7 | 7 | 1 |
| 25 | TGME49_313670.P3718 | AP-1 complex subunit gamma, putative | 10 | 10 | 10 | 10 | 8 | 8 | 8 | 1 |
| 25 | TGME49_313830.P3723 | AARP2CN (NUC121) domain-containing protein | 10 | 10 | 10 | 9 | 8 | 7 | 8 | 1 |
| 28 | TGME49_314055.P3731 | splicing factor, putative | 9 | 8 | 8 | 8 | 8 | 7 | 7 | 1 |
| 23 | TGME49_314100-t26_1-p1 | tetratricopeptide repeat-containing protein | 10 | 9 | 9 | 8 | 8 | 6 | 5 | 1 |
| 23 | TGME49_314240.P3734 | general transcription factor IIIC polypeptide 3 GTF3C3 | 10 | 10 | 9 | 9 | 7 | 7 | 5 | 1 |
| 23 | TGME49_314400-t26_1-p1 | pyruvate dehydrogenase E1 component, beta subunit, putative | 9 | 9 | 9 | 8 | 8 | 8 | 3 | 1 |
| 25 | TGME49_314410-t26_1-p1 | aquarius, putative | 10 | 3 | 10 | 8 | 8 | 10 | 9 | 1 |
| 28 | TGME49_314430.P3743 | serine/threonine specific protein phosphatase | 9 | 9 | 9 | 8 | 8 | 6 | 6 | 1 |
| 28 | TGME49_314480-t26_1-p1 | hypothetical protein | 10 | 10 | 9 | 9 | 7 | 7 | 6 | 1 |
| 28 | TGME49_314500-t26_1-p1 | subtilisin SUB2 | 10 | 10 | 9 | 8 | 8 | 7 | 7 | 1 |
| 25 | TGME49_314740.P3747 | PSP protein | 10 | 9 | 9 | 9 | 8 | 8 | 7 | 1 |
| 3 | TGME49_314780-t26_1-p1 | myosin G | 10 | 10 | 10 | 10 | 10 | 10 | 9 | 1 |
| 28 | TGME49_314810.P3749 | ribosomal protein RPL7 | 8 | 8 | 8 | 8 | 7 | 7 | 7 | 1 |
| 25 | TGME49_314860-t26_1-p1 | zinc knuckle domain-containing protein | 10 | 9 | 9 | 8 | 8 | 8 | 7 | 1 |
| 25 | TGME49_314970-t26_1-p1 | root hair defective 3 gtp-binding protein (rhd3) protein | 10 | 10 | 10 | 7 | 8 | 9 | 9 | 1 |
| 6 | TGME49_315170.P3758 | O-phosphoseryl-tRNA(Sec) selenium transferase | 9 | 9 | 9 | 5 | 7 | 6 | 7 | 1 |
| 25 | TGME49_315220-t26_1-p1 | rhoptry protein ROP14 | 10 | 10 | 10 | 10 | 6 | 8 | 9 | 1 |
| 6 | TGME49_315580-t26_1-p1 | hypothetical protein | 10 | 10 | 9 | 6 | 8 | 8 | 8 | 1 |
| 28 | TGME49_315590-t26_1-p1 | macro domain-containing protein | 10 | 9 | 9 | 7 | 6 | 7 | 8 | 1 |
| 6 | TGME49_315620.P3782 | vacuolar ATP synthase subunit C, putative | 9 | 9 | 9 | 6 | 8 | 8 | 8 | 1 |
| 25 | TGME49_315860.P3790 | EF hand domain-containing protein | 10 | 10 | 10 | 8 | 9 | 9 | 7 | 1 |
| 28 | TGME49_316100-t26_1-p1 | queuine trna-ribosyltransferase domain-containing protein | 10 | 9 | 8 | 7 | 7 | 7 | 6 | 1 |
| 28 | TGME49_316280.P3804 | transporter facilitator protein TFP3 | 9 | 9 | 9 | 9 | 8 | 6 | 7 | 1 |
| 3 | TGME49_316400-t26_1-p1 | alpha tubulin TUBA1 | 9 | 9 | 9 | 9 | 9 | 9 | 9 | 1 |
| 3 | TGME49_316520.P3810 | 1,4-alpha-glucan-branching enzyme | 10 | 10 | 10 | 8 | 10 | 10 | 10 | 1 |
| 28 | TGME49_316660-t26_1-p1 | cullin family protein | 10 | 10 | 9 | 7 | 8 | 6 | 6 | 1 |
| 3 | TGME49_316730-t26_1-p1 | non-discharge partner NDP2 | 10 | 10 | 10 | 7 | 10 | 10 | 10 | 1 |
| 28 | TGME49_316900-t26_1-p1 | Sas10 C-terminal domain-containing protein | 10 | 9 | 9 | 7 | 8 | 7 | 7 | 1 |
| 25 | TGME49_318150.P3821 | apicomplexan amino acid transporter ApiAT3-1 | 10 | 9 | 9 | 9 | 8 | 7 | 8 | 1 |
| 25 | TGME49_318230-t26_1-p1 | phosphoglycerate kinase PGKI | 9 | 9 | 9 | 9 | 9 | 8 | 6 | 1 |
| 23 | TGME49_318260-t26_1-p1 | transcription initiation factor TFIID subunit TAF5 | 10 | 10 | 9 | 8 | 7 | 7 | 5 | 1 |
| 25 | TGME49_318330-t26_1-p1 | histone lysine acetyltransferase MYST-A | 9 | 9 | 9 | 8 | 8 | 8 | 8 | 1 |
| 25 | TGME49_318410-t26_1-p1 | TCP-1 chaperonin, putative | 9 | 9 | 9 | 7 | 9 | 9 | 9 | 1 |
| 28 | TGME49_318430.P3830 | malate dehydrogenase MDH | 8 | 8 | 8 | 8 | 8 | 7 | 7 | 1 |
| 25 | TGME49_318440.P3831 | helicase associated domain (ha2) protein | 10 | 10 | 10 | 10 | 8 | 9 | 7 | 1 |
| 23 | TGME49_318460-t26_1-p1 | P-type ATPase of unknown pump specificity (type V) protein | 10 | 10 | 10 | 10 | 8 | 8 | 4 | 1 |
| 25 | TGME49_318480-t26_1-p1 | SWI2/SNF2-containing protein RAD5 | 10 | 10 | 10 | 9 | 8 | 8 | 7 | 1 |
| 25 | TGME49_318560-t26_1-p1 | ATPase/histidine kinase/DNA gyrase B/HSP90 domain-containing protein | 9 | 9 | 9 | 9 | 9 | 8 | 8 | 1 |
| 23 | TGME49_318580.P3837 | phosphoglucomutase PGM2 | 10 | 10 | 9 | 9 | 7 | 8 | 3 | 1 |
| 3 | TGME49_318620-t26_1-p1 | RNA-directed RNA polymerase | 10 | 10 | 10 | 10 | 8 | 9 | 9 | 1 |
| 6 | TGME49_318650.P3840 | transhydrogenase | 10 | 7 | 10 | 3 | 6 | 10 | 8 | 1 |
| 6 | TGME49_318700.P3843 | eukaryotic initiation factor-6, putative | 8 | 8 | 8 | 3 | 7 | 7 | 7 | 1 |
| 3 | TGME49_318710-t26_1-p1 | ATP-binding cassette sub-family F member 1 | 10 | 10 | 10 | 10 | 8 | 9 | 9 | 1 |

| | | | | | | | | | | |
|---|---|---|---|---|---|---|---|---|---|---|
| 6 | TGME49_319500.P3849 | TMEM121 domain-containing protein | 10 | 10 | 10 | 5 | 8 | 8 | 8 | 1 |
| 25 | TGME49_319640-t26_1-p1 | hypothetical protein | 10 | 10 | 10 | 10 | 9 | 8 | 8 | 1 |
| 28 | TGME49_319850-t26_1-p1 | splicing factor U2AF family SnRNP auxilary factor large subunit, RRM domain-containing protein | 9 | 8 | 8 | 8 | 8 | 7 | 7 | 1 |
| 3 | TGME49_319860-t26_1-p1 | DNA polymerase family B protein | 10 | 10 | 10 | 9 | 9 | 10 | 10 | 1 |
| 25 | TGME49_319920.P3862 | 2-oxo acid dehydrogenases acyltransferase (catalytic domain) domain-containing protein | 9 | 8 | 9 | 8 | 8 | 8 | 8 | 1 |
| 25 | TGME49_320020.P3868 | apicomplexan amino acid transporter ApiAT2 | 10 | 9 | 9 | 9 | 8 | 8 | 7 | 1 |
| 28 | TGME49_320050.P3869 | ribosomal protein RPL5 | 8 | 8 | 8 | 8 | 8 | 7 | 7 | 1 |
| 28 | TGME49_320210-t26_1-p1 | pre-mRNA-processing factor 19, putative | 9 | 9 | 9 | 9 | 7 | 6 | 7 | 1 |
| 25 | TGME49_320300.P3875 | SWI2/SNF2 Brahma-like putative | 10 | 10 | 10 | 7 | 9 | 9 | 8 | 1 |
| 6 | TGME49_320430-t26_1-p1 | cell-cycle-control protein (translation regulation), putative | 10 | 10 | 9 | 6 | 9 | 9 | 8 | 1 |
| 28 | TGME49_320490.P3878 | dense granule protein GRA66 | 10 | 10 | 9 | 9 | 7 | 6 | 6 | 1 |
| 23 | TGME49_320500.P3879 | patched family protein | 10 | 10 | 10 | 9 | 8 | 8 | 3 | 1 |
| 25 | TGME49_320570-t26_1-p1 | elongation factor Tu, putative | 10 | 9 | 9 | 9 | 9 | 6 | 8 | 1 |
| 3 | TGME49_320640-t26_1-p1 | peptidylprolyl isomerase domain-containing protein | 10 | 10 | 10 | 7 | 9 | 9 | 9 | 1 |
| 6 | TGME49_320670-t26_1-p1 | vacuolar protein sorting 16, putative | 10 | 10 | 10 | 6 | 9 | 9 | 7 | 1 |
| 28 | TGME49_320730.P3888 | homoserine O-acetyltransferase, putative | 9 | 9 | 8 | 8 | 7 | 7 | 7 | 1 |
| 3 | TGME49_321440-t26_1-p1 | SWI2/SNF2 ISWI-like SANT | 10 | 10 | 10 | 10 | 10 | 10 | 8 | 1 |
| 28 | TGME49_321500.P3895 | RNA recognition motif-containing protein | 9 | 9 | 9 | 8 | 8 | 7 | 6 | 1 |
| 28 | TGME49_321530-t26_1-p1 | cathepsin CPL | 9 | 9 | 8 | 8 | 8 | 7 | 7 | 1 |
| 23 | TGME49_321620-t26_1-p1 | dynamin-related protein DRPB | 10 | 10 | 10 | 7 | 10 | 9 | 3 | 1 |
| 3 | TGME49_321640.P3898 | cell division protein CDC48AP | 10 | 10 | 10 | 7 | 10 | 9 | 9 | 1 |
| 25 | TGME49_321650.P3899 | rhoptry neck protein RON13 | 10 | 10 | 10 | 8 | 8 | 8 | 7 | 1 |
| 23 | TGME49_500025.P28 | GCC2 and GCC3 domain-containing protein | 10 | 10 | 10 | 9 | 10 | 10 | 4 | 1 |
| 3 | TGME49_500062.P363 | dynein heavy chain, putative | 10 | 10 | 10 | 8 | 10 | 10 | 10 | 1 |
| 3 | TGME49_500072.P373 | dynein heavy chain, putative | 10 | 10 | 10 | 10 | 10 | 10 | 10 | 1 |
| 25 | TGME49_500080.P381 | kinesin-like protein KIN14, putative | 10 | 10 | 10 | 10 | 10 | 9 | 7 | 1 |
| 3 | TGME49_500101.P402 | fatty acyl-CoA reductase, putative | 10 | 9 | 10 | 10 | 9 | 10 | 9 | 1 |
| 28 | TGME49_500202.P207 | DEAD/DEAH box helicase, putative | 10 | 10 | 6 | 8 | 8 | 7 | 8 | 1 |
| 3 | TGME49_500222.P227 | GCC2 and GCC3 domain-containing protein | 10 | 10 | 10 | 9 | 10 | 10 | 7 | 1 |
| 28 | TGME49_500226.P232 | patched domain-containing protein, putative | 10 | 9 | 10 | 9 | 7 | 7 | 6 | 1 |
| 6 | TGME49_500270.P59 | coiled-coil domain-containing protein 39, putative | 10 | 9 | 9 | 6 | 6 | 7 | 7 | 1 |
| 28 | TGME49_500287.P76 | phosphatidylinositol 4-kinase, putative | 10 | 10 | 10 | 9 | 8 | 7 | 6 | 1 |
| 6 | TGME49_500302.P92 | Porphobilinogen Synthase (PBGS) / Aminolevulinic Acid Dehydratase (ALAD) + Pitrilysin (apicoplast stromal processing peptidase) | 9 | 9 | 9 | 3 | 8 | 8 | 8 | 1 |
| 28 | TGME49_500314.P104 | protein phosphatase PPM11C | 10 | 10 | 8 | 7 | 7 | 7 | 7 | 1 |
| 25 | TGME49_500340.P130 | 3'5'-cyclic nucleotide phosphodiesterase, putative | 10 | 9 | 10 | 9 | 8 | 6 | 9 | 1 |
| 23 | TGME49_500366.P265 | hypothetical protein, conserved | 10 | 10 | 10 | 10 | 9 | 7 | 5 | 1 |
| 23 | TGME49_500380.P279 | cilia- and flagella-associated protein 46, putative | 10 | 10 | 10 | 10 | 6 | 7 | 5 | 1 |
| 3 | TGME49_500391.P290 | copper amine oxidase, putative | 10 | 10 | 10 | 10 | 10 | 10 | 10 | 1 |

| Kmean_cluster | GeneID (all protein TG) | Product Description | Decile_bitHH | Decile_bitNcan | Decile_bitBB | Decilebit_CS | Decilebit_SN3 | Decile bit_Ef | Decile bit_CC | Meta_cluster |
|---|---|---|---|---|---|---|---|---|---|---|
| 18 | TGME49_200280-t26_1-p1 | WD domain, G-beta repeat-containing protein | 10 | 10 | 9 | 8 | 6 | 5 | 4 | 2 |
| 18 | TGME49_201200.P466 | cleavage and polyadenylation specificity factor subunit 4 | 10 | 9 | 9 | 9 | 6 | 7 | 5 | 2 |
| 18 | TGME49_201380-t26_1-p1 | chorismate synthase, putative | 9 | 9 | 8 | 8 | 5 | 4 | 5 | 2 |
| 29 | TGME49_201400.P469 | Sin3-associated polypeptide SAP18 | 10 | 8 | 8 | 7 | 7 | 4 | 4 | 2 |
| 29 | TGME49_201840-t26_1-p1 | aspartyl protease ASP1 | 10 | 9 | 8 | 8 | 7 | 3 | 3 | 2 |
| 29 | TGME49_202020.P479 | DnAK-TPR | 10 | 10 | 9 | 7 | 8 | 3 | 3 | 2 |
| 18 | TGME49_202040-t26_1-p1 | hypothetical protein | 10 | 10 | 9 | 6 | 7 | 4 | 5 | 2 |
| 18 | TGME49_202180-t26_1-p1 | hypothetical protein | 10 | 10 | 7 | 8 | 6 | 5 | 5 | 2 |
| 18 | TGME49_202350-t26_1-p1 | 50S ribosomal protein L21, putative | 10 | 8 | 8 | 7 | 5 | 5 | 5 | 2 |
| 18 | TGME49_202400-t26_1-p1 | calcium binding egf domain-containing protein | 10 | 8 | 10 | 6 | 7 | 6 | 3 | 2 |
| 18 | TGME49_202410-t26_1-p1 | CRAL/TRIO domain-containing protein | 10 | 9 | 8 | 7 | 6 | 6 | 5 | 2 |
| 29 | TGME49_202580.P495 | chaperone protein ClpB5, putative | 10 | 10 | 10 | 10 | 8 | 3 | 3 | 2 |
| 18 | TGME49_202720-t26_1-p1 | RNB family domain-containing protein | 10 | 9 | 9 | 7 | 5 | 5 | 4 | 2 |
| 2 | TGME49_202950.P507 | hypothetical protein | 10 | 10 | 10 | 9 | 8 | 3 | 7 | 2 |
| 2 | TGME49_203010-t26_1-p1 | aurora-related kinase ARK3 | 10 | 10 | 8 | 8 | 8 | 3 | 7 | 2 |
| 29 | TGME49_203240-t26_1-p1 | hypothetical protein | 10 | 10 | 9 | 7 | 8 | 4 | 4 | 2 |
| 18 | TGME49_203300-t26_1-p1 | DUF726 domain-containing protein | 10 | 10 | 7 | 7 | 7 | 5 | 6 | 2 |
| 29 | TGME49_203370-t26_1-p1 | Utp12 domain-containing protein | 10 | 9 | 9 | 7 | 7 | 4 | 4 | 2 |
| 18 | TGME49_203510-t26_1-p1 | zinc finger, C3HC4 type (RING finger) domain-containing protein | 10 | 9 | 9 | 8 | 5 | 5 | 4 | 2 |
| 18 | TGME49_203690.P530 | AP2 domain transcription factor AP2VIIa-5 | 10 | 10 | 8 | 7 | 6 | 6 | 3 | 2 |
| 29 | TGME49_203840.P535 | DEAD/DEAH box helicase domain-containing protein | 10 | 10 | 10 | 8 | 7 | 4 | 4 | 2 |
| 18 | TGME49_203950-t26_1-p1 | Myb family DNA-binding domain-containing protein | 10 | 10 | 10 | 7 | 6 | 5 | 4 | 2 |
| 29 | TGME49_204050.P540 | subtilisin SUB1 | 10 | 9 | 7 | 8 | 6 | 3 | 3 | 2 |
| 18 | TGME49_204100-t26_1-p1 | eIF2 kinase IF2K-C | 10 | 10 | 9 | 8 | 5 | 5 | 4 | 2 |
| 18 | TGME49_204410-t26_1-p1 | endonuclease/exonuclease/phosphatase family protein | 10 | 9 | 8 | 7 | 6 | 4 | 5 | 2 |
| 29 | TGME49_204550-t26_1-p1 | hypothetical protein | 10 | 10 | 9 | 7 | 7 | 4 | 3 | 2 |
| 18 | TGME49_205500-t26_1-p1 | U3 small nucleolar RNA-associated protein 14, putative | 10 | 9 | 9 | 7 | 5 | 5 | 4 | 2 |
| 18 | TGME49_205540-t26_1-p1 | DEAD/DEAH box helicase domain-containing protein | 10 | 10 | 9 | 7 | 6 | 5 | 5 | 2 |
| 18 | TGME49_205570-t26_1-p1 | sorting assembly machinery subunit SAM50 | 10 | 9 | 9 | 8 | 6 | 5 | 4 | 2 |
| 18 | TGME49_205580.P574 | nuclear factor NF4 | 9 | 9 | 8 | 7 | 5 | 5 | 5 | 2 |
| 2 | TGME49_206400.P583 | FAD binding domain of DNA photolyase domain-containing protein | 10 | 9 | 9 | 9 | 6 | 4 | 8 | 2 |
| 18 | TGME49_206640-t26_1-p1 | hypothetical protein | 10 | 10 | 10 | 8 | 5 | 4 | 6 | 2 |
| 18 | TGME49_206670-t26_1-p1 | RNA recognition motif-containing protein | 10 | 10 | 8 | 6 | 6 | 5 | 5 | 2 |
| 18 | TGME49_207120.P605 | SUN-like protein SLP2 | 10 | 10 | 7 | 7 | 6 | 6 | 4 | 2 |
| 2 | TGME49_207620-t26_1-p1 | pyridine nucleotide-disulfide oxidoreductase domain-containing protein | 10 | 9 | 9 | 8 | 5 | 3 | 7 | 2 |
| 2 | TGME49_207730-t26_1-p1 | hypothetical protein | 10 | 10 | 9 | 8 | 5 | 3 | 6 | 2 |
| 2 | TGME49_207800-t26_1-p1 | hypothetical protein | 9 | 9 | 8 | 6 | 7 | 4 | 6 | 2 |
| 18 | TGME49_207960.P633 | vacuolar fusion protein CCZ1, putative | 10 | 9 | 9 | 6 | 7 | 6 | 6 | 2 |
| 29 | TGME49_207965-t26_1-p1 | FAD-dependent oxidoreductase, putative | 10 | 9 | 9 | 8 | 7 | 4 | 4 | 2 |
| 18 | TGME49_208320-t26_1-p1 | polynucleotide kinase 3 phosphatase | 10 | 9 | 9 | 8 | 6 | 5 | 3 | 2 |

| | | | | | | | | | | |
|---|---|---|---|---|---|---|---|---|---|---|
| 29 | TGME49_208740.P652 | microneme adhesive repeat protein MCP3 | 9 | 9 | 8 | 6 | 8 | 3 | 3 | 2 |
| 29 | TGME49_208842.P659 | DEAD/DEAH box helicase domain-containing protein | 10 | 10 | 9 | 7 | 7 | 3 | 5 | 2 |
| 18 | TGME49_209150-t26_1-p1 | non-proton pumping type-II NADH dehydrogenase I | 10 | 9 | 9 | 8 | 6 | 6 | 6 | 2 |
| 18 | TGME49_209190-t26_1-p1 | ABC transporter transmembrane region domain-containing protein | 10 | 9 | 9 | 7 | 8 | 6 | 5 | 2 |
| 29 | TGME49_209470.P680 | hypothetical protein | 9 | 9 | 8 | 8 | 8 | 3 | 3 | 2 |
| 18 | TGME49_209500-t26_1-p1 | hypothetical protein | 10 | 10 | 10 | 8 | 6 | 5 | 5 | 2 |
| 29 | TGME49_209550-t26_1-p1 | hypothetical protein | 10 | 10 | 9 | 9 | 7 | 3 | 5 | 2 |
| 18 | TGME49_209770.P692 | helicase, putative | 10 | 10 | 10 | 5 | 7 | 6 | 5 | 2 |
| 2 | TGME49_209800.P693 | formate/nitrite transporter FNT1 | 9 | 9 | 8 | 8 | 7 | 3 | 6 | 2 |
| 18 | TGME49_209970-t26_1-p1 | Spc97 / Spc98 family protein | 10 | 10 | 9 | 6 | 6 | 5 | 4 | 2 |
| 18 | TGME49_210230.P705 | pseudouridine synthase, putative | 10 | 9 | 8 | 8 | 5 | 5 | 6 | 2 |
| 29 | TGME49_210280-t26_1-p1 | aurora-related kinase ARK1 | 9 | 9 | 9 | 8 | 8 | 4 | 3 | 2 |
| 18 | TGME49_210430-t26_1-p1 | DnaJ domain-containing protein | 10 | 9 | 8 | 6 | 5 | 6 | 5 | 2 |
| 18 | TGME49_210440-t26_1-p1 | polynucleotide adenylyltransferase | 10 | 9 | 8 | 8 | 6 | 6 | 3 | 2 |
| 2 | TGME49_210700.P713 | VPS13 domain-containing protein | 10 | 10 | 10 | 10 | 5 | 3 | 6 | 2 |
| 18 | TGME49_210781-t26_1-p1 | ubiquitin carboxyl-terminal hydrolase | 10 | 10 | 7 | 6 | 6 | 6 | 6 | 2 |
| 18 | TGME49_210830.P721 | RIO1 family protein | 10 | 9 | 8 | 7 | 6 | 6 | 4 | 2 |
| 29 | TGME49_210950-t26_1-p1 | oocyst wall protein | 10 | 10 | 10 | 9 | 9 | 3 | 3 | 2 |
| 29 | TGME49_211050.P724 | ATPase, AAA family protein | 10 | 9 | 9 | 9 | 7 | 4 | 4 | 2 |
| 2 | TGME49_211350.P733 | CBS domain-containing protein | 10 | 10 | 9 | 8 | 8 | 3 | 6 | 2 |
| 18 | TGME49_212075-t26_1-p1 | meiosis-specific nuclear structural protein 1, putative | 9 | 9 | 8 | 8 | 6 | 4 | 5 | 2 |
| 18 | TGME49_212100.P753 | ubiquitin activating enzyme UBA1 | 10 | 9 | 8 | 8 | 6 | 5 | 4 | 2 |
| 29 | TGME49_212150.P754 | conserved oligomeric Golgi complex subunit 4 | 10 | 10 | 9 | 9 | 7 | 5 | 4 | 2 |
| 2 | TGME49_213000.P776 | replication factor C, subunit 5, putative | 9 | 8 | 8 | 8 | 5 | 4 | 7 | 2 |
| 29 | TGME49_213340-t26_1-p1 | GMC oxidoreductase | 10 | 10 | 10 | 10 | 9 | 3 | 3 | 2 |
| 18 | TGME49_213650-t26_1-p1 | ATP-binding domain 1 family protein | 9 | 8 | 8 | 8 | 6 | 6 | 5 | 2 |
| 29 | TGME49_213730-t26_1-p1 | lanthionine synthetase C family protein | 9 | 9 | 9 | 8 | 7 | 4 | 4 | 2 |
| 18 | TGME49_213890.P812 | Myb family DNA-binding domain-containing protein | 10 | 9 | 9 | 8 | 7 | 6 | 5 | 2 |
| 18 | TGME49_214140.P816 | hypothetical protein | 10 | 10 | 9 | 6 | 6 | 6 | 5 | 2 |
| 18 | TGME49_214470.P825 | ubiquitin-like protease ULP1 | 10 | 8 | 8 | 7 | 7 | 6 | 5 | 2 |
| 29 | TGME49_214570-t26_1-p1 | hypothetical protein | 10 | 9 | 9 | 9 | 8 | 3 | 3 | 2 |
| 18 | TGME49_214610-t26_1-p1 | trafficking protein particle complex subunit TRS85 | 10 | 10 | 8 | 7 | 7 | 6 | 5 | 2 |
| 18 | TGME49_214830-t26_1-p1 | exportin-T, putative | 10 | 10 | 10 | 9 | 6 | 6 | 4 | 2 |
| 29 | TGME49_214840.P833 | AP2 domain transcription factor AP2X-7 | 10 | 10 | 10 | 7 | 7 | 5 | 3 | 2 |
| 18 | TGME49_214990.P842 | hypothetical protein | 10 | 10 | 9 | 7 | 4 | 6 | 4 | 2 |
| 2 | TGME49_215060-t26_1-p1 | small GTP-binding protein sar1, putative | 7 | 7 | 7 | 7 | 7 | 3 | 7 | 2 |
| 2 | TGME49_215230-t26_1-p1 | RNA recognition motif-containing protein | 10 | 9 | 9 | 8 | 7 | 4 | 6 | 2 |
| 2 | TGME49_215290-t26_1-p1 | saccharopine dehydrogenase domain-containing protein | 9 | 9 | 8 | 8 | 7 | 5 | 7 | 2 |
| 18 | TGME49_215370-t26_1-p1 | hypothetical protein | 10 | 9 | 9 | 8 | 6 | 6 | 5 | 2 |
| 29 | TGME49_215480-t26_1-p1 | Adenosine/AMP deaminase domain-containing protein | 9 | 9 | 8 | 8 | 6 | 3 | 3 | 2 |
| 18 | TGME49_215560.P862 | histone arginine methyltransferase PRMT5 | 10 | 10 | 9 | 8 | 5 | 6 | 5 | 2 |

| | | | | | | | | | | |
|---|---|---|---|---|---|---|---|---|---|---|
| 18 | TGME49_215640-t26_1-p1 | zinc finger, C3HC4 type (RING finger) domain-containing protein | 10 | 10 | 8 | 7 | 6 | 5 | 3 | 2 |
| 29 | TGME49_215690-t26_1-p1 | hypothetical protein | 10 | 10 | 10 | 6 | 8 | 4 | 3 | 2 |
| 18 | TGME49_216250-t26_1-p1 | RecF/RecN/SMC N terminal domain-containing protein | 10 | 10 | 10 | 6 | 6 | 5 | 4 | 2 |
| 2 | TGME49_216260-t26_1-p1 | eukaryotic initiation factor-2B, gamma subunit, putative | 9 | 9 | 9 | 8 | 6 | 5 | 7 | 2 |
| 29 | TGME49_216280.P880 | adenylate cyclase alpha 2 | 10 | 10 | 10 | 10 | 9 | 3 | 3 | 2 |
| 18 | TGME49_216480-t26_1-p1 | NMD3 family protein | 10 | 9 | 9 | 9 | 7 | 6 | 5 | 2 |
| 18 | TGME49_216510-t26_1-p1 | thioredoxin, putative | 10 | 9 | 8 | 7 | 5 | 6 | 5 | 2 |
| 29 | TGME49_216710.P892 | apicomplexan amino acid transporter ApiAT5-4 | 10 | 8 | 9 | 8 | 7 | 4 | 5 | 2 |
| 18 | TGME49_216780.P893 | hypothetical protein | 10 | 10 | 9 | 7 | 6 | 6 | 4 | 2 |
| 29 | TGME49_216810-t26_1-p1 | 5'-nucleotidase, C-terminal domain-containing protein | 10 | 9 | 9 | 7 | 8 | 3 | 3 | 2 |
| 29 | TGME49_217220.P899 | zinc finger, C3HC4 type (RING finger) domain-containing protein | 10 | 10 | 10 | 8 | 7 | 5 | 3 | 2 |
| 29 | TGME49_217480-t26_1-p1 | hypothetical protein | 10 | 10 | 10 | 6 | 7 | 3 | 4 | 2 |
| 29 | TGME49_217610.P912 | hypothetical protein | 10 | 10 | 10 | 9 | 7 | 5 | 3 | 2 |
| 29 | TGME49_217650.P915 | GAS8 domain-containing protein | 9 | 9 | 9 | 7 | 8 | 5 | 4 | 2 |
| 2 | TGME49_217740.P923 | 3-ketoacyl-(acyl-carrier-protein) reductase | 8 | 8 | 8 | 7 | 7 | 3 | 7 | 2 |
| 18 | TGME49_217830-t26_1-p1 | hypothetical protein | 10 | 8 | 9 | 8 | 6 | 5 | 5 | 2 |
| 2 | TGME49_218470.P940 | protein disulfide-isomerase, putative | 10 | 10 | 9 | 9 | 6 | 3 | 6 | 2 |
| 29 | TGME49_218560-t26_1-p1 | acetyl-coA carboxylase ACC2 | 10 | 10 | 10 | 10 | 9 | 3 | 3 | 2 |
| 18 | TGME49_218570-t26_1-p1 | Nin one binding (NOB1) Zn-ribbon family protein | 9 | 9 | 8 | 8 | 6 | 6 | 4 | 2 |
| 2 | TGME49_218880.P952 | striated fiber assemblin SFA3 | 8 | 8 | 8 | 7 | 6 | 3 | 7 | 2 |
| 29 | TGME49_218890.P953 | hypothetical protein | 10 | 10 | 9 | 9 | 8 | 3 | 3 | 2 |
| 2 | TGME49_218920-t26_1-p1 | proteasome subunit beta type-5, putative | 8 | 8 | 8 | 8 | 7 | 3 | 6 | 2 |
| 18 | TGME49_218960.P958 | AP2 domain transcription factor AP2XII-1 | 10 | 8 | 9 | 8 | 5 | 5 | 4 | 2 |
| 18 | TGME49_219250-t26_1-p1 | acetyltransferase, GNAT family protein | 10 | 9 | 8 | 8 | 6 | 5 | 5 | 2 |
| 18 | TGME49_219630-t26_1-p1 | flavodoxin domain-containing protein | 10 | 9 | 9 | 9 | 5 | 6 | 4 | 2 |
| 18 | TGME49_219640-t26_1-p1 | RING zinc finger protein | 10 | 10 | 8 | 8 | 7 | 6 | 4 | 2 |
| 18 | TGME49_220290.P994 | hypothetical protein | 10 | 9 | 9 | 8 | 6 | 5 | 5 | 2 |
| 18 | TGME49_220910-t26_1-p1 | HEAT repeat-containing protein | 10 | 9 | 8 | 7 | 6 | 5 | 5 | 2 |
| 18 | TGME49_221220.P1013 | hypothetical protein | 10 | 8 | 9 | 7 | 5 | 7 | 5 | 2 |
| 18 | TGME49_221260-t26_1-p1 | Class-II DAHP synthetase family protein | 9 | 9 | 8 | 6 | 7 | 4 | 5 | 2 |
| 18 | TGME49_221410.P1021 | actin-like protein ALP4 | 9 | 9 | 8 | 6 | 6 | 5 | 5 | 2 |
| 18 | TGME49_221600-t26_1-p1 | ribosomal silencing factor RsfS, putative | 10 | 9 | 8 | 7 | 6 | 5 | 4 | 2 |
| 2 | TGME49_221620-t26_1-p1 | beta-tubulin, putative | 9 | 9 | 9 | 9 | 9 | 3 | 9 | 2 |
| 29 | TGME49_221720.P1036 | hypothetical protein | 10 | 10 | 9 | 7 | 6 | 4 | 3 | 2 |
| 29 | TGME49_221830.P1038 | subtilisin SUB12 | 10 | 10 | 10 | 10 | 6 | 3 | 3 | 2 |
| 29 | TGME49_221940.P1043 | AP-2 complex subunit alpha | 10 | 10 | 9 | 5 | 8 | 4 | 4 | 2 |
| 18 | TGME49_222030.P1046 | hypothetical protein | 10 | 9 | 9 | 6 | 5 | 5 | 6 | 2 |
| 29 | TGME49_222070-t26_1-p1 | elongation factor Tu GTP binding domain-containing protein | 10 | 9 | 9 | 8 | 6 | 3 | 3 | 2 |
| 29 | TGME49_222200.P1053 | apical annuli protein LMBD3 | 10 | 9 | 9 | 7 | 8 | 3 | 5 | 2 |
| 18 | TGME49_222230-t26_1-p1 | MMS19 nucleotide excision repair protein, putative | 10 | 9 | 8 | 7 | 6 | 4 | 4 | 2 |
| 18 | TGME49_222410.P1059 | lipase maturation factor, putative | 10 | 9 | 9 | 7 | 6 | 6 | 5 | 2 |
| 2 | TGME49_222710-t26_1-p1 | IMP-specific 5'-nucleotidase 1, putative | 10 | 9 | 9 | 8 | 7 | 3 | 7 | 2 |

| | | | | | | | | | | |
|---|---|---|---|---|---|---|---|---|---|---|
| 29 | TGME49_222940-t26_1-p1 | hypothetical protein | 9 | 9 | 9 | 8 | 8 | 3 | 3 | 2 |
| 2 | TGME49_223660-t26_1-p1 | 50S ribosomal protein L4, putative | 10 | 9 | 9 | 9 | 9 | 4 | 7 | 2 |
| 2 | TGME49_223700-t26_1-p1 | F5/8 type C domain-containing protein | 10 | 10 | 10 | 9 | 8 | 3 | 8 | 2 |
| 2 | TGME49_223780-t26_1-p1 | armadillo repeat-containing protein | 10 | 10 | 10 | 10 | 9 | 5 | 8 | 2 |
| 2 | TGME49_223960-t26_1-p1 | ubiquitin interaction motif family protein | 9 | 8 | 8 | 8 | 7 | 4 | 6 | 2 |
| 18 | TGME49_224050-t26_1-p1 | AP2 domain transcription factor AP2X-4 | 10 | 9 | 9 | 6 | 5 | 6 | 6 | 2 |
| 2 | TGME49_224205-t26_1-p1 | ER lumen protein retaining receptor, putative | 9 | 8 | 8 | 6 | 8 | 3 | 7 | 2 |
| 29 | TGME49_224220-t26_1-p1 | serine/threonine-protein phosphatase PP2A catalytic subunit | 9 | 9 | 8 | 6 | 8 | 5 | 3 | 2 |
| 18 | TGME49_224260.P1111 | PHD-finger domain-containing protein | 10 | 10 | 10 | 7 | 7 | 6 | 5 | 2 |
| 18 | TGME49_224510-t26_1-p1 | hypothetical protein | 9 | 10 | 10 | 7 | 5 | 6 | 6 | 2 |
| 2 | TGME49_224530.P1121 | alveolin domain containing intermediate filament IMC5 | 9 | 8 | 7 | 7 | 6 | 3 | 6 | 2 |
| 18 | TGME49_224540-t26_1-p1 | hypothetical protein | 10 | 10 | 9 | 8 | 6 | 5 | 5 | 2 |
| 18 | TGME49_224590-t26_1-p1 | DNA-directed RNA polymerase III POLR3C | 10 | 10 | 9 | 9 | 6 | 7 | 5 | 2 |
| 18 | TGME49_224650.P1123 | mRNA capping enzyme, beta chain protein | 10 | 9 | 9 | 8 | 6 | 5 | 5 | 2 |
| 18 | TGME49_224700.P1125 | intraconoidal microtubule-associated protein ICMAP2 | 10 | 10 | 9 | 8 | 7 | 5 | 5 | 2 |
| 18 | TGME49_224870.P1134 | hypothetical protein | 10 | 10 | 9 | 8 | 5 | 5 | 4 | 2 |
| 29 | TGME49_224890-t26_1-p1 | hypothetical protein | 10 | 10 | 9 | 8 | 7 | 4 | 4 | 2 |
| 18 | TGME49_224920-t26_1-p1 | GPI ethanolamine phosphate transferase 3, putative | 10 | 10 | 6 | 7 | 5 | 6 | 6 | 2 |
| 29 | TGME49_225028.P1143 | myosin heavy chain, putative | 10 | 10 | 10 | 9 | 8 | 4 | 3 | 2 |
| 2 | TGME49_225140.P1153 | BT1 family protein | 10 | 9 | 9 | 9 | 9 | 3 | 6 | 2 |
| 29 | TGME49_225290-t26_1-p1 | GDA1/CD39 (nucleoside phosphatase) family protein | 10 | 8 | 9 | 7 | 7 | 3 | 3 | 2 |
| 18 | TGME49_225380-t26_1-p1 | hypothetical protein | 9 | 9 | 9 | 7 | 7 | 6 | 5 | 2 |
| 2 | TGME49_225800-t26_1-p1 | iron-sulfur cluster assembly protein SufC | 9 | 8 | 8 | 8 | 8 | 5 | 6 | 2 |
| 29 | TGME49_226060-t26_1-p1 | amino acid transporter AAT1 | 9 | 9 | 8 | 8 | 7 | 3 | 3 | 2 |
| 2 | TGME49_226280.P1200 | ribosomal protein L28, putative | 8 | 8 | 8 | 7 | 7 | 3 | 6 | 2 |
| 18 | TGME49_226370.P1203 | dgat2l1-prov protein | 9 | 9 | 8 | 8 | 7 | 5 | 5 | 2 |
| 2 | TGME49_226580.P1215 | hypothetical protein | 9 | 8 | 8 | 8 | 7 | 3 | 6 | 2 |
| 18 | TGME49_226810.P1223 | histone lysine methyltransferase SET1 | 10 | 10 | 9 | 7 | 7 | 6 | 3 | 2 |
| 18 | TGME49_226840-t26_1-p1 | JmjC domain-containing protein | 10 | 9 | 8 | 7 | 6 | 6 | 6 | 2 |
| 18 | TGME49_226950-t26_1-p1 | IgA-specific metalloendopeptidase | 10 | 10 | 9 | 7 | 6 | 5 | 4 | 2 |
| 18 | TGME49_227000.P1235 | hypothetical protein | 10 | 9 | 8 | 7 | 5 | 4 | 5 | 2 |
| 29 | TGME49_227115-t26_1-p1 | hypothetical protein | 10 | 8 | 8 | 8 | 6 | 4 | 4 | 2 |
| 29 | TGME49_227570.P1250 | amino acid transporter AAT3 | 9 | 9 | 9 | 9 | 8 | 3 | 3 | 2 |
| 29 | TGME49_227780.P1258 | P-type ATPase, putative | 10 | 10 | 9 | 6 | 8 | 3 | 3 | 2 |
| 18 | TGME49_227820-t26_1-p1 | hypothetical protein | 10 | 10 | 10 | 7 | 5 | 6 | 4 | 2 |
| 18 | TGME49_228040-t26_1-p1 | PPIC-type PPIASE domain-containing protein | 10 | 9 | 8 | 6 | 5 | 5 | 5 | 2 |
| 18 | TGME49_228120-t26_1-p1 | hypothetical protein | 10 | 10 | 10 | 9 | 6 | 5 | 4 | 2 |
| 18 | TGME49_228130.P1275 | START domain-containing protein | 10 | 9 | 8 | 7 | 6 | 5 | 4 | 2 |
| 29 | TGME49_228170.P1279 | dense granule protein GRA44 | 10 | 9 | 10 | 10 | 9 | 3 | 3 | 2 |
| 2 | TGME49_228190.P1281 | eukaryotic initiation factor-3, subunit 5, putative | 9 | 9 | 8 | 8 | 8 | 5 | 7 | 2 |
| 29 | TGME49_228460.P1296 | hypothetical protein | 10 | 10 | 9 | 8 | 7 | 4 | 4 | 2 |
| 18 | TGME49_228660.P1298 | Sec7 domain-containing protein | 10 | 10 | 8 | 7 | 5 | 6 | 5 | 2 |
| 18 | TGME49_228690.P1301 | phosphatidylinositol 3- and 4-kinase | 10 | 10 | 7 | 7 | 6 | 6 | 5 | 2 |
| 29 | TGME49_228770.P1306 | hypothetical protein | 9 | 9 | 8 | 7 | 7 | 4 | 4 | 2 |
| 29 | TGME49_229290-t26_1-p1 | kelch repeat-containing protein | 10 | 7 | 9 | 7 | 8 | 4 | 3 | 2 |

| | | | | | | | | | | |
|---|---|---|---|---|---|---|---|---|---|---|
| 29 | TGME49_229340.P1317 | hypothetical protein | 10 | 10 | 9 | 9 | 6 | 4 | 3 | 2 |
| 18 | TGME49_229390.P1322 | hypothetical protein | 10 | 9 | 8 | 8 | 7 | 5 | 4 | 2 |
| 29 | TGME49_229405.P1323 | 3'5'-cyclic nucleotide phosphodiesterase PDE4 | 10 | 10 | 10 | 9 | 7 | 3 | 3 | 2 |
| 18 | TGME49_229630.P1329 | eIF2 kinase IF2K-A (incomplete catalytic triad) | 10 | 10 | 9 | 6 | 5 | 7 | 6 | 2 |
| 2 | TGME49_229920.P1344 | cytochrome c oxidase subunit ApiCOX35 | 8 | 7 | 8 | 7 | 7 | 4 | 7 | 2 |
| 18 | TGME49_230000-t26_1-p1 | hypothetical protein | 10 | 10 | 10 | 7 | 6 | 6 | 5 | 2 |
| 29 | TGME49_230020.P1347 | kinesin motor domain-containing protein | 10 | 10 | 10 | 9 | 8 | 4 | 5 | 2 |
| 18 | TGME49_230080.P1350 | DNA polymerase theta, putative | 10 | 9 | 7 | 7 | 5 | 5 | 5 | 2 |
| 18 | TGME49_230140.P1356 | vacuolar protein sorting-associated protein 9 | 10 | 10 | 8 | 7 | 7 | 6 | 5 | 2 |
| 2 | TGME49_230490.P1372 | phosphatidylinositol-4-phosphate 5-kinase | 10 | 10 | 9 | 7 | 6 | 5 | 7 | 2 |
| 29 | TGME49_230710-t26_1-p1 | cell division protein CDC48 | 10 | 10 | 9 | 8 | 8 | 3 | 3 | 2 |
| 29 | TGME49_230820.P1384 | cell division control protein CDC50.1 | 9 | 9 | 9 | 8 | 7 | 3 | 3 | 2 |
| 2 | TGME49_230830.P1385 | ATPase family associated with various cellular activities (AAA) domain-containing protein | 10 | 10 | 10 | 8 | 8 | 3 | 9 | 2 |
| 29 | TGME49_231090-t26_1-p1 | hypothetical protein | 9 | 9 | 9 | 8 | 6 | 3 | 3 | 2 |
| 18 | TGME49_231130.P1404 | peptidase, M28 family protein | 10 | 9 | 9 | 6 | 6 | 6 | 5 | 2 |
| 18 | TGME49_231150.P1406 | hypothetical protein | 9 | 9 | 8 | 7 | 6 | 5 | 5 | 2 |
| 18 | TGME49_231360.P1413 | hypothetical protein | 10 | 9 | 9 | 8 | 7 | 5 | 5 | 2 |
| 18 | TGME49_231370.P1414 | patatin-like phospholipase PL2 | 10 | 10 | 9 | 7 | 6 | 5 | 5 | 2 |
| 2 | TGME49_231410.P1417 | ATP synthase F0 subunit b-like protein | 10 | 9 | 9 | 8 | 8 | 3 | 8 | 2 |
| 18 | TGME49_231840.P1426 | preconoidal region protein PCR3 | 10 | 10 | 8 | 8 | 5 | 6 | 6 | 2 |
| 2 | TGME49_231910.P1430 | ATP synthase F1 gamma subunit | 8 | 8 | 8 | 8 | 8 | 3 | 7 | 2 |
| 18 | TGME49_232150.P1448 | inner membrane complex protein IMC32 | 10 | 10 | 10 | 8 | 6 | 6 | 3 | 2 |
| 18 | TGME49_232190.P1451 | guanine nucleotide exchange factor GBF1 | 10 | 10 | 10 | 6 | 5 | 5 | 6 | 2 |
| 29 | TGME49_232350-t26_1-p1 | lactate dehydrogenase LDH1 | 8 | 8 | 8 | 8 | 8 | 3 | 3 | 2 |
| 18 | TGME49_232520-t26_1-p1 | brix domain-containing protein | 9 | 9 | 8 | 8 | 7 | 6 | 5 | 2 |
| 18 | TGME49_232530.P1470 | ATP-dependent RNA helicase SUV3, putative | 10 | 10 | 9 | 7 | 6 | 6 | 6 | 2 |
| 29 | TGME49_232580.P1472 | acyl-CoA synthetase ACS6, putative | 10 | 10 | 10 | 9 | 9 | 3 | 3 | 2 |
| 18 | TGME49_232680.P1480 | hypothetical protein | 10 | 10 | 9 | 7 | 6 | 4 | 4 | 2 |
| 18 | TGME49_233020-t26_1-p1 | Not1 N-terminal domain, CCR4-Not complex component protein | 10 | 8 | 9 | 7 | 7 | 5 | 5 | 2 |
| 29 | TGME49_233080.P1492 | G-patch domain-containing protein | 10 | 9 | 10 | 8 | 8 | 5 | 4 | 2 |
| 18 | TGME49_233220.P1501 | hypothetical protein | 10 | 10 | 9 | 7 | 7 | 5 | 5 | 2 |
| 2 | TGME49_233300.P1503 | RhoGAP domain-containing protein | 9 | 9 | 8 | 8 | 8 | 5 | 6 | 2 |
| 29 | TGME49_233320-t26_1-p1 | C2 domain-containing protein | 10 | 9 | 10 | 10 | 6 | 3 | 3 | 2 |
| 2 | TGME49_233410.P1506 | Sof1 family domain-containing protein | 9 | 9 | 9 | 7 | 8 | 4 | 8 | 2 |
| 2 | TGME49_233540.P1510 | apicoplast monocarboxylate transporter AMT1 | 9 | 9 | 8 | 8 | 8 | 4 | 6 | 2 |
| 29 | TGME49_233710.P1512 | inorganic anion transporter, sulfate permease (SulP) family protein | 10 | 10 | 8 | 6 | 8 | 3 | 3 | 2 |
| 18 | TGME49_233810.P1517 | Sel1 repeat-containing protein | 10 | 10 | 10 | 8 | 5 | 5 | 4 | 2 |
| 29 | TGME49_234440-t26_1-p1 | aminotransferase, putative | 9 | 9 | 8 | 8 | 8 | 3 | 3 | 2 |
| 18 | TGME49_234490-t26_1-p1 | kelch repeat-containing protein | 10 | 9 | 10 | 8 | 5 | 5 | 6 | 2 |
| 18 | TGME49_234610.P1543 | WD-40 repeat protein | 10 | 9 | 9 | 8 | 6 | 5 | 6 | 2 |
| 18 | TGME49_235340.P1553 | IMC sutures component ISC1 | 10 | 9 | 8 | 7 | 5 | 6 | 5 | 2 |
| 29 | TGME49_235390.P1557 | PAN domain-containing protein | 10 | 10 | 10 | 9 | 9 | 3 | 5 | 2 |
| 29 | TGME49_235490-t26_1-p1 | HID1 domain-containing protein | 10 | 10 | 9 | 8 | 8 | 3 | 4 | 2 |
| 18 | TGME49_235660.P1570 | PAT complex subunit CCDC47, putative | 9 | 8 | 8 | 8 | 7 | 5 | 5 | 2 |
| 2 | TGME49_235670-t26_1-p1 | hypothetical protein | 9 | 8 | 8 | 8 | 6 | 3 | 6 | 2 |
| 29 | TGME49_235680-t26_1-p1 | peptidase M16 inactive domain-containing protein | 10 | 10 | 10 | 7 | 8 | 3 | 3 | 2 |

| | | | | | | | | | | |
|---|---|---|---|---|---|---|---|---|---|---|
| 2 | TGME49_236640.P1600 | zinc finger, C3HC4 type (RING finger) domain-containing protein | 9 | 9 | 8 | 7 | 7 | 4 | 6 | 2 |
| 18 | TGME49_237010.P1613 | RAP domain-containing protein | 10 | 9 | 9 | 8 | 8 | 5 | 5 | 2 |
| 18 | TGME49_237045.P1615 | hypothetical protein | 10 | 8 | 9 | 8 | 6 | 5 | 5 | 2 |
| 18 | TGME49_237100-t26_1-p1 | RAP domain-containing protein | 10 | 10 | 10 | 8 | 6 | 4 | 5 | 2 |
| 29 | TGME49_237140-t26_1-p1 | pyridoxal 5'-phosphate synthase subunit PDX1 | 8 | 8 | 8 | 8 | 7 | 3 | 3 | 2 |
| 2 | TGME49_237280.P1625 | TBC domain-containing protein TBC6 | 10 | 10 | 10 | 6 | 8 | 5 | 6 | 2 |
| 29 | TGME49_237530-t26_1-p1 | hypothetical protein | 10 | 9 | 9 | 8 | 7 | 4 | 5 | 2 |
| 2 | TGME49_238180-t26_1-p1 | 26S proteasome regulatory subunit RPN7, putative | 9 | 9 | 9 | 7 | 8 | 4 | 6 | 2 |
| 18 | TGME49_238510.P1654 | zinc finger protein, putative | 10 | 10 | 9 | 8 | 7 | 6 | 4 | 2 |
| 29 | TGME49_238880-t26_1-p1 | hypothetical protein | 10 | 10 | 10 | 9 | 7 | 5 | 4 | 2 |
| 18 | TGME49_239250.P1666 | diacylglycerol kinase DGK3 | 10 | 10 | 8 | 6 | 6 | 6 | 4 | 2 |
| 29 | TGME49_239300.P1669 | intraconoidal microtubule-associated protein ICMAP1 | 10 | 10 | 10 | 10 | 8 | 3 | 3 | 2 |
| 2 | TGME49_239410-t26_1-p1 | CCR4-NOT transcription complex subunit 4, putative | 10 | 10 | 6 | 7 | 6 | 3 | 6 | 2 |
| 29 | TGME49_239510.P1677 | START domain-containing protein | 9 | 8 | 8 | 8 | 8 | 3 | 3 | 2 |
| 2 | TGME49_239540.P1679 | cell division control protein CDC50.3 | 9 | 9 | 8 | 8 | 8 | 5 | 7 | 2 |
| 18 | TGME49_239590-t26_1-p1 | WD domain, G-beta repeat-containing protein | 10 | 9 | 9 | 9 | 6 | 6 | 5 | 2 |
| 2 | TGME49_240280-t26_1-p1 | S1/P1nuclease | 9 | 9 | 9 | 8 | 7 | 4 | 7 | 2 |
| 18 | TGME49_240390.P1701 | calcium-dependent protein kinase CDPK4B | 10 | 10 | 7 | 6 | 6 | 5 | 5 | 2 |
| 18 | TGME49_240540-t26_1-p1 | conserved oligomeric Golgi complex subunit 8 | 9 | 9 | 8 | 7 | 5 | 5 | 5 | 2 |
| 29 | TGME49_240570.P1708 | hypothetical protein | 9 | 9 | 9 | 8 | 7 | 4 | 4 | 2 |
| 2 | TGME49_241170-t26_1-p1 | KH domain-containing protein | 10 | 10 | 10 | 9 | 9 | 3 | 8 | 2 |
| 2 | TGME49_241820.P1727 | membrane protein, putative | 8 | 8 | 7 | 7 | 7 | 3 | 6 | 2 |
| 29 | TGME49_241840-t26_1-p1 | C2 domain-containing protein | 9 | 9 | 8 | 8 | 7 | 3 | 3 | 2 |
| 2 | TGME49_241860-t26_1-p1 | N-acetylglucosaminyl-phosphatidylinositol biosynthetic protein PigA, family GT4 protein | 10 | 9 | 9 | 8 | 8 | 4 | 8 | 2 |
| 2 | TGME49_242010-t26_1-p1 | glucose inhibited division protein A subfamily protein | 10 | 9 | 9 | 8 | 7 | 4 | 6 | 2 |
| 2 | TGME49_242415-t26_1-p1 | histone lysine-specific demethylase | 10 | 10 | 10 | 8 | 6 | 5 | 7 | 2 |
| 29 | TGME49_242600-t26_1-p1 | 6-phosphogluconate dehydrogenase | 9 | 8 | 9 | 9 | 7 | 3 | 3 | 2 |
| 18 | TGME49_242780.P1746 | hypothetical protein | 10 | 10 | 10 | 7 | 7 | 5 | 5 | 2 |
| 29 | TGME49_242860-t26_1-p1 | hypothetical protein | 9 | 9 | 8 | 8 | 6 | 4 | 4 | 2 |
| 18 | TGME49_243280-t26_1-p1 | Met-10+ like-protein | 10 | 9 | 8 | 7 | 6 | 5 | 6 | 2 |
| 18 | TGME49_243460.P1761 | hypothetical protein | 10 | 10 | 9 | 6 | 6 | 6 | 5 | 2 |
| 18 | TGME49_243590-t26_1-p1 | endonuclease/exonuclease/phosphatase family protein | 10 | 9 | 9 | 6 | 6 | 6 | 6 | 2 |
| 29 | TGME49_243655.P1768 | GMC oxidoreductase | 10 | 10 | 10 | 9 | 8 | 3 | 3 | 2 |
| 18 | TGME49_243930.P1774 | GPI-anchored micronemal antigen GAMA | 10 | 9 | 9 | 8 | 6 | 5 | 5 | 2 |
| 18 | TGME49_243960.P1776 | nuclear transport factor 2 (ntf2) domain-containing protein | 10 | 9 | 9 | 8 | 6 | 6 | 5 | 2 |
| 2 | TGME49_244370.P1788 | TDC1, putative | 9 | 8 | 9 | 8 | 6 | 3 | 7 | 2 |
| 18 | TGME49_244520.P1792 | glycosyl transferase, group 4 family protein | 9 | 9 | 8 | 8 | 6 | 6 | 4 | 2 |
| 18 | TGME49_244580.P1795 | L1P family of ribosomal protein | 9 | 8 | 8 | 8 | 6 | 5 | 5 | 2 |
| 29 | TGME49_245490.P1807 | microneme protein MIC8 | 10 | 9 | 9 | 8 | 8 | 4 | 4 | 2 |
| 18 | TGME49_245570-t26_1-p1 | origin recognition complex subunit 2 protein | 10 | 9 | 9 | 8 | 7 | 6 | 5 | 2 |
| 18 | TGME49_245660.P1815 | zinc finger protein, putative | 10 | 9 | 9 | 7 | 6 | 4 | 4 | 2 |
| 2 | TGME49_245720-t26_1-p1 | SWI2/SNF2-containing protein | 10 | 10 | 10 | 9 | 8 | 5 | 6 | 2 |

| | | | | | | | | | | |
|---|---|---|---|---|---|---|---|---|---|---|
| 18 | TGME49_245730-t26_1-p1 | phosphatidylinositol-4-phosphate 5-Kinase | 10 | 10 | 7 | 6 | 7 | 6 | 5 | 2 |
| 2 | TGME49_246170.P1826 | ARID/BRIGHT DNA binding domain-containing protein | 10 | 10 | 9 | 8 | 6 | 3 | 6 | 2 |
| 2 | TGME49_246178.P1827 | hypothetical protein | 9 | 9 | 8 | 8 | 7 | 3 | 6 | 2 |
| 18 | TGME49_246475-t26_1-p1 | hypothetical protein | 10 | 9 | 8 | 7 | 5 | 5 | 4 | 2 |
| 18 | TGME49_246520-t26_1-p1 | hypothetical protein | 10 | 10 | 9 | 8 | 6 | 7 | 5 | 2 |
| 18 | TGME49_246960-t26_1-p1 | hypothetical protein | 10 | 9 | 8 | 7 | 5 | 6 | 5 | 2 |
| 2 | TGME49_246970-t26_1-p1 | 3'-5' exonuclease domain-containing protein | 10 | 10 | 9 | 9 | 6 | 4 | 7 | 2 |
| 18 | TGME49_247000-t26_1-p1 | tetratricopeptide repeat-containing protein | 10 | 9 | 8 | 7 | 6 | 5 | 5 | 2 |
| 18 | TGME49_247195.P1845 | rhoptry discharge factor RDF1 | 10 | 10 | 10 | 7 | 6 | 6 | 3 | 2 |
| 18 | TGME49_247300-t26_1-p1 | hypothetical protein | 10 | 9 | 9 | 8 | 5 | 5 | 4 | 2 |
| 18 | TGME49_247330-t26_1-p1 | symplekin, putative | 10 | 10 | 10 | 8 | 7 | 6 | 4 | 2 |
| 18 | TGME49_247450-t26_1-p1 | hypothetical protein | 10 | 10 | 9 | 5 | 7 | 6 | 5 | 2 |
| 2 | TGME49_247600-t26_1-p1 | dynein light intermediate chain, putative | 10 | 9 | 9 | 8 | 7 | 3 | 6 | 2 |
| 18 | TGME49_247680-t26_1-p1 | hypothetical protein | 10 | 9 | 8 | 8 | 7 | 6 | 5 | 2 |
| 18 | TGME49_248150-t26_1-p1 | AP-5 complex subunit zeta, putative | 10 | 9 | 9 | 9 | 5 | 5 | 5 | 2 |
| 29 | TGME49_248160.P1876 | ATP-dependent DNA helicase 2 subunit KU70 | 10 | 9 | 9 | 8 | 7 | 4 | 3 | 2 |
| 29 | TGME49_248225.P1879 | PF2 arrest specific protein 8/11, putative | 9 | 7 | 9 | 7 | 7 | 4 | 4 | 2 |
| 18 | TGME49_248290.P1880 | WD domain, G-beta repeat-containing protein | 10 | 10 | 9 | 8 | 5 | 6 | 3 | 2 |
| 29 | TGME49_248300.P1881 | hypothetical protein | 10 | 9 | 8 | 8 | 6 | 3 | 3 | 2 |
| 29 | TGME49_248420.P1888 | apicomplexan amino acid transporter ApiAT3-2 | 10 | 9 | 9 | 7 | 6 | 3 | 3 | 2 |
| 29 | TGME49_248450.P1889 | zinc finger, C3HC4 type (RING finger) domain-containing protein | 10 | 10 | 9 | 9 | 8 | 3 | 3 | 2 |
| 29 | TGME49_248550.P1893 | SPX domain-containing protein | 10 | 7 | 9 | 6 | 8 | 4 | 5 | 2 |
| 29 | TGME49_248610.P1897 | apicomplexan amino acid transporter ApiAT5-1 | 10 | 9 | 9 | 7 | 6 | 3 | 3 | 2 |
| 29 | TGME49_248730-t26_1-p1 | hypothetical protein | 9 | 9 | 9 | 9 | 8 | 3 | 3 | 2 |
| 29 | TGME49_248760-t26_1-p1 | subtilisin SUB7 | 10 | 9 | 9 | 8 | 8 | 4 | 3 | 2 |
| 18 | TGME49_248850.P1909 | methionine aminopeptidase 1b | 9 | 9 | 9 | 8 | 8 | 5 | 5 | 2 |
| 29 | TGME49_249230.P1921 | hypothetical protein | 10 | 10 | 10 | 7 | 7 | 3 | 3 | 2 |
| 18 | TGME49_249360-t26_1-p1 | RED family protein | 10 | 9 | 9 | 6 | 6 | 6 | 6 | 2 |
| 29 | TGME49_249540.P1936 | hypothetical protein | 10 | 10 | 9 | 8 | 7 | 3 | 3 | 2 |
| 18 | TGME49_249560-t26_1-p1 | DNA-directed RNA polymerase alpha chain rpoA | 10 | 10 | 10 | 9 | 4 | 5 | 5 | 2 |
| 18 | TGME49_249610-t26_1-p1 | hypothetical protein | 10 | 10 | 10 | 6 | 7 | 5 | 4 | 2 |
| 29 | TGME49_249990.P1955 | dense granule protein GRA70 | 10 | 9 | 9 | 8 | 8 | 4 | 5 | 2 |
| 18 | TGME49_250680.P1960 | TBC domain-containing protein TBC5 | 10 | 10 | 9 | 8 | 6 | 7 | 5 | 2 |
| 18 | TGME49_250840-t26_1-p1 | myosin light chain MLC3 | 10 | 9 | 8 | 7 | 6 | 5 | 3 | 2 |
| 18 | TGME49_251410-t26_1-p1 | transmembrane pellicle protein PP1 | 10 | 10 | 10 | 7 | 5 | 4 | 5 | 2 |
| 29 | TGME49_251760-t26_1-p1 | subunit of proteaseome activator complex, putative | 8 | 8 | 8 | 8 | 7 | 3 | 3 | 2 |
| 18 | TGME49_252380.P1995 | phosphatase, putative | 10 | 9 | 9 | 8 | 7 | 5 | 5 | 2 |
| 18 | TGME49_252510.P2001 | Vid27 domain-containing protein | 10 | 10 | 9 | 9 | 5 | 5 | 6 | 2 |
| 2 | TGME49_253090.P2010 | DEAD/DEAH box helicase domain-containing protein | 9 | 9 | 9 | 9 | 6 | 5 | 7 | 2 |
| 18 | TGME49_253170.P2014 | zinc carboxypeptidase, putative | 10 | 10 | 9 | 8 | 7 | 5 | 4 | 2 |
| 18 | TGME49_253370.P2019 | rhoptry neck protein RON4L1 | 10 | 10 | 10 | 9 | 6 | 5 | 5 | 2 |
| 18 | TGME49_253650.P2032 | DnaJ C terminal region domain-containing protein | 10 | 8 | 8 | 7 | 6 | 5 | 5 | 2 |
| 18 | TGME49_253710-t26_1-p1 | hypothetical protein | 10 | 9 | 9 | 9 | 7 | 5 | 5 | 2 |
| 18 | TGME49_253750.P2034 | PLU-1 family protein | 10 | 10 | 10 | 7 | 7 | 6 | 5 | 2 |
| 18 | TGME49_253880-t26_1-p1 | GNS1/SUR4 family protein | 10 | 9 | 8 | 7 | 6 | 4 | 5 | 2 |

| | | | | | | | | | | |
|---|---|---|---|---|---|---|---|---|---|---|
| 29 | TGME49_254080-t26_1-p1 | metal cation transporter, ZIP family protein | 9 | 9 | 8 | 8 | 6 | 3 | 3 | 2 |
| 18 | TGME49_254110-t26_1-p1 | tryptophanyl-tRNA synthetase (TrpRS1) | 10 | 9 | 8 | 7 | 6 | 5 | 4 | 2 |
| 18 | TGME49_254200.P2051 | anticodon binding domain-containing protein | 10 | 8 | 9 | 7 | 6 | 7 | 5 | 2 |
| 2 | TGME49_254400.P2055 | LSU ribosomal protein L2P, putative | 9 | 8 | 8 | 8 | 6 | 3 | 6 | 2 |
| 29 | TGME49_254430-t26_1-p1 | microneme protein, putative | 10 | 9 | 9 | 7 | 6 | 3 | 3 | 2 |
| 2 | TGME49_254555.P2060 | histone lysine acetyltransferase GCN5-A | 10 | 9 | 9 | 9 | 8 | 3 | 6 | 2 |
| 2 | TGME49_254580.P2061 | adenine nucleotide transporter ANT | 9 | 8 | 8 | 6 | 7 | 4 | 6 | 2 |
| 18 | TGME49_254730-t26_1-p1 | POPLD (NUC188) domain-containing protein | 10 | 10 | 10 | 8 | 5 | 5 | 4 | 2 |
| 29 | TGME49_255240.P2091 | hypothetical protein | 10 | 10 | 10 | 7 | 7 | 4 | 4 | 2 |
| 29 | TGME49_255340.P2095 | tetratricopeptide repeat-containing protein | 10 | 10 | 10 | 10 | 8 | 5 | 4 | 2 |
| 29 | TGME49_255500.P2107 | tetratricopeptide repeat-containing protein | 10 | 9 | 9 | 9 | 7 | 3 | 5 | 2 |
| 18 | TGME49_255870.P2118 | WD domain, G-beta repeat-containing protein | 10 | 9 | 9 | 7 | 6 | 5 | 5 | 2 |
| 29 | TGME49_255890.P2120 | pyridine nucleotide-disulfide oxidoreductase domain-containing protein | 10 | 9 | 9 | 8 | 7 | 3 | 3 | 2 |
| 18 | TGME49_255960.P2124 | hypothetical protein | 10 | 10 | 9 | 9 | 5 | 5 | 4 | 2 |
| 18 | TGME49_256060-t26_1-p1 | nucleosome assembly protein (nap) protein | 9 | 9 | 8 | 7 | 7 | 5 | 5 | 2 |
| 18 | TGME49_256070.P2131 | CDK-related protein kinase CRK4 | 10 | 9 | 9 | 7 | 7 | 6 | 5 | 2 |
| 29 | TGME49_256865.P2144 | WD repeat domain 35 family protein | 10 | 10 | 10 | 7 | 9 | 5 | 4 | 2 |
| 18 | TGME49_257000.P2149 | solute carrier family 13, putative | 10 | 9 | 9 | 8 | 6 | 6 | 6 | 2 |
| 18 | TGME49_257180.P2162 | DNA repair protein RAD50, putative | 10 | 10 | 10 | 6 | 7 | 5 | 6 | 2 |
| 29 | TGME49_257510.P2177 | CDP-alcohol phosphatidyltransferase superfamily protein | 9 | 9 | 9 | 8 | 7 | 5 | 3 | 2 |
| 18 | TGME49_257710.P2184 | actin-like protein ALP 5 | 10 | 9 | 8 | 6 | 6 | 5 | 5 | 2 |
| 29 | TGME49_257720.P2185 | proton ATPase, putative | 10 | 10 | 10 | 9 | 9 | 3 | 3 | 2 |
| 29 | TGME49_257770-t26_1-p1 | histone lysine methyltransferase SET2 | 10 | 10 | 10 | 9 | 6 | 3 | 3 | 2 |
| 18 | TGME49_257960.P2194 | GDP-D-mannose pyrophosphorylase | 9 | 9 | 9 | 8 | 7 | 6 | 5 | 2 |
| 18 | TGME49_258070.P2203 | signal recognition particle subunit SRP68, putative | 10 | 9 | 9 | 5 | 7 | 6 | 4 | 2 |
| 18 | TGME49_258090.P2205 | hypothetical protein | 10 | 10 | 9 | 9 | 6 | 6 | 3 | 2 |
| 18 | TGME49_258560.P2228 | hypothetical protein | 10 | 9 | 9 | 8 | 7 | 5 | 4 | 2 |
| 29 | TGME49_259190.P2269 | hypothetical protein | 10 | 9 | 9 | 7 | 7 | 4 | 4 | 2 |
| 18 | TGME49_259550.P2278 | hydroxymethyldihydropterin pyrophosphokinase-dihydropteroate synthase | 10 | 9 | 9 | 8 | 5 | 4 | 6 | 2 |
| 18 | TGME49_259560.P2279 | endonuclease/exonuclease/phosphatase family protein | 10 | 10 | 10 | 7 | 5 | 6 | 6 | 2 |
| 18 | TGME49_259640.P2284 | nucleoporin NUP302 | 10 | 10 | 10 | 8 | 5 | 5 | 5 | 2 |
| 18 | TGME49_259830.P2289 | diacylglycerol kinase DGK2 | 10 | 9 | 8 | 8 | 7 | 5 | 4 | 2 |
| 18 | TGME49_259960.P2298 | Nucleoside-diphosphatase | 9 | 9 | 8 | 7 | 5 | 5 | 5 | 2 |
| 18 | TGME49_260230.P2311 | hypothetical protein | 9 | 8 | 9 | 6 | 5 | 5 | 5 | 2 |
| 18 | TGME49_260360.P2316 | GTP-binding protein | 10 | 9 | 9 | 8 | 6 | 5 | 4 | 2 |
| 18 | TGME49_260410.P2320 | hypothetical protein | 10 | 10 | 9 | 7 | 6 | 4 | 4 | 2 |
| 29 | TGME49_260450-t26_1-p1 | DEAD/DEAH box helicase domain-containing protein | 10 | 10 | 10 | 8 | 6 | 5 | 3 | 2 |
| 18 | TGME49_260480.P2325 | leucine rich repeat-containing protein | 10 | 10 | 10 | 7 | 5 | 6 | 3 | 2 |
| 18 | TGME49_260570.P2331 | replication protein A, putative | 10 | 9 | 9 | 8 | 7 | 5 | 5 | 2 |
| 2 | TGME49_260680.P2339 | small subunit DNA primase, putative | 9 | 9 | 9 | 7 | 7 | 3 | 8 | 2 |
| 18 | TGME49_261400-t26_1-p1 | hypothetical protein | 9 | 9 | 8 | 6 | 6 | 5 | 5 | 2 |
| 18 | TGME49_261680.P2376 | hypothetical protein | 10 | 10 | 9 | 8 | 5 | 6 | 4 | 2 |
| 2 | TGME49_261760.P2382 | CDP-alcohol phosphatidyltransferase superfamily protein | 9 | 9 | 8 | 8 | 8 | 5 | 7 | 2 |

| | | | | | | | | | | |
|---|---|---|---|---|---|---|---|---|---|---|
| 18 | TGME49_261940.P2385 | hydrolase, alpha/beta fold family protein | 10 | 9 | 8 | 6 | 7 | 5 | 5 | 2 |
| 18 | TGME49_262100-t26_1-p1 | tetratricopeptide repeat-containing protein | 9 | 9 | 8 | 7 | 6 | 4 | 5 | 2 |
| 2 | TGME49_262380-t26_1-p1 | elongation factor Tu, putative | 9 | 9 | 9 | 9 | 9 | 3 | 9 | 2 |
| 2 | TGME49_262390.P2400 | TLD protein | 10 | 10 | 8 | 7 | 7 | 4 | 6 | 2 |
| 18 | TGME49_262400.P2401 | lipase | 10 | 10 | 9 | 8 | 6 | 7 | 4 | 2 |
| 18 | TGME49_262420.P2402 | AP2 domain transcription factor AP2VIIb-1/ADA2-B | 10 | 10 | 9 | 6 | 6 | 5 | 4 | 2 |
| 18 | TGME49_262650.P2417 | WD domain, G-beta repeat-containing protein | 10 | 9 | 9 | 7 | 5 | 6 | 4 | 2 |
| 18 | TGME49_262700.P2423 | tetratricopeptide repeat-containing protein | 10 | 9 | 9 | 8 | 5 | 6 | 6 | 2 |
| 18 | TGME49_262760.P2426 | poly(ADP-ribose) glycohydrolase | 10 | 9 | 9 | 7 | 5 | 5 | 4 | 2 |
| 18 | TGME49_262950-t26_1-p1 | hypothetical protein | 10 | 9 | 9 | 8 | 6 | 4 | 5 | 2 |
| 29 | TGME49_263260.P2457 | apicomplexan amino acid transporter ApiAT7-2 | 10 | 9 | 9 | 7 | 7 | 3 | 3 | 2 |
| 29 | TGME49_263510.P2473 | Spc97 / Spc98 family protein | 10 | 10 | 10 | 7 | 8 | 5 | 4 | 2 |
| 18 | TGME49_263560.P2476 | hypothetical protein | 9 | 9 | 9 | 7 | 6 | 4 | 4 | 2 |
| 18 | TGME49_263610.P2482 | tetratricopeptide repeat-containing protein | 10 | 10 | 9 | 8 | 7 | 5 | 5 | 2 |
| 18 | TGME49_263740.P2490 | ABC transporter transmembrane region domain-containing protein | 10 | 9 | 8 | 7 | 5 | 7 | 3 | 2 |
| 18 | TGME49_264020.P2502 | bromodomain-containing protein BDP5 | 10 | 9 | 8 | 7 | 5 | 6 | 5 | 2 |
| 18 | TGME49_264090-t26_1-p1 | TRAPPC11-like protein | 10 | 10 | 9 | 7 | 6 | 6 | 6 | 2 |
| 18 | TGME49_264120.P2506 | Myb family DNA-binding domain-containing protein | 10 | 10 | 9 | 8 | 6 | 5 | 3 | 2 |
| 18 | TGME49_264640.P2518 | bromodomain-containing protein BDP2 | 10 | 10 | 8 | 7 | 5 | 5 | 4 | 2 |
| 29 | TGME49_264720-t26_1-p1 | hypothetical protein | 10 | 9 | 9 | 7 | 6 | 4 | 3 | 2 |
| 18 | TGME49_264740.P2522 | hypothetical protein | 10 | 10 | 9 | 7 | 5 | 6 | 6 | 2 |
| 18 | TGME49_264752.P2523 | HEAT repeat-containing protein | 10 | 10 | 10 | 6 | 5 | 5 | 5 | 2 |
| 29 | TGME49_264840-t26_1-p1 | ATP-dependent DNA helicase, RecQ family protein | 10 | 10 | 10 | 6 | 7 | 3 | 3 | 2 |
| 18 | TGME49_264860.P2525 | zinc finger, C3HC4 type (RING finger) domain-containing protein | 10 | 10 | 10 | 6 | 7 | 4 | 5 | 2 |
| 29 | TGME49_265520.P2541 | hypothetical protein | 9 | 9 | 8 | 8 | 7 | 3 | 3 | 2 |
| 18 | TGME49_265850.P2550 | tetratricopeptide repeat-containing protein | 10 | 10 | 9 | 6 | 6 | 5 | 5 | 2 |
| 29 | TGME49_265870.P2552 | pantoate-beta-alanine ligase | 10 | 8 | 9 | 8 | 6 | 4 | 4 | 2 |
| 29 | TGME49_266110-t26_1-p1 | DEAD/DEAH box helicase domain-containing protein | 10 | 9 | 9 | 7 | 8 | 5 | 4 | 2 |
| 29 | TGME49_266366.P2564 | BT1 family protein | 10 | 10 | 9 | 9 | 9 | 3 | 3 | 2 |
| 29 | TGME49_266660.P2575 | mediator complex subunit MED9 | 10 | 9 | 9 | 9 | 7 | 4 | 3 | 2 |
| 2 | TGME49_266860.P2585 | BTB/POZ domain-containing protein | 9 | 8 | 8 | 8 | 8 | 4 | 6 | 2 |
| 29 | TGME49_267070-t26_1-p1 | aquaporin 2 | 9 | 9 | 8 | 8 | 7 | 3 | 5 | 2 |
| 18 | TGME49_267370.P2599 | kinesin A | 10 | 10 | 8 | 7 | 6 | 5 | 5 | 2 |
| 2 | TGME49_267380.P2600 | nucleotide-sugar transporter NST1 | 9 | 8 | 8 | 7 | 5 | 3 | 7 | 2 |
| 2 | TGME49_267390-t26_1-p1 | DNA-directed RNA polymerase I RPAC1 | 9 | 8 | 8 | 5 | 8 | 3 | 7 | 2 |
| 18 | TGME49_267580-t26_1-p1 | cyclin2 related protein | 10 | 9 | 7 | 7 | 6 | 6 | 5 | 2 |
| 18 | TGME49_267660-t26_1-p1 | hypothetical protein | 10 | 9 | 10 | 8 | 7 | 5 | 4 | 2 |
| 18 | TGME49_268010.P2621 | hypothetical protein | 10 | 10 | 8 | 6 | 5 | 6 | 4 | 2 |
| 18 | TGME49_268250-t26_1-p1 | WD domain, G-beta repeat-containing protein | 9 | 9 | 9 | 8 | 6 | 7 | 5 | 2 |
| 29 | TGME49_268290.P2630 | DEAD/DEAH box helicase domain-containing protein | 10 | 9 | 9 | 8 | 8 | 3 | 3 | 2 |
| 29 | TGME49_268310-t26_1-p1 | oocyst wall protein OWP3 | 10 | 9 | 9 | 9 | 8 | 3 | 3 | 2 |
| 18 | TGME49_268330.P2633 | trafficking protein particle complex subunit TRS120, putative | 10 | 10 | 9 | 8 | 5 | 6 | 6 | 2 |

| | | | | | | | | | | |
|---|---|---|---|---|---|---|---|---|---|---|
| 18 | TGME49_268560.P2638 | XPG N-terminal domain-containing protein | 10 | 10 | 9 | 6 | 7 | 5 | 5 | 2 |
| 29 | TGME49_268580.P2640 | hypothetical protein | 9 | 9 | 9 | 8 | 8 | 4 | 3 | 2 |
| 18 | TGME49_268600-t26_1-p1 | DNA polymerase epsilon subunit B protein | 10 | 10 | 9 | 8 | 7 | 6 | 5 | 2 |
| 29 | TGME49_268650.P2642 | chaperone protein ClpB2 | 10 | 10 | 10 | 10 | 10 | 3 | 3 | 2 |
| 18 | TGME49_268770-t26_1-p1 | dual specificity phosphatase, catalytic domain-containing protein | 10 | 9 | 9 | 8 | 7 | 6 | 4 | 2 |
| 29 | TGME49_268860.P2654 | enolase 1 | 9 | 9 | 9 | 9 | 9 | 3 | 3 | 2 |
| 29 | TGME49_269120.P2664 | oxidoreductase, putative | 9 | 9 | 8 | 8 | 8 | 4 | 4 | 2 |
| 29 | TGME49_269320.P2680 | hypothetical protein | 10 | 10 | 10 | 9 | 9 | 5 | 5 | 2 |
| 29 | TGME49_269400.P2681 | oxidoreductase, short chain dehydrogenase/reductase family protein | 9 | 9 | 8 | 7 | 8 | 3 | 4 | 2 |
| 18 | TGME49_269460.P2685 | basal complex component BCC5 | 10 | 10 | 6 | 7 | 5 | 6 | 6 | 2 |
| 2 | TGME49_269660.P2693 | TFIIH basal transcription factor complex helicase XPB subunit | 10 | 10 | 10 | 9 | 9 | 3 | 9 | 2 |
| 29 | TGME49_269690.P2695 | dense granule protein GRA29 | 10 | 10 | 9 | 9 | 7 | 3 | 3 | 2 |
| 29 | TGME49_269860.P2706 | hypothetical protein | 8 | 8 | 8 | 8 | 8 | 3 | 3 | 2 |
| 18 | TGME49_269900.P2708 | WD domain, G-beta repeat-containing protein | 10 | 10 | 9 | 5 | 6 | 5 | 5 | 2 |
| 18 | TGME49_270330.P2733 | cyclin-dependent kinase CDK7 | 9 | 8 | 8 | 8 | 6 | 6 | 5 | 2 |
| 18 | TGME49_270340.P2734 | BTB/POZ domain-containing protein | 10 | 9 | 8 | 8 | 5 | 5 | 4 | 2 |
| 29 | TGME49_270670-t26_1-p1 | hypothetical protein | 10 | 10 | 10 | 9 | 6 | 3 | 3 | 2 |
| 2 | TGME49_270750.P2781 | hypothetical protein | 10 | 9 | 9 | 8 | 5 | 3 | 6 | 2 |
| 18 | TGME49_270770.P2782 | PWI domain-containing protein | 10 | 9 | 8 | 6 | 6 | 7 | 5 | 2 |
| 29 | TGME49_270900-t26_1-p1 | ATPase, AAA family protein | 10 | 10 | 10 | 8 | 6 | 3 | 4 | 2 |
| 2 | TGME49_270960.P2793 | U3 small nucleolar RNA-associated protein 25, putative | 10 | 10 | 10 | 9 | 8 | 3 | 7 | 2 |
| 18 | TGME49_271100.P2800 | NUC153 domain-containing protein | 9 | 9 | 8 | 8 | 7 | 6 | 5 | 2 |
| 18 | TGME49_271250.P2806 | KH domain-containing protein | 9 | 9 | 8 | 7 | 6 | 5 | 5 | 2 |
| 2 | TGME49_271460.P2816 | protein c14orf29, putative | 10 | 9 | 9 | 8 | 7 | 5 | 7 | 2 |
| 29 | TGME49_271470.P2817 | hypothetical protein | 10 | 10 | 10 | 10 | 10 | 4 | 4 | 2 |
| 18 | TGME49_271625.P2820 | serine--tRNA ligase | 10 | 9 | 8 | 7 | 6 | 6 | 3 | 2 |
| 2 | TGME49_271820-t26_1-p1 | lipoate-protein ligase 1, putative | 9 | 8 | 8 | 8 | 7 | 3 | 6 | 2 |
| 29 | TGME49_271840-t26_1-p1 | RAP domain-containing protein | 10 | 10 | 9 | 9 | 8 | 4 | 3 | 2 |
| 18 | TGME49_271960-t26_1-p1 | haloacid dehalogenase-like hydrolase, putative | 10 | 9 | 9 | 7 | 7 | 5 | 5 | 2 |
| 29 | TGME49_272030.P2838 | kelch repeat-containing protein | 10 | 8 | 9 | 8 | 6 | 4 | 3 | 2 |
| 2 | TGME49_272270.P2847 | radical SAM domain-containing protein | 10 | 10 | 9 | 6 | 8 | 3 | 7 | 2 |
| 29 | TGME49_272430.P2863 | perforin-like protein PLP2 | 10 | 10 | 9 | 9 | 8 | 3 | 3 | 2 |
| 18 | TGME49_272670.P2879 | mitochondrial intermediate peptidase, putative | 10 | 10 | 9 | 8 | 6 | 6 | 4 | 2 |
| 29 | TGME49_273360.P2898 | ABC transporter transmembrane region domain-containing protein | 10 | 10 | 10 | 9 | 9 | 3 | 3 | 2 |
| 29 | TGME49_273500.P2906 | O-linked N-acetylglucosamine transferase SPINDLY | 10 | 10 | 10 | 10 | 6 | 3 | 3 | 2 |
| 18 | TGME49_273510-t26_1-p1 | hypothetical protein | 10 | 9 | 9 | 8 | 5 | 5 | 5 | 2 |
| 29 | TGME49_273690.P2916 | Wee kinase | 10 | 9 | 9 | 9 | 8 | 5 | 5 | 2 |
| 18 | TGME49_273780.P2919 | SWI2/SNF2-containing protein | 10 | 10 | 9 | 8 | 6 | 5 | 5 | 2 |
| 18 | TGME49_273840.P2922 | brix domain-containing protein | 10 | 9 | 8 | 6 | 6 | 6 | 4 | 2 |
| 29 | TGME49_273970.P2929 | CorA family Mg2+ transporter protein | 9 | 9 | 9 | 8 | 8 | 3 | 3 | 2 |
| 18 | TGME49_274070.P2935 | ThiF family protein | 10 | 9 | 9 | 8 | 6 | 6 | 6 | 2 |
| 18 | TGME49_274130.P2941 | TBC domain-containing protein TBC2 | 10 | 10 | 8 | 7 | 7 | 7 | 5 | 2 |
| 29 | TGME49_274150.P2943 | hypothetical protein | 10 | 8 | 9 | 7 | 7 | 3 | 5 | 2 |

| | | | | | | | | | | |
|---|---|---|---|---|---|---|---|---|---|---|
| 18 | TGME49_275420-t26_1-p1 | histone lysine-specific demethylase LSD1/BHC110/KDMA1A | 10 | 9 | 8 | 6 | 7 | 7 | 5 | 2 |
| 18 | TGME49_275610-t26_1-p1 | protein kinase, other | 10 | 9 | 9 | 8 | 5 | 7 | 4 | 2 |
| 18 | TGME49_275670.P2957 | alveolin domain containing intermediate filament IMC15 | 10 | 8 | 8 | 8 | 7 | 6 | 5 | 2 |
| 18 | TGME49_275680.P2958 | ELM2 domain-containing protein | 10 | 8 | 8 | 7 | 6 | 5 | 6 | 2 |
| 18 | TGME49_275802-t26_1-p1 | SRP72 RNA-binding domain-containing protein | 10 | 9 | 9 | 8 | 6 | 7 | 4 | 2 |
| 18 | TGME49_276190-t26_1-p1 | hypothetical protein | 9 | 8 | 8 | 8 | 7 | 5 | 5 | 2 |
| 18 | TGME49_277030-t26_1-p1 | isoleucyl-tRNA synthetase, putative | 10 | 10 | 9 | 7 | 5 | 5 | 5 | 2 |
| 2 | TGME49_277070-t26_1-p1 | SWI2/SNF2-containing protein | 9 | 9 | 3 | 9 | 9 | 3 | 6 | 2 |
| 18 | TGME49_277090.P2977 | mitochondrial iron transporter MIT | 10 | 9 | 8 | 7 | 6 | 5 | 4 | 2 |
| 18 | TGME49_277910-t26_1-p1 | thrombospondin type 1 domain-containing protein | 10 | 9 | 8 | 6 | 7 | 5 | 5 | 2 |
| 29 | TGME49_278050-t26_1-p1 | proteasome subunit alpha type-1, putative | 8 | 8 | 8 | 8 | 7 | 3 | 4 | 2 |
| 29 | TGME49_278120-t26_1-p1 | SCP family extracellular subfamily protein | 10 | 9 | 8 | 9 | 8 | 3 | 3 | 2 |
| 18 | TGME49_278250-t26_1-p1 | hypothetical protein | 9 | 9 | 9 | 5 | 6 | 6 | 5 | 2 |
| 29 | TGME49_278280-t26_1-p1 | WD domain, G-beta repeat-containing protein | 10 | 9 | 9 | 7 | 7 | 3 | 4 | 2 |
| 2 | TGME49_278640.P3006 | MIZ/SP-RING zinc finger domain-containing protein | 10 | 9 | 9 | 8 | 8 | 5 | 6 | 2 |
| 18 | TGME49_278700.P3009 | hypothetical protein | 9 | 9 | 8 | 8 | 7 | 5 | 5 | 2 |
| 18 | TGME49_278800-t26_1-p1 | zinc finger protein 36 family 3 protein | 10 | 9 | 8 | 7 | 6 | 5 | 5 | 2 |
| 18 | TGME49_278910-t26_1-p1 | O-acetylserine (thiol) lyase 2, putative | 9 | 9 | 8 | 9 | 6 | 5 | 5 | 2 |
| 29 | TGME49_278940-t26_1-p1 | HECT-domain (ubiquitin-transferase) domain-containing protein | 10 | 10 | 9 | 6 | 8 | 3 | 3 | 2 |
| 18 | TGME49_279320.P3029 | hypothetical protein | 10 | 10 | 8 | 7 | 6 | 7 | 3 | 2 |
| 29 | TGME49_279380-t26_1-p1 | hypothetical protein | 8 | 8 | 8 | 8 | 7 | 3 | 3 | 2 |
| 29 | TGME49_280380-t26_1-p1 | poly(ADP-ribose) glycohydrolase | 9 | 9 | 9 | 8 | 6 | 3 | 3 | 2 |
| 18 | TGME49_280390.P3033 | kinetochore protein STU2, putative | 10 | 10 | 10 | 8 | 6 | 5 | 4 | 2 |
| 29 | TGME49_280435.P3038 | zinc finger protein, putative | 9 | 9 | 9 | 9 | 8 | 3 | 3 | 2 |
| 18 | TGME49_280540.P3043 | HEAT repeat-containing protein | 10 | 10 | 10 | 9 | 5 | 6 | 5 | 2 |
| 18 | TGME49_280560.P3045 | selenide, water dikinase | 10 | 10 | 9 | 6 | 5 | 6 | 5 | 2 |
| 18 | TGME49_280690.P3049 | DNA polymerase epsilon subunit B protein | 10 | 9 | 9 | 7 | 6 | 6 | 5 | 2 |
| 18 | TGME49_280770.P3055 | regulator of chromosome condensation (RCC1) repeat-containing protein | 10 | 10 | 10 | 8 | 7 | 6 | 4 | 2 |
| 18 | TGME49_281370-t26_1-p1 | polynucleotide adenylyltransferase | 10 | 9 | 9 | 8 | 7 | 5 | 5 | 2 |
| 29 | TGME49_281400.P3057 | phosphofructokinase domain-containing protein | 10 | 10 | 10 | 9 | 9 | 3 | 3 | 2 |
| 18 | TGME49_281545.P3064 | tRNA (adenine(58)-N(1))-methyltransferase non-catalytic subunit TRM6, putative | 9 | 9 | 8 | 7 | 5 | 5 | 4 | 2 |
| 18 | TGME49_282040-t26_1-p1 | hypothetical protein | 10 | 10 | 10 | 9 | 6 | 5 | 4 | 2 |
| 29 | TGME49_282150-t26_1-p1 | hypothetical protein | 9 | 9 | 8 | 8 | 6 | 4 | 4 | 2 |
| 18 | TGME49_282220-t26_1-p1 | AP2 domain transcription factor AP2VIIa-9 | 10 | 9 | 9 | 7 | 6 | 5 | 5 | 2 |
| 18 | TGME49_283820-t26_1-p1 | glycine cleavage T-protein (aminomethyl transferase) domain-containing protein | 10 | 9 | 9 | 7 | 6 | 5 | 3 | 2 |
| 29 | TGME49_284190-t26_1-p1 | pyruvate carboxylase | 10 | 10 | 10 | 9 | 9 | 3 | 3 | 2 |
| 29 | TGME49_284580.P3109 | ribose-phosphate diphosphokinase subfamily protein | 9 | 9 | 8 | 8 | 8 | 3 | 3 | 2 |
| 29 | TGME49_284620.P3111 | hypothetical protein | 10 | 10 | 10 | 9 | 6 | 3 | 5 | 2 |
| 18 | TGME49_284670.P3115 | flagellar associated protein | 9 | 8 | 8 | 8 | 6 | 6 | 5 | 2 |
| 18 | TGME49_285210.P3122 | centrosomal protein CEP120, putative | 10 | 10 | 8 | 6 | 6 | 5 | 4 | 2 |

| | | | | | | | | | | |
|---|---|---|---|---|---|---|---|---|---|---|
| 18 | TGME49_285220.P3123 | CAP-Gly domain-containing protein | 9 | 9 | 9 | 7 | 5 | 4 | 5 | 2 |
| 2 | TGME49_285240.P3125 | 3-oxo-5-alpha-steroid 4-dehydrogenase | 8 | 8 | 8 | 8 | 7 | 3 | 7 | 2 |
| 18 | TGME49_285250.P3126 | hypothetical protein | 10 | 10 | 10 | 6 | 6 | 5 | 5 | 2 |
| 18 | TGME49_285540-t26_1-p1 | DNA-directed DNA polymerase | 10 | 9 | 9 | 7 | 5 | 5 | 5 | 2 |
| 2 | TGME49_285690-t26_1-p1 | notch (dsl) domain-containing protein | 9 | 9 | 8 | 8 | 6 | 3 | 6 | 2 |
| 18 | TGME49_285810-t26_1-p1 | MYND finger domain-containing protein | 10 | 10 | 9 | 8 | 5 | 7 | 5 | 2 |
| 18 | TGME49_286140-t26_1-p1 | DNA replication complex GINS protein Psf2, putative | 10 | 9 | 9 | 8 | 5 | 6 | 5 | 2 |
| 2 | TGME49_286420-t26_1-p1 | elongation factor 1-alpha (EF-1-ALPHA), putative | 9 | 9 | 9 | 9 | 9 | 3 | 9 | 2 |
| 2 | TGME49_286440-t26_1-p1 | malic enzyme | 9 | 8 | 8 | 8 | 8 | 3 | 7 | 2 |
| 2 | TGME49_286620-t26_1-p1 | S1 RNA binding domain-containing protein | 10 | 10 | 10 | 10 | 8 | 5 | 6 | 2 |
| 2 | TGME49_286660.P3160 | kinesin, putative | 10 | 9 | 8 | 8 | 7 | 3 | 6 | 2 |
| 18 | TGME49_286800.P3165 | deubiquitinating enzyme MINDY, putative | 10 | 9 | 9 | 6 | 6 | 6 | 6 | 2 |
| 2 | TGME49_286920-t26_1-p1 | SWIRM domain-containing protein | 10 | 9 | 9 | 5 | 7 | 3 | 6 | 2 |
| 29 | TGME49_287480.P3174 | hypothetical protein | 10 | 10 | 10 | 9 | 7 | 5 | 4 | 2 |
| 29 | TGME49_287490.P3175 | hypothetical protein | 9 | 9 | 9 | 9 | 7 | 3 | 3 | 2 |
| 29 | TGME49_287960.P3176 | hypothetical protein | 9 | 9 | 8 | 8 | 7 | 3 | 3 | 2 |
| 18 | TGME49_287980-t26_1-p1 | FHA domain-containing protein | 10 | 9 | 9 | 6 | 6 | 6 | 6 | 2 |
| 18 | TGME49_288040-t26_1-p1 | hypothetical protein | 10 | 10 | 9 | 9 | 5 | 5 | 4 | 2 |
| 2 | TGME49_288540-t26_1-p1 | nucleoside transporter protein | 10 | 9 | 9 | 8 | 8 | 3 | 6 | 2 |
| 2 | TGME49_288550-t26_1-p1 | ankyrin repeat-containing protein | 9 | 8 | 8 | 7 | 7 | 4 | 6 | 2 |
| 29 | TGME49_289000-t26_1-p1 | hypothetical protein | 10 | 9 | 9 | 7 | 6 | 3 | 3 | 2 |
| 2 | TGME49_289380.P3202 | dense granule protein GRA39 | 10 | 10 | 9 | 7 | 6 | 4 | 6 | 2 |
| 18 | TGME49_289570-t26_1-p1 | phosphatidylinositol transfer protein | 10 | 10 | 10 | 6 | 6 | 6 | 5 | 2 |
| 2 | TGME49_289880-t26_1-p1 | hydrolase, putative | 10 | 10 | 10 | 10 | 8 | 5 | 8 | 2 |
| 2 | TGME49_289910-t26_1-p1 | haloacid dehalogenase-like hydrolase HAD2a | 9 | 9 | 8 | 8 | 7 | 3 | 6 | 2 |
| 18 | TGME49_289980-t26_1-p1 | HAD hydrolase, family IIA protein | 10 | 9 | 8 | 6 | 6 | 5 | 6 | 2 |
| 18 | TGME49_289990-t26_1-p1 | hypothetical protein | 10 | 9 | 8 | 7 | 6 | 5 | 5 | 2 |
| 29 | TGME49_290020-t26_1-p1 | cyclin dependent kinase binding protein | 10 | 9 | 8 | 7 | 7 | 3 | 3 | 2 |
| 18 | TGME49_290460-t26_1-p1 | hypothetical protein | 9 | 9 | 8 | 6 | 6 | 6 | 5 | 2 |
| 29 | TGME49_290620.P3227 | centrosomal protein CEP250L1 | 10 | 8 | 9 | 7 | 6 | 4 | 3 | 2 |
| 2 | TGME49_290860.P3236 | apicomplexan amino acid transporter ApiAT6-2 | 10 | 9 | 8 | 8 | 7 | 3 | 6 | 2 |
| 2 | TGME49_290870-t26_1-p1 | patched family protein | 10 | 10 | 10 | 10 | 8 | 3 | 8 | 2 |
| 18 | TGME49_291000-t26_1-p1 | PHD-finger domain-containing protein | 9 | 8 | 8 | 8 | 5 | 6 | 5 | 2 |
| 18 | TGME49_291030-t26_1-p1 | zinc finger, C3HC4 type (RING finger) domain-containing protein | 10 | 10 | 9 | 7 | 5 | 6 | 5 | 2 |
| 29 | TGME49_291670-t26_1-p1 | DEAD/DEAH box helicase domain-containing protein | 10 | 9 | 9 | 8 | 8 | 3 | 5 | 2 |
| 29 | TGME49_291950-t26_1-p1 | RNA recognition motif-containing protein | 10 | 9 | 9 | 8 | 8 | 3 | 3 | 2 |
| 18 | TGME49_291980.P3256 | HECT-domain (ubiquitin-transferase) domain-containing protein | 10 | 10 | 9 | 7 | 7 | 5 | 5 | 2 |
| 18 | TGME49_292000-t26_1-p1 | hypothetical protein | 10 | 9 | 9 | 8 | 5 | 5 | 4 | 2 |
| 18 | TGME49_292055.P3259 | calcium dependent protein kinase CDPK8 | 10 | 10 | 7 | 6 | 6 | 5 | 4 | 2 |
| 18 | TGME49_292200.P3262 | RNA recognition motif-containing protein | 10 | 9 | 9 | 7 | 5 | 6 | 5 | 2 |
| 18 | TGME49_292330-t26_1-p1 | TMEM121 domain-containing protein | 10 | 10 | 9 | 6 | 8 | 6 | 5 | 2 |
| 2 | TGME49_292935.P3268 | coiled-coil domain-containing protein | 10 | 9 | 9 | 9 | 9 | 5 | 6 | 2 |
| 29 | TGME49_292975-t26_1-p1 | hypothetical protein | 10 | 10 | 10 | 9 | 8 | 3 | 3 | 2 |
| 18 | TGME49_293060.P3273 | SPRY domain-containing protein | 10 | 9 | 9 | 6 | 6 | 5 | 5 | 2 |
| 2 | TGME49_293460.P3300 | ATP-dependent DNA ligase domain-containing protein | 10 | 9 | 10 | 9 | 9 | 3 | 6 | 2 |

| | | | | | | | | | | |
|---|---|---|---|---|---|---|---|---|---|---|
| 18 | TGME49_293630.P3314 | zinc finger, C2H2 type domain-containing protein | 10 | 10 | 9 | 9 | 8 | 5 | 5 | 2 |
| 18 | TGME49_293680.P3317 | hypothetical protein | 10 | 9 | 8 | 7 | 6 | 4 | 5 | 2 |
| 18 | TGME49_293770.P3326 | chitinase-like protein CLP1 | 10 | 9 | 9 | 6 | 7 | 5 | 4 | 2 |
| 2 | TGME49_294240.P3348 | hypothetical protein | 10 | 10 | 10 | 8 | 6 | 3 | 6 | 2 |
| 2 | TGME49_294600.P3361 | Toxoplasma gondii family D protein | 10 | 9 | 9 | 9 | 9 | 3 | 8 | 2 |
| 18 | TGME49_294700-t26_1-p1 | polynucleotide 5'-hydroxyl-kinase | 10 | 9 | 9 | 7 | 6 | 7 | 5 | 2 |
| 2 | TGME49_295010-t26_1-p1 | DEAD/DEAH box helicase domain-containing protein | 10 | 9 | 9 | 9 | 7 | 3 | 7 | 2 |
| 18 | TGME49_295420-t26_1-p1 | hypothetical protein | 10 | 9 | 9 | 7 | 6 | 5 | 5 | 2 |
| 29 | TGME49_295658-t26_1-p1 | zinc finger in N-recognin protein | 10 | 10 | 10 | 8 | 6 | 3 | 3 | 2 |
| 29 | TGME49_295720.P3388 | sulfite oxidase, putative | 10 | 8 | 9 | 9 | 7 | 3 | 3 | 2 |
| 18 | TGME49_295730.P3389 | tetratricopeptide repeat-containing protein | 9 | 9 | 8 | 7 | 5 | 5 | 5 | 2 |
| 18 | TGME49_295850.P3392 | apical annuli protein AAP2 | 10 | 10 | 9 | 7 | 5 | 5 | 5 | 2 |
| 18 | TGME49_297230-t26_1-p1 | vacuolar protein sorting-associated protein 53 | 10 | 10 | 9 | 6 | 5 | 6 | 6 | 2 |
| 29 | TGME49_297850-t26_1-p1 | Branched-chain-amino-acid aminotransferase | 9 | 9 | 9 | 8 | 6 | 3 | 5 | 2 |
| 29 | TGME49_297870.P3422 | inner membrane complex protein IMC36 | 10 | 10 | 9 | 7 | 7 | 3 | 3 | 2 |
| 18 | TGME49_299080.P3434 | vacuolar transporter chaperone VTC4 | 10 | 10 | 8 | 6 | 7 | 6 | 6 | 2 |
| 2 | TGME49_299110.P3435 | cleft lip and palate transmembrane protein 1 (clptm1) protein | 10 | 9 | 9 | 7 | 9 | 5 | 6 | 2 |
| 2 | TGME49_299210.P3438 | CTP synthase | 10 | 10 | 9 | 9 | 9 | 5 | 9 | 2 |
| 29 | TGME49_299230.P3439 | F-box domain-containing protein | 10 | 9 | 9 | 8 | 8 | 4 | 4 | 2 |
| 2 | TGME49_300020-t26_1-p1 | ATP-dependent metallopeptidase HflB subfamily protein | 10 | 10 | 9 | 9 | 9 | 5 | 7 | 2 |
| 18 | TGME49_300120.P3455 | aminotransferase, class V superfamily protein | 10 | 9 | 9 | 7 | 6 | 6 | 6 | 2 |
| 18 | TGME49_300330-t26_1-p1 | hypothetical protein | 10 | 10 | 9 | 7 | 6 | 6 | 3 | 2 |
| 18 | TGME49_300380-t26_1-p1 | endoplasmic reticulum oxidoreductin, putative | 10 | 9 | 9 | 8 | 6 | 5 | 5 | 2 |
| 29 | TGME49_300650-t26_1-p1 | iron-sulfur cluster assembly protein SufB | 8 | 3 | 9 | 9 | 8 | 3 | 3 | 2 |
| 29 | TGME49_300660-t26_1-p1 | RNA polymerase B | 9 | 3 | 10 | 10 | 10 | 3 | 3 | 2 |
| 29 | TGME49_300680-t26_1-p1 | RNA polymerase C1, putative | 8 | 3 | 9 | 9 | 9 | 3 | 3 | 2 |
| 29 | TGME49_300690-t26_1-p1 | RNA polymerase C2, putative | 9 | 3 | 9 | 9 | 9 | 3 | 3 | 2 |
| 29 | TGME49_302000-t26_1-p1 | chaperone clp, putative | 10 | 3 | 10 | 9 | 9 | 3 | 3 | 2 |
| 2 | TGME49_304470.P3481 | 3-ketosphinganine reductase, putative | 9 | 8 | 8 | 7 | 6 | 3 | 6 | 2 |
| 18 | TGME49_304630.P3484 | THO complex subunit 2, putative | 10 | 10 | 8 | 7 | 5 | 5 | 4 | 2 |
| 18 | TGME49_305140.P3496 | patatin-like phospholipase PL3 | 10 | 10 | 8 | 8 | 5 | 6 | 6 | 2 |
| 2 | TGME49_305180-t26_1-p1 | Na+/H+ exchanger NHE3 | 10 | 9 | 9 | 5 | 8 | 3 | 6 | 2 |
| 18 | TGME49_305310.P3500 | ERCC4 domain-containing protein | 10 | 9 | 9 | 6 | 7 | 6 | 4 | 2 |
| 18 | TGME49_305470-t26_1-p1 | hypothetical protein | 10 | 10 | 8 | 6 | 7 | 6 | 4 | 2 |
| 18 | TGME49_305510.P3505 | hypothetical protein | 9 | 9 | 8 | 8 | 6 | 6 | 4 | 2 |
| 18 | TGME49_305540-t26_1-p1 | DUF1769 domain-containing protein | 10 | 9 | 9 | 5 | 7 | 6 | 5 | 2 |
| 18 | TGME49_305560-t26_1-p1 | Vps51/Vps67 protein | 10 | 10 | 10 | 8 | 6 | 6 | 6 | 2 |
| 18 | TGME49_306000.P3512 | AP2 domain transcription factor AP2IX-8 | 10 | 10 | 10 | 6 | 5 | 6 | 6 | 2 |
| 29 | TGME49_306330-t26_1-p1 | phospholipase | 9 | 9 | 9 | 8 | 7 | 4 | 4 | 2 |
| 18 | TGME49_306430.P3522 | spindle assembly abnormal protein 6 | 10 | 9 | 9 | 6 | 7 | 5 | 4 | 2 |
| 29 | TGME49_306910-t26_1-p1 | hypothetical protein | 9 | 9 | 8 | 7 | 7 | 4 | 3 | 2 |
| 18 | TGME49_307040.P3535 | shikimate dehydrogenase substrate binding domain-containing protein | 10 | 10 | 9 | 8 | 6 | 5 | 5 | 2 |
| 29 | TGME49_307570-t26_1-p1 | glycerol-3-phosphate dehydrogenase (gpdh), putative | 9 | 10 | 9 | 9 | 8 | 5 | 3 | 2 |
| 2 | TGME49_307580-t26_1-p1 | CBS domain-containing protein | 10 | 9 | 9 | 6 | 8 | 3 | 6 | 2 |

| | | | | | | | | | | |
|---|---|---|---|---|---|---|---|---|---|---|
| 2 | TGME49_307640.P3537 | CMGC kinase, CK2 family | 8 | 8 | 8 | 8 | 6 | 3 | 6 | 2 |
| 18 | TGME49_307760-t26_1-p1 | Tubulin-tyrosine ligase family protein | 10 | 10 | 5 | 8 | 7 | 6 | 5 | 2 |
| 18 | TGME49_308000-t26_1-p1 | Gpi16 subunit, GPI transamidase component protein | 10 | 9 | 9 | 7 | 5 | 5 | 4 | 2 |
| 2 | TGME49_308030.P3541 | hypothetical protein | 10 | 8 | 8 | 8 | 7 | 4 | 6 | 2 |
| 29 | TGME49_308990-t26_1-p1 | transporter, solute:sodium symporter (SSS) family protein | 9 | 9 | 8 | 8 | 6 | 4 | 4 | 2 |
| 29 | TGME49_309030-t26_1-p1 | hypothetical protein | 10 | 10 | 9 | 9 | 7 | 3 | 3 | 2 |
| 18 | TGME49_309380-t26_1-p1 | kinetochore protein NUF2 | 10 | 9 | 9 | 8 | 6 | 6 | 4 | 2 |
| 18 | TGME49_309400.P3568 | RecF/RecN/SMC N terminal domain-containing protein | 10 | 10 | 10 | 9 | 5 | 5 | 5 | 2 |
| 18 | TGME49_309910.P3583 | amino acid transporter, putative | 10 | 10 | 8 | 9 | 6 | 5 | 6 | 2 |
| 29 | TGME49_309950.P3587 | NLE (NUC135) domain-containing protein | 10 | 9 | 9 | 8 | 7 | 4 | 4 | 2 |
| 18 | TGME49_310010.P3589 | rhoptry neck protein RON1 | 10 | 10 | 7 | 7 | 5 | 6 | 6 | 2 |
| 29 | TGME49_310150.P3598 | acyl-CoA synthetase ACS2, putative | 10 | 10 | 10 | 9 | 7 | 3 | 3 | 2 |
| 18 | TGME49_310190.P3601 | VPS15-like kinase | 10 | 7 | 10 | 8 | 5 | 5 | 4 | 2 |
| 18 | TGME49_310250-t26_1-p1 | hypothetical protein | 10 | 9 | 9 | 6 | 6 | 5 | 4 | 2 |
| 29 | TGME49_310290-t26_1-p1 | regulator of chromosome condensation (RCC1) repeat-containing protein | 10 | 9 | 9 | 8 | 8 | 3 | 5 | 2 |
| 18 | TGME49_310310.P3604 | WD domain, G-beta repeat-containing protein | 9 | 9 | 8 | 7 | 5 | 5 | 5 | 2 |
| 29 | TGME49_310370.P3606 | hypothetical protein | 9 | 9 | 8 | 8 | 6 | 4 | 3 | 2 |
| 18 | TGME49_310450.P3609 | IMC-associated protein IAP2 | 10 | 10 | 10 | 7 | 8 | 6 | 5 | 2 |
| 18 | TGME49_310530.P3614 | SNF2 family N-terminal domain-containing protein | 10 | 9 | 10 | 7 | 5 | 6 | 5 | 2 |
| 18 | TGME49_311030-t26_1-p1 | U3 small nucleolar RNA-associated protein 10, putative | 10 | 10 | 10 | 8 | 6 | 5 | 4 | 2 |
| 29 | TGME49_311080.P3633 | two pore calcium channel protein TPC | 10 | 9 | 8 | 8 | 8 | 5 | 3 | 2 |
| 18 | TGME49_311150-t26_1-p1 | SURF1 domain-containing protein | 9 | 9 | 8 | 8 | 5 | 6 | 5 | 2 |
| 18 | TGME49_311180.P3637 | hypothetical protein | 10 | 10 | 10 | 6 | 5 | 5 | 5 | 2 |
| 18 | TGME49_311230.P3639 | basal complex component BCC7 | 10 | 10 | 8 | 8 | 7 | 6 | 5 | 2 |
| 2 | TGME49_311310-t26_1-p1 | serine/threonine protein phosphatase 2B catalytic subunit A | 9 | 9 | 9 | 8 | 9 | 5 | 9 | 2 |
| 29 | TGME49_311410-t26_1-p1 | M phase phosphoprotein MPP10, putative | 10 | 9 | 9 | 8 | 7 | 3 | 3 | 2 |
| 18 | TGME49_311625.P3652 | bromodomain-containing protein BDP6, putative | 10 | 10 | 10 | 8 | 6 | 5 | 4 | 2 |
| 18 | TGME49_312065.P3661 | hypothetical protein | 10 | 9 | 9 | 6 | 5 | 5 | 5 | 2 |
| 2 | TGME49_312230-t26_1-p1 | DNA topoisomerase 2, putative | 10 | 10 | 10 | 9 | 9 | 3 | 10 | 2 |
| 18 | TGME49_312510.P3673 | ATP-dependent DNA helicase 2 subunit KU80 | 10 | 10 | 10 | 8 | 6 | 6 | 5 | 2 |
| 18 | TGME49_312590-t26_1-p1 | CLASP domain-containing protein | 10 | 8 | 8 | 7 | 6 | 5 | 5 | 2 |
| 29 | TGME49_312650.P3679 | TH1 domain-containing protein | 10 | 9 | 9 | 7 | 6 | 3 | 4 | 2 |
| 29 | TGME49_313000-t26_1-p1 | Toxoplasma gondii family D protein | 10 | 9 | 9 | 8 | 8 | 3 | 3 | 2 |
| 2 | TGME49_313150.P3692 | DUF89/Fructose bisphosphatase | 9 | 9 | 9 | 8 | 7 | 5 | 6 | 2 |
| 18 | TGME49_313280-t26_1-p1 | WD domain, G-beta repeat-containing protein | 10 | 9 | 8 | 8 | 5 | 5 | 4 | 2 |
| 18 | TGME49_313370.P3701 | leucine rich repeat-containing protein | 10 | 10 | 10 | 9 | 5 | 7 | 4 | 2 |
| 18 | TGME49_313430.P3705 | nucleoporin, putative | 10 | 10 | 10 | 8 | 5 | 5 | 4 | 2 |
| 29 | TGME49_313480.P3709 | apical annuli protein AAP3 | 10 | 10 | 9 | 8 | 8 | 3 | 5 | 2 |
| 2 | TGME49_313600-t26_1-p1 | DDHD domain-containing protein | 10 | 10 | 9 | 8 | 7 | 3 | 6 | 2 |
| 18 | TGME49_313690-t26_1-p1 | Sel1 repeat-containing protein | 10 | 10 | 9 | 8 | 6 | 5 | 3 | 2 |
| 18 | TGME49_313725.P3719 | RNA binding protein, putative | 10 | 9 | 9 | 8 | 4 | 7 | 4 | 2 |

| | | | | | | | | | | |
|---|---|---|---|---|---|---|---|---|---|---|
| 18 | TGME49_313860-t26_1-p1 | regulator of chromosome condensation (RCC1) repeat-containing protein | 10 | 10 | 9 | 6 | 6 | 6 | 3 | 2 |
| 18 | TGME49_314320-t26_1-p1 | GTP-binding protein, putative | 10 | 9 | 9 | 9 | 6 | 6 | 5 | 2 |
| 29 | TGME49_314330.P3738 | ABC transporter, ATP-binding domain-containing protein | 10 | 10 | 10 | 7 | 7 | 3 | 3 | 2 |
| 18 | TGME49_314875-t26_1-p1 | RIC1 protein | 10 | 10 | 8 | 6 | 7 | 5 | 5 | 2 |
| 18 | TGME49_315140-t26_1-p1 | WD domain, G-beta repeat-containing protein | 10 | 10 | 10 | 7 | 7 | 5 | 5 | 2 |
| 2 | TGME49_315260.P3760 | alanine dehydrogenase | 9 | 8 | 8 | 8 | 7 | 3 | 7 | 2 |
| 18 | TGME49_315360.P3764 | inner centromere protein INCENP2 | 10 | 10 | 9 | 8 | 5 | 6 | 5 | 2 |
| 29 | TGME49_315530.P3777 | vacuolar protein sorting-associated protein 39 | 10 | 10 | 10 | 10 | 9 | 5 | 4 | 2 |
| 18 | TGME49_315600.P3781 | MCM2/3/5 family protein | 10 | 10 | 10 | 7 | 6 | 6 | 3 | 2 |
| 18 | TGME49_315670.P3784 | serine/threonine-protein phosphatase PP2A subunit A | 10 | 9 | 9 | 6 | 5 | 7 | 6 | 2 |
| 29 | TGME49_315770.P3788 | cytochrome p450 | 10 | 9 | 9 | 9 | 8 | 3 | 3 | 2 |
| 18 | TGME49_315810-t26_1-p1 | tRNA nucleotidyltransferase/poly(A) polymerase family protein | 10 | 9 | 9 | 6 | 7 | 6 | 5 | 2 |
| 18 | TGME49_316200-t26_1-p1 | phosphoglycerate mutase family protein | 9 | 9 | 8 | 8 | 5 | 5 | 4 | 2 |
| 18 | TGME49_316230-t26_1-p1 | SAC1 phosphoinositide phosphatase, putative | 10 | 8 | 9 | 7 | 5 | 4 | 5 | 2 |
| 2 | TGME49_316270.P3803 | prenyltransferase GGT-2 | 8 | 8 | 8 | 8 | 7 | 3 | 7 | 2 |
| 29 | TGME49_316300-t26_1-p1 | ATP-dependent DNA helicase, RecQ family protein | 10 | 10 | 10 | 9 | 8 | 3 | 3 | 2 |
| 18 | TGME49_316480-t26_1-p1 | XRN 5'-3' exonuclease N-terminus protein | 10 | 9 | 9 | 6 | 5 | 7 | 5 | 2 |
| 18 | TGME49_316530.P3811 | apicomplexan kinetochore protein AKIT7, putative | 10 | 10 | 9 | 6 | 5 | 6 | 5 | 2 |
| 29 | TGME49_316560-t26_1-p1 | Toxoplasma gondii family D protein | 10 | 9 | 9 | 9 | 8 | 3 | 3 | 2 |
| 29 | TGME49_316680.P3817 | RNA pseudouridine synthase superfamily protein | 10 | 10 | 9 | 8 | 7 | 3 | 3 | 2 |
| 2 | TGME49_317720-t26_1-p1 | eukaryotic translation initiation factor 3 subunit 7, putative | 10 | 9 | 9 | 8 | 9 | 5 | 8 | 2 |
| 18 | TGME49_318140.P3820 | SART-1 family protein | 10 | 9 | 9 | 8 | 6 | 5 | 5 | 2 |
| 29 | TGME49_318190-t26_1-p1 | phosphoglycerate mutase family protein | 10 | 9 | 8 | 8 | 7 | 3 | 3 | 2 |
| 18 | TGME49_318400.P3829 | hypothetical protein | 10 | 10 | 10 | 7 | 7 | 5 | 4 | 2 |
| 2 | TGME49_318510.P3833 | N-ethylmaleimide-sensitive fusion protein, putative | 10 | 10 | 10 | 8 | 10 | 3 | 9 | 2 |
| 18 | TGME49_318525.P3834 | hypothetical protein | 10 | 10 | 9 | 8 | 5 | 5 | 5 | 2 |
| 29 | TGME49_318540.P3835 | calcium-binding egf domain-containing protein CBDP2 | 10 | 9 | 9 | 9 | 7 | 3 | 3 | 2 |
| 2 | TGME49_318660.P3841 | protein phosphatase PPM5B | 9 | 9 | 9 | 8 | 7 | 3 | 7 | 2 |
| 2 | TGME49_318730-t26_1-p1 | glycosyl transferase | 10 | 10 | 9 | 9 | 8 | 4 | 9 | 2 |
| 18 | TGME49_319610-t26_1-p1 | eIF2 kinase IF2K-D (incomplete catalytic triad) | 10 | 10 | 8 | 8 | 5 | 5 | 4 | 2 |
| 29 | TGME49_319620-t26_1-p1 | hypothetical protein | 10 | 10 | 9 | 9 | 8 | 3 | 3 | 2 |
| 18 | TGME49_319880-t26_1-p1 | MORN repeat-containing protein | 10 | 10 | 7 | 6 | 5 | 6 | 6 | 2 |
| 18 | TGME49_319900.P3861 | apical annuli protein AAP5 | 10 | 9 | 8 | 6 | 7 | 5 | 4 | 2 |
| 18 | TGME49_319930.P3863 | serine/threonine protein phosphatase 6 regulatory subunit PP6R | 10 | 9 | 8 | 7 | 6 | 6 | 4 | 2 |
| 29 | TGME49_319970.P3866 | subtilisin SUB10 | 10 | 9 | 9 | 8 | 7 | 3 | 4 | 2 |
| 18 | TGME49_320000-t26_1-p1 | SCY kinase (incomplete catalytic triad) | 10 | 9 | 9 | 7 | 8 | 5 | 5 | 2 |
| 2 | TGME49_320220-t26_1-p1 | cytochrome b-c1 complex subunit Rieske, putative | 9 | 9 | 9 | 8 | 8 | 5 | 7 | 2 |
| 29 | TGME49_320280.P3873 | hypothetical protein | 9 | 8 | 8 | 8 | 7 | 3 | 3 | 2 |

| | | | | | | | | | | |
|---|---|---|---|---|---|---|---|---|---|---|
| 18 | TGME49_320550.P3882 | AP-5 complex subunit beta-1, putative | 10 | 10 | 9 | 9 | 5 | 5 | 5 | 2 |
| 29 | TGME49_320592-t26_1-p1 | hypothetical protein | 10 | 10 | 10 | 8 | 8 | 5 | 4 | 2 |
| 29 | TGME49_320620-t26_1-p1 | queuine tRNA ribosyl transferase | 10 | 7 | 9 | 7 | 7 | 3 | 3 | 2 |
| 18 | TGME49_321560-t26_1-p1 | zinc knuckle domain-containing protein | 10 | 9 | 9 | 7 | 8 | 6 | 5 | 2 |
| 29 | TGME49_321660-t26_1-p1 | mannosyltransferase, putative | 10 | 9 | 9 | 8 | 6 | 3 | 3 | 2 |
| 18 | TGME49_321690-t26_1-p1 | hypothetical protein | 10 | 10 | 9 | 9 | 6 | 5 | 6 | 2 |
| 18 | TGME49_500057.P358 | EF-hand domain-containing protein | 10 | 10 | 8 | 6 | 5 | 7 | 3 | 2 |
| 29 | TGME49_500061.P362 | PAN domain-containing protein | 9 | 9 | 10 | 8 | 6 | 3 | 3 | 2 |
| 18 | TGME49_500098.P399 | regulator of chromosome condensation, putative | 10 | 10 | 10 | 9 | 5 | 5 | 5 | 2 |
| 29 | TGME49_500109.P410 | GPI transamidase component PIG-S, putative | 10 | 9 | 9 | 8 | 6 | 4 | 4 | 2 |
| 2 | TGME49_500125.P427 | mechanosensitive ion channel MscS, putative | 10 | 10 | 10 | 8 | 7 | 5 | 7 | 2 |
| 29 | TGME49_500127.P429 | hypothetical protein, conserved | 10 | 9 | 9 | 9 | 6 | 3 | 3 | 2 |
| 29 | TGME49_500168.P172 | cilia- and flagella-associated protein 54, putative | 10 | 10 | 10 | 9 | 7 | 4 | 4 | 2 |
| 29 | TGME49_500188.P192 | WD repeat-containing protein | 10 | 9 | 9 | 9 | 6 | 5 | 3 | 2 |
| 29 | TGME49_500224.P229 | GAF domain-containing protein | 9 | 7 | 9 | 9 | 6 | 3 | 3 | 2 |
| 29 | TGME49_500241.P247 | cysteine dioxygenase, putative | 10 | 9 | 9 | 7 | 6 | 3 | 3 | 2 |
| 18 | TGME49_500246.P252 | intraflagellar transport protein IFT88, putative | 10 | 8 | 9 | 8 | 5 | 5 | 5 | 2 |
| 18 | TGME49_500260.P49 | hypothetical protein, conserved | 10 | 8 | 9 | 7 | 6 | 5 | 3 | 2 |
| 2 | TGME49_500263.P52 | dynein assembly factor 3, putative | 9 | 7 | 8 | 8 | 6 | 4 | 6 | 2 |
| 29 | TGME49_500271.P60 | vacuolar protein 8, putative | 10 | 10 | 9 | 10 | 7 | 3 | 3 | 2 |
| 29 | TGME49_500283.P72 | ATP:guanido phosphotransferase, putative | 10 | 9 | 10 | 10 | 6 | 3 | 3 | 2 |
| 2 | TGME49_500303.P93 | PAN domain-containing protein | 10 | 10 | 10 | 9 | 9 | 4 | 9 | 2 |
| 18 | TGME49_500307.P97 | DEAD/DEAH box helicase, putative | 10 | 9 | 9 | 8 | 5 | 5 | 5 | 2 |
| 18 | TGME49_500341.P131 | hypothetical protein, conserved | 10 | 10 | 10 | 9 | 5 | 6 | 5 | 2 |
| 29 | TGME49_500342.P132 | 3'5'-cyclic nucleotide phosphodiesterase PDE3 | 10 | 10 | 10 | 8 | 7 | 3 | 3 | 2 |
| 29 | TGME49_500345.P135 | hypothetical protein, conserved | 9 | 7 | 9 | 9 | 7 | 3 | 3 | 2 |
| 29 | TGME49_500404.P303 | nucleoside diphosphate kinase, putative | 10 | 10 | 10 | 9 | 7 | 5 | 4 | 2 |
| 18 | TGME49_500435.P334 | HECT-domain (ubiquitin-transferase) -containing protein | 10 | 10 | 10 | 9 | 5 | 6 | 4 | 2 |

| Kmean_cluster | GeneID (all protein TG) | Product Description | Decile_bitHH | Decile_bitNcan | Decile_bitBB | Decilebit_CS | Decilebit_SN3 | Decile bit_Ef | Decile bit_CC | Meta_cluster |
|---|---|---|---|---|---|---|---|---|---|---|
| 21 | TGME49_200290-t26_1-p1 | rhomboid protease ROM1 | 8 | 7 | 8 | 8 | 7 | 5 | 7 | 3 |
| 17 | TGME49_200320.P452 | hypoxanthine-xanthine-guanine phosphoribosyl transferase HXGPRT | 3 | 7 | 7 | 6 | 3 | 6 | 5 | 3 |
| 21 | TGME49_200370.P456 | protein farnesyltransferase subunit beta, putative | 9 | 8 | 8 | 8 | 6 | 7 | 5 | 3 |
| 21 | TGME49_200460-t26_1-p1 | hypothetical protein | 9 | 9 | 8 | 7 | 7 | 5 | 6 | 3 |
| 30 | TGME49_201720.P470 | alpha N-terminal protein methyltransferase NTM1 | 8 | 8 | 7 | 5 | 7 | 3 | 3 | 3 |
| 1 | TGME49_201790.P474 | centrosomal protein CEP170, putative | 9 | 7 | 7 | 6 | 6 | 6 | 3 | 3 |
| 17 | TGME49_201880.P478 | cytochrome b-c1 complex subunit 9 | 6 | 6 | 6 | 5 | 5 | 5 | 5 | 3 |
| 1 | TGME49_202160-t26_1-p1 | ABC1 family protein | 10 | 7 | 7 | 6 | 6 | 6 | 5 | 3 |
| 17 | TGME49_202300-t26_1-p1 | inosine triphosphate pyrophosphatase, putative | 7 | 7 | 7 | 6 | 5 | 5 | 5 | 3 |
| 1 | TGME49_202470-t26_1-p1 | rRNA metabolism protein, SBDS family protein | 10 | 8 | 7 | 7 | 6 | 5 | 4 | 3 |
| 17 | TGME49_202690-t26_1-p1 | DNA-directed RNA polymerase II RPB9 | 7 | 6 | 6 | 5 | 5 | 5 | 5 | 3 |
| 21 | TGME49_202770.P500 | RNA recognition motif-containing protein | 9 | 9 | 9 | 6 | 6 | 7 | 6 | 3 |
| 1 | TGME49_202800-t26_1-p1 | cytochrome c oxidase assembly protein COX11 protein, putative | 8 | 7 | 7 | 6 | 5 | 6 | 4 | 3 |
| 1 | TGME49_202850-t26_1-p1 | ATP-binding domain-containing protein | 10 | 9 | 7 | 7 | 6 | 6 | 3 | 3 |
| 17 | TGME49_202870.P502 | SAP domain-containing protein | 8 | 7 | 6 | 5 | 5 | 5 | 5 | 3 |
| 1 | TGME49_202880.P503 | carrier superfamily protein | 9 | 6 | 7 | 6 | 5 | 5 | 4 | 3 |
| 21 | TGME49_203030-t26_1-p1 | N-methyl-D-aspartate receptor-associated protein | 8 | 7 | 8 | 7 | 6 | 4 | 6 | 3 |
| 21 | TGME49_203080.P510 | RNA recognition motif-containing protein | 8 | 8 | 8 | 7 | 7 | 7 | 7 | 3 |
| 30 | TGME49_203330-t26_1-p1 | hypothetical protein | 9 | 7 | 7 | 7 | 4 | 4 | 4 | 3 |
| 21 | TGME49_203350-t26_1-p1 | hypothetical protein | 9 | 8 | 8 | 7 | 6 | 6 | 6 | 3 |
| 1 | TGME49_203358.P519 | hypothetical protein | 9 | 8 | 8 | 6 | 7 | 6 | 3 | 3 |
| 21 | TGME49_203390.P520 | CRAL/TRIO domain-containing protein | 9 | 8 | 7 | 6 | 6 | 6 | 6 | 3 |
| 17 | TGME49_203450.P523 | brain colonization protein BCP1 | 3 | 7 | 7 | 4 | 5 | 6 | 6 | 3 |
| 17 | TGME49_203610-t26_1-p1 | 3'-5' exoribonuclease csl4, putative | 8 | 7 | 6 | 6 | 6 | 4 | 5 | 3 |
| 1 | TGME49_203620.P528 | ribosomal protein RPS35 | 9 | 7 | 7 | 7 | 6 | 5 | 4 | 3 |
| 30 | TGME49_203700-t26_1-p1 | SFT2 family protein | 8 | 7 | 6 | 6 | 5 | 4 | 4 | 3 |
| 1 | TGME49_203830.P534 | FHA domain-containing protein | 10 | 3 | 8 | 7 | 6 | 6 | 5 | 3 |
| 21 | TGME49_203870-t26_1-p1 | pre-mRNA-splicing factor ISY1, putative | 8 | 8 | 7 | 7 | 7 | 6 | 5 | 3 |
| 1 | TGME49_203970.P538 | dolichyl-diphosphooligosaccharide--protein glycosyltransferase | 10 | 9 | 8 | 5 | 6 | 6 | 4 | 3 |
| 1 | TGME49_204060.P541 | syntaxin-12 | 9 | 8 | 8 | 6 | 7 | 6 | 4 | 3 |
| 1 | TGME49_204070-t26_1-p1 | YT521-B family protein | 8 | 8 | 7 | 6 | 5 | 5 | 4 | 3 |
| 30 | TGME49_204350-t26_1-p1 | GTP-binding protein, putative | 10 | 8 | 7 | 6 | 6 | 4 | 3 | 3 |
| 17 | TGME49_204390.P550 | hypothetical protein | 8 | 7 | 7 | 6 | 5 | 5 | 5 | 3 |
| 1 | TGME49_204540-t26_1-p1 | DUF367 domain-containing protein | 9 | 8 | 7 | 6 | 6 | 5 | 5 | 3 |
| 21 | TGME49_205180-t26_1-p1 | RNA recognition motif-containing protein | 8 | 8 | 7 | 7 | 7 | 6 | 6 | 3 |
| 1 | TGME49_205330-t26_1-p1 | hypothetical protein | 8 | 8 | 8 | 7 | 6 | 5 | 3 | 3 |
| 17 | TGME49_205340-t26_1-p1 | ribosomal protein RPS12 | 6 | 6 | 6 | 6 | 6 | 6 | 5 | 3 |
| 30 | TGME49_205410-t26_1-p1 | RNA pseudouridine synthase superfamily protein | 10 | 8 | 7 | 6 | 5 | 3 | 4 | 3 |
| 30 | TGME49_205550-t26_1-p1 | AGC kinase | 9 | 9 | 6 | 6 | 6 | 3 | 4 | 3 |
| 17 | TGME49_205558-t26_1-p1 | NAC domain-containing protein | 7 | 7 | 6 | 6 | 5 | 5 | 5 | 3 |

| | | | | | | | | | | |
|---|---|---|---|---|---|---|---|---|---|---|
| 21 | TGME49_205670.P578 | striated fiber assemblin SFA2 | 8 | 8 | 8 | 8 | 7 | 7 | 6 | 3 |
| 30 | TGME49_205690.P579 | hypothetical protein | 9 | 8 | 8 | 7 | 6 | 3 | 5 | 3 |
| 21 | TGME49_205720-t26_1-p1 | Adenosine/AMP deaminase domain-containing protein | 9 | 8 | 8 | 7 | 6 | 7 | 6 | 3 |
| 17 | TGME49_205730-t26_1-p1 | hypothetical protein | 6 | 6 | 6 | 6 | 6 | 5 | 3 | 3 |
| 1 | TGME49_206310-t26_1-p1 | prenyltransferase, UbiA family protein | 10 | 8 | 8 | 7 | 5 | 6 | 3 | 3 |
| 21 | TGME49_206350-t26_1-p1 | ER-retrevial receptor Ter1p, putative | 7 | 7 | 7 | 7 | 7 | 7 | 5 | 3 |
| 1 | TGME49_206390-t26_1-p1 | checkpoint protein HUS1, putative | 8 | 8 | 8 | 5 | 6 | 6 | 4 | 3 |
| 1 | TGME49_206450.P585 | autophagy-related cysteine peptidase atg4, putative | 10 | 9 | 7 | 6 | 5 | 5 | 4 | 3 |
| 30 | TGME49_206500.P587 | hypothetical protein | 9 | 8 | 7 | 6 | 5 | 4 | 5 | 3 |
| 17 | TGME49_206560.P591 | EF hand domain-containing protein | 7 | 7 | 6 | 6 | 6 | 5 | 4 | 3 |
| 30 | TGME49_206600-t26_1-p1 | sigma-70, region 3 protein | 10 | 7 | 7 | 5 | 5 | 4 | 3 | 3 |
| 21 | TGME49_206620.P595 | OST-HTH domain-containing protein | 10 | 3 | 9 | 8 | 7 | 7 | 8 | 3 |
| 21 | TGME49_206690.P597 | glideosome-associated protein with multiple-membrane spans GAPM2B | 9 | 9 | 8 | 7 | 7 | 5 | 6 | 3 |
| 17 | TGME49_206700-t26_1-p1 | hypothetical protein | 8 | 7 | 7 | 6 | 5 | 5 | 5 | 3 |
| 1 | TGME49_207100-t26_1-p1 | hypothetical protein | 10 | 8 | 8 | 6 | 6 | 6 | 5 | 3 |
| 17 | TGME49_207680.P624 | suppressor of kinetochore protein 1 | 7 | 7 | 7 | 6 | 6 | 5 | 5 | 3 |
| 17 | TGME49_207710-t26_1-p1 | phosphatidylinositol synthase | 7 | 7 | 6 | 5 | 5 | 5 | 5 | 3 |
| 21 | TGME49_207770-t26_1-p1 | PCI domain-containing protein | 9 | 9 | 8 | 6 | 7 | 6 | 6 | 3 |
| 21 | TGME49_207830-t26_1-p1 | MORN repeat-containing protein | 9 | 8 | 8 | 7 | 7 | 7 | 7 | 3 |
| 17 | TGME49_207840.P628 | ribosomal protein RPS17 | 6 | 6 | 6 | 6 | 6 | 4 | 5 | 3 |
| 1 | TGME49_207940-t26_1-p1 | ribosomal protein S9, putative | 8 | 6 | 7 | 6 | 6 | 6 | 3 | 3 |
| 30 | TGME49_208030-t26_1-p1 | microneme protein MIC4 | 9 | 8 | 7 | 6 | 5 | 3 | 3 | 3 |
| 21 | TGME49_208070-t26_1-p1 | inositol polyphosphate kinase | 10 | 9 | 5 | 7 | 7 | 7 | 6 | 3 |
| 30 | TGME49_208200-t26_1-p1 | PHD-finger domain-containing protein | 9 | 8 | 7 | 6 | 5 | 4 | 4 | 3 |
| 30 | TGME49_208310-t26_1-p1 | FHA domain-containing protein | 9 | 8 | 8 | 6 | 5 | 4 | 3 | 3 |
| 1 | TGME49_208390.P640 | F-actin-capping protein subunit alpha, putative | 9 | 8 | 7 | 7 | 5 | 5 | 4 | 3 |
| 1 | TGME49_208400-t26_1-p1 | hypothetical protein | 10 | 7 | 6 | 6 | 5 | 5 | 5 | 3 |
| 21 | TGME49_208500.P646 | protein phosphatase PPM4 | 9 | 9 | 8 | 6 | 8 | 7 | 5 | 3 |
| 1 | TGME49_208560.P648 | carrier superfamily protein | 9 | 8 | 8 | 7 | 5 | 5 | 5 | 3 |
| 30 | TGME49_208710-t26_1-p1 | DNA/RNA non-specific endonuclease | 9 | 8 | 7 | 5 | 5 | 3 | 3 | 3 |
| 1 | TGME49_208718-t26_1-p1 | protein tyrosine phosphatase type IVA A, putative | 9 | 7 | 6 | 6 | 6 | 5 | 5 | 3 |
| 30 | TGME49_208730.P651 | microneme adhesive repeat protein MCP4 | 10 | 9 | 7 | 7 | 5 | 3 | 4 | 3 |
| 30 | TGME49_208750.P653 | RWD domain-containing protein | 8 | 7 | 7 | 6 | 5 | 3 | 4 | 3 |
| 21 | TGME49_208780.P655 | ubiquitin-conjugating enzyme subfamily protein | 7 | 7 | 7 | 6 | 7 | 6 | 6 | 3 |
| 1 | TGME49_209010.P662 | elongation factor TS protein | 9 | 8 | 7 | 5 | 5 | 5 | 3 | 3 |
| 17 | TGME49_209080-t26_1-p1 | transport protein particle (trapp) component, bet3 protein | 7 | 6 | 6 | 5 | 5 | 3 | 5 | 3 |
| 17 | TGME49_209140.P671 | anti-silencing protein, ASF1 family protein | 8 | 5 | 7 | 6 | 5 | 6 | 6 | 3 |
| 30 | TGME49_209240-t26_1-p1 | RNA methyltransferase | 9 | 8 | 8 | 7 | 5 | 4 | 4 | 3 |
| 1 | TGME49_209280.P675 | FAM76 domain-containing protein | 10 | 9 | 4 | 7 | 7 | 6 | 3 | 3 |
| 21 | TGME49_209430-t26_1-p1 | 40S ribosomal protein S4, putative | 7 | 7 | 7 | 7 | 6 | 6 | 6 | 3 |
| 17 | TGME49_209570-t26_1-p1 | suppressor of mitotic defects protein | 8 | 8 | 7 | 6 | 5 | 5 | 6 | 3 |
| 21 | TGME49_209580-t26_1-p1 | RNA methylase family UPF0020 protein | 10 | 7 | 9 | 6 | 8 | 7 | 5 | 3 |
| 30 | TGME49_209680.P690 | hypothetical protein | 9 | 7 | 6 | 6 | 5 | 4 | 4 | 3 |

| | | | | | | | | | | |
|---|---|---|---|---|---|---|---|---|---|---|
| 1 | TGME49_209710-t26_1-p1 | ribosomal protein RPL28 | 8 | 7 | 6 | 6 | 5 | 5 | 4 | 3 |
| 1 | TGME49_209720-t26_1-p1 | DUF4149 domain-containing protein | 7 | 7 | 7 | 7 | 5 | 5 | 4 | 3 |
| 30 | TGME49_209810-t26_1-p1 | hypothetical protein | 8 | 8 | 8 | 7 | 7 | 3 | 4 | 3 |
| 1 | TGME49_209820.P694 | syntaxin 1 | 8 | 7 | 8 | 7 | 5 | 6 | 4 | 3 |
| 30 | TGME49_209830.P695 | leucine-rich repeat-containing protein LRRC45, putative | 9 | 9 | 8 | 6 | 6 | 4 | 3 | 3 |
| 17 | TGME49_209860.P696 | SRP40, C-terminal domain-containing protein | 7 | 6 | 6 | 5 | 5 | 5 | 4 | 3 |
| 1 | TGME49_209870.P697 | HAD hydrolase, family IIA protein | 9 | 8 | 7 | 7 | 5 | 6 | 5 | 3 |
| 21 | TGME49_209890.P698 | hypothetical protein | 7 | 6 | 7 | 7 | 7 | 7 | 6 | 3 |
| 17 | TGME49_209910-t26_1-p1 | histone H2Bv | 6 | 6 | 6 | 6 | 3 | 4 | 6 | 3 |
| 30 | TGME49_210390-t26_1-p1 | WD domain, G-beta repeat-containing protein | 9 | 8 | 8 | 6 | 5 | 3 | 6 | 3 |
| 17 | TGME49_210412-t26_1-p1 | hypothetical protein | 7 | 7 | 7 | 5 | 6 | 3 | 5 | 3 |
| 30 | TGME49_210760.P716 | glutamine amidotransferase-related, putative | 9 | 8 | 8 | 7 | 5 | 3 | 3 | 3 |
| 21 | TGME49_210778.P717 | hemimethylated DNA binding domain-containing protein | 10 | 3 | 9 | 7 | 7 | 8 | 5 | 3 |
| 1 | TGME49_210800.P718 | activator of hsp90 ATPase, putative | 9 | 8 | 8 | 7 | 7 | 5 | 4 | 3 |
| 30 | TGME49_211020.P723 | polyadenylate-binding protein PABPN | 8 | 7 | 6 | 6 | 6 | 3 | 3 | 3 |
| 30 | TGME49_211030-t26_1-p1 | hypothetical protein | 7 | 7 | 7 | 6 | 5 | 3 | 3 | 3 |
| 1 | TGME49_211220.P727 | origin recognition complex subunit 4, putative | 9 | 7 | 7 | 6 | 5 | 5 | 5 | 3 |
| 30 | TGME49_211400.P737 | DEAD/DEAH box helicase domain-containing protein | 10 | 8 | 7 | 5 | 6 | 4 | 4 | 3 |
| 1 | TGME49_211600.P744 | hypothetical protein | 10 | 9 | 7 | 6 | 5 | 5 | 5 | 3 |
| 1 | TGME49_211700-t26_1-p1 | SAC3 domain-containing protein | 10 | 9 | 6 | 6 | 7 | 8 | 4 | 3 |
| 17 | TGME49_211850.P749 | hypothetical protein | 7 | 6 | 6 | 6 | 5 | 5 | 5 | 3 |
| 30 | TGME49_212110-t26_1-p1 | histone lysine demethylase JMJD5 | 10 | 8 | 7 | 6 | 5 | 4 | 4 | 3 |
| 1 | TGME49_212140-t26_1-p1 | hypothetical protein | 10 | 8 | 6 | 6 | 6 | 6 | 5 | 3 |
| 17 | TGME49_212290.P763 | ribosomal protein RPS19 | 7 | 7 | 7 | 6 | 6 | 6 | 5 | 3 |
| 30 | TGME49_212300.P764 | dense granule protein GRA32 | 10 | 9 | 7 | 7 | 6 | 4 | 4 | 3 |
| 17 | TGME49_212310.P765 | vacuolar ATP synthetase | 6 | 6 | 6 | 6 | 6 | 5 | 6 | 3 |
| 17 | TGME49_212725-t26_1-p1 | hypothetical protein | 7 | 6 | 6 | 6 | 5 | 4 | 4 | 3 |
| 30 | TGME49_212840-t26_1-p1 | HIT zinc finger protein | 9 | 8 | 8 | 7 | 6 | 4 | 4 | 3 |
| 17 | TGME49_212930.P769 | NifU family domain-containing protein | 8 | 6 | 7 | 6 | 6 | 5 | 5 | 3 |
| 17 | TGME49_212950.P772 | ADP-ribosylation factor family protein 2, putative | 7 | 6 | 7 | 7 | 6 | 4 | 6 | 3 |
| 1 | TGME49_212960.P773 | pre-mRNA-splicing regulator WTAP | 9 | 8 | 7 | 7 | 6 | 5 | 4 | 3 |
| 17 | TGME49_213069.P781 | ATP synthase mitochondrial F1 complex assembly factor 1, putative | 5 | 7 | 6 | 5 | 4 | 5 | 5 | 3 |
| 17 | TGME49_213350.P788 | ribosomal protein RPS15 | 3 | 6 | 6 | 3 | 6 | 6 | 6 | 3 |
| 1 | TGME49_213360.P789 | leucine rich repeat-containing protein | 9 | 7 | 8 | 7 | 5 | 5 | 4 | 3 |
| 1 | TGME49_213420.P791 | RAP domain-containing protein | 9 | 8 | 8 | 7 | 6 | 6 | 3 | 3 |
| 30 | TGME49_213600.P798 | hypothetical protein | 10 | 8 | 6 | 7 | 6 | 4 | 3 | 3 |
| 30 | TGME49_213670-t26_1-p1 | MIC60 domain-containing protein | 10 | 5 | 8 | 7 | 5 | 4 | 3 | 3 |
| 30 | TGME49_213780-t26_1-p1 | radical SAM domain-containing protein | 9 | 8 | 7 | 5 | 6 | 3 | 3 | 3 |
| 21 | TGME49_213800.P808 | protein phosphatase 2b regulatory subunit, putative | 7 | 7 | 7 | 6 | 7 | 7 | 7 | 3 |
| 17 | TGME49_213830-t26_1-p1 | Ufm1-conjugating enzyme 1 | 7 | 6 | 6 | 5 | 5 | 5 | 6 | 3 |
| 1 | TGME49_213910-t26_1-p1 | hypothetical protein | 8 | 8 | 8 | 7 | 7 | 6 | 4 | 3 |
| 1 | TGME49_213930-t26_1-p1 | exosome complex component RRP45, putative | 9 | 8 | 8 | 7 | 7 | 5 | 3 | 3 |

| | | | | | | | | | | |
|---|---|---|---|---|---|---|---|---|---|---|
| 17 | TGME49_214150-t26_1-p1 | mitochondrial import inner membrane translocase subunit TIM23 | 7 | 6 | 6 | 5 | 5 | 5 | 5 | 3 |
| 21 | TGME49_214180.P818 | epsin-like protein EPSL | 9 | 8 | 8 | 7 | 7 | 6 | 6 | 3 |
| 1 | TGME49_214290-t26_1-p1 | DJ-1 family protein | 8 | 7 | 7 | 6 | 6 | 5 | 5 | 3 |
| 1 | TGME49_214500-t26_1-p1 | ankyrin repeat-containing protein | 9 | 7 | 7 | 6 | 5 | 5 | 3 | 3 |
| 1 | TGME49_214590-t26_1-p1 | micro-fibrillar-associated protein 1 | 9 | 7 | 8 | 7 | 8 | 6 | 3 | 3 |
| 21 | TGME49_214620.P830 | prolyl hydroxylase PHYb | 9 | 8 | 8 | 7 | 7 | 7 | 6 | 3 |
| 21 | TGME49_214760-t26_1-p1 | erythronate-4-phosphate dehydrogenase domain-containing protein | 9 | 8 | 8 | 7 | 5 | 6 | 6 | 3 |
| 21 | TGME49_214770-t26_1-p1 | ras-related protein RAB1B | 7 | 7 | 7 | 7 | 7 | 5 | 7 | 3 |
| 21 | TGME49_214790.P832 | glycoprotein | 8 | 8 | 7 | 7 | 6 | 6 | 6 | 3 |
| 21 | TGME49_215020-t26_1-p1 | hypothetical protein | 9 | 8 | 8 | 6 | 7 | 6 | 6 | 3 |
| 21 | TGME49_215090-t26_1-p1 | ATP binding protein, putative | 8 | 7 | 8 | 8 | 7 | 7 | 6 | 3 |
| 17 | TGME49_215130-t26_1-p1 | adaptor-related protein complex 3, sigma 2 subunit | 6 | 6 | 6 | 6 | 6 | 6 | 6 | 3 |
| 30 | TGME49_215160.P846 | hypothetical protein | 8 | 7 | 7 | 7 | 5 | 3 | 3 | 3 |
| 17 | TGME49_215320.P851 | ODAD3 domain-containing protein | 8 | 3 | 6 | 7 | 5 | 6 | 6 | 3 |
| 30 | TGME49_215400-t26_1-p1 | RNA recognition motif-containing protein | 9 | 8 | 7 | 6 | 5 | 3 | 5 | 3 |
| 17 | TGME49_215420.P854 | synaptobrevin homolog YKT6, putative | 8 | 7 | 7 | 7 | 5 | 5 | 6 | 3 |
| 21 | TGME49_215430.P855 | mitochondrial marker protein MYS | 8 | 8 | 7 | 6 | 7 | 6 | 5 | 3 |
| 21 | TGME49_215450.P856 | aquaporin 1 | 8 | 8 | 7 | 7 | 7 | 6 | 5 | 3 |
| 17 | TGME49_215460.P857 | ribosomal protein RPS24 | 6 | 6 | 6 | 6 | 6 | 5 | 5 | 3 |
| 21 | TGME49_215470-t26_1-p1 | ribosomal protein RPL10A | 7 | 7 | 7 | 7 | 7 | 7 | 6 | 3 |
| 30 | TGME49_215550-t26_1-p1 | exonuclease V, putative | 10 | 8 | 8 | 6 | 7 | 4 | 4 | 3 |
| 1 | TGME49_215730-t26_1-p1 | Gas41, putative | 9 | 8 | 7 | 6 | 6 | 6 | 3 | 3 |
| 17 | TGME49_215920.P868 | hypothetical protein | 6 | 7 | 7 | 6 | 6 | 5 | 4 | 3 |
| 1 | TGME49_216010-t26_1-p1 | ribosomal protein RPL24 | 9 | 7 | 7 | 6 | 5 | 6 | 5 | 3 |
| 1 | TGME49_216030.P873 | VHS domain-containing protein | 9 | 7 | 7 | 7 | 6 | 5 | 4 | 3 |
| 21 | TGME49_216040.P874 | 30S ribosomal protein S15, putative | 6 | 8 | 8 | 8 | 6 | 7 | 7 | 3 |
| 30 | TGME49_216050-t26_1-p1 | tetratricopeptide repeat-containing protein | 9 | 8 | 8 | 6 | 5 | 4 | 3 | 3 |
| 21 | TGME49_216130-t26_1-p1 | ubiquitin conjugating enzyme E2, putative | 7 | 7 | 7 | 6 | 7 | 6 | 6 | 3 |
| 17 | TGME49_216410.P884 | hypothetical protein | 7 | 7 | 7 | 6 | 6 | 6 | 5 | 3 |
| 21 | TGME49_216450-t26_1-p1 | proteasome subunit alpha type-3, putative | 8 | 8 | 8 | 7 | 7 | 6 | 6 | 3 |
| 30 | TGME49_216530-t26_1-p1 | ribosome recycling factor protein | 7 | 7 | 7 | 6 | 5 | 4 | 4 | 3 |
| 1 | TGME49_216660-t26_1-p1 | hypothetical protein | 9 | 8 | 7 | 7 | 5 | 5 | 5 | 3 |
| 1 | TGME49_216680-t26_1-p1 | serine/threonine protein phosphatase 6 structural subunit PP6S | 9 | 8 | 8 | 7 | 5 | 6 | 4 | 3 |
| 30 | TGME49_216740-t26_1-p1 | riboflavin kinase | 10 | 8 | 6 | 5 | 5 | 4 | 4 | 3 |
| 30 | TGME49_216750-t26_1-p1 | Paf1/RNA polymerase II complex component LEO1 | 8 | 6 | 7 | 6 | 6 | 3 | 4 | 3 |
| 17 | TGME49_216800-t26_1-p1 | flagellar/basal body protein | 7 | 3 | 7 | 3 | 7 | 6 | 5 | 3 |
| 30 | TGME49_216920-t26_1-p1 | mediator complex subunit MED8 | 7 | 7 | 7 | 6 | 6 | 4 | 4 | 3 |
| 21 | TGME49_216930-t26_1-p1 | cholinephosphate cytidylyltransferase | 8 | 8 | 8 | 6 | 7 | 7 | 7 | 3 |
| 30 | TGME49_216965.P897 | leucine rich repeat-containing protein | 8 | 7 | 7 | 6 | 5 | 4 | 4 | 3 |
| 1 | TGME49_217010-t26_1-p1 | CD2BP2-like protein | 9 | 8 | 7 | 6 | 5 | 5 | 4 | 3 |
| 30 | TGME49_217440-t26_1-p1 | HCNGP family protein | 9 | 8 | 7 | 6 | 5 | 3 | 5 | 3 |
| 1 | TGME49_217540.P909 | serine/arginine-rich splicing factor SR2 | 8 | 8 | 7 | 7 | 6 | 6 | 5 | 3 |
| 17 | TGME49_217570.P911 | ribosomal protein RPS27 | 5 | 5 | 5 | 5 | 5 | 5 | 5 | 3 |
| 21 | TGME49_217580-t26_1-p1 | DNA-directed RNA polymerase III RPC6 | 8 | 8 | 8 | 8 | 7 | 5 | 5 | 3 |
| 21 | TGME49_217710.P922 | DnaJ domain-containing protein | 8 | 8 | 8 | 8 | 7 | 5 | 6 | 3 |

| | | | | | | | | | | |
|---|---|---|---|---|---|---|---|---|---|---|
| 21 | TGME49_217760.P924 | GTP-binding protein | 8 | 8 | 8 | 7 | 7 | 6 | 5 | 3 |
| 1 | TGME49_217780-t26_1-p1 | Sec20 protein | 9 | 8 | 6 | 5 | 5 | 5 | 4 | 3 |
| 30 | TGME49_217870.P927 | palmitoyltransferase DHHC3 | 7 | 8 | 7 | 7 | 6 | 3 | 3 | 3 |
| 17 | TGME49_217900-t26_1-p1 | hypothetical protein | 7 | 5 | 6 | 6 | 3 | 5 | 5 | 3 |
| 1 | TGME49_218250.P934 | TAP42 family protein | 9 | 7 | 8 | 7 | 7 | 5 | 5 | 3 |
| 17 | TGME49_218260-t26_1-p1 | histone H3.3 | 6 | 6 | 6 | 6 | 6 | 3 | 6 | 3 |
| 1 | TGME49_218510-t26_1-p1 | hypothetical protein | 9 | 7 | 7 | 6 | 4 | 7 | 3 | 3 |
| 17 | TGME49_218530.P944 | proteasome-interacting thioredoxin domain-containing protein | 7 | 5 | 6 | 6 | 5 | 5 | 5 | 3 |
| 1 | TGME49_218600-t26_1-p1 | RNA recognition motif-containing protein | 10 | 7 | 8 | 7 | 6 | 5 | 4 | 3 |
| 21 | TGME49_218760.P950 | alpha-soluble NSF attachment protein | 8 | 8 | 8 | 8 | 6 | 6 | 6 | 3 |
| 17 | TGME49_218820-t26_1-p1 | alba 2 | 6 | 6 | 6 | 6 | 5 | 5 | 5 | 3 |
| 1 | TGME49_218870-t26_1-p1 | TBC domain-containing protein TBC3 | 9 | 8 | 8 | 6 | 7 | 7 | 4 | 3 |
| 17 | TGME49_218940.P957 | ATP synthase-associated protein | 7 | 6 | 6 | 6 | 5 | 5 | 3 | 3 |
| 1 | TGME49_219080-t26_1-p1 | Golgi to ER traffic protein GET4, putative | 8 | 8 | 7 | 6 | 6 | 6 | 4 | 3 |
| 17 | TGME49_219100.P960 | cyclin-dependent kinase regulatory subunit protein | 7 | 8 | 7 | 6 | 6 | 5 | 5 | 3 |
| 17 | TGME49_219140-t26_1-p1 | EF-1 guanine nucleotide exchange domain-containing protein | 7 | 6 | 6 | 6 | 5 | 5 | 5 | 3 |
| 30 | TGME49_219190.P964 | nuclear movement protein domain containing protein | 7 | 7 | 6 | 6 | 6 | 3 | 3 | 3 |
| 1 | TGME49_219290-t26_1-p1 | F-actin-capping protein subunit beta, putative | 8 | 8 | 7 | 6 | 5 | 5 | 4 | 3 |
| 17 | TGME49_219300-t26_1-p1 | ran binding protein | 7 | 6 | 6 | 6 | 5 | 5 | 3 | 3 |
| 30 | TGME49_219720-t26_1-p1 | ras-related protein RAB5C | 8 | 7 | 7 | 7 | 5 | 3 | 6 | 3 |
| 30 | TGME49_219742.P982 | hypothetical protein | 9 | 8 | 6 | 6 | 5 | 3 | 3 | 3 |
| 17 | TGME49_219750-t26_1-p1 | cytochrome c, putative | 6 | 6 | 6 | 6 | 6 | 5 | 5 | 3 |
| 21 | TGME49_219760-t26_1-p1 | N-acetyltransferase family protein | 7 | 7 | 7 | 6 | 6 | 6 | 6 | 3 |
| 17 | TGME49_220150-t26_1-p1 | 50S ribosomal protein L16, putative | 7 | 7 | 6 | 6 | 5 | 5 | 5 | 3 |
| 1 | TGME49_220160.P989 | ATG18/WIPI-like protein PROP2 | 9 | 8 | 7 | 5 | 5 | 6 | 4 | 3 |
| 21 | TGME49_220200-t26_1-p1 | hypothetical protein | 9 | 7 | 7 | 6 | 6 | 6 | 6 | 3 |
| 1 | TGME49_220260-t26_1-p1 | hypothetical protein | 9 | 8 | 8 | 6 | 6 | 5 | 4 | 3 |
| 21 | TGME49_220300.P995 | ribosomal protein L15 protein | 9 | 9 | 8 | 7 | 7 | 6 | 6 | 3 |
| 30 | TGME49_220390-t26_1-p1 | hypothetical protein | 8 | 8 | 7 | 7 | 6 | 4 | 5 | 3 |
| 17 | TGME49_220400.P997 | actin depolymerizing factor ADF | 6 | 6 | 6 | 5 | 5 | 5 | 5 | 3 |
| 1 | TGME49_220490-t26_1-p1 | pre-rRNA-processing protein PNO1, putative | 8 | 7 | 7 | 7 | 6 | 6 | 5 | 3 |
| 30 | TGME49_220500-t26_1-p1 | UBX domain-containing protein | 8 | 7 | 7 | 7 | 7 | 4 | 3 | 3 |
| 17 | TGME49_220880-t26_1-p1 | dynactin subunit 3, putative | 7 | 7 | 7 | 6 | 5 | 5 | 5 | 3 |
| 21 | TGME49_220940.P1009 | ribosomal RNA methyltransferase (FtsJ) family protein | 9 | 8 | 8 | 7 | 7 | 7 | 7 | 3 |
| 1 | TGME49_221180.P1011 | hypothetical protein | 9 | 7 | 6 | 6 | 5 | 5 | 5 | 3 |
| 1 | TGME49_221190-t26_1-p1 | mrna cleavage factor family protein, putative | 9 | 7 | 7 | 5 | 6 | 6 | 5 | 3 |
| 17 | TGME49_221210.P1012 | cyclophilin | 7 | 6 | 6 | 6 | 3 | 5 | 5 | 3 |
| 30 | TGME49_221230.P1014 | protein tyrosine phosphatase, putative | 9 | 9 | 8 | 7 | 6 | 4 | 4 | 3 |
| 1 | TGME49_221270-t26_1-p1 | oxidoreductase, 2OG-Fe(II) oxygenase family protein | 9 | 8 | 7 | 7 | 6 | 5 | 5 | 3 |
| 17 | TGME49_221380.P1019 | alba 1 | 8 | 7 | 6 | 6 | 6 | 5 | 5 | 3 |
| 1 | TGME49_221450-t26_1-p1 | SPRY domain-containing protein | 9 | 7 | 7 | 6 | 5 | 6 | 5 | 3 |
| 17 | TGME49_221510.P1026 | cytochrome c oxidase subunit ApiCOX18 | 7 | 6 | 6 | 6 | 5 | 5 | 5 | 3 |
| 30 | TGME49_221700-t26_1-p1 | golgi apparatus membrane protein TVP23, putative | 8 | 7 | 7 | 7 | 6 | 3 | 5 | 3 |

| | | | | | | | | | | |
|---|---|---|---|---|---|---|---|---|---|---|
| 21 | TGME49_221710-t26_1-p1 | TBC domain-containing protein TBC13 | 10 | 9 | 7 | 7 | 7 | 6 | 6 | 3 |
| 30 | TGME49_221850-t26_1-p1 | prohibitin family protein, putative | 7 | 7 | 7 | 6 | 6 | 4 | 4 | 3 |
| 17 | TGME49_221970-t26_1-p1 | DUF2423 domain-containing protein | 7 | 7 | 6 | 6 | 5 | 4 | 4 | 3 |
| 21 | TGME49_222000-t26_1-p1 | ribosome biogenesis protein BRX1, putative | 9 | 8 | 8 | 6 | 7 | 5 | 6 | 3 |
| 30 | TGME49_222010-t26_1-p1 | HAD hydrolase, family IIID protein | 9 | 8 | 8 | 4 | 7 | 3 | 3 | 3 |
| 30 | TGME49_222060.P1048 | transporter, cation channel family protein | 6 | 8 | 6 | 8 | 6 | 3 | 3 | 3 |
| 21 | TGME49_222110-t26_1-p1 | PUB domain-containing protein | 9 | 8 | 8 | 6 | 7 | 7 | 6 | 3 |
| 1 | TGME49_222188-t26_1-p1 | microneme protein MIC21L1 | 9 | 7 | 6 | 6 | 5 | 5 | 5 | 3 |
| 30 | TGME49_222220.P1054 | alveolin domain containing intermediate filament IMC7 | 9 | 8 | 7 | 7 | 5 | 3 | 3 | 3 |
| 1 | TGME49_222270-t26_1-p1 | hypothetical protein | 9 | 7 | 7 | 6 | 5 | 5 | 5 | 3 |
| 1 | TGME49_222350-t26_1-p1 | hypothetical protein | 9 | 8 | 7 | 5 | 5 | 6 | 5 | 3 |
| 17 | TGME49_222920-t26_1-p1 | trafficking protein particle complex subunit 2, putative | 6 | 6 | 6 | 6 | 6 | 3 | 6 | 3 |
| 1 | TGME49_222952-t26_1-p1 | phosphohistidine phosphatase | 8 | 7 | 6 | 6 | 6 | 5 | 4 | 3 |
| 30 | TGME49_222970-t26_1-p1 | inositol(myo)-1(or 4)-monophosphatase 2, putative | 8 | 7 | 8 | 6 | 5 | 4 | 5 | 3 |
| 1 | TGME49_222990-t26_1-p1 | ribosomal protein S18, putative | 10 | 8 | 8 | 5 | 6 | 6 | 3 | 3 |
| 21 | TGME49_223020-t26_1-p1 | coproporphyrinogen III oxidase | 9 | 8 | 7 | 8 | 7 | 7 | 6 | 3 |
| 1 | TGME49_223040-t26_1-p1 | hypothetical protein | 8 | 8 | 7 | 7 | 7 | 5 | 5 | 3 |
| 17 | TGME49_223050.P1071 | ribosomal protein RPS20 | 6 | 6 | 6 | 6 | 5 | 5 | 5 | 3 |
| 17 | TGME49_223070.P1073 | hypothetical protein | 7 | 6 | 7 | 6 | 5 | 5 | 5 | 3 |
| 1 | TGME49_223110.P1075 | glycolipid transfer protein GLTP | 8 | 7 | 7 | 7 | 4 | 5 | 3 | 3 |
| 21 | TGME49_223410-t26_1-p1 | eukaryotic initiation factor eIF4E1 | 8 | 8 | 7 | 7 | 7 | 7 | 6 | 3 |
| 1 | TGME49_223580-t26_1-p1 | mediator complex subunit MED4 | 9 | 8 | 7 | 7 | 5 | 5 | 5 | 3 |
| 30 | TGME49_223590.P1085 | proteasome subunit beta type-6, putative | 8 | 7 | 7 | 7 | 5 | 3 | 6 | 3 |
| 30 | TGME49_223680.P1089 | ubiquitin domain-containing protein DSK2 | 9 | 8 | 8 | 6 | 6 | 3 | 5 | 3 |
| 30 | TGME49_223690-t26_1-p1 | hypothetical protein | 7 | 7 | 7 | 6 | 5 | 3 | 3 | 3 |
| 21 | TGME49_223930.P1098 | RNA recognition motif-containing protein | 9 | 8 | 8 | 6 | 7 | 5 | 6 | 3 |
| 30 | TGME49_223985.P1102 | protein phosphatase PPM11B, putative | 10 | 6 | 6 | 7 | 6 | 4 | 5 | 3 |
| 1 | TGME49_224040.P1104 | flagellar associated protein | 8 | 8 | 7 | 7 | 6 | 5 | 5 | 3 |
| 30 | TGME49_224090.P1107 | enoyl-CoA hydratase/isomerase family protein | 8 | 8 | 8 | 6 | 6 | 3 | 4 | 3 |
| 30 | TGME49_224200-t26_1-p1 | tRNA pseudouridine synthase | 10 | 8 | 8 | 6 | 6 | 4 | 5 | 3 |
| 21 | TGME49_224240-t26_1-p1 | protein phosphatase PPM1, putative | 10 | 9 | 5 | 8 | 7 | 6 | 6 | 3 |
| 30 | TGME49_224330.P1115 | proteasome activator pa28 beta subunit protein | 9 | 8 | 8 | 7 | 6 | 3 | 3 | 3 |
| 21 | TGME49_224490-t26_1-p1 | farnesyl/geranylgeranyl diphosphate synthase | 9 | 8 | 8 | 6 | 5 | 6 | 6 | 3 |
| 17 | TGME49_224520.P1120 | alveolin domain containing intermediate filament IMC8 | 7 | 7 | 6 | 6 | 5 | 5 | 5 | 3 |
| 21 | TGME49_224580-t26_1-p1 | RNA recognition motif-containing protein | 9 | 8 | 7 | 7 | 7 | 7 | 7 | 3 |
| 1 | TGME49_224620-t26_1-p1 | hypothetical protein | 8 | 7 | 7 | 6 | 6 | 5 | 4 | 3 |
| 21 | TGME49_224630-t26_1-p1 | zinc finger (CCCH type) motif-containing protein | 10 | 9 | 6 | 6 | 6 | 6 | 6 | 3 |
| 1 | TGME49_224740-t26_1-p1 | hypothetical protein | 9 | 8 | 7 | 6 | 6 | 5 | 5 | 3 |
| 21 | TGME49_224860-t26_1-p1 | exosome component 2, putative | 9 | 8 | 8 | 6 | 7 | 6 | 5 | 3 |
| 21 | TGME49_224880.P1135 | kinesin motor domain-containing protein | 10 | 10 | 3 | 8 | 8 | 6 | 7 | 3 |
| 21 | TGME49_224900-t26_1-p1 | adenylate kinase, putative | 8 | 8 | 8 | 7 | 7 | 7 | 6 | 3 |
| 17 | TGME49_224932-t26_1-p1 | hypothetical protein | 7 | 3 | 6 | 6 | 6 | 5 | 5 | 3 |

| | | | | | | | | | | |
|---|---|---|---|---|---|---|---|---|---|---|
| 30 | TGME49_224960.P1137 | Sec66 domain-containing protein | 8 | 7 | 7 | 6 | 5 | 3 | 5 | 3 |
| 30 | TGME49_225060.P1145 | nucleoredoxin family protein | 8 | 7 | 7 | 6 | 7 | 3 | 3 | 3 |
| 30 | TGME49_225070.P1146 | hypothetical protein | 9 | 9 | 8 | 7 | 6 | 3 | 3 | 3 |
| 17 | TGME49_225080.P1147 | ribosomal protein RPS18 | 6 | 6 | 6 | 6 | 6 | 6 | 6 | 3 |
| 1 | TGME49_225090.P1148 | WD repeat-containing protein | 10 | 9 | 7 | 6 | 5 | 5 | 4 | 3 |
| 1 | TGME49_225240-t26_1-p1 | 50S ribosomal protein L13, putative | 8 | 7 | 7 | 7 | 7 | 6 | 4 | 3 |
| 17 | TGME49_225410-t26_1-p1 | histone H3 centromeric CENH3 | 7 | 6 | 6 | 5 | 4 | 5 | 5 | 3 |
| 30 | TGME49_225480-t26_1-p1 | hypothetical protein | 10 | 8 | 7 | 6 | 5 | 4 | 4 | 3 |
| 17 | TGME49_225710-t26_1-p1 | mitochondrial import inner membrane translocase subunit TIM22 | 7 | 6 | 6 | 5 | 5 | 5 | 4 | 3 |
| 30 | TGME49_225780.P1180 | hypothetical protein | 9 | 9 | 7 | 7 | 5 | 4 | 4 | 3 |
| 17 | TGME49_225810-t26_1-p1 | BIR protein | 8 | 7 | 7 | 7 | 5 | 3 | 7 | 3 |
| 21 | TGME49_225880-t26_1-p1 | UDP-glucose 4-epimerase | 9 | 8 | 8 | 6 | 7 | 6 | 6 | 3 |
| 30 | TGME49_225890-t26_1-p1 | zinc finger (CCCH type) motif-containing protein | 10 | 8 | 7 | 6 | 5 | 3 | 3 | 3 |
| 1 | TGME49_225920.P1184 | hypothetical protein | 10 | 7 | 8 | 5 | 6 | 5 | 3 | 3 |
| 21 | TGME49_225930-t26_1-p1 | triose-phosphate isomerase TPI-I | 8 | 8 | 7 | 8 | 7 | 7 | 7 | 3 |
| 1 | TGME49_225940.P1185 | sialic acid binding protein SABP1 | 8 | 7 | 7 | 6 | 6 | 5 | 5 | 3 |
| 21 | TGME49_225990-t26_1-p1 | acyl transferase domain-containing protein | 9 | 8 | 8 | 8 | 7 | 6 | 6 | 3 |
| 17 | TGME49_226000-t26_1-p1 | ATP synthase subunit delta | 7 | 7 | 6 | 6 | 5 | 5 | 4 | 3 |
| 30 | TGME49_226050-t26_1-p1 | hypothetical protein | 10 | 8 | 7 | 6 | 6 | 3 | 3 | 3 |
| 21 | TGME49_226068-t26_1-p1 | DnaJ domain-containing protein | 9 | 8 | 8 | 6 | 7 | 7 | 7 | 3 |
| 1 | TGME49_226270.P1199 | TMEM147 domain-containing protein | 8 | 7 | 7 | 5 | 5 | 5 | 4 | 3 |
| 1 | TGME49_226410.P1206 | EF-1 guanine nucleotide exchange domain-containing protein | 8 | 8 | 7 | 7 | 6 | 6 | 5 | 3 |
| 21 | TGME49_226550.P1213 | TBC domain-containing protein TBC9 | 8 | 8 | 8 | 8 | 6 | 7 | 6 | 3 |
| 21 | TGME49_226560-t26_1-p1 | pre-mRNA-splicing factor CWC24, putative | 9 | 8 | 8 | 7 | 7 | 7 | 6 | 3 |
| 17 | TGME49_226590.P1216 | cytochrome c oxidase subunit COX2a | 7 | 7 | 7 | 6 | 6 | 5 | 6 | 3 |
| 21 | TGME49_226600-t26_1-p1 | syntaxin 5, putative | 8 | 8 | 8 | 6 | 6 | 6 | 6 | 3 |
| 17 | TGME49_226640.P1219 | zinc binding protein, putative | 6 | 6 | 6 | 6 | 6 | 5 | 3 | 3 |
| 30 | TGME49_226720.P1221 | translin family protein | 8 | 7 | 7 | 6 | 5 | 3 | 5 | 3 |
| 1 | TGME49_226940-t26_1-p1 | ubiquitin carboxyl-terminal hydrolase | 9 | 8 | 8 | 7 | 6 | 6 | 3 | 3 |
| 17 | TGME49_226980-t26_1-p1 | hypothetical protein | 7 | 7 | 6 | 6 | 6 | 5 | 5 | 3 |
| 21 | TGME49_227020.P1237 | histone deacetylase SIR2 | 8 | 8 | 8 | 7 | 6 | 6 | 7 | 3 |
| 17 | TGME49_227030-t26_1-p1 | IMP1-like protein, putative | 7 | 6 | 6 | 6 | 5 | 5 | 5 | 3 |
| 30 | TGME49_227270.P1241 | hypothetical protein | 9 | 8 | 7 | 6 | 5 | 3 | 4 | 3 |
| 21 | TGME49_227560-t26_1-p1 | IWS1 transcription factor, putative | 9 | 9 | 8 | 7 | 6 | 7 | 6 | 3 |
| 21 | TGME49_227650.P1256 | microtubule-associated protein EB1 | 9 | 8 | 7 | 7 | 7 | 6 | 5 | 3 |
| 30 | TGME49_227660.P1257 | DNA methyltransferase DNMTa | 10 | 8 | 6 | 5 | 5 | 3 | 5 | 3 |
| 21 | TGME49_227790.P1259 | glycosyltransferase, group 1 family protein | 9 | 8 | 8 | 7 | 5 | 6 | 6 | 3 |
| 1 | TGME49_227800.P1260 | Eps15-like protein | 10 | 10 | 6 | 6 | 6 | 5 | 5 | 3 |
| 30 | TGME49_227840-t26_1-p1 | Memo family protein | 8 | 7 | 8 | 6 | 6 | 4 | 5 | 3 |
| 21 | TGME49_227850-t26_1-p1 | peptidyl-prolyl cis-trans isomerase, cyclophilin-type domain-containing protein | 9 | 9 | 8 | 6 | 7 | 6 | 7 | 3 |
| 21 | TGME49_228000-t26_1-p1 | splicing factor 3A subunit 2, putative | 8 | 7 | 7 | 8 | 7 | 7 | 7 | 3 |
| 21 | TGME49_228180-t26_1-p1 | cytochrome C oxidase assembly factor COX15, putative | 8 | 8 | 8 | 7 | 7 | 6 | 7 | 3 |
| 17 | TGME49_228290.P1288 | serine hydrolase ASH1 | 8 | 7 | 6 | 5 | 6 | 5 | 5 | 3 |
| 17 | TGME49_228300-t26_1-p1 | CCDC25 protein | 7 | 7 | 7 | 4 | 5 | 3 | 6 | 3 |
| 1 | TGME49_228330-t26_1-p1 | NLI interacting factor family phosphatase | 9 | 7 | 7 | 6 | 6 | 5 | 5 | 3 |

| | | | | | | | | | | |
|---|---|---|---|---|---|---|---|---|---|---|
| 30 | TGME49_228350.P1290 | elongation factor Tu GTP binding domain-containing protein | 10 | 7 | 9 | 7 | 6 | 4 | 4 | 3 |
| 30 | TGME49_228370.P1292 | hypothetical protein | 9 | 8 | 7 | 6 | 7 | 3 | 3 | 3 |
| 21 | TGME49_228470-t26_1-p1 | ribosomal protein RPL15 | 7 | 7 | 7 | 7 | 7 | 5 | 6 | 3 |
| 1 | TGME49_228490.P1297 | protein BCP1, putative | 8 | 8 | 7 | 7 | 6 | 5 | 5 | 3 |
| 30 | TGME49_228630-t26_1-p1 | MIR domain-containing protein | 8 | 6 | 7 | 5 | 6 | 3 | 5 | 3 |
| 30 | TGME49_228760-t26_1-p1 | hypothetical protein | 9 | 8 | 7 | 7 | 6 | 3 | 5 | 3 |
| 21 | TGME49_229210.P1315 | small nuclear ribonucleoprotein polypeptide A', putative | 8 | 7 | 7 | 7 | 6 | 6 | 6 | 3 |
| 30 | TGME49_229270-t26_1-p1 | hypothetical protein | 7 | 7 | 7 | 6 | 5 | 3 | 3 | 3 |
| 21 | TGME49_229360.P1319 | transaldolase | 8 | 8 | 8 | 6 | 7 | 7 | 7 | 3 |
| 30 | TGME49_229420-t26_1-p1 | cytochrome c, putative | 7 | 6 | 6 | 6 | 6 | 3 | 3 | 3 |
| 30 | TGME49_229440.P1324 | zinc finger, C3HC4 type (RING finger) domain-containing protein | 9 | 5 | 8 | 8 | 6 | 3 | 3 | 3 |
| 21 | TGME49_229480-t26_1-p1 | calcium binding protein precursor, putative | 8 | 8 | 7 | 7 | 7 | 6 | 6 | 3 |
| 21 | TGME49_229490.P1326 | tetratricopeptide repeat-containing protein | 10 | 9 | 7 | 7 | 7 | 6 | 6 | 3 |
| 1 | TGME49_229500.P1327 | hypothetical protein | 8 | 7 | 7 | 7 | 6 | 5 | 3 | 3 |
| 21 | TGME49_229650.P1331 | josephin protein | 9 | 9 | 8 | 6 | 7 | 6 | 6 | 3 |
| 17 | TGME49_229670.P1332 | ribosomal protein RPS23 | 6 | 6 | 6 | 6 | 6 | 6 | 6 | 3 |
| 1 | TGME49_229690.P1334 | autophagy-related protein ATG7 | 10 | 8 | 8 | 8 | 5 | 7 | 3 | 3 |
| 1 | TGME49_229740.P1337 | hypothetical protein | 9 | 8 | 8 | 5 | 7 | 6 | 4 | 3 |
| 1 | TGME49_229780.P1340 | GHMP kinase, N-terminal domain-containing protein | 10 | 6 | 7 | 6 | 6 | 6 | 5 | 3 |
| 17 | TGME49_229930-t26_1-p1 | p25-alpha family protein | 7 | 6 | 6 | 6 | 3 | 5 | 5 | 3 |
| 30 | TGME49_230100.P1351 | ER membrane protein complex subunit 3, putative | 8 | 7 | 7 | 7 | 5 | 4 | 3 | 3 |
| 1 | TGME49_230410-t26_1-p1 | peroxiredoxin PRX3 | 8 | 7 | 7 | 6 | 6 | 6 | 5 | 3 |
| 1 | TGME49_230440.P1369 | CHY zinc finger domain-containing protein | 7 | 7 | 8 | 7 | 6 | 7 | 4 | 3 |
| 30 | TGME49_230520.P1375 | cyclophilin 1, putative | 7 | 7 | 7 | 6 | 6 | 3 | 3 | 3 |
| 21 | TGME49_230535-t26_1-p1 | hypothetical protein | 9 | 7 | 6 | 8 | 7 | 7 | 5 | 3 |
| 17 | TGME49_230890.P1389 | PHD-finger domain-containing protein | 8 | 7 | 7 | 5 | 5 | 5 | 5 | 3 |
| 21 | TGME49_231010.P1397 | general transcription factor IIE polypeptide 1 GTF2E1 | 10 | 8 | 8 | 7 | 7 | 6 | 6 | 3 |
| 1 | TGME49_231100-t26_1-p1 | hypothetical protein | 9 | 9 | 7 | 7 | 6 | 5 | 5 | 3 |
| 1 | TGME49_231120-t26_1-p1 | ribosomal protein S11, putative | 9 | 6 | 7 | 6 | 6 | 6 | 3 | 3 |
| 21 | TGME49_231220.P1410 | hypothetical protein | 9 | 9 | 8 | 7 | 7 | 6 | 6 | 3 |
| 1 | TGME49_231280.P1411 | WD domain, G-beta repeat-containing protein | 8 | 8 | 7 | 6 | 6 | 6 | 5 | 3 |
| 30 | TGME49_231470.P1419 | hypothetical protein | 9 | 8 | 8 | 7 | 6 | 3 | 3 | 3 |
| 17 | TGME49_231590-t26_1-p1 | SGS domain-containing protein | 8 | 7 | 7 | 6 | 5 | 5 | 5 | 3 |
| 30 | TGME49_231750.P1422 | hypothetical protein | 9 | 9 | 8 | 7 | 6 | 4 | 3 | 3 |
| 1 | TGME49_232130.P1446 | thioredoxin-like associated protein TLAP2 | 8 | 8 | 7 | 7 | 5 | 6 | 3 | 3 |
| 30 | TGME49_232160.P1449 | zinc finger, C3HC4 type (RING finger) domain-containing protein | 9 | 8 | 8 | 5 | 6 | 3 | 5 | 3 |
| 1 | TGME49_232260.P1454 | transverse sutures component TSC5 | 8 | 8 | 7 | 7 | 6 | 6 | 5 | 3 |
| 21 | TGME49_232300.P1456 | ribosomal protein RPS3 | 8 | 8 | 7 | 7 | 7 | 7 | 7 | 3 |
| 1 | TGME49_232390.P1463 | 18S rRNA biogenesis protein RCL1 protein | 9 | 8 | 8 | 8 | 5 | 6 | 4 | 3 |
| 21 | TGME49_232410.P1465 | thioredoxin-like protein 1 | 7 | 7 | 7 | 7 | 7 | 6 | 6 | 3 |
| 30 | TGME49_232430.P1466 | kinetochore protein SPC25, putative | 9 | 7 | 7 | 6 | 5 | 4 | 3 | 3 |
| 30 | TGME49_232510.P1469 | RING zinc finger protein | 10 | 9 | 7 | 7 | 5 | 4 | 4 | 3 |

| | | | | | | | | | | |
|---|---|---|---|---|---|---|---|---|---|---|
| 30 | TGME49_232550-t26_1-p1 | hypothetical protein | 7 | 7 | 7 | 6 | 5 | 4 | 4 | 3 |
| 30 | TGME49_232650.P1478 | EELM2 domain-containing protein | 8 | 9 | 8 | 7 | 5 | 3 | 5 | 3 |
| 1 | TGME49_232660-t26_1-p1 | Hsp70 interacting protein HIP | 9 | 8 | 8 | 7 | 7 | 6 | 4 | 3 |
| 30 | TGME49_232690-t26_1-p1 | DP-fucose transporter, putative | 9 | 7 | 7 | 7 | 6 | 3 | 3 | 3 |
| 17 | TGME49_232815-t26_1-p1 | Erv1 / Alr family protein | 7 | 7 | 6 | 5 | 6 | 5 | 5 | 3 |
| 17 | TGME49_232940-t26_1-p1 | heat shock protein HSP20 | 3 | 7 | 7 | 6 | 6 | 5 | 5 | 3 |
| 21 | TGME49_233100-t26_1-p1 | SPFH domain / Band 7 family protein | 8 | 7 | 8 | 7 | 8 | 7 | 7 | 3 |
| 21 | TGME49_233130-t26_1-p1 | nucleoside transporter protein | 9 | 9 | 8 | 6 | 6 | 5 | 6 | 3 |
| 17 | TGME49_233140-t26_1-p1 | deoxyuridine 5'-triphosphate nucleotidohydrolase, putative | 7 | 6 | 6 | 6 | 5 | 5 | 5 | 3 |
| 17 | TGME49_233230-t26_1-p1 | RNA-binding protein 8A family protein | 7 | 6 | 6 | 5 | 5 | 5 | 4 | 3 |
| 17 | TGME49_233350-t26_1-p1 | nuclear transport factor 2, putative | 6 | 6 | 6 | 5 | 6 | 5 | 5 | 3 |
| 17 | TGME49_233400-t26_1-p1 | TAF9 RNA polymerase II, TATA box binding protein (TBP)-associated factor isoform 2 family protein | 7 | 7 | 6 | 6 | 6 | 3 | 6 | 3 |
| 30 | TGME49_233500-t26_1-p1 | triose-phosphate isomerase TPI-II | 9 | 8 | 8 | 6 | 7 | 3 | 3 | 3 |
| 17 | TGME49_233720-t26_1-p1 | DNA-directed RNA polymerase II RPBABC8 | 6 | 6 | 6 | 6 | 6 | 3 | 5 | 3 |
| 30 | TGME49_233850.P1520 | coiled-coil domain-containing protein | 7 | 7 | 7 | 7 | 5 | 4 | 4 | 3 |
| 1 | TGME49_233940.P1525 | centrosomal protein CEP72, putative | 9 | 7 | 8 | 7 | 7 | 5 | 5 | 3 |
| 30 | TGME49_234350.P1532 | hypothetical protein | 10 | 7 | 7 | 5 | 5 | 3 | 3 | 3 |
| 17 | TGME49_234450.P1536 | ribosomal protein RPS15A | 6 | 6 | 6 | 6 | 6 | 6 | 6 | 3 |
| 1 | TGME49_234560-t26_1-p1 | hypothetical protein | 10 | 7 | 8 | 7 | 5 | 6 | 5 | 3 |
| 21 | TGME49_235000-t26_1-p1 | phosphorylase family protein | 10 | 7 | 8 | 5 | 6 | 7 | 6 | 3 |
| 21 | TGME49_235315.P1552 | PAN domain-containing protein | 9 | 3 | 10 | 7 | 7 | 5 | 6 | 3 |
| 21 | TGME49_235350-t26_1-p1 | cwfj family protein | 10 | 8 | 8 | 7 | 6 | 6 | 6 | 3 |
| 30 | TGME49_235402.P1558 | CorA family Mg2+ transporter protein | 10 | 8 | 8 | 7 | 6 | 3 | 3 | 3 |
| 30 | TGME49_235450.P1561 | ubiquitin-conjugating enzyme subfamily protein | 7 | 7 | 7 | 7 | 6 | 3 | 5 | 3 |
| 30 | TGME49_235478.P1563 | pantothenate kinase 2 | 10 | 8 | 6 | 5 | 5 | 4 | 4 | 3 |
| 21 | TGME49_235540-t26_1-p1 | eukaryotic initiation factor-2 beta, putative | 8 | 7 | 7 | 5 | 7 | 7 | 7 | 3 |
| 1 | TGME49_235550-t26_1-p1 | PHD-finger domain-containing protein | 9 | 6 | 7 | 6 | 4 | 6 | 5 | 3 |
| 30 | TGME49_236030.P1584 | hypothetical protein | 7 | 7 | 7 | 6 | 6 | 3 | 3 | 3 |
| 1 | TGME49_236070.P1587 | pyrroline-5-carboxylate reductase | 8 | 8 | 7 | 7 | 7 | 5 | 5 | 3 |
| 21 | TGME49_236110-t26_1-p1 | autophagy-related protein 3 atg3, putative | 9 | 8 | 8 | 6 | 7 | 6 | 6 | 3 |
| 21 | TGME49_236200-t26_1-p1 | DNA mismatch repair protein, C-terminal domain-containing protein | 9 | 7 | 7 | 5 | 6 | 7 | 7 | 3 |
| 30 | TGME49_236250-t26_1-p1 | regulator of chromosome condensation (RCC1) repeat-containing protein | 10 | 8 | 7 | 6 | 5 | 4 | 4 | 3 |
| 30 | TGME49_236520.P1593 | hypothetical protein | 8 | 7 | 7 | 5 | 6 | 3 | 3 | 3 |
| 1 | TGME49_236530-t26_1-p1 | DUF298 domain-containing protein | 8 | 7 | 7 | 7 | 5 | 5 | 5 | 3 |
| 30 | TGME49_236540.P1594 | RNA recognition motif-containing protein | 9 | 8 | 5 | 7 | 5 | 3 | 6 | 3 |
| 21 | TGME49_236570.P1596 | lysine decarboxylase family protein | 9 | 9 | 7 | 7 | 8 | 7 | 6 | 3 |
| 17 | TGME49_236580.P1597 | Prp31-15.5k-U4 Snrna Complex family protein | 6 | 6 | 6 | 6 | 6 | 5 | 5 | 3 |
| 1 | TGME49_236595.P1598 | ankyrin repeat-containing protein | 9 | 5 | 8 | 7 | 7 | 5 | 4 | 3 |
| 30 | TGME49_236800-t26_1-p1 | hypothetical protein | 9 | 8 | 7 | 6 | 5 | 4 | 4 | 3 |
| 21 | TGME49_236910.P1606 | U2 snRNP auxiliary factor, putative | 8 | 4 | 5 | 7 | 7 | 6 | 7 | 3 |
| 1 | TGME49_237000.P1612 | polyphosphoinositide binding protein, putative | 8 | 8 | 8 | 6 | 7 | 6 | 3 | 3 |
| 21 | TGME49_237150-t26_1-p1 | minor histocompatibility antigen h13, putative | 9 | 7 | 8 | 6 | 7 | 7 | 7 | 3 |

| | | | | | | | | | | |
|---|---|---|---|---|---|---|---|---|---|---|
| 30 | TGME49_237250.P1624 | hypothetical protein | 8 | 7 | 6 | 6 | 5 | 3 | 5 | 3 |
| 17 | TGME49_237260-t26_1-p1 | Coiled-coil domain containing 124 family protein | 7 | 7 | 6 | 6 | 5 | 5 | 5 | 3 |
| 30 | TGME49_237410.P1627 | protein phosphatase PPM16, putative | 10 | 8 | 6 | 5 | 5 | 4 | 4 | 3 |
| 30 | TGME49_237460-t26_1-p1 | DER1-like protein | 9 | 7 | 7 | 6 | 5 | 4 | 4 | 3 |
| 21 | TGME49_237480.P1629 | BRCA1 C Terminus (BRCT) domain-containing protein | 10 | 7 | 9 | 5 | 6 | 6 | 6 | 3 |
| 17 | TGME49_237560.P1633 | iron-sulfur cluster assembly protein ISU1 | 7 | 6 | 6 | 6 | 6 | 6 | 6 | 3 |
| 17 | TGME49_237820-t26_1-p1 | IMC sub-compartment protein ISP2 | 6 | 6 | 6 | 5 | 5 | 5 | 5 | 3 |
| 21 | TGME49_238000-t26_1-p1 | peptidyl-prolyl isomerase | 8 | 8 | 8 | 7 | 7 | 6 | 6 | 3 |
| 17 | TGME49_238010.P1638 | ribosomal protein RPL23A | 7 | 7 | 6 | 6 | 5 | 5 | 4 | 3 |
| 1 | TGME49_238070.P1642 | glutaredoxin domain-containing protein | 8 | 7 | 7 | 6 | 5 | 5 | 4 | 3 |
| 21 | TGME49_238100-t26_1-p1 | transmembrane protein | 8 | 7 | 7 | 7 | 6 | 7 | 6 | 3 |
| 30 | TGME49_238380-t26_1-p1 | hypothetical protein | 10 | 9 | 7 | 7 | 6 | 3 | 4 | 3 |
| 17 | TGME49_239100-t26_1-p1 | ribosomal protein RPS7 | 7 | 7 | 7 | 7 | 7 | 4 | 6 | 3 |
| 21 | TGME49_239310-t26_1-p1 | ribulose 5-phosphate isomerase | 8 | 7 | 7 | 6 | 6 | 5 | 6 | 3 |
| 1 | TGME49_239720.P1682 | 50S ribosomal protein l24, putative | 8 | 7 | 7 | 7 | 7 | 6 | 4 | 3 |
| 17 | TGME49_239760.P1684 | ribosomal protein RPL22 | 6 | 6 | 5 | 5 | 5 | 5 | 5 | 3 |
| 21 | TGME49_239790-t26_1-p1 | BRCA1 C Terminus (BRCT) domain-containing protein | 10 | 8 | 8 | 8 | 6 | 6 | 6 | 3 |
| 1 | TGME49_239800.P1687 | transverse sutures component TSC2 | 8 | 7 | 7 | 7 | 5 | 5 | 4 | 3 |
| 30 | TGME49_239830-t26_1-p1 | TBC domain-containing protein TBC11 | 10 | 7 | 8 | 7 | 5 | 4 | 4 | 3 |
| 30 | TGME49_239855-t26_1-p1 | Ras-associated protein Rap1 isoform 1 family protein | 10 | 8 | 7 | 6 | 5 | 4 | 4 | 3 |
| 30 | TGME49_239870.P1690 | 5'-AMP-activated protein kinase subunit gamma | 10 | 7 | 7 | 6 | 5 | 3 | 3 | 3 |
| 30 | TGME49_240070.P1693 | RNA recognition motif-containing protein | 8 | 7 | 7 | 6 | 6 | 3 | 3 | 3 |
| 30 | TGME49_240080.P1694 | transporter, major facilitator family protein | 10 | 7 | 6 | 6 | 6 | 3 | 4 | 3 |
| 30 | TGME49_240220-t26_1-p1 | golgi localizing protein GLP4 | 10 | 8 | 7 | 6 | 5 | 4 | 3 | 3 |
| 30 | TGME49_240300-t26_1-p1 | zinc finger domain, LSD1 subclass domain-containing protein | 10 | 6 | 6 | 5 | 6 | 4 | 3 | 3 |
| 30 | TGME49_240520-t26_1-p1 | hypothetical protein | 9 | 8 | 6 | 6 | 6 | 4 | 4 | 3 |
| 21 | TGME49_240590-t26_1-p1 | DNA-directed RNA polymerase II RPB5 | 7 | 5 | 7 | 7 | 7 | 6 | 7 | 3 |
| 30 | TGME49_240660.P1711 | rRNA-processing protein EFG1, putative | 9 | 8 | 7 | 6 | 5 | 4 | 3 | 3 |
| 17 | TGME49_240670-t26_1-p1 | adrenodoxin-type ferredoxin, putative | 7 | 6 | 6 | 5 | 5 | 5 | 5 | 3 |
| 17 | TGME49_240730-t26_1-p1 | hypothetical protein | 7 | 6 | 6 | 6 | 6 | 5 | 5 | 3 |
| 30 | TGME49_240830.P1717 | hydrolase, alpha/beta fold family protein | 8 | 8 | 7 | 6 | 5 | 3 | 4 | 3 |
| 21 | TGME49_240860.P1718 | acyl-glycerol-3-phosphate acyltransferase AGPAT | 8 | 7 | 8 | 6 | 7 | 6 | 5 | 3 |
| 21 | TGME49_240980-t26_1-p1 | WD repeat-containing protein | 10 | 9 | 7 | 7 | 7 | 5 | 6 | 3 |
| 30 | TGME49_241000.P1722 | rhoptry protein ROP51 | 8 | 7 | 7 | 6 | 6 | 3 | 3 | 3 |
| 21 | TGME49_241160-t26_1-p1 | hemolysin-III related subfamily protein | 8 | 8 | 7 | 7 | 7 | 6 | 5 | 3 |
| 30 | TGME49_241180-t26_1-p1 | hypothetical protein | 8 | 7 | 7 | 6 | 6 | 3 | 5 | 3 |
| 1 | TGME49_241870-t26_1-p1 | tRNA ligase class I (E and Q), catalytic domain-containing protein | 9 | 8 | 7 | 6 | 5 | 5 | 5 | 3 |
| 21 | TGME49_242000-t26_1-p1 | AMMECR1 domain-containing protein | 8 | 7 | 7 | 7 | 5 | 6 | 6 | 3 |
| 1 | TGME49_242380-t26_1-p1 | fatty acid elongase | 9 | 8 | 8 | 7 | 7 | 6 | 3 | 3 |
| 21 | TGME49_242800.P1748 | ribosome biogenesis protein NSA2, putative | 8 | 6 | 8 | 7 | 7 | 7 | 7 | 3 |
| 1 | TGME49_242870-t26_1-p1 | histone lysine methyltransferase, SET, putative | 9 | 8 | 8 | 7 | 5 | 5 | 5 | 3 |

| | | | | | | | | | | |
|---|---|---|---|---|---|---|---|---|---|---|
| 30 | TGME49_242880.P1752 | phosphopantothenoylcysteine decarboxylase | 8 | 7 | 6 | 6 | 5 | 3 | 4 | 3 |
| 30 | TGME49_243240-t26_1-p1 | WD domain, G-beta repeat-containing protein | 9 | 9 | 8 | 5 | 7 | 3 | 3 | 3 |
| 1 | TGME49_243340.P1758 | atypical MEK-related kinase (incomplete catalytic triad) | 9 | 8 | 8 | 6 | 5 | 5 | 4 | 3 |
| 30 | TGME49_243350-t26_1-p1 | gamma-glutamyl hydrolase | 10 | 7 | 7 | 6 | 5 | 4 | 4 | 3 |
| 21 | TGME49_243480-t26_1-p1 | 50S ribosomal protein L3, putative | 9 | 8 | 8 | 6 | 7 | 6 | 6 | 3 |
| 30 | TGME49_243510-t26_1-p1 | OTU domain-containing protein OTUD6B | 8 | 7 | 7 | 5 | 5 | 3 | 5 | 3 |
| 21 | TGME49_243550.P1764 | H-SHIPPO 1, putative | 8 | 7 | 8 | 8 | 7 | 6 | 5 | 3 |
| 17 | TGME49_243570-t26_1-p1 | ribosomal protein RPS26 | 6 | 6 | 5 | 5 | 5 | 5 | 5 | 3 |
| 17 | TGME49_243580.P1766 | Hit family protein involved in cell-cycle regulation, putative | 6 | 6 | 6 | 6 | 5 | 5 | 5 | 3 |
| 30 | TGME49_243680.P1770 | dihydrodipicolinate reductase | 8 | 8 | 7 | 7 | 6 | 3 | 3 | 3 |
| 1 | TGME49_243730.P1772 | rhoptry protein ROP9 | 8 | 7 | 6 | 6 | 5 | 5 | 3 | 3 |
| 30 | TGME49_243910-t26_1-p1 | Cof family hydrolase subfamily protein | 9 | 8 | 7 | 6 | 6 | 4 | 3 | 3 |
| 21 | TGME49_243950-t26_1-p1 | prohibitin, putative | 8 | 8 | 8 | 7 | 8 | 7 | 7 | 3 |
| 1 | TGME49_244050.P1780 | pentatricopeptide repeat-containing protein PPR1 | 10 | 5 | 9 | 8 | 7 | 5 | 4 | 3 |
| 21 | TGME49_244110.P1782 | nucleosome assembly protein (nap) protein | 9 | 8 | 8 | 7 | 7 | 6 | 6 | 3 |
| 30 | TGME49_244150-t26_1-p1 | glycerate kinase | 8 | 7 | 7 | 6 | 5 | 3 | 3 | 3 |
| 17 | TGME49_244240.P1785 | methyltransferase, putative | 8 | 7 | 6 | 5 | 4 | 5 | 5 | 3 |
| 30 | TGME49_244250-t26_1-p1 | hypothetical protein | 9 | 7 | 6 | 6 | 5 | 4 | 4 | 3 |
| 17 | TGME49_244320.P1786 | ribosomal protein RPL24 | 6 | 6 | 6 | 6 | 5 | 3 | 5 | 3 |
| 30 | TGME49_244390.P1789 | coatomer epsilon subunit protein | 8 | 8 | 7 | 6 | 6 | 4 | 4 | 3 |
| 1 | TGME49_244430-t26_1-p1 | pseudouridylate synthase, putative | 10 | 7 | 7 | 7 | 6 | 5 | 5 | 3 |
| 1 | TGME49_244660-t26_1-p1 | hypothetical protein | 10 | 7 | 7 | 6 | 6 | 5 | 5 | 3 |
| 21 | TGME49_244700-t26_1-p1 | NAD(+)/NADH kinase domain-containing protein | 10 | 9 | 5 | 7 | 6 | 7 | 6 | 3 |
| 17 | TGME49_244840-t26_1-p1 | zinc knuckle domain-containing protein | 7 | 6 | 6 | 5 | 5 | 5 | 4 | 3 |
| 17 | TGME49_244860-t26_1-p1 | PH domain-containing protein | 8 | 7 | 7 | 5 | 5 | 4 | 5 | 3 |
| 1 | TGME49_244870-t26_1-p1 | quinone oxidoreductase, putative | 9 | 6 | 6 | 7 | 4 | 6 | 5 | 3 |
| 17 | TGME49_245450.P1805 | ATP synthase-associated protein | 6 | 6 | 6 | 6 | 6 | 5 | 4 | 3 |
| 21 | TGME49_245460-t26_1-p1 | ribosomal protein RPS8 | 7 | 7 | 7 | 7 | 7 | 7 | 5 | 3 |
| 17 | TGME49_245620.P1813 | ribosomal-ubiquitin protein RPS27A | 6 | 6 | 6 | 6 | 5 | 3 | 5 | 3 |
| 30 | TGME49_245640.P1814 | hypothetical protein | 9 | 8 | 7 | 6 | 6 | 4 | 5 | 3 |
| 17 | TGME49_245680-t26_1-p1 | ribosomal protein RPL21 | 6 | 6 | 6 | 5 | 6 | 3 | 6 | 3 |
| 1 | TGME49_246050-t26_1-p1 | hypothetical protein | 10 | 7 | 7 | 8 | 5 | 5 | 5 | 3 |
| 17 | TGME49_246100.P1824 | phosducin, putative | 8 | 6 | 6 | 7 | 6 | 3 | 6 | 3 |
| 1 | TGME49_246110-t26_1-p1 | haloacid dehalogenase-like hydrolase, putative | 9 | 8 | 8 | 5 | 7 | 6 | 3 | 3 |
| 21 | TGME49_246120-t26_1-p1 | tetratricopeptide repeat-containing protein | 9 | 7 | 8 | 7 | 6 | 5 | 6 | 3 |
| 21 | TGME49_246130-t26_1-p1 | serpin (serine proteinase inhibitor) superfamily protein | 9 | 8 | 8 | 7 | 6 | 6 | 6 | 3 |
| 17 | TGME49_246560.P1836 | vacuolar ATP synthase subunit g, putative | 6 | 6 | 6 | 6 | 3 | 5 | 5 | 3 |
| 30 | TGME49_246570-t26_1-p1 | hypothetical protein | 7 | 7 | 7 | 6 | 5 | 3 | 5 | 3 |
| 17 | TGME49_246620.P1837 | G10 family protein | 6 | 5 | 6 | 6 | 6 | 4 | 5 | 3 |
| 1 | TGME49_246720-t26_1-p1 | hypothetical protein | 9 | 8 | 8 | 7 | 6 | 5 | 3 | 3 |
| 30 | TGME49_246730.P1839 | rRNA-processing protein EBP2, putative | 8 | 8 | 7 | 6 | 5 | 3 | 5 | 3 |
| 30 | TGME49_246780-t26_1-p1 | hypothetical protein | 10 | 8 | 7 | 6 | 5 | 3 | 3 | 3 |
| 30 | TGME49_247020-t26_1-p1 | high mobility group protein b1, putative | 8 | 7 | 6 | 6 | 5 | 3 | 5 | 3 |

| | | | | | | | | | | |
|---|---|---|---|---|---|---|---|---|---|---|
| 30 | TGME49_247220-t26_1-p1 | nudix -type motif 9 isoform a family protein | 9 | 8 | 8 | 7 | 7 | 3 | 5 | 3 |
| 21 | TGME49_247230.P1846 | centrin 1 | 7 | 7 | 7 | 6 | 6 | 6 | 6 | 3 |
| 21 | TGME49_247240-t26_1-p1 | ubiquitin carboxyl-terminal hydrolase, family 1 protein | 9 | 8 | 8 | 8 | 6 | 6 | 6 | 3 |
| 1 | TGME49_247270.P1847 | GEX1 domain-containing protein | 10 | 9 | 7 | 7 | 6 | 6 | 3 | 3 |
| 1 | TGME49_247350.P1849 | thioredoxin domain-containing protein | 8 | 7 | 7 | 5 | 6 | 6 | 5 | 3 |
| 21 | TGME49_247460-t26_1-p1 | proliferating cell nuclear antigen PCNA1 | 8 | 8 | 8 | 8 | 7 | 6 | 6 | 3 |
| 17 | TGME49_247630-t26_1-p1 | membrane protein, putative | 7 | 7 | 6 | 5 | 5 | 5 | 5 | 3 |
| 17 | TGME49_247670.P1863 | ribulose-phosphate 3 epimerase family protein | 7 | 7 | 7 | 5 | 6 | 6 | 6 | 3 |
| 30 | TGME49_247780.P1867 | hypothetical protein | 8 | 7 | 7 | 7 | 6 | 4 | 4 | 3 |
| 21 | TGME49_247930.P1869 | SNARE domain-containing protein | 8 | 8 | 8 | 8 | 7 | 6 | 5 | 3 |
| 30 | TGME49_248100.P1872 | synaptobrevin protein | 8 | 7 | 7 | 7 | 5 | 3 | 5 | 3 |
| 30 | TGME49_248110.P1873 | repressor of RNA polymerase III transcription MAF1, putative | 9 | 8 | 7 | 7 | 5 | 4 | 4 | 3 |
| 30 | TGME49_248130.P1875 | hypothetical protein | 10 | 8 | 8 | 6 | 5 | 3 | 3 | 3 |
| 21 | TGME49_248200-t26_1-p1 | ribosomal RNA (adenine(1779)-N(6)/adenine(1780)-N(6))- dimethyltransferase, putative | 9 | 8 | 8 | 6 | 8 | 6 | 7 | 3 |
| 1 | TGME49_248310-t26_1-p1 | vacuolar protein sorting-associated protein 32, putative | 9 | 8 | 7 | 7 | 6 | 5 | 4 | 3 |
| 30 | TGME49_248320-t26_1-p1 | carrier superfamily protein | 9 | 8 | 8 | 7 | 5 | 3 | 4 | 3 |
| 21 | TGME49_248340.P1883 | GTP-binding nuclear protein ran/tc4 | 7 | 7 | 7 | 7 | 7 | 7 | 6 | 3 |
| 17 | TGME49_248370.P1885 | prefoldin subunit 6, putative | 6 | 6 | 6 | 5 | 5 | 5 | 5 | 3 |
| 17 | TGME49_248390.P1886 | ribosomal protein RPL26 | 6 | 6 | 6 | 6 | 6 | 3 | 6 | 3 |
| 17 | TGME49_248460.P1890 | ubiquitin, putative | 7 | 7 | 6 | 6 | 5 | 5 | 5 | 3 |
| 21 | TGME49_248480-t26_1-p1 | ribosomal protein RPS9 | 7 | 7 | 7 | 7 | 7 | 7 | 7 | 3 |
| 30 | TGME49_248500-t26_1-p1 | nucleoporin NUP37 | 8 | 8 | 7 | 7 | 6 | 3 | 4 | 3 |
| 30 | TGME49_248510.P1892 | hypothetical protein | 10 | 10 | 7 | 7 | 6 | 3 | 5 | 3 |
| 30 | TGME49_248770.P1905 | hypothetical protein | 9 | 8 | 7 | 5 | 5 | 3 | 4 | 3 |
| 21 | TGME49_248810.P1906 | nuclear factor NF7 | 9 | 9 | 8 | 7 | 6 | 6 | 6 | 3 |
| 17 | TGME49_248880.P1910 | ras-related protein RAB7 | 7 | 7 | 7 | 5 | 4 | 3 | 6 | 3 |
| 21 | TGME49_248890.P1911 | actin-like protein ALP3b | 10 | 9 | 8 | 6 | 6 | 6 | 6 | 3 |
| 1 | TGME49_248980-t26_1-p1 | NudC family protein | 9 | 8 | 7 | 7 | 6 | 5 | 5 | 3 |
| 21 | TGME49_249030.P1917 | endonuclease/exonuclease/phosphatase family protein | 9 | 9 | 8 | 7 | 7 | 6 | 6 | 3 |
| 17 | TGME49_249240.P1922 | calmodulin, putative | 6 | 6 | 6 | 6 | 6 | 6 | 6 | 3 |
| 17 | TGME49_249250-t26_1-p1 | ribosomal protein RPL35A | 6 | 6 | 6 | 5 | 5 | 5 | 4 | 3 |
| 1 | TGME49_249300.P1925 | hypothetical protein | 8 | 7 | 7 | 6 | 7 | 5 | 3 | 3 |
| 17 | TGME49_249330-t26_1-p1 | DNA repair protein rad10 subfamily protein | 5 | 7 | 7 | 5 | 6 | 5 | 5 | 3 |
| 30 | TGME49_249350.P1927 | esterase/lipase/thioesterase domain-containing protein | 9 | 7 | 7 | 6 | 6 | 4 | 4 | 3 |
| 1 | TGME49_249650.P1939 | apolipoprotein A-I binding protein, putative | 8 | 7 | 7 | 6 | 6 | 6 | 3 | 3 |
| 30 | TGME49_249680-t26_1-p1 | RNA polymerase II associated Paf1 complex component PAF1 | 9 | 8 | 7 | 5 | 5 | 4 | 3 | 3 |
| 30 | TGME49_249690-t26_1-p1 | hypothetical protein | 9 | 9 | 8 | 7 | 6 | 4 | 4 | 3 |
| 30 | TGME49_249710-t26_1-p1 | Parkinson disease 7 domain containing 1 family protein | 10 | 7 | 7 | 6 | 5 | 4 | 4 | 3 |
| 17 | TGME49_249720-t26_1-p1 | ATP synthase F0 subunit 9, putative | 7 | 6 | 6 | 5 | 6 | 5 | 5 | 3 |
| 30 | TGME49_249770.P1944 | Nmda1 protein | 7 | 7 | 7 | 7 | 4 | 3 | 3 | 3 |

| | | | | | | | | | | |
|---|---|---|---|---|---|---|---|---|---|---|
| 30 | TGME49_249870-t26_1-p1 | hypothetical protein | 8 | 8 | 7 | 6 | 5 | 3 | 4 | 3 |
| 17 | TGME49_249910-t26_1-p1 | mitochondrial import inner membrane translocase subunit TIM16, putative | 6 | 6 | 5 | 5 | 5 | 5 | 5 | 3 |
| 21 | TGME49_249950-t26_1-p1 | Mak16 protein | 8 | 8 | 7 | 7 | 6 | 5 | 6 | 3 |
| 17 | TGME49_249970.P1953 | acylated pleckstrin-homology domain-containing protein APH | 7 | 7 | 6 | 6 | 5 | 5 | 6 | 3 |
| 30 | TGME49_249980.P1954 | leucine rich repeat-containing protein | 8 | 8 | 7 | 6 | 5 | 4 | 4 | 3 |
| 17 | TGME49_250330-t26_1-p1 | DDRGK domain-containing protein | 8 | 6 | 7 | 6 | 4 | 5 | 5 | 3 |
| 21 | TGME49_250340.P1958 | centrin 2 | 7 | 7 | 7 | 6 | 6 | 6 | 6 | 3 |
| 1 | TGME49_250730.P1962 | 60S ribosomal protein L7-B, putative | 8 | 8 | 8 | 6 | 7 | 5 | 4 | 3 |
| 17 | TGME49_250810.P1964 | ribosomal protein RPL35 | 6 | 6 | 6 | 5 | 5 | 5 | 5 | 3 |
| 30 | TGME49_250830-t26_1-p1 | 26S proteasome regulatory subunit RPN12, putative | 9 | 8 | 7 | 7 | 6 | 3 | 4 | 3 |
| 21 | TGME49_250870.P1967 | palmitoyltransferase DHHC1 | 9 | 7 | 7 | 6 | 7 | 6 | 7 | 3 |
| 30 | TGME49_250890-t26_1-p1 | 3'-5' exonuclease domain-containing protein | 9 | 8 | 7 | 6 | 5 | 3 | 4 | 3 |
| 30 | TGME49_250920.P1971 | tetratricopeptide repeat-containing protein | 8 | 7 | 8 | 6 | 6 | 3 | 4 | 3 |
| 17 | TGME49_251440-t26_1-p1 | troponin c, isotype gamma, putative | 6 | 6 | 6 | 6 | 5 | 5 | 5 | 3 |
| 21 | TGME49_251600-t26_1-p1 | IMP4 family U3 small nucleolar ribonucleoprotein (snoRNP), putative | 9 | 8 | 8 | 8 | 8 | 5 | 5 | 3 |
| 21 | TGME49_251630.P1980 | zinc transporter ZNT | 8 | 7 | 7 | 7 | 6 | 6 | 6 | 3 |
| 17 | TGME49_251680.P1981 | translationally-controlled tumor protein homolog | 7 | 7 | 6 | 6 | 6 | 5 | 5 | 3 |
| 1 | TGME49_251750-t26_1-p1 | MORN repeat-containing protein | 7 | 7 | 7 | 7 | 6 | 5 | 3 | 3 |
| 30 | TGME49_251840-t26_1-p1 | hypothetical protein | 9 | 8 | 7 | 6 | 6 | 3 | 4 | 3 |
| 1 | TGME49_251940-t26_1-p1 | leucine rich repeat-containing protein | 8 | 8 | 7 | 7 | 6 | 6 | 4 | 3 |
| 17 | TGME49_251950-t26_1-p1 | ribosomal protein l7/l12 c-terminal domain-containing protein | 8 | 7 | 6 | 5 | 5 | 5 | 4 | 3 |
| 30 | TGME49_252200.P1987 | palmitoyltransferase DHHC7 | 9 | 5 | 8 | 8 | 6 | 3 | 3 | 3 |
| 30 | TGME49_252260.P1989 | hypothetical protein | 9 | 7 | 7 | 7 | 4 | 4 | 4 | 3 |
| 21 | TGME49_252270-t26_1-p1 | L1P family of ribosomal protein | 9 | 8 | 8 | 7 | 6 | 5 | 6 | 3 |
| 1 | TGME49_252440.P1999 | GPI-anchor transamidase, putative | 8 | 7 | 8 | 7 | 6 | 5 | 5 | 3 |
| 21 | TGME49_252465-t26_1-p1 | radical SAM domain-containing protein | 9 | 6 | 8 | 5 | 6 | 7 | 7 | 3 |
| 21 | TGME49_252490-t26_1-p1 | vacuolar protein sorting 29, putative | 7 | 7 | 7 | 6 | 7 | 6 | 6 | 3 |
| 17 | TGME49_253070.P2009 | hydrolase, TatD family protein | 7 | 7 | 3 | 5 | 5 | 6 | 6 | 3 |
| 30 | TGME49_253080-t26_1-p1 | hypothetical protein | 8 | 7 | 6 | 6 | 5 | 4 | 4 | 3 |
| 1 | TGME49_253110.P2011 | kinesin-8, putative | 9 | 9 | 3 | 9 | 8 | 7 | 4 | 3 |
| 30 | TGME49_253300-t26_1-p1 | hypothetical protein | 9 | 7 | 8 | 7 | 6 | 3 | 3 | 3 |
| 1 | TGME49_253360.P2018 | SNARE protein STX20 | 8 | 8 | 7 | 7 | 6 | 5 | 5 | 3 |
| 17 | TGME49_253420.P2021 | PQ loop repeat-containing protein | 8 | 5 | 5 | 5 | 5 | 5 | 5 | 3 |
| 30 | TGME49_253570.P2028 | hypothetical protein | 8 | 7 | 7 | 6 | 5 | 4 | 4 | 3 |
| 17 | TGME49_253640-t26_1-p1 | hypothetical protein | 7 | 6 | 7 | 6 | 6 | 4 | 6 | 3 |
| 17 | TGME49_253820-t26_1-p1 | hypothetical protein | 7 | 7 | 6 | 6 | 5 | 4 | 4 | 3 |
| 21 | TGME49_253860.P2036 | tyrosine kinase-like protein TKL3 | 10 | 8 | 8 | 6 | 6 | 7 | 6 | 3 |
| 30 | TGME49_253940-t26_1-p1 | CAM Kinase family, incomplete catalytic triad | 9 | 8 | 8 | 6 | 7 | 3 | 3 | 3 |
| 30 | TGME49_254050-t26_1-p1 | optic atrophy 3 protein (opa3) protein | 8 | 7 | 7 | 6 | 6 | 3 | 5 | 3 |
| 30 | TGME49_254070.P2045 | hypothetical protein | 8 | 7 | 6 | 6 | 6 | 3 | 3 | 3 |
| 17 | TGME49_254120-t26_1-p1 | autophagy-related protein ATG8 | 6 | 6 | 6 | 6 | 6 | 6 | 6 | 3 |
| 1 | TGME49_254180.P2049 | radial spoke protein 3 protein | 9 | 8 | 7 | 7 | 6 | 5 | 4 | 3 |
| 21 | TGME49_254190.P2050 | calcium signaling protein kinase MARK, putative | 8 | 8 | 8 | 6 | 6 | 7 | 6 | 3 |

| | | | | | | | | | | |
|---|---|---|---|---|---|---|---|---|---|---|
| 30 | TGME49_254250.P2052 | PRELI domain-containing protein PRELID | 9 | 8 | 7 | 6 | 6 | 3 | 3 | 3 |
| 30 | TGME49_254365-t26_1-p1 | phosphatidate cytidylyltransferase | 8 | 7 | 7 | 7 | 6 | 3 | 5 | 3 |
| 21 | TGME49_254380-t26_1-p1 | ribosomal protein L11, putative | 7 | 7 | 7 | 6 | 6 | 6 | 6 | 3 |
| 17 | TGME49_254440-t26_1-p1 | ribosomal protein RPL12 | 7 | 7 | 6 | 6 | 6 | 6 | 5 | 3 |
| 21 | TGME49_254450.P2057 | Pelota, putative | 9 | 8 | 8 | 6 | 6 | 6 | 6 | 3 |
| 30 | TGME49_254500-t26_1-p1 | WD repeat-containing protein | 9 | 8 | 6 | 6 | 5 | 4 | 3 | 3 |
| 17 | TGME49_254510-t26_1-p1 | ankyrin repeat-containing protein | 8 | 5 | 5 | 7 | 5 | 5 | 5 | 3 |
| 1 | TGME49_254760.P2072 | hypothetical protein | 9 | 8 | 8 | 7 | 7 | 5 | 5 | 3 |
| 21 | TGME49_254770.P2073 | shewanella-like protein phosphatase SLP | 9 | 8 | 8 | 7 | 7 | 6 | 6 | 3 |
| 30 | TGME49_254860-t26_1-p1 | hypothetical protein | 10 | 8 | 7 | 6 | 5 | 4 | 3 | 3 |
| 1 | TGME49_254870.P2078 | hypothetical protein | 9 | 8 | 8 | 7 | 7 | 5 | 5 | 3 |
| 1 | TGME49_254900.P2079 | proteasome subunit beta type-2, putative | 7 | 7 | 7 | 6 | 6 | 6 | 3 | 3 |
| 30 | TGME49_255370-t26_1-p1 | apicoplast import protein TIC20 | 9 | 7 | 6 | 6 | 5 | 3 | 4 | 3 |
| 1 | TGME49_255420.P2099 | inner membrane complex protein IMC39 | 8 | 7 | 7 | 7 | 6 | 5 | 5 | 3 |
| 1 | TGME49_255480.P2105 | thioredoxin domain-containing protein | 9 | 8 | 8 | 8 | 7 | 5 | 4 | 3 |
| 1 | TGME49_255635.P2109 | hypothetical protein | 9 | 8 | 8 | 7 | 6 | 6 | 5 | 3 |
| 1 | TGME49_255680.P2113 | YbaK/proline-tRNA ligase associated domain-containing protein | 7 | 7 | 6 | 6 | 6 | 5 | 3 | 3 |
| 17 | TGME49_255900-t26_1-p1 | Bax inhibitor-1, putative | 8 | 3 | 7 | 7 | 6 | 3 | 6 | 3 |
| 1 | TGME49_255920.P2123 | GINS complex subunit Psf3, putative | 9 | 8 | 7 | 6 | 5 | 5 | 3 | 3 |
| 21 | TGME49_256030.P2130 | doublecortin domain-containing protein DCX | 8 | 8 | 7 | 7 | 7 | 6 | 6 | 3 |
| 1 | TGME49_256810.P2139 | DnaJ domain-containing protein | 9 | 7 | 7 | 5 | 6 | 5 | 4 | 3 |
| 1 | TGME49_256900.P2146 | DEAD/DEAH box helicase domain-containing protein | 10 | 8 | 6 | 7 | 6 | 5 | 5 | 3 |
| 21 | TGME49_257040-t26_1-p1 | TB2/DP1, HVA22 family protein | 7 | 7 | 7 | 6 | 7 | 6 | 5 | 3 |
| 30 | TGME49_257060.P2152 | translation initiation factor sui1 protein | 8 | 7 | 7 | 6 | 5 | 3 | 3 | 3 |
| 30 | TGME49_257070.P2153 | Arf-GAP domain and FG repeat-containing protein AGFG | 9 | 8 | 7 | 6 | 5 | 4 | 4 | 3 |
| 30 | TGME49_257090.P2155 | NAC domain-containing protein | 7 | 7 | 7 | 6 | 6 | 3 | 3 | 3 |
| 1 | TGME49_257110.P2157 | mitochondrial inner membrane protease ATP23, putative | 9 | 8 | 7 | 7 | 6 | 6 | 5 | 3 |
| 21 | TGME49_257150.P2160 | NOT2 / NOT3 / NOT5 family protein | 9 | 8 | 8 | 7 | 7 | 6 | 5 | 3 |
| 17 | TGME49_257290.P2163 | RNA 2'-phosphotransferase, Tpt1/KptA family protein | 6 | 8 | 6 | 6 | 6 | 5 | 4 | 3 |
| 17 | TGME49_257310.P2165 | hypothetical protein | 8 | 7 | 7 | 6 | 4 | 4 | 5 | 3 |
| 17 | TGME49_257340.P2167 | ras-related protein RAB4 | 7 | 7 | 7 | 5 | 6 | 5 | 7 | 3 |
| 21 | TGME49_257350-t26_1-p1 | eukaryotic translation initiation factor, putative | 10 | 9 | 3 | 8 | 8 | 7 | 7 | 3 |
| 21 | TGME49_257440.P2174 | SCO1/SenC protein | 9 | 7 | 8 | 7 | 7 | 6 | 6 | 3 |
| 30 | TGME49_257520-t26_1-p1 | synaptobrevin protein | 8 | 7 | 7 | 6 | 5 | 4 | 4 | 3 |
| 21 | TGME49_257680.P2182 | myosin light chain MLC1 | 7 | 7 | 7 | 6 | 6 | 6 | 6 | 3 |
| 21 | TGME49_257730.P2186 | methionine aminopeptidase 1a | 9 | 8 | 8 | 7 | 6 | 7 | 5 | 3 |
| 1 | TGME49_257740.P2187 | UMP-CMP kinase | 9 | 7 | 7 | 7 | 6 | 5 | 5 | 3 |
| 30 | TGME49_257750.P2188 | homocysteine s-methyltransferase domain-containing protein | 9 | 8 | 8 | 7 | 6 | 3 | 3 | 3 |
| 30 | TGME49_258020.P2199 | glutaminyl-peptide cyclotransferase, putative | 9 | 8 | 7 | 5 | 5 | 4 | 4 | 3 |
| 21 | TGME49_258050.P2201 | actin like protein ALP2a | 9 | 9 | 8 | 7 | 7 | 6 | 6 | 3 |
| 17 | TGME49_258060.P2202 | ATP synthase subunit ATPTG8 | 7 | 6 | 7 | 6 | 5 | 5 | 4 | 3 |
| 17 | TGME49_258110.P2207 | N-acetylglucosaminylphosphatidylinositol deacetylase | 8 | 6 | 7 | 6 | 4 | 6 | 5 | 3 |

| | | | | | | | | | | |
|---|---|---|---|---|---|---|---|---|---|---|
| 21 | TGME49_258130.P2209 | ras-related protein RAB1A | 7 | 7 | 7 | 7 | 7 | 6 | 6 | 3 |
| 21 | TGME49_258150-t26_1-p1 | proteasome subunit alpha type-7, putative | 8 | 8 | 8 | 7 | 7 | 7 | 7 | 3 |
| 30 | TGME49_258160.P2210 | carbohydrate kinase | 9 | 8 | 8 | 7 | 5 | 4 | 4 | 3 |
| 1 | TGME49_258170-t26_1-p1 | hypothetical protein | 9 | 7 | 7 | 7 | 6 | 5 | 4 | 3 |
| 30 | TGME49_258380.P2218 | elongation factor P | 9 | 6 | 6 | 7 | 6 | 3 | 3 | 3 |
| 17 | TGME49_258410-t26_1-p1 | photosensitized INA-labeled protein PHIL1 | 7 | 6 | 6 | 6 | 3 | 5 | 5 | 3 |
| 30 | TGME49_258500.P2224 | ribosomal RNA-processing protein 7, putative | 8 | 8 | 7 | 6 | 5 | 4 | 4 | 3 |
| 21 | TGME49_258590.P2229 | ribonuclease Z, putative | 10 | 7 | 9 | 8 | 7 | 6 | 6 | 3 |
| 30 | TGME49_258610-t26_1-p1 | RNA recognition motif-containing protein | 9 | 8 | 7 | 6 | 6 | 3 | 4 | 3 |
| 30 | TGME49_258630.P2231 | hypothetical protein | 9 | 8 | 7 | 5 | 5 | 3 | 3 | 3 |
| 30 | TGME49_258640-t26_1-p1 | adenylate kinase superfamily protein | 9 | 7 | 7 | 6 | 5 | 4 | 3 | 3 |
| 1 | TGME49_258680.P2234 | TATA-box binding protein TBP2 | 9 | 8 | 7 | 7 | 5 | 6 | 5 | 3 |
| 1 | TGME49_258710.P2237 | spindle assembly abnormal protein 4, putative | 10 | 9 | 6 | 6 | 5 | 5 | 5 | 3 |
| 30 | TGME49_258780-t26_1-p1 | OTU domain-containing protein OTUD3A | 9 | 8 | 7 | 6 | 6 | 3 | 6 | 3 |
| 21 | TGME49_258790.P2240 | protein essential for chromosome replication ECR1 | 9 | 8 | 8 | 7 | 6 | 7 | 5 | 3 |
| 17 | TGME49_258820-t26_1-p1 | hypothetical protein | 6 | 6 | 5 | 5 | 5 | 5 | 4 | 3 |
| 17 | TGME49_258850.P2244 | hypothetical protein | 8 | 7 | 6 | 6 | 5 | 5 | 5 | 3 |
| 1 | TGME49_258920.P2250 | hypothetical protein | 8 | 7 | 7 | 6 | 6 | 5 | 3 | 3 |
| 17 | TGME49_259050.P2258 | hypothetical protein | 8 | 7 | 6 | 5 | 6 | 5 | 6 | 3 |
| 1 | TGME49_259090.P2261 | ubiquitin-conjugating enzyme subfamily protein | 9 | 7 | 6 | 6 | 6 | 5 | 5 | 3 |
| 30 | TGME49_259115.P2262 | ABC1 family protein | 9 | 8 | 6 | 5 | 5 | 4 | 4 | 3 |
| 1 | TGME49_259172.P2267 | mRNA export factor GLE1-like, putative | 9 | 3 | 8 | 7 | 7 | 6 | 4 | 3 |
| 21 | TGME49_259220.P2273 | DEAD/DEAH box helicase domain-containing protein | 10 | 8 | 7 | 7 | 7 | 7 | 6 | 3 |
| 21 | TGME49_259230.P2274 | site-specific recombinase, phage integrase family protein | 10 | 8 | 8 | 7 | 8 | 7 | 5 | 3 |
| 21 | TGME49_259250.P2275 | ATP-dependent DNA helicase, RecQ family protein | 10 | 9 | 3 | 6 | 7 | 6 | 7 | 3 |
| 17 | TGME49_259630.P2283 | inner membrane complex protein IMC27 | 7 | 6 | 6 | 5 | 5 | 4 | 5 | 3 |
| 1 | TGME49_259660-t26_1-p1 | orotate phosphoribosyltransferase | 8 | 8 | 7 | 7 | 6 | 6 | 3 | 3 |
| 1 | TGME49_259690.P2287 | orotidine-5-phosphate decarboxylase/orotate phosphoribosyltransferase | 8 | 8 | 7 | 6 | 7 | 5 | 5 | 3 |
| 30 | TGME49_259840.P2290 | hypothetical protein | 10 | 7 | 6 | 5 | 5 | 3 | 4 | 3 |
| 1 | TGME49_259850.P2291 | RNA recognition motif-containing protein | 8 | 8 | 8 | 8 | 7 | 6 | 3 | 3 |
| 21 | TGME49_259980.P2301 | DnaJ domain-containing protein | 9 | 8 | 8 | 8 | 7 | 6 | 6 | 3 |
| 17 | TGME49_260010.P2302 | acetyltransferase, GNAT family protein | 8 | 7 | 7 | 6 | 4 | 5 | 5 | 3 |
| 1 | TGME49_260160.P2307 | hypothetical protein | 10 | 8 | 6 | 6 | 5 | 5 | 4 | 3 |
| 17 | TGME49_260180-t26_1-p1 | ATP synthase-associated protein | 6 | 6 | 6 | 6 | 6 | 5 | 5 | 3 |
| 1 | TGME49_260440-t26_1-p1 | nuclear factor NF3 | 8 | 7 | 7 | 7 | 5 | 5 | 5 | 3 |
| 1 | TGME49_260510.P2327 | OTU domain-containing protein OTUB1 | 8 | 8 | 7 | 5 | 6 | 6 | 4 | 3 |
| 30 | TGME49_260530.P2329 | Sel1 repeat-containing protein | 9 | 7 | 7 | 6 | 5 | 3 | 3 | 3 |
| 1 | TGME49_260540-t26_1-p1 | alveolin domain containing intermediate filament IMC14 | 8 | 7 | 8 | 7 | 5 | 5 | 5 | 3 |
| 1 | TGME49_260610.P2333 | methyltransferase | 9 | 8 | 8 | 6 | 6 | 6 | 5 | 3 |
| 17 | TGME49_260670.P2338 | centrin 3 | 7 | 7 | 6 | 5 | 6 | 6 | 6 | 3 |
| 30 | TGME49_260810.P2342 | hypothetical protein | 8 | 7 | 7 | 6 | 5 | 3 | 3 | 3 |

| | | | | | | | | | | |
|---|---|---|---|---|---|---|---|---|---|---|
| 17 | TGME49_260820-t26_1-p1 | IMC sub-compartment protein ISP1 | 7 | 6 | 6 | 4 | 5 | 5 | 5 | 3 |
| 1 | TGME49_261200.P2354 | TBC domain-containing protein TBC8 | 10 | 8 | 8 | 6 | 7 | 6 | 3 | 3 |
| 17 | TGME49_261220.P2356 | transcription elongation factor SPT4 | 6 | 6 | 6 | 5 | 6 | 3 | 5 | 3 |
| 17 | TGME49_261240-t26_1-p1 | histone H3 | 6 | 6 | 6 | 6 | 6 | 3 | 5 | 3 |
| 17 | TGME49_261250.P2358 | histone H2A1 | 6 | 6 | 6 | 6 | 6 | 3 | 6 | 3 |
| 21 | TGME49_261370-t26_1-p1 | F-box domain-containing protein | 10 | 10 | 4 | 8 | 6 | 6 | 6 | 3 |
| 17 | TGME49_261380-t26_1-p1 | ribosomal protein S8, putative | 6 | 6 | 6 | 6 | 6 | 5 | 5 | 3 |
| 30 | TGME49_261410.P2360 | protein-tyrosine-phosphatase | 9 | 8 | 8 | 6 | 6 | 3 | 5 | 3 |
| 21 | TGME49_261440.P2362 | armadillo repeats only protein ARO | 8 | 8 | 8 | 5 | 8 | 6 | 7 | 3 |
| 1 | TGME49_261510.P2369 | hypothetical protein | 9 | 8 | 8 | 7 | 7 | 5 | 5 | 3 |
| 1 | TGME49_261710-t26_1-p1 | ankyrin repeat-containing protein | 7 | 7 | 6 | 6 | 4 | 6 | 3 | 3 |
| 1 | TGME49_261720.P2378 | metal cation transporter, ZIP family protein | 9 | 8 | 7 | 5 | 5 | 5 | 4 | 3 |
| 21 | TGME49_261780-t26_1-p1 | microneme protein MIC7 | 8 | 8 | 7 | 5 | 7 | 6 | 6 | 3 |
| 30 | TGME49_262000.P2390 | AP2 domain transcription factor AP2VIIb-2 | 8 | 7 | 6 | 6 | 5 | 4 | 4 | 3 |
| 17 | TGME49_262010-t26_1-p1 | calmodulin CAM2 | 6 | 6 | 6 | 5 | 5 | 5 | 5 | 3 |
| 1 | TGME49_262040.P2391 | SAC3/GANP family protein | 8 | 8 | 7 | 7 | 6 | 5 | 5 | 3 |
| 30 | TGME49_262080.P2393 | nucleus export protein BRR6, putative | 10 | 8 | 7 | 6 | 5 | 4 | 3 | 3 |
| 1 | TGME49_262410-t26_1-p1 | WD domain, G-beta repeat-containing protein | 10 | 7 | 6 | 6 | 6 | 5 | 5 | 3 |
| 17 | TGME49_262480.P2407 | dynein light chain roadblock-type 2, putative | 5 | 5 | 5 | 5 | 5 | 5 | 5 | 3 |
| 30 | TGME49_262490.P2408 | serine hydrolase ASH4 | 8 | 8 | 7 | 7 | 7 | 3 | 5 | 3 |
| 1 | TGME49_262520.P2410 | cyclophilin, putative | 8 | 8 | 8 | 6 | 5 | 5 | 3 | 3 |
| 21 | TGME49_262530.P2411 | leucine rich repeat-containing protein | 9 | 8 | 7 | 6 | 5 | 6 | 6 | 3 |
| 17 | TGME49_262640.P2416 | cytochrome c oxidase subunit ApiCOX23 | 7 | 7 | 7 | 6 | 6 | 5 | 5 | 3 |
| 21 | TGME49_262670.P2419 | ribosomal protein RPL18A | 7 | 7 | 7 | 7 | 7 | 6 | 6 | 3 |
| 17 | TGME49_262690.P2422 | ribosomal protein RPL27 | 6 | 6 | 6 | 6 | 6 | 5 | 5 | 3 |
| 17 | TGME49_262720-t26_1-p1 | eukaryotic initiation factor-1A, putative | 6 | 6 | 6 | 6 | 6 | 6 | 5 | 3 |
| 1 | TGME49_262850.P2429 | histone methyltransferase complex subunit ASH2, putative | 10 | 8 | 7 | 6 | 6 | 5 | 5 | 3 |
| 21 | TGME49_262910.P2431 | NADH-cytochrome b5 reductase 1, putative | 8 | 8 | 7 | 7 | 6 | 6 | 6 | 3 |
| 1 | TGME49_262940-t26_1-p1 | aspartyl protease ASP2 | 9 | 9 | 9 | 5 | 7 | 6 | 3 | 3 |
| 17 | TGME49_263040.P2439 | ribosomal protein RPS16 | 6 | 6 | 6 | 6 | 6 | 6 | 6 | 3 |
| 21 | TGME49_263050.P2440 | ribosomal protein RPL13 | 7 | 7 | 7 | 7 | 7 | 7 | 6 | 3 |
| 30 | TGME49_263150.P2449 | tetratricopeptide repeat-containing protein | 8 | 8 | 8 | 7 | 5 | 3 | 5 | 3 |
| 1 | TGME49_263290-t26_1-p1 | rhomboid protease ROM2 | 8 | 8 | 7 | 7 | 7 | 5 | 3 | 3 |
| 21 | TGME49_263300.P2459 | voltage-dependent anion-selective channel protein | 8 | 8 | 8 | 8 | 7 | 5 | 6 | 3 |
| 1 | TGME49_263310-t26_1-p1 | step ii splicing factor, putative | 8 | 7 | 6 | 6 | 5 | 5 | 4 | 3 |
| 17 | TGME49_263323.P2460 | mitochondrial fission 1 protein | 6 | 6 | 6 | 6 | 6 | 5 | 5 | 3 |
| 1 | TGME49_263380.P2465 | Dullard family phosphatase domain-containing protein | 8 | 8 | 7 | 7 | 5 | 5 | 5 | 3 |
| 30 | TGME49_263430.P2468 | 3-hydroxyisobutyrate dehydrogenase | 8 | 8 | 8 | 6 | 6 | 3 | 6 | 3 |
| 1 | TGME49_263470-t26_1-p1 | ubiquitin carboxyl-terminal hydrolase UCHL3 | 8 | 7 | 7 | 6 | 6 | 5 | 3 | 3 |
| 1 | TGME49_263490-t26_1-p1 | ubiquitin conjugating enzyme E2, putative | 7 | 7 | 6 | 5 | 6 | 6 | 3 | 3 |
| 21 | TGME49_263500.P2472 | vacuolar protein sorting-associated protein 26, putative | 8 | 8 | 8 | 6 | 7 | 7 | 7 | 3 |
| 21 | TGME49_263550-t26_1-p1 | 39S ribosomal protein L47, mitochondrial precursor, putative | 8 | 7 | 7 | 6 | 7 | 6 | 6 | 3 |
| 30 | TGME49_263570.P2477 | hypothetical protein | 9 | 8 | 8 | 7 | 6 | 3 | 3 | 3 |
| 30 | TGME49_263595.P2481 | RNA-binding protein | 8 | 7 | 7 | 7 | 6 | 3 | 6 | 3 |

| | | | | | | | | | | |
|---|---|---|---|---|---|---|---|---|---|---|
| 17 | TGME49_263680.P2487 | hypothetical protein | 7 | 7 | 6 | 6 | 5 | 5 | 5 | 3 |
| 17 | TGME49_263700-t26_1-p1 | ribosomal protein RPS14 | 6 | 6 | 6 | 6 | 6 | 6 | 5 | 3 |
| 17 | TGME49_263850.P2497 | hypothetical protein | 7 | 7 | 7 | 5 | 6 | 5 | 5 | 3 |
| 1 | TGME49_263860-t26_1-p1 | telomere length and silencing protein 1, putative | 8 | 7 | 7 | 6 | 6 | 5 | 5 | 3 |
| 17 | TGME49_264040.P2503 | cytochrome c oxidase subunit ApiCOX25 | 7 | 7 | 7 | 6 | 5 | 3 | 6 | 3 |
| 30 | TGME49_264050.P2504 | fused toes protein homolog FTS | 8 | 8 | 8 | 7 | 6 | 4 | 3 | 3 |
| 17 | TGME49_264080-t26_1-p1 | acyl carrier protein ACP | 7 | 7 | 5 | 5 | 5 | 3 | 5 | 3 |
| 1 | TGME49_264130.P2507 | pre-mRNA-splicing factor CWC26, putative | 9 | 8 | 8 | 6 | 5 | 5 | 5 | 3 |
| 1 | TGME49_264440-t26_1-p1 | signal recognition particle receptor beta subunit protein | 8 | 7 | 7 | 6 | 6 | 5 | 5 | 3 |
| 30 | TGME49_264770-t26_1-p1 | hypothetical protein | 8 | 8 | 7 | 6 | 5 | 4 | 4 | 3 |
| 21 | TGME49_265010.P2530 | glutamate 5-kinase domain-containing protein | 10 | 8 | 7 | 7 | 7 | 6 | 5 | 3 |
| 21 | TGME49_265020-t26_1-p1 | arabinogalactan protein, putative | 7 | 7 | 7 | 7 | 6 | 6 | 6 | 3 |
| 1 | TGME49_265220-t26_1-p1 | co-chaperone GrpE protein | 8 | 7 | 7 | 6 | 5 | 5 | 3 | 3 |
| 21 | TGME49_265360-t26_1-p1 | hypothetical protein | 9 | 7 | 9 | 8 | 7 | 6 | 5 | 3 |
| 21 | TGME49_265380-t26_1-p1 | tetratricopeptide repeat (TPR)-/ U-box domain-containing protein | 8 | 7 | 7 | 7 | 7 | 6 | 5 | 3 |
| 21 | TGME49_265390-t26_1-p1 | CTLH domain-containing protein | 9 | 8 | 8 | 7 | 8 | 6 | 6 | 3 |
| 21 | TGME49_265420-t26_1-p1 | hypothetical protein | 8 | 7 | 8 | 7 | 7 | 6 | 6 | 3 |
| 30 | TGME49_265440-t26_1-p1 | hypothetical protein | 10 | 8 | 7 | 6 | 5 | 4 | 3 | 3 |
| 1 | TGME49_265470.P2539 | hypothetical protein | 8 | 8 | 8 | 7 | 6 | 6 | 4 | 3 |
| 1 | TGME49_265500.P2540 | chloride transporter, chloride channel (ClC) family protein | 10 | 9 | 5 | 7 | 7 | 5 | 5 | 3 |
| 30 | TGME49_265530.P2542 | RNA recognition motif-containing protein | 9 | 8 | 7 | 6 | 6 | 4 | 4 | 3 |
| 1 | TGME49_265650-t26_1-p1 | protein phosphatase 2C domain-containing protein | 8 | 9 | 8 | 7 | 5 | 6 | 4 | 3 |
| 30 | TGME49_265770.P2543 | apicomplexan kinetochore protein AKIT5, putative | 10 | 7 | 8 | 5 | 5 | 3 | 3 | 3 |
| 1 | TGME49_265780.P2544 | flagellar/basal body protein | 10 | 8 | 7 | 6 | 6 | 5 | 5 | 3 |
| 21 | TGME49_265790.P2545 | claudin-like apicomplexan microneme protein | 9 | 8 | 8 | 8 | 7 | 6 | 6 | 3 |
| 30 | TGME49_265860.P2551 | ribonuclease ZC3H12D, putative | 9 | 8 | 8 | 7 | 6 | 3 | 3 | 3 |
| 21 | TGME49_266030.P2554 | pre-mRNA splicing factor, putative | 9 | 8 | 4 | 7 | 7 | 6 | 6 | 3 |
| 21 | TGME49_266090.P2558 | transmembrane protein 43, putative | 9 | 8 | 8 | 8 | 8 | 6 | 6 | 3 |
| 30 | TGME49_266140-t26_1-p1 | peptidase, M50 family protein | 9 | 7 | 8 | 7 | 5 | 3 | 4 | 3 |
| 1 | TGME49_266260-t26_1-p1 | hypothetical protein | 8 | 7 | 7 | 6 | 6 | 5 | 5 | 3 |
| 30 | TGME49_266372-t26_1-p1 | hypothetical protein | 10 | 7 | 7 | 6 | 5 | 4 | 4 | 3 |
| 17 | TGME49_266470-t26_1-p1 | hypothetical protein | 6 | 6 | 6 | 5 | 5 | 5 | 4 | 3 |
| 17 | TGME49_266620.P2572 | thioredoxin CTRX1 | 7 | 7 | 6 | 6 | 5 | 4 | 4 | 3 |
| 21 | TGME49_266750-t26_1-p1 | transporter/permease protein, putative | 9 | 8 | 8 | 7 | 7 | 7 | 6 | 3 |
| 17 | TGME49_266760.P2581 | isocitrate dehydrogenase 1 | 3 | 3 | 3 | 3 | 3 | 8 | 8 | 3 |
| 30 | TGME49_266810.P2583 | peroxisome assembly protein 22, putative | 8 | 7 | 6 | 5 | 6 | 4 | 4 | 3 |
| 30 | TGME49_266850-t26_1-p1 | 3-demethylubiquinone-9 3-O-methyltransferase | 9 | 7 | 7 | 7 | 4 | 3 | 4 | 3 |
| 21 | TGME49_266880-t26_1-p1 | dihydrouridine synthase, putative | 9 | 8 | 8 | 7 | 7 | 7 | 6 | 3 |
| 1 | TGME49_266930.P2587 | general transcription factor IIH polypeptide 3 GTF2H3 | 9 | 8 | 8 | 7 | 6 | 6 | 5 | 3 |
| 1 | TGME49_266970.P2591 | ubiquitin family protein | 8 | 8 | 7 | 6 | 6 | 5 | 4 | 3 |

| | | | | | | | | | | |
|---|---|---|---|---|---|---|---|---|---|---|
| 30 | TGME49_267050-t26_1-p1 | hydrolase, alpha/beta fold family protein | 8 | 8 | 8 | 5 | 7 | 3 | 3 | 3 |
| 17 | TGME49_267060.P2595 | ribosomal protein RPL14 | 6 | 6 | 6 | 6 | 6 | 5 | 5 | 3 |
| 30 | TGME49_267360-t26_1-p1 | histone deacetylase SIR2-like | 10 | 8 | 8 | 5 | 6 | 3 | 5 | 3 |
| 17 | TGME49_267420-t26_1-p1 | mago nashi family protein 2, putative | 6 | 6 | 3 | 6 | 3 | 6 | 6 | 3 |
| 1 | TGME49_267440-t26_1-p1 | RING zinc finger protein | 9 | 8 | 8 | 7 | 5 | 5 | 3 | 3 |
| 30 | TGME49_267500.P2603 | conserved basal apical peripheral protein CBAP | 8 | 7 | 7 | 7 | 6 | 3 | 6 | 3 |
| 17 | TGME49_267510-t26_1-p1 | hypothetical protein | 8 | 7 | 6 | 5 | 5 | 5 | 5 | 3 |
| 30 | TGME49_267530.P2604 | syntaxin-18 | 9 | 7 | 7 | 5 | 5 | 3 | 3 | 3 |
| 21 | TGME49_267550-t26_1-p1 | leucine-rich repeat protein LRR1 | 8 | 8 | 8 | 7 | 6 | 5 | 6 | 3 |
| 21 | TGME49_267720.P2612 | transporter, cation channel family protein | 10 | 3 | 9 | 8 | 8 | 7 | 7 | 3 |
| 30 | TGME49_267760-t26_1-p1 | zinc finger CCHC domain-containing protein | 10 | 7 | 7 | 5 | 5 | 4 | 4 | 3 |
| 30 | TGME49_267775-t26_1-p1 | hypothetical protein | 10 | 8 | 8 | 6 | 5 | 3 | 3 | 3 |
| 21 | TGME49_267810-t26_1-p1 | ras-related protein RAB5A | 8 | 7 | 7 | 7 | 7 | 7 | 6 | 3 |
| 1 | TGME49_267840.P2618 | ER membrane protein complex subunit 2, putative | 8 | 7 | 7 | 6 | 5 | 5 | 4 | 3 |
| 30 | TGME49_268350-t26_1-p1 | WD repeat-containing protein | 9 | 8 | 7 | 5 | 5 | 3 | 4 | 3 |
| 1 | TGME49_268620-t26_1-p1 | blood stage antigen 41-3 precursor, putative | 9 | 8 | 8 | 6 | 4 | 6 | 3 | 3 |
| 1 | TGME49_268690.P2644 | OTU domain-containing protein OTU10 | 8 | 8 | 7 | 7 | 5 | 5 | 4 | 3 |
| 1 | TGME49_268720-t26_1-p1 | Hrf1 family protein | 9 | 8 | 8 | 7 | 6 | 5 | 5 | 3 |
| 17 | TGME49_268740-t26_1-p1 | hypothetical protein | 7 | 6 | 6 | 5 | 5 | 5 | 4 | 3 |
| 17 | TGME49_268750.P2646 | peptidyl-prolyl cis-trans isomerase E, putative | 6 | 6 | 6 | 6 | 6 | 5 | 5 | 3 |
| 21 | TGME49_268810.P2649 | ck2 beta subunit | 8 | 8 | 7 | 6 | 7 | 6 | 6 | 3 |
| 21 | TGME49_268840.P2652 | N6-adenosine-methyltransferase non-catalytic subunit METTL14 | 9 | 9 | 9 | 6 | 6 | 6 | 7 | 3 |
| 1 | TGME49_268960-t26_1-p1 | 5'-AMP-activated protein kinase subunit beta | 9 | 8 | 7 | 5 | 6 | 7 | 3 | 3 |
| 21 | TGME49_269140.P2666 | trafficking protein particle complex subunit 5, putative | 7 | 7 | 7 | 7 | 7 | 7 | 6 | 3 |
| 17 | TGME49_269410-t26_1-p1 | hypothetical protein | 7 | 7 | 7 | 6 | 6 | 5 | 5 | 3 |
| 21 | TGME49_269430.P2682 | heptaprenyl diphosphate synthase COQ1 | 9 | 8 | 8 | 8 | 6 | 6 | 6 | 3 |
| 17 | TGME49_269442.P2683 | essential light chain ELC1 | 5 | 6 | 5 | 5 | 5 | 5 | 5 | 3 |
| 30 | TGME49_269650.P2692 | FFD and TFG box motifs protein | 9 | 8 | 8 | 7 | 5 | 4 | 4 | 3 |
| 30 | TGME49_269830.P2704 | RAP domain-binding protein RA15 | 10 | 8 | 7 | 5 | 5 | 3 | 4 | 3 |
| 21 | TGME49_269850.P2705 | protein c21orf59, putative | 8 | 3 | 8 | 7 | 7 | 6 | 6 | 3 |
| 30 | TGME49_269960.P2711 | inner membrane complex protein IMC40 | 9 | 8 | 7 | 6 | 5 | 4 | 4 | 3 |
| 1 | TGME49_270010.P2714 | hypothetical protein | 9 | 7 | 7 | 5 | 6 | 5 | 4 | 3 |
| 21 | TGME49_270070.P2717 | protein transport protein SEC22, putative | 8 | 7 | 7 | 7 | 6 | 6 | 6 | 3 |
| 17 | TGME49_270140.P2721 | splicing factor DIM1, putative | 6 | 6 | 6 | 6 | 6 | 4 | 6 | 3 |
| 1 | TGME49_270270-t26_1-p1 | TMEM33 domain-containing protein | 9 | 8 | 8 | 7 | 5 | 5 | 5 | 3 |
| 21 | TGME49_270273.P2730 | hypothetical protein | 7 | 3 | 7 | 8 | 7 | 6 | 6 | 3 |
| 21 | TGME49_270320.P2732 | protein phosphatase PPM3C | 9 | 8 | 8 | 7 | 6 | 6 | 6 | 3 |
| 17 | TGME49_270370.P2736 | AP-1 complex subunit sigma, putative | 7 | 7 | 6 | 6 | 6 | 3 | 6 | 3 |
| 17 | TGME49_270380-t26_1-p1 | ribosomal protein RPS13 | 6 | 6 | 6 | 6 | 6 | 6 | 6 | 3 |
| 17 | TGME49_270540.P2771 | Trm112p family domain-containing protein | 6 | 6 | 6 | 5 | 5 | 5 | 4 | 3 |
| 17 | TGME49_270560.P2773 | peptidyl-prolyl cis-trans isomerase family 1 | 7 | 7 | 7 | 6 | 5 | 4 | 6 | 3 |
| 17 | TGME49_270780-t26_1-p1 | DNA-directed RNA polymerase II RPB6 | 6 | 6 | 6 | 3 | 5 | 5 | 5 | 3 |
| 21 | TGME49_270800-t26_1-p1 | GAF domain-containing protein | 9 | 8 | 8 | 5 | 5 | 7 | 7 | 3 |
| 17 | TGME49_270830.P2785 | small nuclear ribonucleoprotein | 5 | 5 | 5 | 5 | 5 | 5 | 5 | 3 |
| 30 | TGME49_270950-t26_1-p1 | hypothetical protein | 9 | 8 | 8 | 7 | 6 | 3 | 3 | 3 |
| 17 | TGME49_271070-t26_1-p1 | OTU domain-containing protein OTUD6A | 8 | 7 | 5 | 6 | 5 | 4 | 5 | 3 |

| | | | | | | | | | | |
|---|---|---|---|---|---|---|---|---|---|---|
| 21 | TGME49_271280.P2807 | 60S ribosome subunit biogenesis protein NIP7, putative | 7 | 7 | 7 | 7 | 7 | 6 | 6 | 3 |
| 21 | TGME49_271300-t26_1-p1 | DNA-directed RNA polymerase II RPB7 | 7 | 7 | 7 | 7 | 7 | 6 | 6 | 3 |
| 21 | TGME49_271350.P2810 | bifunctional protein FolC subfamily protein | 9 | 9 | 9 | 7 | 7 | 6 | 6 | 3 |
| 30 | TGME49_271610.P2819 | pyrroline-5-carboxylate reductase | 8 | 8 | 8 | 7 | 7 | 3 | 3 | 3 |
| 1 | TGME49_271880.P2826 | preconoidal ring protein PCR7 | 10 | 10 | 4 | 6 | 8 | 6 | 3 | 3 |
| 1 | TGME49_271888.P2827 | 3-ketoacyl-CoA reductase, putative | 9 | 8 | 8 | 6 | 7 | 5 | 5 | 3 |
| 17 | TGME49_272010.P2836 | Gar1 protein RNA binding region protein | 7 | 5 | 6 | 5 | 5 | 5 | 4 | 3 |
| 1 | TGME49_272230.P2844 | heat shock protein hslv, putative | 8 | 7 | 7 | 7 | 7 | 5 | 5 | 3 |
| 1 | TGME49_272260.P2845 | Dpy-30 motif protein | 10 | 8 | 7 | 7 | 5 | 5 | 4 | 3 |
| 21 | TGME49_272350.P2853 | mRNA export protein | 9 | 8 | 9 | 6 | 7 | 7 | 6 | 3 |
| 30 | TGME49_272360.P2854 | EF hand family protein | 9 | 3 | 9 | 9 | 7 | 3 | 5 | 3 |
| 21 | TGME49_272400.P2859 | casein kinase ii regulatory subunit protein | 9 | 8 | 8 | 7 | 7 | 7 | 6 | 3 |
| 30 | TGME49_272695.P2880 | hypothetical protein | 9 | 8 | 6 | 6 | 5 | 3 | 3 | 3 |
| 30 | TGME49_272760.P2885 | hypothetical protein | 9 | 7 | 6 | 7 | 4 | 4 | 4 | 3 |
| 17 | TGME49_273060-t26_1-p1 | ribosomal protein S17, putative | 6 | 6 | 6 | 6 | 5 | 5 | 5 | 3 |
| 21 | TGME49_273070.P2891 | GTPase activating protein for adp ribosylation factor, putative | 9 | 8 | 8 | 7 | 6 | 6 | 6 | 3 |
| 1 | TGME49_273390.P2901 | mitochondrial carrier superfamily protein | 9 | 8 | 7 | 5 | 6 | 6 | 4 | 3 |
| 30 | TGME49_273530-t26_1-p1 | flagellar associated protein | 8 | 7 | 7 | 7 | 6 | 3 | 5 | 3 |
| 21 | TGME49_273905.P2925 | armadillo repeat-containing protein | 9 | 8 | 8 | 6 | 6 | 6 | 6 | 3 |
| 21 | TGME49_273920.P2926 | aldose reductase, putative | 9 | 9 | 8 | 7 | 7 | 6 | 6 | 3 |
| 1 | TGME49_273960.P2928 | chaperonin GroS protein | 8 | 7 | 7 | 7 | 6 | 5 | 5 | 3 |
| 1 | TGME49_273990-t26_1-p1 | CobW/P47K family protein | 9 | 8 | 7 | 6 | 5 | 5 | 5 | 3 |
| 1 | TGME49_274050.P2934 | hypothetical protein | 8 | 8 | 7 | 6 | 5 | 5 | 3 | 3 |
| 1 | TGME49_275310.P2947 | U4/U6.U5 small nuclear ribonucleoprotein component Snu23, putative | 9 | 8 | 7 | 7 | 6 | 5 | 5 | 3 |
| 17 | TGME49_275590.P2955 | mono- or diacylglycerol acyltransferase | 6 | 6 | 7 | 7 | 3 | 5 | 5 | 3 |
| 17 | TGME49_275750.P2961 | small nuclear ribonucleoprotein E, putative | 5 | 5 | 5 | 5 | 5 | 5 | 5 | 3 |
| 30 | TGME49_275790-t26_1-p1 | EGF family domain-containing protein | 9 | 8 | 7 | 7 | 5 | 3 | 3 | 3 |
| 17 | TGME49_275810-t26_1-p1 | ribosomal protein RPS10 | 6 | 6 | 6 | 5 | 5 | 5 | 4 | 3 |
| 21 | TGME49_275980-t26_1-p1 | coenzyme q (ubiquinone) biosynthesis protein coq4 protein | 9 | 8 | 7 | 6 | 6 | 5 | 6 | 3 |
| 1 | TGME49_276140.P2966 | ADP ribosylation factor ARF1 | 7 | 7 | 7 | 7 | 6 | 7 | 3 | 3 |
| 1 | TGME49_276155.P2967 | acyl-CoA synthetase ACS7, putative | 10 | 8 | 7 | 8 | 6 | 7 | 3 | 3 |
| 30 | TGME49_276210.P2969 | phosphoglycerate mutase family protein | 10 | 8 | 7 | 7 | 5 | 3 | 4 | 3 |
| 30 | TGME49_276900-t26_1-p1 | hypothetical protein | 10 | 7 | 8 | 8 | 5 | 4 | 4 | 3 |
| 1 | TGME49_276920.P2973 | protein phosphatase PPM3B, putative | 9 | 8 | 8 | 8 | 6 | 5 | 4 | 3 |
| 30 | TGME49_276970-t26_1-p1 | hypothetical protein | 9 | 8 | 8 | 7 | 6 | 4 | 4 | 3 |
| 1 | TGME49_277690-t26_1-p1 | hypothetical protein | 9 | 7 | 7 | 6 | 5 | 5 | 3 | 3 |
| 17 | TGME49_277700.P2984 | ribosomal protein S14 precursor, putative | 6 | 6 | 6 | 5 | 5 | 5 | 5 | 3 |
| 30 | TGME49_277740-t26_1-p1 | zinc finger, C3HC4 type (RING finger) domain-containing protein | 10 | 8 | 7 | 6 | 5 | 3 | 3 | 3 |
| 17 | TGME49_277770-t26_1-p1 | zinc finger (CCCH type) motif-containing protein | 7 | 5 | 6 | 6 | 5 | 5 | 5 | 3 |
| 30 | TGME49_277820.P2986 | hypothetical protein | 10 | 8 | 8 | 7 | 5 | 3 | 3 | 3 |
| 1 | TGME49_277840-t26_1-p1 | Ras family protein | 8 | 7 | 7 | 7 | 6 | 5 | 5 | 3 |
| 1 | TGME49_277895.P2988 | ubiquitin carboxyl-terminal hydrolase UBP1 | 10 | 8 | 4 | 7 | 7 | 6 | 3 | 3 |
| 30 | TGME49_277970.P2992 | dolichol-phosphate-mannose synthase family protein | 8 | 8 | 7 | 7 | 6 | 3 | 3 | 3 |

| | | | | | | | | | | |
|---|---|---|---|---|---|---|---|---|---|---|
| 1 | TGME49_277990.P2993 | OTU domain-containing protein OTUD2 | 9 | 8 | 7 | 7 | 6 | 5 | 5 | 3 |
| 30 | TGME49_278170-t26_1-p1 | hypothetical protein | 8 | 9 | 7 | 7 | 4 | 3 | 5 | 3 |
| 30 | TGME49_278220.P2997 | ribosomal protein L46, putative | 9 | 8 | 7 | 7 | 4 | 3 | 5 | 3 |
| 21 | TGME49_278470-t26_1-p1 | START domain-containing protein | 9 | 8 | 8 | 5 | 6 | 6 | 6 | 3 |
| 1 | TGME49_278490-t26_1-p1 | Zn-finger, RING domain containing protein | 8 | 8 | 8 | 7 | 6 | 6 | 4 | 3 |
| 17 | TGME49_278530-t26_1-p1 | multiprotein bridging factor type 1 family transcriptional co-activator, putative | 6 | 6 | 6 | 6 | 6 | 6 | 5 | 3 |
| 30 | TGME49_278730-t26_1-p1 | guanine nucleotide-binding protein, putative | 10 | 7 | 8 | 7 | 6 | 4 | 4 | 3 |
| 21 | TGME49_278850.P3020 | palmitoyltransferase DHHC2 | 9 | 8 | 8 | 6 | 7 | 7 | 5 | 3 |
| 30 | TGME49_278930.P3023 | Tubulin-tyrosine ligase family protein | 9 | 7 | 9 | 6 | 7 | 3 | 4 | 3 |
| 1 | TGME49_279360-t26_1-p1 | hypothetical protein | 9 | 9 | 8 | 6 | 6 | 5 | 4 | 3 |
| 17 | TGME49_279370-t26_1-p1 | SNARE associated Golgi protein | 3 | 8 | 8 | 7 | 5 | 4 | 4 | 3 |
| 17 | TGME49_280440.P3039 | ubiquitin conjugating enzyme E2, putative | 7 | 6 | 6 | 6 | 6 | 6 | 5 | 3 |
| 21 | TGME49_280480.P3041 | EF hand domain-containing protein | 8 | 8 | 8 | 8 | 7 | 6 | 7 | 3 |
| 17 | TGME49_280550.P3044 | clathrin adaptor complex small chain subfamily protein | 7 | 7 | 6 | 6 | 5 | 5 | 4 | 3 |
| 1 | TGME49_280670.P3048 | hypothetical protein | 9 | 8 | 8 | 6 | 6 | 5 | 4 | 3 |
| 1 | TGME49_280710.P3051 | proteasome subunit beta type-4, putative | 8 | 8 | 7 | 7 | 7 | 6 | 3 | 3 |
| 21 | TGME49_280730-t26_1-p1 | cytosolic fe-s cluster assembling factor nbp35, putative | 8 | 8 | 8 | 7 | 7 | 6 | 6 | 3 |
| 17 | TGME49_280740.P3053 | signal peptidase | 7 | 7 | 7 | 6 | 6 | 5 | 6 | 3 |
| 17 | TGME49_280760-t26_1-p1 | RUB1 conjugating enzyme | 7 | 6 | 6 | 5 | 5 | 5 | 5 | 3 |
| 21 | TGME49_280780-t26_1-p1 | dihydrouridine synthase (dus) protein | 9 | 9 | 8 | 7 | 7 | 6 | 6 | 3 |
| 30 | TGME49_281380.P3056 | hypothetical protein | 10 | 9 | 6 | 6 | 5 | 4 | 4 | 3 |
| 1 | TGME49_281420.P3058 | histone deacetylase HDAC1 | 10 | 9 | 7 | 7 | 5 | 6 | 4 | 3 |
| 30 | TGME49_281490-t26_1-p1 | pyridoxal 5'-phosphate synthase subunit PDX2 | 8 | 7 | 7 | 6 | 5 | 3 | 3 | 3 |
| 1 | TGME49_281510-t26_1-p1 | ribonuclease H1 large subunit, putative | 8 | 6 | 8 | 5 | 6 | 6 | 5 | 3 |
| 21 | TGME49_281520.P3063 | zinc finger, C3HC4 type (RING finger) domain-containing protein | 8 | 8 | 8 | 5 | 6 | 7 | 6 | 3 |
| 21 | TGME49_281620-t26_1-p1 | aldo/keto reductase family oxidoreductase | 9 | 7 | 8 | 6 | 6 | 5 | 6 | 3 |
| 21 | TGME49_281920-t26_1-p1 | V-type ATPase, D subunit protein | 8 | 8 | 7 | 7 | 7 | 6 | 6 | 3 |
| 1 | TGME49_281950-t26_1-p1 | membrane protein, putative | 8 | 8 | 8 | 6 | 6 | 5 | 5 | 3 |
| 30 | TGME49_281970-t26_1-p1 | adenylate kinase superfamily protein | 8 | 8 | 7 | 6 | 6 | 3 | 4 | 3 |
| 30 | TGME49_281990.P3076 | Nicotinamidase | 9 | 8 | 7 | 7 | 5 | 4 | 4 | 3 |
| 30 | TGME49_282170-t26_1-p1 | hypothetical protein | 9 | 8 | 7 | 6 | 5 | 3 | 3 | 3 |
| 1 | TGME49_282180-t26_1-p1 | ATP synthase-associated protein | 8 | 8 | 8 | 6 | 6 | 6 | 5 | 3 |
| 1 | TGME49_283590.P3094 | mitochondrial import inner membrane translocase subunit TIM50 | 9 | 8 | 8 | 7 | 7 | 5 | 5 | 3 |
| 21 | TGME49_283710.P3095 | ceramide synthase CERS1, putative | 9 | 8 | 8 | 8 | 8 | 6 | 6 | 3 |
| 30 | TGME49_283860-t26_1-p1 | leucine rich repeat-containing protein | 10 | 8 | 7 | 7 | 5 | 3 | 5 | 3 |
| 1 | TGME49_284530.P3107 | hypothetical protein | 10 | 9 | 6 | 7 | 5 | 5 | 5 | 3 |
| 1 | TGME49_284540-t26_1-p1 | ATP synthase subunit O | 8 | 7 | 7 | 6 | 6 | 5 | 5 | 3 |
| 21 | TGME49_284550.P3108 | EF hand domain-containing protein | 8 | 3 | 8 | 9 | 6 | 5 | 6 | 3 |
| 21 | TGME49_284560-t26_1-p1 | ribosomal protein RPL9 | 7 | 7 | 7 | 7 | 6 | 6 | 6 | 3 |
| 21 | TGME49_284570-t26_1-p1 | hypothetical protein | 8 | 8 | 7 | 7 | 6 | 6 | 6 | 3 |
| 21 | TGME49_285268.P3128 | DnaJ domain-containing protein | 9 | 8 | 8 | 8 | 7 | 6 | 6 | 3 |
| 30 | TGME49_285272-t26_1-p1 | FAS1 domain-containing protein | 9 | 7 | 7 | 6 | 5 | 4 | 3 | 3 |
| 17 | TGME49_285350.P3129 | dynein light chain, putative | 5 | 5 | 6 | 6 | 6 | 6 | 6 | 3 |
| 17 | TGME49_285510-t26_1-p1 | ATP synthase subunit ATPTG9 | 7 | 6 | 6 | 6 | 5 | 5 | 5 | 3 |

| | | | | | | | | | | |
|---|---|---|---|---|---|---|---|---|---|---|
| 1 | TGME49_285670-t26_1-p1 | hypothetical protein | 8 | 7 | 7 | 7 | 7 | 6 | 3 | 3 |
| 30 | TGME49_285700-t26_1-p1 | ubiquitin fusion degradation protein UFD1AP | 9 | 7 | 7 | 6 | 6 | 3 | 5 | 3 |
| 21 | TGME49_285720-t26_1-p1 | ATP binding protein, putative | 8 | 8 | 8 | 8 | 8 | 7 | 6 | 3 |
| 1 | TGME49_285730-t26_1-p1 | TBC domain-containing protein TBC4 | 9 | 8 | 8 | 7 | 5 | 7 | 5 | 3 |
| 30 | TGME49_285750.P3136 | 6,7-dihydropteridine reductase | 8 | 7 | 6 | 6 | 5 | 3 | 3 | 3 |
| 30 | TGME49_285760-t26_1-p1 | U-snRNP-associated cyclophilin family protein | 7 | 7 | 7 | 7 | 7 | 4 | 4 | 3 |
| 1 | TGME49_285850-t26_1-p1 | peptidyl-prolyl cis-trans isomerase, FKBP-type domain-containing protein | 9 | 8 | 7 | 6 | 5 | 5 | 4 | 3 |
| 30 | TGME49_286050.P3145 | apicoplast TIC22 protein | 9 | 8 | 7 | 6 | 5 | 4 | 3 | 3 |
| 17 | TGME49_286090.P3147 | translation initiation factor SUI1, putative | 6 | 6 | 6 | 5 | 6 | 5 | 5 | 3 |
| 1 | TGME49_286430-t26_1-p1 | FAM221 domain-containing protein | 8 | 7 | 7 | 7 | 5 | 5 | 5 | 3 |
| 30 | TGME49_286550.P3156 | exosome complex component RRP43, putative | 8 | 8 | 7 | 6 | 6 | 3 | 4 | 3 |
| 30 | TGME49_286630.P3159 | alkyl hydroperoxide reductase AHP1 | 8 | 7 | 7 | 6 | 5 | 3 | 3 | 3 |
| 21 | TGME49_286640-t26_1-p1 | GTPase | 10 | 6 | 8 | 8 | 7 | 7 | 7 | 3 |
| 30 | TGME49_286680-t26_1-p1 | hypothetical protein | 9 | 7 | 7 | 6 | 5 | 3 | 3 | 3 |
| 21 | TGME49_286710.P3161 | zinc finger protein ZNF2 | 8 | 8 | 8 | 7 | 7 | 6 | 6 | 3 |
| 30 | TGME49_286720-t26_1-p1 | heat shock protein HSP28 | 8 | 7 | 7 | 6 | 5 | 4 | 4 | 3 |
| 17 | TGME49_286730.P3162 | signal recognition particle subunit SRP19, putative | 7 | 6 | 6 | 6 | 5 | 5 | 5 | 3 |
| 21 | TGME49_286790-t26_1-p1 | nuclear factor NF2 | 8 | 7 | 7 | 6 | 6 | 6 | 6 | 3 |
| 21 | TGME49_287210.P3170 | proteasome subunit alpha type-2, putative | 8 | 8 | 7 | 8 | 8 | 6 | 7 | 3 |
| 1 | TGME49_287270.P3172 | peripheral plastid protein 1 | 10 | 7 | 7 | 5 | 5 | 5 | 4 | 3 |
| 1 | TGME49_288260-t26_1-p1 | ADP-ribosylation factor-like protein ArlX3 | 9 | 8 | 6 | 6 | 5 | 5 | 4 | 3 |
| 1 | TGME49_288340-t26_1-p1 | UBX domain-containing protein | 9 | 7 | 7 | 6 | 6 | 5 | 4 | 3 |
| 30 | TGME49_288510-t26_1-p1 | ubiquitin carboxyl-terminal hydrolase | 10 | 6 | 8 | 6 | 5 | 3 | 3 | 3 |
| 21 | TGME49_288570-t26_1-p1 | hypothetical protein | 9 | 8 | 7 | 7 | 6 | 6 | 6 | 3 |
| 1 | TGME49_288610-t26_1-p1 | 50S ribosomal protein L21, putative | 9 | 7 | 7 | 7 | 6 | 6 | 5 | 3 |
| 21 | TGME49_288630.P3185 | nucleosome assembly protein (NAP), putative | 8 | 7 | 7 | 7 | 7 | 5 | 6 | 3 |
| 21 | TGME49_288680-t26_1-p1 | endonuclease IV APN | 10 | 8 | 8 | 6 | 5 | 7 | 7 | 3 |
| 30 | TGME49_288710-t26_1-p1 | hypothetical protein | 9 | 8 | 7 | 7 | 6 | 3 | 4 | 3 |
| 21 | TGME49_288720.P3187 | ribosomal protein RPL10 | 7 | 7 | 7 | 7 | 7 | 6 | 6 | 3 |
| 1 | TGME49_288730.P3188 | DNA polymerase, putative | 10 | 5 | 7 | 7 | 5 | 5 | 4 | 3 |
| 21 | TGME49_288750.P3189 | cytochrome b-c1 complex subunit 7, putative | 8 | 7 | 7 | 7 | 7 | 6 | 6 | 3 |
| 30 | TGME49_288990-t26_1-p1 | homeobox domain-containing protein | 9 | 7 | 8 | 8 | 6 | 3 | 3 | 3 |
| 21 | TGME49_289250-t26_1-p1 | cyclophilin, putative | 7 | 7 | 7 | 6 | 6 | 6 | 6 | 3 |
| 30 | TGME49_289360-t26_1-p1 | hypothetical protein | 8 | 7 | 7 | 7 | 6 | 3 | 4 | 3 |
| 21 | TGME49_289530-t26_1-p1 | ribosomal protein RPL19 | 7 | 7 | 7 | 6 | 6 | 6 | 6 | 3 |
| 1 | TGME49_289580-t26_1-p1 | strictosidine synthase subfamily protein | 9 | 8 | 8 | 7 | 6 | 5 | 5 | 3 |
| 1 | TGME49_289600-t26_1-p1 | heat shock protein HSP29 | 8 | 7 | 7 | 7 | 5 | 5 | 3 | 3 |
| 1 | TGME49_289630-t26_1-p1 | microneme protein MIC16 | 10 | 6 | 9 | 8 | 7 | 6 | 3 | 3 |
| 30 | TGME49_289670-t26_1-p1 | DNA repair metallo-beta-lactamase | 9 | 7 | 7 | 8 | 5 | 4 | 4 | 3 |
| 21 | TGME49_289680.P3208 | ras-related protein RAB11A | 7 | 7 | 7 | 7 | 7 | 7 | 7 | 3 |
| 17 | TGME49_289750.P3214 | ribosomal-ubiquitin protein RPL40 | 6 | 6 | 6 | 5 | 3 | 6 | 6 | 3 |
| 30 | TGME49_289760-t26_1-p1 | biotin--protein ligase, putative | 10 | 8 | 7 | 7 | 6 | 4 | 4 | 3 |
| 21 | TGME49_289820-t26_1-p1 | TBC domain-containing protein TBC14 | 9 | 9 | 9 | 6 | 6 | 7 | 6 | 3 |
| 30 | TGME49_289890.P3216 | hypothetical protein | 9 | 7 | 7 | 6 | 5 | 4 | 4 | 3 |
| 17 | TGME49_289900-t26_1-p1 | n-acetyltransferase family protein | 7 | 6 | 7 | 6 | 6 | 5 | 5 | 3 |
| 21 | TGME49_289950-t26_1-p1 | hypothetical protein | 9 | 8 | 8 | 7 | 7 | 6 | 5 | 3 |
| 30 | TGME49_290005.P3222 | proteasome subunit beta type-1, putative | 8 | 7 | 7 | 7 | 7 | 4 | 3 | 3 |

| | | | | | | | | | | |
|---|---|---|---|---|---|---|---|---|---|---|
| 21 | TGME49_290030-t26_1-p1 | ATP synthase-associated protein | 7 | 7 | 7 | 7 | 6 | 6 | 6 | 3 |
| 17 | TGME49_290260-t26_1-p1 | thioredoxin family Trp26 protein | 7 | 7 | 7 | 7 | 5 | 6 | 5 | 3 |
| 1 | TGME49_290340-t26_1-p1 | HEAT repeat-containing protein | 9 | 8 | 7 | 6 | 6 | 6 | 5 | 3 |
| 17 | TGME49_290900.P3238 | hydrolase, NUDIX family protein | 7 | 7 | 7 | 6 | 6 | 5 | 5 | 3 |
| 30 | TGME49_290920.P3239 | oxidoreductase, 2OG-Fe(II) oxygenase family protein | 10 | 8 | 7 | 7 | 6 | 3 | 3 | 3 |
| 21 | TGME49_291050-t26_1-p1 | serine/threonine-protein kinase KIN | 10 | 10 | 3 | 7 | 7 | 7 | 7 | 3 |
| 1 | TGME49_291080-t26_1-p1 | TATA-box binding protein TBP1 | 8 | 8 | 7 | 6 | 6 | 6 | 3 | 3 |
| 21 | TGME49_291090-t26_1-p1 | SWI2/SNF2-containing protein | 10 | 10 | 3 | 7 | 8 | 7 | 6 | 3 |
| 17 | TGME49_291300.P3248 | RER1 protein, putative | 7 | 7 | 6 | 6 | 6 | 3 | 6 | 3 |
| 30 | TGME49_291650-t26_1-p1 | hypothetical protein | 9 | 8 | 7 | 7 | 5 | 3 | 4 | 3 |
| 21 | TGME49_291800-t26_1-p1 | ADP-ribosylation factor-like protein ArlX2 | 7 | 7 | 7 | 6 | 7 | 6 | 6 | 3 |
| 17 | TGME49_291810-t26_1-p1 | thioredoxin domain-containing protein | 7 | 7 | 7 | 5 | 6 | 3 | 5 | 3 |
| 30 | TGME49_291900-t26_1-p1 | Ydr279p family (RNase H2 complex component) protein | 9 | 8 | 6 | 6 | 5 | 4 | 4 | 3 |
| 21 | TGME49_291930-t26_1-p1 | RNA recognition motif-containing protein | 10 | 8 | 7 | 7 | 6 | 6 | 6 | 3 |
| 21 | TGME49_292130.P3261 | ribosomal protein RPL13A | 7 | 7 | 7 | 7 | 7 | 6 | 6 | 3 |
| 21 | TGME49_292160-t26_1-p1 | replication termination factor, putative | 8 | 8 | 8 | 6 | 7 | 6 | 6 | 3 |
| 30 | TGME49_292190-t26_1-p1 | rRNA-processing protein FCF1, putative | 8 | 8 | 7 | 5 | 5 | 4 | 4 | 3 |
| 30 | TGME49_292310.P3264 | RNA polymerase Rpb7, N-terminal domain-containing protein | 7 | 7 | 7 | 6 | 6 | 3 | 5 | 3 |
| 17 | TGME49_293050.P3272 | sybindin family protein | 6 | 6 | 6 | 3 | 6 | 5 | 5 | 3 |
| 1 | TGME49_293220.P3278 | palmitoyltransferase DHHC15 | 10 | 8 | 7 | 7 | 6 | 5 | 4 | 3 |
| 30 | TGME49_293230.P3279 | FAM98 domain-containing protein | 9 | 7 | 7 | 6 | 5 | 4 | 3 | 3 |
| 30 | TGME49_293320.P3288 | hypothetical protein | 10 | 5 | 9 | 8 | 5 | 3 | 3 | 3 |
| 21 | TGME49_293340.P3290 | ran binding family protein 1, putative | 7 | 7 | 7 | 6 | 6 | 6 | 6 | 3 |
| 1 | TGME49_293380.P3293 | histone lysine acetyltransferase HAT1 | 9 | 8 | 8 | 6 | 5 | 5 | 5 | 3 |
| 30 | TGME49_293390.P3294 | cytochrome C-type heme lyase, putative | 9 | 7 | 7 | 6 | 5 | 4 | 5 | 3 |
| 21 | TGME49_293450.P3299 | protein phosphatase PPM6 | 8 | 9 | 8 | 8 | 7 | 6 | 6 | 3 |
| 1 | TGME49_293470.P3301 | immune mapped protein IMP1 | 9 | 8 | 7 | 6 | 6 | 5 | 4 | 3 |
| 30 | TGME49_293510.P3305 | poly(ADP-ribose) polymerase and DNA-Ligase Zn-finger region domain-containing protein | 9 | 8 | 7 | 7 | 5 | 3 | 3 | 3 |
| 21 | TGME49_293610-t26_1-p1 | dihydroorotase | 9 | 8 | 8 | 8 | 6 | 7 | 5 | 3 |
| 1 | TGME49_293660.P3315 | hypothetical protein | 10 | 5 | 8 | 7 | 6 | 5 | 5 | 3 |
| 21 | TGME49_293670.P3316 | transcription elongation factor A TFIIS | 9 | 8 | 8 | 7 | 7 | 6 | 6 | 3 |
| 17 | TGME49_293690.P3318 | profilin PRF | 7 | 7 | 6 | 6 | 6 | 4 | 5 | 3 |
| 1 | TGME49_293900.P3339 | sporozoite protein with an altered thrombospondin repeat SPATR | 9 | 8 | 7 | 6 | 5 | 5 | 4 | 3 |
| 21 | TGME49_294040.P3342 | KDEL endoplasmic reticulum proteinretention receptor 2 (KDELR2), putative | 9 | 8 | 8 | 6 | 8 | 6 | 7 | 3 |
| 30 | TGME49_294220-t26_1-p1 | MYND-like zinc finger protein | 9 | 7 | 8 | 7 | 6 | 3 | 3 | 3 |
| 21 | TGME49_294250-t26_1-p1 | WD domain, G-beta repeat-containing protein | 9 | 5 | 8 | 8 | 7 | 7 | 6 | 3 |
| 21 | TGME49_294290-t26_1-p1 | Der1ER1 | 8 | 8 | 6 | 7 | 7 | 6 | 5 | 3 |
| 30 | TGME49_294420-t26_1-p1 | programmed cell death protein 2, c-terminal domain-containing protein | 9 | 8 | 8 | 7 | 5 | 4 | 4 | 3 |
| 21 | TGME49_294670.P3364 | eukaryotic translation initiation factor 3 subunit G, putative | 8 | 8 | 8 | 6 | 7 | 6 | 6 | 3 |
| 30 | TGME49_294780.P3366 | hypothetical protein | 8 | 7 | 7 | 6 | 6 | 3 | 4 | 3 |
| 21 | TGME49_294850.P3371 | actin-like protein ALP7 | 8 | 8 | 8 | 7 | 7 | 5 | 5 | 3 |

| | | | | | | | | | | |
|---|---|---|---|---|---|---|---|---|---|---|
| 30 | TGME49_294870.P3373 | universal stress family protein | 9 | 8 | 7 | 6 | 6 | 4 | 3 | 3 |
| 21 | TGME49_295070-t26_1-p1 | helicase associated domain (ha2) protein | 10 | 10 | 3 | 7 | 7 | 9 | 7 | 3 |
| 1 | TGME49_295340-t26_1-p1 | UV excision repair protein Rad23 protein | 8 | 8 | 8 | 7 | 7 | 6 | 4 | 3 |
| 17 | TGME49_295350-t26_1-p1 | nucleoside diphosphate kinase, putative | 7 | 6 | 6 | 6 | 5 | 6 | 6 | 3 |
| 30 | TGME49_295390.P3384 | hypothetical protein | 9 | 8 | 6 | 7 | 6 | 4 | 4 | 3 |
| 21 | TGME49_295760.P3391 | calcium-dependent protein kinase CDPK4A | 9 | 8 | 9 | 6 | 7 | 5 | 6 | 3 |
| 30 | TGME49_295920.P3393 | dynein axonemal assembly factor 4, putative | 9 | 7 | 8 | 6 | 5 | 4 | 4 | 3 |
| 1 | TGME49_295990.P3394 | ubiquitin conjugating enzyme E2, putative | 9 | 8 | 7 | 6 | 6 | 5 | 5 | 3 |
| 17 | TGME49_295995-t26_1-p1 | hypothetical protein | 8 | 3 | 7 | 6 | 6 | 6 | 6 | 3 |
| 17 | TGME49_297130-t26_1-p1 | hypothetical protein | 7 | 7 | 7 | 6 | 6 | 5 | 5 | 3 |
| 21 | TGME49_297150-t26_1-p1 | MORN repeat-containing protein | 9 | 7 | 7 | 7 | 6 | 5 | 6 | 3 |
| 1 | TGME49_297170-t26_1-p1 | 50S ribosomal protein L17, putative | 8 | 7 | 7 | 7 | 7 | 6 | 3 | 3 |
| 17 | TGME49_297270-t26_1-p1 | hypothetical protein | 8 | 7 | 7 | 5 | 5 | 6 | 5 | 3 |
| 21 | TGME49_297470.P3409 | myosin light chain MLC2 | 9 | 8 | 8 | 6 | 5 | 6 | 6 | 3 |
| 30 | TGME49_297640.P3413 | acyltransferase ATS2 | 9 | 7 | 7 | 6 | 5 | 4 | 4 | 3 |
| 1 | TGME49_297650.P3414 | inorganic phosphate transporter PIT | 10 | 9 | 7 | 6 | 5 | 5 | 5 | 3 |
| 17 | TGME49_297680-t26_1-p1 | calmodulin, putative | 7 | 5 | 6 | 6 | 6 | 5 | 5 | 3 |
| 17 | TGME49_297730.P3415 | transcription elongation factor 1-like divergent protein ELD | 7 | 6 | 6 | 5 | 5 | 5 | 4 | 3 |
| 17 | TGME49_297810.P3420 | cytochrome c oxidase subunit ApiCOX30 | 7 | 7 | 7 | 6 | 6 | 5 | 5 | 3 |
| 21 | TGME49_298630-t26_1-p1 | vacuolar transporter chaperone VTC2 | 10 | 3 | 10 | 9 | 8 | 6 | 6 | 3 |
| 1 | TGME49_298990-t26_1-p1 | ferredoxin NADP+ oxidoreductase FNR | 9 | 8 | 7 | 7 | 5 | 5 | 5 | 3 |
| 30 | TGME49_299040-t26_1-p1 | phosphatidylinositol n-acetylglucosaminyltransferase | 9 | 8 | 8 | 7 | 6 | 3 | 5 | 3 |
| 17 | TGME49_299200-t26_1-p1 | Bet3 transport protein, putative | 7 | 7 | 7 | 6 | 7 | 5 | 5 | 3 |
| 1 | TGME49_299970-t26_1-p1 | tetratricopeptide repeat-containing protein | 9 | 8 | 7 | 7 | 6 | 5 | 5 | 3 |
| 21 | TGME49_300000.P3449 | ribosomal protein RPL18 | 7 | 7 | 7 | 6 | 6 | 6 | 6 | 3 |
| 21 | TGME49_300040-t26_1-p1 | ubiquitin-conjugating enzyme subfamily protein | 8 | 7 | 7 | 7 | 7 | 6 | 5 | 3 |
| 17 | TGME49_300060-t26_1-p1 | signal peptidase subunit protein | 7 | 6 | 6 | 6 | 6 | 5 | 5 | 3 |
| 30 | TGME49_300110.P3454 | RNA transcription, translation and transport factor protein, putative | 8 | 7 | 6 | 6 | 5 | 3 | 4 | 3 |
| 30 | TGME49_300150-t26_1-p1 | hypothetical protein | 8 | 7 | 6 | 6 | 6 | 3 | 5 | 3 |
| 17 | TGME49_300200.P3460 | histone H2AZ | 6 | 6 | 6 | 6 | 6 | 6 | 5 | 3 |
| 30 | TGME49_300320-t26_1-p1 | dimethyladenosine transferase | 10 | 6 | 8 | 7 | 7 | 4 | 5 | 3 |
| 1 | TGME49_300340-t26_1-p1 | CAF1 family ribonuclease | 9 | 8 | 8 | 6 | 5 | 5 | 5 | 3 |
| 30 | TGME49_300621-t26_1-p1 | ribosomal protein L2 | 6 | 5 | 7 | 7 | 7 | 3 | 3 | 3 |
| 1 | TGME49_301340.P3473 | DnaJ domain-containing protein | 9 | 8 | 7 | 5 | 5 | 6 | 5 | 3 |
| 1 | TGME49_301350.P3474 | SNARE associated protein | 9 | 8 | 7 | 6 | 6 | 5 | 5 | 3 |
| 21 | TGME49_301380-t26_1-p1 | elongation factor Tu GTP binding domain-containing protein | 10 | 3 | 9 | 8 | 8 | 6 | 6 | 3 |
| 30 | TGME49_301410-t26_1-p1 | AP-4 complex accessory subunit Tepsin, putative | 10 | 9 | 7 | 6 | 6 | 4 | 3 | 3 |
| 21 | TGME49_301420.P3478 | SAS6-like protein | 8 | 7 | 7 | 7 | 6 | 6 | 6 | 3 |
| 1 | TGME49_301430.P3479 | septum formation protein maf, putative | 9 | 7 | 6 | 6 | 5 | 5 | 4 | 3 |
| 30 | TGME49_302050-t26_1-p1 | elongation factor tu, apicoplast, putative | 6 | 3 | 9 | 9 | 8 | 3 | 3 | 3 |
| 21 | TGME49_304490-t26_1-p1 | hypothetical protein | 8 | 8 | 7 | 7 | 6 | 6 | 6 | 3 |
| 21 | TGME49_304500.P3483 | hypothetical protein | 8 | 8 | 7 | 7 | 7 | 5 | 6 | 3 |
| 1 | TGME49_304740.P3490 | rhoptry kinase family protein ROP35 | 9 | 8 | 8 | 7 | 6 | 6 | 5 | 3 |

| | | | | | | | | | | |
|---|---|---|---|---|---|---|---|---|---|---|
| 17 | TGME49_304770.P3492 | ubiquitin-conjugating enzyme subfamily protein | 6 | 6 | 6 | 6 | 6 | 6 | 6 | 3 |
| 21 | TGME49_304890-t26_1-p1 | hypothetical protein | 8 | 7 | 7 | 5 | 6 | 6 | 6 | 3 |
| 17 | TGME49_305010.P3494 | pre-mRNA branch site protein p14, putative | 6 | 6 | 6 | 6 | 5 | 5 | 5 | 3 |
| 1 | TGME49_305190-t26_1-p1 | CorA family Mg2+ transporter protein | 10 | 9 | 7 | 7 | 6 | 6 | 3 | 3 |
| 21 | TGME49_305290.P3499 | vacuolar atp synthase subunit e, putative | 8 | 7 | 7 | 7 | 7 | 7 | 6 | 3 |
| 1 | TGME49_305450-t26_1-p1 | acetyltransferase, GNAT family protein | 9 | 8 | 8 | 7 | 7 | 5 | 4 | 3 |
| 17 | TGME49_305455.P3503 | hypothetical protein | 7 | 3 | 6 | 6 | 5 | 5 | 5 | 3 |
| 30 | TGME49_305490.P3504 | programmed cell death protein 2, c-terminal domain-containing protein | 9 | 8 | 7 | 7 | 5 | 4 | 5 | 3 |
| 21 | TGME49_305520-t26_1-p1 | ribosomal protein RPS2 | 8 | 8 | 8 | 7 | 7 | 7 | 7 | 3 |
| 17 | TGME49_305600-t26_1-p1 | hypothetical protein | 7 | 7 | 6 | 6 | 5 | 5 | 5 | 3 |
| 21 | TGME49_305610-t26_1-p1 | hypothetical protein | 9 | 8 | 8 | 6 | 6 | 6 | 6 | 3 |
| 30 | TGME49_305810-t26_1-p1 | hypothetical protein | 8 | 8 | 7 | 6 | 5 | 3 | 4 | 3 |
| 30 | TGME49_305820-t26_1-p1 | SGS domain-containing protein | 8 | 7 | 7 | 6 | 5 | 4 | 4 | 3 |
| 1 | TGME49_305850-t26_1-p1 | RNA recognition motif-containing protein | 9 | 8 | 6 | 5 | 5 | 5 | 4 | 3 |
| 21 | TGME49_305920-t26_1-p1 | endonuclease III family 1 protein | 10 | 7 | 8 | 7 | 6 | 6 | 6 | 3 |
| 1 | TGME49_306030.P3514 | glutathione S-transferase 2 | 8 | 3 | 8 | 7 | 8 | 6 | 3 | 3 |
| 17 | TGME49_306050-t26_1-p1 | FAS1 domain-containing protein | 7 | 3 | 7 | 6 | 5 | 5 | 5 | 3 |
| 17 | TGME49_306070.P3516 | DUF4201 domain-containing protein | 7 | 3 | 6 | 6 | 6 | 4 | 5 | 3 |
| 30 | TGME49_306480-t26_1-p1 | CAM kinase, CDPK family | 10 | 7 | 8 | 6 | 6 | 3 | 6 | 3 |
| 1 | TGME49_306560-t26_1-p1 | nucleoporin NUP67, putative | 9 | 8 | 8 | 7 | 5 | 5 | 5 | 3 |
| 17 | TGME49_306600.P3525 | RNA recognition motif-containing protein | 7 | 5 | 6 | 5 | 5 | 5 | 4 | 3 |
| 30 | TGME49_306610-t26_1-p1 | 3' exoribonuclease family, domain 1 domain-containing protein | 9 | 8 | 7 | 5 | 5 | 4 | 4 | 3 |
| 30 | TGME49_306640.P3526 | SNARE protein STX21 | 8 | 8 | 7 | 6 | 5 | 3 | 3 | 3 |
| 17 | TGME49_306650-t26_1-p1 | succinate dehydrogenase subunit SDH31 | 8 | 7 | 6 | 6 | 5 | 4 | 5 | 3 |
| 17 | TGME49_306670.P3528 | cytochrome c oxidase subunit ApiCOX26 | 7 | 7 | 7 | 6 | 6 | 3 | 6 | 3 |
| 21 | TGME49_306930.P3531 | proteasome subunit beta type-7, putative | 8 | 8 | 8 | 6 | 7 | 7 | 6 | 3 |
| 30 | TGME49_306980.P3533 | hypothetical protein | 7 | 7 | 6 | 6 | 6 | 3 | 4 | 3 |
| 1 | TGME49_307020-t26_1-p1 | hypothetical protein | 8 | 7 | 6 | 6 | 5 | 5 | 4 | 3 |
| 17 | TGME49_307030-t26_1-p1 | Purine nucleoside phosphorylase | 3 | 7 | 7 | 7 | 3 | 5 | 6 | 3 |
| 30 | TGME49_307610-t26_1-p1 | elongation factor TS, putative | 10 | 8 | 7 | 6 | 6 | 4 | 3 | 3 |
| 30 | TGME49_307650-t26_1-p1 | uracil-DNA glycosylase | 8 | 7 | 8 | 5 | 6 | 3 | 4 | 3 |
| 1 | TGME49_307840.P3539 | striated fiber assemblin SFA1 | 7 | 7 | 8 | 7 | 6 | 7 | 3 | 3 |
| 17 | TGME49_308050-t26_1-p1 | thioredoxin domain-containing protein | 6 | 6 | 6 | 6 | 4 | 4 | 5 | 3 |
| 1 | TGME49_308860.P3545 | apical cap protein AC3 | 9 | 8 | 7 | 6 | 5 | 5 | 4 | 3 |
| 17 | TGME49_308930-t26_1-p1 | 50S ribosomal protein L33, putative | 6 | 6 | 6 | 6 | 5 | 3 | 5 | 3 |
| 17 | TGME49_309010.P3555 | elongation factor P, putative | 7 | 6 | 6 | 6 | 4 | 5 | 4 | 3 |
| 21 | TGME49_309170.P3562 | TAF7-like RNA polymerase II TAF7L | 10 | 7 | 8 | 7 | 7 | 7 | 6 | 3 |
| 21 | TGME49_309210.P3565 | peroxiredoxin 6, putative | 7 | 7 | 7 | 7 | 7 | 5 | 6 | 3 |
| 1 | TGME49_309220-t26_1-p1 | GTPase activating protein for Arf protein | 9 | 7 | 7 | 6 | 6 | 6 | 5 | 3 |
| 1 | TGME49_309240-t26_1-p1 | CorA family Mg2+ transporter protein | 10 | 7 | 7 | 7 | 6 | 5 | 4 | 3 |
| 30 | TGME49_309390-t26_1-p1 | hypothetical protein | 9 | 7 | 7 | 6 | 5 | 4 | 4 | 3 |
| 30 | TGME49_309550.P3570 | hypothetical protein | 8 | 8 | 8 | 7 | 6 | 3 | 3 | 3 |
| 21 | TGME49_309560.P3571 | nmda receptor glutamate-binding chain | 8 | 8 | 7 | 7 | 7 | 6 | 6 | 3 |
| 17 | TGME49_309740-t26_1-p1 | LSM domain-containing protein | 6 | 6 | 6 | 5 | 5 | 5 | 4 | 3 |
| 1 | TGME49_309790.P3576 | hypothetical protein | 8 | 8 | 8 | 7 | 6 | 5 | 4 | 3 |
| 1 | TGME49_309800-t26_1-p1 | RNA recognition motif-containing protein | 9 | 8 | 7 | 7 | 5 | 6 | 5 | 3 |
| 21 | TGME49_309820-t26_1-p1 | ribosomal protein RPL11 | 7 | 7 | 7 | 7 | 7 | 6 | 6 | 3 |

| | | | | | | | | | | |
|---|---|---|---|---|---|---|---|---|---|---|
| 1 | TGME49_309940.P3586 | phospholipase D active site domain-containing protein | 10 | 9 | 7 | 6 | 7 | 5 | 4 | 3 |
| 17 | TGME49_310040.P3592 | ubiquitin conjugating enzyme E2, putative | 6 | 6 | 6 | 3 | 6 | 3 | 6 | 3 |
| 30 | TGME49_310050-t26_1-p1 | RNA recognition motif-containing protein | 9 | 8 | 8 | 7 | 6 | 4 | 4 | 3 |
| 1 | TGME49_310070.P3593 | apical annuli methyltransferase AAMT | 8 | 7 | 7 | 7 | 7 | 5 | 5 | 3 |
| 1 | TGME49_310100.P3594 | mannosyltransferase (pig-m) protein | 8 | 8 | 7 | 6 | 5 | 5 | 5 | 3 |
| 30 | TGME49_310200-t26_1-p1 | hypothetical protein | 10 | 7 | 8 | 7 | 5 | 4 | 4 | 3 |
| 30 | TGME49_310300-t26_1-p1 | hypothetical protein | 10 | 7 | 8 | 6 | 6 | 3 | 4 | 3 |
| 21 | TGME49_310380-t26_1-p1 | U3 small nucleolar ribonucleoprotein protein IMP4, putative | 8 | 7 | 7 | 8 | 8 | 7 | 7 | 3 |
| 17 | TGME49_310460-t26_1-p1 | ras-related protein RAB6 | 7 | 7 | 7 | 7 | 7 | 3 | 6 | 3 |
| 17 | TGME49_310470.P3610 | cytochrome c oxidase subunit COX2b | 7 | 7 | 7 | 6 | 6 | 3 | 6 | 3 |
| 17 | TGME49_310490.P3611 | ribosomal protein RPL27A | 6 | 6 | 6 | 6 | 6 | 6 | 5 | 3 |
| 30 | TGME49_310500-t26_1-p1 | hypothetical protein | 7 | 7 | 7 | 6 | 5 | 3 | 4 | 3 |
| 21 | TGME49_310580-t26_1-p1 | 1,3(4)-beta-glucanase | 10 | 7 | 9 | 8 | 6 | 6 | 6 | 3 |
| 1 | TGME49_310640.P3618 | phosphorylase family protein | 8 | 8 | 7 | 7 | 6 | 5 | 5 | 3 |
| 30 | TGME49_310680.P3620 | aminotransferase, class I/II superfamily protein | 9 | 7 | 6 | 8 | 6 | 3 | 3 | 3 |
| 21 | TGME49_310760-t26_1-p1 | protein phosphatase PPM7, putative | 9 | 7 | 8 | 6 | 5 | 7 | 6 | 3 |
| 30 | TGME49_310770.P3622 | apicoplast-associated thioredoxin family protein Atrx2 | 8 | 7 | 6 | 6 | 6 | 3 | 4 | 3 |
| 21 | TGME49_310830-t26_1-p1 | enoyl-CoA hydratase/isomerase family protein | 8 | 8 | 7 | 6 | 7 | 5 | 6 | 3 |
| 30 | TGME49_310850-t26_1-p1 | MYND finger domain-containing protein | 8 | 7 | 7 | 6 | 6 | 3 | 5 | 3 |
| 17 | TGME49_310870-t26_1-p1 | integral membrane protein, putative | 7 | 7 | 6 | 6 | 5 | 4 | 4 | 3 |
| 17 | TGME49_310960.P3629 | vacuolar atp synthase subunit f, putative | 6 | 6 | 6 | 6 | 6 | 5 | 4 | 3 |
| 30 | TGME49_311250-t26_1-p1 | hypothetical protein | 10 | 6 | 9 | 7 | 5 | 4 | 4 | 3 |
| 1 | TGME49_311290-t26_1-p1 | protein tyrosine phosphatase family protein, ptpla protein | 8 | 7 | 7 | 6 | 6 | 5 | 4 | 3 |
| 1 | TGME49_311660-t26_1-p1 | histone lysine methyltransferase, SET, putative | 10 | 8 | 7 | 7 | 6 | 6 | 4 | 3 |
| 21 | TGME49_311690-t26_1-p1 | UBA/TS-N domain-containing protein | 9 | 8 | 8 | 8 | 7 | 5 | 6 | 3 |
| 17 | TGME49_311750-t26_1-p1 | mago binding protein | 7 | 6 | 6 | 4 | 4 | 5 | 5 | 3 |
| 21 | TGME49_311780-t26_1-p1 | Zn-containing alcohol dehydrogenase | 9 | 8 | 8 | 8 | 6 | 7 | 6 | 3 |
| 30 | TGME49_311820.P3657 | hypothetical protein | 8 | 9 | 8 | 8 | 6 | 4 | 4 | 3 |
| 21 | TGME49_312050.P3660 | ras-related protein RAB2 | 7 | 7 | 7 | 7 | 7 | 5 | 7 | 3 |
| 17 | TGME49_312090-t26_1-p1 | ribosomal protein RPL23 | 6 | 6 | 6 | 6 | 6 | 6 | 3 | 3 |
| 17 | TGME49_312220-t26_1-p1 | mitochondrial inner membrane translocase subunit TIM17, putative | 7 | 7 | 7 | 5 | 6 | 6 | 6 | 3 |
| 30 | TGME49_312390-t26_1-p1 | hypothetical protein | 8 | 8 | 8 | 6 | 6 | 3 | 4 | 3 |
| 1 | TGME49_312400.P3669 | exosomal 3'-5' exoribonuclease complex subunit, putative | 8 | 7 | 7 | 6 | 6 | 5 | 3 | 3 |
| 21 | TGME49_312480-t26_1-p1 | uracil phosphoribosyltransferase FUR1, putative | 8 | 7 | 7 | 6 | 7 | 4 | 6 | 3 |
| 17 | TGME49_312490-t26_1-p1 | hypothetical protein | 8 | 6 | 4 | 6 | 6 | 3 | 5 | 3 |
| 30 | TGME49_312520-t26_1-p1 | tRNA dimethylallyltransferase | 10 | 7 | 8 | 8 | 6 | 3 | 5 | 3 |
| 1 | TGME49_312618-t26_1-p1 | hypothetical protein | 9 | 8 | 8 | 6 | 6 | 5 | 5 | 3 |
| 17 | TGME49_312960-t26_1-p1 | ribosome biogenesis protein NOP16, putative | 7 | 6 | 6 | 6 | 5 | 4 | 4 | 3 |
| 1 | TGME49_312980.P3686 | flagella associated protein, putative | 9 | 8 | 8 | 8 | 7 | 5 | 4 | 3 |

| | | | | | | | | | | |
|---|---|---|---|---|---|---|---|---|---|---|
| 30 | TGME49_313050-t26_1-p1 | oxidoreductase, short chain dehydrogenase/reductase family protein | 9 | 8 | 8 | 7 | 6 | 3 | 3 | 3 |
| 30 | TGME49_313080-t26_1-p1 | hypothetical protein | 9 | 8 | 7 | 7 | 6 | 4 | 4 | 3 |
| 30 | TGME49_313130-t26_1-p1 | hypothetical protein | 9 | 7 | 6 | 7 | 5 | 4 | 3 | 3 |
| 21 | TGME49_313190-t26_1-p1 | ras-related protein RAB18 | 7 | 7 | 7 | 6 | 7 | 6 | 6 | 3 |
| 17 | TGME49_313200-t26_1-p1 | F-Box Protein L2 | 8 | 6 | 7 | 5 | 5 | 5 | 4 | 3 |
| 21 | TGME49_313290.P3697 | MORN repeat-containing protein | 9 | 8 | 8 | 8 | 7 | 5 | 6 | 3 |
| 21 | TGME49_313380.P3702 | IMC localizing protein ILP1 | 8 | 8 | 8 | 5 | 6 | 7 | 7 | 3 |
| 17 | TGME49_313390.P3703 | ribosomal protein RPL6 | 7 | 7 | 7 | 6 | 6 | 3 | 6 | 3 |
| 1 | TGME49_313400-t26_1-p1 | DnaJ domain-containing protein | 8 | 7 | 7 | 6 | 6 | 5 | 4 | 3 |
| 30 | TGME49_313550-t26_1-p1 | ribosomal protein RPL22 | 9 | 6 | 6 | 6 | 5 | 3 | 4 | 3 |
| 17 | TGME49_313560-t26_1-p1 | 60S ribosomal protein L7a, putative | 7 | 6 | 6 | 6 | 5 | 3 | 5 | 3 |
| 30 | TGME49_313640-t26_1-p1 | hypothetical protein | 7 | 7 | 6 | 6 | 5 | 3 | 4 | 3 |
| 30 | TGME49_313665-t26_1-p1 | hypothetical protein | 9 | 7 | 7 | 6 | 5 | 3 | 3 | 3 |
| 1 | TGME49_313790.P3722 | centrosomal protein CEP44, putative | 10 | 8 | 7 | 6 | 5 | 5 | 4 | 3 |
| 30 | TGME49_313820-t26_1-p1 | dynactin p25, putative | 9 | 7 | 7 | 7 | 4 | 3 | 5 | 3 |
| 30 | TGME49_313880-t26_1-p1 | nuclear protein-like family protein | 8 | 8 | 7 | 6 | 5 | 3 | 6 | 3 |
| 21 | TGME49_313960-t26_1-p1 | ribosomal protein L19 protein | 9 | 7 | 8 | 8 | 7 | 7 | 6 | 3 |
| 17 | TGME49_314042.P3730 | cytochrome c1 heme lyase, putative | 7 | 8 | 7 | 6 | 6 | 5 | 5 | 3 |
| 1 | TGME49_314090-t26_1-p1 | proteasome subunit beta type-3, putative | 7 | 4 | 7 | 7 | 7 | 6 | 3 | 3 |
| 1 | TGME49_314270.P3735 | peptidyl-tRNA hydrolase domain-containing protein | 9 | 8 | 8 | 7 | 5 | 6 | 5 | 3 |
| 30 | TGME49_314280-t26_1-p1 | AAR2 protein | 9 | 8 | 8 | 7 | 7 | 3 | 4 | 3 |
| 17 | TGME49_314418.P3741 | AP-4 complex subunit sigma, putative | 6 | 6 | 6 | 6 | 6 | 6 | 5 | 3 |
| 1 | TGME49_314540-t26_1-p1 | hypothetical protein | 9 | 9 | 8 | 7 | 5 | 6 | 3 | 3 |
| 17 | TGME49_314720-t26_1-p1 | trafficking protein particle complex subunit 2, putative | 7 | 6 | 6 | 6 | 5 | 5 | 4 | 3 |
| 30 | TGME49_314750-t26_1-p1 | ribosomal protein L1, putative | 9 | 8 | 7 | 7 | 6 | 3 | 5 | 3 |
| 17 | TGME49_314890.P3751 | ThiF family protein | 3 | 9 | 9 | 6 | 6 | 4 | 5 | 3 |
| 1 | TGME49_314900.P3752 | LisH protein | 9 | 8 | 8 | 7 | 6 | 6 | 3 | 3 |
| 21 | TGME49_315300.P3761 | transcription factor IIB, putative | 7 | 7 | 7 | 7 | 7 | 7 | 5 | 3 |
| 30 | TGME49_315440-t26_1-p1 | hypothetical protein | 10 | 7 | 6 | 6 | 5 | 3 | 3 | 3 |
| 17 | TGME49_315500-t26_1-p1 | hypothetical protein | 7 | 6 | 6 | 6 | 3 | 5 | 5 | 3 |
| 30 | TGME49_315560-t26_1-p1 | ATP-binding cassette G family transporter ABCG77 | 10 | 10 | 5 | 7 | 7 | 3 | 4 | 3 |
| 30 | TGME49_315640-t26_1-p1 | lipoyl(octanoyl) transferase | 9 | 7 | 7 | 6 | 5 | 4 | 4 | 3 |
| 1 | TGME49_315690-t26_1-p1 | DnaJ domain-containing protein | 8 | 7 | 7 | 7 | 6 | 5 | 5 | 3 |
| 30 | TGME49_315710.P3785 | hypothetical protein | 7 | 7 | 6 | 6 | 6 | 3 | 3 | 3 |
| 21 | TGME49_315730-t26_1-p1 | apical membrane antigen AMA3 | 9 | 8 | 8 | 8 | 7 | 6 | 6 | 3 |
| 17 | TGME49_315780.P3789 | myosin light chain MLC7 | 7 | 7 | 6 | 5 | 6 | 5 | 4 | 3 |
| 17 | TGME49_315920.P3794 | DNA-directed RNA polymerase II RPB11A | 6 | 6 | 6 | 6 | 6 | 5 | 5 | 3 |
| 17 | TGME49_315930-t26_1-p1 | succinate dehydrogenase subunit SDH18 | 7 | 6 | 6 | 5 | 6 | 5 | 5 | 3 |
| 17 | TGME49_315958-t26_1-p1 | radical SAM domain-containing protein | 3 | 8 | 3 | 8 | 8 | 5 | 3 | 3 |
| 17 | TGME49_316140-t26_1-p1 | MGR2 domain-containing protein | 7 | 7 | 6 | 6 | 5 | 5 | 5 | 3 |
| 30 | TGME49_316220-t26_1-p1 | mediator complex subunit MED6 | 8 | 8 | 7 | 6 | 5 | 4 | 4 | 3 |
| 30 | TGME49_316250.P3800 | dense granule protein GRA45 | 9 | 8 | 8 | 7 | 6 | 3 | 3 | 3 |
| 17 | TGME49_316310.P3805 | superoxide dismutase SOD1 | 7 | 7 | 7 | 7 | 6 | 5 | 5 | 3 |
| 17 | TGME49_316340.P3807 | inner membrane complex protein IMC22 | 8 | 7 | 6 | 6 | 6 | 5 | 5 | 3 |
| 21 | TGME49_316450.P3809 | ceramide synthase CERS2, putative | 9 | 8 | 8 | 7 | 6 | 6 | 6 | 3 |
| 17 | TGME49_316540.P3812 | IMC sub-compartment protein ISP3 | 7 | 7 | 6 | 6 | 6 | 5 | 4 | 3 |

| | | | | | | | | | | |
|---|---|---|---|---|---|---|---|---|---|---|
| 21 | TGME49_318160.P3822 | MSP (Major sperm protein) domain-containing protein | 8 | 7 | 7 | 7 | 6 | 6 | 6 | 3 |
| 30 | TGME49_318290.P3826 | PDZ domain-containing protein | 8 | 7 | 7 | 6 | 5 | 3 | 4 | 3 |
| 17 | TGME49_318420-t26_1-p1 | 30S ribosomal protein S16, putative | 7 | 6 | 6 | 6 | 6 | 5 | 3 | 3 |
| 21 | TGME49_318570.P3836 | SFT2 family protein | 8 | 8 | 7 | 7 | 6 | 6 | 6 | 3 |
| 1 | TGME49_318720-t26_1-p1 | pyridoxal phosphate enzyme, YggS family protein | 8 | 7 | 7 | 5 | 6 | 5 | 4 | 3 |
| 30 | TGME49_318750-t26_1-p1 | deoxyribose-phosphate aldolase | 8 | 8 | 7 | 6 | 6 | 3 | 3 | 3 |
| 1 | TGME49_318760-t26_1-p1 | hypothetical protein | 8 | 8 | 8 | 5 | 7 | 6 | 3 | 3 |
| 1 | TGME49_319370-t26_1-p1 | hypothetical protein | 10 | 9 | 6 | 5 | 6 | 6 | 4 | 3 |
| 17 | TGME49_319570.P3852 | WD domain, G-beta repeat-containing protein | 3 | 9 | 9 | 7 | 6 | 3 | 6 | 3 |
| 1 | TGME49_319650-t26_1-p1 | 3' exoribonuclease family, domain 1 domain-containing protein | 8 | 7 | 7 | 6 | 5 | 6 | 4 | 3 |
| 30 | TGME49_319700.P3855 | NEK kinase | 8 | 8 | 8 | 7 | 5 | 3 | 6 | 3 |
| 30 | TGME49_319740.P3858 | transporter, major facilitator family protein | 8 | 8 | 7 | 7 | 5 | 3 | 4 | 3 |
| 17 | TGME49_319870.P3859 | ubiquitin-conjugating enzyme subfamily protein | 6 | 6 | 6 | 6 | 6 | 6 | 6 | 3 |
| 30 | TGME49_319910-t26_1-p1 | WD domain, G-beta repeat-containing protein | 9 | 7 | 7 | 5 | 6 | 4 | 4 | 3 |
| 21 | TGME49_319950-t26_1-p1 | rRNA-processing protein FCF1, putative | 6 | 7 | 7 | 7 | 6 | 6 | 6 | 3 |
| 21 | TGME49_320070-t26_1-p1 | CDK-activating kinase assembly factor MAT1 protein | 8 | 7 | 8 | 7 | 6 | 6 | 5 | 3 |
| 21 | TGME49_320110-t26_1-p1 | proliferating cell nuclear antigen PCNA2 | 8 | 7 | 8 | 8 | 7 | 6 | 6 | 3 |
| 17 | TGME49_320120-t26_1-p1 | diadenosine tetraphosphatase family protein | 7 | 6 | 6 | 6 | 5 | 4 | 4 | 3 |
| 30 | TGME49_320150-t26_1-p1 | elongation factor Tu GTP binding domain-containing protein | 9 | 9 | 8 | 6 | 7 | 3 | 3 | 3 |
| 1 | TGME49_320450-t26_1-p1 | ribosome biogenesis regulatory protein (rrs1) protein | 8 | 8 | 7 | 7 | 6 | 5 | 4 | 3 |
| 21 | TGME49_320480-t26_1-p1 | ras-related protein RAB11B | 8 | 7 | 7 | 5 | 7 | 7 | 7 | 3 |
| 17 | TGME49_320560-t26_1-p1 | p25-alpha family protein | 6 | 6 | 6 | 3 | 6 | 5 | 5 | 3 |
| 30 | TGME49_320700-t26_1-p1 | AP2 domain transcription factor AP2IV-1 | 9 | 7 | 6 | 6 | 6 | 3 | 4 | 3 |
| 1 | TGME49_320740.P3889 | hypothetical protein | 8 | 7 | 7 | 7 | 6 | 5 | 5 | 3 |
| 17 | TGME49_321340.P3891 | membrane protein, putative | 8 | 5 | 8 | 6 | 7 | 4 | 6 | 3 |
| 1 | TGME49_321360.P3892 | clustered-asparagine-rich protein | 8 | 7 | 7 | 6 | 7 | 6 | 4 | 3 |
| 30 | TGME49_321430-t26_1-p1 | DNA repair protein recA , putative | 9 | 8 | 8 | 7 | 6 | 4 | 4 | 3 |
| 17 | TGME49_321520-t26_1-p1 | HSP20-like chaperone | 7 | 6 | 6 | 5 | 5 | 5 | 5 | 3 |
| 21 | TGME49_321570.P3896 | beta-hydroxyacyl-acyl carrier protein dehydratase (FABZ) | 7 | 7 | 7 | 6 | 6 | 6 | 6 | 3 |
| 1 | TGME49_321590-t26_1-p1 | hypothetical protein | 9 | 7 | 8 | 7 | 6 | 5 | 4 | 3 |
| 30 | TGME49_355200-t26_1-p1 | hypothetical protein | 8 | 3 | 8 | 8 | 8 | 3 | 3 | 3 |
| 21 | TGME49_500022.P25 | cyclic nucleotide-binding domain-containing protein | 7 | 9 | 6 | 8 | 7 | 7 | 5 | 3 |
| 30 | TGME49_500053.P354 | hypothetical protein, conserved | 8 | 7 | 7 | 7 | 6 | 4 | 5 | 3 |
| 30 | TGME49_500070.P371 | coiled-coil domain-containing protein, putative | 7 | 7 | 6 | 6 | 5 | 3 | 4 | 3 |
| 1 | TGME49_500076.P377 | tubulin epsilon chain | 9 | 7 | 8 | 5 | 5 | 5 | 4 | 3 |
| 17 | TGME49_500136.P438 | kelch repeat protein | 7 | 8 | 3 | 8 | 7 | 5 | 6 | 3 |
| 17 | TGME49_500155.P159 | hypothetical protein, conserved | 8 | 3 | 7 | 6 | 7 | 5 | 5 | 3 |
| 30 | TGME49_500160.P164 | histone arginine methyltransferase PRMT2 | 9 | 8 | 7 | 7 | 5 | 3 | 5 | 3 |

| | | | | | | | | | | |
|---|---|---|---|---|---|---|---|---|---|---|
| 1 | TGME49_500206.P211 | coiled-coil domain-containing protein | 8 | 8 | 8 | 6 | 5 | 6 | 5 | 3 |
| 17 | TGME49_500252.P41 | AP2 domain transcription factor AP2XII-10 | 7 | 3 | 7 | 7 | 6 | 6 | 5 | 3 |
| 17 | TGME49_500301.P90 | ubiquitin-associated domain-containing protein | 3 | 9 | 8 | 6 | 5 | 7 | 5 | 3 |
| 30 | TGME49_500335.P125 | CFAP53 domain-containing protein | 7 | 9 | 8 | 7 | 6 | 4 | 3 | 3 |
| 30 | TGME49_500352.P142 | hypothetical protein, conserved | 10 | 8 | 7 | 7 | 5 | 4 | 4 | 3 |

| Kmean_cluster | GeneID (all protein TG) | Product Description | Decile_bitHH | Decile_bitNcan | Decile_bitBB | Decilebit_CS | Decilebit_SN3 | Decile bit_Ef | Decile bit_CC | Meta_cluster |
|---|---|---|---|---|---|---|---|---|---|---|
| 11 | TGME49_200430.P460 | cytidine and deoxycytidylate deaminase zinc-binding region domain-containing protein | 9 | 8 | 8 | 7 | 4 | 6 | 6 | 4 |
| 7 | TGME49_201270-t26_1-p1 | hypothetical protein | 10 | 8 | 8 | 7 | 4 | 4 | 3 | 4 |
| 27 | TGME49_201780.P473 | microneme protein MIC2 | 10 | 9 | 9 | 8 | 3 | 7 | 7 | 4 |
| 7 | TGME49_201870-t26_1-p1 | tetratricopeptide repeat-containing protein | 10 | 10 | 9 | 8 | 3 | 4 | 4 | 4 |
| 7 | TGME49_202070.P481 | hypothetical protein | 10 | 10 | 8 | 7 | 5 | 4 | 4 | 4 |
| 7 | TGME49_202190.P485 | hypothetical protein | 10 | 9 | 9 | 8 | 5 | 4 | 4 | 4 |
| 27 | TGME49_202230-t26_1-p1 | histone deacetylase HDAC5 | 10 | 10 | 10 | 7 | 5 | 8 | 7 | 4 |
| 7 | TGME49_202240-t26_1-p1 | RAP domain-containing protein | 10 | 10 | 9 | 6 | 4 | 3 | 3 | 4 |
| 11 | TGME49_202290-t26_1-p1 | membrane protein, putative | 9 | 8 | 7 | 8 | 3 | 6 | 6 | 4 |
| 11 | TGME49_202340-t26_1-p1 | proline dehydrogenase | 9 | 8 | 8 | 8 | 3 | 6 | 6 | 4 |
| 7 | TGME49_202390-t26_1-p1 | mature soluble cytoskeleton protein MSC1B | 9 | 9 | 8 | 8 | 3 | 3 | 3 | 4 |
| 27 | TGME49_202840-t26_1-p1 | FHA domain-containing protein | 10 | 10 | 9 | 8 | 3 | 7 | 7 | 4 |
| 7 | TGME49_203040.P509 | hypothetical protein | 10 | 9 | 9 | 9 | 3 | 3 | 3 | 4 |
| 11 | TGME49_203060-t26_1-p1 | hypothetical protein | 9 | 8 | 8 | 7 | 3 | 6 | 6 | 4 |
| 11 | TGME49_203500-t26_1-p1 | alanine dehydrogenase/pyridine nucleotide transhydrogenase domain-containing protein | 9 | 9 | 8 | 8 | 3 | 3 | 6 | 4 |
| 11 | TGME49_203520-t26_1-p1 | hypothetical protein | 10 | 10 | 7 | 7 | 3 | 6 | 6 | 4 |
| 7 | TGME49_203665-t26_1-p1 | hypothetical protein | 10 | 10 | 10 | 7 | 3 | 4 | 4 | 4 |
| 7 | TGME49_203750-t26_1-p1 | hypothetical protein | 10 | 10 | 10 | 7 | 4 | 5 | 4 | 4 |
| 7 | TGME49_203800-t26_1-p1 | hypothetical protein | 10 | 10 | 10 | 6 | 3 | 3 | 3 | 4 |
| 11 | TGME49_203810.P533 | kelch repeat-containing protein | 9 | 9 | 9 | 9 | 3 | 4 | 7 | 4 |
| 7 | TGME49_203820-t26_1-p1 | hypothetical protein | 10 | 9 | 9 | 8 | 4 | 4 | 4 | 4 |
| 7 | TGME49_203900.P537 | FAM184 domain-containing protein | 10 | 9 | 9 | 8 | 4 | 4 | 4 | 4 |
| 7 | TGME49_204290.P546 | hypothetical protein | 10 | 9 | 8 | 7 | 3 | 3 | 3 | 4 |
| 7 | TGME49_204300-t26_1-p1 | hypothetical protein | 10 | 9 | 8 | 8 | 4 | 3 | 3 | 4 |
| 7 | TGME49_204420.P552 | oocyst wall protein OWP1 | 9 | 9 | 9 | 9 | 3 | 3 | 3 | 4 |
| 11 | TGME49_204440.P553 | cyclic nucleotide-binding domain-containing protein | 10 | 8 | 8 | 6 | 3 | 6 | 5 | 4 |
| 7 | TGME49_205090-t26_1-p1 | Toxoplasma gondii family D protein | 10 | 9 | 9 | 9 | 3 | 3 | 3 | 4 |
| 7 | TGME49_205140.P561 | hypothetical protein | 10 | 8 | 9 | 8 | 4 | 4 | 4 | 4 |
| 27 | TGME49_205265.P563 | transporter, cation channel family protein | 10 | 10 | 10 | 10 | 3 | 6 | 6 | 4 |
| 7 | TGME49_205420.P569 | aspartate-semialdehyde dehydrogenase | 8 | 9 | 8 | 8 | 5 | 3 | 3 | 4 |
| 7 | TGME49_205520.P573 | apicomplexan amino acid transporter ApiAT5-2 | 10 | 9 | 9 | 7 | 3 | 3 | 3 | 4 |
| 11 | TGME49_205590.P575 | centrosomal protein CEP131 | 8 | 9 | 8 | 8 | 3 | 5 | 5 | 4 |
| 7 | TGME49_205650.P577 | AP2 domain transcription factor AP2VIIa-3 | 10 | 9 | 9 | 8 | 4 | 4 | 4 | 4 |
| 11 | TGME49_205700.P580 | cyclophilin precursor | 7 | 7 | 7 | 6 | 3 | 6 | 6 | 4 |
| 7 | TGME49_206300-t26_1-p1 | hypothetical protein | 10 | 9 | 8 | 8 | 4 | 5 | 4 | 4 |
| 7 | TGME49_206570.P592 | kinesin, putative | 9 | 9 | 8 | 7 | 3 | 4 | 4 | 4 |
| 7 | TGME49_206710-t26_1-p1 | hypothetical protein | 10 | 9 | 8 | 7 | 5 | 4 | 4 | 4 |
| 27 | TGME49_207080-t26_1-p1 | histone lysine acetyltransferase MYST-B | 10 | 9 | 9 | 6 | 5 | 8 | 8 | 4 |
| 11 | TGME49_207460-t26_1-p1 | ras-related protein RAB5B | 7 | 7 | 7 | 7 | 3 | 6 | 6 | 4 |
| 27 | TGME49_207590.P619 | protein phosphatase PPM12 | 9 | 9 | 9 | 8 | 5 | 7 | 6 | 4 |
| 11 | TGME49_207630.P622 | peptidyl-tRNA hydrolase domain-containing protein | 9 | 8 | 8 | 7 | 4 | 6 | 5 | 4 |
| 27 | TGME49_207750-t26_1-p1 | glycosyltransferase family 17 protein | 10 | 9 | 9 | 9 | 3 | 8 | 7 | 4 |
| 7 | TGME49_208370.P639 | dense granule protein GRA46 | 10 | 9 | 9 | 8 | 4 | 4 | 4 | 4 |
| 7 | TGME49_208410.P641 | transporter, putative | 10 | 10 | 10 | 6 | 3 | 3 | 3 | 4 |

| | | | | | | | | | | |
|---|---|---|---|---|---|---|---|---|---|---|
| 7 | TGME49_208760.P654 | hypothetical protein | 10 | 8 | 9 | 8 | 5 | 3 | 3 | 4 |
| 27 | TGME49_209450.P679 | TLD protein | 10 | 10 | 9 | 7 | 4 | 7 | 5 | 4 |
| 7 | TGME49_209460-t26_1-p1 | hypothetical protein | 10 | 9 | 8 | 7 | 5 | 3 | 3 | 4 |
| 27 | TGME49_209490-t26_1-p1 | hypothetical protein | 9 | 8 | 8 | 8 | 5 | 7 | 7 | 4 |
| 11 | TGME49_209520-t26_1-p1 | WD domain, G-beta repeat-containing protein | 10 | 8 | 7 | 7 | 3 | 5 | 5 | 4 |
| 11 | TGME49_209610.P685 | oocyst wall protein OWP2 | 9 | 9 | 8 | 7 | 3 | 5 | 4 | 4 |
| 11 | TGME49_209940.P699 | transporter/permease protein | 9 | 8 | 8 | 8 | 3 | 6 | 3 | 4 |
| 11 | TGME49_210260-t26_1-p1 | NAD-dependent glycerol-3-phosphate dehydrogenase | 9 | 9 | 9 | 9 | 3 | 3 | 8 | 4 |
| 7 | TGME49_210380.P709 | transmembrane protein 184, putative | 10 | 9 | 9 | 7 | 3 | 3 | 3 | 4 |
| 11 | TGME49_210690.P712 | ribosomal protein RPS6 | 8 | 8 | 7 | 7 | 3 | 6 | 7 | 4 |
| 27 | TGME49_210750-t26_1-p1 | Ferredoxin-fold anticodon binding domain-containing protein | 10 | 9 | 9 | 8 | 4 | 6 | 6 | 4 |
| 7 | TGME49_211150-t26_1-p1 | hypothetical protein | 10 | 10 | 10 | 7 | 4 | 4 | 4 | 4 |
| 11 | TGME49_211330.P732 | methionine aminopeptidase 1c | 10 | 8 | 8 | 7 | 3 | 4 | 5 | 4 |
| 27 | TGME49_211410-t26_1-p1 | translation initiation factor sui1 protein | 10 | 8 | 9 | 7 | 5 | 8 | 7 | 4 |
| 27 | TGME49_211670.P746 | S1 RNA binding domain-containing protein | 9 | 8 | 8 | 7 | 4 | 7 | 6 | 4 |
| 27 | TGME49_212240.P758 | beta-1 tubulin, putative | 9 | 9 | 9 | 9 | 3 | 9 | 3 | 4 |
| 27 | TGME49_212250.P759 | XPG N-terminal domain-containing protein | 10 | 10 | 10 | 6 | 5 | 8 | 8 | 4 |
| 7 | TGME49_212770.P766 | inner membrane complex protein IMC34 | 10 | 10 | 9 | 8 | 5 | 3 | 4 | 4 |
| 7 | TGME49_212820-t26_1-p1 | ubiquitin family protein | 10 | 9 | 8 | 8 | 5 | 4 | 4 | 4 |
| 7 | TGME49_212880.P768 | centrosomal protein CEP250 | 10 | 10 | 10 | 8 | 4 | 4 | 3 | 4 |
| 7 | TGME49_212970-t26_1-p1 | protein kinase (incomplete catalytic triad) | 10 | 8 | 9 | 7 | 3 | 3 | 3 | 4 |
| 11 | TGME49_212980.P774 | KH domain-containing protein | 10 | 9 | 8 | 8 | 4 | 6 | 6 | 4 |
| 11 | TGME49_213530.P794 | hypothetical protein | 9 | 3 | 9 | 7 | 3 | 7 | 7 | 4 |
| 27 | TGME49_214350-t26_1-p1 | GTP-binding protein, putative | 9 | 9 | 9 | 7 | 3 | 8 | 9 | 4 |
| 7 | TGME49_214750.P831 | hypothetical protein | 10 | 10 | 10 | 8 | 4 | 4 | 4 | 4 |
| 7 | TGME49_214800-t26_1-p1 | hypothetical protein | 10 | 9 | 8 | 8 | 5 | 4 | 3 | 4 |
| 7 | TGME49_215210.P848 | F-box domain-containing protein | 10 | 9 | 8 | 7 | 3 | 4 | 4 | 4 |
| 11 | TGME49_215440-t26_1-p1 | WWE domain-containing protein | 10 | 9 | 8 | 7 | 3 | 7 | 5 | 4 |
| 7 | TGME49_215700-t26_1-p1 | phosphatidylinositol 3- and 4-kinase | 10 | 10 | 8 | 7 | 3 | 5 | 3 | 4 |
| 7 | TGME49_215890.P867 | hypothetical protein | 10 | 10 | 9 | 7 | 5 | 3 | 3 | 4 |
| 11 | TGME49_216150-t26_1-p1 | peptidase family M3 protein | 9 | 9 | 9 | 8 | 3 | 6 | 6 | 4 |
| 7 | TGME49_216335-t26_1-p1 | hypothetical protein | 10 | 10 | 9 | 7 | 3 | 3 | 3 | 4 |
| 7 | TGME49_216380.P883 | phospholipid-translocating P-type ATPase, flippase subfamily protein | 10 | 9 | 9 | 8 | 3 | 3 | 3 | 4 |
| 7 | TGME49_216390-t26_1-p1 | RNA methyltransferase, TrmH family protein | 10 | 8 | 8 | 7 | 4 | 3 | 5 | 4 |
| 7 | TGME49_216610.P889 | ribosomal RNA large subunit methyltransferase J protein | 10 | 10 | 9 | 9 | 3 | 3 | 3 | 4 |
| 11 | TGME49_216900.P896 | cytokine induced apoptosis inhibitor 1 family protein | 8 | 7 | 7 | 6 | 3 | 5 | 5 | 4 |
| 7 | TGME49_217680.P919 | dense granule protein GRA57 | 10 | 10 | 9 | 7 | 5 | 4 | 4 | 4 |
| 7 | TGME49_217790.P925 | S1 RNA binding domain-containing protein | 10 | 10 | 9 | 7 | 5 | 4 | 4 | 4 |
| 27 | TGME49_217850-t26_1-p1 | Sucrose-6F-phosphate phosphohydrolase | 9 | 9 | 9 | 8 | 5 | 7 | 6 | 4 |
| 27 | TGME49_217880-t26_1-p1 | RNA-binding protein Nova-1, putative | 9 | 8 | 8 | 8 | 3 | 7 | 6 | 4 |
| 11 | TGME49_217890.P928 | peroxiredoxin PRX1 | 7 | 7 | 7 | 7 | 3 | 5 | 6 | 4 |
| 11 | TGME49_218215.P932 | hypothetical protein | 10 | 9 | 8 | 7 | 3 | 6 | 5 | 4 |
| 7 | TGME49_218300.P938 | zinc finger (CCCH type) motif-containing protein | 10 | 10 | 9 | 7 | 5 | 5 | 3 | 4 |

| | | | | | | | | | | |
|---|---|---|---|---|---|---|---|---|---|---|
| 11 | TGME49_218400.P939 | NEK kinase | 6 | 9 | 8 | 8 | 3 | 3 | 7 | 4 |
| 11 | TGME49_218540-t26_1-p1 | peptidase S15, putative | 9 | 8 | 8 | 7 | 4 | 5 | 5 | 4 |
| 27 | TGME49_218580-t26_1-p1 | RNA methyltransferase, TrmH family protein | 10 | 10 | 9 | 9 | 3 | 7 | 7 | 4 |
| 11 | TGME49_218780-t26_1-p1 | phosphoserine aminotransferase, putative | 9 | 9 | 8 | 6 | 3 | 5 | 6 | 4 |
| 11 | TGME49_218790-t26_1-p1 | elongation factor G C-terminus domain-containing protein | 10 | 9 | 9 | 6 | 3 | 5 | 4 | 4 |
| 11 | TGME49_218910.P955 | hypothetical protein | 10 | 10 | 9 | 8 | 3 | 5 | 5 | 4 |
| 7 | TGME49_219180-t26_1-p1 | acetyltransferase, GNAT family protein | 10 | 9 | 8 | 7 | 3 | 3 | 3 | 4 |
| 7 | TGME49_219230.P965 | AMP-binding enzyme domain-containing protein | 10 | 9 | 10 | 9 | 3 | 3 | 3 | 4 |
| 11 | TGME49_219260.P967 | cation-transporting ATPase, putative | 10 | 10 | 9 | 6 | 3 | 6 | 5 | 4 |
| 11 | TGME49_219430.P969 | tRNA ligase class II core domain (G, H, P, S and T) domain-containing protein | 8 | 8 | 8 | 7 | 4 | 6 | 5 | 4 |
| 7 | TGME49_219450.P970 | WD domain, G-beta repeat-containing protein | 10 | 10 | 8 | 9 | 3 | 3 | 3 | 4 |
| 7 | TGME49_219500.P972 | hypothetical protein | 10 | 9 | 8 | 8 | 5 | 3 | 3 | 4 |
| 27 | TGME49_219650.P979 | transporter, small conductance mechanosensitive ion channel (MscS) family protein | 10 | 10 | 9 | 8 | 3 | 7 | 7 | 4 |
| 7 | TGME49_219660.P980 | hypothetical protein | 10 | 10 | 9 | 6 | 4 | 3 | 3 | 4 |
| 7 | TGME49_219710-t26_1-p1 | hypothetical protein | 10 | 10 | 9 | 7 | 4 | 4 | 4 | 4 |
| 7 | TGME49_219810.P984 | dense granule protein GRA40 | 10 | 9 | 8 | 7 | 3 | 4 | 4 | 4 |
| 11 | TGME49_220280.P993 | mucin family glycoprotein | 9 | 6 | 8 | 8 | 3 | 3 | 5 | 4 |
| 11 | TGME49_220420.P998 | 3'5'-cyclic nucleotide phosphodiesterase PDE5 | 10 | 9 | 9 | 8 | 3 | 6 | 4 | 4 |
| 7 | TGME49_221310.P1018 | aminopeptidase N1 | 10 | 8 | 10 | 9 | 3 | 3 | 3 | 4 |
| 11 | TGME49_221360-t26_1-p1 | autophagy-related protein ATG6, putative | 9 | 9 | 8 | 7 | 3 | 4 | 6 | 4 |
| 27 | TGME49_221420.P1022 | Chromosome-associated kinesin KLP1, putative | 10 | 10 | 10 | 10 | 3 | 9 | 8 | 4 |
| 27 | TGME49_221470-t26_1-p1 | hypothetical protein | 9 | 9 | 9 | 8 | 3 | 7 | 7 | 4 |
| 27 | TGME49_222100-t26_1-p1 | effector protein SOS1 | 10 | 10 | 10 | 10 | 3 | 8 | 6 | 4 |
| 27 | TGME49_222660.P1061 | non-discharge partner NDP1 | 10 | 10 | 10 | 9 | 3 | 10 | 3 | 4 |
| 11 | TGME49_222910.P1069 | phosphoglycerate mutase PGM3 | 9 | 8 | 8 | 8 | 4 | 5 | 4 | 4 |
| 7 | TGME49_223060.P1072 | MORN repeat-containing protein | 10 | 10 | 10 | 10 | 3 | 3 | 3 | 4 |
| 11 | TGME49_223140-t26_1-p1 | tRNA import protein tRIP, putative | 9 | 8 | 8 | 8 | 3 | 6 | 6 | 4 |
| 7 | TGME49_223430.P1078 | hypothetical protein | 10 | 10 | 10 | 9 | 3 | 3 | 3 | 4 |
| 11 | TGME49_223672-t26_1-p1 | 3'(2'),5'-bisphosphate nucleotidase | 8 | 8 | 8 | 7 | 3 | 5 | 6 | 4 |
| 7 | TGME49_223810.P1091 | magnesium transporter, putative | 10 | 9 | 8 | 7 | 4 | 3 | 3 | 4 |
| 11 | TGME49_223830.P1092 | fasciclin domain-containing protein | 10 | 7 | 9 | 8 | 3 | 5 | 5 | 4 |
| 7 | TGME49_223900.P1097 | hypothetical protein | 10 | 9 | 9 | 7 | 3 | 3 | 3 | 4 |
| 11 | TGME49_223910-t26_1-p1 | acyltransferase domain-containing protein | 9 | 6 | 8 | 6 | 4 | 5 | 5 | 4 |
| 7 | TGME49_223950.P1100 | hypothetical protein | 10 | 9 | 9 | 8 | 4 | 4 | 4 | 4 |
| 7 | TGME49_224460.P1117 | aminopeptidase N3 | 10 | 10 | 9 | 9 | 3 | 3 | 3 | 4 |
| 7 | TGME49_224610.P1122 | GYF domain-containing protein | 10 | 10 | 9 | 8 | 4 | 4 | 4 | 4 |
| 27 | TGME49_224660-t26_1-p1 | transcription factor s-ii (tfiis), central domain-containing protein | 10 | 10 | 9 | 9 | 5 | 6 | 6 | 4 |
| 7 | TGME49_225200.P1157 | hypothetical protein | 10 | 10 | 9 | 7 | 3 | 3 | 4 | 4 |
| 7 | TGME49_225560-t26_1-p1 | inner membrane complex protein IMC41 | 10 | 8 | 8 | 8 | 3 | 3 | 3 | 4 |
| 7 | TGME49_225745.P1178 | hypothetical protein | 10 | 10 | 10 | 8 | 3 | 4 | 5 | 4 |
| 27 | TGME49_225770.P1179 | tyrosine kinase-like protein TKL8 | 10 | 9 | 8 | 8 | 4 | 7 | 6 | 4 |

| | | | | | | | | | | |
|---|---|---|---|---|---|---|---|---|---|---|
| 7 | TGME49_225850-t26_1-p1 | peptidase, M28 family protein | 10 | 10 | 10 | 8 | 5 | 3 | 5 | 4 |
| 7 | TGME49_225910-t26_1-p1 | hypothetical protein | 10 | 9 | 9 | 7 | 3 | 3 | 3 | 4 |
| 11 | TGME49_225950.P1186 | zinc finger (CCCH type) motif-containing protein | 10 | 10 | 9 | 8 | 3 | 6 | 5 | 4 |
| 27 | TGME49_226090-t26_1-p1 | DEAD/DEAH box helicase domain-containing protein | 10 | 9 | 9 | 8 | 4 | 6 | 6 | 4 |
| 11 | TGME49_226220-t26_1-p1 | alveolin domain containing intermediate filament IMC9 | 10 | 9 | 8 | 7 | 3 | 6 | 6 | 4 |
| 11 | TGME49_226300-t26_1-p1 | 2,4-dienoyl CoA reductase 2, peroxisomal family protein | 9 | 8 | 8 | 7 | 3 | 6 | 5 | 4 |
| 11 | TGME49_226320-t26_1-p1 | RNA binding protein, putative | 10 | 8 | 7 | 6 | 3 | 5 | 5 | 4 |
| 7 | TGME49_226390.P1205 | hypothetical protein | 10 | 9 | 8 | 8 | 4 | 3 | 5 | 4 |
| 7 | TGME49_226420.P1207 | dense granule protein GRA69 | 10 | 9 | 9 | 8 | 4 | 3 | 3 | 4 |
| 7 | TGME49_226540-t26_1-p1 | protein kinase | 9 | 9 | 9 | 8 | 3 | 3 | 3 | 4 |
| 7 | TGME49_226690-t26_1-p1 | hypothetical protein | 10 | 9 | 9 | 7 | 3 | 3 | 3 | 4 |
| 27 | TGME49_226910.P1230 | Amylo-alpha-1,6-glucosidase | 10 | 10 | 10 | 9 | 4 | 10 | 9 | 4 |
| 7 | TGME49_227080-t26_1-p1 | PCIF1 WW domain-containing protein | 10 | 9 | 8 | 8 | 3 | 3 | 4 | 4 |
| 7 | TGME49_227090.P1239 | aspartokinase | 9 | 9 | 9 | 8 | 3 | 3 | 3 | 4 |
| 11 | TGME49_227260.P1240 | RIO1 family protein | 9 | 9 | 8 | 8 | 3 | 7 | 3 | 4 |
| 7 | TGME49_227390-t26_1-p1 | SET domain-containing protein | 10 | 10 | 7 | 8 | 5 | 3 | 3 | 4 |
| 27 | TGME49_227450.P1249 | hydrolase, NUDIX family protein | 10 | 10 | 9 | 8 | 3 | 6 | 6 | 4 |
| 7 | TGME49_227952.P1268 | 14-3-3 superfamily protein | 9 | 8 | 8 | 7 | 4 | 4 | 4 | 4 |
| 7 | TGME49_228410.P1294 | Tubulin-tyrosine ligase family protein | 10 | 10 | 9 | 8 | 3 | 4 | 4 | 4 |
| 11 | TGME49_228680.P1300 | mitochondrial carrier superfamily protein | 9 | 8 | 8 | 6 | 3 | 5 | 5 | 4 |
| 11 | TGME49_229150.P1311 | transporter, major facilitator family protein | 10 | 9 | 8 | 7 | 3 | 5 | 5 | 4 |
| 11 | TGME49_229200.P1314 | cyclin CYC1 | 9 | 8 | 8 | 6 | 4 | 5 | 4 | 4 |
| 7 | TGME49_229260.P1316 | basal complex component BCC4 | 10 | 10 | 9 | 6 | 3 | 4 | 3 | 4 |
| 7 | TGME49_229310-t26_1-p1 | mediator complex subunit MED14 | 10 | 10 | 8 | 7 | 5 | 4 | 4 | 4 |
| 11 | TGME49_229915.P1343 | dynein regulatory complex protein 9, putative | 9 | 8 | 8 | 8 | 3 | 6 | 5 | 4 |
| 7 | TGME49_230030.P1348 | WD domain, G-beta repeat-containing protein | 10 | 9 | 8 | 7 | 3 | 4 | 4 | 4 |
| 11 | TGME49_230150.P1357 | ChAPs (Chs5p-Arf1p-binding proteins) protein | 10 | 10 | 8 | 6 | 3 | 5 | 4 | 4 |
| 27 | TGME49_230220.P1362 | vacuolar protein sorting-associated protein 11 | 10 | 10 | 10 | 9 | 3 | 6 | 6 | 4 |
| 7 | TGME49_230500.P1373 | hypothetical protein | 10 | 8 | 9 | 7 | 4 | 3 | 3 | 4 |
| 7 | TGME49_230555.P1376 | adenylate cyclase alpha 3 | 10 | 9 | 9 | 9 | 5 | 3 | 3 | 4 |
| 7 | TGME49_230580.P1378 | hypothetical protein | 10 | 8 | 9 | 7 | 5 | 4 | 4 | 4 |
| 7 | TGME49_230630.P1380 | hypothetical protein | 10 | 10 | 9 | 7 | 5 | 3 | 4 | 4 |
| 7 | TGME49_230650.P1381 | DUF2362 domain-containing protein | 10 | 10 | 9 | 8 | 3 | 3 | 4 | 4 |
| 7 | TGME49_230700-t26_1-p1 | hypothetical protein | 10 | 10 | 9 | 7 | 5 | 4 | 4 | 4 |
| 11 | TGME49_230905-t26_1-p1 | hydrolase, alpha/beta fold family protein | 10 | 7 | 8 | 7 | 3 | 6 | 6 | 4 |
| 7 | TGME49_230980.P1394 | myosin I | 10 | 10 | 10 | 9 | 3 | 3 | 3 | 4 |
| 7 | TGME49_231400.P1416 | tubulin/FtsZ family, GTPase domain-containing protein | 9 | 9 | 8 | 8 | 3 | 3 | 3 | 4 |
| 7 | TGME49_231440-t26_1-p1 | LsmAD domain-containing protein | 10 | 10 | 10 | 8 | 4 | 4 | 4 | 4 |
| 7 | TGME49_231770-t26_1-p1 | alpha-tubulin I, putative | 9 | 7 | 9 | 9 | 3 | 3 | 3 | 4 |
| 7 | TGME49_231780-t26_1-p1 | hypothetical protein | 9 | 9 | 8 | 7 | 4 | 3 | 3 | 4 |
| 7 | TGME49_231800.P1424 | helicase, putative | 10 | 10 | 9 | 7 | 3 | 5 | 4 | 4 |
| 27 | TGME49_232540.P1471 | FAD binding domain-containing protein | 10 | 10 | 9 | 7 | 5 | 6 | 7 | 4 |

| | | | | | | | | | | |
|---|---|---|---|---|---|---|---|---|---|---|
| 11 | TGME49_232600.P1474 | patatin-like phospholipase PL1 | 9 | 9 | 8 | 8 | 3 | 5 | 5 | 4 |
| 11 | TGME49_232620.P1476 | hypothetical protein | 10 | 9 | 9 | 8 | 3 | 6 | 4 | 4 |
| 11 | TGME49_232750-t26_1-p1 | 23S rRNA (adenine(1618)-N(6))-methyltransferase, putative | 10 | 9 | 8 | 7 | 3 | 3 | 6 | 4 |
| 7 | TGME49_232970.P1486 | origin recognition complex subunit 5, putative | 10 | 10 | 9 | 7 | 5 | 4 | 4 | 4 |
| 7 | TGME49_233180.P1497 | hypothetical protein | 10 | 8 | 8 | 7 | 4 | 3 | 3 | 4 |
| 7 | TGME49_233340.P1505 | mini-chromosome maintenance complex-binding protein, putative | 10 | 9 | 8 | 8 | 5 | 3 | 3 | 4 |
| 11 | TGME49_233370-t26_1-p1 | hypothetical protein | 7 | 7 | 7 | 6 | 3 | 5 | 5 | 4 |
| 7 | TGME49_233390-t26_1-p1 | hypothetical protein | 10 | 9 | 8 | 8 | 5 | 3 | 3 | 4 |
| 27 | TGME49_233790.P1516 | serine/threonine protein kinase AktR, putative | 10 | 8 | 9 | 6 | 3 | 7 | 7 | 4 |
| 11 | TGME49_233830.P1519 | THUMP domain-containing protein | 8 | 7 | 7 | 7 | 4 | 4 | 5 | 4 |
| 7 | TGME49_233860.P1521 | DALR anticodon binding domain-containing protein | 10 | 9 | 9 | 7 | 5 | 3 | 3 | 4 |
| 27 | TGME49_233905.P1523 | 5'-AMP-activated protein kinase subunit alpha | 10 | 9 | 9 | 8 | 5 | 8 | 8 | 4 |
| 27 | TGME49_233930.P1524 | CAF1 family ribonuclease | 10 | 9 | 9 | 9 | 3 | 8 | 7 | 4 |
| 11 | TGME49_234220-t26_1-p1 | hypothetical protein | 10 | 9 | 8 | 6 | 3 | 5 | 5 | 4 |
| 27 | TGME49_235150.P1550 | phosphate transporter PT2 | 10 | 9 | 9 | 9 | 3 | 8 | 8 | 4 |
| 11 | TGME49_235580.P1564 | hypothetical protein | 9 | 8 | 7 | 7 | 3 | 5 | 5 | 4 |
| 11 | TGME49_235650.P1569 | carrier superfamily protein | 9 | 9 | 8 | 7 | 4 | 5 | 5 | 4 |
| 11 | TGME49_235700-t26_1-p1 | sedoheptulose-1,7-bisphosphatase | 8 | 6 | 8 | 8 | 3 | 5 | 6 | 4 |
| 27 | TGME49_235950.P1580 | subtilisin SUB8 | 10 | 10 | 10 | 10 | 5 | 8 | 6 | 4 |
| 27 | TGME49_235990.P1582 | hypothetical protein | 10 | 9 | 9 | 7 | 5 | 6 | 7 | 4 |
| 7 | TGME49_236040.P1585 | fructose-1,6-bisphosphate aldolase | 8 | 9 | 8 | 8 | 3 | 3 | 3 | 4 |
| 11 | TGME49_236840-t26_1-p1 | zinc finger (C-x8-C-x5-C-x3-H)-2, putative | 9 | 9 | 8 | 6 | 3 | 4 | 6 | 4 |
| 11 | TGME49_236860.P1603 | haloacid dehalogenase family hydrolase domain-containing protein | 10 | 10 | 10 | 10 | 3 | 3 | 7 | 4 |
| 7 | TGME49_236930.P1608 | hypothetical protein | 10 | 10 | 10 | 7 | 4 | 4 | 3 | 4 |
| 27 | TGME49_236975-t26_1-p1 | Toxoplasma gondii family D protein | 10 | 9 | 9 | 9 | 3 | 8 | 8 | 4 |
| 11 | TGME49_237020-t26_1-p1 | exonuclease | 10 | 10 | 9 | 7 | 3 | 5 | 5 | 4 |
| 11 | TGME49_237500.P1631 | protein phosphatase PPM3A, putative | 10 | 9 | 8 | 7 | 3 | 5 | 5 | 4 |
| 11 | TGME49_237570-t26_1-p1 | leucine carboxyl methyltransferase | 9 | 8 | 8 | 7 | 4 | 6 | 6 | 4 |
| 7 | TGME49_237585-t26_1-p1 | hypothetical protein | 10 | 10 | 10 | 9 | 3 | 3 | 3 | 4 |
| 11 | TGME49_237600-t26_1-p1 | Armadillo/beta-catenin family repeat-containing protein | 10 | 10 | 7 | 7 | 3 | 6 | 4 | 4 |
| 11 | TGME49_237830.P1635 | DNA polymerase I domain-containing protein | 10 | 10 | 9 | 7 | 4 | 6 | 6 | 4 |
| 27 | TGME49_237890-t26_1-p1 | calcium-dependent protein kinase CDPK4 | 10 | 10 | 9 | 9 | 3 | 6 | 8 | 4 |
| 27 | TGME49_238040.P1640 | protein disulfide-isomerase domain-containing protein | 10 | 10 | 10 | 9 | 5 | 8 | 8 | 4 |
| 11 | TGME49_238140-t26_1-p1 | WD repeat-containing protein | 10 | 8 | 7 | 6 | 4 | 5 | 5 | 4 |
| 11 | TGME49_238170.P1644 | IMC-associated protein IAP3 | 10 | 9 | 8 | 6 | 4 | 5 | 5 | 4 |
| 11 | TGME49_238915-t26_1-p1 | hypothetical protein | 10 | 10 | 10 | 9 | 3 | 5 | 5 | 4 |
| 27 | TGME49_239420.P1674 | protein kinase | 10 | 10 | 9 | 8 | 5 | 7 | 6 | 4 |
| 27 | TGME49_239475-t26_1-p1 | hypothetical protein | 10 | 10 | 10 | 10 | 4 | 6 | 5 | 4 |
| 27 | TGME49_239480-t26_1-p1 | RNB family domain-containing protein | 10 | 10 | 10 | 7 | 5 | 7 | 6 | 4 |
| 27 | TGME49_239670-t26_1-p1 | hypothetical protein | 10 | 8 | 9 | 8 | 3 | 7 | 7 | 4 |
| 7 | TGME49_239700-t26_1-p1 | regulator of chromosome condensation (RCC1) repeat-containing protein | 10 | 10 | 9 | 9 | 5 | 3 | 3 | 4 |

| | | | | | | | | | | |
|---|---|---|---|---|---|---|---|---|---|---|
| 27 | TGME49_240380.P1700 | conoid gliding protein CGP | 10 | 5 | 3 | 10 | 4 | 9 | 9 | 4 |
| 7 | TGME49_240460.P1703 | AP2 domain transcription factor AP2VI-1 | 10 | 8 | 8 | 8 | 3 | 3 | 3 | 4 |
| 11 | TGME49_240560.P1707 | MAPEG family protein | 7 | 7 | 7 | 6 | 3 | 5 | 5 | 4 |
| 7 | TGME49_240880.P1719 | hypothetical protein | 10 | 10 | 9 | 6 | 3 | 3 | 3 | 4 |
| 7 | TGME49_242020-t26_1-p1 | mannosyl-oligosaccharide glucosidase | 10 | 10 | 9 | 9 | 3 | 3 | 3 | 4 |
| 11 | TGME49_242390-t26_1-p1 | enoyl-coa hydratase/isomerase family protein | 8 | 7 | 7 | 6 | 3 | 3 | 6 | 4 |
| 11 | TGME49_242570-t26_1-p1 | armadillo repeat-containing protein | 10 | 9 | 9 | 8 | 3 | 5 | 5 | 4 |
| 7 | TGME49_242580-t26_1-p1 | iron only hydrogenase large subunit, c-terminal domain-containing protein | 10 | 10 | 9 | 8 | 5 | 3 | 3 | 4 |
| 7 | TGME49_243330.P1757 | hypothetical protein | 9 | 10 | 10 | 8 | 3 | 3 | 3 | 4 |
| 11 | TGME49_243430.P1760 | OTU domain-containing protein OTUD3C | 10 | 9 | 8 | 7 | 4 | 5 | 4 | 4 |
| 11 | TGME49_243500.P1762 | CAM kinase, SNF1/AMK1 family ToxPK1 | 8 | 9 | 9 | 8 | 3 | 7 | 4 | 4 |
| 7 | TGME49_243750-t26_1-p1 | tetratricopeptide repeat-containing protein | 10 | 8 | 8 | 8 | 5 | 5 | 3 | 4 |
| 11 | TGME49_244040.P1779 | PIKfyve regulatory protein | 10 | 10 | 8 | 7 | 3 | 6 | 4 | 4 |
| 7 | TGME49_244060-t26_1-p1 | ankyrin repeat-containing protein | 10 | 8 | 8 | 7 | 4 | 3 | 5 | 4 |
| 11 | TGME49_244270-t26_1-p1 | ATP-binding cassette G family transporter ABCG87 | 10 | 10 | 9 | 6 | 3 | 3 | 6 | 4 |
| 11 | TGME49_244360-t26_1-p1 | OST-HTH associated domain-containing protein | 9 | 7 | 7 | 7 | 3 | 6 | 5 | 4 |
| 11 | TGME49_244440-t26_1-p1 | adenosine transporter AT1 | 9 | 9 | 8 | 7 | 3 | 3 | 6 | 4 |
| 7 | TGME49_244470.P1790 | apical ring protein RNG2 | 10 | 10 | 10 | 9 | 4 | 4 | 4 | 4 |
| 11 | TGME49_244510.P1791 | AP2 domain transcription factor AP2VI-3 | 10 | 10 | 8 | 6 | 3 | 5 | 5 | 4 |
| 11 | TGME49_244590.P1796 | katanin-like family protein | 9 | 9 | 9 | 7 | 3 | 7 | 4 | 4 |
| 27 | TGME49_244620.P1798 | NEK kinase | 8 | 8 | 8 | 8 | 3 | 7 | 6 | 4 |
| 7 | TGME49_244850-t26_1-p1 | GMC oxidoreductase | 10 | 10 | 10 | 6 | 3 | 3 | 3 | 4 |
| 11 | TGME49_245590-t26_1-p1 | rhomboid protease ROM6 | 9 | 8 | 7 | 7 | 3 | 4 | 5 | 4 |
| 27 | TGME49_245610-t26_1-p1 | hypothetical protein | 10 | 9 | 8 | 7 | 4 | 7 | 6 | 4 |
| 27 | TGME49_246040-t26_1-p1 | MIF4G domain-containing protein | 10 | 10 | 10 | 9 | 3 | 8 | 8 | 4 |
| 7 | TGME49_246080.P1822 | NAD dependent epimerase/dehydratase family protein | 9 | 9 | 8 | 7 | 3 | 4 | 3 | 4 |
| 11 | TGME49_246460-t26_1-p1 | hypothetical protein | 9 | 7 | 7 | 6 | 4 | 5 | 5 | 4 |
| 11 | TGME49_246580-t26_1-p1 | hypothetical protein | 9 | 8 | 8 | 7 | 3 | 4 | 5 | 4 |
| 27 | TGME49_246760.P1840 | hypothetical protein | 10 | 10 | 8 | 9 | 3 | 7 | 6 | 4 |
| 11 | TGME49_246910-t26_1-p1 | histone lysine methyltransferase, SET, putative | 9 | 9 | 7 | 7 | 3 | 5 | 4 | 4 |
| 27 | TGME49_246920-t26_1-p1 | glutathione reductase | 9 | 9 | 9 | 9 | 3 | 8 | 7 | 4 |
| 7 | TGME49_246982-t26_1-p1 | hypothetical protein | 10 | 9 | 8 | 7 | 3 | 4 | 4 | 4 |
| 7 | TGME49_247040-t26_1-p1 | hypothetical protein | 10 | 10 | 9 | 7 | 5 | 4 | 4 | 4 |
| 7 | TGME49_247970.P1871 | hypothetical protein | 10 | 10 | 10 | 9 | 4 | 4 | 3 | 4 |
| 11 | TGME49_248270-t26_1-p1 | zinc finger (CCCH type) motif-containing protein | 10 | 8 | 7 | 6 | 3 | 4 | 5 | 4 |
| 7 | TGME49_248590.P1895 | hypothetical protein | 10 | 10 | 9 | 9 | 3 | 3 | 3 | 4 |
| 27 | TGME49_248600.P1896 | aspartate aminotransferase | 9 | 9 | 9 | 8 | 3 | 7 | 7 | 4 |
| 7 | TGME49_248750.P1904 | hypothetical protein | 10 | 10 | 9 | 8 | 3 | 3 | 3 | 4 |
| 11 | TGME49_248950-t26_1-p1 | carrier superfamily protein | 9 | 8 | 8 | 6 | 4 | 6 | 6 | 4 |
| 11 | TGME49_248970.P1912 | leucine rich repeat-containing protein | 8 | 8 | 8 | 7 | 4 | 6 | 6 | 4 |
| 7 | TGME49_249580.P1937 | apicomplexan amino acid transporter ApiAT6-3 | 10 | 9 | 8 | 8 | 3 | 6 | 3 | 4 |
| 11 | TGME49_249702-t26_1-p1 | MC family transporter, putative | 8 | 8 | 7 | 7 | 3 | 6 | 5 | 4 |

| | | | | | | | | | | |
|---|---|---|---|---|---|---|---|---|---|---|
| 11 | TGME49_249860-t26_1-p1 | hypothetical protein | 10 | 9 | 9 | 6 | 3 | 6 | 6 | 4 |
| 27 | TGME49_249900.P1950 | adenine nucleotide translocator | 8 | 8 | 8 | 8 | 3 | 6 | 8 | 4 |
| 11 | TGME49_249960.P1952 | inner dynein arm light chain, putative | 8 | 7 | 7 | 7 | 3 | 6 | 5 | 4 |
| 7 | TGME49_250010-t26_1-p1 | SUN-like protein SLP1 | 10 | 9 | 8 | 8 | 4 | 5 | 4 | 4 |
| 11 | TGME49_250360-t26_1-p1 | alpha/beta hydrolase fold domain-containing protein | 10 | 9 | 8 | 8 | 4 | 4 | 6 | 4 |
| 7 | TGME49_251460-t26_1-p1 | hypothetical protein | 10 | 9 | 8 | 7 | 3 | 4 | 4 | 4 |
| 27 | TGME49_251480.P1975 | DEAD/DEAH box helicase domain-containing protein | 10 | 10 | 9 | 9 | 3 | 8 | 6 | 4 |
| 27 | TGME49_252210-t26_1-p1 | pentatricopeptide repeat domain-containing protein | 10 | 10 | 9 | 9 | 3 | 7 | 7 | 4 |
| 11 | TGME49_252300.P1991 | hypothetical protein | 10 | 10 | 9 | 8 | 3 | 5 | 5 | 4 |
| 7 | TGME49_252340.P1993 | MBOAT domain-containing protein | 10 | 10 | 9 | 7 | 3 | 3 | 3 | 4 |
| 11 | TGME49_253120.P2012 | mandelonitrile lyase, putative | 10 | 9 | 10 | 10 | 3 | 3 | 8 | 4 |
| 27 | TGME49_253150.P2013 | hypothetical protein | 10 | 10 | 10 | 10 | 3 | 8 | 3 | 4 |
| 7 | TGME49_253400-t26_1-p1 | hypothetical protein | 10 | 9 | 8 | 7 | 3 | 3 | 3 | 4 |
| 11 | TGME49_253470-t26_1-p1 | alveolin domain containing intermediate filament IMC13 | 9 | 8 | 8 | 8 | 3 | 5 | 6 | 4 |
| 11 | TGME49_253510-t26_1-p1 | transporter/permease protein | 8 | 8 | 8 | 7 | 3 | 6 | 5 | 4 |
| 11 | TGME49_253580.P2029 | CMGC kinase, CDK family | 10 | 9 | 9 | 6 | 3 | 5 | 4 | 4 |
| 7 | TGME49_253910.P2038 | hypothetical protein | 10 | 9 | 9 | 8 | 5 | 3 | 3 | 4 |
| 27 | TGME49_254010.P2042 | serine carboxypeptidase s28 protein | 10 | 9 | 9 | 8 | 3 | 7 | 5 | 4 |
| 11 | TGME49_254480-t26_1-p1 | WD domain, G-beta repeat-containing protein | 10 | 9 | 8 | 6 | 3 | 4 | 5 | 4 |
| 7 | TGME49_254810-t26_1-p1 | hypothetical protein | 10 | 10 | 9 | 7 | 5 | 4 | 4 | 4 |
| 27 | TGME49_254930-t26_1-p1 | hypothetical protein | 10 | 10 | 9 | 8 | 3 | 6 | 6 | 4 |
| 7 | TGME49_254950.P2082 | RNA cap guanine-N2 methyltransferase | 10 | 10 | 8 | 7 | 5 | 4 | 4 | 4 |
| 7 | TGME49_255170.P2084 | hypothetical protein | 10 | 10 | 9 | 6 | 3 | 3 | 3 | 4 |
| 11 | TGME49_255180.P2085 | ubiquitin carboxyl-terminal hydrolase | 10 | 10 | 8 | 7 | 3 | 6 | 4 | 4 |
| 7 | TGME49_255450.P2102 | inner membrane complex protein IMC31 | 10 | 9 | 8 | 7 | 4 | 4 | 4 | 4 |
| 7 | TGME49_255460.P2103 | EGF family domain-containing protein | 10 | 10 | 10 | 9 | 3 | 3 | 3 | 4 |
| 11 | TGME49_256000-t26_1-p1 | endoplasmic reticulum retention receptor | 7 | 7 | 7 | 7 | 3 | 5 | 7 | 4 |
| 11 | TGME49_256090.P2133 | glycerophosphodiester phosphodiesterase family protein | 9 | 8 | 8 | 8 | 3 | 3 | 6 | 4 |
| 27 | TGME49_257010-t26_1-p1 | toxolysin TLN3 | 10 | 9 | 10 | 10 | 3 | 10 | 9 | 4 |
| 27 | TGME49_257020.P2150 | trafficking protein particle complex subunit 13, putative | 9 | 7 | 8 | 8 | 4 | 7 | 6 | 4 |
| 7 | TGME49_257100.P2156 | hypothetical protein | 9 | 9 | 9 | 7 | 3 | 4 | 3 | 4 |
| 7 | TGME49_257120.P2158 | sugar transporter ST1 | 9 | 9 | 8 | 8 | 3 | 3 | 4 | 4 |
| 11 | TGME49_257300.P2164 | CPH1-interacting protein CIP2 | 10 | 8 | 7 | 7 | 3 | 5 | 5 | 4 |
| 27 | TGME49_257530-t26_1-p1 | apicomplexan amino acid transporter ApiAT5-3 | 9 | 9 | 9 | 8 | 3 | 7 | 7 | 4 |
| 11 | TGME49_257790.P2189 | lysine deacetylase KDAC4 | 10 | 8 | 7 | 7 | 4 | 5 | 5 | 4 |
| 27 | TGME49_257990.P2196 | chaperone protein ClpB1 | 10 | 10 | 10 | 9 | 3 | 10 | 8 | 4 |
| 27 | TGME49_258240.P2214 | chromodomain helicase DNA binding protein CHD1/SWI2/SNF2 | 10 | 10 | 10 | 10 | 3 | 10 | 10 | 4 |
| 11 | TGME49_258360.P2216 | hypothetical protein | 9 | 8 | 8 | 7 | 3 | 5 | 5 | 4 |
| 7 | TGME49_258420.P2220 | hypothetical protein | 10 | 9 | 8 | 7 | 5 | 3 | 3 | 4 |
| 7 | TGME49_258490.P2223 | hypothetical protein | 10 | 9 | 8 | 7 | 5 | 4 | 4 | 4 |

| | | | | | | | | | | |
|---|---|---|---|---|---|---|---|---|---|---|
| 27 | TGME49_258520.P2225 | WD domain, G-beta repeat-containing protein | 10 | 8 | 9 | 9 | 3 | 7 | 6 | 4 |
| 27 | TGME49_258530.P2226 | flagellar associated protein | 9 | 10 | 9 | 8 | 5 | 8 | 7 | 4 |
| 7 | TGME49_258650.P2232 | ferrochelatase | 10 | 9 | 8 | 8 | 5 | 3 | 5 | 4 |
| 7 | TGME49_258860.P2245 | hypothetical protein | 10 | 9 | 9 | 8 | 5 | 4 | 3 | 4 |
| 27 | TGME49_258970.P2252 | nucleoporin NUP155, putative | 10 | 10 | 10 | 10 | 5 | 7 | 6 | 4 |
| 11 | TGME49_258990.P2254 | bromodomain-containing protein BDP3 | 10 | 7 | 8 | 6 | 3 | 5 | 5 | 4 |
| 7 | TGME49_259670.P2286 | von Willebrand factor type A domain-containing protein | 10 | 8 | 8 | 8 | 3 | 3 | 4 | 4 |
| 11 | TGME49_259710-t26_1-p1 | EF hand domain-containing protein | 8 | 7 | 7 | 6 | 3 | 4 | 5 | 4 |
| 7 | TGME49_259880.P2294 | hypothetical protein | 10 | 10 | 9 | 7 | 4 | 3 | 3 | 4 |
| 7 | TGME49_259920.P2296 | GTP-binding protein, putative | 10 | 10 | 9 | 9 | 3 | 4 | 4 | 4 |
| 7 | TGME49_259940-t26_1-p1 | hypothetical protein | 10 | 9 | 8 | 8 | 3 | 3 | 3 | 4 |
| 27 | TGME49_259990-t26_1-p1 | SAC3/GANP family protein | 10 | 10 | 9 | 9 | 4 | 6 | 6 | 4 |
| 11 | TGME49_260140.P2305 | phosphomannose isomerase type I protein | 10 | 9 | 9 | 7 | 4 | 4 | 5 | 4 |
| 7 | TGME49_260310-t26_1-p1 | ATP-binding cassette transporter ABC.B1 | 10 | 10 | 10 | 10 | 3 | 3 | 3 | 4 |
| 27 | TGME49_260320-t26_1-p1 | Noc2p family protein | 10 | 10 | 9 | 8 | 3 | 6 | 6 | 4 |
| 7 | TGME49_260460.P2323 | oxidoreductase, putative | 9 | 8 | 8 | 8 | 4 | 4 | 4 | 4 |
| 27 | TGME49_260600.P2332 | mRNA-binding protein PUF1 | 10 | 10 | 10 | 8 | 4 | 6 | 6 | 4 |
| 11 | TGME49_261070.P2352 | apicoplast triosephosphate translocator APT1 | 8 | 8 | 8 | 8 | 3 | 3 | 7 | 4 |
| 27 | TGME49_261260.P2359 | histone lysine demethylase JMJD6a | 10 | 9 | 9 | 8 | 5 | 7 | 8 | 4 |
| 7 | TGME49_261610.P2373 | crossover junction endonuclease MUS81, putative | 10 | 9 | 8 | 7 | 4 | 4 | 3 | 4 |
| 7 | TGME49_261660.P2374 | Rab3 GTPase-activating protein catalytic subunit, putative | 10 | 10 | 8 | 7 | 4 | 4 | 4 | 4 |
| 7 | TGME49_261750.P2381 | rhoptry neck protein RON10 | 10 | 8 | 9 | 8 | 5 | 3 | 5 | 4 |
| 27 | TGME49_262120.P2394 | IQ calmodulin-binding motif domain-containing protein | 10 | 9 | 8 | 7 | 4 | 7 | 6 | 4 |
| 27 | TGME49_262140.P2396 | chain a, pp2a-specific methylesterase apo form (pme), putative | 9 | 8 | 9 | 8 | 3 | 7 | 7 | 4 |
| 11 | TGME49_262800.P2428 | leucine rich repeat-containing protein | 7 | 7 | 8 | 7 | 3 | 5 | 5 | 4 |
| 7 | TGME49_262900-t26_1-p1 | hypothetical protein | 10 | 10 | 9 | 9 | 5 | 4 | 4 | 4 |
| 11 | TGME49_262970-t26_1-p1 | zinc finger (CCCH type) motif-containing protein | 10 | 7 | 8 | 6 | 3 | 4 | 5 | 4 |
| 11 | TGME49_263110.P2446 | hypothetical protein | 8 | 7 | 7 | 7 | 3 | 5 | 6 | 4 |
| 7 | TGME49_263180.P2451 | myosin D | 10 | 10 | 10 | 8 | 3 | 3 | 3 | 4 |
| 7 | TGME49_263270-t26_1-p1 | glycerophosphodiester phosphodiesterase family protein | 10 | 8 | 9 | 8 | 4 | 6 | 3 | 4 |
| 11 | TGME49_263360.P2464 | WD domain, G-beta repeat-containing protein | 10 | 9 | 8 | 8 | 3 | 7 | 3 | 4 |
| 27 | TGME49_263670.P2486 | ribosomal RNA large subunit methyltransferase FTSJ1, putative | 8 | 8 | 7 | 7 | 3 | 7 | 7 | 4 |
| 7 | TGME49_263840.P2496 | hypothetical protein | 10 | 9 | 9 | 7 | 5 | 4 | 3 | 4 |
| 27 | TGME49_264030-t26_1-p1 | aminotransferase, putative | 10 | 9 | 9 | 8 | 3 | 8 | 8 | 4 |
| 7 | TGME49_264160.P2508 | DEAD/DEAH box helicase domain-containing protein | 10 | 10 | 9 | 8 | 3 | 5 | 3 | 4 |
| 7 | TGME49_264220.P2514 | hypothetical protein | 10 | 8 | 8 | 7 | 4 | 3 | 3 | 4 |
| 27 | TGME49_264460.P2515 | DNA-directed RNA polymerase III RPC5 | 10 | 10 | 9 | 9 | 5 | 8 | 7 | 4 |
| 27 | TGME49_264650-t26_1-p1 | phosphoacetylglucosamine mutase | 10 | 10 | 9 | 8 | 3 | 6 | 6 | 4 |
| 7 | TGME49_264660-t26_1-p1 | SAG-related sequence SRS44 | 10 | 10 | 10 | 8 | 3 | 3 | 3 | 4 |

| | | | | | | | | | | |
|---|---|---|---|---|---|---|---|---|---|---|
| 27 | TGME49_264670.P2519 | DNA polymerase family B protein | 10 | 10 | 10 | 9 | 5 | 6 | 7 | 4 |
| 7 | TGME49_264780-t26_1-p1 | UTP-glucose-1-phosphate uridylyltransferase subfamily protein | 10 | 9 | 8 | 8 | 5 | 4 | 4 | 4 |
| 7 | TGME49_264830-t26_1-p1 | hypothetical protein | 10 | 10 | 10 | 9 | 3 | 5 | 4 | 4 |
| 11 | TGME49_265040-t26_1-p1 | EGF-like domain | 8 | 7 | 7 | 7 | 3 | 3 | 6 | 4 |
| 7 | TGME49_265090-t26_1-p1 | hypothetical protein | 10 | 9 | 8 | 7 | 3 | 4 | 4 | 4 |
| 7 | TGME49_265100.P2533 | hypothetical protein | 10 | 8 | 8 | 8 | 3 | 3 | 3 | 4 |
| 11 | TGME49_265190.P2535 | Ulp1 protease family, C-terminal catalytic domain-containing protein | 10 | 7 | 6 | 6 | 4 | 5 | 5 | 4 |
| 11 | TGME49_265320-t26_1-p1 | pre-mRNA-splicing factor CWC2, putative | 9 | 9 | 8 | 8 | 4 | 3 | 6 | 4 |
| 11 | TGME49_265510-t26_1-p1 | hypothetical protein | 8 | 7 | 7 | 7 | 3 | 6 | 6 | 4 |
| 7 | TGME49_265830.P2548 | phosphodiesterase/alkaline phosphatase D family protein | 10 | 8 | 8 | 7 | 4 | 3 | 5 | 4 |
| 27 | TGME49_266435.P2568 | hypothetical protein | 10 | 10 | 10 | 10 | 3 | 7 | 9 | 4 |
| 7 | TGME49_266680.P2577 | DNA-directed RNA polymerase III subunit RPC4, putative | 9 | 9 | 8 | 7 | 3 | 4 | 3 | 4 |
| 7 | TGME49_266710.P2578 | NEK kinase | 8 | 8 | 8 | 8 | 3 | 3 | 3 | 4 |
| 27 | TGME49_266785-t26_1-p1 | zinc finger (CCCH type) motif-containing protein | 10 | 10 | 10 | 9 | 3 | 7 | 7 | 4 |
| 7 | TGME49_266890-t26_1-p1 | hypothetical protein | 10 | 10 | 9 | 8 | 3 | 3 | 3 | 4 |
| 11 | TGME49_266900-t26_1-p1 | cyclin, N-terminal domain-containing protein | 8 | 7 | 7 | 7 | 3 | 6 | 6 | 4 |
| 11 | TGME49_266920.P2586 | 3'5'-cyclic nucleotide phosphodiesterase PDE6 | 10 | 10 | 10 | 10 | 3 | 3 | 7 | 4 |
| 7 | TGME49_266940.P2588 | palmitoyltransferase DHHC16 | 10 | 9 | 8 | 8 | 4 | 3 | 4 | 4 |
| 27 | TGME49_267540.P2605 | AGC kinase | 10 | 8 | 8 | 8 | 3 | 7 | 7 | 4 |
| 7 | TGME49_268000.P2620 | hypothetical protein | 10 | 10 | 8 | 7 | 5 | 4 | 4 | 4 |
| 7 | TGME49_268780.P2647 | hypothetical protein | 10 | 10 | 8 | 7 | 4 | 5 | 4 | 4 |
| 27 | TGME49_268850.P2653 | enolase 2 | 9 | 9 | 9 | 9 | 3 | 9 | 9 | 4 |
| 27 | TGME49_268900.P2657 | dense granule protein GRA10 | 10 | 10 | 9 | 6 | 4 | 7 | 6 | 4 |
| 11 | TGME49_269240.P2673 | actin-like family protein | 9 | 9 | 8 | 8 | 3 | 5 | 5 | 4 |
| 7 | TGME49_269290.P2677 | nucleoporin NUP503 | 10 | 10 | 10 | 9 | 5 | 4 | 4 | 4 |
| 7 | TGME49_269300-t26_1-p1 | lipase | 10 | 10 | 9 | 9 | 4 | 5 | 4 | 4 |
| 11 | TGME49_269310.P2678 | zinc finger (CCCH type) motif-containing protein | 10 | 10 | 9 | 7 | 3 | 5 | 5 | 4 |
| 7 | TGME49_269480.P2686 | AP-4 complex subunit beta, putative | 9 | 9 | 9 | 8 | 5 | 4 | 3 | 4 |
| 7 | TGME49_270190.P2724 | protein phosphatase PPM18 | 10 | 9 | 9 | 8 | 3 | 4 | 4 | 4 |
| 7 | TGME49_270200.P2725 | transporter, major facilitator family protein | 10 | 9 | 8 | 8 | 3 | 3 | 3 | 4 |
| 11 | TGME49_270620-t26_1-p1 | DEAD/DEAH box helicase domain-containing protein | 10 | 10 | 9 | 9 | 3 | 5 | 5 | 4 |
| 7 | TGME49_270920.P2790 | rhoptry kinase family protein ROP32 | 9 | 9 | 8 | 8 | 3 | 3 | 3 | 4 |
| 11 | TGME49_271810.P2825 | lanp, putative | 8 | 7 | 7 | 6 | 3 | 5 | 5 | 4 |
| 7 | TGME49_271980-t26_1-p1 | SAG-related sequence SRS32 | 10 | 9 | 8 | 8 | 3 | 3 | 3 | 4 |
| 27 | TGME49_272200.P2842 | AGC kinase | 10 | 7 | 9 | 9 | 3 | 6 | 7 | 4 |
| 7 | TGME49_272280.P2848 | protein phosphatase PPM5A | 10 | 8 | 8 | 7 | 4 | 3 | 3 | 4 |
| 7 | TGME49_272370.P2855 | tartrolon E responsive gene | 10 | 10 | 9 | 7 | 5 | 4 | 4 | 4 |
| 27 | TGME49_272550.P2871 | hypothetical protein | 10 | 9 | 10 | 9 | 4 | 8 | 6 | 4 |
| 7 | TGME49_272590.P2873 | hypothetical protein | 10 | 10 | 10 | 7 | 5 | 3 | 3 | 4 |
| 7 | TGME49_272600-t26_1-p1 | K13 complex-associated alpha-ear containing protein KAE | 10 | 10 | 9 | 7 | 5 | 5 | 3 | 4 |

| | | | | | | | | | | |
|---|---|---|---|---|---|---|---|---|---|---|
| 7 | TGME49_272640.P2876 | eukaryotic initiation factor-2B, epsilon subunit, putative | 10 | 9 | 8 | 7 | 3 | 3 | 3 | 4 |
| 7 | TGME49_272650.P2877 | 3'5'-cyclic nucleotide phosphodiesterase PDE10 | 10 | 10 | 9 | 8 | 3 | 3 | 3 | 4 |
| 11 | TGME49_272890.P2886 | RNA recognition motif-containing protein | 9 | 9 | 8 | 6 | 3 | 5 | 5 | 4 |
| 7 | TGME49_273200-t26_1-p1 | hypothetical protein | 10 | 9 | 8 | 7 | 3 | 6 | 3 | 4 |
| 7 | TGME49_273550.P2908 | GDP-fucose protein O-fucosyltransferase 2 | 10 | 9 | 8 | 8 | 5 | 4 | 4 | 4 |
| 7 | TGME49_274010.P2932 | hypothetical protein | 10 | 10 | 8 | 7 | 4 | 4 | 4 | 4 |
| 11 | TGME49_274120.P2940 | hypothetical protein | 10 | 9 | 8 | 7 | 4 | 3 | 6 | 4 |
| 7 | TGME49_276120.P2964 | histone lysine methyltransferase, SET, putative | 10 | 8 | 8 | 7 | 4 | 3 | 4 | 4 |
| 7 | TGME49_276890-t26_1-p1 | hypothetical protein | 10 | 10 | 9 | 8 | 4 | 5 | 4 | 4 |
| 27 | TGME49_277572-t26_1-p1 | Toxoplasma gondii family D protein | 10 | 9 | 9 | 9 | 3 | 6 | 6 | 4 |
| 11 | TGME49_277870-t26_1-p1 | hypothetical protein | 9 | 9 | 8 | 8 | 4 | 6 | 6 | 4 |
| 7 | TGME49_277930.P2990 | WD40 repeat-containing protein | 10 | 9 | 10 | 8 | 3 | 4 | 3 | 4 |
| 7 | TGME49_278030-t26_1-p1 | hypothetical protein | 10 | 10 | 8 | 7 | 3 | 4 | 3 | 4 |
| 27 | TGME49_278060-t26_1-p1 | double-strand break repair protein MRE11, putative | 10 | 10 | 9 | 9 | 5 | 8 | 8 | 4 |
| 11 | TGME49_278230-t26_1-p1 | prenyltransferase GGT-1 | 9 | 8 | 8 | 7 | 4 | 4 | 5 | 4 |
| 11 | TGME49_278240-t26_1-p1 | Zn-finger in Ran binding protein and others domain-containing protein | 10 | 10 | 9 | 6 | 3 | 5 | 5 | 4 |
| 27 | TGME49_278518.P3001 | N-acetylgalactosaminyl transferase | 10 | 10 | 3 | 9 | 3 | 8 | 8 | 4 |
| 11 | TGME49_278670-t26_1-p1 | ubiquitin-conjugating enzyme subfamily protein | 9 | 8 | 7 | 7 | 3 | 6 | 6 | 4 |
| 7 | TGME49_278710.P3010 | cilia- and flagella-associated protein 221, putative | 10 | 8 | 9 | 8 | 4 | 4 | 3 | 4 |
| 7 | TGME49_278740.P3011 | diaminopimelate decarboxylase | 10 | 9 | 9 | 8 | 5 | 3 | 3 | 4 |
| 11 | TGME49_278780-t26_1-p1 | hypothetical protein | 9 | 7 | 6 | 6 | 3 | 5 | 5 | 4 |
| 7 | TGME49_278882-t26_1-p1 | GDA1/CD39 (nucleoside phosphatase) family protein | 10 | 9 | 9 | 9 | 3 | 3 | 3 | 4 |
| 7 | TGME49_280370-t26_1-p1 | hypothetical protein | 10 | 9 | 9 | 8 | 4 | 4 | 3 | 4 |
| 27 | TGME49_280420.P3036 | HEAT repeat-containing protein | 10 | 10 | 10 | 8 | 4 | 8 | 8 | 4 |
| 27 | TGME49_281440.P3060 | hypothetical protein, DUF3712 domain-containing protein | 10 | 3 | 3 | 9 | 3 | 10 | 8 | 4 |
| 11 | TGME49_281480-t26_1-p1 | WD domain, G-beta repeat-containing protein | 9 | 7 | 7 | 7 | 3 | 6 | 5 | 4 |
| 27 | TGME49_281900-t26_1-p1 | SET domain containing lysine methyltransferase KMTox | 10 | 9 | 9 | 7 | 3 | 7 | 7 | 4 |
| 11 | TGME49_283730.P3097 | endomembrane protein 70 subfamily protein | 10 | 10 | 9 | 7 | 3 | 4 | 7 | 4 |
| 7 | TGME49_285160.P3117 | cyclin-dependent kinase family 5 protein | 9 | 8 | 8 | 8 | 3 | 4 | 3 | 4 |
| 27 | TGME49_285230.P3124 | PRP38 family protein | 10 | 9 | 9 | 5 | 3 | 7 | 8 | 4 |
| 7 | TGME49_285800-t26_1-p1 | transporter, putative | 10 | 10 | 9 | 7 | 4 | 3 | 3 | 4 |
| 11 | TGME49_285920.P3140 | glutathione S-transferase 3 | 8 | 8 | 7 | 6 | 3 | 5 | 5 | 4 |
| 11 | TGME49_285940.P3142 | male gamete fusion factor HAP2, putative | 10 | 9 | 9 | 7 | 3 | 6 | 6 | 4 |
| 27 | TGME49_286060-t26_1-p1 | tricarboxylate carrier protein | 9 | 8 | 8 | 8 | 4 | 7 | 6 | 4 |
| 11 | TGME49_286220.P3152 | GDP-L-fucose synthetase | 8 | 6 | 8 | 7 | 4 | 6 | 6 | 4 |
| 27 | TGME49_286470-t26_1-p1 | cAMP-dependent protein kinase subunit PKAc3 | 9 | 9 | 9 | 9 | 3 | 8 | 8 | 4 |
| 11 | TGME49_286500-t26_1-p1 | transporter, putative | 10 | 7 | 9 | 8 | 3 | 3 | 6 | 4 |
| 7 | TGME49_286580.P3157 | inner membrane complex protein IMC17 | 10 | 10 | 10 | 9 | 5 | 3 | 3 | 4 |

| | | | | | | | | | | |
|---|---|---|---|---|---|---|---|---|---|---|
| 11 | TGME49_286600.P3158 | thioredoxin-like associated protein TLAP1 | 9 | 8 | 7 | 7 | 3 | 5 | 5 | 4 |
| 7 | TGME49_287180.P3168 | DNA polymerase V, putative | 10 | 10 | 10 | 8 | 4 | 5 | 3 | 4 |
| 7 | TGME49_287230.P3171 | inorganic anion transporter, sulfate permease (SulP) family protein | 10 | 9 | 9 | 8 | 5 | 3 | 3 | 4 |
| 7 | TGME49_288000-t26_1-p1 | hypothetical protein | 10 | 10 | 10 | 10 | 3 | 3 | 3 | 4 |
| 27 | TGME49_288050.P3177 | intraflagellar transport protein 172, putative | 10 | 10 | 9 | 8 | 5 | 8 | 6 | 4 |
| 11 | TGME49_288290-t26_1-p1 | hypothetical protein | 9 | 8 | 7 | 7 | 4 | 5 | 5 | 4 |
| 11 | TGME49_288330-t26_1-p1 | histone lysine methyltransferase, SET, putative | 10 | 9 | 9 | 7 | 4 | 6 | 5 | 4 |
| 27 | TGME49_288360-t26_1-p1 | tryptophanyl-tRNA synthetase (TrpRS2) | 10 | 9 | 9 | 6 | 3 | 9 | 8 | 4 |
| 27 | TGME49_288450-t26_1-p1 | aldehyde dehydrogenase (NAD) family protein | 10 | 9 | 9 | 9 | 3 | 7 | 6 | 4 |
| 11 | TGME49_288920-t26_1-p1 | ATP-binding cassette G family transporter ABCG96 | 10 | 9 | 9 | 6 | 3 | 3 | 7 | 4 |
| 27 | TGME49_289050.P3197 | FIKK kinase, putative | 10 | 9 | 9 | 9 | 5 | 7 | 7 | 4 |
| 7 | TGME49_289240.P3200 | DnaJ domain-containing protein | 10 | 9 | 9 | 9 | 5 | 3 | 3 | 4 |
| 27 | TGME49_289490.P3203 | protein phosphatase PPM11A, putative | 10 | 9 | 9 | 8 | 5 | 7 | 7 | 4 |
| 7 | TGME49_289540-t26_1-p1 | hypothetical protein | 10 | 10 | 9 | 8 | 3 | 3 | 3 | 4 |
| 11 | TGME49_290180-t26_1-p1 | AP2 domain transcription factor AP2IX-6 | 9 | 8 | 7 | 7 | 3 | 3 | 6 | 4 |
| 7 | TGME49_290330-t26_1-p1 | chloride transporter, chloride channel (ClC) family protein | 10 | 10 | 9 | 8 | 3 | 3 | 3 | 4 |
| 11 | TGME49_290890.P3237 | carbonyl reductase 1, putative | 8 | 8 | 7 | 7 | 3 | 6 | 6 | 4 |
| 7 | TGME49_290990-t26_1-p1 | HEAT repeat-containing protein | 10 | 10 | 10 | 8 | 5 | 4 | 3 | 4 |
| 27 | TGME49_291040-t26_1-p1 | lactate dehydrogenase LDH2 | 8 | 8 | 8 | 7 | 3 | 7 | 7 | 4 |
| 11 | TGME49_291640-t26_1-p1 | aspartate carbamoyltransferase | 9 | 8 | 8 | 6 | 5 | 4 | 6 | 4 |
| 7 | TGME49_291880.P3253 | hypothetical protein | 10 | 9 | 8 | 7 | 4 | 3 | 3 | 4 |
| 7 | TGME49_293420.P3295 | apicomplexan amino acid transporter ApiAT5-5 | 9 | 9 | 8 | 8 | 3 | 3 | 3 | 4 |
| 7 | TGME49_293480.P3302 | MoeA N-terminal region (domain I and II) domain-containing protein | 10 | 10 | 9 | 7 | 3 | 3 | 3 | 4 |
| 7 | TGME49_293500.P3304 | hypothetical protein | 10 | 10 | 10 | 9 | 3 | 4 | 5 | 4 |
| 11 | TGME49_293540.P3306 | hypothetical protein | 10 | 8 | 8 | 7 | 3 | 5 | 5 | 4 |
| 11 | TGME49_293550.P3307 | hypothetical protein | 10 | 9 | 7 | 8 | 4 | 6 | 5 | 4 |
| 11 | TGME49_293560.P3308 | hypothetical protein | 10 | 10 | 8 | 7 | 3 | 6 | 6 | 4 |
| 7 | TGME49_293710.P3320 | Zn-finger in Ran binding protein and others domain-containing protein | 10 | 9 | 9 | 8 | 5 | 3 | 4 | 4 |
| 11 | TGME49_293720.P3321 | HAD-like superfamily | 9 | 8 | 8 | 8 | 3 | 6 | 6 | 4 |
| 11 | TGME49_293890.P3338 | intraflagellar transport 80 family protein | 10 | 9 | 9 | 6 | 3 | 5 | 4 | 4 |
| 11 | TGME49_294010.P3340 | yrdC domain-containing protein | 10 | 9 | 8 | 7 | 4 | 5 | 4 | 4 |
| 7 | TGME49_294320.P3350 | Oxysterol-binding protein | 10 | 10 | 9 | 8 | 5 | 3 | 3 | 4 |
| 27 | TGME49_294380.P3356 | thiocytidine biosynthesis protein TTCA, putative | 10 | 10 | 10 | 8 | 3 | 7 | 6 | 4 |
| 27 | TGME49_294400.P3357 | rhoptry neck protein RON2L1 | 10 | 10 | 10 | 9 | 5 | 8 | 7 | 4 |
| 27 | TGME49_294440-t26_1-p1 | NOL1/NOP2/sun family protein | 10 | 10 | 9 | 8 | 4 | 8 | 7 | 4 |
| 7 | TGME49_294570.P3359 | rhodanese family domain-containing protein | 10 | 8 | 9 | 8 | 3 | 3 | 3 | 4 |
| 27 | TGME49_294610.P3362 | inner membrane complex protein IMC30 | 10 | 10 | 10 | 9 | 3 | 8 | 7 | 4 |
| 27 | TGME49_294620-t26_1-p1 | eukaryotic initiation factor-3, subunit 8, putative | 10 | 10 | 10 | 9 | 3 | 9 | 9 | 4 |
| 11 | TGME49_294690-t26_1-p1 | rhomboid protease ROM5 | 10 | 9 | 8 | 8 | 3 | 6 | 6 | 4 |
| 11 | TGME49_294710.P3365 | RNA recognition motif-containing protein | 10 | 9 | 7 | 7 | 3 | 6 | 5 | 4 |

| | | | | | | | | | | |
|---|---|---|---|---|---|---|---|---|---|---|
| 7 | TGME49_294830-t26_1-p1 | methyltransferase domain-containing protein | 10 | 9 | 8 | 7 | 3 | 4 | 4 | 4 |
| 7 | TGME49_295090.P3380 | oligosaccharyl transferase complex subunit, putative | 10 | 9 | 8 | 7 | 4 | 4 | 3 | 4 |
| 7 | TGME49_295590-t26_1-p1 | hypothetical protein | 10 | 9 | 9 | 7 | 4 | 3 | 3 | 4 |
| 7 | TGME49_295640-t26_1-p1 | peptidase family M13 protein | 10 | 10 | 10 | 10 | 3 | 3 | 3 | 4 |
| 27 | TGME49_295680.P3387 | periodic tryptophan protein PWP2, putative | 10 | 10 | 10 | 10 | 3 | 9 | 8 | 4 |
| 27 | TGME49_295690-t26_1-p1 | ubiquinone/menaquinone biosynthesis methyltransferase subfamily protein | 9 | 8 | 8 | 8 | 5 | 7 | 7 | 4 |
| 11 | TGME49_297245-t26_1-p1 | apicoplast monocarboxylate transporter AMT2 | 9 | 9 | 8 | 7 | 4 | 6 | 6 | 4 |
| 7 | TGME49_297330-t26_1-p1 | hypothetical protein | 10 | 10 | 8 | 7 | 5 | 4 | 4 | 4 |
| 7 | TGME49_297360-t26_1-p1 | hypothetical protein | 10 | 9 | 10 | 9 | 3 | 4 | 4 | 4 |
| 27 | TGME49_297745-t26_1-p1 | hypothetical protein | 10 | 10 | 10 | 7 | 5 | 8 | 7 | 4 |
| 27 | TGME49_297960-t26_1-p1 | rhoptry neck protein RON6 | 9 | 9 | 8 | 8 | 5 | 7 | 6 | 4 |
| 27 | TGME49_298010.P3428 | CLU domain-containing protein | 10 | 10 | 10 | 8 | 3 | 9 | 7 | 4 |
| 11 | TGME49_298030-t26_1-p1 | Ubiquinol-cytochrome c chaperone, putative | 10 | 9 | 9 | 8 | 3 | 4 | 5 | 4 |
| 11 | TGME49_298980-t26_1-p1 | RNA pseudouridine synthase superfamily protein | 10 | 9 | 9 | 6 | 4 | 5 | 5 | 4 |
| 11 | TGME49_299050-t26_1-p1 | ribosomal protein RPL17 | 7 | 7 | 7 | 7 | 3 | 6 | 6 | 4 |
| 7 | TGME49_299060-t26_1-p1 | sodium/hydrogen exchanger NHE2 | 10 | 10 | 10 | 9 | 3 | 3 | 3 | 4 |
| 7 | TGME49_299820.P3446 | hypothetical protein | 10 | 10 | 7 | 7 | 3 | 3 | 4 | 4 |
| 7 | TGME49_300220-t26_1-p1 | hypothetical protein | 10 | 9 | 10 | 9 | 3 | 6 | 3 | 4 |
| 7 | TGME49_300360.P3466 | ADP/ATP translocase | 8 | 8 | 8 | 8 | 3 | 3 | 3 | 4 |
| 7 | TGME49_301210.P3468 | NAD(P) transhydrogenase subunit beta, putative | 10 | 10 | 10 | 10 | 3 | 3 | 3 | 4 |
| 7 | TGME49_304520-t26_1-p1 | hypothetical protein | 10 | 9 | 8 | 7 | 4 | 3 | 3 | 4 |
| 11 | TGME49_305030-t26_1-p1 | kinase, pfkB family protein | 9 | 8 | 7 | 7 | 3 | 4 | 5 | 4 |
| 7 | TGME49_305150.P3497 | tetratricopeptide repeat-containing protein | 10 | 10 | 10 | 9 | 3 | 4 | 3 | 4 |
| 27 | TGME49_305260-t26_1-p1 | hydrolase, putative | 9 | 9 | 9 | 8 | 5 | 7 | 7 | 4 |
| 11 | TGME49_305790-t26_1-p1 | protein-tyrosine phosphatase, putative | 10 | 9 | 9 | 7 | 3 | 4 | 5 | 4 |
| 11 | TGME49_305950-t26_1-p1 | tetratricopeptide repeat-containing protein | 10 | 6 | 8 | 7 | 3 | 5 | 4 | 4 |
| 11 | TGME49_306040.P3515 | CHY zinc finger protein | 10 | 8 | 9 | 8 | 4 | 5 | 5 | 4 |
| 11 | TGME49_306210.P3517 | RNA polymerase II accessory factor CDC73 | 10 | 9 | 9 | 6 | 4 | 5 | 5 | 4 |
| 7 | TGME49_306500-t26_1-p1 | hypothetical protein | 10 | 10 | 10 | 8 | 4 | 3 | 3 | 4 |
| 11 | TGME49_306900-t26_1-p1 | NOL1/NOP2/sun family protein | 10 | 9 | 8 | 7 | 3 | 6 | 6 | 4 |
| 7 | TGME49_307860-t26_1-p1 | hypothetical protein | 10 | 10 | 10 | 8 | 5 | 4 | 4 | 4 |
| 27 | TGME49_307980-t26_1-p1 | GTP-binding protein lepA, putative | 10 | 10 | 9 | 9 | 5 | 8 | 8 | 4 |
| 11 | TGME49_308960.P3551 | ATPase, AAA family protein | 10 | 8 | 8 | 6 | 3 | 7 | 6 | 4 |
| 11 | TGME49_309020.P3556 | tRNA 2-thiolation protein NCS6, putative | 9 | 9 | 8 | 8 | 5 | 4 | 6 | 4 |
| 7 | TGME49_309265.P3566 | oxidoreductase, short chain dehydrogenase/reductase family protein | 9 | 9 | 8 | 8 | 5 | 3 | 3 | 4 |
| 11 | TGME49_309850-t26_1-p1 | insulin-degrading enzyme | 9 | 9 | 8 | 8 | 3 | 6 | 6 | 4 |
| 7 | TGME49_309890-t26_1-p1 | alpha/beta hydrolase, putative | 10 | 10 | 9 | 7 | 3 | 3 | 4 | 4 |
| 7 | TGME49_309930.P3585 | dense granule protein GRA56 | 10 | 9 | 9 | 8 | 3 | 3 | 3 | 4 |
| 7 | TGME49_310080-t26_1-p1 | acyl-CoA synthetase ACS3, putative | 10 | 10 | 10 | 7 | 3 | 3 | 3 | 4 |
| 27 | TGME49_310220.P3602 | basal complex component BCC10 | 10 | 10 | 9 | 9 | 5 | 6 | 6 | 4 |
| 27 | TGME49_310270-t26_1-p1 | regulator of MON1-CCZ1 complex, putative | 10 | 10 | 10 | 9 | 5 | 6 | 6 | 4 |
| 7 | TGME49_310410-t26_1-p1 | WD domain, G-beta repeat-containing protein | 10 | 10 | 10 | 8 | 4 | 4 | 4 | 4 |

| | | | | | | | | | | |
|---|---|---|---|---|---|---|---|---|---|---|
| 11 | TGME49_310480-t26_1-p1 | flagellar/basal body protein, PACRG family protein | 8 | 7 | 6 | 7 | 3 | 5 | 4 | 4 |
| 11 | TGME49_310520.P3613 | 3'5'-cyclic nucleotide phosphodiesterase PDE12 | 10 | 10 | 8 | 8 | 3 | 6 | 6 | 4 |
| 11 | TGME49_310660-t26_1-p1 | Dullard family phosphatase domain-containing protein | 10 | 8 | 8 | 7 | 3 | 5 | 4 | 4 |
| 27 | TGME49_310710-t26_1-p1 | small ribosomal subunit Rsm22 protein | 10 | 9 | 9 | 9 | 3 | 7 | 8 | 4 |
| 11 | TGME49_310930.P3628 | F-box protein FBXO1 | 10 | 9 | 9 | 8 | 3 | 5 | 6 | 4 |
| 7 | TGME49_310950-t26_1-p1 | AP2 domain transcription factor AP2XI-3 | 10 | 9 | 9 | 7 | 5 | 5 | 3 | 4 |
| 7 | TGME49_310970-t26_1-p1 | hypothetical protein | 10 | 9 | 8 | 7 | 5 | 4 | 4 | 4 |
| 27 | TGME49_311070-t26_1-p1 | RNA recognition motif-containing protein | 10 | 9 | 8 | 8 | 4 | 7 | 7 | 4 |
| 11 | TGME49_311160.P3636 | PWI domain-containing protein | 8 | 7 | 7 | 7 | 3 | 3 | 6 | 4 |
| 7 | TGME49_311210-t26_1-p1 | hypothetical protein | 10 | 9 | 9 | 8 | 5 | 4 | 3 | 4 |
| 11 | TGME49_311370.P3645 | methylmalonate-semialdehyde dehydrogenase [acylating], putative | 10 | 8 | 9 | 9 | 3 | 8 | 3 | 4 |
| 11 | TGME49_311380.P3647 | hypothetical protein | 10 | 10 | 9 | 6 | 3 | 5 | 5 | 4 |
| 27 | TGME49_311400-t26_1-p1 | protein transport protein SEC31, putative | 10 | 10 | 10 | 9 | 5 | 9 | 6 | 4 |
| 27 | TGME49_311470.P3650 | rhoptry neck protein RON5 | 10 | 10 | 10 | 10 | 4 | 7 | 6 | 4 |
| 11 | TGME49_311480.P3651 | apical cap protein AC1 | 9 | 9 | 8 | 8 | 3 | 6 | 6 | 4 |
| 7 | TGME49_311710-t26_1-p1 | hypothetical protein | 10 | 9 | 9 | 8 | 3 | 3 | 3 | 4 |
| 11 | TGME49_312150-t26_1-p1 | hypothetical protein | 9 | 8 | 8 | 7 | 3 | 5 | 5 | 4 |
| 11 | TGME49_312300-t26_1-p1 | TBC domain-containing protein TBC16 | 10 | 10 | 7 | 6 | 3 | 6 | 6 | 4 |
| 27 | TGME49_312420.P3670 | dense granule protein GRA38 | 10 | 9 | 9 | 8 | 4 | 6 | 6 | 4 |
| 11 | TGME49_312530.P3674 | splicing factor, CC1 family protein | 10 | 9 | 9 | 8 | 3 | 6 | 5 | 4 |
| 11 | TGME49_312580-t26_1-p1 | hypothetical protein | 10 | 8 | 8 | 7 | 4 | 5 | 4 | 4 |
| 7 | TGME49_313030.P3688 | hypothetical protein | 10 | 10 | 8 | 7 | 5 | 3 | 5 | 4 |
| 27 | TGME49_313310.P3698 | DnaJ domain-containing protein | 10 | 9 | 9 | 9 | 3 | 7 | 8 | 4 |
| 27 | TGME49_313330.P3700 | rhoptry kinase family protein ROP27 | 10 | 9 | 9 | 8 | 5 | 7 | 6 | 4 |
| 7 | TGME49_313340-t26_1-p1 | hypothetical protein | 10 | 10 | 8 | 8 | 4 | 4 | 4 | 4 |
| 27 | TGME49_313630.P3715 | VPS13 domain-containing protein | 10 | 10 | 9 | 9 | 3 | 7 | 7 | 4 |
| 27 | TGME49_313950.P3728 | GMC oxidoreductase | 10 | 10 | 9 | 7 | 3 | 7 | 7 | 4 |
| 11 | TGME49_313980-t26_1-p1 | ribosome biogenesis protein nep1, putative | 8 | 7 | 7 | 6 | 4 | 6 | 6 | 4 |
| 11 | TGME49_314295.P3736 | ribosomal l25 family protein | 8 | 7 | 8 | 7 | 4 | 6 | 6 | 4 |
| 7 | TGME49_314340.P3739 | Sodium:neurotransmitter symporter family protein | 10 | 9 | 8 | 8 | 3 | 3 | 3 | 4 |
| 7 | TGME49_314530-t26_1-p1 | RPAP1 family, C-terminal protein | 10 | 10 | 10 | 7 | 4 | 4 | 3 | 4 |
| 7 | TGME49_314700.P3746 | hypothetical protein | 10 | 10 | 8 | 7 | 4 | 3 | 3 | 4 |
| 7 | TGME49_314730-t26_1-p1 | ALG6, ALG8 glycosyltransferase family protein | 10 | 9 | 9 | 8 | 3 | 3 | 3 | 4 |
| 11 | TGME49_314850-t26_1-p1 | peptidase, M16 family potein | 10 | 10 | 10 | 7 | 3 | 5 | 5 | 4 |
| 27 | TGME49_315100-t26_1-p1 | hypothetical protein | 10 | 10 | 10 | 9 | 3 | 7 | 6 | 4 |
| 11 | TGME49_315250-t26_1-p1 | GAMM1 protein, putative | 9 | 9 | 8 | 8 | 3 | 5 | 5 | 4 |
| 7 | TGME49_315270-t26_1-p1 | hypothetical protein | 10 | 9 | 8 | 7 | 5 | 4 | 4 | 4 |
| 7 | TGME49_315460.P3770 | hypothetical protein | 10 | 8 | 9 | 7 | 5 | 4 | 3 | 4 |
| 7 | TGME49_315480.P3772 | acyl-CoA dehydrogenase, middle domain-containing protein | 10 | 9 | 10 | 9 | 3 | 3 | 3 | 4 |
| 11 | TGME49_315720.P3786 | Smg-4/UPF3 family protein | 9 | 7 | 7 | 6 | 4 | 5 | 5 | 4 |
| 27 | TGME49_316180-t26_1-p1 | hypothetical protein | 10 | 9 | 9 | 8 | 4 | 8 | 6 | 4 |
| 7 | TGME49_316240-t26_1-p1 | dynactin subunit 4, putative | 10 | 9 | 9 | 8 | 3 | 5 | 4 | 4 |
| 11 | TGME49_316330-t26_1-p1 | superoxide dismutase SOD2 | 8 | 8 | 7 | 7 | 3 | 3 | 6 | 4 |
| 7 | TGME49_316430.P3808 | target of rapamycin (TOR), putative | 10 | 10 | 10 | 10 | 3 | 3 | 4 | 4 |

| | | | | | | | | | | |
|---|---|---|---|---|---|---|---|---|---|---|
| 7 | TGME49_316670-t26_1-p1 | Toxoplasma gondii family D protein | 10 | 10 | 9 | 9 | 3 | 3 | 3 | 4 |
| 7 | TGME49_316750-t26_1-p1 | DEAD/DEAH box helicase domain-containing protein | 10 | 9 | 9 | 7 | 5 | 3 | 3 | 4 |
| 27 | TGME49_318310-t26_1-p1 | transketolase | 10 | 10 | 10 | 10 | 3 | 9 | 9 | 4 |
| 11 | TGME49_318490.P3832 | protein arginine N-methyltransferase SFM1-like, putative | 7 | 7 | 7 | 6 | 3 | 5 | 5 | 4 |
| 27 | TGME49_318590.P3838 | Fe-S cluster assembly factor HCF101 | 10 | 9 | 9 | 8 | 5 | 7 | 7 | 4 |
| 11 | TGME49_318640-t26_1-p1 | hypothetical protein | 10 | 6 | 9 | 7 | 4 | 5 | 4 | 4 |
| 27 | TGME49_318675.P3842 | 3'5'-cyclic nucleotide phosphodiesterase PDE8 | 10 | 10 | 10 | 8 | 3 | 8 | 6 | 4 |
| 7 | TGME49_319340.P3846 | dense granule protein GRA52 | 9 | 8 | 8 | 7 | 4 | 4 | 4 | 4 |
| 7 | TGME49_319580-t26_1-p1 | hypothetical protein | 10 | 8 | 9 | 7 | 4 | 4 | 4 | 4 |
| 27 | TGME49_319658-t26_1-p1 | CBF/Mak21 family protein | 10 | 9 | 8 | 8 | 4 | 6 | 7 | 4 |
| 27 | TGME49_319710.P3856 | kinesin motor domain-containing protein | 10 | 10 | 10 | 8 | 4 | 6 | 6 | 4 |
| 7 | TGME49_319720.P3857 | hypothetical protein | 10 | 9 | 9 | 7 | 5 | 4 | 3 | 4 |
| 11 | TGME49_320080-t26_1-p1 | KH domain-containing protein | 10 | 10 | 8 | 7 | 3 | 5 | 5 | 4 |
| 7 | TGME49_320270.P3872 | nucleoside transporter, putative | 10 | 9 | 8 | 7 | 3 | 3 | 3 | 4 |
| 11 | TGME49_320290.P3874 | hypothetical protein | 9 | 9 | 8 | 8 | 3 | 6 | 6 | 4 |
| 27 | TGME49_320460.P3876 | ABC transporter transmembrane region domain-containing protein | 10 | 10 | 10 | 9 | 3 | 9 | 9 | 4 |
| 7 | TGME49_321420-t26_1-p1 | kelch repeat-containing protein | 9 | 9 | 8 | 7 | 3 | 4 | 3 | 4 |
| 27 | TGME49_321450-t26_1-p1 | Myb family DNA-binding domain-containing protein | 10 | 10 | 9 | 8 | 4 | 8 | 7 | 4 |
| 11 | TGME49_321630-t26_1-p1 | RNA recognition motif-containing protein | 9 | 5 | 8 | 6 | 3 | 6 | 6 | 4 |
| 7 | TGME49_500044.P345 | 3'5'-cyclic nucleotide phosphodiesterase PDE16 | 10 | 10 | 10 | 10 | 3 | 3 | 3 | 4 |
| 27 | TGME49_500045.P346 | endo-1,3(4)-beta-glucanase, putative | 9 | 10 | 10 | 10 | 3 | 10 | 9 | 4 |
| 11 | TGME49_500046.P347 | hypothetical protein, conserved | 10 | 6 | 9 | 9 | 3 | 3 | 6 | 4 |
| 7 | TGME49_500047.P348 | Dpy-30 motif protein | 10 | 9 | 9 | 9 | 3 | 5 | 4 | 4 |
| 7 | TGME49_500058.P359 | ferlin 3, putative | 10 | 10 | 10 | 10 | 5 | 3 | 3 | 4 |
| 7 | TGME49_500079.P380 | cilia- and flagella-associated protein 69, putative | 10 | 8 | 9 | 7 | 4 | 4 | 4 | 4 |
| 27 | TGME49_500113.P415 | WD repeat-containing protein | 10 | 7 | 10 | 9 | 3 | 8 | 6 | 4 |
| 27 | TGME49_500142.P444 | serine-protein kinase ATM, putative | 10 | 10 | 10 | 9 | 3 | 8 | 5 | 4 |
| 11 | TGME49_500143.P445 | hypothetical protein, conserved | 9 | 7 | 9 | 6 | 3 | 5 | 4 | 4 |
| 27 | TGME49_500148.P152 | GCC2 and GCC3 domain-containing protein | 10 | 10 | 10 | 8 | 5 | 10 | 9 | 4 |
| 11 | TGME49_500153.P157 | WD repeat-containing protein | 10 | 8 | 9 | 9 | 3 | 6 | 5 | 4 |
| 11 | TGME49_500179.P183 | FYVE zinc finger domain-containing protein | 10 | 9 | 8 | 6 | 4 | 5 | 5 | 4 |
| 7 | TGME49_500182.P186 | regulator of chromosome condensation, putative | 10 | 9 | 8 | 7 | 3 | 4 | 4 | 4 |
| 27 | TGME49_500185.P189 | WD domain, G-beta repeat-containing protein | 10 | 9 | 10 | 9 | 5 | 8 | 7 | 4 |
| 27 | TGME49_500189.P193 | E3 ubiquitin-protein ligase, putative | 10 | 10 | 10 | 8 | 3 | 7 | 7 | 4 |
| 11 | TGME49_500223.P228 | oocyst wall protein | 10 | 9 | 9 | 7 | 3 | 5 | 5 | 4 |
| 27 | TGME49_500227.P233 | dynein heavy chain, putative | 10 | 10 | 10 | 10 | 3 | 10 | 10 | 4 |
| 27 | TGME49_500245.P251 | rhoptry discharge factor RDF2 | 10 | 10 | 9 | 9 | 3 | 8 | 7 | 4 |
| 11 | TGME49_500254.P43 | poly [ADP-ribose] polymerase, putative | 10 | 8 | 7 | 7 | 3 | 5 | 5 | 4 |
| 27 | TGME49_500261.P50 | alanine--glyoxylate aminotransferase | 9 | 9 | 8 | 8 | 3 | 7 | 7 | 4 |
| 7 | TGME49_500267.P56 | intraflagellar transport protein IFT140, putative | 10 | 9 | 10 | 8 | 4 | 5 | 4 | 4 |

| | | | | | | | | | | |
|---|---|---|---|---|---|---|---|---|---|---|
| 7 | TGME49_500272.P61 | P-type ATPase, putative | 10 | 10 | 10 | 10 | 3 | 3 | 3 | 4 |
| 27 | TGME49_500291.P80 | WD repeat-containing protein 19, putative | 10 | 9 | 10 | 10 | 5 | 7 | 5 | 4 |
| 11 | TGME49_500302.P91 | Porphobilinogen Synthase (PBGS) / Aminolevulinic Acid Dehydratase (ALAD) + Pitrilysin (apicoplast stromal processing peptidase) | 10 | 9 | 10 | 8 | 3 | 3 | 7 | 4 |
| 27 | TGME49_500306.P96 | adaptor complexes medium subunit family protein | 10 | 9 | 9 | 7 | 3 | 7 | 6 | 4 |
| 7 | TGME49_500333.P123 | tetratricopeptide repeat-containing protein | 10 | 9 | 10 | 9 | 3 | 5 | 4 | 4 |
| 7 | TGME49_500343.P133 | thrombospondin type 1 domain-containing protein | 10 | 10 | 10 | 9 | 4 | 3 | 4 | 4 |
| 27 | TGME49_500358.P148 | WD domain, G-beta repeat-containing protein | 9 | 9 | 9 | 8 | 5 | 8 | 6 | 4 |
| 11 | TGME49_500361.P260 | aminopeptidase, putative | 9 | 9 | 9 | 7 | 3 | 5 | 4 | 4 |
| 7 | TGME49_500363.P262 | regulator of chromosome condensation, putative | 9 | 10 | 8 | 7 | 5 | 3 | 3 | 4 |
| 7 | TGME49_500376.P275 | rhoptry kinase family protein | 9 | 9 | 8 | 7 | 4 | 3 | 3 | 4 |
| 27 | TGME49_500383.P282 | cilia- and flagella-associated protein 61, putative | 10 | 9 | 10 | 9 | 4 | 9 | 6 | 4 |
| 11 | TGME49_500385.P284 | cilia- and flagella-associated protein 58, putative | 10 | 10 | 8 | 8 | 3 | 6 | 6 | 4 |
| 27 | TGME49_500389.P288 | ATPase, AAA family protein | 10 | 8 | 9 | 7 | 3 | 7 | 7 | 4 |
| 27 | TGME49_500398.P297 | rhoptry neck protein RON2L2 (sporoRON2) | 7 | 9 | 10 | 10 | 3 | 8 | 3 | 4 |
| 7 | TGME49_500401.P300 | PAN domain-containing protein | 10 | 9 | 9 | 8 | 3 | 4 | 4 | 4 |
| 11 | TGME49_500411.P310 | hypothetical protein, conserved | 9 | 5 | 8 | 8 | 3 | 5 | 5 | 4 |
| 7 | TGME49_500438.P337 | SPAG17 domain-containing protein | 10 | 9 | 9 | 6 | 3 | 5 | 3 | 4 |

| Kmean_cluster | GeneID (all protein TG) | Product Description | Decile_bitHH | Decile_bitNcan | Decile_bitBB | Decilebit_CS | Decilebit_SN3 | Decile bit_Ef | Decile bit_CC | Meta_cluster |
|---|---|---|---|---|---|---|---|---|---|---|
| 13 | TGME49_200340.P454 | hypothetical protein | 8 | 7 | 7 | 7 | 3 | 3 | 3 | 5 |
| 8 | TGME49_200385.P457 | bradyzoite-formation deficient protein BFD1 | 10 | 10 | 7 | 3 | 5 | 4 | 3 | 5 |
| 13 | TGME49_200470-t26_1-p1 | hypothetical protein | 9 | 6 | 7 | 6 | 3 | 3 | 3 | 5 |
| 14 | TGME49_201120-t26_1-p1 | ELMO/CED-12 family protein | 10 | 10 | 3 | 3 | 7 | 5 | 4 | 5 |
| 14 | TGME49_201150-t26_1-p1 | heavy metal translocating P-type ATPase subfamily protein | 9 | 10 | 8 | 3 | 7 | 5 | 5 | 5 |
| 13 | TGME49_201230-t26_1-p1 | kinesin motor domain-containing protein | 10 | 9 | 8 | 5 | 3 | 3 | 3 | 5 |
| 8 | TGME49_201240-t26_1-p1 | hypothetical protein | 10 | 8 | 6 | 3 | 3 | 3 | 3 | 5 |
| 13 | TGME49_201260.P468 | sugar transporter ST3 | 10 | 8 | 8 | 5 | 4 | 3 | 3 | 5 |
| 15 | TGME49_201670-t26_1-p1 | DnaJ domain-containing protein | 10 | 9 | 7 | 5 | 5 | 3 | 4 | 5 |
| 8 | TGME49_201690-t26_1-p1 | BT1 family protein | 10 | 9 | 9 | 3 | 3 | 3 | 3 | 5 |
| 14 | TGME49_201700-t26_1-p1 | protein transport protein SEC13 | 9 | 8 | 8 | 3 | 6 | 6 | 5 | 5 |
| 14 | TGME49_201710-t26_1-p1 | WD domain, G-beta repeat-containing protein | 9 | 9 | 8 | 5 | 7 | 4 | 6 | 5 |
| 13 | TGME49_201900-t26_1-p1 | hypothetical protein | 10 | 8 | 7 | 7 | 4 | 4 | 4 | 5 |
| 13 | TGME49_202080-t26_1-p1 | hypothetical protein | 10 | 8 | 9 | 6 | 4 | 3 | 3 | 5 |
| 8 | TGME49_202140.P483 | enoyl-CoA hydratase/isomerase family protein | 10 | 9 | 8 | 3 | 5 | 3 | 3 | 5 |
| 8 | TGME49_202170.P484 | hypothetical protein | 10 | 8 | 8 | 4 | 4 | 4 | 4 | 5 |
| 13 | TGME49_202200-t26_1-p1 | hypothetical protein | 10 | 7 | 7 | 7 | 3 | 4 | 4 | 5 |
| 8 | TGME49_202220.P486 | IMC suture-associated protein ISAP1 | 10 | 9 | 7 | 4 | 4 | 3 | 3 | 5 |
| 15 | TGME49_202250-t26_1-p1 | hypothetical protein | 10 | 9 | 9 | 3 | 5 | 4 | 3 | 5 |
| 19 | TGME49_202360-t26_1-p1 | methylase, putative | 9 | 8 | 8 | 3 | 4 | 5 | 4 | 5 |
| 15 | TGME49_202490-t26_1-p1 | AP2 domain transcription factor AP2VIIa-7 | 10 | 9 | 9 | 5 | 5 | 5 | 4 | 5 |
| 19 | TGME49_202510-t26_1-p1 | glideosome-associated protein with multiple-membrane spans GAPM1B | 8 | 8 | 3 | 3 | 3 | 6 | 7 | 5 |
| 13 | TGME49_202550.P494 | basal complex component BCC6 | 10 | 9 | 7 | 5 | 3 | 4 | 4 | 5 |
| 8 | TGME49_202640.P498 | tRNA pseudouridine synthase 1 | 10 | 9 | 8 | 4 | 3 | 3 | 3 | 5 |
| 14 | TGME49_202740-t26_1-p1 | pseudouridine synthase, putative | 10 | 8 | 6 | 4 | 5 | 5 | 4 | 5 |
| 14 | TGME49_202750-t26_1-p1 | 3' exoribonuclease family, domain 1 domain-containing protein | 8 | 8 | 8 | 4 | 6 | 5 | 5 | 5 |
| 13 | TGME49_202780-t26_1-p1 | rhoptry kinase family protein ROP25 | 10 | 9 | 8 | 6 | 3 | 3 | 3 | 5 |
| 19 | TGME49_202790-t26_1-p1 | dihydrouridine synthase (dus) protein | 9 | 9 | 8 | 4 | 5 | 6 | 5 | 5 |
| 14 | TGME49_202820-t26_1-p1 | ubiquitin-conjugating enzyme subfamily protein | 7 | 7 | 7 | 3 | 6 | 6 | 6 | 5 |
| 13 | TGME49_202900-t26_1-p1 | zinc finger (CCCH type) motif-containing protein | 10 | 10 | 7 | 5 | 3 | 4 | 4 | 5 |
| 19 | TGME49_202970-t26_1-p1 | hypothetical protein | 10 | 10 | 9 | 4 | 3 | 6 | 6 | 5 |
| 13 | TGME49_203120-t26_1-p1 | hypothetical protein | 9 | 8 | 6 | 6 | 4 | 4 | 3 | 5 |
| 13 | TGME49_203180.P514 | leucine rich repeat-containing protein | 10 | 9 | 8 | 5 | 3 | 3 | 3 | 5 |
| 15 | TGME49_203200.P515 | hypothetical protein | 10 | 9 | 9 | 6 | 5 | 3 | 3 | 5 |
| 13 | TGME49_203260-t26_1-p1 | cell cycle checkpoint protein RAD17, putative | 9 | 8 | 7 | 5 | 4 | 4 | 3 | 5 |
| 13 | TGME49_203420-t26_1-p1 | 4'-phosphopantetheinyl transferase domain-containing protein | 10 | 7 | 7 | 5 | 4 | 4 | 4 | 5 |
| 13 | TGME49_203440.P522 | anaphase-promoting complex subunit APC10 | 9 | 7 | 7 | 6 | 3 | 5 | 3 | 5 |
| 15 | TGME49_203480-t26_1-p1 | hypothetical protein | 10 | 9 | 8 | 4 | 4 | 4 | 4 | 5 |
| 15 | TGME49_203490-t26_1-p1 | GNL3L/Grn1 GTPase | 10 | 9 | 9 | 6 | 5 | 3 | 5 | 5 |
| 19 | TGME49_203540.P526 | RNA binding protein, putative | 10 | 10 | 9 | 5 | 3 | 6 | 5 | 5 |
| 15 | TGME49_203710-t26_1-p1 | AP2 domain transcription factor AP2VIIa-4 | 10 | 10 | 10 | 6 | 4 | 4 | 4 | 5 |
| 15 | TGME49_203760-t26_1-p1 | hypothetical protein | 10 | 10 | 8 | 5 | 4 | 4 | 4 | 5 |
| 8 | TGME49_203780-t26_1-p1 | hypothetical protein | 10 | 10 | 8 | 4 | 4 | 3 | 3 | 5 |
| 15 | TGME49_203960-t26_1-p1 | hypothetical protein | 10 | 9 | 8 | 4 | 4 | 4 | 4 | 5 |
| 15 | TGME49_203980.P539 | hypothetical protein | 10 | 10 | 10 | 3 | 7 | 5 | 3 | 5 |

| | | | | | | | | | | |
|---|---|---|---|---|---|---|---|---|---|---|
| 8 | TGME49_204140-t26_1-p1 | PHD-finger domain-containing protein | 10 | 8 | 7 | 4 | 4 | 3 | 4 | 5 |
| 8 | TGME49_204380-t26_1-p1 | hypothetical protein | 10 | 8 | 6 | 4 | 5 | 3 | 4 | 5 |
| 8 | TGME49_204460.P554 | hydroxymethylglutaryl-CoA lyase, putative | 9 | 8 | 8 | 3 | 4 | 3 | 3 | 5 |
| 8 | TGME49_204510.P556 | WD domain, G-beta repeat-containing protein | 10 | 9 | 9 | 3 | 4 | 3 | 3 | 5 |
| 15 | TGME49_204870.P558 | thrombospondin type 1 domain-containing protein | 10 | 9 | 9 | 3 | 5 | 5 | 3 | 5 |
| 13 | TGME49_205000-t26_1-p1 | phosphoglycerate mutase family protein | 10 | 9 | 7 | 7 | 4 | 4 | 4 | 5 |
| 15 | TGME49_205040-t26_1-p1 | PGAP1 family protein | 10 | 10 | 10 | 4 | 4 | 4 | 3 | 5 |
| 8 | TGME49_205050-t26_1-p1 | hypothetical protein | 10 | 9 | 8 | 4 | 3 | 3 | 3 | 5 |
| 14 | TGME49_205150-t26_1-p1 | ACR, YagE family COG1723 domain-containing protein | 10 | 9 | 5 | 5 | 6 | 6 | 5 | 5 |
| 8 | TGME49_205160-t26_1-p1 | zinc finger CCHC domain-containing protein | 10 | 8 | 7 | 3 | 4 | 3 | 3 | 5 |
| 15 | TGME49_205190-t26_1-p1 | hypothetical protein | 10 | 10 | 10 | 5 | 6 | 4 | 4 | 5 |
| 8 | TGME49_205200-t26_1-p1 | hypothetical protein | 10 | 9 | 6 | 3 | 3 | 4 | 4 | 5 |
| 13 | TGME49_205280-t26_1-p1 | hypothetical protein | 9 | 8 | 7 | 5 | 3 | 4 | 4 | 5 |
| 19 | TGME49_205300-t26_1-p1 | hypothetical protein | 10 | 9 | 8 | 4 | 5 | 6 | 5 | 5 |
| 15 | TGME49_205320-t26_1-p1 | hypothetical protein | 9 | 9 | 8 | 5 | 5 | 3 | 3 | 5 |
| 13 | TGME49_205370.P566 | hypothetical protein | 9 | 9 | 6 | 5 | 3 | 3 | 3 | 5 |
| 14 | TGME49_205430.P570 | isovaleryl-CoA dehydrogenase | 8 | 7 | 8 | 3 | 6 | 3 | 3 | 5 |
| 15 | TGME49_205562-t26_1-p1 | hypothetical protein | 10 | 10 | 8 | 5 | 4 | 4 | 3 | 5 |
| 13 | TGME49_205640-t26_1-p1 | hypothetical protein | 10 | 7 | 6 | 5 | 3 | 3 | 3 | 5 |
| 19 | TGME49_205680-t26_1-p1 | hypothetical protein | 8 | 7 | 6 | 3 | 3 | 5 | 5 | 5 |
| 14 | TGME49_205760-t26_1-p1 | membrane protein | 9 | 8 | 8 | 3 | 7 | 7 | 3 | 5 |
| 8 | TGME49_206380.P582 | hypothetical protein | 10 | 10 | 7 | 4 | 3 | 3 | 3 | 5 |
| 19 | TGME49_206415-t26_1-p1 | myosin K | 10 | 10 | 10 | 6 | 3 | 7 | 3 | 5 |
| 15 | TGME49_206630.P596 | hypothetical protein | 10 | 9 | 8 | 6 | 6 | 4 | 4 | 5 |
| 13 | TGME49_206650-t26_1-p1 | zinc finger, c2h2 type domain-containing protein | 10 | 9 | 7 | 5 | 3 | 3 | 3 | 5 |
| 14 | TGME49_207040.P602 | hypothetical protein | 10 | 8 | 8 | 5 | 6 | 5 | 5 | 5 |
| 15 | TGME49_207200-t26_1-p1 | hypothetical protein | 10 | 10 | 8 | 4 | 5 | 4 | 3 | 5 |
| 19 | TGME49_207400.P614 | splicing factor Yju2, putative | 9 | 8 | 8 | 4 | 4 | 4 | 5 | 5 |
| 8 | TGME49_207600.P620 | tubulin/FtsZ family, GTPase domain-containing protein | 10 | 9 | 8 | 3 | 4 | 4 | 4 | 5 |
| 14 | TGME49_207610.P621 | rhoptry kinase family protein ROP36 (incomplete catalytic triad) | 10 | 9 | 9 | 3 | 8 | 4 | 3 | 5 |
| 8 | TGME49_207700-t26_1-p1 | rhoptry kinase family protein ROP22 (incomplete catalytic triad) | 10 | 8 | 9 | 3 | 3 | 3 | 3 | 5 |
| 15 | TGME49_207970.P634 | HEAT repeat-containing protein | 10 | 9 | 9 | 4 | 3 | 4 | 4 | 5 |
| 8 | TGME49_208020.P636 | AP2 domain transcription factor AP2Ib-1 | 10 | 10 | 5 | 3 | 4 | 4 | 4 | 5 |
| 19 | TGME49_208050-t26_1-p1 | ABC transporter, putative | 10 | 9 | 8 | 3 | 3 | 6 | 6 | 5 |
| 13 | TGME49_208360-t26_1-p1 | hypothetical protein | 10 | 9 | 7 | 6 | 4 | 4 | 4 | 5 |
| 19 | TGME49_208490-t26_1-p1 | GTP-binding protein, putative | 10 | 9 | 8 | 4 | 5 | 7 | 4 | 5 |
| 13 | TGME49_208510-t26_1-p1 | hypothetical protein | 10 | 7 | 7 | 5 | 3 | 4 | 5 | 5 |
| 8 | TGME49_208570.P649 | ubiquitin conjugating enzyme E2, putative | 10 | 8 | 7 | 3 | 3 | 3 | 3 | 5 |
| 13 | TGME49_208800-t26_1-p1 | mediator complex subunit MED18 | 8 | 8 | 7 | 6 | 4 | 4 | 4 | 5 |
| 15 | TGME49_209020.P663 | hypothetical protein | 10 | 9 | 9 | 5 | 4 | 4 | 4 | 5 |
| 19 | TGME49_209060-t26_1-p1 | thrombospondin type 1 domain-containing protein | 10 | 9 | 9 | 4 | 4 | 6 | 5 | 5 |
| 14 | TGME49_209100-t26_1-p1 | PUB domain-containing protein | 9 | 9 | 9 | 5 | 7 | 5 | 4 | 5 |
| 13 | TGME49_209108.P668 | mRNA-decapping enzyme subunit 1, putative | 9 | 8 | 6 | 6 | 4 | 4 | 4 | 5 |
| 13 | TGME49_209170-t26_1-p1 | hypothetical protein | 9 | 7 | 7 | 5 | 4 | 3 | 3 | 5 |

| | | | | | | | | | | |
|---|---|---|---|---|---|---|---|---|---|---|
| 8 | TGME49_209180-t26_1-p1 | PAN domain-containing protein | 9 | 8 | 7 | 3 | 5 | 3 | 3 | 5 |
| 15 | TGME49_209200-t26_1-p1 | hypothetical protein | 10 | 9 | 7 | 4 | 5 | 4 | 4 | 5 |
| 15 | TGME49_209210.P672 | KH domain-containing protein | 10 | 10 | 8 | 6 | 5 | 4 | 4 | 5 |
| 15 | TGME49_209250-t26_1-p1 | hypothetical protein | 10 | 9 | 8 | 5 | 4 | 3 | 4 | 5 |
| 8 | TGME49_209540-t26_1-p1 | hypothetical protein | 10 | 8 | 7 | 4 | 4 | 3 | 3 | 5 |
| 13 | TGME49_209650.P689 | telomerase reverse transcriptase, putative | 10 | 9 | 6 | 5 | 4 | 4 | 4 | 5 |
| 14 | TGME49_209690-t26_1-p1 | small nuclear ribonucleoprotein | 8 | 7 | 7 | 3 | 7 | 6 | 6 | 5 |
| 15 | TGME49_209730-t26_1-p1 | hypothetical protein | 10 | 9 | 9 | 4 | 5 | 4 | 4 | 5 |
| 13 | TGME49_209790-t26_1-p1 | radical SAM domain-containing protein | 9 | 8 | 8 | 7 | 3 | 3 | 3 | 5 |
| 13 | TGME49_209920-t26_1-p1 | PAN domain-containing protein | 9 | 10 | 8 | 6 | 3 | 3 | 3 | 5 |
| 8 | TGME49_210270.P707 | hypothetical protein | 10 | 10 | 7 | 3 | 3 | 3 | 3 | 5 |
| 8 | TGME49_210300-t26_1-p1 | zinc finger protein, putative | 10 | 10 | 6 | 3 | 5 | 3 | 4 | 5 |
| 15 | TGME49_210310-t26_1-p1 | hypothetical protein | 10 | 10 | 10 | 5 | 5 | 4 | 4 | 5 |
| 13 | TGME49_210345-t26_1-p1 | hypothetical protein | 10 | 6 | 7 | 5 | 3 | 4 | 3 | 5 |
| 15 | TGME49_210730.P714 | ATP-dependent protease ATPase subunit ClpY, putative | 10 | 9 | 9 | 4 | 8 | 3 | 3 | 5 |
| 15 | TGME49_210820.P720 | transporter, putative | 10 | 9 | 9 | 5 | 3 | 4 | 4 | 5 |
| 14 | TGME49_210980.P722 | alternative splicing type 3 and, putative | 9 | 8 | 8 | 3 | 7 | 3 | 7 | 5 |
| 8 | TGME49_211260-t26_1-p1 | rhoptry kinase family protein ROP26 (incomplete catalytic triad) | 9 | 8 | 6 | 3 | 4 | 3 | 3 | 5 |
| 8 | TGME49_211280.P730 | hypothetical protein | 10 | 9 | 6 | 4 | 3 | 3 | 3 | 5 |
| 15 | TGME49_211310.P731 | regulator of chromosome condensation, putative | 10 | 9 | 9 | 4 | 4 | 4 | 4 | 5 |
| 13 | TGME49_211360.P734 | hypothetical protein | 10 | 9 | 7 | 6 | 3 | 3 | 3 | 5 |
| 15 | TGME49_211390.P736 | ubiquitin-activating enzyme, putative | 9 | 8 | 8 | 5 | 5 | 5 | 4 | 5 |
| 14 | TGME49_211430.P740 | centrosomal protein CEP110, putative | 10 | 9 | 9 | 4 | 7 | 5 | 5 | 5 |
| 15 | TGME49_211440-t26_1-p1 | hypothetical protein | 10 | 10 | 9 | 6 | 5 | 4 | 4 | 5 |
| 13 | TGME49_211450-t26_1-p1 | pseudouridine synthase, putative | 10 | 8 | 8 | 5 | 4 | 3 | 3 | 5 |
| 13 | TGME49_211470.P742 | Fcf2 pre-rRNA processing protein | 9 | 8 | 7 | 5 | 4 | 3 | 4 | 5 |
| 13 | TGME49_211640-t26_1-p1 | NGX6 domain-containing protein | 10 | 10 | 6 | 5 | 3 | 4 | 4 | 5 |
| 8 | TGME49_211720.P748 | AP2 domain transcription factor AP2IV-5 | 10 | 10 | 6 | 3 | 4 | 4 | 4 | 5 |
| 15 | TGME49_211890-t26_1-p1 | RAP domain-containing protein | 10 | 10 | 10 | 3 | 5 | 4 | 3 | 5 |
| 13 | TGME49_212090-t26_1-p1 | hypothetical protein | 9 | 8 | 7 | 6 | 4 | 3 | 3 | 5 |
| 13 | TGME49_212200-t26_1-p1 | hypothetical protein | 9 | 7 | 8 | 7 | 4 | 3 | 3 | 5 |
| 8 | TGME49_212220-t26_1-p1 | zinc finger (CCCH type) motif-containing protein | 10 | 10 | 7 | 3 | 3 | 4 | 4 | 5 |
| 13 | TGME49_212740-t26_1-p1 | aromatic amino acid hydrolase AAH2 | 5 | 8 | 8 | 8 | 3 | 3 | 3 | 5 |
| 15 | TGME49_212780-t26_1-p1 | hypothetical protein | 10 | 10 | 8 | 6 | 4 | 4 | 3 | 5 |
| 15 | TGME49_212800.P767 | kinetochore protein SPC24, putative | 10 | 9 | 7 | 6 | 5 | 4 | 3 | 5 |
| 13 | TGME49_212900-t26_1-p1 | hypothetical protein | 10 | 9 | 6 | 6 | 3 | 3 | 3 | 5 |
| 8 | TGME49_212940.P771 | condensin-2 complex subunit H2, putative | 10 | 9 | 8 | 3 | 3 | 4 | 4 | 5 |
| 15 | TGME49_213010-t26_1-p1 | hypothetical protein | 10 | 9 | 6 | 5 | 5 | 4 | 4 | 5 |
| 19 | TGME49_213300.P787 | hypothetical protein | 10 | 10 | 10 | 5 | 4 | 5 | 5 | 5 |
| 15 | TGME49_213310-t26_1-p1 | hypothetical protein | 10 | 9 | 8 | 6 | 4 | 4 | 4 | 5 |
| 14 | TGME49_213325-t26_1-p1 | TBC domain-containing protein TBC18 | 10 | 7 | 8 | 4 | 6 | 6 | 6 | 5 |
| 8 | TGME49_213388-t26_1-p1 | hypothetical protein | 10 | 8 | 6 | 4 | 4 | 4 | 3 | 5 |
| 15 | TGME49_213392.P790 | surface antigen repeat-containing protein | 10 | 10 | 9 | 6 | 4 | 4 | 4 | 5 |
| 19 | TGME49_213620-t26_1-p1 | ABC1 family protein | 10 | 10 | 8 | 3 | 3 | 6 | 6 | 5 |
| 13 | TGME49_213660-t26_1-p1 | zinc finger (CCCH type) motif-containing protein | 10 | 9 | 7 | 6 | 3 | 3 | 3 | 5 |
| 8 | TGME49_213700-t26_1-p1 | coiled-coil domain-containing protein | 10 | 9 | 7 | 3 | 3 | 4 | 3 | 5 |
| 13 | TGME49_213710.P802 | WD domain, G-beta repeat-containing protein | 10 | 9 | 9 | 5 | 3 | 3 | 3 | 5 |
| 8 | TGME49_213760.P806 | hypothetical protein | 9 | 8 | 6 | 4 | 3 | 3 | 3 | 5 |

| | | | | | | | | | | |
|---|---|---|---|---|---|---|---|---|---|---|
| 19 | TGME49_213790.P807 | CRAL/TRIO domain-containing protein | 10 | 8 | 9 | 5 | 3 | 5 | 4 | 5 |
| 13 | TGME49_213840.P809 | choline transporter-like protein, putative | 9 | 8 | 8 | 7 | 3 | 3 | 3 | 5 |
| 8 | TGME49_213950.P814 | hypothetical protein | 10 | 8 | 8 | 3 | 4 | 4 | 4 | 5 |
| 14 | TGME49_214110-t26_1-p1 | protein yipf5, putative | 9 | 8 | 7 | 3 | 6 | 5 | 5 | 5 |
| 14 | TGME49_214280-t26_1-p1 | phosphoadenosine phosphosulfate reductase family protein | 9 | 8 | 7 | 4 | 6 | 6 | 5 | 5 |
| 13 | TGME49_214310-t26_1-p1 | hypothetical protein | 9 | 9 | 7 | 6 | 4 | 3 | 3 | 5 |
| 15 | TGME49_214370-t26_1-p1 | hypothetical protein | 10 | 9 | 8 | 4 | 5 | 3 | 3 | 5 |
| 8 | TGME49_214510-t26_1-p1 | hypothetical protein | 10 | 9 | 7 | 3 | 3 | 3 | 3 | 5 |
| 13 | TGME49_214530-t26_1-p1 | DnaJ domain-containing protein | 10 | 8 | 7 | 5 | 5 | 3 | 3 | 5 |
| 15 | TGME49_214600-t26_1-p1 | hypothetical protein | 10 | 10 | 9 | 4 | 4 | 3 | 3 | 5 |
| 14 | TGME49_214810-t26_1-p1 | PRA1 family protein, putative | 8 | 7 | 6 | 3 | 5 | 5 | 4 | 5 |
| 13 | TGME49_214910-t26_1-p1 | hypothetical protein | 9 | 8 | 7 | 5 | 4 | 3 | 3 | 5 |
| 19 | TGME49_214920.P836 | GPI-anchored wall transfer protein GWT1, putative | 10 | 9 | 8 | 4 | 4 | 5 | 4 | 5 |
| 15 | TGME49_214960.P840 | AP2 domain transcription factor AP2X-8 | 10 | 10 | 10 | 5 | 5 | 5 | 4 | 5 |
| 8 | TGME49_214980.P841 | hypothetical protein | 10 | 9 | 6 | 3 | 3 | 3 | 3 | 5 |
| 13 | TGME49_215010.P843 | hypothetical protein | 9 | 8 | 7 | 5 | 4 | 4 | 3 | 5 |
| 14 | TGME49_215030-t26_1-p1 | helicase, putative | 10 | 8 | 7 | 4 | 6 | 5 | 3 | 5 |
| 15 | TGME49_215040.P844 | HEAT repeat-containing protein | 10 | 9 | 10 | 4 | 3 | 3 | 4 | 5 |
| 13 | TGME49_215100-t26_1-p1 | PP-loop family protein | 10 | 8 | 7 | 5 | 3 | 4 | 3 | 5 |
| 15 | TGME49_215150-t26_1-p1 | AP2 domain transcription factor AP2X-9 | 10 | 10 | 9 | 5 | 5 | 5 | 5 | 5 |
| 14 | TGME49_215490.P858 | novel putative transporter NPT1 | 9 | 9 | 9 | 3 | 8 | 3 | 7 | 5 |
| 15 | TGME49_215510.P860 | MraW methylase family protein | 10 | 9 | 7 | 6 | 5 | 4 | 4 | 5 |
| 13 | TGME49_215520-t26_1-p1 | hypothetical protein | 9 | 8 | 6 | 5 | 4 | 3 | 3 | 5 |
| 13 | TGME49_215570.P863 | AP2 domain transcription factor AP2X-11 | 10 | 7 | 7 | 5 | 4 | 4 | 4 | 5 |
| 19 | TGME49_215670-t26_1-p1 | cAMP-dependent protein kinase | 10 | 10 | 7 | 3 | 3 | 5 | 5 | 5 |
| 15 | TGME49_215710-t26_1-p1 | hypothetical protein | 10 | 8 | 9 | 6 | 5 | 3 | 4 | 5 |
| 15 | TGME49_215750.P866 | cytidine and deoxycytidylate deaminase zinc-binding region domain-containing protein | 9 | 9 | 8 | 5 | 4 | 4 | 4 | 5 |
| 8 | TGME49_215910-t26_1-p1 | hypothetical protein | 10 | 10 | 10 | 3 | 3 | 3 | 3 | 5 |
| 13 | TGME49_215950.P871 | hypothetical protein | 10 | 9 | 7 | 6 | 3 | 3 | 3 | 5 |
| 19 | TGME49_215990.P872 | helicase, putative | 10 | 9 | 8 | 5 | 3 | 6 | 5 | 5 |
| 15 | TGME49_216020-t26_1-p1 | Ufm1-specific protease UFSP | 10 | 9 | 9 | 4 | 6 | 3 | 4 | 5 |
| 15 | TGME49_216120-t26_1-p1 | hypothetical protein | 10 | 9 | 8 | 6 | 5 | 4 | 3 | 5 |
| 14 | TGME49_216140.P877 | tetratricopeptide repeat-containing protein ANK1 | 9 | 9 | 8 | 3 | 8 | 3 | 5 | 5 |
| 14 | TGME49_216220.P879 | AP2 domain transcription factor AP2XI-5 | 10 | 9 | 7 | 4 | 5 | 4 | 5 | 5 |
| 8 | TGME49_216240-t26_1-p1 | hypothetical protein | 10 | 8 | 8 | 4 | 5 | 3 | 3 | 5 |
| 13 | TGME49_216285.P881 | zinc finger, c2h2 type domain-containing protein | 8 | 7 | 9 | 7 | 3 | 4 | 3 | 5 |
| 14 | TGME49_216400-t26_1-p1 | meiotic recombination protein DMC1 family protein | 9 | 9 | 8 | 3 | 8 | 8 | 3 | 5 |
| 13 | TGME49_216460.P885 | hypothetical protein | 9 | 8 | 7 | 6 | 4 | 3 | 4 | 5 |
| 15 | TGME49_216500-t26_1-p1 | tRNA synthetase, putative | 10 | 10 | 7 | 5 | 5 | 4 | 4 | 5 |
| 13 | TGME49_216640-t26_1-p1 | GHMP kinase, putative | 10 | 8 | 9 | 6 | 3 | 3 | 3 | 5 |
| 15 | TGME49_216760-t26_1-p1 | RNA pseudouridine synthase superfamily protein | 10 | 9 | 8 | 5 | 5 | 3 | 3 | 5 |
| 13 | TGME49_216830-t26_1-p1 | DTW domain-containing protein | 9 | 8 | 7 | 6 | 3 | 3 | 5 | 5 |
| 14 | TGME49_217160.P898 | Der1 family protein | 7 | 7 | 7 | 3 | 6 | 7 | 5 | 5 |
| 15 | TGME49_217360-t26_1-p1 | hypothetical protein | 10 | 10 | 9 | 6 | 5 | 4 | 4 | 5 |
| 8 | TGME49_217380.P901 | hypothetical protein | 10 | 10 | 9 | 3 | 3 | 3 | 3 | 5 |
| 8 | TGME49_217390.P902 | hypothetical protein | 9 | 10 | 10 | 3 | 3 | 3 | 4 | 5 |

| | | | | | | | | | | |
|---|---|---|---|---|---|---|---|---|---|---|
| 13 | TGME49_217400-t26_1-p1 | hypothetical protein | 10 | 9 | 8 | 5 | 3 | 3 | 3 | 5 |
| 13 | TGME49_217410-t26_1-p1 | hypothetical protein | 10 | 8 | 7 | 5 | 3 | 3 | 3 | 5 |
| 13 | TGME49_217420.P903 | hypothetical protein | 10 | 7 | 8 | 7 | 4 | 3 | 3 | 5 |
| 13 | TGME49_217430-t26_1-p1 | protease inhibitor PI1 | 9 | 8 | 8 | 7 | 3 | 3 | 3 | 5 |
| 13 | TGME49_217470.P905 | hypothetical protein | 10 | 10 | 8 | 5 | 3 | 3 | 3 | 5 |
| 8 | TGME49_217692.P920 | hypothetical protein | 10 | 10 | 8 | 4 | 3 | 3 | 3 | 5 |
| 15 | TGME49_217700.P921 | AP2 domain transcription factor AP2XII-2 | 10 | 10 | 9 | 3 | 5 | 4 | 4 | 5 |
| 15 | TGME49_218000-t26_1-p1 | hypothetical protein | 10 | 8 | 7 | 5 | 5 | 4 | 3 | 5 |
| 15 | TGME49_218070-t26_1-p1 | hypothetical protein | 10 | 10 | 9 | 5 | 7 | 5 | 3 | 5 |
| 13 | TGME49_218230-t26_1-p1 | histone lysine methyltransferase, SET, putative | 9 | 8 | 6 | 6 | 3 | 3 | 5 | 5 |
| 15 | TGME49_218240-t26_1-p1 | inner membrane complex protein IMC25 | 10 | 9 | 7 | 5 | 5 | 3 | 4 | 5 |
| 8 | TGME49_218270.P935 | rhoptry protein ROP48 | 9 | 8 | 7 | 3 | 3 | 3 | 3 | 5 |
| 19 | TGME49_218358-t26_1-p1 | zinc finger protein ZFP1 | 9 | 7 | 7 | 3 | 4 | 5 | 5 | 5 |
| 13 | TGME49_218362-t26_1-p1 | zinc finger protein | 10 | 9 | 7 | 6 | 3 | 4 | 3 | 5 |
| 8 | TGME49_218500.P942 | hypothetical protein | 10 | 9 | 7 | 4 | 3 | 4 | 4 | 5 |
| 14 | TGME49_218590.P945 | protein phosphatase PPM8 | 10 | 9 | 3 | 3 | 7 | 6 | 6 | 5 |
| 8 | TGME49_218750.P949 | nucleoside triphosphate hydrolase, putative | 10 | 10 | 9 | 3 | 3 | 3 | 3 | 5 |
| 8 | TGME49_218800-t26_1-p1 | hypothetical protein | 10 | 9 | 7 | 4 | 4 | 4 | 3 | 5 |
| 13 | TGME49_218830-t26_1-p1 | hypothetical protein | 10 | 9 | 8 | 6 | 4 | 3 | 4 | 5 |
| 8 | TGME49_218900.P954 | hypothetical protein | 10 | 9 | 7 | 3 | 3 | 4 | 3 | 5 |
| 8 | TGME49_218950-t26_1-p1 | hypothetical protein | 10 | 8 | 7 | 3 | 4 | 4 | 3 | 5 |
| 19 | TGME49_219120.P961 | zinc finger (CCCH type) motif-containing protein | 10 | 9 | 6 | 3 | 3 | 5 | 5 | 5 |
| 15 | TGME49_219130.P962 | NADPH-glutathione reductase | 10 | 9 | 9 | 3 | 6 | 3 | 3 | 5 |
| 15 | TGME49_219440-t26_1-p1 | hypothetical protein | 10 | 10 | 9 | 4 | 4 | 4 | 4 | 5 |
| 15 | TGME49_219460.P971 | hypothetical protein | 10 | 10 | 10 | 6 | 5 | 4 | 4 | 5 |
| 13 | TGME49_219485-t26_1-p1 | peptidase, putative | 10 | 8 | 6 | 5 | 4 | 4 | 4 | 5 |
| 8 | TGME49_219600.P976 | hypothetical protein | 10 | 8 | 6 | 4 | 4 | 3 | 4 | 5 |
| 8 | TGME49_219620.P978 | hypothetical protein | 9 | 8 | 6 | 4 | 4 | 3 | 3 | 5 |
| 13 | TGME49_219670-t26_1-p1 | zinc finger (CCCH type) motif-containing protein | 9 | 9 | 8 | 6 | 3 | 3 | 3 | 5 |
| 13 | TGME49_219738-t26_1-p1 | armadillo repeat-containing protein | 10 | 8 | 8 | 5 | 4 | 3 | 3 | 5 |
| 14 | TGME49_219770.P983 | 30S ribosomal protein S12, putative | 8 | 7 | 7 | 5 | 7 | 4 | 6 | 5 |
| 19 | TGME49_219820-t26_1-p1 | polyubiquitin UbC, putative | 8 | 9 | 10 | 3 | 3 | 3 | 6 | 5 |
| 15 | TGME49_220110-t26_1-p1 | hypothetical protein | 10 | 10 | 9 | 4 | 4 | 4 | 4 | 5 |
| 8 | TGME49_220132.P988 | hypothetical protein | 10 | 10 | 7 | 3 | 4 | 3 | 3 | 5 |
| 8 | TGME49_220208-t26_1-p1 | hypothetical protein | 10 | 9 | 7 | 4 | 4 | 3 | 4 | 5 |
| 19 | TGME49_220212-t26_1-p1 | pseudouridine synthase | 9 | 8 | 8 | 5 | 3 | 6 | 6 | 5 |
| 15 | TGME49_220230-t26_1-p1 | leucine rich repeat-containing protein | 10 | 9 | 9 | 6 | 5 | 3 | 3 | 5 |
| 19 | TGME49_220340.P996 | transporter, putative | 10 | 9 | 8 | 3 | 5 | 5 | 5 | 5 |
| 15 | TGME49_220360-t26_1-p1 | FAD binding domain-containing protein | 10 | 9 | 9 | 5 | 6 | 5 | 4 | 5 |
| 8 | TGME49_220370-t26_1-p1 | hypothetical protein | 10 | 9 | 6 | 4 | 3 | 3 | 3 | 5 |
| 8 | TGME49_220380-t26_1-p1 | hypothetical protein | 10 | 8 | 6 | 4 | 4 | 4 | 4 | 5 |
| 13 | TGME49_220430-t26_1-p1 | hypothetical protein | 10 | 9 | 8 | 5 | 3 | 3 | 4 | 5 |
| 8 | TGME49_220440-t26_1-p1 | cyclin-dependent kinase regulatory subunit protein | 10 | 7 | 6 | 3 | 3 | 5 | 3 | 5 |
| 8 | TGME49_220480.P1000 | hypothetical protein | 10 | 8 | 6 | 4 | 4 | 3 | 3 | 5 |
| 8 | TGME49_220560.P1001 | transmembrane protein TauE, putative | 9 | 8 | 8 | 3 | 5 | 3 | 4 | 5 |
| 15 | TGME49_220570.P1002 | zinc finger protein, putative | 9 | 9 | 8 | 4 | 5 | 4 | 4 | 5 |
| 14 | TGME49_220610.P1004 | protein phosphatase PPM9 | 9 | 8 | 6 | 5 | 5 | 5 | 5 | 5 |
| 13 | TGME49_220840.P1005 | threonine synthase | 9 | 9 | 8 | 6 | 3 | 4 | 4 | 5 |
| 13 | TGME49_220890.P1006 | dense granule protein GRA12A | 9 | 8 | 7 | 5 | 3 | 3 | 3 | 5 |

| | | | | | | | | | | |
|---|---|---|---|---|---|---|---|---|---|---|
| 13 | TGME49_220900.P1007 | apical cap protein AC13 | 10 | 9 | 7 | 6 | 3 | 3 | 3 | 5 |
| 15 | TGME49_220920-t26_1-p1 | hypothetical protein | 10 | 9 | 9 | 5 | 4 | 4 | 4 | 5 |
| 8 | TGME49_221200-t26_1-p1 | CW-type Zinc Finger protein | 10 | 9 | 6 | 4 | 5 | 4 | 3 | 5 |
| 8 | TGME49_221280.P1017 | hypothetical protein | 10 | 10 | 7 | 4 | 4 | 4 | 3 | 5 |
| 13 | TGME49_221340-t26_1-p1 | CRAL-TRIO domain-containing protein | 9 | 7 | 7 | 6 | 3 | 4 | 3 | 5 |
| 13 | TGME49_221370-t26_1-p1 | hypothetical protein | 10 | 8 | 8 | 6 | 3 | 4 | 3 | 5 |
| 13 | TGME49_221390.P1020 | hypothetical protein | 10 | 9 | 6 | 7 | 4 | 3 | 4 | 5 |
| 8 | TGME49_221400-t26_1-p1 | hypothetical protein | 10 | 10 | 5 | 3 | 3 | 3 | 3 | 5 |
| 14 | TGME49_221490.P1024 | cell cycle regulator protein | 7 | 7 | 7 | 3 | 6 | 6 | 5 | 5 |
| 14 | TGME49_221500.P1025 | dual specificity phosphatase, catalytic domain-containing protein | 10 | 7 | 7 | 5 | 5 | 5 | 5 | 5 |
| 14 | TGME49_221590.P1031 | dual specificity phosphatase, catalytic domain-containing protein | 8 | 7 | 6 | 3 | 5 | 5 | 5 | 5 |
| 13 | TGME49_221610.P1032 | ubiquitin carboxyl-terminal hydrolase | 10 | 9 | 8 | 6 | 3 | 3 | 3 | 5 |
| 13 | TGME49_221630.P1033 | IMC-associated protein IAP4 | 10 | 9 | 7 | 6 | 3 | 3 | 3 | 5 |
| 15 | TGME49_221870-t26_1-p1 | hypothetical protein | 10 | 9 | 8 | 6 | 5 | 4 | 3 | 5 |
| 8 | TGME49_222050.P1047 | hypothetical protein | 10 | 9 | 8 | 4 | 3 | 3 | 3 | 5 |
| 13 | TGME49_222120-t26_1-p1 | pseudouridine synthase, putative | 10 | 8 | 6 | 5 | 5 | 3 | 3 | 5 |
| 14 | TGME49_222130-t26_1-p1 | hypothetical protein | 7 | 7 | 7 | 3 | 6 | 3 | 6 | 5 |
| 13 | TGME49_222192.P1052 | protein HGH1, putative | 9 | 8 | 8 | 5 | 3 | 4 | 4 | 5 |
| 13 | TGME49_222250-t26_1-p1 | hypothetical protein | 10 | 9 | 7 | 5 | 3 | 4 | 4 | 5 |
| 15 | TGME49_222300-t26_1-p1 | hypothetical protein | 10 | 8 | 7 | 4 | 5 | 4 | 4 | 5 |
| 14 | TGME49_222310-t26_1-p1 | THH1 domain-containing protein | 9 | 7 | 7 | 4 | 6 | 6 | 4 | 5 |
| 15 | TGME49_222360.P1056 | E3 UFM1-protein ligase 1 | 10 | 10 | 8 | 6 | 5 | 5 | 4 | 5 |
| 8 | TGME49_222400-t26_1-p1 | hypothetical protein | 10 | 9 | 7 | 4 | 4 | 3 | 3 | 5 |
| 8 | TGME49_222420-t26_1-p1 | hypothetical protein | 9 | 8 | 7 | 3 | 3 | 3 | 3 | 5 |
| 13 | TGME49_222700.P1062 | hypothetical protein | 9 | 8 | 8 | 5 | 3 | 3 | 3 | 5 |
| 14 | TGME49_222840.P1063 | Ser/Thr phosphatase family protein | 8 | 7 | 7 | 3 | 6 | 5 | 4 | 5 |
| 15 | TGME49_222880.P1067 | hypothetical protein | 10 | 9 | 8 | 5 | 4 | 4 | 3 | 5 |
| 15 | TGME49_222900.P1068 | phosphoserine phosphatase | 10 | 10 | 9 | 5 | 6 | 5 | 3 | 5 |
| 8 | TGME49_223030-t26_1-p1 | tetratricopeptide repeat-containing protein | 9 | 8 | 6 | 3 | 3 | 3 | 4 | 5 |
| 13 | TGME49_223095-t26_1-p1 | hypothetical protein | 10 | 6 | 7 | 6 | 4 | 3 | 4 | 5 |
| 19 | TGME49_223130-t26_1-p1 | tRNA:m(4)X modification enzyme TRM13, putative | 10 | 7 | 8 | 5 | 4 | 5 | 5 | 5 |
| 13 | TGME49_223270-t26_1-p1 | hypothetical protein | 10 | 8 | 8 | 6 | 4 | 3 | 3 | 5 |
| 13 | TGME49_223440.P1079 | DEAD/DEAH box helicase domain-containing protein | 10 | 8 | 9 | 6 | 3 | 4 | 3 | 5 |
| 15 | TGME49_223500.P1083 | hypothetical protein | 10 | 9 | 9 | 6 | 5 | 4 | 3 | 5 |
| 13 | TGME49_223570-t26_1-p1 | zinc finger, C2H2 type domain-containing protein | 9 | 8 | 7 | 6 | 3 | 3 | 4 | 5 |
| 13 | TGME49_223600-t26_1-p1 | hypothetical protein | 9 | 8 | 8 | 6 | 4 | 3 | 3 | 5 |
| 15 | TGME49_223640.P1087 | TBC domain-containing protein TBC12 | 10 | 9 | 8 | 5 | 4 | 4 | 4 | 5 |
| 8 | TGME49_223750-t26_1-p1 | hypothetical protein | 9 | 8 | 6 | 4 | 3 | 3 | 3 | 5 |
| 15 | TGME49_223760-t26_1-p1 | hypothetical protein | 10 | 10 | 9 | 5 | 4 | 4 | 3 | 5 |
| 13 | TGME49_223870.P1095 | haloacid dehalogenase-like hydrolase, putative | 10 | 8 | 7 | 7 | 3 | 3 | 3 | 5 |
| 8 | TGME49_224000-t26_1-p1 | IMC-associated protein IAP5 | 10 | 9 | 7 | 4 | 4 | 4 | 4 | 5 |
| 8 | TGME49_224010-t26_1-p1 | hypothetical protein | 10 | 9 | 6 | 3 | 3 | 3 | 3 | 5 |
| 8 | TGME49_224030.P1103 | hypothetical protein | 10 | 8 | 7 | 4 | 3 | 4 | 4 | 5 |
| 13 | TGME49_224070.P1105 | hypothetical protein | 10 | 8 | 7 | 6 | 3 | 4 | 3 | 5 |
| 13 | TGME49_224110.P1108 | 26S proteasome regulatory subunit RPN13, putative | 8 | 8 | 7 | 5 | 3 | 4 | 4 | 5 |
| 13 | TGME49_224140-t26_1-p1 | hypothetical protein | 10 | 6 | 8 | 5 | 3 | 3 | 3 | 5 |

| | | | | | | | | | | |
|---|---|---|---|---|---|---|---|---|---|---|
| 15 | TGME49_224150-t26_1-p1 | conserved oligomeric Golgi complex subunit 6 | 10 | 10 | 7 | 6 | 5 | 4 | 3 | 5 |
| 15 | TGME49_224210.P1110 | hypothetical protein | 10 | 10 | 9 | 5 | 6 | 4 | 4 | 5 |
| 15 | TGME49_224230-t26_1-p1 | AP2 domain transcription factor AP2X-3 | 10 | 10 | 9 | 4 | 5 | 4 | 4 | 5 |
| 15 | TGME49_224270.P1112 | vacuolar protein sorting-associated protein 41 | 10 | 10 | 8 | 6 | 4 | 4 | 4 | 5 |
| 8 | TGME49_224290.P1113 | palmitoyltransferase DHHC5 | 9 | 8 | 7 | 3 | 5 | 3 | 3 | 5 |
| 13 | TGME49_224320-t26_1-p1 | hypothetical protein | 10 | 9 | 7 | 6 | 4 | 4 | 4 | 5 |
| 8 | TGME49_224470.P1118 | hypothetical protein | 8 | 8 | 8 | 3 | 4 | 3 | 3 | 5 |
| 13 | TGME49_224570-t26_1-p1 | hypothetical protein | 9 | 8 | 8 | 7 | 4 | 3 | 3 | 5 |
| 13 | TGME49_224670-t26_1-p1 | DnaJ domain-containing protein | 10 | 9 | 7 | 5 | 4 | 4 | 3 | 5 |
| 13 | TGME49_224680.P1124 | hypothetical protein | 9 | 9 | 8 | 5 | 3 | 4 | 3 | 5 |
| 14 | TGME49_224710.P1126 | PA domain-containing protein | 9 | 9 | 9 | 3 | 9 | 3 | 3 | 5 |
| 19 | TGME49_224720-t26_1-p1 | SPOC domain-containing protein | 10 | 9 | 8 | 5 | 3 | 5 | 4 | 5 |
| 8 | TGME49_224800-t26_1-p1 | hypothetical protein | 8 | 8 | 7 | 3 | 3 | 3 | 5 | 5 |
| 13 | TGME49_224928-t26_1-p1 | histone lysine demethylase JMJD4 | 10 | 9 | 7 | 6 | 3 | 3 | 3 | 5 |
| 13 | TGME49_224935-t26_1-p1 | hypothetical protein | 10 | 7 | 7 | 5 | 4 | 5 | 4 | 5 |
| 15 | TGME49_224970.P1138 | nucleolar protein Nop52, putative | 10 | 9 | 8 | 6 | 4 | 4 | 4 | 5 |
| 15 | TGME49_224980-t26_1-p1 | hypothetical protein | 10 | 9 | 9 | 6 | 6 | 4 | 3 | 5 |
| 13 | TGME49_225010.P1141 | hypothetical protein | 10 | 8 | 7 | 6 | 4 | 4 | 3 | 5 |
| 13 | TGME49_225105.P1149 | hypothetical protein | 10 | 8 | 8 | 6 | 3 | 3 | 3 | 5 |
| 13 | TGME49_225150.P1154 | hypothetical protein | 10 | 8 | 7 | 6 | 3 | 3 | 3 | 5 |
| 15 | TGME49_225220.P1158 | SCP family extracellular subfamily protein | 10 | 8 | 9 | 5 | 5 | 3 | 4 | 5 |
| 8 | TGME49_225270.P1161 | hypothetical protein | 10 | 8 | 6 | 4 | 3 | 3 | 3 | 5 |
| 15 | TGME49_225280.P1162 | hypothetical protein | 10 | 10 | 8 | 3 | 6 | 3 | 3 | 5 |
| 19 | TGME49_225320.P1164 | hypothetical protein | 9 | 8 | 8 | 3 | 3 | 5 | 6 | 5 |
| 13 | TGME49_225330.P1165 | hypothetical protein | 9 | 8 | 7 | 6 | 3 | 3 | 3 | 5 |
| 14 | TGME49_225340-t26_1-p1 | hypothetical protein | 10 | 10 | 5 | 3 | 6 | 4 | 4 | 5 |
| 13 | TGME49_225400-t26_1-p1 | hypothetical protein | 10 | 9 | 6 | 6 | 3 | 3 | 3 | 5 |
| 13 | TGME49_225450.P1168 | COP9 signalosome complex subunit 3, putative | 9 | 9 | 8 | 6 | 4 | 3 | 3 | 5 |
| 14 | TGME49_225470.P1170 | peptide methionine sulfoxide reductase | 8 | 7 | 7 | 3 | 6 | 5 | 3 | 5 |
| 15 | TGME49_225530.P1173 | metal cation transporter, ZIP family protein | 10 | 9 | 8 | 5 | 6 | 5 | 4 | 5 |
| 14 | TGME49_225580-t26_1-p1 | proteasome (prosome, macropain) 26S subunit, non-ATPase, 9, putative | 8 | 7 | 6 | 3 | 6 | 5 | 4 | 5 |
| 13 | TGME49_225690.P1176 | apical cap protein AC7 | 10 | 9 | 6 | 5 | 4 | 4 | 4 | 5 |
| 8 | TGME49_225820-t26_1-p1 | hypothetical protein | 9 | 8 | 7 | 3 | 3 | 3 | 3 | 5 |
| 13 | TGME49_225840-t26_1-p1 | hypothetical protein | 9 | 8 | 7 | 5 | 4 | 3 | 4 | 5 |
| 8 | TGME49_225870-t26_1-p1 | hypothetical protein | 10 | 10 | 8 | 4 | 3 | 3 | 3 | 5 |
| 15 | TGME49_225900.P1183 | hypothetical protein | 10 | 9 | 9 | 3 | 4 | 4 | 4 | 5 |
| 19 | TGME49_225960.P1190 | STE kinase | 10 | 10 | 9 | 5 | 5 | 7 | 6 | 5 |
| 15 | TGME49_225980-t26_1-p1 | hypothetical protein | 10 | 10 | 9 | 4 | 4 | 4 | 3 | 5 |
| 13 | TGME49_226230-t26_1-p1 | hypothetical protein | 9 | 8 | 8 | 7 | 3 | 3 | 3 | 5 |
| 13 | TGME49_226310-t26_1-p1 | zinc finger (CCCH type) motif-containing protein | 10 | 9 | 7 | 6 | 3 | 3 | 5 | 5 |
| 13 | TGME49_226350-t26_1-p1 | hypothetical protein | 10 | 9 | 6 | 5 | 3 | 4 | 3 | 5 |
| 8 | TGME49_226450.P1208 | hypothetical protein | 9 | 8 | 6 | 4 | 3 | 3 | 3 | 5 |
| 15 | TGME49_226470.P1209 | hypothetical protein | 10 | 10 | 7 | 5 | 5 | 4 | 4 | 5 |
| 13 | TGME49_226610.P1217 | centrosomal protein CEP76, putative | 10 | 8 | 8 | 7 | 3 | 3 | 3 | 5 |
| 15 | TGME49_226620.P1218 | hypothetical protein | 10 | 10 | 9 | 5 | 4 | 3 | 3 | 5 |
| 15 | TGME49_226680-t26_1-p1 | hypothetical protein | 10 | 9 | 7 | 6 | 5 | 4 | 4 | 5 |
| 13 | TGME49_226700-t26_1-p1 | nuclease, putative | 9 | 8 | 7 | 7 | 3 | 5 | 3 | 5 |
| 8 | TGME49_226710-t26_1-p1 | hypothetical protein | 8 | 7 | 7 | 3 | 3 | 3 | 3 | 5 |

| | | | | | | | | | | |
|---|---|---|---|---|---|---|---|---|---|---|
| 14 | TGME49_226740-t26_1-p1 | zinc finger, C3HC4 type (RING finger) domain-containing protein | 8 | 8 | 8 | 5 | 7 | 4 | 6 | 5 |
| 19 | TGME49_226755.P1222 | 3'5'-cyclic nucleotide phosphodiesterase PDE18 | 10 | 9 | 8 | 3 | 4 | 6 | 6 | 5 |
| 15 | TGME49_226790-t26_1-p1 | ABC transporter, ATP-binding domain-containing protein | 10 | 9 | 8 | 6 | 4 | 5 | 4 | 5 |
| 13 | TGME49_226800-t26_1-p1 | eukaryotic glutathione synthase, atp binding domain-containing protein | 10 | 9 | 7 | 5 | 4 | 4 | 3 | 5 |
| 15 | TGME49_226890.P1228 | origin recognition complex subunit 3, putative | 10 | 10 | 8 | 5 | 5 | 4 | 4 | 5 |
| 14 | TGME49_226900.P1229 | hypothetical protein | 10 | 7 | 6 | 4 | 5 | 5 | 4 | 5 |
| 14 | TGME49_226970.P1232 | ribosomal protein RPS11 | 7 | 7 | 6 | 3 | 6 | 6 | 6 | 5 |
| 15 | TGME49_227010.P1236 | rhoptry kinase family protein ROP30 | 9 | 9 | 9 | 3 | 6 | 3 | 4 | 5 |
| 13 | TGME49_227050.P1238 | ATPase/histidine kinase/DNA gyrase B/HSP90 domain-containing protein | 10 | 8 | 8 | 5 | 4 | 3 | 3 | 5 |
| 15 | TGME49_227060-t26_1-p1 | hypothetical protein | 10 | 10 | 9 | 5 | 4 | 4 | 4 | 5 |
| 19 | TGME49_227140-t26_1-p1 | PUA domain-containing protein | 10 | 9 | 8 | 4 | 4 | 5 | 4 | 5 |
| 15 | TGME49_227300.P1242 | hypothetical protein | 10 | 10 | 8 | 5 | 4 | 3 | 3 | 5 |
| 13 | TGME49_227320-t26_1-p1 | hypothetical protein | 10 | 7 | 7 | 5 | 4 | 4 | 4 | 5 |
| 15 | TGME49_227350.P1244 | hypothetical protein | 10 | 10 | 9 | 3 | 5 | 5 | 4 | 5 |
| 13 | TGME49_227400.P1247 | hypothetical protein | 9 | 7 | 7 | 6 | 3 | 3 | 3 | 5 |
| 13 | TGME49_227430-t26_1-p1 | amino acid transporter AAT2 | 10 | 8 | 8 | 6 | 3 | 3 | 3 | 5 |
| 13 | TGME49_227580.P1251 | amino acid transporter AAT4 | 9 | 9 | 8 | 6 | 3 | 3 | 3 | 5 |
| 14 | TGME49_227860-t26_1-p1 | hypothetical protein | 8 | 7 | 7 | 4 | 5 | 5 | 5 | 5 |
| 8 | TGME49_227900.P1265 | AP2 domain transcription factor AP2X-1 | 10 | 8 | 6 | 4 | 4 | 3 | 3 | 5 |
| 15 | TGME49_227990.P1271 | hypothetical protein | 10 | 10 | 9 | 5 | 6 | 5 | 3 | 5 |
| 8 | TGME49_228030-t26_1-p1 | hypothetical protein | 10 | 8 | 7 | 4 | 4 | 3 | 3 | 5 |
| 8 | TGME49_228150.P1277 | hypothetical protein | 10 | 10 | 7 | 4 | 3 | 3 | 3 | 5 |
| 13 | TGME49_228200.P1282 | vacuolar (h+)-atpase g subunit protein | 9 | 8 | 7 | 6 | 4 | 4 | 4 | 5 |
| 15 | TGME49_228230.P1285 | hypothetical protein | 10 | 9 | 8 | 5 | 5 | 4 | 4 | 5 |
| 14 | TGME49_228240.P1286 | hypothetical protein | 10 | 9 | 7 | 3 | 6 | 5 | 5 | 5 |
| 14 | TGME49_228320-t26_1-p1 | ribosome-binding factor A, putative | 9 | 8 | 7 | 4 | 6 | 6 | 6 | 5 |
| 8 | TGME49_228340-t26_1-p1 | hypothetical protein | 10 | 9 | 7 | 4 | 4 | 3 | 3 | 5 |
| 8 | TGME49_228380-t26_1-p1 | hypothetical protein | 10 | 10 | 6 | 4 | 3 | 3 | 3 | 5 |
| 13 | TGME49_228430-t26_1-p1 | hypothetical protein | 9 | 8 | 6 | 5 | 3 | 3 | 3 | 5 |
| 15 | TGME49_228670.P1299 | zinc finger, C2H2 type domain-containing protein | 10 | 9 | 7 | 5 | 5 | 4 | 3 | 5 |
| 15 | TGME49_228710.P1302 | hypothetical protein | 10 | 8 | 8 | 5 | 5 | 3 | 4 | 5 |
| 13 | TGME49_228990.P1307 | zinc finger (CCCH type) motif-containing protein | 8 | 7 | 7 | 6 | 3 | 3 | 3 | 5 |
| 19 | TGME49_229010-t26_1-p1 | rhoptry neck protein RON4 | 10 | 9 | 9 | 5 | 3 | 5 | 4 | 5 |
| 14 | TGME49_229030.P1309 | signal peptidase complex subunit 2, putative | 7 | 7 | 6 | 3 | 6 | 4 | 4 | 5 |
| 13 | TGME49_229140.P1310 | MaoC family domain-containing protein | 8 | 8 | 7 | 7 | 3 | 3 | 3 | 5 |
| 8 | TGME49_229280-t26_1-p1 | hypothetical protein | 10 | 9 | 7 | 3 | 4 | 3 | 3 | 5 |
| 8 | TGME49_229320-t26_1-p1 | Cof family hydrolase subfamily protein | 8 | 7 | 8 | 3 | 3 | 3 | 5 | 5 |
| 19 | TGME49_229330-t26_1-p1 | Cof family hydrolase subfamily protein | 8 | 8 | 8 | 3 | 3 | 6 | 5 | 5 |
| 15 | TGME49_229350.P1318 | HEAT repeat-containing protein | 10 | 8 | 9 | 5 | 7 | 4 | 4 | 5 |
| 8 | TGME49_229380.P1321 | hypothetical protein | 10 | 7 | 7 | 4 | 5 | 3 | 3 | 5 |
| 13 | TGME49_229640.P1330 | apical cap protein AC8 | 10 | 9 | 6 | 5 | 4 | 4 | 4 | 5 |
| 15 | TGME49_229750.P1338 | hypothetical protein | 10 | 10 | 9 | 4 | 4 | 3 | 3 | 5 |
| 8 | TGME49_229770.P1339 | hypothetical protein | 10 | 8 | 6 | 3 | 4 | 3 | 3 | 5 |
| 8 | TGME49_229790.P1341 | hypothetical protein | 10 | 8 | 7 | 4 | 4 | 3 | 4 | 5 |
| 14 | TGME49_230050.P1349 | 50S ribosomal protein L3, putative | 9 | 8 | 8 | 5 | 6 | 5 | 5 | 5 |
| 13 | TGME49_230060-t26_1-p1 | acetyltransferase, GNAT family protein | 8 | 8 | 7 | 6 | 3 | 4 | 4 | 5 |

| | | | | | | | | | | |
|---|---|---|---|---|---|---|---|---|---|---|
| 13 | TGME49_230110.P1352 | hypothetical protein | 10 | 9 | 6 | 5 | 4 | 3 | 3 | 5 |
| 15 | TGME49_230170.P1358 | hypothetical protein | 10 | 10 | 9 | 4 | 3 | 4 | 4 | 5 |
| 13 | TGME49_230200.P1361 | hypothetical protein | 9 | 6 | 8 | 6 | 3 | 4 | 4 | 5 |
| 19 | TGME49_230400-t26_1-p1 | WIP-related protein domain-containing protein WPDP | 10 | 10 | 9 | 4 | 5 | 5 | 5 | 5 |
| 14 | TGME49_230430.P1368 | vesicle-associated membrane protein, putative | 7 | 7 | 7 | 4 | 6 | 3 | 6 | 5 |
| 13 | TGME49_230510.P1374 | hypothetical protein | 9 | 8 | 6 | 5 | 3 | 3 | 3 | 5 |
| 14 | TGME49_230570.P1377 | transporter, major facilitator family protein | 10 | 10 | 9 | 3 | 6 | 6 | 6 | 5 |
| 14 | TGME49_230590.P1379 | chitobiosyldiphosphodolichol beta-mannosyltransferase | 10 | 9 | 8 | 3 | 5 | 5 | 4 | 5 |
| 8 | TGME49_230840.P1386 | hypothetical protein | 9 | 8 | 7 | 4 | 3 | 3 | 3 | 5 |
| 8 | TGME49_230860-t26_1-p1 | autophagy-related protein ATG5, putative | 10 | 8 | 6 | 4 | 4 | 4 | 4 | 5 |
| 8 | TGME49_230880.P1388 | hypothetical protein | 10 | 7 | 6 | 4 | 4 | 4 | 3 | 5 |
| 15 | TGME49_230940-t26_1-p1 | hypothetical protein | 10 | 10 | 8 | 5 | 5 | 4 | 4 | 5 |
| 15 | TGME49_230990.P1395 | hypothetical protein | 10 | 9 | 8 | 4 | 4 | 4 | 4 | 5 |
| 15 | TGME49_231200.P1408 | hypothetical protein | 10 | 10 | 10 | 5 | 4 | 3 | 3 | 5 |
| 19 | TGME49_231420-t26_1-p1 | folate/biopterin transporter, putative | 9 | 9 | 9 | 3 | 3 | 8 | 3 | 5 |
| 14 | TGME49_231460.P1418 | hypothetical protein | 10 | 7 | 7 | 5 | 5 | 4 | 5 | 5 |
| 8 | TGME49_231790.P1423 | hypothetical protein | 10 | 10 | 5 | 4 | 4 | 4 | 4 | 5 |
| 15 | TGME49_231815.P1425 | hypothetical protein | 10 | 10 | 9 | 4 | 5 | 5 | 4 | 5 |
| 13 | TGME49_231920-t26_1-p1 | oxidoreductase, short chain dehydrogenase/reductase family protein | 8 | 7 | 7 | 6 | 3 | 3 | 3 | 5 |
| 15 | TGME49_231930.P1431 | RING zinc finger protein | 10 | 8 | 8 | 5 | 5 | 4 | 4 | 5 |
| 19 | TGME49_231940.P1432 | ubiquitin-like modifier-activating enzyme UBA5 | 8 | 8 | 8 | 3 | 5 | 7 | 7 | 5 |
| 15 | TGME49_231980.P1435 | hypothetical protein | 10 | 10 | 9 | 6 | 4 | 4 | 4 | 5 |
| 8 | TGME49_232000.P1437 | dense granule protein GRA30 | 10 | 8 | 7 | 3 | 3 | 4 | 4 | 5 |
| 13 | TGME49_232010.P1438 | protein phosphatase PPM14, putative | 10 | 9 | 6 | 5 | 4 | 3 | 4 | 5 |
| 15 | TGME49_232020.P1439 | rhoptry neck protein RON12 | 10 | 9 | 8 | 5 | 4 | 4 | 4 | 5 |
| 13 | TGME49_232060.P1443 | hypothetical protein | 9 | 8 | 7 | 6 | 4 | 3 | 3 | 5 |
| 8 | TGME49_232100-t26_1-p1 | RAP domain-containing protein | 10 | 8 | 7 | 4 | 4 | 4 | 4 | 5 |
| 15 | TGME49_232120.P1445 | hypothetical protein | 10 | 10 | 10 | 4 | 8 | 3 | 3 | 5 |
| 13 | TGME49_232270-t26_1-p1 | histidine acid phosphatase superfamily protein | 10 | 8 | 8 | 6 | 3 | 3 | 3 | 5 |
| 15 | TGME49_232310.P1457 | endonuclease/exonuclease/phosphatase family protein | 10 | 9 | 8 | 6 | 5 | 3 | 6 | 5 |
| 8 | TGME49_232370.P1461 | CW-type Zinc Finger protein | 10 | 8 | 6 | 4 | 4 | 3 | 3 | 5 |
| 13 | TGME49_232420-t26_1-p1 | hypothetical protein | 9 | 7 | 8 | 7 | 3 | 4 | 4 | 5 |
| 19 | TGME49_232500.P1468 | inner centromere protein INCENP1 | 10 | 10 | 9 | 3 | 3 | 7 | 5 | 5 |
| 15 | TGME49_232560-t26_1-p1 | hypothetical protein | 10 | 9 | 8 | 6 | 4 | 4 | 4 | 5 |
| 13 | TGME49_232670.P1479 | hypothetical protein | 10 | 8 | 6 | 6 | 4 | 3 | 3 | 5 |
| 13 | TGME49_232720-t26_1-p1 | Lsm12, putative | 8 | 7 | 7 | 6 | 3 | 3 | 3 | 5 |
| 19 | TGME49_232740-t26_1-p1 | hypothetical protein | 9 | 8 | 9 | 3 | 3 | 8 | 3 | 5 |
| 15 | TGME49_232780.P1482 | basal complex component BCC1 | 10 | 8 | 8 | 5 | 5 | 4 | 4 | 5 |
| 8 | TGME49_232800.P1483 | hypothetical protein | 10 | 9 | 8 | 4 | 4 | 4 | 3 | 5 |
| 15 | TGME49_233160.P1495 | hypothetical protein | 10 | 10 | 8 | 5 | 4 | 3 | 4 | 5 |
| 13 | TGME49_233210.P1500 | hypothetical protein | 10 | 10 | 6 | 5 | 4 | 3 | 3 | 5 |
| 15 | TGME49_233245-t26_1-p1 | hypothetical protein | 10 | 9 | 8 | 6 | 5 | 4 | 4 | 5 |
| 13 | TGME49_233270-t26_1-p1 | MORN repeat-containing protein | 9 | 8 | 7 | 6 | 3 | 3 | 3 | 5 |
| 8 | TGME49_233380-t26_1-p1 | hypothetical protein | 10 | 10 | 8 | 3 | 4 | 4 | 4 | 5 |
| 8 | TGME49_233430.P1507 | C2 domain containing protein | 10 | 10 | 7 | 4 | 4 | 3 | 3 | 5 |
| 13 | TGME49_233530-t26_1-p1 | hypothetical protein | 10 | 9 | 6 | 5 | 4 | 4 | 4 | 5 |

| | | | | | | | | | | |
|---|---|---|---|---|---|---|---|---|---|---|
| 15 | TGME49_233760.P1514 | GPR180 domain-containing protein | 9 | 9 | 8 | 5 | 4 | 5 | 4 | 5 |
| 15 | TGME49_233890-t26_1-p1 | hypothetical protein | 10 | 8 | 8 | 6 | 5 | 4 | 4 | 5 |
| 8 | TGME49_233920-t26_1-p1 | zinc finger, C3HC4 type (RING finger) domain-containing protein | 10 | 8 | 7 | 3 | 3 | 4 | 3 | 5 |
| 13 | TGME49_234165-t26_1-p1 | hypothetical protein | 8 | 10 | 9 | 5 | 3 | 3 | 3 | 5 |
| 15 | TGME49_234230.P1528 | hypothetical protein | 10 | 10 | 8 | 5 | 4 | 4 | 4 | 5 |
| 13 | TGME49_234390-t26_1-p1 | hypothetical protein | 9 | 8 | 6 | 5 | 3 | 3 | 3 | 5 |
| 13 | TGME49_234460.P1537 | zinc finger Ran-binding domain-containing protein | 8 | 7 | 7 | 6 | 3 | 3 | 3 | 5 |
| 13 | TGME49_234540-t26_1-p1 | hypothetical protein | 9 | 7 | 8 | 7 | 3 | 3 | 3 | 5 |
| 14 | TGME49_234580.P1542 | ImpB/MucB/SamB family protein | 10 | 8 | 6 | 5 | 5 | 5 | 5 | 5 |
| 13 | TGME49_234670-t26_1-p1 | actin-like family protein | 9 | 8 | 7 | 7 | 3 | 5 | 3 | 5 |
| 8 | TGME49_234940-t26_1-p1 | hypothetical protein | 9 | 8 | 6 | 3 | 3 | 3 | 3 | 5 |
| 15 | TGME49_234990-t26_1-p1 | hypothetical protein | 10 | 9 | 8 | 5 | 5 | 3 | 3 | 5 |
| 14 | TGME49_235200.P1551 | PAN domain-containing protein | 10 | 10 | 3 | 3 | 7 | 6 | 5 | 5 |
| 13 | TGME49_235380.P1556 | apical cap protein AC5 | 9 | 8 | 7 | 6 | 4 | 4 | 4 | 5 |
| 15 | TGME49_235420.P1559 | hypothetical protein | 10 | 10 | 9 | 4 | 4 | 4 | 3 | 5 |
| 8 | TGME49_235560-t26_1-p1 | hypothetical protein | 9 | 8 | 7 | 4 | 4 | 4 | 4 | 5 |
| 8 | TGME49_235640.P1568 | formyl transferase domain-containing protein | 10 | 8 | 8 | 3 | 3 | 3 | 3 | 5 |
| 13 | TGME49_235740.P1574 | hypothetical protein | 9 | 8 | 7 | 5 | 4 | 4 | 4 | 5 |
| 19 | TGME49_235905-t26_1-p1 | ribonuclease z, putative | 10 | 10 | 9 | 5 | 4 | 7 | 6 | 5 |
| 8 | TGME49_235960-t26_1-p1 | hypothetical protein | 9 | 8 | 7 | 3 | 3 | 4 | 4 | 5 |
| 14 | TGME49_235980.P1581 | ARIADNE family protein | 10 | 9 | 9 | 5 | 8 | 5 | 5 | 5 |
| 15 | TGME49_236010.P1583 | prenylcysteine oxidase | 10 | 9 | 8 | 3 | 6 | 3 | 3 | 5 |
| 19 | TGME49_236050.P1586 | fructose-bisphosphate aldolase, putative | 9 | 8 | 8 | 3 | 3 | 8 | 8 | 5 |
| 15 | TGME49_236140-t26_1-p1 | hypothetical protein | 10 | 9 | 9 | 5 | 3 | 4 | 4 | 5 |
| 15 | TGME49_236220.P1591 | PCI domain-containing protein | 9 | 9 | 8 | 4 | 5 | 4 | 4 | 5 |
| 14 | TGME49_236240.P1592 | tyrosine kinase-like protein TKL6 | 9 | 8 | 8 | 5 | 5 | 5 | 5 | 5 |
| 13 | TGME49_236260-t26_1-p1 | protein phosphatase PPM15, putative | 9 | 9 | 8 | 5 | 3 | 4 | 4 | 5 |
| 14 | TGME49_236510-t26_1-p1 | TMEM65 domain-containing protein | 8 | 8 | 8 | 4 | 6 | 5 | 6 | 5 |
| 8 | TGME49_236560.P1595 | hypothetical protein | 10 | 10 | 6 | 4 | 3 | 3 | 3 | 5 |
| 15 | TGME49_236780-t26_1-p1 | zinc finger, C3HC4 type (RING finger) domain-containing protein | 10 | 8 | 9 | 6 | 5 | 4 | 3 | 5 |
| 13 | TGME49_236790-t26_1-p1 | hypothetical protein | 9 | 7 | 7 | 6 | 4 | 4 | 4 | 5 |
| 8 | TGME49_236850-t26_1-p1 | hypothetical protein | 10 | 10 | 5 | 4 | 4 | 4 | 4 | 5 |
| 13 | TGME49_237080-t26_1-p1 | hypothetical protein | 9 | 8 | 7 | 6 | 3 | 3 | 3 | 5 |
| 15 | TGME49_237090.P1616 | AP2 domain transcription factor AP2X-5 | 10 | 10 | 8 | 5 | 5 | 5 | 5 | 5 |
| 8 | TGME49_237160.P1617 | hypothetical protein | 10 | 8 | 7 | 4 | 3 | 3 | 3 | 5 |
| 13 | TGME49_237195.P1620 | hypothetical protein | 10 | 9 | 8 | 6 | 3 | 3 | 3 | 5 |
| 19 | TGME49_237200.P1621 | sphingolipid delta(4)-desaturase DES | 9 | 9 | 8 | 3 | 3 | 7 | 7 | 5 |
| 13 | TGME49_237210.P1622 | tyrosine kinase-like protein TKL4 | 10 | 8 | 7 | 7 | 3 | 3 | 3 | 5 |
| 15 | TGME49_237290.P1626 | hypothetical protein | 10 | 10 | 8 | 6 | 6 | 3 | 3 | 5 |
| 13 | TGME49_237520-t26_1-p1 | SANT domain-containing protein | 10 | 9 | 7 | 7 | 3 | 5 | 3 | 5 |
| 13 | TGME49_237550.P1632 | KH domain-containing protein | 10 | 9 | 8 | 5 | 3 | 3 | 3 | 5 |
| 13 | TGME49_237860.P1636 | calcium-dependent protein kinase CDPK7A | 10 | 9 | 6 | 6 | 3 | 4 | 4 | 5 |
| 15 | TGME49_237870-t26_1-p1 | FYVE zinc finger domain-containing protein | 9 | 8 | 8 | 4 | 5 | 3 | 4 | 5 |
| 13 | TGME49_238130.P1643 | HXXEE motif protein | 9 | 8 | 7 | 7 | 3 | 3 | 3 | 5 |
| 13 | TGME49_238150-t26_1-p1 | hypothetical protein | 8 | 8 | 7 | 6 | 4 | 3 | 4 | 5 |
| 13 | TGME49_238160-t26_1-p1 | hypothetical protein | 8 | 7 | 7 | 6 | 3 | 3 | 3 | 5 |
| 19 | TGME49_238200-t26_1-p1 | alpha/beta hydrolase fold domain-containing protein | 10 | 9 | 8 | 5 | 3 | 6 | 6 | 5 |

| | | | | | | | | | | |
|---|---|---|---|---|---|---|---|---|---|---|
| 8 | TGME49_238210-t26_1-p1 | EGF family domain-containing protein | 8 | 7 | 7 | 3 | 3 | 3 | 3 | 5 |
| 13 | TGME49_238220-t26_1-p1 | EGF family domain-containing protein | 8 | 7 | 7 | 6 | 3 | 3 | 3 | 5 |
| 8 | TGME49_238895-t26_1-p1 | inner membrane complex protein IMC43 | 10 | 9 | 8 | 4 | 4 | 3 | 3 | 5 |
| 8 | TGME49_239050.P1663 | hypothetical protein | 10 | 8 | 7 | 4 | 4 | 3 | 3 | 5 |
| 19 | TGME49_239070.P1664 | zinc finger, C2H2 type domain-containing protein | 9 | 9 | 8 | 4 | 4 | 5 | 5 | 5 |
| 14 | TGME49_239080-t26_1-p1 | carrier superfamily protein | 9 | 8 | 5 | 5 | 5 | 5 | 6 | 5 |
| 19 | TGME49_239130.P1665 | tyrosine kinase-like protein TKL7 | 10 | 10 | 7 | 5 | 5 | 7 | 6 | 5 |
| 13 | TGME49_239350.P1671 | RAP domain-containing protein | 10 | 9 | 6 | 5 | 4 | 4 | 3 | 5 |
| 13 | TGME49_239520.P1678 | hypothetical protein | 10 | 8 | 7 | 7 | 3 | 3 | 3 | 5 |
| 13 | TGME49_239570-t26_1-p1 | hypothetical protein | 9 | 7 | 7 | 6 | 3 | 3 | 3 | 5 |
| 8 | TGME49_239600.P1680 | rhoptry kinase family protein ROP23 (incomplete catalytic triad) | 9 | 8 | 7 | 4 | 3 | 3 | 3 | 5 |
| 13 | TGME49_239630-t26_1-p1 | cytidine and deoxycytidylate deaminase zinc-binding region domain-containing protein | 9 | 8 | 7 | 6 | 3 | 4 | 4 | 5 |
| 13 | TGME49_239885-t26_1-p1 | protein kinase, putative | 10 | 8 | 6 | 5 | 4 | 5 | 4 | 5 |
| 8 | TGME49_239910.P1691 | cyclin-dependent kinase | 9 | 8 | 6 | 3 | 3 | 5 | 4 | 5 |
| 13 | TGME49_240050-t26_1-p1 | hypothetical protein | 10 | 10 | 6 | 5 | 3 | 3 | 3 | 5 |
| 13 | TGME49_240090.P1695 | rhoptry kinase family protein ROP34, putative | 9 | 7 | 7 | 6 | 3 | 3 | 3 | 5 |
| 13 | TGME49_240250.P1698 | macro domain-containing protein | 10 | 9 | 8 | 6 | 3 | 3 | 3 | 5 |
| 13 | TGME49_240410.P1702 | protein kinase (incomplete catalytic triad) | 10 | 9 | 6 | 5 | 4 | 4 | 3 | 5 |
| 8 | TGME49_240450-t26_1-p1 | Maf family protein | 8 | 7 | 7 | 3 | 3 | 4 | 3 | 5 |
| 14 | TGME49_240500.P1705 | hypothetical protein | 8 | 8 | 7 | 3 | 6 | 3 | 4 | 5 |
| 8 | TGME49_240510.P1706 | nucleoporin NUP134 | 10 | 10 | 9 | 3 | 4 | 3 | 3 | 5 |
| 19 | TGME49_240610-t26_1-p1 | hypothetical protein | 8 | 8 | 7 | 3 | 3 | 7 | 4 | 5 |
| 13 | TGME49_240700-t26_1-p1 | ubiquitin family protein | 9 | 8 | 7 | 5 | 3 | 4 | 3 | 5 |
| 13 | TGME49_240800-t26_1-p1 | MORN repeat-containing protein | 8 | 7 | 7 | 7 | 3 | 3 | 5 | 5 |
| 14 | TGME49_240840-t26_1-p1 | histone lysine demethylase JmjC NO66 | 9 | 8 | 7 | 5 | 6 | 4 | 6 | 5 |
| 13 | TGME49_240850-t26_1-p1 | WD domain, G-beta repeat-containing protein | 9 | 8 | 7 | 6 | 4 | 5 | 3 | 5 |
| 13 | TGME49_241130.P1724 | CKK domain-containing protein | 10 | 7 | 6 | 5 | 3 | 3 | 3 | 5 |
| 15 | TGME49_241140-t26_1-p1 | DEAD/DEAH box helicase domain-containing protein | 10 | 9 | 9 | 5 | 4 | 4 | 4 | 5 |
| 15 | TGME49_241850.P1728 | hypothetical protein | 10 | 10 | 8 | 6 | 5 | 5 | 4 | 5 |
| 15 | TGME49_242030-t26_1-p1 | conserved oligomeric Golgi complex subunit 7 | 10 | 8 | 9 | 6 | 4 | 5 | 4 | 5 |
| 15 | TGME49_242055.P1729 | DEAD/DEAH box helicase domain-containing protein | 10 | 8 | 7 | 4 | 5 | 4 | 4 | 5 |
| 14 | TGME49_242090.P1732 | zinc finger (CCCH type) motif-containing protein | 10 | 7 | 7 | 3 | 6 | 4 | 6 | 5 |
| 8 | TGME49_242240-t26_1-p1 | rhoptry kinase family protein ROP19A | 9 | 8 | 6 | 3 | 4 | 3 | 3 | 5 |
| 8 | TGME49_242260.P1733 | hypothetical protein | 10 | 10 | 5 | 4 | 4 | 3 | 3 | 5 |
| 13 | TGME49_242300-t26_1-p1 | dihydrodipicolinate synthase | 8 | 8 | 8 | 7 | 3 | 3 | 3 | 5 |
| 13 | TGME49_242370.P1737 | DUF4200 domain-containing protein | 9 | 8 | 8 | 6 | 3 | 3 | 3 | 5 |
| 15 | TGME49_242435-t26_1-p1 | hypothetical protein | 10 | 10 | 8 | 6 | 5 | 4 | 4 | 5 |
| 13 | TGME49_242460-t26_1-p1 | mitochondrial carrier superfamily protein | 9 | 8 | 7 | 6 | 3 | 3 | 5 | 5 |
| 13 | TGME49_242610-t26_1-p1 | WD domain, G-beta repeat-containing protein | 9 | 8 | 7 | 6 | 3 | 3 | 4 | 5 |
| 15 | TGME49_242640-t26_1-p1 | hypothetical protein | 10 | 10 | 10 | 6 | 5 | 4 | 4 | 5 |
| 14 | TGME49_242680-t26_1-p1 | ribosomal protein RPL24e | 7 | 6 | 6 | 3 | 6 | 5 | 4 | 5 |
| 8 | TGME49_242700.P1741 | WD repeat-containing protein | 10 | 7 | 8 | 4 | 4 | 5 | 3 | 5 |
| 14 | TGME49_242730.P1744 | guanylate kinase family protein | 8 | 7 | 7 | 4 | 6 | 3 | 5 | 5 |
| 8 | TGME49_242750.P1745 | hypothetical protein | 10 | 9 | 8 | 3 | 3 | 3 | 3 | 5 |
| 8 | TGME49_242790.P1747 | apical annuli protein AAP1 | 10 | 9 | 6 | 3 | 4 | 4 | 4 | 5 |
| 8 | TGME49_242810-t26_1-p1 | hypothetical protein | 9 | 7 | 7 | 4 | 4 | 3 | 3 | 5 |

| | | | | | | | | | | |
|---|---|---|---|---|---|---|---|---|---|---|
| 15 | TGME49_243200.P1755 | inner membrane complex protein IMC29 | 10 | 10 | 9 | 3 | 5 | 5 | 4 | 5 |
| 14 | TGME49_243210-t26_1-p1 | DUF862 domain-containing protein | 10 | 9 | 7 | 4 | 6 | 6 | 6 | 5 |
| 8 | TGME49_243265-t26_1-p1 | protamine P1 protein | 10 | 9 | 8 | 3 | 4 | 4 | 4 | 5 |
| 8 | TGME49_243298.P1756 | metacaspase MCA3, putative | 10 | 9 | 6 | 3 | 3 | 4 | 3 | 5 |
| 8 | TGME49_243310-t26_1-p1 | hypothetical protein | 9 | 9 | 6 | 4 | 4 | 3 | 3 | 5 |
| 8 | TGME49_243400.P1759 | PX domain-containing protein | 10 | 9 | 7 | 4 | 3 | 3 | 3 | 5 |
| 14 | TGME49_243410-t26_1-p1 | tetratricopeptide repeat-containing protein | 9 | 8 | 7 | 5 | 6 | 4 | 5 | 5 |
| 14 | TGME49_243450-t26_1-p1 | Ras-related protein Rab2BV, putative | 8 | 8 | 7 | 5 | 6 | 5 | 5 | 5 |
| 14 | TGME49_243600-t26_1-p1 | acetyltransferase, GNAT family protein | 9 | 8 | 6 | 3 | 5 | 3 | 5 | 5 |
| 19 | TGME49_243610.P1767 | DNA methyltransferase DNMTb | 10 | 9 | 9 | 3 | 5 | 5 | 5 | 5 |
| 14 | TGME49_243620-t26_1-p1 | pre-mRNA-splicing factor CWF7, putative | 8 | 8 | 7 | 3 | 5 | 5 | 5 | 5 |
| 13 | TGME49_243690.P1771 | hypothetical protein | 9 | 7 | 6 | 6 | 3 | 4 | 3 | 5 |
| 19 | TGME49_243740.P1773 | WD domain, G-beta repeat-containing protein | 10 | 8 | 9 | 5 | 3 | 7 | 3 | 5 |
| 13 | TGME49_243760-t26_1-p1 | hypothetical protein | 9 | 8 | 7 | 6 | 4 | 3 | 4 | 5 |
| 15 | TGME49_243780-t26_1-p1 | hypothetical protein | 10 | 9 | 8 | 4 | 5 | 3 | 3 | 5 |
| 13 | TGME49_243990-t26_1-p1 | Dullard family phosphatase domain-containing protein | 9 | 8 | 7 | 6 | 3 | 5 | 3 | 5 |
| 8 | TGME49_244090-t26_1-p1 | hypothetical protein | 10 | 8 | 7 | 4 | 3 | 3 | 3 | 5 |
| 13 | TGME49_244120-t26_1-p1 | zinc finger protein, putative | 10 | 8 | 6 | 5 | 4 | 3 | 3 | 5 |
| 14 | TGME49_244180.P1783 | microneme-like protein | 10 | 9 | 9 | 3 | 6 | 3 | 5 | 5 |
| 8 | TGME49_244230.P1784 | hypothetical protein | 8 | 8 | 7 | 4 | 4 | 3 | 3 | 5 |
| 8 | TGME49_244300-t26_1-p1 | hypothetical protein | 10 | 9 | 8 | 4 | 4 | 3 | 3 | 5 |
| 13 | TGME49_244335.P1787 | hypothetical protein | 9 | 6 | 8 | 5 | 3 | 3 | 3 | 5 |
| 14 | TGME49_244350-t26_1-p1 | DnaJ domain-containing protein | 9 | 8 | 8 | 5 | 5 | 3 | 6 | 5 |
| 13 | TGME49_244400-t26_1-p1 | hypothetical protein | 9 | 8 | 7 | 6 | 3 | 3 | 5 | 5 |
| 15 | TGME49_244460-t26_1-p1 | hypothetical protein | 10 | 10 | 8 | 4 | 4 | 4 | 3 | 5 |
| 8 | TGME49_244490-t26_1-p1 | insulysin, putative | 9 | 9 | 8 | 3 | 3 | 3 | 3 | 5 |
| 13 | TGME49_244540.P1793 | mitochondrial carrier superfamily protein | 9 | 8 | 7 | 5 | 3 | 4 | 4 | 5 |
| 13 | TGME49_244550-t26_1-p1 | hypothetical protein | 9 | 8 | 6 | 5 | 3 | 3 | 3 | 5 |
| 8 | TGME49_244640-t26_1-p1 | hypothetical protein | 10 | 9 | 5 | 4 | 4 | 4 | 4 | 5 |
| 8 | TGME49_244680-t26_1-p1 | hypothetical protein | 10 | 8 | 7 | 3 | 3 | 3 | 3 | 5 |
| 13 | TGME49_244910-t26_1-p1 | MIZ/SP-RING zinc finger domain-containing protein | 10 | 6 | 9 | 5 | 3 | 4 | 4 | 5 |
| 8 | TGME49_245440.P1804 | hypothetical protein | 10 | 10 | 6 | 4 | 3 | 4 | 4 | 5 |
| 8 | TGME49_245485-t26_1-p1 | microneme protein MIC9 | 8 | 8 | 6 | 3 | 3 | 3 | 3 | 5 |
| 13 | TGME49_245530.P1808 | hypothetical protein | 10 | 10 | 6 | 5 | 3 | 3 | 3 | 5 |
| 15 | TGME49_245580.P1811 | protein kinase, putative | 10 | 10 | 7 | 6 | 4 | 4 | 4 | 5 |
| 14 | TGME49_246020.P1820 | SprT domain-containing protein | 9 | 7 | 7 | 4 | 6 | 4 | 4 | 5 |
| 13 | TGME49_246160.P1825 | hypothetical protein | 10 | 9 | 7 | 5 | 3 | 4 | 4 | 5 |
| 13 | TGME49_246200.P1829 | zinc finger (CCCH type) motif-containing protein | 10 | 9 | 7 | 5 | 3 | 3 | 3 | 5 |
| 13 | TGME49_246490.P1831 | sphingolipid synthase | 7 | 8 | 8 | 7 | 3 | 3 | 4 | 5 |
| 13 | TGME49_246630-t26_1-p1 | DNA-directed RNA polymerase I RPA43 | 9 | 8 | 8 | 7 | 3 | 3 | 3 | 5 |
| 8 | TGME49_246660-t26_1-p1 | AP2 domain transcription factor AP2XII-3 | 10 | 8 | 6 | 3 | 3 | 4 | 3 | 5 |
| 19 | TGME49_246740-t26_1-p1 | dynactin subunit 2, putative | 10 | 9 | 9 | 5 | 4 | 7 | 7 | 5 |
| 19 | TGME49_246770.P1841 | hypothetical protein | 9 | 8 | 7 | 4 | 4 | 5 | 4 | 5 |
| 13 | TGME49_247010.P1844 | EF-hand domain-containing protein | 8 | 8 | 6 | 6 | 3 | 3 | 4 | 5 |
| 15 | TGME49_247050-t26_1-p1 | hypothetical protein | 10 | 10 | 9 | 6 | 4 | 4 | 3 | 5 |
| 15 | TGME49_247290-t26_1-p1 | hypothetical protein | 10 | 9 | 7 | 5 | 5 | 3 | 3 | 5 |
| 14 | TGME49_247360.P1850 | PAP2 superfamily protein | 8 | 8 | 7 | 3 | 6 | 4 | 5 | 5 |
| 14 | TGME49_247380.P1852 | cell division control protein CDC50, putative | 10 | 9 | 8 | 4 | 6 | 6 | 6 | 5 |

| | | | | | | | | | | |
|---|---|---|---|---|---|---|---|---|---|---|
| 8 | TGME49_247420-t26_1-p1 | hypothetical protein | 10 | 8 | 7 | 3 | 3 | 3 | 3 | 5 |
| 13 | TGME49_247485-t26_1-p1 | zinc finger, C3HC4 type (RING finger) domain-containing protein | 10 | 8 | 7 | 6 | 3 | 4 | 3 | 5 |
| 13 | TGME49_247590-t26_1-p1 | methyltransferase domain-containing protein | 10 | 7 | 7 | 5 | 3 | 3 | 3 | 5 |
| 13 | TGME49_247652-t26_1-p1 | peroxisome assembly protein PEX12, putative | 9 | 8 | 7 | 5 | 4 | 5 | 3 | 5 |
| 14 | TGME49_247660.P1862 | thioredoxin domain-containing protein | 10 | 9 | 9 | 3 | 8 | 4 | 4 | 5 |
| 15 | TGME49_247700-t26_1-p1 | AP2 domain transcription factor AP2XII-4 | 10 | 9 | 8 | 5 | 4 | 4 | 3 | 5 |
| 15 | TGME49_247730-t26_1-p1 | AP2 domain transcription factor AP2XII-5 | 10 | 10 | 9 | 5 | 5 | 4 | 4 | 5 |
| 8 | TGME49_247920-t26_1-p1 | methyltransferase domain-containing protein | 8 | 8 | 7 | 3 | 3 | 5 | 3 | 5 |
| 8 | TGME49_248240-t26_1-p1 | leucine rich repeat-containing protein | 10 | 8 | 8 | 3 | 4 | 3 | 3 | 5 |
| 19 | TGME49_248380-t26_1-p1 | hypothetical protein | 8 | 7 | 7 | 3 | 3 | 6 | 3 | 5 |
| 15 | TGME49_248440-t26_1-p1 | hypothetical protein | 10 | 10 | 8 | 6 | 4 | 5 | 4 | 5 |
| 15 | TGME49_248680-t26_1-p1 | hypothetical protein | 10 | 10 | 9 | 6 | 5 | 4 | 3 | 5 |
| 13 | TGME49_248710-t26_1-p1 | peptidase, putative | 10 | 9 | 8 | 6 | 3 | 3 | 3 | 5 |
| 15 | TGME49_248720-t26_1-p1 | hypothetical protein | 10 | 10 | 8 | 4 | 4 | 5 | 4 | 5 |
| 13 | TGME49_248740.P1903 | hypothetical protein | 10 | 8 | 6 | 5 | 4 | 4 | 5 | 5 |
| 14 | TGME49_248790-t26_1-p1 | hypothetical protein | 8 | 7 | 7 | 5 | 6 | 5 | 5 | 5 |
| 15 | TGME49_248840.P1908 | transporter facilitator protein TFP4 | 10 | 9 | 9 | 6 | 6 | 3 | 5 | 5 |
| 8 | TGME49_248900-t26_1-p1 | hypothetical protein | 10 | 9 | 7 | 4 | 3 | 3 | 3 | 5 |
| 14 | TGME49_249000.P1914 | regulator of chromosome condensation (RCC1) repeat-containing protein | 10 | 9 | 9 | 3 | 6 | 6 | 6 | 5 |
| 13 | TGME49_249010.P1915 | methyltransferase, putative | 9 | 8 | 6 | 5 | 4 | 3 | 4 | 5 |
| 8 | TGME49_249160.P1918 | UAA transporter family protein | 10 | 8 | 6 | 4 | 3 | 3 | 3 | 5 |
| 15 | TGME49_249215-t26_1-p1 | hypothetical protein | 10 | 10 | 9 | 4 | 4 | 3 | 4 | 5 |
| 19 | TGME49_249320.P1926 | flavodoxin domain-containing protein | 10 | 9 | 9 | 4 | 5 | 6 | 5 | 5 |
| 13 | TGME49_249340-t26_1-p1 | hypothetical protein | 10 | 8 | 7 | 6 | 3 | 4 | 4 | 5 |
| 15 | TGME49_249440.P1932 | apical cap protein AC12 | 10 | 9 | 8 | 4 | 4 | 4 | 4 | 5 |
| 13 | TGME49_249450-t26_1-p1 | hypothetical protein | 10 | 8 | 7 | 5 | 4 | 3 | 3 | 5 |
| 14 | TGME49_249460-t26_1-p1 | WD domain, G-beta repeat-containing protein | 10 | 9 | 8 | 4 | 6 | 5 | 5 | 5 |
| 8 | TGME49_249470-t26_1-p1 | Rhoptry kinase family protein, truncated (incomplete catalytic triad) | 9 | 7 | 7 | 3 | 3 | 3 | 3 | 5 |
| 15 | TGME49_249490.P1934 | hypothetical protein | 10 | 10 | 9 | 5 | 4 | 3 | 3 | 5 |
| 13 | TGME49_249510-t26_1-p1 | hypothetical protein | 8 | 8 | 7 | 7 | 3 | 4 | 4 | 5 |
| 15 | TGME49_249550-t26_1-p1 | hypothetical protein | 10 | 9 | 8 | 6 | 6 | 3 | 3 | 5 |
| 19 | TGME49_249660-t26_1-p1 | hypothetical protein | 10 | 9 | 8 | 3 | 3 | 4 | 6 | 5 |
| 14 | TGME49_249790-t26_1-p1 | hypothetical protein | 8 | 8 | 7 | 5 | 7 | 6 | 5 | 5 |
| 15 | TGME49_249880.P1949 | hypothetical protein | 10 | 10 | 9 | 5 | 5 | 4 | 4 | 5 |
| 13 | TGME49_249890-t26_1-p1 | hypothetical protein | 10 | 9 | 9 | 5 | 3 | 3 | 3 | 5 |
| 19 | TGME49_249920-t26_1-p1 | diphthine synthase, putative | 8 | 8 | 8 | 3 | 4 | 7 | 7 | 5 |
| 13 | TGME49_250060-t26_1-p1 | DNA-directed RNA polymerase I RPA12 | 8 | 7 | 7 | 6 | 3 | 3 | 5 | 5 |
| 8 | TGME49_250090-t26_1-p1 | hypothetical protein | 10 | 7 | 7 | 3 | 4 | 4 | 4 | 5 |
| 15 | TGME49_250700-t26_1-p1 | hypothetical protein | 10 | 10 | 9 | 6 | 6 | 5 | 3 | 5 |
| 13 | TGME49_250720-t26_1-p1 | hypothetical protein | 9 | 6 | 8 | 5 | 3 | 3 | 3 | 5 |
| 14 | TGME49_250740-t26_1-p1 | Yip1 domain-containing protein | 7 | 7 | 7 | 3 | 6 | 3 | 5 | 5 |
| 19 | TGME49_250750-t26_1-p1 | tRNA (uracil-5-)-methyltransferase | 10 | 10 | 9 | 4 | 5 | 6 | 4 | 5 |
| 13 | TGME49_250780-t26_1-p1 | Zn-finger in ubiquitin-hydrolases domain-containing protein | 10 | 8 | 6 | 5 | 4 | 4 | 4 | 5 |
| 15 | TGME49_250800-t26_1-p1 | AP2 domain transcription factor AP2XII-8 | 10 | 10 | 9 | 4 | 5 | 3 | 3 | 5 |
| 13 | TGME49_250820.P1965 | apical cap protein AC2 | 10 | 7 | 6 | 5 | 4 | 3 | 3 | 5 |
| 15 | TGME49_250850.P1966 | CMGC kinase, putative | 10 | 10 | 8 | 5 | 4 | 4 | 3 | 5 |

| | | | | | | | | | | |
|---|---|---|---|---|---|---|---|---|---|---|
| 14 | TGME49_250880.P1968 | adenosine kinase | 8 | 8 | 7 | 4 | 6 | 3 | 5 | 5 |
| 13 | TGME49_251450-t26_1-p1 | hypothetical protein | 10 | 9 | 6 | 5 | 3 | 4 | 4 | 5 |
| 13 | TGME49_251490-t26_1-p1 | ACR, COG2135 domain-containing protein | 9 | 7 | 6 | 6 | 4 | 3 | 3 | 5 |
| 14 | TGME49_251570-t26_1-p1 | CAAX amino terminal protease family protein | 9 | 8 | 7 | 3 | 6 | 5 | 5 | 5 |
| 13 | TGME49_251610-t26_1-p1 | hypothetical protein | 10 | 7 | 8 | 6 | 4 | 4 | 4 | 5 |
| 8 | TGME49_251670-t26_1-p1 | werner helicase interacting protein 1, putative | 10 | 7 | 9 | 4 | 3 | 3 | 3 | 5 |
| 8 | TGME49_251730-t26_1-p1 | conserved oligomeric Golgi complex subunit 5 | 10 | 10 | 9 | 4 | 3 | 3 | 3 | 5 |
| 14 | TGME49_251810-t26_1-p1 | translation initiation factor eIF-5A, putative | 7 | 7 | 7 | 3 | 6 | 3 | 6 | 5 |
| 15 | TGME49_251890.P1985 | hypothetical protein | 10 | 10 | 9 | 6 | 4 | 4 | 3 | 5 |
| 15 | TGME49_251920.P1986 | hypothetical protein | 10 | 9 | 8 | 6 | 5 | 4 | 3 | 5 |
| 15 | TGME49_252250-t26_1-p1 | ATPase, AAA family protein | 10 | 9 | 8 | 5 | 4 | 4 | 3 | 5 |
| 8 | TGME49_252370.P1994 | AP2 domain transcription factor AP2III-1 | 10 | 8 | 6 | 3 | 3 | 5 | 4 | 5 |
| 8 | TGME49_252500.P2000 | polo kinase | 10 | 8 | 6 | 3 | 3 | 3 | 3 | 5 |
| 8 | TGME49_252870.P2004 | hypothetical protein | 10 | 9 | 6 | 3 | 4 | 3 | 3 | 5 |
| 13 | TGME49_253000.P2005 | ELMO/CED-12 family protein | 9 | 8 | 7 | 5 | 4 | 3 | 4 | 5 |
| 15 | TGME49_253040.P2007 | actin-related protein ARP4A | 10 | 9 | 8 | 5 | 5 | 5 | 4 | 5 |
| 15 | TGME49_253060.P2008 | WD domain, G-beta repeat-containing protein | 10 | 9 | 8 | 6 | 5 | 4 | 4 | 5 |
| 14 | TGME49_253320-t26_1-p1 | hypothetical protein | 9 | 7 | 7 | 4 | 6 | 5 | 5 | 5 |
| 15 | TGME49_253380-t26_1-p1 | AP2 domain transcription factor AP2III-2 | 10 | 10 | 7 | 4 | 5 | 4 | 3 | 5 |
| 13 | TGME49_253480-t26_1-p1 | topoisomerase VIA, putative | 8 | 8 | 7 | 5 | 3 | 3 | 3 | 5 |
| 15 | TGME49_253490.P2024 | hypothetical protein | 10 | 8 | 8 | 5 | 5 | 5 | 4 | 5 |
| 19 | TGME49_253520-t26_1-p1 | U3 small nucleolar RNA-associated protein 6 protein | 10 | 10 | 9 | 3 | 3 | 5 | 4 | 5 |
| 8 | TGME49_253540.P2026 | hypothetical protein | 10 | 8 | 6 | 4 | 3 | 4 | 3 | 5 |
| 8 | TGME49_253560-t26_1-p1 | hypothetical protein | 10 | 10 | 5 | 3 | 3 | 3 | 3 | 5 |
| 14 | TGME49_253600.P2030 | preconoidal ring protein PCR6 | 10 | 9 | 8 | 4 | 6 | 6 | 6 | 5 |
| 15 | TGME49_253615.P2031 | hypothetical protein | 10 | 10 | 10 | 4 | 4 | 4 | 3 | 5 |
| 8 | TGME49_253740-t26_1-p1 | hypothetical protein | 10 | 8 | 6 | 4 | 4 | 3 | 3 | 5 |
| 8 | TGME49_253780.P2035 | GTP cyclohydrolase I, putative | 9 | 8 | 6 | 4 | 4 | 5 | 3 | 5 |
| 14 | TGME49_253800-t26_1-p1 | ribosomal protein L15, putative | 10 | 8 | 8 | 5 | 8 | 4 | 6 | 5 |
| 8 | TGME49_253810-t26_1-p1 | hypothetical protein | 10 | 9 | 7 | 4 | 4 | 4 | 3 | 5 |
| 8 | TGME49_253870.P2037 | hypothetical protein | 10 | 10 | 5 | 4 | 3 | 3 | 3 | 5 |
| 14 | TGME49_253950-t26_1-p1 | protein fam50a, putative | 9 | 9 | 8 | 3 | 8 | 5 | 7 | 5 |
| 8 | TGME49_254090.P2046 | hypothetical protein | 10 | 8 | 7 | 3 | 3 | 4 | 4 | 5 |
| 13 | TGME49_254150-t26_1-p1 | hypothetical protein | 10 | 7 | 6 | 5 | 3 | 3 | 3 | 5 |
| 8 | TGME49_254210-t26_1-p1 | RNA recognition motif-containing protein | 10 | 7 | 7 | 4 | 4 | 4 | 3 | 5 |
| 8 | TGME49_254290-t26_1-p1 | hypothetical protein | 10 | 9 | 6 | 4 | 3 | 3 | 4 | 5 |
| 13 | TGME49_254330-t26_1-p1 | lipase | 10 | 9 | 7 | 6 | 3 | 3 | 4 | 5 |
| 14 | TGME49_254410.P2056 | protein phosphatase PPM10, putative | 9 | 8 | 7 | 4 | 7 | 3 | 4 | 5 |
| 19 | TGME49_254420-t26_1-p1 | phospholipase, patatin family protein | 10 | 8 | 8 | 3 | 3 | 4 | 6 | 5 |
| 13 | TGME49_254470.P2059 | Myc-regulation protein MYR1 | 10 | 8 | 7 | 5 | 3 | 3 | 3 | 5 |
| 15 | TGME49_254790-t26_1-p1 | hypothetical protein | 10 | 7 | 10 | 5 | 4 | 3 | 3 | 5 |
| 19 | TGME49_254830.P2076 | hypothetical protein | 10 | 8 | 7 | 4 | 4 | 5 | 4 | 5 |
| 13 | TGME49_254835-t26_1-p1 | RNA recognition motif-containing protein | 10 | 7 | 7 | 5 | 3 | 4 | 4 | 5 |
| 19 | TGME49_254840.P2077 | tetratricopeptide repeat-containing protein | 9 | 8 | 7 | 3 | 3 | 5 | 4 | 5 |
| 8 | TGME49_254880-t26_1-p1 | Alpha-galactosidase | 10 | 9 | 7 | 3 | 3 | 3 | 4 | 5 |
| 13 | TGME49_255220.P2090 | AP2 domain transcription factor AP2VIIb-3 | 10 | 10 | 5 | 5 | 3 | 3 | 3 | 5 |
| 15 | TGME49_255300-t26_1-p1 | hypothetical protein | 10 | 10 | 8 | 5 | 4 | 3 | 3 | 5 |
| 13 | TGME49_255310-t26_1-p1 | zinc finger (CCCH type) motif-containing protein | 10 | 9 | 9 | 6 | 3 | 3 | 3 | 5 |
| 14 | TGME49_255320.P2094 | mRNA turnover 4 (MRT4) family protein | 7 | 7 | 7 | 3 | 6 | 6 | 6 | 5 |

| | | | | | | | | | | |
|---|---|---|---|---|---|---|---|---|---|---|
| 15 | TGME49_255390.P2097 | HEAT repeat-containing protein | 10 | 10 | 10 | 3 | 3 | 4 | 4 | 5 |
| 15 | TGME49_255430.P2100 | Rad9 protein | 10 | 9 | 8 | 4 | 6 | 3 | 4 | 5 |
| 14 | TGME49_255650.P2110 | palmitoyltransferase DHHC8 | 9 | 8 | 8 | 3 | 7 | 6 | 4 | 5 |
| 19 | TGME49_255710-t26_1-p1 | ULK kinase | 9 | 7 | 8 | 3 | 3 | 6 | 3 | 5 |
| 8 | TGME49_255730.P2117 | hypothetical protein | 9 | 9 | 8 | 3 | 3 | 3 | 3 | 5 |
| 13 | TGME49_255895.P2121 | hypothetical protein | 9 | 8 | 6 | 5 | 4 | 3 | 4 | 5 |
| 19 | TGME49_255980.P2126 | hypothetical protein | 10 | 9 | 9 | 3 | 3 | 5 | 6 | 5 |
| 19 | TGME49_256040-t26_1-p1 | PA14 domain-containing protein | 10 | 8 | 6 | 3 | 5 | 8 | 6 | 5 |
| 8 | TGME49_256080.P2132 | glycosyltransferase, group 2 family protein | 9 | 8 | 7 | 3 | 3 | 3 | 3 | 5 |
| 15 | TGME49_256100.P2134 | tetratricopeptide repeat-containing protein | 10 | 10 | 8 | 3 | 5 | 4 | 4 | 5 |
| 14 | TGME49_256780.P2136 | PPPDE peptidase, putative | 9 | 8 | 7 | 5 | 6 | 5 | 5 | 5 |
| 14 | TGME49_256820.P2140 | zinc finger (CCCH type) motif-containing protein | 9 | 9 | 8 | 5 | 7 | 6 | 5 | 5 |
| 15 | TGME49_256840.P2142 | hypothetical protein | 10 | 9 | 9 | 5 | 6 | 6 | 3 | 5 |
| 13 | TGME49_256850.P2143 | hypothetical protein | 9 | 9 | 8 | 6 | 3 | 3 | 3 | 5 |
| 19 | TGME49_256910.P2147 | vacuolar protein sorting-associated protein 31, putative | 10 | 10 | 9 | 4 | 3 | 5 | 4 | 5 |
| 13 | TGME49_257030-t26_1-p1 | hypothetical protein | 9 | 6 | 8 | 7 | 3 | 4 | 3 | 5 |
| 13 | TGME49_257130.P2159 | zinc finger (CCCH type) motif-containing protein | 10 | 9 | 8 | 6 | 3 | 3 | 5 | 5 |
| 13 | TGME49_257170.P2161 | hypothetical protein | 9 | 8 | 7 | 5 | 4 | 3 | 3 | 5 |
| 13 | TGME49_257360.P2168 | hypothetical protein | 10 | 10 | 8 | 5 | 3 | 3 | 3 | 5 |
| 13 | TGME49_257370.P2169 | preconoidal region protein PCR2 | 10 | 8 | 6 | 6 | 3 | 3 | 3 | 5 |
| 13 | TGME49_257430.P2173 | hypothetical protein | 10 | 9 | 7 | 6 | 4 | 3 | 3 | 5 |
| 15 | TGME49_257500.P2176 | hypothetical protein | 10 | 9 | 8 | 5 | 5 | 4 | 3 | 5 |
| 8 | TGME49_257540-t26_1-p1 | hypothetical protein | 9 | 8 | 7 | 4 | 4 | 3 | 3 | 5 |
| 15 | TGME49_257550.P2178 | hypothetical protein | 10 | 10 | 7 | 6 | 5 | 3 | 3 | 5 |
| 13 | TGME49_257640.P2180 | hypothetical protein | 9 | 8 | 6 | 5 | 3 | 4 | 4 | 5 |
| 13 | TGME49_257690-t26_1-p1 | hypothetical protein | 9 | 8 | 7 | 5 | 3 | 3 | 3 | 5 |
| 8 | TGME49_257780-t26_1-p1 | hypothetical protein | 10 | 9 | 6 | 4 | 4 | 3 | 3 | 5 |
| 14 | TGME49_257800.P2190 | polynucleotide adenylyltransferase | 10 | 8 | 8 | 5 | 6 | 6 | 5 | 5 |
| 15 | TGME49_257945.P2193 | 3'5'-cyclic nucleotide phosphodiesterase PDE17 | 10 | 9 | 8 | 6 | 6 | 3 | 3 | 5 |
| 8 | TGME49_258080.P2204 | golgi localizing protein GLP1 | 10 | 8 | 7 | 4 | 4 | 3 | 3 | 5 |
| 13 | TGME49_258120.P2208 | Pex2 / Pex12 amino terminal region protein | 9 | 8 | 7 | 5 | 3 | 4 | 4 | 5 |
| 13 | TGME49_258180.P2211 | ELMO domain-containing protein | 9 | 9 | 8 | 6 | 3 | 4 | 4 | 5 |
| 8 | TGME49_258200.P2212 | hypothetical protein | 10 | 8 | 6 | 3 | 3 | 3 | 3 | 5 |
| 14 | TGME49_258350.P2215 | START domain-containing protein | 10 | 8 | 8 | 3 | 6 | 4 | 6 | 5 |
| 13 | TGME49_258370.P2217 | rhoptry kinase family protein ROP28 | 10 | 8 | 7 | 6 | 3 | 3 | 3 | 5 |
| 13 | TGME49_258450.P2221 | apicomplexan kinetochore protein AKIT8, putative | 10 | 9 | 6 | 5 | 3 | 4 | 3 | 5 |
| 13 | TGME49_258480-t26_1-p1 | BRCT domain-containing protein | 10 | 9 | 7 | 6 | 4 | 4 | 4 | 5 |
| 8 | TGME49_258540.P2227 | phosphoglycerate mutase family protein | 10 | 9 | 7 | 4 | 3 | 3 | 3 | 5 |
| 13 | TGME49_258580-t26_1-p1 | rhoptry protein ROP17 | 9 | 8 | 7 | 7 | 3 | 3 | 3 | 5 |
| 15 | TGME49_258826.P2242 | hypothetical protein | 10 | 9 | 9 | 6 | 4 | 4 | 4 | 5 |
| 8 | TGME49_258870.P2246 | cyst wall protein CST7 | 10 | 7 | 7 | 3 | 3 | 3 | 3 | 5 |
| 13 | TGME49_258880.P2247 | axonemal dynein light chain p33, putative | 8 | 8 | 7 | 7 | 3 | 3 | 4 | 5 |
| 13 | TGME49_258900.P2248 | hypothetical protein | 9 | 7 | 7 | 6 | 4 | 3 | 3 | 5 |
| 14 | TGME49_259130.P2263 | 4-hydroxybenzoate polyprenyl transferase | 9 | 8 | 8 | 5 | 5 | 5 | 5 | 5 |
| 13 | TGME49_259160.P2265 | hypothetical protein | 8 | 8 | 7 | 6 | 4 | 3 | 3 | 5 |
| 8 | TGME49_259520.P2276 | spindle and kinetochore-associated protein SKA2 | 9 | 8 | 7 | 4 | 3 | 3 | 3 | 5 |
| 13 | TGME49_259870.P2293 | hypothetical protein | 10 | 9 | 6 | 5 | 3 | 4 | 3 | 5 |
| 15 | TGME49_259890.P2295 | Kazal-type serine protease inhibitor domain-containing protein | 10 | 10 | 10 | 4 | 8 | 3 | 3 | 5 |

| | | | | | | | | | | |
|---|---|---|---|---|---|---|---|---|---|---|
| 13 | TGME49_259950.P2297 | carbonate dehydratase, eukaryotic-type domain-containing protein | 10 | 9 | 7 | 7 | 4 | 4 | 4 | 5 |
| 8 | TGME49_259965.P2299 | Bardet-Biedl syndrome 5, putative | 10 | 7 | 8 | 3 | 3 | 3 | 3 | 5 |
| 8 | TGME49_260000-t26_1-p1 | hypothetical protein | 10 | 9 | 8 | 3 | 3 | 3 | 3 | 5 |
| 14 | TGME49_260250-t26_1-p1 | cyclin domain protein, cyclin H family protein | 9 | 8 | 8 | 3 | 5 | 5 | 5 | 5 |
| 19 | TGME49_260270.P2314 | HEAT repeat-containing protein | 10 | 10 | 8 | 3 | 3 | 5 | 4 | 5 |
| 14 | TGME49_260420.P2321 | kinetochore protein NDC80 | 10 | 9 | 9 | 4 | 8 | 6 | 5 | 5 |
| 14 | TGME49_260430.P2322 | DHH phosphoesterase, putative | 9 | 7 | 7 | 5 | 6 | 5 | 5 | 5 |
| 8 | TGME49_260490.P2326 | hypothetical protein | 10 | 10 | 6 | 3 | 4 | 3 | 4 | 5 |
| 13 | TGME49_260520.P2328 | dense granule protein GRA53 | 10 | 8 | 7 | 5 | 4 | 3 | 3 | 5 |
| 13 | TGME49_260550.P2330 | acetyltransferase, GNAT family protein | 10 | 9 | 6 | 6 | 4 | 3 | 4 | 5 |
| 14 | TGME49_260660.P2337 | 50S ribosomal protein L23, putative | 8 | 8 | 7 | 4 | 7 | 6 | 6 | 5 |
| 13 | TGME49_260830.P2344 | AMP-activated protein kinase, putative | 10 | 8 | 6 | 5 | 3 | 3 | 3 | 5 |
| 15 | TGME49_260840-t26_1-p1 | tetratricopeptide repeat-containing protein | 10 | 10 | 9 | 6 | 5 | 4 | 4 | 5 |
| 14 | TGME49_261030.P2348 | pyridine nucleotide-disulfide oxidoreductase domain-containing protein | 10 | 9 | 9 | 4 | 8 | 5 | 5 | 5 |
| 15 | TGME49_261430.P2361 | hypothetical protein | 10 | 9 | 8 | 6 | 4 | 4 | 3 | 5 |
| 13 | TGME49_261790.P2383 | hypothetical protein | 10 | 8 | 7 | 5 | 4 | 3 | 3 | 5 |
| 15 | TGME49_261955.P2386 | hypothetical protein | 10 | 8 | 7 | 5 | 5 | 4 | 3 | 5 |
| 15 | TGME49_261960.P2387 | zinc finger, C2H2 type domain-containing protein | 10 | 9 | 7 | 5 | 5 | 4 | 3 | 5 |
| 14 | TGME49_261980.P2388 | gorasp2-prov protein | 9 | 8 | 8 | 5 | 6 | 4 | 5 | 5 |
| 13 | TGME49_262030-t26_1-p1 | ALG6, ALG8 glycosyltransferase family protein | 10 | 8 | 7 | 5 | 3 | 3 | 3 | 5 |
| 8 | TGME49_262060.P2392 | hypothetical protein | 9 | 9 | 8 | 4 | 3 | 3 | 3 | 5 |
| 13 | TGME49_262110-t26_1-p1 | hypothetical protein | 9 | 9 | 7 | 5 | 3 | 3 | 3 | 5 |
| 8 | TGME49_262440.P2404 | hypothetical protein | 9 | 7 | 8 | 3 | 3 | 5 | 3 | 5 |
| 15 | TGME49_262450.P2405 | dymeclin, putative | 10 | 10 | 7 | 6 | 6 | 4 | 4 | 5 |
| 15 | TGME49_262460.P2406 | PX domain-containing protein | 10 | 10 | 7 | 5 | 4 | 4 | 4 | 5 |
| 13 | TGME49_262510.P2409 | GTP-binding protein engB, putative | 9 | 8 | 7 | 7 | 3 | 4 | 3 | 5 |
| 15 | TGME49_262540.P2412 | protein kinase, putative | 10 | 9 | 8 | 6 | 5 | 3 | 3 | 5 |
| 8 | TGME49_262550.P2413 | hypothetical protein | 10 | 8 | 6 | 3 | 3 | 3 | 3 | 5 |
| 15 | TGME49_262610.P2414 | hypothetical protein | 10 | 9 | 8 | 6 | 5 | 3 | 3 | 5 |
| 14 | TGME49_262620-t26_1-p1 | RNA recognition motif-containing protein | 8 | 8 | 7 | 5 | 5 | 5 | 5 | 5 |
| 13 | TGME49_262660.P2418 | transporter, major facilitator family protein | 10 | 9 | 8 | 6 | 3 | 3 | 3 | 5 |
| 15 | TGME49_262680.P2420 | hypothetical protein | 10 | 9 | 9 | 4 | 4 | 4 | 3 | 5 |
| 14 | TGME49_262750.P2425 | histone lysine methyltransferase, SET, putative | 9 | 8 | 6 | 3 | 5 | 5 | 5 | 5 |
| 13 | TGME49_262780.P2427 | FHA domain-containing protein | 10 | 10 | 8 | 5 | 3 | 3 | 3 | 5 |
| 13 | TGME49_262825-t26_1-p1 | peptidase family c50 protein | 10 | 8 | 5 | 6 | 3 | 3 | 3 | 5 |
| 14 | TGME49_262860-t26_1-p1 | ADP-ribosylation factor family protein 1, putative | 7 | 7 | 7 | 3 | 6 | 6 | 5 | 5 |
| 14 | TGME49_262960-t26_1-p1 | U1 snRNP-associated protein Usp106, putative | 9 | 8 | 8 | 3 | 6 | 5 | 6 | 5 |
| 13 | TGME49_262990.P2436 | hypothetical protein | 10 | 9 | 6 | 5 | 4 | 3 | 3 | 5 |
| 15 | TGME49_263200.P2453 | hypothetical protein | 10 | 9 | 8 | 5 | 6 | 5 | 4 | 5 |
| 14 | TGME49_263220.P2455 | rhoptry kinase family protein ROP21 | 9 | 9 | 8 | 3 | 8 | 6 | 6 | 5 |
| 13 | TGME49_263320-t26_1-p1 | hypothetical protein | 9 | 7 | 7 | 6 | 4 | 4 | 3 | 5 |
| 13 | TGME49_263340.P2462 | RNA methyltransferase, TrmH family protein | 9 | 9 | 8 | 5 | 3 | 3 | 3 | 5 |
| 15 | TGME49_263390-t26_1-p1 | hypothetical protein | 9 | 9 | 8 | 5 | 5 | 5 | 3 | 5 |
| 15 | TGME49_263440.P2469 | hypothetical protein | 9 | 9 | 7 | 5 | 5 | 5 | 4 | 5 |
| 8 | TGME49_263450.P2470 | hypothetical protein | 10 | 8 | 7 | 4 | 4 | 4 | 4 | 5 |
| 14 | TGME49_263520-t26_1-p1 | microtubule associated protein SPM1 | 9 | 8 | 7 | 3 | 7 | 6 | 6 | 5 |
| 19 | TGME49_263580.P2478 | bromodomain-containing protein BDP1 | 10 | 9 | 9 | 3 | 3 | 8 | 4 | 5 |
| 13 | TGME49_263690-t26_1-p1 | hypothetical protein | 10 | 7 | 7 | 5 | 3 | 3 | 3 | 5 |

| | | | | | | | | | | |
|---|---|---|---|---|---|---|---|---|---|---|
| 8 | TGME49_263760.P2492 | hypothetical protein | 10 | 8 | 7 | 3 | 3 | 3 | 3 | 5 |
| 14 | TGME49_263800.P2494 | RNase P subunit p30, putative | 9 | 7 | 7 | 3 | 5 | 4 | 4 | 5 |
| 8 | TGME49_264140-t26_1-p1 | hypothetical protein | 10 | 10 | 6 | 4 | 3 | 3 | 3 | 5 |
| 8 | TGME49_264170.P2509 | hypothetical protein | 10 | 10 | 9 | 3 | 3 | 3 | 3 | 5 |
| 15 | TGME49_264190.P2511 | pseudouridine synthase, putative | 10 | 10 | 8 | 6 | 4 | 4 | 4 | 5 |
| 13 | TGME49_264200.P2512 | hypothetical protein | 10 | 8 | 7 | 6 | 4 | 3 | 3 | 5 |
| 8 | TGME49_264210.P2513 | hypothetical protein | 10 | 9 | 6 | 4 | 3 | 3 | 3 | 5 |
| 14 | TGME49_264610.P2517 | RNA recognition motif-containing protein | 9 | 8 | 7 | 4 | 5 | 4 | 4 | 5 |
| 15 | TGME49_264790.P2524 | hypothetical protein | 10 | 10 | 7 | 5 | 5 | 5 | 5 | 5 |
| 15 | TGME49_264890-t26_1-p1 | hypothetical protein | 10 | 9 | 8 | 4 | 4 | 4 | 4 | 5 |
| 13 | TGME49_264940-t26_1-p1 | kelch repeat-containing protein | 10 | 8 | 6 | 5 | 4 | 4 | 3 | 5 |
| 14 | TGME49_264970-t26_1-p1 | hypothetical protein | 10 | 7 | 4 | 4 | 6 | 5 | 5 | 5 |
| 13 | TGME49_265000-t26_1-p1 | phosphotransferase enzyme family protein | 10 | 8 | 6 | 5 | 3 | 3 | 3 | 5 |
| 8 | TGME49_265070-t26_1-p1 | hypothetical protein | 9 | 8 | 7 | 3 | 3 | 3 | 3 | 5 |
| 13 | TGME49_265150-t26_1-p1 | ankyrin repeat-containing protein | 9 | 8 | 8 | 6 | 3 | 3 | 3 | 5 |
| 13 | TGME49_265170-t26_1-p1 | hypothetical protein | 9 | 8 | 7 | 5 | 4 | 3 | 4 | 5 |
| 14 | TGME49_265180.P2534 | lasso maintenance factor LMF1 | 9 | 8 | 7 | 5 | 5 | 4 | 5 | 5 |
| 14 | TGME49_265230-t26_1-p1 | electron transfer flavoprotein-ubiquinone oxidoreductase | 10 | 9 | 9 | 3 | 6 | 5 | 6 | 5 |
| 13 | TGME49_265240-t26_1-p1 | hypothetical protein | 10 | 8 | 6 | 5 | 4 | 5 | 3 | 5 |
| 15 | TGME49_265250-t26_1-p1 | RNA recognition motif-containing protein | 10 | 10 | 9 | 5 | 4 | 4 | 3 | 5 |
| 13 | TGME49_265280-t26_1-p1 | hypothetical protein | 10 | 9 | 7 | 7 | 4 | 4 | 3 | 5 |
| 8 | TGME49_265400-t26_1-p1 | hypothetical protein | 10 | 8 | 6 | 3 | 4 | 4 | 3 | 5 |
| 13 | TGME49_265460-t26_1-p1 | hypothetical protein | 9 | 8 | 7 | 5 | 3 | 3 | 3 | 5 |
| 8 | TGME49_265820.P2547 | Las1 family protein | 10 | 8 | 6 | 3 | 4 | 4 | 4 | 5 |
| 15 | TGME49_266010.P2553 | phosphatidylinositol 3- and 4-kinase | 10 | 9 | 7 | 6 | 5 | 5 | 3 | 5 |
| 13 | TGME49_266080.P2557 | apical cap protein AC11 | 10 | 8 | 6 | 5 | 3 | 3 | 3 | 5 |
| 8 | TGME49_266100.P2559 | rhoptry kinase family protein ROP41 | 9 | 9 | 7 | 3 | 3 | 3 | 4 | 5 |
| 14 | TGME49_266120-t26_1-p1 | thioredoxin-dependent peroxidase TPX1/2 | 8 | 7 | 6 | 3 | 6 | 5 | 4 | 5 |
| 14 | TGME49_266270-t26_1-p1 | short-chain-acyl-CoA dehydrogenase | 9 | 8 | 8 | 4 | 8 | 7 | 4 | 5 |
| 13 | TGME49_266310-t26_1-p1 | Alpha-1,3-mannosyl-glycoprotein 2-beta-N-acetylglucosaminyltransferase | 9 | 8 | 8 | 5 | 3 | 3 | 3 | 5 |
| 15 | TGME49_266320.P2562 | glycosyltransferase family protein | 10 | 9 | 8 | 6 | 4 | 4 | 4 | 5 |
| 8 | TGME49_266390.P2566 | DNA mismatch repair protein, C-terminal domain-containing protein | 10 | 8 | 6 | 4 | 5 | 5 | 3 | 5 |
| 13 | TGME49_266450-t26_1-p1 | lysine decarboxylase family protein | 9 | 8 | 8 | 7 | 3 | 3 | 3 | 5 |
| 13 | TGME49_266690-t26_1-p1 | hypothetical protein | 10 | 10 | 8 | 5 | 3 | 3 | 3 | 5 |
| 14 | TGME49_266760.P2580 | isocitrate dehydrogenase 1 | 10 | 9 | 9 | 3 | 9 | 3 | 3 | 5 |
| 8 | TGME49_266770-t26_1-p1 | cpw-wpc domain-containing protein | 9 | 9 | 6 | 3 | 3 | 3 | 6 | 5 |
| 19 | TGME49_266830.P2584 | TBC domain-containing protein TBC17 | 10 | 10 | 7 | 3 | 3 | 5 | 5 | 5 |
| 14 | TGME49_267020.P2592 | tetratricopeptide repeat-containing protein | 9 | 8 | 8 | 5 | 5 | 4 | 5 | 5 |
| 19 | TGME49_267100.P2596 | protein phosphatase PPM2B | 10 | 9 | 7 | 5 | 3 | 7 | 6 | 5 |
| 8 | TGME49_267280-t26_1-p1 | hypothetical protein | 10 | 8 | 6 | 4 | 4 | 4 | 3 | 5 |
| 15 | TGME49_267460.P2601 | AP2 domain transcription factor AP2IX-1 | 10 | 10 | 9 | 5 | 3 | 4 | 4 | 5 |
| 15 | TGME49_267560.P2606 | folate-binding protein YgfZ protein | 10 | 9 | 8 | 5 | 4 | 4 | 4 | 5 |
| 8 | TGME49_267570.P2607 | hypothetical protein | 10 | 8 | 7 | 4 | 4 | 3 | 3 | 5 |
| 8 | TGME49_267590-t26_1-p1 | hypothetical protein | 10 | 7 | 8 | 4 | 4 | 4 | 3 | 5 |
| 8 | TGME49_267650.P2610 | hypothetical protein | 10 | 10 | 9 | 3 | 3 | 3 | 3 | 5 |
| 13 | TGME49_267670-t26_1-p1 | hypothetical protein | 10 | 9 | 6 | 6 | 3 | 3 | 3 | 5 |
| 8 | TGME49_267690-t26_1-p1 | hypothetical protein | 10 | 8 | 8 | 4 | 4 | 4 | 3 | 5 |

| | | | | | | | | | | |
|---|---|---|---|---|---|---|---|---|---|---|
| 13 | TGME49_267730.P2613 | GDP-fucose transporter NST2 | 9 | 8 | 8 | 7 | 3 | 3 | 4 | 5 |
| 14 | TGME49_267820.P2617 | electron transfer flavoprotein subunit beta, putative | 8 | 7 | 7 | 3 | 5 | 5 | 5 | 5 |
| 15 | TGME49_267875.P2619 | NBP2b protein, putative | 10 | 9 | 9 | 5 | 5 | 4 | 5 | 5 |
| 8 | TGME49_268020.P2622 | transporter, major facilitator family protein | 10 | 9 | 7 | 3 | 4 | 5 | 3 | 5 |
| 8 | TGME49_268240.P2627 | hypothetical protein | 10 | 9 | 6 | 4 | 3 | 4 | 3 | 5 |
| 13 | TGME49_268260.P2628 | hypothetical protein | 10 | 8 | 8 | 5 | 4 | 4 | 3 | 5 |
| 8 | TGME49_268320.P2632 | hypothetical protein | 10 | 9 | 8 | 4 | 3 | 3 | 3 | 5 |
| 19 | TGME49_268380.P2635 | RNA recognition motif-containing protein | 10 | 9 | 7 | 5 | 3 | 6 | 5 | 5 |
| 19 | TGME49_268400.P2636 | hypothetical protein | 9 | 8 | 8 | 4 | 5 | 6 | 5 | 5 |
| 15 | TGME49_268430.P2637 | hypothetical protein | 10 | 9 | 7 | 5 | 4 | 4 | 4 | 5 |
| 13 | TGME49_268570.P2639 | zinc finger (CCCH type) motif-containing protein | 9 | 8 | 8 | 7 | 3 | 4 | 3 | 5 |
| 13 | TGME49_268820.P2650 | hypothetical protein | 10 | 6 | 6 | 6 | 3 | 4 | 4 | 5 |
| 13 | TGME49_268880.P2656 | hypothetical protein | 9 | 8 | 7 | 5 | 3 | 3 | 3 | 5 |
| 13 | TGME49_268920-t26_1-p1 | inositol polyphosphate kinase | 9 | 8 | 6 | 6 | 3 | 4 | 4 | 5 |
| 13 | TGME49_268930.P2659 | hypothetical protein | 9 | 8 | 6 | 5 | 3 | 3 | 3 | 5 |
| 13 | TGME49_268940.P2660 | hypothetical protein | 6 | 8 | 6 | 8 | 3 | 3 | 3 | 5 |
| 15 | TGME49_268990.P2661 | hypothetical protein | 10 | 10 | 9 | 6 | 4 | 4 | 4 | 5 |
| 15 | TGME49_269010-t26_1-p1 | AP2 domain transcription factor AP2VIII-7 | 10 | 9 | 8 | 5 | 5 | 4 | 4 | 5 |
| 15 | TGME49_269020.P2663 | hypothetical protein | 9 | 9 | 8 | 6 | 4 | 4 | 4 | 5 |
| 13 | TGME49_269060-t26_1-p1 | inositol polyphosphate kinase, putative | 10 | 9 | 7 | 6 | 3 | 3 | 3 | 5 |
| 14 | TGME49_269150.P2667 | palmitoyltransferase DHHC9 | 9 | 8 | 8 | 3 | 6 | 5 | 4 | 5 |
| 13 | TGME49_269260.P2675 | suppressor of calcium-dependent egress SCE1 | 10 | 7 | 8 | 7 | 3 | 3 | 3 | 5 |
| 19 | TGME49_269330-t26_1-p1 | hypothetical protein | 8 | 7 | 7 | 3 | 4 | 6 | 6 | 5 |
| 8 | TGME49_269380-t26_1-p1 | hypothetical protein | 10 | 9 | 6 | 3 | 3 | 3 | 4 | 5 |
| 19 | TGME49_269390-t26_1-p1 | CRAL/TRIO domain-containing protein | 10 | 9 | 8 | 3 | 3 | 6 | 4 | 5 |
| 13 | TGME49_269420-t26_1-p1 | hypothetical protein | 10 | 9 | 8 | 5 | 4 | 3 | 3 | 5 |
| 14 | TGME49_269450.P2684 | hypothetical protein | 8 | 7 | 6 | 3 | 6 | 6 | 5 | 5 |
| 19 | TGME49_269582.P2687 | 14-3-3 protein | 8 | 7 | 7 | 5 | 3 | 5 | 5 | 5 |
| 8 | TGME49_269670.P2694 | hypothetical protein | 10 | 8 | 6 | 3 | 3 | 3 | 3 | 5 |
| 8 | TGME49_269680-t26_1-p1 | acyl-CoA carboxyltransferase beta chain, putative | 9 | 9 | 9 | 3 | 4 | 3 | 3 | 5 |
| 8 | TGME49_269720.P2698 | hypothetical protein | 10 | 10 | 8 | 3 | 3 | 3 | 3 | 5 |
| 13 | TGME49_269750.P2701 | CrcB family protein | 10 | 8 | 8 | 6 | 3 | 3 | 3 | 5 |
| 19 | TGME49_269780-t26_1-p1 | ADP-ribosylation factor-like protein ArlX1 | 7 | 7 | 7 | 3 | 4 | 5 | 5 | 5 |
| 15 | TGME49_269940-t26_1-p1 | zinc finger motif, C2HC5-type protein | 10 | 9 | 8 | 4 | 5 | 5 | 4 | 5 |
| 15 | TGME49_269950.P2710 | dense granule protein GRA61 | 10 | 9 | 9 | 5 | 3 | 4 | 4 | 5 |
| 14 | TGME49_270000.P2713 | NAD dependent epimerase/dehydratase family protein | 8 | 8 | 8 | 5 | 6 | 6 | 5 | 5 |
| 15 | TGME49_270090.P2718 | hypothetical protein | 10 | 10 | 8 | 4 | 4 | 4 | 4 | 5 |
| 15 | TGME49_270150.P2722 | hypothetical protein | 10 | 10 | 10 | 6 | 5 | 4 | 4 | 5 |
| 8 | TGME49_270210.P2726 | kinesin motor domain-containing protein | 9 | 10 | 9 | 3 | 3 | 3 | 3 | 5 |
| 13 | TGME49_270220.P2727 | hypothetical protein | 10 | 9 | 7 | 5 | 3 | 3 | 3 | 5 |
| 13 | TGME49_270260.P2729 | hypothetical protein | 9 | 9 | 7 | 5 | 3 | 3 | 3 | 5 |
| 14 | TGME49_270300-t26_1-p1 | RING zinc finger protein | 9 | 9 | 7 | 3 | 6 | 6 | 4 | 5 |
| 14 | TGME49_270530.P2770 | ubiquitin fusion degradation protein UFD1CY | 8 | 8 | 8 | 5 | 7 | 3 | 6 | 5 |
| 13 | TGME49_270570.P2774 | hypothetical protein | 10 | 9 | 6 | 5 | 4 | 3 | 3 | 5 |
| 13 | TGME49_270650-t26_1-p1 | deoxyribose-phosphate aldolase | 8 | 7 | 7 | 6 | 3 | 3 | 3 | 5 |
| 14 | TGME49_270710.P2779 | WW domain binding protein 11 | 9 | 7 | 6 | 3 | 6 | 5 | 5 | 5 |
| 14 | TGME49_270740-t26_1-p1 | Cwf15/Cwc15 cell cycle control protein | 7 | 7 | 7 | 4 | 6 | 5 | 4 | 5 |
| 13 | TGME49_270820.P2784 | hypothetical protein | 7 | 8 | 7 | 6 | 3 | 4 | 3 | 5 |

| | | | | | | | | | | |
|---|---|---|---|---|---|---|---|---|---|---|
| 15 | TGME49_270880.P2788 | RNA recognition motif-containing protein | 10 | 9 | 8 | 5 | 5 | 4 | 4 | 5 |
| 13 | TGME49_270940.P2792 | hypothetical protein | 10 | 9 | 7 | 5 | 4 | 3 | 3 | 5 |
| 15 | TGME49_271000-t26_1-p1 | hypothetical protein | 10 | 9 | 8 | 5 | 6 | 5 | 4 | 5 |
| 13 | TGME49_271030.P2797 | AP2 domain transcription factor AP2VIII-6 | 10 | 9 | 6 | 5 | 3 | 3 | 3 | 5 |
| 15 | TGME49_271110.P2801 | hypothetical protein | 10 | 9 | 10 | 4 | 7 | 3 | 4 | 5 |
| 8 | TGME49_271145.P2803 | hypothetical protein | 10 | 8 | 6 | 4 | 4 | 3 | 3 | 5 |
| 13 | TGME49_271170.P2804 | dolichol kinase | 10 | 6 | 6 | 6 | 3 | 4 | 4 | 5 |
| 13 | TGME49_271190-t26_1-p1 | bicoid-interacting protein BIN3 | 10 | 9 | 7 | 6 | 3 | 3 | 3 | 5 |
| 15 | TGME49_271200-t26_1-p1 | AP2 domain transcription factor AP2VIII-5 | 10 | 10 | 9 | 6 | 5 | 5 | 3 | 5 |
| 15 | TGME49_271270-t26_1-p1 | hypothetical protein | 10 | 9 | 8 | 5 | 6 | 4 | 3 | 5 |
| 15 | TGME49_271290.P2808 | hypothetical protein | 10 | 9 | 7 | 5 | 5 | 4 | 4 | 5 |
| 8 | TGME49_271370.P2812 | hypothetical protein | 10 | 9 | 8 | 4 | 3 | 4 | 4 | 5 |
| 15 | TGME49_271380.P2813 | PUB domain-containing protein | 10 | 9 | 9 | 6 | 5 | 4 | 3 | 5 |
| 14 | TGME49_271420-t26_1-p1 | porphobilinogen deaminase | 9 | 8 | 8 | 3 | 5 | 5 | 4 | 5 |
| 8 | TGME49_271430.P2814 | hypothetical protein | 10 | 8 | 7 | 4 | 4 | 4 | 3 | 5 |
| 14 | TGME49_271760.P2822 | seryl-tRNA synthetase (SerRS2) | 10 | 9 | 8 | 3 | 7 | 3 | 4 | 5 |
| 13 | TGME49_271800.P2824 | serine esterase (DUF676) protein | 10 | 9 | 7 | 7 | 4 | 3 | 3 | 5 |
| 15 | TGME49_271860-t26_1-p1 | tRNA (Uracil-5-)-methyltransferase | 10 | 9 | 8 | 5 | 5 | 3 | 3 | 5 |
| 19 | TGME49_271870-t26_1-p1 | zinc carboxypeptidase superfamily protein | 10 | 9 | 9 | 4 | 5 | 7 | 5 | 5 |
| 13 | TGME49_271910.P2829 | DnaJ domain-containing protein | 9 | 8 | 7 | 6 | 4 | 3 | 4 | 5 |
| 8 | TGME49_271950.P2833 | hypothetical protein | 10 | 8 | 6 | 4 | 4 | 3 | 3 | 5 |
| 15 | TGME49_272000.P2835 | hypothetical protein | 10 | 10 | 7 | 5 | 5 | 4 | 3 | 5 |
| 15 | TGME49_272040-t26_1-p1 | WD domain, G-beta repeat-containing protein | 10 | 8 | 7 | 5 | 5 | 4 | 4 | 5 |
| 15 | TGME49_272155.P2839 | hypothetical protein | 10 | 10 | 7 | 4 | 5 | 4 | 3 | 5 |
| 15 | TGME49_272210.P2843 | RNA methyltransferase, putative | 10 | 9 | 8 | 5 | 6 | 5 | 4 | 5 |
| 14 | TGME49_272320.P2851 | palmitoyltransferase DHHC17 | 10 | 9 | 9 | 5 | 7 | 5 | 5 | 5 |
| 15 | TGME49_272390.P2858 | protein TOC75 | 10 | 9 | 7 | 5 | 4 | 4 | 4 | 5 |
| 8 | TGME49_272410.P2860 | phosphogluconate dehydrogenase (decarboxylating), NAD binding domain-containing protein | 8 | 8 | 7 | 3 | 4 | 3 | 3 | 5 |
| 8 | TGME49_272475.P2866 | protein kinase domain-containing protein | 10 | 10 | 7 | 3 | 5 | 3 | 4 | 5 |
| 15 | TGME49_272490.P2867 | protoporphyrinogen oxidase | 10 | 9 | 7 | 5 | 4 | 4 | 4 | 5 |
| 19 | TGME49_272510-t26_1-p1 | aspartyl protease ASP6 | 9 | 5 | 9 | 3 | 3 | 6 | 3 | 5 |
| 13 | TGME49_272540.P2870 | protein kinase (incomplete catalytic triad) | 10 | 9 | 8 | 6 | 3 | 5 | 3 | 5 |
| 15 | TGME49_272570.P2872 | dihydrouridine synthase (dus) protein | 10 | 9 | 8 | 6 | 5 | 4 | 3 | 5 |
| 15 | TGME49_272710.P2881 | AP2 domain transcription factor AP2VIII-4 | 10 | 10 | 7 | 6 | 5 | 3 | 3 | 5 |
| 19 | TGME49_273040-t26_1-p1 | hypothetical protein | 8 | 8 | 7 | 3 | 3 | 6 | 4 | 5 |
| 14 | TGME49_273100-t26_1-p1 | 3'-5' exonuclease domain-containing protein | 9 | 8 | 8 | 3 | 5 | 4 | 6 | 5 |
| 13 | TGME49_273150.P2894 | zinc finger, C3HC4 type (RING finger) domain-containing protein | 9 | 8 | 7 | 5 | 3 | 5 | 4 | 5 |
| 13 | TGME49_273290.P2896 | Mpv17 / PMP22 family protein | 10 | 8 | 7 | 5 | 3 | 3 | 4 | 5 |
| 8 | TGME49_273330-t26_1-p1 | hypothetical protein | 10 | 8 | 6 | 4 | 4 | 3 | 3 | 5 |
| 15 | TGME49_273410-t26_1-p1 | tyrosine-tRNA ligase | 10 | 9 | 8 | 6 | 4 | 5 | 4 | 5 |
| 13 | TGME49_273430-t26_1-p1 | SAM-dependent methyltransferase | 10 | 7 | 8 | 6 | 3 | 4 | 3 | 5 |
| 13 | TGME49_273580.P2910 | conserved oligomeric Golgi complex subunit 2 | 10 | 9 | 8 | 6 | 4 | 3 | 3 | 5 |
| 15 | TGME49_273660.P2914 | AP2 domain transcription factor AP2VIII-3 | 10 | 10 | 6 | 5 | 5 | 4 | 4 | 5 |
| 19 | TGME49_273705-t26_1-p1 | hypothetical protein | 9 | 6 | 8 | 3 | 3 | 5 | 4 | 5 |
| 14 | TGME49_273850.P2923 | nucleoporin NUP68, putative | 8 | 8 | 7 | 5 | 5 | 5 | 5 | 5 |
| 15 | TGME49_273885-t26_1-p1 | hypothetical protein | 10 | 8 | 8 | 5 | 5 | 5 | 4 | 5 |
| 13 | TGME49_274020.P2933 | kinesin motor domain-containing protein | 10 | 7 | 8 | 7 | 3 | 4 | 4 | 5 |

| | | | | | | | | | | |
|---|---|---|---|---|---|---|---|---|---|---|
| 13 | TGME49_274080.P2936 | PIH1 domain-containing protein | 10 | 7 | 7 | 7 | 3 | 4 | 4 | 5 |
| 15 | TGME49_274110.P2939 | glycoprotease family protein | 10 | 9 | 9 | 4 | 5 | 3 | 4 | 5 |
| 14 | TGME49_274140.P2942 | RNA binding protein, putative | 9 | 9 | 8 | 3 | 5 | 5 | 4 | 5 |
| 14 | TGME49_274160.P2944 | preconoidal ring protein PCR1 | 9 | 8 | 7 | 5 | 5 | 5 | 5 | 5 |
| 8 | TGME49_274170-t26_1-p1 | protein kinase (incomplete catalytic triad) | 8 | 8 | 6 | 3 | 3 | 3 | 3 | 5 |
| 15 | TGME49_275490-t26_1-p1 | nucleoporin, putative | 10 | 10 | 10 | 4 | 4 | 3 | 3 | 5 |
| 15 | TGME49_275568-t26_1-p1 | GPI transamidase subunit PIG-U protein | 9 | 9 | 8 | 6 | 5 | 4 | 4 | 5 |
| 13 | TGME49_275660-t26_1-p1 | methyltransferase, putative | 9 | 8 | 8 | 7 | 3 | 3 | 3 | 5 |
| 15 | TGME49_275740-t26_1-p1 | hypothetical protein | 10 | 9 | 7 | 5 | 5 | 4 | 4 | 5 |
| 13 | TGME49_275798-t26_1-p1 | microneme protein, putative | 9 | 8 | 7 | 6 | 3 | 3 | 3 | 5 |
| 8 | TGME49_275830-t26_1-p1 | 3'-5' exonuclease, putative | 10 | 9 | 6 | 4 | 4 | 4 | 4 | 5 |
| 8 | TGME49_275840-t26_1-p1 | protein phosphatase PPM19, putative | 9 | 7 | 7 | 4 | 4 | 3 | 3 | 5 |
| 8 | TGME49_275860-t26_1-p1 | dense granule protein GRA12B | 9 | 7 | 7 | 3 | 3 | 3 | 3 | 5 |
| 8 | TGME49_276100.P2962 | hypothetical protein | 9 | 8 | 6 | 3 | 4 | 3 | 3 | 5 |
| 8 | TGME49_276860.P2971 | LEA domain-containing protein LEA860 | 9 | 8 | 6 | 4 | 3 | 3 | 3 | 5 |
| 14 | TGME49_276910-t26_1-p1 | endoplasmic reticulum lumen protein retaining receptor (ERD2) family protein | 7 | 7 | 7 | 3 | 7 | 3 | 3 | 5 |
| 19 | TGME49_276960.P2974 | tetratricopeptide repeat-containing protein | 10 | 10 | 8 | 3 | 3 | 5 | 4 | 5 |
| 15 | TGME49_277050-t26_1-p1 | SPRY domain-containing protein | 10 | 10 | 8 | 6 | 5 | 5 | 4 | 5 |
| 8 | TGME49_277260.P2980 | hypothetical protein | 10 | 9 | 7 | 4 | 3 | 4 | 4 | 5 |
| 8 | TGME49_277490-t26_1-p1 | hypothetical protein | 10 | 8 | 8 | 3 | 3 | 3 | 3 | 5 |
| 8 | TGME49_277530.P2982 | DNA topoisomerase domain-containing protein | 10 | 9 | 8 | 3 | 3 | 3 | 3 | 5 |
| 15 | TGME49_277550.P2983 | UvrD/REP helicase domain-containing protein | 10 | 8 | 9 | 6 | 6 | 4 | 3 | 5 |
| 8 | TGME49_277560-t26_1-p1 | hypothetical protein | 10 | 9 | 7 | 4 | 4 | 4 | 3 | 5 |
| 13 | TGME49_277710-t26_1-p1 | hypothetical protein | 10 | 7 | 6 | 5 | 4 | 3 | 3 | 5 |
| 14 | TGME49_277790.P2985 | hypothetical protein | 10 | 10 | 5 | 5 | 8 | 4 | 4 | 5 |
| 15 | TGME49_277940.P2991 | apicomplexan kinetochore protein AKIT1 | 10 | 9 | 7 | 5 | 5 | 4 | 4 | 5 |
| 19 | TGME49_277950-t26_1-p1 | lipase | 10 | 8 | 8 | 4 | 3 | 5 | 5 | 5 |
| 15 | TGME49_278020.P2994 | hypothetical protein | 10 | 9 | 8 | 5 | 4 | 4 | 4 | 5 |
| 15 | TGME49_278130.P2995 | basal complex component BCC11 | 10 | 10 | 9 | 5 | 4 | 3 | 3 | 5 |
| 19 | TGME49_278145-t26_1-p1 | hypothetical protein | 9 | 6 | 8 | 3 | 4 | 6 | 3 | 5 |
| 13 | TGME49_278205.P2996 | hypothetical protein | 10 | 9 | 7 | 6 | 4 | 5 | 3 | 5 |
| 15 | TGME49_278260-t26_1-p1 | polynucleotide adenylyltransferase | 10 | 10 | 8 | 5 | 5 | 4 | 4 | 5 |
| 13 | TGME49_278600.P3005 | hypothetical protein | 10 | 9 | 7 | 6 | 4 | 3 | 3 | 5 |
| 13 | TGME49_278720-t26_1-p1 | hypothetical protein | 10 | 9 | 8 | 6 | 3 | 3 | 3 | 5 |
| 13 | TGME49_278770-t26_1-p1 | hypothetical protein | 10 | 9 | 7 | 6 | 4 | 4 | 4 | 5 |
| 13 | TGME49_278840.P3019 | hypothetical protein | 10 | 9 | 8 | 6 | 3 | 3 | 3 | 5 |
| 13 | TGME49_278890.P3022 | protein CASP, putative | 10 | 9 | 8 | 6 | 3 | 4 | 4 | 5 |
| 13 | TGME49_278920-t26_1-p1 | hypothetical protein | 9 | 7 | 7 | 6 | 4 | 3 | 3 | 5 |
| 8 | TGME49_278960-t26_1-p1 | hypothetical protein | 9 | 8 | 7 | 4 | 3 | 3 | 3 | 5 |
| 8 | TGME49_278965.P3025 | hypothetical protein | 9 | 8 | 7 | 4 | 3 | 3 | 3 | 5 |
| 19 | TGME49_279440-t26_1-p1 | PA14 domain-containing protein | 10 | 9 | 10 | 3 | 3 | 7 | 6 | 5 |
| 13 | TGME49_280470.P3040 | AP2 domain transcription factor AP2VIIa-1 | 10 | 9 | 7 | 5 | 3 | 3 | 4 | 5 |
| 8 | TGME49_280590-t26_1-p1 | hypothetical protein | 10 | 10 | 6 | 4 | 3 | 3 | 3 | 5 |
| 15 | TGME49_280790-t26_1-p1 | phosphatidylinositol n-acetylglucosaminyltransferase | 10 | 10 | 9 | 5 | 4 | 4 | 4 | 5 |
| 19 | TGME49_281430-t26_1-p1 | protein kinase, putative | 9 | 8 | 8 | 5 | 3 | 4 | 5 | 5 |
| 14 | TGME49_281440.P3059 | hypothetical protein, DUF3712 domain-containing protein | 3 | 10 | 10 | 3 | 7 | 3 | 5 | 5 |
| 13 | TGME49_281555.P3065 | hypothetical protein | 8 | 9 | 7 | 7 | 4 | 3 | 3 | 5 |

| | | | | | | | | | | |
|---|---|---|---|---|---|---|---|---|---|---|
| 14 | TGME49_281570.P3067 | VPS9 domain-containing protein | 9 | 8 | 7 | 4 | 5 | 5 | 4 | 5 |
| 14 | TGME49_281580.P3068 | protein phosphatase PPM5C, putative | 10 | 9 | 8 | 5 | 6 | 6 | 6 | 5 |
| 14 | TGME49_281630-t26_1-p1 | hydroxyacylglutathione hydrolase | 8 | 7 | 7 | 4 | 6 | 7 | 6 | 5 |
| 8 | TGME49_281640-t26_1-p1 | hypothetical protein | 10 | 9 | 8 | 4 | 4 | 3 | 3 | 5 |
| 15 | TGME49_281960.P3074 | hypothetical protein | 10 | 9 | 8 | 5 | 5 | 3 | 4 | 5 |
| 13 | TGME49_282000.P3077 | hypothetical protein | 10 | 9 | 9 | 6 | 3 | 3 | 3 | 5 |
| 19 | TGME49_282030.P3078 | coiled-coil domain-containing protein 151, putative | 10 | 10 | 8 | 3 | 3 | 4 | 5 | 5 |
| 15 | TGME49_282070.P3079 | inner membrane complex protein IMC33 | 10 | 9 | 8 | 6 | 6 | 3 | 3 | 5 |
| 8 | TGME49_282090-t26_1-p1 | hypothetical protein | 9 | 8 | 7 | 3 | 4 | 3 | 3 | 5 |
| 8 | TGME49_282130.P3080 | hypothetical protein | 10 | 6 | 7 | 4 | 3 | 3 | 3 | 5 |
| 13 | TGME49_282210.P3082 | AP2 domain transcription factor AP2VIIa-8 | 10 | 6 | 8 | 6 | 3 | 4 | 4 | 5 |
| 15 | TGME49_283520.P3089 | hypothetical protein | 10 | 10 | 10 | 6 | 6 | 4 | 4 | 5 |
| 15 | TGME49_283550.P3092 | hypothetical protein | 10 | 10 | 7 | 6 | 5 | 4 | 3 | 5 |
| 14 | TGME49_283720.P3096 | phosphotyrosyl phosphate activator (ptpa) protein | 9 | 8 | 8 | 3 | 7 | 6 | 5 | 5 |
| 14 | TGME49_283740.P3098 | RNA recognition motif-containing protein | 9 | 8 | 7 | 5 | 5 | 4 | 5 | 5 |
| 13 | TGME49_283760.P3099 | FAD-binding domain-containing protein | 10 | 8 | 7 | 5 | 4 | 4 | 3 | 5 |
| 13 | TGME49_283790-t26_1-p1 | protein kinase, putative | 10 | 8 | 8 | 6 | 3 | 4 | 3 | 5 |
| 14 | TGME49_283830-t26_1-p1 | type I inorganic pyrophosphatase PPase | 8 | 6 | 7 | 4 | 6 | 5 | 4 | 5 |
| 19 | TGME49_283878-t26_1-p1 | RNA recognition motif-containing protein | 9 | 8 | 7 | 5 | 4 | 6 | 5 | 5 |
| 13 | TGME49_283890.P3103 | histone lysine demethylase JMJD6b | 9 | 9 | 8 | 6 | 3 | 3 | 3 | 5 |
| 13 | TGME49_284170.P3106 | palmitoyltransferase DHHC11 | 10 | 8 | 7 | 5 | 4 | 4 | 4 | 5 |
| 8 | TGME49_284180-t26_1-p1 | hypothetical protein | 10 | 9 | 7 | 3 | 3 | 4 | 3 | 5 |
| 8 | TGME49_284645.P3113 | hypothetical protein | 10 | 10 | 5 | 4 | 3 | 3 | 3 | 5 |
| 15 | TGME49_285170.P3118 | ribosomal RNA large subunit methyltransferase I, putative | 10 | 8 | 8 | 6 | 4 | 5 | 4 | 5 |
| 15 | TGME49_285180.P3119 | hypothetical protein | 9 | 9 | 8 | 4 | 5 | 3 | 3 | 5 |
| 8 | TGME49_285190.P3120 | zinc finger, C3HC4 type (RING finger) domain-containing protein | 9 | 8 | 7 | 4 | 5 | 4 | 3 | 5 |
| 13 | TGME49_285290-t26_1-p1 | hypothetical protein | 8 | 8 | 7 | 6 | 3 | 3 | 4 | 5 |
| 8 | TGME49_285480-t26_1-p1 | hypothetical protein | 10 | 9 | 8 | 4 | 4 | 3 | 3 | 5 |
| 13 | TGME49_285490-t26_1-p1 | helix-hairpin-helix motif domain-containing protein | 10 | 8 | 7 | 5 | 4 | 3 | 3 | 5 |
| 15 | TGME49_285500.P3133 | hypothetical protein | 10 | 10 | 9 | 6 | 6 | 5 | 3 | 5 |
| 15 | TGME49_285520-t26_1-p1 | RNA cap guanine-N2 methyltransferase | 10 | 9 | 7 | 5 | 4 | 5 | 4 | 5 |
| 13 | TGME49_285650.P3134 | hypothetical protein | 10 | 9 | 6 | 6 | 4 | 3 | 3 | 5 |
| 13 | TGME49_285780-t26_1-p1 | gamma-tubulin complex component, putative | 10 | 7 | 8 | 7 | 4 | 4 | 3 | 5 |
| 15 | TGME49_285830.P3138 | hypothetical protein | 10 | 10 | 10 | 4 | 4 | 3 | 3 | 5 |
| 14 | TGME49_285840-t26_1-p1 | RAP domain-containing protein | 8 | 7 | 6 | 3 | 6 | 5 | 6 | 5 |
| 13 | TGME49_285895.P3139 | AP2 domain transcription factor AP2V-2 | 10 | 8 | 6 | 5 | 3 | 4 | 4 | 5 |
| 13 | TGME49_286130.P3148 | guanylylate cyclase, putative | 9 | 8 | 5 | 6 | 3 | 3 | 3 | 5 |
| 8 | TGME49_286150.P3149 | PAN/Apple domain-containing protein | 9 | 9 | 8 | 3 | 3 | 3 | 3 | 5 |
| 8 | TGME49_286180-t26_1-p1 | tRNA ligases class I (M) protein | 10 | 9 | 6 | 4 | 4 | 3 | 3 | 5 |
| 15 | TGME49_286230-t26_1-p1 | hypothetical protein | 10 | 10 | 10 | 4 | 4 | 3 | 3 | 5 |
| 15 | TGME49_286270-t26_1-p1 | hypothetical protein | 10 | 10 | 8 | 5 | 4 | 4 | 4 | 5 |
| 14 | TGME49_286530.P3155 | cytochrome c oxidase subunit ApiCOX24 | 7 | 7 | 7 | 3 | 6 | 3 | 6 | 5 |
| 8 | TGME49_286740-t26_1-p1 | microneme-like protein | 10 | 9 | 9 | 3 | 3 | 3 | 4 | 5 |
| 8 | TGME49_286760.P3164 | hypothetical protein | 9 | 8 | 6 | 4 | 3 | 3 | 3 | 5 |
| 15 | TGME49_287260-t26_1-p1 | hypothetical protein | 10 | 10 | 9 | 3 | 5 | 4 | 4 | 5 |

| | | | | | | | | | | |
|---|---|---|---|---|---|---|---|---|---|---|
| 15 | TGME49_287970-t26_1-p1 | hypothetical protein | 10 | 10 | 7 | 6 | 4 | 4 | 4 | 5 |
| 13 | TGME49_288240-t26_1-p1 | GRIP domain-containing protein | 10 | 8 | 7 | 5 | 4 | 3 | 3 | 5 |
| 15 | TGME49_288250-t26_1-p1 | WD repeat-containing protein | 10 | 8 | 9 | 5 | 4 | 4 | 4 | 5 |
| 13 | TGME49_288280-t26_1-p1 | hypothetical protein | 10 | 9 | 8 | 5 | 3 | 3 | 3 | 5 |
| 19 | TGME49_288350-t26_1-p1 | hypothetical protein | 10 | 6 | 6 | 4 | 3 | 5 | 5 | 5 |
| 19 | TGME49_288420-t26_1-p1 | hypothetical protein | 10 | 8 | 7 | 4 | 3 | 5 | 4 | 5 |
| 19 | TGME49_288440-t26_1-p1 | NEK kinase | 10 | 10 | 8 | 4 | 3 | 3 | 6 | 5 |
| 13 | TGME49_288560-t26_1-p1 | hypothetical protein | 10 | 7 | 7 | 5 | 3 | 3 | 3 | 5 |
| 8 | TGME49_288640-t26_1-p1 | radical SAM domain-containing protein | 9 | 8 | 8 | 3 | 3 | 4 | 4 | 5 |
| 13 | TGME49_288670-t26_1-p1 | hypothetical protein | 10 | 9 | 8 | 6 | 3 | 3 | 3 | 5 |
| 13 | TGME49_288820.P3191 | magnesium transporter, putative | 10 | 9 | 8 | 6 | 3 | 3 | 3 | 5 |
| 15 | TGME49_288890-t26_1-p1 | hypothetical protein | 10 | 10 | 7 | 5 | 4 | 4 | 3 | 5 |
| 15 | TGME49_288930-t26_1-p1 | condensin complex subunit 2, putative | 10 | 10 | 9 | 4 | 6 | 5 | 4 | 5 |
| 13 | TGME49_288950-t26_1-p1 | AP2 domain transcription factor AP2IX-4 | 10 | 10 | 8 | 6 | 3 | 3 | 3 | 5 |
| 13 | TGME49_289020.P3196 | hypothetical protein | 9 | 8 | 8 | 7 | 3 | 3 | 3 | 5 |
| 14 | TGME49_289140-t26_1-p1 | ribosomal protein l22/l43, putative | 7 | 7 | 7 | 3 | 6 | 6 | 4 | 5 |
| 15 | TGME49_289150-t26_1-p1 | hypothetical protein | 10 | 9 | 9 | 5 | 5 | 5 | 5 | 5 |
| 15 | TGME49_289510.P3204 | H/ACA ribonucleoprotein complex subunit 1, putative | 10 | 8 | 7 | 5 | 5 | 4 | 4 | 5 |
| 15 | TGME49_289520.P3205 | vacuolar protein sorting-associated protein 8 | 10 | 10 | 10 | 5 | 5 | 4 | 4 | 5 |
| 14 | TGME49_289560-t26_1-p1 | RNA recognition motif-containing protein | 8 | 7 | 7 | 3 | 6 | 5 | 5 | 5 |
| 14 | TGME49_289640-t26_1-p1 | hypothetical protein | 9 | 8 | 3 | 3 | 6 | 4 | 5 | 5 |
| 8 | TGME49_289700.P3210 | apicomplexan kinetochore protein AKIT6, putative | 9 | 8 | 6 | 3 | 4 | 3 | 3 | 5 |
| 15 | TGME49_289710.P3211 | AP2 domain transcription factor AP2IX-5 | 10 | 10 | 8 | 5 | 5 | 4 | 5 | 5 |
| 14 | TGME49_289800-t26_1-p1 | transmembrane emp24 domain-containing protein | 8 | 7 | 7 | 3 | 6 | 3 | 6 | 5 |
| 8 | TGME49_289930-t26_1-p1 | phosphoenolpyruvate carboxykinase, putative | 10 | 9 | 9 | 3 | 3 | 3 | 3 | 5 |
| 14 | TGME49_289960.P3219 | hypothetical protein | 9 | 8 | 8 | 5 | 6 | 5 | 5 | 5 |
| 8 | TGME49_290190.P3224 | protein SPT2, putative | 10 | 8 | 7 | 4 | 4 | 4 | 4 | 5 |
| 15 | TGME49_290310-t26_1-p1 | conserved oligomeric Golgi complex subunit 1 | 10 | 8 | 9 | 5 | 4 | 4 | 3 | 5 |
| 15 | TGME49_290630-t26_1-p1 | AP2 domain transcription factor AP2IX-7 | 10 | 10 | 9 | 5 | 4 | 4 | 4 | 5 |
| 13 | TGME49_290650-t26_1-p1 | hypothetical protein | 9 | 8 | 7 | 6 | 3 | 4 | 4 | 5 |
| 15 | TGME49_290678.P3230 | hypothetical protein | 10 | 10 | 8 | 6 | 5 | 3 | 3 | 5 |
| 14 | TGME49_290730.P3233 | CS domain-containing protein | 8 | 6 | 7 | 4 | 6 | 3 | 5 | 5 |
| 13 | TGME49_290910-t26_1-p1 | hypothetical protein | 10 | 7 | 6 | 6 | 4 | 3 | 3 | 5 |
| 19 | TGME49_291120-t26_1-p1 | trafficking protein mon1 subfamily protein | 10 | 10 | 9 | 5 | 4 | 5 | 6 | 5 |
| 13 | TGME49_291130-t26_1-p1 | hypothetical protein | 10 | 9 | 8 | 6 | 3 | 4 | 4 | 5 |
| 15 | TGME49_291150-t26_1-p1 | zinc finger (CCCH type) motif-containing protein | 10 | 8 | 9 | 6 | 5 | 4 | 4 | 5 |
| 19 | TGME49_291310-t26_1-p1 | V-ATPase subunit c' proteolipid, putative | 7 | 7 | 7 | 3 | 3 | 6 | 6 | 5 |
| 14 | TGME49_291320-t26_1-p1 | hypothetical protein | 9 | 7 | 7 | 4 | 6 | 5 | 5 | 5 |
| 8 | TGME49_291340.P3249 | hypothetical protein | 10 | 8 | 6 | 4 | 3 | 4 | 3 | 5 |
| 15 | TGME49_291350-t26_1-p1 | hypothetical protein | 10 | 9 | 9 | 5 | 5 | 4 | 4 | 5 |
| 13 | TGME49_291590-t26_1-p1 | MYND-like zinc finger protein | 10 | 8 | 8 | 6 | 3 | 5 | 3 | 5 |
| 13 | TGME49_291690-t26_1-p1 | hypothetical protein | 10 | 8 | 8 | 7 | 3 | 3 | 4 | 5 |
| 13 | TGME49_291830.P3251 | hypothetical protein, conserved | 10 | 9 | 8 | 6 | 3 | 3 | 3 | 5 |
| 13 | TGME49_291940-t26_1-p1 | hypothetical protein | 10 | 9 | 7 | 7 | 3 | 4 | 4 | 5 |
| 15 | TGME49_292035.P3258 | hypothetical protein | 10 | 10 | 8 | 6 | 5 | 4 | 4 | 5 |
| 14 | TGME49_292100-t26_1-p1 | 2-oxoisovalerate dehydrogenase subunit alpha, putative | 9 | 8 | 8 | 3 | 7 | 3 | 3 | 5 |
| 19 | TGME49_292110.P3260 | formate/nitrite transporter FNT2 | 9 | 8 | 8 | 3 | 3 | 3 | 7 | 5 |

| | | | | | | | | | | |
|---|---|---|---|---|---|---|---|---|---|---|
| 15 | TGME49_292235-t26_1-p1 | E3 ubiquitin-protein ligase ZNF598, putative | 10 | 9 | 8 | 5 | 6 | 5 | 4 | 5 |
| 8 | TGME49_292340-t26_1-p1 | zinc finger, C3HC4 type (RING finger) domain-containing protein | 10 | 9 | 8 | 3 | 4 | 4 | 4 | 5 |
| 15 | TGME49_292360.P3266 | hypothetical protein | 10 | 9 | 9 | 6 | 5 | 4 | 3 | 5 |
| 15 | TGME49_292950.P3269 | apical cap protein AC10 | 10 | 10 | 9 | 6 | 5 | 4 | 4 | 5 |
| 8 | TGME49_292960-t26_1-p1 | hypothetical protein | 9 | 9 | 6 | 3 | 5 | 3 | 3 | 5 |
| 15 | TGME49_293170.P3274 | CH domain-containing protein | 10 | 10 | 9 | 5 | 5 | 5 | 5 | 5 |
| 15 | TGME49_293240.P3280 | triacylglycerol lipase | 10 | 9 | 8 | 5 | 5 | 5 | 5 | 5 |
| 13 | TGME49_293258.P3283 | hypothetical protein | 10 | 8 | 6 | 5 | 3 | 4 | 4 | 5 |
| 15 | TGME49_293260.P3284 | ATPase/histidine kinase/DNA gyrase B/HSP90 domain-containing protein | 10 | 9 | 8 | 6 | 5 | 3 | 4 | 5 |
| 8 | TGME49_293330.P3289 | protein of unknown function UPF0301 | 10 | 9 | 7 | 4 | 4 | 3 | 3 | 5 |
| 19 | TGME49_293350.P3291 | mitochondrial carrier superfamily protein | 9 | 8 | 8 | 5 | 4 | 5 | 5 | 5 |
| 8 | TGME49_293425.P3296 | apicomplexan amino acid transporter ApiAT5-6 | 6 | 9 | 8 | 3 | 3 | 3 | 3 | 5 |
| 13 | TGME49_293430.P3297 | hypothetical protein | 9 | 8 | 7 | 5 | 3 | 3 | 4 | 5 |
| 13 | TGME49_293440.P3298 | hypothetical protein | 9 | 8 | 7 | 6 | 4 | 3 | 3 | 5 |
| 13 | TGME49_293620.P3313 | hypothetical protein | 10 | 9 | 7 | 5 | 3 | 3 | 3 | 5 |
| 8 | TGME49_293760.P3325 | EF hand domain-containing protein | 9 | 8 | 7 | 4 | 3 | 3 | 3 | 5 |
| 8 | TGME49_293780.P3327 | hypothetical protein | 8 | 8 | 7 | 4 | 3 | 3 | 3 | 5 |
| 8 | TGME49_293790.P3328 | hypothetical protein | 8 | 8 | 7 | 3 | 4 | 3 | 3 | 5 |
| 13 | TGME49_293830.P3332 | methyltransferase domain-containing protein | 9 | 7 | 7 | 6 | 3 | 3 | 3 | 5 |
| 13 | TGME49_293850.P3335 | clusterin-associated protein 1, putative | 9 | 7 | 7 | 6 | 3 | 3 | 3 | 5 |
| 14 | TGME49_294180-t26_1-p1 | tRNA -methyltransferase family protein | 8 | 8 | 8 | 4 | 7 | 6 | 5 | 5 |
| 14 | TGME49_294190.P3345 | enoyl-CoA hydratase/isomerase family protein | 10 | 8 | 9 | 4 | 7 | 7 | 3 | 5 |
| 14 | TGME49_294210.P3347 | G-protein gamma-like domain-containing protein | 10 | 9 | 8 | 3 | 7 | 3 | 4 | 5 |
| 14 | TGME49_294260.P3349 | NEK kinase | 9 | 7 | 8 | 3 | 6 | 4 | 3 | 5 |
| 13 | TGME49_294560.P3358 | rhoptry kinase family protein ROP37 (incomplete catalytic triad) | 9 | 8 | 7 | 5 | 3 | 3 | 3 | 5 |
| 13 | TGME49_294585-t26_1-p1 | hypothetical protein | 10 | 7 | 8 | 5 | 3 | 3 | 3 | 5 |
| 15 | TGME49_294730-t26_1-p1 | Sec16-like protein | 10 | 9 | 8 | 5 | 5 | 4 | 4 | 5 |
| 8 | TGME49_294740-t26_1-p1 | hypothetical protein | 10 | 9 | 7 | 4 | 4 | 3 | 3 | 5 |
| 15 | TGME49_294840.P3370 | zinc finger (CCCH type) motif-containing protein | 10 | 9 | 7 | 6 | 4 | 5 | 4 | 5 |
| 13 | TGME49_294860.P3372 | basal complex component BCC0 | 10 | 9 | 7 | 7 | 3 | 3 | 3 | 5 |
| 8 | TGME49_294902-t26_1-p1 | lipoate-protein ligase 2, putative | 10 | 8 | 7 | 4 | 4 | 4 | 3 | 5 |
| 14 | TGME49_294920-t26_1-p1 | general transcription factor IIH polypeptide 2 GTF2H2 | 9 | 9 | 9 | 4 | 6 | 5 | 6 | 5 |
| 14 | TGME49_294930-t26_1-p1 | leucine rich repeat-containing protein | 10 | 8 | 8 | 5 | 6 | 5 | 5 | 5 |
| 13 | TGME49_294960.P3375 | hypothetical protein | 9 | 8 | 8 | 5 | 4 | 4 | 4 | 5 |
| 13 | TGME49_295030.P3378 | hypothetical protein | 10 | 8 | 7 | 5 | 3 | 3 | 3 | 5 |
| 13 | TGME49_295050-t26_1-p1 | tRNA ligase class II core domain (G, H, P, S and T) domain-containing protein | 10 | 8 | 7 | 6 | 3 | 4 | 4 | 5 |
| 13 | TGME49_295060-t26_1-p1 | exonuclease | 9 | 8 | 7 | 5 | 3 | 3 | 3 | 5 |
| 8 | TGME49_295100.P3381 | hypothetical protein | 10 | 9 | 6 | 3 | 3 | 3 | 3 | 5 |
| 15 | TGME49_295330-t26_1-p1 | Vps54 family protein | 10 | 10 | 9 | 6 | 5 | 4 | 4 | 5 |
| 15 | TGME49_295380-t26_1-p1 | hypothetical protein | 10 | 10 | 10 | 6 | 5 | 5 | 4 | 5 |
| 15 | TGME49_295410-t26_1-p1 | transcription initiation factor TFIID complex subunit TAF6 | 10 | 9 | 9 | 6 | 5 | 4 | 4 | 5 |
| 13 | TGME49_295440-t26_1-p1 | hypothetical protein | 9 | 7 | 6 | 6 | 3 | 3 | 3 | 5 |
| 19 | TGME49_295610-t26_1-p1 | histone lysine methyltransferase, SET, putative | 10 | 10 | 8 | 5 | 4 | 6 | 6 | 5 |
| 15 | TGME49_295620-t26_1-p1 | zinc finger (CCCH type) motif-containing protein | 10 | 10 | 8 | 4 | 4 | 4 | 3 | 5 |

| | | | | | | | | | | |
|---|---|---|---|---|---|---|---|---|---|---|
| 8 | TGME49_295770-t26_1-p1 | hypothetical protein | 10 | 10 | 6 | 4 | 3 | 3 | 3 | 5 |
| 13 | TGME49_297080-t26_1-p1 | pyridoxal kinase | 9 | 7 | 8 | 7 | 4 | 3 | 3 | 5 |
| 15 | TGME49_297120.P3400 | inner membrane complex protein IMC44 | 10 | 9 | 7 | 5 | 4 | 4 | 4 | 5 |
| 13 | TGME49_297160-t26_1-p1 | hypothetical protein | 9 | 8 | 7 | 5 | 4 | 4 | 3 | 5 |
| 15 | TGME49_297180-t26_1-p1 | CRAL-TRIO domain-containing protein | 10 | 9 | 8 | 6 | 5 | 5 | 4 | 5 |
| 8 | TGME49_297210-t26_1-p1 | E3 ubiquitin-protein ligase RBBP6, putative | 10 | 8 | 6 | 4 | 4 | 3 | 3 | 5 |
| 13 | TGME49_297310.P3403 | hypothetical protein | 9 | 7 | 8 | 6 | 3 | 3 | 3 | 5 |
| 13 | TGME49_297410.P3405 | hypothetical protein | 9 | 8 | 7 | 6 | 3 | 3 | 3 | 5 |
| 14 | TGME49_297460.P3407 | WD repeat-containing protein | 9 | 9 | 8 | 5 | 6 | 5 | 5 | 5 |
| 15 | TGME49_297480.P3410 | hypothetical protein | 10 | 9 | 9 | 4 | 4 | 3 | 4 | 5 |
| 14 | TGME49_297520.P3412 | proteophosphoglycan PPG1 | 10 | 9 | 8 | 4 | 7 | 4 | 4 | 5 |
| 8 | TGME49_297710-t26_1-p1 | hypothetical protein | 9 | 9 | 8 | 3 | 3 | 4 | 4 | 5 |
| 15 | TGME49_297890-t26_1-p1 | hypothetical protein | 10 | 10 | 9 | 5 | 5 | 3 | 4 | 5 |
| 8 | TGME49_297900-t26_1-p1 | dense granule protein GRA83 | 10 | 9 | 8 | 3 | 4 | 3 | 3 | 5 |
| 13 | TGME49_298050-t26_1-p1 | hypothetical protein | 9 | 7 | 7 | 6 | 4 | 4 | 3 | 5 |
| 15 | TGME49_299020.P3433 | AP2 domain transcription factor AP2III-4 | 10 | 9 | 9 | 5 | 4 | 5 | 3 | 5 |
| 8 | TGME49_299030-t26_1-p1 | RNA recognition motif 2 protein | 9 | 7 | 7 | 3 | 3 | 3 | 3 | 5 |
| 8 | TGME49_299130-t26_1-p1 | hypothetical protein | 10 | 10 | 7 | 4 | 3 | 3 | 3 | 5 |
| 13 | TGME49_299150-t26_1-p1 | AP2 domain transcription factor AP2III-3 | 10 | 8 | 8 | 6 | 3 | 3 | 3 | 5 |
| 8 | TGME49_299168.P3436 | ATPase, AAA family protein | 9 | 7 | 7 | 3 | 3 | 3 | 3 | 5 |
| 19 | TGME49_299180-t26_1-p1 | synaptobrevin homolog YKT6, putative | 7 | 7 | 7 | 4 | 3 | 5 | 5 | 5 |
| 15 | TGME49_299190-t26_1-p1 | B-box zinc finger domain-containing protein | 10 | 9 | 7 | 5 | 5 | 4 | 5 | 5 |
| 8 | TGME49_299240-t26_1-p1 | hypothetical protein | 10 | 8 | 8 | 4 | 3 | 4 | 4 | 5 |
| 8 | TGME49_299250.P3440 | hypothetical protein | 10 | 8 | 6 | 4 | 4 | 3 | 3 | 5 |
| 15 | TGME49_299900-t26_1-p1 | hypothetical protein | 10 | 9 | 9 | 6 | 5 | 3 | 3 | 5 |
| 15 | TGME49_300030.P3451 | hypothetical protein | 10 | 9 | 9 | 6 | 6 | 3 | 3 | 5 |
| 14 | TGME49_300270-t26_1-p1 | hypothetical protein | 9 | 7 | 6 | 5 | 5 | 5 | 5 | 5 |
| 8 | TGME49_301216.P3469 | endonuclease/exonuclease/phosphatase family protein | 8 | 8 | 8 | 4 | 3 | 3 | 3 | 5 |
| 14 | TGME49_301222-t26_1-p1 | DNA repair protein Rad4 domain-containing protein | 10 | 7 | 7 | 4 | 6 | 5 | 4 | 5 |
| 15 | TGME49_301270.P3471 | tyrosine kinase-like protein TKL1 | 10 | 10 | 9 | 6 | 4 | 5 | 3 | 5 |
| 15 | TGME49_301400.P3477 | hypothetical protein | 10 | 10 | 9 | 4 | 4 | 3 | 5 | 5 |
| 19 | TGME49_304640-t26_1-p1 | hypothetical protein | 10 | 9 | 9 | 3 | 3 | 5 | 4 | 5 |
| 8 | TGME49_304660-t26_1-p1 | prostaglandin-E synthase | 9 | 8 | 7 | 3 | 5 | 3 | 3 | 5 |
| 15 | TGME49_304700-t26_1-p1 | hypothetical protein | 10 | 10 | 9 | 4 | 4 | 3 | 3 | 5 |
| 15 | TGME49_304720.P3488 | SWIM zinc finger domain-containing protein | 10 | 10 | 10 | 5 | 5 | 5 | 3 | 5 |
| 13 | TGME49_304750.P3491 | zinc finger, C3HC4 type (RING finger) domain-containing protein | 10 | 9 | 7 | 5 | 4 | 3 | 4 | 5 |
| 8 | TGME49_304760-t26_1-p1 | RNA recognition motif-containing protein | 10 | 9 | 7 | 3 | 3 | 4 | 4 | 5 |
| 8 | TGME49_304880-t26_1-p1 | hypothetical protein | 10 | 10 | 8 | 4 | 4 | 3 | 3 | 5 |
| 14 | TGME49_304900-t26_1-p1 | hypothetical protein | 8 | 8 | 7 | 5 | 6 | 5 | 5 | 5 |
| 15 | TGME49_304910-t26_1-p1 | hypothetical protein | 10 | 10 | 9 | 5 | 5 | 4 | 4 | 5 |
| 8 | TGME49_305200-t26_1-p1 | tetratricopeptide repeat-containing protein | 10 | 8 | 8 | 4 | 3 | 4 | 3 | 5 |
| 19 | TGME49_305270-t26_1-p1 | hypothetical protein | 10 | 9 | 8 | 3 | 3 | 4 | 6 | 5 |
| 8 | TGME49_305330.P3501 | cyclin, N-terminal domain-containing protein | 10 | 9 | 6 | 3 | 4 | 4 | 5 | 5 |
| 8 | TGME49_305570.P3506 | intraflagellar transport protein IFT81 | 9 | 8 | 8 | 3 | 3 | 3 | 3 | 5 |
| 8 | TGME49_305580.P3507 | ankyrin repeat-containing protein | 8 | 7 | 7 | 3 | 3 | 4 | 3 | 5 |
| 8 | TGME49_305630.P3508 | F-box domain-containing protein | 10 | 8 | 7 | 3 | 3 | 3 | 4 | 5 |

| | | | | | | | | | | |
|---|---|---|---|---|---|---|---|---|---|---|
| 13 | TGME49_305770-t26_1-p1 | ABC transporter, ATP-binding domain-containing protein | 9 | 7 | 8 | 6 | 3 | 4 | 4 | 5 |
| 8 | TGME49_305890.P3510 | hypothetical protein | 10 | 10 | 10 | 3 | 3 | 3 | 3 | 5 |
| 13 | TGME49_305910-t26_1-p1 | hypothetical protein | 9 | 8 | 7 | 7 | 3 | 4 | 4 | 5 |
| 13 | TGME49_306260.P3518 | 2-C-methyl-D-erythritol 4-phosphate cytidylyltransferase, putative | 10 | 7 | 7 | 5 | 4 | 3 | 3 | 5 |
| 13 | TGME49_306400-t26_1-p1 | zinc finger (CCCH type) motif-containing protein | 10 | 9 | 6 | 5 | 4 | 4 | 4 | 5 |
| 14 | TGME49_306410-t26_1-p1 | conserved oligomeric Golgi complex subunit 3 | 10 | 9 | 9 | 4 | 6 | 5 | 5 | 5 |
| 8 | TGME49_306440-t26_1-p1 | hypothetical protein | 10 | 7 | 6 | 4 | 4 | 4 | 3 | 5 |
| 15 | TGME49_306460.P3523 | bromodomain-containing protein BDP4 | 10 | 9 | 8 | 6 | 5 | 4 | 5 | 5 |
| 13 | TGME49_306520-t26_1-p1 | tRNA pseudouridine synthase B, putative | 10 | 7 | 7 | 5 | 4 | 5 | 4 | 5 |
| 13 | TGME49_306540-t26_1-p1 | phosphotransferase enzyme family protein | 9 | 8 | 8 | 5 | 3 | 3 | 4 | 5 |
| 13 | TGME49_306550-t26_1-p1 | hypothetical protein | 10 | 7 | 7 | 5 | 4 | 3 | 4 | 5 |
| 19 | TGME49_306630-t26_1-p1 | tRNA methyltransferase complex GCD14 subunit protein | 9 | 8 | 8 | 3 | 5 | 6 | 6 | 5 |
| 13 | TGME49_306920.P3530 | HOOK interacting protein HIP | 9 | 8 | 7 | 5 | 4 | 4 | 4 | 5 |
| 15 | TGME49_306950.P3532 | RAP domain-containing protein | 10 | 10 | 9 | 5 | 5 | 3 | 3 | 5 |
| 14 | TGME49_306970-t26_1-p1 | thymidylate kinase | 8 | 7 | 7 | 3 | 5 | 5 | 5 | 5 |
| 13 | TGME49_307810-t26_1-p1 | hypothetical protein | 9 | 8 | 7 | 5 | 4 | 3 | 3 | 5 |
| 8 | TGME49_307990-t26_1-p1 | arrestin (or s-antigen), n-terminal domain-containing protein | 9 | 8 | 8 | 3 | 4 | 4 | 4 | 5 |
| 15 | TGME49_308060-t26_1-p1 | PH domain-containing protein | 10 | 10 | 9 | 4 | 4 | 4 | 4 | 5 |
| 15 | TGME49_308070.P3542 | hypothetical protein | 10 | 9 | 7 | 5 | 4 | 4 | 4 | 5 |
| 13 | TGME49_308590-t26_1-p1 | Mov34/MPN/PAD-1 family protein | 9 | 8 | 8 | 7 | 4 | 3 | 4 | 5 |
| 13 | TGME49_308850-t26_1-p1 | hypothetical protein | 10 | 8 | 7 | 5 | 4 | 3 | 3 | 5 |
| 15 | TGME49_308870.P3546 | zinc finger (CCCH type) motif-containing protein | 10 | 8 | 7 | 4 | 5 | 4 | 4 | 5 |
| 13 | TGME49_308880-t26_1-p1 | ImpB/MucB/SamB family protein | 10 | 8 | 6 | 6 | 4 | 3 | 4 | 5 |
| 8 | TGME49_308940.P3549 | hypothetical protein | 10 | 9 | 8 | 3 | 4 | 3 | 3 | 5 |
| 13 | TGME49_308950.P3550 | histidine acid phosphatase superfamily protein | 9 | 9 | 9 | 5 | 3 | 3 | 3 | 5 |
| 8 | TGME49_308970.P3553 | dense granule protein GRA12D | 8 | 8 | 7 | 3 | 5 | 3 | 3 | 5 |
| 8 | TGME49_308980.P3554 | hypothetical protein | 10 | 9 | 6 | 3 | 4 | 3 | 3 | 5 |
| 13 | TGME49_309000-t26_1-p1 | hypothetical protein | 10 | 7 | 7 | 5 | 3 | 4 | 3 | 5 |
| 8 | TGME49_309060-t26_1-p1 | ubiquitin carboxyl-terminal hydrolase | 10 | 8 | 7 | 3 | 3 | 3 | 3 | 5 |
| 13 | TGME49_309090-t26_1-p1 | hypothetical protein | 10 | 8 | 7 | 6 | 4 | 4 | 4 | 5 |
| 13 | TGME49_309100.P3558 | peroxisomal biogenesis factor 3, putative | 10 | 9 | 7 | 5 | 4 | 4 | 3 | 5 |
| 8 | TGME49_309200.P3564 | zinc finger (CCCH type) motif-containing protein | 10 | 8 | 7 | 4 | 3 | 3 | 3 | 5 |
| 13 | TGME49_309410-t26_1-p1 | AP2 domain transcription factor AP2XI-1 | 10 | 9 | 7 | 5 | 3 | 3 | 3 | 5 |
| 15 | TGME49_309600.P3573 | dense granule protein GRA71 | 10 | 9 | 9 | 5 | 5 | 5 | 5 | 5 |
| 8 | TGME49_309720-t26_1-p1 | hypothetical protein | 10 | 9 | 8 | 3 | 4 | 4 | 4 | 5 |
| 8 | TGME49_309770-t26_1-p1 | hypothetical protein | 10 | 9 | 7 | 4 | 3 | 4 | 4 | 5 |
| 8 | TGME49_309870.P3580 | PX domain-containing protein | 10 | 10 | 9 | 3 | 3 | 4 | 3 | 5 |
| 13 | TGME49_309920.P3584 | hypothetical protein | 10 | 8 | 7 | 5 | 4 | 3 | 3 | 5 |
| 14 | TGME49_309960-t26_1-p1 | hypothetical protein | 10 | 9 | 8 | 3 | 6 | 5 | 5 | 5 |
| 14 | TGME49_310060-t26_1-p1 | small nuclease | 9 | 7 | 7 | 3 | 6 | 3 | 5 | 5 |
| 8 | TGME49_310170-t26_1-p1 | hypothetical protein | 10 | 9 | 7 | 3 | 3 | 3 | 3 | 5 |
| 14 | TGME49_310280.P3603 | Phosphoethanolamine-cytidyltransferase | 10 | 9 | 7 | 3 | 6 | 6 | 6 | 5 |
| 13 | TGME49_310320-t26_1-p1 | calreticulin family protein | 9 | 8 | 8 | 7 | 3 | 4 | 4 | 5 |
| 13 | TGME49_310350-t26_1-p1 | PGAP1 family protein | 10 | 9 | 6 | 6 | 5 | 3 | 3 | 5 |
| 13 | TGME49_310400.P3607 | glycosyltransferase GAT1 | 9 | 8 | 7 | 7 | 4 | 4 | 4 | 5 |
| 19 | TGME49_310560.P3615 | TRP cation channel TRPPL2 | 10 | 9 | 9 | 5 | 4 | 7 | 5 | 5 |

| | | | | | | | | | | |
|---|---|---|---|---|---|---|---|---|---|---|
| 13 | TGME49_310620.P3616 | starch binding domain-containing protein | 10 | 7 | 6 | 5 | 3 | 4 | 4 | 5 |
| 13 | TGME49_310630.P3617 | CDK5 regulatory subunit-associated protein 3, putative | 9 | 9 | 8 | 6 | 4 | 3 | 3 | 5 |
| 15 | TGME49_310720-t26_1-p1 | hypothetical protein | 10 | 10 | 9 | 5 | 5 | 5 | 3 | 5 |
| 13 | TGME49_310730.P3621 | apicomplexan kinetochore protein AKIT4, putative | 9 | 8 | 7 | 7 | 4 | 3 | 4 | 5 |
| 13 | TGME49_310740-t26_1-p1 | hypothetical protein | 10 | 8 | 7 | 5 | 3 | 3 | 3 | 5 |
| 14 | TGME49_310750-t26_1-p1 | emp24/gp25L/p24 family protein | 9 | 3 | 7 | 3 | 6 | 6 | 6 | 5 |
| 13 | TGME49_310802.P3624 | CRAL/TRIO domain-containing protein | 9 | 9 | 7 | 7 | 3 | 3 | 3 | 5 |
| 13 | TGME49_310810-t26_1-p1 | apyrase | 9 | 8 | 8 | 5 | 3 | 3 | 3 | 5 |
| 8 | TGME49_310900.P3626 | AP2 domain transcription factor AP2XI-2 | 10 | 10 | 8 | 3 | 3 | 4 | 3 | 5 |
| 14 | TGME49_310910.P3627 | WD domain, G-beta repeat-containing protein | 10 | 9 | 4 | 3 | 5 | 5 | 5 | 5 |
| 8 | TGME49_311100.P3634 | bradyzoite formation deficient protein BFD2 | 10 | 10 | 7 | 3 | 3 | 3 | 3 | 5 |
| 14 | TGME49_311390-t26_1-p1 | tRNA (guanine(9)-N(1))-methyltransferase | 9 | 8 | 7 | 3 | 5 | 5 | 4 | 5 |
| 19 | TGME49_311450.P3649 | zinc finger, c2h2 type domain-containing protein | 9 | 8 | 8 | 5 | 4 | 5 | 5 | 5 |
| 15 | TGME49_311700-t26_1-p1 | hypothetical protein | 10 | 10 | 7 | 5 | 6 | 3 | 3 | 5 |
| 13 | TGME49_311740-t26_1-p1 | hypothetical protein | 10 | 9 | 7 | 5 | 4 | 3 | 4 | 5 |
| 13 | TGME49_311810.P3656 | hypothetical protein | 10 | 9 | 7 | 5 | 4 | 3 | 3 | 5 |
| 8 | TGME49_311860.P3658 | DUF789 domain-containing protein | 10 | 10 | 6 | 4 | 4 | 4 | 4 | 5 |
| 8 | TGME49_311890.P3659 | hypothetical protein | 9 | 8 | 7 | 3 | 3 | 3 | 3 | 5 |
| 15 | TGME49_311920-t26_1-p1 | GRAM domain-containing protein | 10 | 10 | 7 | 6 | 5 | 5 | 4 | 5 |
| 19 | TGME49_312140-t26_1-p1 | hypothetical protein | 10 | 9 | 9 | 4 | 5 | 7 | 6 | 5 |
| 15 | TGME49_312260-t26_1-p1 | hypothetical protein | 10 | 9 | 8 | 6 | 5 | 4 | 4 | 5 |
| 15 | TGME49_312330.P3667 | cyst wall protein CST10 | 10 | 10 | 9 | 5 | 3 | 3 | 3 | 5 |
| 13 | TGME49_312350-t26_1-p1 | IMPACT, putative | 9 | 8 | 7 | 5 | 3 | 3 | 3 | 5 |
| 8 | TGME49_312360.P3668 | hypothetical protein | 10 | 8 | 7 | 3 | 4 | 4 | 4 | 5 |
| 8 | TGME49_312370-t26_1-p1 | RNA pseudouridine synthase superfamily protein | 10 | 9 | 8 | 4 | 3 | 3 | 3 | 5 |
| 19 | TGME49_312380-t26_1-p1 | tetratricopeptide repeat-containing protein | 9 | 9 | 9 | 5 | 5 | 6 | 5 | 5 |
| 14 | TGME49_312430.P3671 | mitochondrial inner membrane protein OXA1, putative | 9 | 9 | 8 | 5 | 6 | 5 | 5 | 5 |
| 15 | TGME49_312560-t26_1-p1 | eukaryotic initiation factor eIF4E3 | 10 | 10 | 8 | 6 | 5 | 4 | 4 | 5 |
| 19 | TGME49_312660.P3680 | hydrolase, putative | 9 | 9 | 8 | 3 | 3 | 6 | 6 | 5 |
| 15 | TGME49_312700-t26_1-p1 | hypothetical protein | 10 | 10 | 9 | 6 | 5 | 4 | 3 | 5 |
| 8 | TGME49_312820-t26_1-p1 | hypothetical protein | 9 | 8 | 6 | 4 | 4 | 3 | 3 | 5 |
| 15 | TGME49_312840-t26_1-p1 | hypothetical protein | 10 | 9 | 8 | 6 | 6 | 4 | 4 | 5 |
| 8 | TGME49_312875-t26_1-p1 | hypothetical protein | 10 | 8 | 7 | 4 | 4 | 3 | 3 | 5 |
| 15 | TGME49_313040.P3689 | hypothetical protein | 9 | 9 | 8 | 4 | 6 | 3 | 4 | 5 |
| 8 | TGME49_313220.P3695 | hypothetical protein | 10 | 9 | 6 | 3 | 3 | 4 | 4 | 5 |
| 15 | TGME49_313270.P3696 | hypothetical protein | 10 | 10 | 9 | 6 | 4 | 4 | 3 | 5 |
| 15 | TGME49_313300-t26_1-p1 | YL1 nuclear protein C-terminal domain-containing protein | 10 | 9 | 8 | 6 | 5 | 4 | 4 | 5 |
| 14 | TGME49_313350-t26_1-p1 | WD repeat-containing protein | 9 | 7 | 8 | 3 | 5 | 5 | 4 | 5 |
| 15 | TGME49_313470.P3708 | helix-hairpin-helix motif domain-containing protein | 10 | 9 | 8 | 6 | 5 | 3 | 4 | 5 |
| 8 | TGME49_313495.P3710 | THO complex subunit 2, putative | 10 | 9 | 8 | 4 | 4 | 3 | 3 | 5 |
| 8 | TGME49_313540-t26_1-p1 | hypothetical protein | 10 | 9 | 7 | 4 | 4 | 3 | 3 | 5 |
| 13 | TGME49_313680-t26_1-p1 | hypothetical protein | 9 | 8 | 7 | 5 | 3 | 3 | 4 | 5 |
| 13 | TGME49_313760.P3721 | RING zinc finger protein | 10 | 9 | 6 | 5 | 4 | 3 | 3 | 5 |
| 8 | TGME49_313770-t26_1-p1 | hypothetical protein | 10 | 9 | 8 | 4 | 3 | 3 | 3 | 5 |
| 13 | TGME49_313810-t26_1-p1 | transcription initiation factor TFIID complex subunit TAF8 | 9 | 8 | 7 | 6 | 4 | 4 | 4 | 5 |

| | | | | | | | | | | |
|---|---|---|---|---|---|---|---|---|---|---|
| 13 | TGME49_313870-t26_1-p1 | zinc finger, C3HC4 type (RING finger) domain-containing protein | 10 | 8 | 8 | 5 | 4 | 3 | 4 | 5 |
| 8 | TGME49_313910.P3726 | RNA recognition motif 2 protein | 10 | 10 | 7 | 3 | 3 | 3 | 3 | 5 |
| 14 | TGME49_313930.P3727 | chloroquine resistance transporter CRT | 10 | 9 | 8 | 4 | 6 | 4 | 6 | 5 |
| 19 | TGME49_313970-t26_1-p1 | Phytanoyl-CoA dioxygenase (PhyH) superfamily protein | 10 | 8 | 9 | 5 | 4 | 6 | 5 | 5 |
| 19 | TGME49_313990-t26_1-p1 | CBF/Mak21 family protein | 10 | 10 | 10 | 5 | 3 | 6 | 4 | 5 |
| 8 | TGME49_314020-t26_1-p1 | hypothetical protein | 10 | 9 | 7 | 4 | 3 | 3 | 3 | 5 |
| 15 | TGME49_314030.P3729 | hypothetical protein | 10 | 9 | 8 | 4 | 5 | 5 | 4 | 5 |
| 8 | TGME49_314110-t26_1-p1 | hypothetical protein | 10 | 10 | 9 | 3 | 3 | 3 | 3 | 5 |
| 8 | TGME49_314220.P3733 | hypothetical protein | 10 | 7 | 7 | 4 | 4 | 3 | 3 | 5 |
| 14 | TGME49_314710-t26_1-p1 | carrier superfamily protein | 9 | 8 | 7 | 5 | 7 | 5 | 5 | 5 |
| 8 | TGME49_314840.P3750 | ubiquitin carboxyl-terminal hydrolase | 10 | 9 | 6 | 4 | 3 | 4 | 4 | 5 |
| 8 | TGME49_314955-t26_1-p1 | DNA ligase (NAD+), putative | 10 | 8 | 8 | 3 | 4 | 4 | 4 | 5 |
| 14 | TGME49_315130.P3755 | L-isoaspartyl protein carboxyl methyltransferase family protein | 8 | 7 | 7 | 3 | 6 | 6 | 6 | 5 |
| 8 | TGME49_315150-t26_1-p1 | eukaryotic initiation factor eIF4E2 | 9 | 8 | 7 | 4 | 3 | 3 | 3 | 5 |
| 14 | TGME49_315160.P3756 | rhoptry apical surface protein RASP2 | 9 | 3 | 8 | 3 | 8 | 7 | 3 | 5 |
| 19 | TGME49_315190-t26_1-p1 | CAM kinase, SNF1 family | 10 | 10 | 9 | 4 | 3 | 8 | 5 | 5 |
| 15 | TGME49_315210.P3759 | rhoptry protein, putative | 10 | 10 | 10 | 3 | 4 | 4 | 3 | 5 |
| 8 | TGME49_315230-t26_1-p1 | hypothetical protein | 10 | 9 | 8 | 3 | 3 | 3 | 3 | 5 |
| 13 | TGME49_315240-t26_1-p1 | hypothetical protein | 9 | 8 | 7 | 5 | 4 | 3 | 3 | 5 |
| 15 | TGME49_315280-t26_1-p1 | hypothetical protein | 10 | 9 | 7 | 5 | 4 | 4 | 4 | 5 |
| 8 | TGME49_315290-t26_1-p1 | hypothetical protein | 10 | 9 | 7 | 4 | 3 | 3 | 3 | 5 |
| 13 | TGME49_315310.P3762 | hypothetical protein | 10 | 8 | 7 | 6 | 4 | 4 | 3 | 5 |
| 13 | TGME49_315420.P3768 | hypothetical protein | 10 | 9 | 8 | 5 | 4 | 3 | 3 | 5 |
| 15 | TGME49_315450.P3769 | hypothetical protein | 9 | 9 | 9 | 3 | 6 | 3 | 3 | 5 |
| 13 | TGME49_315510.P3775 | apical polar ring protein APR1 | 9 | 8 | 7 | 6 | 3 | 3 | 4 | 5 |
| 8 | TGME49_315520.P3776 | calcium-binding egf domain-containing protein CBDP1 | 9 | 8 | 7 | 4 | 3 | 3 | 3 | 5 |
| 8 | TGME49_315550.P3779 | calcium-binding egf domain-containing protein CBDP4 | 8 | 7 | 7 | 3 | 4 | 3 | 3 | 5 |
| 13 | TGME49_315660-t26_1-p1 | hypothetical protein | 10 | 9 | 7 | 6 | 3 | 4 | 3 | 5 |
| 19 | TGME49_315680-t26_1-p1 | vacuolar protein sorting-associated protein vps4, putative | 9 | 8 | 9 | 4 | 3 | 8 | 7 | 5 |
| 15 | TGME49_315760-t26_1-p1 | AP2 domain transcription factor AP2XI-4 | 10 | 10 | 8 | 5 | 5 | 5 | 4 | 5 |
| 8 | TGME49_315900.P3792 | hypothetical protein | 10 | 9 | 8 | 3 | 3 | 3 | 3 | 5 |
| 8 | TGME49_315940.P3795 | rhoptry protein, putative | 10 | 10 | 9 | 3 | 3 | 3 | 3 | 5 |
| 15 | TGME49_315962.P3797 | hypothetical protein | 10 | 10 | 9 | 3 | 8 | 3 | 3 | 5 |
| 13 | TGME49_316120-t26_1-p1 | hypothetical protein | 10 | 8 | 6 | 5 | 3 | 4 | 4 | 5 |
| 13 | TGME49_316130-t26_1-p1 | hypothetical protein | 9 | 8 | 7 | 6 | 3 | 3 | 4 | 5 |
| 13 | TGME49_316150-t26_1-p1 | ULK kinase | 10 | 8 | 8 | 7 | 3 | 3 | 3 | 5 |
| 13 | TGME49_316190-t26_1-p1 | superoxide dismutase SOD3 | 8 | 7 | 7 | 7 | 3 | 5 | 3 | 5 |
| 13 | TGME49_316260.P3802 | hypothetical protein | 10 | 9 | 8 | 6 | 3 | 3 | 3 | 5 |
| 8 | TGME49_316320.P3806 | ABC transporter transmembrane region domain-containing protein | 10 | 7 | 9 | 3 | 3 | 3 | 3 | 5 |
| 13 | TGME49_316350-t26_1-p1 | hypothetical protein | 8 | 8 | 7 | 5 | 3 | 4 | 3 | 5 |
| 8 | TGME49_316370-t26_1-p1 | hypothetical protein | 10 | 9 | 8 | 3 | 4 | 3 | 3 | 5 |
| 13 | TGME49_316490-t26_1-p1 | hypothetical protein | 9 | 8 | 7 | 6 | 4 | 3 | 4 | 5 |
| 15 | TGME49_316570.P3813 | hypothetical protein | 10 | 8 | 8 | 5 | 4 | 5 | 4 | 5 |

| | | | | | | | | | | |
|---|---|---|---|---|---|---|---|---|---|---|
| 15 | TGME49_316610.P3815 | hypothetical protein | 10 | 8 | 9 | 5 | 5 | 3 | 3 | 5 |
| 19 | TGME49_316635-t26_1-p1 | hypothetical protein | 10 | 8 | 7 | 5 | 3 | 6 | 6 | 5 |
| 14 | TGME49_316770-t26_1-p1 | undecaprenyl diphosphate synthase | 9 | 6 | 7 | 4 | 5 | 5 | 4 | 5 |
| 13 | TGME49_316890.P3818 | hypothetical oocyst wall protein HOWP1 | 9 | 8 | 8 | 7 | 3 | 4 | 4 | 5 |
| 8 | TGME49_317705-t26_1-p1 | enoyl-CoA hydratase/isomerase family protein | 9 | 6 | 7 | 3 | 3 | 3 | 3 | 5 |
| 19 | TGME49_318240.P3825 | Tubulin-tyrosine ligase family protein | 10 | 10 | 10 | 3 | 3 | 7 | 6 | 5 |
| 19 | TGME49_318350.P3828 | Pumilio-family RNA binding repeat-containing protein | 10 | 10 | 9 | 3 | 3 | 5 | 7 | 5 |
| 19 | TGME49_318470-t26_1-p1 | AP2 domain transcription factor AP2IV-4 | 10 | 10 | 8 | 4 | 3 | 5 | 4 | 5 |
| 13 | TGME49_318610.P3839 | AP2 domain transcription factor AP2IV-3 | 10 | 9 | 7 | 5 | 3 | 3 | 3 | 5 |
| 14 | TGME49_318690-t26_1-p1 | RNA recognition motif-containing protein | 10 | 9 | 9 | 3 | 7 | 3 | 7 | 5 |
| 14 | TGME49_318770-t26_1-p1 | aurora-related kinase ARK2 | 10 | 8 | 8 | 4 | 6 | 3 | 5 | 5 |
| 15 | TGME49_319300.P3845 | glycosylphosphatidylinositol anchor attachment 1 protein, putative | 10 | 10 | 10 | 4 | 5 | 5 | 4 | 5 |
| 8 | TGME49_319390.P3848 | hypothetical protein | 10 | 9 | 7 | 3 | 3 | 3 | 3 | 5 |
| 8 | TGME49_319530-t26_1-p1 | serine/arginine-rich splicing factor SR1 | 9 | 7 | 7 | 4 | 4 | 3 | 3 | 5 |
| 13 | TGME49_319540-t26_1-p1 | EF hand domain-containing protein | 9 | 8 | 7 | 6 | 3 | 4 | 4 | 5 |
| 13 | TGME49_319600-t26_1-p1 | alpha-tubulin N-acetyltransferase, putative | 10 | 7 | 6 | 5 | 3 | 3 | 3 | 5 |
| 19 | TGME49_319680-t26_1-p1 | hypothetical protein | 9 | 8 | 7 | 3 | 3 | 5 | 5 | 5 |
| 8 | TGME49_320010-t26_1-p1 | hypothetical protein | 10 | 9 | 7 | 3 | 3 | 4 | 4 | 5 |
| 8 | TGME49_320030-t26_1-p1 | hypothetical protein | 10 | 8 | 6 | 3 | 3 | 4 | 4 | 5 |
| 13 | TGME49_320090-t26_1-p1 | hypothetical protein | 9 | 9 | 7 | 5 | 4 | 3 | 3 | 5 |
| 13 | TGME49_320100-t26_1-p1 | RNA recognition motif-containing protein | 10 | 9 | 6 | 6 | 5 | 3 | 3 | 5 |
| 15 | TGME49_320160-t26_1-p1 | hypothetical protein | 10 | 9 | 8 | 5 | 5 | 4 | 4 | 5 |
| 19 | TGME49_320260-t26_1-p1 | hypothetical protein | 10 | 9 | 6 | 3 | 3 | 5 | 5 | 5 |
| 13 | TGME49_320540-t26_1-p1 | hypothetical protein | 8 | 8 | 7 | 6 | 3 | 3 | 3 | 5 |
| 13 | TGME49_320588.P3883 | glycosyl hydrolases family 35 protein | 10 | 10 | 8 | 6 | 3 | 4 | 3 | 5 |
| 15 | TGME49_320610.P3885 | SAC3 domain-containing protein | 10 | 9 | 8 | 6 | 5 | 3 | 3 | 5 |
| 14 | TGME49_320630-t26_1-p1 | phosphotransferase enzyme family protein | 9 | 9 | 8 | 3 | 5 | 5 | 5 | 5 |
| 13 | TGME49_320680-t26_1-p1 | AP2 domain transcription factor AP2IV-2 | 10 | 9 | 6 | 6 | 3 | 5 | 3 | 5 |
| 13 | TGME49_320690-t26_1-p1 | gamma-soluble NSF attachment protein, putative | 9 | 8 | 7 | 7 | 3 | 4 | 3 | 5 |
| 15 | TGME49_320750-t26_1-p1 | hypothetical protein | 10 | 10 | 8 | 6 | 5 | 5 | 4 | 5 |
| 13 | TGME49_320760-t26_1-p1 | hypothetical protein | 9 | 8 | 6 | 5 | 3 | 4 | 4 | 5 |
| 14 | TGME49_321290-t26_1-p1 | Prp18 domain-containing protein | 9 | 8 | 8 | 4 | 8 | 3 | 6 | 5 |
| 14 | TGME49_321310-t26_1-p1 | 16S rRNA processing protein RimM domain-containing protein | 10 | 9 | 8 | 3 | 6 | 4 | 5 | 5 |
| 8 | TGME49_321410.P3894 | hypothetical protein | 10 | 9 | 7 | 4 | 4 | 3 | 3 | 5 |
| 13 | TGME49_321600-t26_1-p1 | hypothetical protein | 10 | 8 | 7 | 6 | 4 | 4 | 4 | 5 |
| 13 | TGME49_321610.P3897 | HROB domain-containing protein | 10 | 7 | 7 | 5 | 4 | 4 | 3 | 5 |
| 13 | TGME49_500007.P10 | hypothetical protein, conserved | 9 | 9 | 8 | 5 | 3 | 3 | 3 | 5 |
| 13 | TGME49_500008.P11 | CCCH-type zinc finger protein, putative | 9 | 8 | 7 | 6 | 3 | 3 | 3 | 5 |
| 19 | TGME49_500012.P15 | thrombospondin type 1 domain-containing protein | 7 | 9 | 8 | 5 | 3 | 5 | 4 | 5 |
| 13 | TGME49_500013.P16 | hypothetical protein, conserved | 10 | 9 | 7 | 5 | 4 | 4 | 3 | 5 |
| 8 | TGME49_500017.P20 | hypothetical protein, conserved | 9 | 9 | 8 | 3 | 3 | 3 | 3 | 5 |
| 14 | TGME49_500030.P33 | tubulin-tyrosine ligase, putative | 10 | 9 | 9 | 3 | 8 | 5 | 3 | 5 |
| 8 | TGME49_500039.P2 | hypothetical protein | 9 | 8 | 6 | 4 | 3 | 3 | 3 | 5 |
| 19 | TGME49_500040.P340 | MORN repeat-containing protein | 10 | 9 | 9 | 4 | 3 | 5 | 5 | 5 |
| 15 | TGME49_500042.P342 | AP2 domain transcription factor AP2X-10 | 10 | 9 | 7 | 5 | 5 | 3 | 3 | 5 |
| 15 | TGME49_500082.P383 | armadillo repeat-containing protein | 10 | 9 | 9 | 4 | 3 | 4 | 4 | 5 |

| | | | | | | | | | | |
|---|---|---|---|---|---|---|---|---|---|---|
| 19 | TGME49_500085.P386 | P-type ATPase, putative | 10 | 10 | 7 | 5 | 3 | 5 | 5 | 5 |
| 13 | TGME49_500092.P393 | hypothetical protein, conserved | 10 | 9 | 6 | 5 | 4 | 3 | 3 | 5 |
| 14 | TGME49_500116.P418 | aspartyl/glutamyl-tRNA(Asn/Gln) amidotransferase subunit B, putative | 10 | 8 | 8 | 4 | 5 | 5 | 5 | 5 |
| 15 | TGME49_500124.P426 | anaphase-promoting complex subunit 1 | 10 | 10 | 8 | 5 | 3 | 4 | 4 | 5 |
| 13 | TGME49_500128.P430 | hypothetical protein, conserved | 10 | 7 | 7 | 5 | 3 | 4 | 4 | 5 |
| 8 | TGME49_500144.P446 | hypothetical protein, conserved | 9 | 8 | 8 | 4 | 3 | 3 | 4 | 5 |
| 15 | TGME49_500146.P150 | guanine-nucleotide exchange factor, putative | 10 | 10 | 10 | 5 | 4 | 4 | 3 | 5 |
| 13 | TGME49_500163.P167 | zinc transporter ZIP, putative | 10 | 9 | 7 | 6 | 4 | 4 | 3 | 5 |
| 13 | TGME49_500169.P173 | CFAP99 domain-containing protein | 5 | 10 | 8 | 6 | 3 | 4 | 4 | 5 |
| 19 | TGME49_500181.P185 | calpain family cysteine protease, putative | 10 | 10 | 10 | 3 | 3 | 3 | 6 | 5 |
| 14 | TGME49_500183.P187 | glyoxalase, putative | 9 | 8 | 8 | 4 | 7 | 4 | 4 | 5 |
| 8 | TGME49_500200.P204 | intraconoidal microtubule-associated protein ICMAP3 | 10 | 9 | 6 | 4 | 4 | 3 | 3 | 5 |
| 13 | TGME49_500204.P209 | PIBF1 domain-containing protein | 10 | 7 | 9 | 6 | 3 | 4 | 3 | 5 |
| 19 | TGME49_500210.P215 | leucine rich repeat-containing protein | 7 | 9 | 7 | 5 | 3 | 7 | 3 | 5 |
| 13 | TGME49_500221.P226 | cyclin-dependent kinase regulatory subunit protein | 9 | 8 | 6 | 5 | 3 | 4 | 3 | 5 |
| 8 | TGME49_500232.P238 | PIH1 domain-containing protein | 10 | 7 | 8 | 4 | 3 | 5 | 3 | 5 |
| 19 | TGME49_500237.P243 | WD repeat-containing protein | 10 | 9 | 8 | 4 | 4 | 6 | 4 | 5 |
| 8 | TGME49_500256.P45 | hypothetical protein, conserved | 10 | 8 | 8 | 3 | 3 | 3 | 3 | 5 |
| 13 | TGME49_500276.P65 | Rtr1/RPAP2 family protein | 10 | 9 | 7 | 5 | 3 | 4 | 4 | 5 |
| 13 | TGME49_500284.P73 | hypothetical protein, conserved | 9 | 8 | 8 | 7 | 3 | 3 | 3 | 5 |
| 14 | TGME49_500322.P112 | fatty acyl-AMP ligase, putative | 10 | 10 | 10 | 3 | 7 | 7 | 3 | 5 |
| 19 | TGME49_500325.P115 | WD repeat-containing protein | 10 | 8 | 9 | 5 | 4 | 6 | 6 | 5 |
| 8 | TGME49_500327.P117 | hypothetical protein, conserved | 10 | 8 | 7 | 4 | 4 | 3 | 4 | 5 |
| 8 | TGME49_500339.P129 | thrombospondin type 1 domain-containing protein | 9 | 7 | 8 | 4 | 3 | 3 | 3 | 5 |
| 15 | TGME49_500344.P134 | AP2 domain transcription factor AP2X-6 | 10 | 10 | 7 | 5 | 5 | 5 | 5 | 5 |
| 14 | TGME49_500355.P145 | ankyrin repeat-containing protein | 7 | 8 | 8 | 3 | 5 | 5 | 5 | 5 |
| 13 | TGME49_500362.P261 | hypothetical protein, conserved | 9 | 8 | 6 | 5 | 3 | 3 | 3 | 5 |
| 14 | TGME49_500365.P264 | tRNA (guanine-N1-)-methyltransferase, putative | 8 | 7 | 3 | 3 | 7 | 5 | 5 | 5 |
| 13 | TGME49_500374.P273 | hypothetical protein, conserved | 10 | 10 | 7 | 6 | 3 | 3 | 4 | 5 |
| 13 | TGME49_500384.P283 | F-box domain-containing protein | 9 | 10 | 7 | 6 | 3 | 4 | 4 | 5 |
| 13 | TGME49_500392.P291 | IQ calmodulin-binding motif domain-containing protein | 10 | 9 | 7 | 6 | 3 | 4 | 3 | 5 |
| 8 | TGME49_500393.P292 | hypothetical protein, conserved | 7 | 9 | 6 | 3 | 4 | 4 | 4 | 5 |
| 8 | TGME49_500408.P307 | hypothetical protein, conserved | 9 | 9 | 8 | 4 | 3 | 4 | 4 | 5 |
| 13 | TGME49_500416.P315 | hypothetical protein, conserved | 9 | 7 | 6 | 6 | 3 | 4 | 4 | 5 |
| 8 | TGME49_500421.P320 | hypothetical protein, conserved | 6 | 10 | 10 | 3 | 3 | 3 | 3 | 5 |
| 13 | TGME49_500423.P322 | MYCBP-associated protein, putative | 9 | 8 | 6 | 6 | 3 | 4 | 3 | 5 |
| 8 | TGME49_500439.P338 | hypothetical protein, conserved | 10 | 9 | 6 | 3 | 3 | 3 | 3 | 5 |

| Kmean_cluster | GeneID (all protein TG) | Product Description | Decile_bitHH | Decile_bitNcan | Decile_bitBB | Decilebit_CS | Decilebit_SN3 | Decile bit_Ef | Decile bit_CC | Meta_cluster |
|---|---|---|---|---|---|---|---|---|---|---|
| 4 | TGME49_200270-t26_1-p1 | PAN/Apple domain-containing protein | 7 | 7 | 7 | 4 | 3 | 3 | 3 | 6 |
| 5 | TGME49_200410.P459 | hypothetical protein | 10 | 7 | 5 | 4 | 3 | 3 | 3 | 6 |
| 4 | TGME49_200440.P461 | hypothetical protein | 9 | 7 | 7 | 4 | 4 | 4 | 3 | 6 |
| 5 | TGME49_200450.P462 | hypothetical protein | 9 | 6 | 3 | 4 | 3 | 3 | 3 | 6 |
| 10 | TGME49_201130-t26_1-p1 | rhoptry kinase family protein ROP33 | 10 | 8 | 3 | 7 | 3 | 3 | 4 | 6 |
| 10 | TGME49_201140-t26_1-p1 | hypothetical protein | 10 | 9 | 3 | 7 | 5 | 3 | 3 | 6 |
| 4 | TGME49_201210-t26_1-p1 | hypothetical protein | 8 | 7 | 6 | 5 | 3 | 3 | 3 | 6 |
| 5 | TGME49_201660-t26_1-p1 | hypothetical protein | 9 | 7 | 5 | 4 | 3 | 3 | 3 | 6 |
| 4 | TGME49_201740-t26_1-p1 | hypothetical protein | 8 | 6 | 6 | 4 | 4 | 3 | 4 | 6 |
| 4 | TGME49_201760.P471 | thioredoxin-like associated protein TLAP4 | 8 | 7 | 6 | 4 | 5 | 4 | 4 | 6 |
| 4 | TGME49_201800.P475 | ATP synthase-associated protein | 8 | 7 | 6 | 5 | 5 | 4 | 4 | 6 |
| 5 | TGME49_201820-t26_1-p1 | hypothetical protein | 10 | 7 | 5 | 3 | 4 | 4 | 4 | 6 |
| 4 | TGME49_201830.P476 | ribosomal protein L37 | 9 | 5 | 5 | 5 | 3 | 4 | 4 | 6 |
| 10 | TGME49_201850-t26_1-p1 | WD domain, G-beta repeat-containing protein | 9 | 8 | 3 | 7 | 3 | 5 | 5 | 6 |
| 5 | TGME49_202010-t26_1-p1 | hypothetical protein | 10 | 7 | 3 | 4 | 3 | 3 | 3 | 6 |
| 4 | TGME49_202050.P480 | hypothetical protein | 7 | 8 | 6 | 4 | 3 | 3 | 3 | 6 |
| 10 | TGME49_202210-t26_1-p1 | nucleoside diphosphate kinase | 9 | 8 | 3 | 8 | 5 | 5 | 5 | 6 |
| 4 | TGME49_202420.P488 | hypothetical protein | 8 | 7 | 6 | 5 | 3 | 3 | 3 | 6 |
| 5 | TGME49_202430-t26_1-p1 | hypothetical protein | 9 | 6 | 4 | 4 | 3 | 3 | 3 | 6 |
| 4 | TGME49_202440.P489 | hypothetical protein | 7 | 7 | 6 | 5 | 4 | 4 | 4 | 6 |
| 4 | TGME49_202480-t26_1-p1 | hypothetical protein | 7 | 6 | 6 | 4 | 4 | 4 | 3 | 6 |
| 5 | TGME49_202520-t26_1-p1 | hypothetical protein | 10 | 7 | 5 | 4 | 4 | 3 | 3 | 6 |
| 4 | TGME49_202590-t26_1-p1 | hypothetical protein | 9 | 7 | 6 | 5 | 4 | 5 | 4 | 6 |
| 10 | TGME49_202610.P496 | protein phosphatase PPM3D | 9 | 8 | 3 | 7 | 3 | 3 | 3 | 6 |
| 4 | TGME49_202700-t26_1-p1 | hypothetical protein | 9 | 5 | 6 | 4 | 4 | 3 | 3 | 6 |
| 4 | TGME49_202830.P501 | Elicitor-responsive protein, putative | 7 | 6 | 6 | 5 | 5 | 3 | 3 | 6 |
| 10 | TGME49_202910-t26_1-p1 | zinc carboxypeptidase superfamily protein | 8 | 7 | 3 | 5 | 4 | 4 | 4 | 6 |
| 10 | TGME49_203050-t26_1-p1 | AP2 domain transcription factor AP2VIIa-6 | 10 | 10 | 3 | 5 | 3 | 5 | 5 | 6 |
| 5 | TGME49_203090-t26_1-p1 | hypothetical protein | 9 | 7 | 4 | 3 | 4 | 3 | 3 | 6 |
| 5 | TGME49_203150-t26_1-p1 | hypothetical protein | 10 | 9 | 5 | 4 | 4 | 3 | 3 | 6 |
| 4 | TGME49_203230-t26_1-p1 | hypothetical protein | 7 | 6 | 6 | 5 | 4 | 3 | 3 | 6 |
| 4 | TGME49_203280-t26_1-p1 | hypothetical protein | 7 | 6 | 6 | 5 | 5 | 5 | 3 | 6 |
| 4 | TGME49_203362-t26_1-p1 | prefoldin, putative | 7 | 7 | 6 | 6 | 4 | 4 | 3 | 6 |
| 4 | TGME49_203400.P521 | hypothetical protein | 8 | 6 | 6 | 4 | 4 | 4 | 4 | 6 |
| 5 | TGME49_203590-t26_1-p1 | hypothetical protein | 9 | 7 | 4 | 5 | 3 | 3 | 3 | 6 |
| 4 | TGME49_203600.P527 | dense granule protein GRA50 | 8 | 7 | 6 | 5 | 4 | 3 | 3 | 6 |
| 10 | TGME49_203730.P531 | hypothetical protein | 10 | 10 | 3 | 8 | 7 | 5 | 4 | 6 |
| 4 | TGME49_203860-t26_1-p1 | hypothetical protein | 9 | 7 | 6 | 5 | 4 | 4 | 3 | 6 |
| 10 | TGME49_203880.P536 | hypothetical protein | 10 | 10 | 5 | 6 | 4 | 4 | 3 | 6 |
| 4 | TGME49_203890-t26_1-p1 | hypothetical protein | 8 | 7 | 5 | 5 | 3 | 3 | 3 | 6 |
| 10 | TGME49_204040-t26_1-p1 | hypothetical protein | 10 | 9 | 3 | 5 | 3 | 3 | 3 | 6 |
| 4 | TGME49_204120.P542 | pre-mRNA-splicing factor CWC25, putative | 9 | 6 | 6 | 5 | 5 | 4 | 4 | 6 |
| 4 | TGME49_204530.P557 | microneme protein MIC11 | 7 | 6 | 6 | 5 | 3 | 3 | 3 | 6 |
| 5 | TGME49_204880-t26_1-p1 | hypothetical protein | 10 | 8 | 5 | 4 | 4 | 4 | 4 | 6 |
| 10 | TGME49_205060-t26_1-p1 | hypothetical protein | 9 | 8 | 3 | 6 | 3 | 3 | 3 | 6 |
| 5 | TGME49_205100.P560 | hypothetical protein | 9 | 6 | 6 | 3 | 4 | 3 | 3 | 6 |
| 5 | TGME49_205120-t26_1-p1 | hypothetical protein | 10 | 6 | 5 | 4 | 4 | 4 | 4 | 6 |
| 4 | TGME49_205390.P568 | hypothetical protein | 8 | 7 | 6 | 5 | 4 | 3 | 3 | 6 |
| 4 | TGME49_205460-t26_1-p1 | AN1 family Zinc finger domain-containing protein | 7 | 7 | 6 | 6 | 4 | 5 | 3 | 6 |

| | | | | | | | | | | |
|---|---|---|---|---|---|---|---|---|---|---|
| 4 | TGME49_205480-t26_1-p1 | IMC subcompartment protein ISP4 | 7 | 6 | 5 | 5 | 4 | 3 | 3 | 6 |
| 4 | TGME49_205600-t26_1-p1 | zinc finger, C3HC4 type (RING finger) domain-containing protein | 9 | 6 | 6 | 6 | 5 | 4 | 3 | 6 |
| 5 | TGME49_205615.P576 | hypothetical protein | 10 | 9 | 5 | 4 | 4 | 3 | 3 | 6 |
| 4 | TGME49_206320-t26_1-p1 | hypothetical protein | 8 | 7 | 6 | 3 | 4 | 5 | 3 | 6 |
| 5 | TGME49_206540.P589 | hypothetical protein | 9 | 7 | 5 | 4 | 4 | 3 | 3 | 6 |
| 4 | TGME49_207070.P604 | UDP-N-acetylglucosamine transferase subunit ALG14, putative | 8 | 6 | 6 | 4 | 3 | 5 | 4 | 6 |
| 5 | TGME49_207230-t26_1-p1 | hypothetical protein | 10 | 6 | 4 | 3 | 3 | 3 | 3 | 6 |
| 10 | TGME49_207370.P613 | golgi localizing protein GLP3 | 10 | 10 | 4 | 6 | 3 | 4 | 3 | 6 |
| 5 | TGME49_207740-t26_1-p1 | hypothetical protein | 10 | 9 | 4 | 3 | 3 | 3 | 3 | 6 |
| 4 | TGME49_207930.P631 | phosphatidylethanolamine-binding protein | 7 | 6 | 6 | 5 | 4 | 5 | 3 | 6 |
| 4 | TGME49_207980-t26_1-p1 | PIG-P protein | 8 | 7 | 5 | 5 | 3 | 4 | 3 | 6 |
| 4 | TGME49_208010.P635 | syntaxin-19 | 9 | 6 | 6 | 5 | 4 | 4 | 3 | 6 |
| 4 | TGME49_208090.P637 | 5-formyltetrahydrofolate cyclo-ligase | 8 | 5 | 6 | 3 | 4 | 4 | 4 | 6 |
| 10 | TGME49_208340-t26_1-p1 | PH domain-containing protein | 10 | 10 | 4 | 8 | 6 | 5 | 4 | 6 |
| 4 | TGME49_208350.P638 | RNA-binding protein, putative | 8 | 7 | 6 | 5 | 3 | 4 | 4 | 6 |
| 4 | TGME49_208440.P643 | ATP synthase-associated protein | 7 | 6 | 6 | 5 | 5 | 4 | 3 | 6 |
| 10 | TGME49_208810.P656 | peptidyl-prolyl cis-trans isomerase, putative | 10 | 9 | 3 | 5 | 3 | 4 | 3 | 6 |
| 5 | TGME49_208830.P658 | dense granule protein GRA16 | 9 | 6 | 3 | 3 | 3 | 3 | 3 | 6 |
| 5 | TGME49_208910-t26_1-p1 | hypothetical protein | 10 | 8 | 5 | 4 | 3 | 3 | 3 | 6 |
| 4 | TGME49_208970-t26_1-p1 | RNA recognition motif-containing protein | 7 | 7 | 6 | 5 | 5 | 4 | 3 | 6 |
| 4 | TGME49_208990-t26_1-p1 | hypothetical protein | 8 | 7 | 6 | 5 | 4 | 3 | 4 | 6 |
| 4 | TGME49_209090.P667 | proteasome maturation factor ump1 protein | 7 | 6 | 6 | 5 | 5 | 3 | 4 | 6 |
| 5 | TGME49_209270.P674 | zinc finger (CCCH type) motif-containing protein | 10 | 9 | 4 | 4 | 3 | 3 | 3 | 6 |
| 4 | TGME49_209420-t26_1-p1 | hypothetical protein | 8 | 6 | 5 | 5 | 4 | 3 | 3 | 6 |
| 5 | TGME49_209440.P678 | hypothetical protein | 10 | 8 | 4 | 3 | 3 | 3 | 3 | 6 |
| 5 | TGME49_209510.P681 | hypothetical protein | 10 | 6 | 5 | 4 | 3 | 3 | 3 | 6 |
| 5 | TGME49_209560.P683 | hypothetical protein | 9 | 7 | 3 | 3 | 3 | 3 | 3 | 6 |
| 4 | TGME49_209590-t26_1-p1 | hypothetical protein | 7 | 7 | 6 | 5 | 3 | 3 | 3 | 6 |
| 10 | TGME49_209620.P686 | aspartyl protease ASP4 | 9 | 8 | 5 | 7 | 3 | 4 | 3 | 6 |
| 4 | TGME49_209630.P687 | coiled-coil domain-containing protein | 8 | 5 | 5 | 5 | 4 | 3 | 3 | 6 |
| 4 | TGME49_209700.P691 | hypothetical protein | 7 | 6 | 5 | 5 | 4 | 3 | 4 | 6 |
| 4 | TGME49_209850-t26_1-p1 | RNA recognition motif-containing protein | 8 | 6 | 6 | 5 | 3 | 3 | 4 | 6 |
| 5 | TGME49_210210.P704 | hypothetical protein | 10 | 10 | 3 | 3 | 3 | 3 | 3 | 6 |
| 5 | TGME49_210370.P708 | rhoptry protein ROP54 | 8 | 7 | 5 | 3 | 3 | 3 | 3 | 6 |
| 5 | TGME49_210400-t26_1-p1 | hypothetical protein | 10 | 9 | 5 | 3 | 3 | 3 | 3 | 6 |
| 4 | TGME49_210420-t26_1-p1 | hypothetical protein | 8 | 5 | 5 | 5 | 3 | 3 | 4 | 6 |
| 5 | TGME49_210450-t26_1-p1 | hypothetical protein | 9 | 7 | 3 | 3 | 3 | 3 | 3 | 6 |
| 10 | TGME49_211010-t26_1-p1 | SANT/Myb domain-containing protein | 10 | 8 | 3 | 5 | 4 | 4 | 4 | 6 |
| 4 | TGME49_211250.P728 | hypothetical protein | 8 | 7 | 6 | 5 | 3 | 3 | 3 | 6 |
| 4 | TGME49_211300-t26_1-p1 | DUF74 family protein, putative | 8 | 7 | 5 | 5 | 3 | 3 | 4 | 6 |
| 5 | TGME49_211340-t26_1-p1 | hypothetical protein | 10 | 7 | 4 | 4 | 3 | 3 | 3 | 6 |
| 5 | TGME49_211460.P741 | Myc-regulation protein MYR4 | 10 | 9 | 5 | 4 | 3 | 3 | 3 | 6 |
| 4 | TGME49_211610-t26_1-p1 | hypothetical protein | 10 | 6 | 6 | 4 | 4 | 4 | 4 | 6 |
| 5 | TGME49_211860.P750 | hypothetical protein | 9 | 6 | 3 | 3 | 3 | 3 | 3 | 6 |
| 4 | TGME49_211870.P751 | LSU ribosomal protein L18P, putative | 9 | 7 | 5 | 5 | 5 | 3 | 3 | 6 |
| 10 | TGME49_212030-t26_1-p1 | hypothetical protein | 7 | 8 | 3 | 6 | 3 | 3 | 3 | 6 |
| 5 | TGME49_212170-t26_1-p1 | GIY-YIG catalytic domain-containing protein | 10 | 7 | 5 | 5 | 3 | 4 | 4 | 6 |
| 4 | TGME49_212180.P755 | hypothetical protein | 8 | 7 | 6 | 3 | 4 | 3 | 3 | 6 |

| | | | | | | | | | | |
|---|---|---|---|---|---|---|---|---|---|---|
| 5 | TGME49_212190.P756 | RNA recognition motif-containing protein | 10 | 6 | 5 | 4 | 5 | 3 | 3 | 6 |
| 4 | TGME49_212260.P760 | zinc finger protein ZFP2 | 8 | 7 | 6 | 3 | 4 | 4 | 3 | 6 |
| 10 | TGME49_212270.P761 | CLAMP-linked invasion protein CLIP | 9 | 8 | 5 | 6 | 3 | 4 | 3 | 6 |
| 5 | TGME49_212735-t26_1-p1 | transporter, major facilitator family protein | 10 | 7 | 5 | 5 | 4 | 4 | 3 | 6 |
| 5 | TGME49_212810-t26_1-p1 | hypothetical protein | 10 | 8 | 4 | 3 | 3 | 3 | 3 | 6 |
| 4 | TGME49_212850-t26_1-p1 | PI31 domain-containing protein | 9 | 7 | 6 | 4 | 3 | 3 | 4 | 6 |
| 4 | TGME49_212920-t26_1-p1 | hypothetical protein | 7 | 6 | 5 | 5 | 4 | 4 | 4 | 6 |
| 10 | TGME49_213020-t26_1-p1 | RING zinc finger protein | 10 | 8 | 5 | 6 | 4 | 3 | 3 | 6 |
| 4 | TGME49_213030-t26_1-p1 | clathrin light chain, putative | 8 | 7 | 6 | 5 | 4 | 4 | 3 | 6 |
| 4 | TGME49_213040.P777 | centromere protein CENP-C | 10 | 6 | 6 | 5 | 5 | 3 | 3 | 6 |
| 4 | TGME49_213100-t26_1-p1 | hypothetical protein | 8 | 5 | 5 | 4 | 5 | 3 | 4 | 6 |
| 5 | TGME49_213255.P784 | hypothetical protein | 10 | 6 | 4 | 3 | 3 | 3 | 3 | 6 |
| 4 | TGME49_213400-t26_1-p1 | zinc finger (CCCH type) motif-containing protein | 9 | 7 | 6 | 4 | 4 | 4 | 4 | 6 |
| 5 | TGME49_213480.P793 | hypothetical protein | 10 | 8 | 3 | 3 | 3 | 3 | 3 | 6 |
| 4 | TGME49_213490-t26_1-p1 | hypothetical protein | 8 | 6 | 5 | 5 | 4 | 3 | 4 | 6 |
| 10 | TGME49_213550.P795 | palmitoyltransferase DHHC4 | 10 | 8 | 3 | 8 | 3 | 7 | 3 | 6 |
| 4 | TGME49_213570-t26_1-p1 | hypothetical protein | 7 | 6 | 5 | 6 | 3 | 3 | 3 | 6 |
| 4 | TGME49_213752.P805 | herpesviridae ul52/ul70 dna primase | 10 | 6 | 6 | 5 | 4 | 3 | 4 | 6 |
| 5 | TGME49_213920-t26_1-p1 | hypothetical protein | 10 | 7 | 5 | 4 | 3 | 3 | 3 | 6 |
| 10 | TGME49_214120-t26_1-p1 | hypothetical protein | 10 | 9 | 3 | 5 | 5 | 3 | 3 | 6 |
| 4 | TGME49_214170.P817 | hypothetical protein | 10 | 7 | 6 | 4 | 4 | 4 | 4 | 6 |
| 5 | TGME49_214190-t26_1-p1 | SAG-related sequence SRS46 | 8 | 7 | 6 | 3 | 3 | 3 | 3 | 6 |
| 4 | TGME49_214340-t26_1-p1 | hypothetical protein | 8 | 7 | 6 | 3 | 5 | 3 | 4 | 6 |
| 5 | TGME49_214440-t26_1-p1 | 4'-phosphopantetheinyl transferase superfamily protein | 9 | 7 | 5 | 3 | 4 | 3 | 3 | 6 |
| 4 | TGME49_214540.P828 | hypothetical protein | 8 | 7 | 5 | 4 | 4 | 4 | 3 | 6 |
| 5 | TGME49_214630-t26_1-p1 | hypothetical protein | 10 | 6 | 5 | 3 | 3 | 3 | 3 | 6 |
| 5 | TGME49_214740-t26_1-p1 | hypothetical protein | 10 | 8 | 5 | 3 | 3 | 3 | 3 | 6 |
| 4 | TGME49_214780-t26_1-p1 | hydrolase, NUDIX family protein | 7 | 6 | 5 | 5 | 4 | 3 | 3 | 6 |
| 4 | TGME49_214870-t26_1-p1 | ribosomal protein L9, N-terminal domain-containing protein | 9 | 7 | 7 | 5 | 4 | 3 | 4 | 6 |
| 5 | TGME49_214880.P835 | apical cap protein AC4 | 10 | 8 | 5 | 4 | 3 | 3 | 3 | 6 |
| 10 | TGME49_214900-t26_1-p1 | hypothetical protein | 10 | 9 | 3 | 6 | 3 | 3 | 3 | 6 |
| 4 | TGME49_214940.P838 | MIC2-associated protein M2AP | 8 | 6 | 5 | 5 | 4 | 3 | 3 | 6 |
| 4 | TGME49_215070.P845 | ferredoxin FD | 7 | 6 | 6 | 5 | 5 | 5 | 3 | 6 |
| 5 | TGME49_215140-t26_1-p1 | hypothetical protein | 8 | 7 | 6 | 3 | 3 | 3 | 3 | 6 |
| 5 | TGME49_215360.P853 | dense granule protein GRA62 | 9 | 7 | 5 | 3 | 3 | 3 | 3 | 6 |
| 5 | TGME49_215380-t26_1-p1 | hypothetical protein | 10 | 8 | 5 | 3 | 3 | 3 | 3 | 6 |
| 4 | TGME49_215410-t26_1-p1 | DPCD family protein | 7 | 7 | 6 | 5 | 3 | 3 | 3 | 6 |
| 5 | TGME49_215620-t26_1-p1 | hypothetical protein | 10 | 6 | 6 | 4 | 3 | 3 | 3 | 6 |
| 4 | TGME49_215660-t26_1-p1 | hypothetical protein | 9 | 7 | 6 | 5 | 4 | 3 | 3 | 6 |
| 5 | TGME49_215775-t26_1-p1 | rhoptry protein ROP8 | 9 | 7 | 4 | 3 | 3 | 3 | 4 | 6 |
| 5 | TGME49_215895-t26_1-p1 | AP2 domain-containing protein | 10 | 6 | 3 | 5 | 3 | 4 | 4 | 6 |
| 5 | TGME49_215900-t26_1-p1 | hypothetical protein | 10 | 9 | 3 | 3 | 3 | 3 | 3 | 6 |
| 5 | TGME49_216060-t26_1-p1 | hypothetical protein | 10 | 7 | 4 | 3 | 3 | 3 | 3 | 6 |
| 5 | TGME49_216160-t26_1-p1 | hypothetical protein | 9 | 7 | 3 | 3 | 3 | 3 | 3 | 6 |
| 10 | TGME49_216190-t26_1-p1 | hypothetical protein | 10 | 10 | 4 | 6 | 5 | 4 | 4 | 6 |
| 4 | TGME49_216440-t26_1-p1 | OTU domain-containing protein OTU8 | 9 | 6 | 6 | 4 | 3 | 3 | 3 | 6 |
| 5 | TGME49_216470.P886 | hypothetical protein | 9 | 7 | 4 | 3 | 3 | 3 | 3 | 6 |
| 4 | TGME49_216490-t26_1-p1 | hypothetical protein | 7 | 6 | 6 | 5 | 4 | 3 | 3 | 6 |

| | | | | | | | | | | |
|---|---|---|---|---|---|---|---|---|---|---|
| 10 | TGME49_216540.P887 | dolichyl-phosphate beta-glucosyltransferase | 9 | 8 | 4 | 6 | 3 | 3 | 3 | 6 |
| 10 | TGME49_216560.P888 | methyltransferase domain-containing protein | 9 | 7 | 3 | 6 | 3 | 6 | 6 | 6 |
| 5 | TGME49_216580-t26_1-p1 | hypothetical protein | 10 | 9 | 5 | 4 | 3 | 3 | 3 | 6 |
| 10 | TGME49_216600-t26_1-p1 | exonuclease III APE | 9 | 8 | 3 | 7 | 5 | 3 | 6 | 6 |
| 10 | TGME49_216840-t26_1-p1 | hypothetical protein | 10 | 9 | 3 | 6 | 7 | 4 | 4 | 6 |
| 4 | TGME49_216850-t26_1-p1 | hypothetical protein | 8 | 7 | 6 | 3 | 3 | 4 | 4 | 6 |
| 4 | TGME49_216980-t26_1-p1 | WDR8 protein isoform 3 family protein | 9 | 6 | 7 | 4 | 6 | 3 | 3 | 6 |
| 4 | TGME49_216990-t26_1-p1 | LysM domain-containing protein | 8 | 7 | 6 | 4 | 4 | 4 | 4 | 6 |
| 10 | TGME49_217000-t26_1-p1 | hypothetical protein | 9 | 8 | 3 | 7 | 6 | 4 | 4 | 6 |
| 4 | TGME49_217340-t26_1-p1 | hypothetical protein | 9 | 5 | 5 | 4 | 5 | 4 | 4 | 6 |
| 4 | TGME49_217500.P906 | HMG (high mobility group) box domain-containing protein | 8 | 7 | 6 | 5 | 3 | 3 | 3 | 6 |
| 4 | TGME49_217510.P907 | inner membrane complex protein IMC19 | 9 | 7 | 6 | 5 | 5 | 3 | 4 | 6 |
| 4 | TGME49_217550.P910 | dynactin subunit 6, putative | 7 | 6 | 6 | 5 | 4 | 4 | 3 | 6 |
| 4 | TGME49_217590-t26_1-p1 | 50S ribosomal protein L17, putative | 9 | 6 | 6 | 5 | 3 | 5 | 3 | 6 |
| 5 | TGME49_217620.P913 | translation initiation factor IF-1, putative | 9 | 7 | 5 | 3 | 3 | 3 | 4 | 6 |
| 10 | TGME49_217630-t26_1-p1 | hypothetical protein | 10 | 9 | 5 | 6 | 3 | 4 | 4 | 6 |
| 5 | TGME49_217670.P918 | ribosomal protein RPS6 | 10 | 6 | 6 | 3 | 3 | 4 | 4 | 6 |
| 5 | TGME49_217688-t26_1-p1 | hypothetical protein | 10 | 6 | 5 | 4 | 3 | 3 | 3 | 6 |
| 5 | TGME49_217770-t26_1-p1 | hypothetical protein | 10 | 8 | 5 | 4 | 3 | 3 | 3 | 6 |
| 10 | TGME49_217860-t26_1-p1 | hypothetical protein | 10 | 10 | 5 | 5 | 4 | 3 | 3 | 6 |
| 5 | TGME49_218450-t26_1-p1 | hypothetical protein | 9 | 8 | 3 | 3 | 3 | 3 | 3 | 6 |
| 5 | TGME49_218490.P941 | hypothetical protein | 10 | 7 | 5 | 4 | 4 | 4 | 3 | 6 |
| 4 | TGME49_218730.P947 | hypothetical protein | 8 | 7 | 5 | 5 | 3 | 3 | 3 | 6 |
| 4 | TGME49_218740.P948 | membrane protein, putative | 7 | 6 | 6 | 6 | 5 | 3 | 3 | 6 |
| 4 | TGME49_218770.P951 | carrier superfamily protein | 9 | 5 | 6 | 5 | 4 | 4 | 3 | 6 |
| 4 | TGME49_218810-t26_1-p1 | histidyl-tRNA synthetase | 7 | 6 | 6 | 4 | 5 | 3 | 5 | 6 |
| 4 | TGME49_218850-t26_1-p1 | ribosomal protein RPS9 | 8 | 7 | 6 | 3 | 4 | 3 | 4 | 6 |
| 4 | TGME49_219170.P963 | IMC sutures component ISC2 | 8 | 7 | 6 | 5 | 4 | 4 | 4 | 6 |
| 4 | TGME49_219210-t26_1-p1 | ankyrin repeat-containing protein | 8 | 6 | 7 | 6 | 4 | 4 | 4 | 6 |
| 4 | TGME49_219370.P968 | hypothetical protein | 8 | 6 | 6 | 4 | 3 | 3 | 3 | 6 |
| 4 | TGME49_219530.P974 | hypothetical protein | 8 | 6 | 6 | 4 | 3 | 3 | 3 | 6 |
| 5 | TGME49_219678-t26_1-p1 | hypothetical protein | 8 | 8 | 4 | 5 | 4 | 3 | 3 | 6 |
| 4 | TGME49_220090.P987 | DIX domain-containing protein | 7 | 7 | 6 | 5 | 5 | 4 | 4 | 6 |
| 4 | TGME49_220120-t26_1-p1 | AP-5 complex subunit sigma, putative | 8 | 7 | 6 | 4 | 4 | 4 | 4 | 6 |
| 5 | TGME49_220170-t26_1-p1 | coiled-coil domain-containing protein | 9 | 8 | 4 | 4 | 5 | 3 | 4 | 6 |
| 4 | TGME49_220190-t26_1-p1 | hypothetical protein | 9 | 6 | 5 | 4 | 4 | 3 | 4 | 6 |
| 4 | TGME49_220250.P991 | Nucleotide-sensitive chloride conductance regulator (ICln) protein | 7 | 7 | 6 | 5 | 3 | 3 | 4 | 6 |
| 4 | TGME49_220320-t26_1-p1 | BSD domain-containing protein | 8 | 7 | 6 | 5 | 5 | 5 | 3 | 6 |
| 4 | TGME49_220460-t26_1-p1 | SNF7 family protein | 8 | 6 | 6 | 6 | 3 | 4 | 3 | 6 |
| 5 | TGME49_220470-t26_1-p1 | hypothetical protein | 10 | 8 | 4 | 4 | 3 | 3 | 3 | 6 |
| 5 | TGME49_220510-t26_1-p1 | hypothetical protein | 9 | 6 | 5 | 4 | 3 | 3 | 3 | 6 |
| 5 | TGME49_220520-t26_1-p1 | hypothetical protein | 10 | 9 | 4 | 4 | 3 | 4 | 4 | 6 |
| 5 | TGME49_220585-t26_1-p1 | hypothetical protein | 10 | 7 | 4 | 3 | 3 | 3 | 3 | 6 |
| 4 | TGME49_221160.P1010 | acetyltransferase, GNAT family protein | 8 | 5 | 6 | 5 | 5 | 3 | 3 | 6 |
| 4 | TGME49_221350-t26_1-p1 | Ctr copper transporter family protein | 8 | 7 | 6 | 3 | 4 | 3 | 4 | 6 |
| 5 | TGME49_221430-t26_1-p1 | hypothetical protein | 9 | 6 | 4 | 4 | 3 | 3 | 3 | 6 |
| 5 | TGME49_221440-t26_1-p1 | RPGR, putative | 10 | 6 | 5 | 4 | 3 | 3 | 3 | 6 |
| 5 | TGME49_221460-t26_1-p1 | phosphoglycerate mutase family protein | 9 | 7 | 6 | 3 | 3 | 3 | 3 | 6 |

| | | | | | | | | | | |
|---|---|---|---|---|---|---|---|---|---|---|
| 4 | TGME49_221530-t26_1-p1 | hypothetical protein | 9 | 7 | 5 | 5 | 3 | 3 | 4 | 6 |
| 5 | TGME49_221540-t26_1-p1 | DUSP domain-containing protein | 10 | 7 | 3 | 3 | 3 | 3 | 3 | 6 |
| 5 | TGME49_221560.P1028 | hypothetical protein | 9 | 6 | 4 | 3 | 3 | 3 | 3 | 6 |
| 4 | TGME49_221570.P1029 | pre-rRNA-processing protein, putative | 8 | 7 | 6 | 3 | 5 | 3 | 4 | 6 |
| 10 | TGME49_221580.P1030 | ribosomal RNA large subunit methyltransferase J protein | 10 | 9 | 3 | 6 | 4 | 5 | 4 | 6 |
| 10 | TGME49_221640.P1034 | hypothetical protein | 10 | 8 | 5 | 5 | 5 | 4 | 4 | 6 |
| 5 | TGME49_221840.P1039 | hypothetical protein | 9 | 8 | 3 | 3 | 3 | 3 | 3 | 6 |
| 4 | TGME49_221900-t26_1-p1 | hypothetical protein | 7 | 5 | 6 | 5 | 4 | 3 | 3 | 6 |
| 4 | TGME49_221910.P1041 | AN1 family Zinc finger domain-containing protein | 7 | 6 | 6 | 5 | 5 | 4 | 3 | 6 |
| 4 | TGME49_221922-t26_1-p1 | NifU family domain-containing protein | 9 | 7 | 6 | 6 | 4 | 4 | 4 | 6 |
| 4 | TGME49_221980-t26_1-p1 | U1 zinc finger protein | 9 | 5 | 5 | 5 | 4 | 4 | 4 | 6 |
| 5 | TGME49_221990-t26_1-p1 | hypothetical protein | 9 | 7 | 6 | 3 | 3 | 3 | 3 | 6 |
| 4 | TGME49_222040-t26_1-p1 | Ran-interacting Mog1 protein | 7 | 6 | 6 | 5 | 4 | 3 | 4 | 6 |
| 4 | TGME49_222170.P1051 | dense granule protein GRA17 | 8 | 6 | 5 | 5 | 3 | 4 | 4 | 6 |
| 5 | TGME49_222240.P1055 | hypothetical protein | 10 | 7 | 4 | 3 | 3 | 3 | 3 | 6 |
| 5 | TGME49_222330-t26_1-p1 | hypothetical protein | 10 | 9 | 4 | 3 | 4 | 3 | 4 | 6 |
| 4 | TGME49_222340-t26_1-p1 | NOL1/NOP2/sun family protein | 9 | 7 | 7 | 4 | 4 | 3 | 5 | 6 |
| 4 | TGME49_222930-t26_1-p1 | hypothetical protein | 8 | 7 | 6 | 5 | 3 | 4 | 3 | 6 |
| 10 | TGME49_223080.P1074 | hypothetical protein | 10 | 10 | 4 | 6 | 3 | 3 | 4 | 6 |
| 10 | TGME49_223150-t26_1-p1 | START domain-containing protein | 9 | 9 | 3 | 8 | 5 | 6 | 5 | 6 |
| 4 | TGME49_223420.P1077 | DnaJ domain-containing protein | 8 | 7 | 5 | 5 | 4 | 3 | 4 | 6 |
| 4 | TGME49_223490-t26_1-p1 | hypothetical protein | 8 | 6 | 6 | 5 | 5 | 3 | 3 | 6 |
| 5 | TGME49_223550-t26_1-p1 | hypothetical protein | 8 | 7 | 5 | 4 | 3 | 3 | 3 | 6 |
| 5 | TGME49_223560-t26_1-p1 | hypothetical protein | 9 | 7 | 5 | 5 | 3 | 3 | 3 | 6 |
| 4 | TGME49_223620-t26_1-p1 | protein transport protein BOS1, putative | 7 | 6 | 6 | 6 | 5 | 3 | 4 | 6 |
| 4 | TGME49_223630-t26_1-p1 | E2F-associated phosphoprotein, putative | 8 | 7 | 6 | 5 | 4 | 4 | 4 | 6 |
| 5 | TGME49_223725-t26_1-p1 | hypothetical protein | 10 | 7 | 6 | 3 | 3 | 3 | 3 | 6 |
| 4 | TGME49_223800-t26_1-p1 | hypothetical protein | 9 | 6 | 7 | 4 | 4 | 5 | 4 | 6 |
| 10 | TGME49_223880.P1096 | zinc finger, C3HC4 type (RING finger) domain-containing protein | 10 | 10 | 6 | 7 | 4 | 4 | 4 | 6 |
| 5 | TGME49_224080.P1106 | Kazal-type serine protease inhibitor domain-containing protein | 10 | 8 | 5 | 3 | 3 | 3 | 3 | 6 |
| 10 | TGME49_224160-t26_1-p1 | AP-5 complex subunit mu, putative | 10 | 10 | 5 | 6 | 5 | 4 | 3 | 6 |
| 4 | TGME49_224310.P1114 | palmitoyltransferase DHHC6 | 8 | 5 | 6 | 4 | 4 | 5 | 4 | 6 |
| 4 | TGME49_224730-t26_1-p1 | hypothetical protein | 8 | 6 | 6 | 3 | 3 | 5 | 5 | 6 |
| 4 | TGME49_224750-t26_1-p1 | SAG-related sequence SRS40F | 8 | 7 | 6 | 5 | 3 | 3 | 3 | 6 |
| 4 | TGME49_224760-t26_1-p1 | SAG-related sequence SRS40E | 8 | 7 | 6 | 4 | 3 | 3 | 3 | 6 |
| 4 | TGME49_224770.P1127 | SAG-related sequence SRS40D | 8 | 7 | 6 | 5 | 3 | 3 | 3 | 6 |
| 4 | TGME49_224790.P1128 | SAG-related sequence SRS40A | 8 | 8 | 6 | 5 | 3 | 3 | 3 | 6 |
| 5 | TGME49_224810.P1129 | hypothetical protein | 10 | 6 | 3 | 4 | 3 | 3 | 3 | 6 |
| 5 | TGME49_224830-t26_1-p1 | hypothetical protein | 10 | 7 | 5 | 5 | 3 | 4 | 4 | 6 |
| 4 | TGME49_224910-t26_1-p1 | divalent cation tolerance protein, CutA1 family protein | 8 | 6 | 5 | 5 | 5 | 3 | 3 | 6 |
| 5 | TGME49_224940-t26_1-p1 | hypothetical protein | 10 | 7 | 3 | 5 | 3 | 3 | 3 | 6 |
| 5 | TGME49_225108.P1150 | SNF7 family protein | 9 | 7 | 5 | 4 | 3 | 3 | 3 | 6 |
| 5 | TGME49_225110.P1151 | AP2 domain transcription factor AP2X-2 | 10 | 8 | 4 | 4 | 3 | 4 | 4 | 6 |
| 5 | TGME49_225130-t26_1-p1 | hypothetical protein | 10 | 6 | 6 | 4 | 3 | 3 | 3 | 6 |
| 5 | TGME49_225190.P1156 | hypothetical protein | 10 | 6 | 6 | 3 | 4 | 3 | 3 | 6 |
| 5 | TGME49_225230.P1159 | hypothetical protein | 10 | 8 | 5 | 5 | 3 | 3 | 3 | 6 |

| | | | | | | | | | | |
|---|---|---|---|---|---|---|---|---|---|---|
| 4 | TGME49_225250-t26_1-p1 | LSU ribosomal protein L14P, putative | 6 | 6 | 6 | 6 | 3 | 3 | 3 | 6 |
| 10 | TGME49_225300-t26_1-p1 | hypothetical protein | 8 | 7 | 3 | 6 | 3 | 3 | 3 | 6 |
| 4 | TGME49_225420-t26_1-p1 | histidine triad domain-containing protein | 7 | 7 | 6 | 5 | 4 | 3 | 3 | 6 |
| 10 | TGME49_225430-t26_1-p1 | Ras-related GTP binding A family protein | 9 | 8 | 3 | 6 | 3 | 3 | 3 | 6 |
| 4 | TGME49_225500-t26_1-p1 | protein SYS1, putative | 9 | 6 | 5 | 5 | 4 | 4 | 4 | 6 |
| 4 | TGME49_225540-t26_1-p1 | hypothetical protein | 9 | 7 | 5 | 5 | 4 | 3 | 3 | 6 |
| 4 | TGME49_225590.P1175 | hypothetical protein | 7 | 5 | 7 | 4 | 4 | 3 | 3 | 6 |
| 4 | TGME49_225830.P1182 | hypothetical protein | 8 | 7 | 6 | 4 | 4 | 3 | 3 | 6 |
| 5 | TGME49_225860-t26_1-p1 | hypothetical protein | 10 | 7 | 6 | 3 | 3 | 3 | 3 | 6 |
| 4 | TGME49_226040.P1194 | calmodulin CAM3 | 7 | 6 | 6 | 6 | 3 | 3 | 4 | 6 |
| 4 | TGME49_226290.P1201 | hypothetical protein | 8 | 7 | 6 | 4 | 3 | 3 | 4 | 6 |
| 4 | TGME49_226430-t26_1-p1 | reticulon protein | 7 | 6 | 6 | 3 | 5 | 3 | 3 | 6 |
| 4 | TGME49_226460-t26_1-p1 | hypothetical protein | 9 | 6 | 7 | 4 | 4 | 4 | 3 | 6 |
| 4 | TGME49_226520-t26_1-p1 | LSM domain-containing protein | 8 | 5 | 6 | 6 | 4 | 4 | 3 | 6 |
| 10 | TGME49_226780-t26_1-p1 | zinc finger, C3HC4 type (RING finger) domain-containing protein | 10 | 9 | 3 | 6 | 3 | 5 | 5 | 6 |
| 10 | TGME49_226850-t26_1-p1 | TBC domain-containing protein TBC15 | 10 | 9 | 3 | 6 | 7 | 3 | 3 | 6 |
| 5 | TGME49_226990.P1234 | hypothetical protein | 10 | 6 | 5 | 3 | 3 | 3 | 3 | 6 |
| 4 | TGME49_227150-t26_1-p1 | glutaredoxin 4 | 8 | 6 | 5 | 5 | 3 | 3 | 3 | 6 |
| 10 | TGME49_227380.P1246 | hypothetical protein | 10 | 8 | 3 | 6 | 4 | 3 | 3 | 6 |
| 5 | TGME49_227410-t26_1-p1 | hypothetical protein | 10 | 7 | 5 | 3 | 4 | 4 | 3 | 6 |
| 5 | TGME49_227810-t26_1-p1 | rhoptry kinase family protein ROP11 (incomplete catalytic triad) | 9 | 7 | 5 | 5 | 3 | 3 | 3 | 6 |
| 5 | TGME49_227880.P1263 | hypothetical protein | 9 | 7 | 5 | 4 | 3 | 4 | 3 | 6 |
| 4 | TGME49_227890.P1264 | hypothetical protein | 7 | 6 | 6 | 5 | 4 | 4 | 4 | 6 |
| 5 | TGME49_227930-t26_1-p1 | hypothetical protein | 10 | 6 | 5 | 4 | 4 | 3 | 3 | 6 |
| 4 | TGME49_227980.P1270 | hypothetical protein | 10 | 6 | 5 | 5 | 4 | 4 | 4 | 6 |
| 5 | TGME49_228050.P1272 | hypothetical protein | 9 | 7 | 3 | 3 | 3 | 3 | 3 | 6 |
| 5 | TGME49_228060-t26_1-p1 | hypothetical protein | 10 | 7 | 6 | 4 | 4 | 3 | 3 | 6 |
| 10 | TGME49_228220.P1284 | MYND finger domain-containing protein | 9 | 8 | 3 | 8 | 4 | 6 | 6 | 6 |
| 4 | TGME49_228360-t26_1-p1 | peptidyl-prolyl isomerase FKBP12, putative | 8 | 7 | 7 | 4 | 5 | 3 | 4 | 6 |
| 4 | TGME49_228440-t26_1-p1 | hypothetical protein | 8 | 5 | 6 | 3 | 3 | 4 | 4 | 6 |
| 4 | TGME49_228720.P1303 | hypothetical protein | 8 | 7 | 7 | 5 | 5 | 4 | 3 | 6 |
| 4 | TGME49_228730.P1304 | CRT domain-containing protein | 9 | 7 | 6 | 6 | 4 | 4 | 4 | 6 |
| 4 | TGME49_229160.P1312 | palmitoyltransferase DHHC12 | 9 | 5 | 5 | 5 | 4 | 4 | 3 | 6 |
| 5 | TGME49_229230-t26_1-p1 | hypothetical protein | 8 | 7 | 5 | 3 | 3 | 3 | 3 | 6 |
| 4 | TGME49_229240-t26_1-p1 | hypothetical protein | 8 | 7 | 6 | 5 | 3 | 3 | 3 | 6 |
| 10 | TGME49_229370.P1320 | AP2 domain transcription factor AP2VIII-1 | 10 | 8 | 5 | 5 | 3 | 5 | 5 | 6 |
| 4 | TGME49_229470-t26_1-p1 | Btz domain-containing protein | 8 | 6 | 5 | 4 | 4 | 4 | 4 | 6 |
| 10 | TGME49_229710.P1335 | OTU domain-containing protein OTUD3B | 10 | 7 | 3 | 5 | 5 | 5 | 4 | 6 |
| 5 | TGME49_229720.P1336 | hypothetical protein | 10 | 7 | 4 | 3 | 3 | 3 | 3 | 6 |
| 4 | TGME49_229910-t26_1-p1 | EF hand domain-containing protein | 7 | 7 | 6 | 6 | 3 | 4 | 4 | 6 |
| 4 | TGME49_230190-t26_1-p1 | hypothetical protein | 8 | 8 | 6 | 6 | 4 | 3 | 4 | 6 |
| 4 | TGME49_230230.P1363 | pre-mRNA polyadenylation factor FIP1 | 8 | 6 | 5 | 4 | 5 | 3 | 3 | 6 |
| 4 | TGME49_230850.P1387 | transverse sutures component TSC3 | 9 | 7 | 6 | 4 | 4 | 3 | 3 | 6 |
| 4 | TGME49_230910-t26_1-p1 | hypothetical protein | 8 | 7 | 7 | 5 | 4 | 5 | 4 | 6 |
| 5 | TGME49_230950.P1391 | hypothetical protein | 10 | 7 | 6 | 3 | 4 | 3 | 3 | 6 |
| 10 | TGME49_231030.P1398 | protein ARV1, putative | 10 | 9 | 4 | 5 | 5 | 4 | 4 | 6 |
| 4 | TGME49_231040.P1399 | 3' exoribonuclease family, domain 1 domain-containing protein | 8 | 7 | 7 | 4 | 5 | 4 | 3 | 6 |

| | | | | | | | | | | |
|---|---|---|---|---|---|---|---|---|---|---|
| 4 | TGME49_231390.P1415 | hypothetical protein | 8 | 7 | 5 | 6 | 3 | 3 | 3 | 6 |
| 4 | TGME49_232140.P1447 | hypothetical protein | 8 | 7 | 6 | 3 | 4 | 3 | 3 | 6 |
| 4 | TGME49_232210-t26_1-p1 | hypothetical protein | 8 | 6 | 6 | 4 | 4 | 4 | 3 | 6 |
| 4 | TGME49_232220.P1452 | SWIB/MDM2 domain-containing protein | 9 | 6 | 6 | 5 | 4 | 4 | 3 | 6 |
| 4 | TGME49_232240.P1453 | snRNA-activating protein complex subunit 3, putative | 10 | 7 | 6 | 5 | 5 | 3 | 3 | 6 |
| 10 | TGME49_232280.P1455 | hypothetical protein | 8 | 7 | 3 | 5 | 3 | 4 | 4 | 6 |
| 4 | TGME49_232290-t26_1-p1 | hypothetical protein | 8 | 6 | 8 | 3 | 3 | 5 | 3 | 6 |
| 10 | TGME49_232360.P1460 | exonuclease | 9 | 7 | 3 | 7 | 3 | 6 | 3 | 6 |
| 5 | TGME49_232475-t26_1-p1 | hypothetical protein | 10 | 8 | 3 | 3 | 3 | 4 | 3 | 6 |
| 4 | TGME49_232610.P1475 | histone acetyltransferase subunit nua4 protein | 8 | 7 | 6 | 5 | 5 | 4 | 4 | 6 |
| 10 | TGME49_232640-t26_1-p1 | RNA 2'-phosphotransferase, Tpt1/KptA family protein | 9 | 8 | 4 | 6 | 4 | 3 | 5 | 6 |
| 10 | TGME49_233120.P1494 | AP2 domain transcription factor AP2VIII-2 | 10 | 5 | 3 | 7 | 5 | 4 | 3 | 6 |
| 4 | TGME49_233280-t26_1-p1 | hypothetical protein | 9 | 5 | 5 | 5 | 3 | 3 | 3 | 6 |
| 5 | TGME49_233470-t26_1-p1 | hypothetical protein | 9 | 6 | 3 | 4 | 3 | 3 | 3 | 6 |
| 5 | TGME49_233490-t26_1-p1 | hypothetical protein | 10 | 8 | 5 | 4 | 4 | 3 | 3 | 6 |
| 4 | TGME49_233550-t26_1-p1 | polyketide cyclase/dehydrase | 10 | 6 | 6 | 4 | 4 | 4 | 3 | 6 |
| 5 | TGME49_233735.P1513 | hypothetical protein | 10 | 6 | 6 | 4 | 3 | 3 | 3 | 6 |
| 5 | TGME49_233870.P1522 | hypothetical protein | 10 | 6 | 4 | 3 | 3 | 3 | 3 | 6 |
| 5 | TGME49_233960.P1526 | hypothetical protein | 10 | 5 | 5 | 3 | 3 | 3 | 3 | 6 |
| 4 | TGME49_234180-t26_1-p1 | hypothetical protein | 7 | 6 | 5 | 5 | 4 | 3 | 3 | 6 |
| 5 | TGME49_234200.P1527 | hypothetical protein | 10 | 7 | 5 | 4 | 3 | 3 | 3 | 6 |
| 5 | TGME49_234250.P1529 | CPH1-interacting protein CIP1 | 10 | 9 | 4 | 4 | 3 | 4 | 3 | 6 |
| 10 | TGME49_234270.P1530 | hypothetical protein | 10 | 9 | 5 | 5 | 4 | 3 | 3 | 6 |
| 5 | TGME49_234300-t26_1-p1 | hypothetical protein | 10 | 6 | 4 | 3 | 3 | 4 | 3 | 6 |
| 4 | TGME49_234370-t26_1-p1 | SAG-related sequence SRS42 | 7 | 6 | 6 | 5 | 3 | 3 | 3 | 6 |
| 4 | TGME49_234380.P1533 | microneme protein MIC21 | 8 | 7 | 6 | 5 | 3 | 3 | 3 | 6 |
| 5 | TGME49_234430-t26_1-p1 | hypothetical protein | 10 | 9 | 5 | 4 | 3 | 3 | 3 | 6 |
| 4 | TGME49_234600-t26_1-p1 | hypothetical protein | 9 | 7 | 5 | 5 | 4 | 3 | 4 | 6 |
| 5 | TGME49_234980.P1547 | hypothetical protein | 9 | 6 | 4 | 3 | 3 | 3 | 3 | 6 |
| 4 | TGME49_235360.P1554 | hypothetical protein | 7 | 6 | 6 | 5 | 5 | 4 | 4 | 6 |
| 4 | TGME49_235690.P1571 | inner membrane complex protein IMC37 | 7 | 7 | 6 | 4 | 5 | 4 | 4 | 6 |
| 4 | TGME49_236000-t26_1-p1 | ferredoxin, putative | 8 | 6 | 5 | 5 | 4 | 5 | 4 | 6 |
| 10 | TGME49_236020-t26_1-p1 | BT1 family protein | 9 | 9 | 3 | 7 | 7 | 3 | 3 | 6 |
| 4 | TGME49_236120-t26_1-p1 | hypothetical protein | 8 | 6 | 5 | 5 | 3 | 3 | 3 | 6 |
| 4 | TGME49_236160.P1589 | hypothetical protein | 9 | 6 | 7 | 4 | 4 | 4 | 4 | 6 |
| 4 | TGME49_236270-t26_1-p1 | RING zinc finger protein | 8 | 6 | 5 | 4 | 4 | 4 | 4 | 6 |
| 10 | TGME49_236660.P1602 | START domain-containing protein | 10 | 9 | 4 | 5 | 5 | 6 | 5 | 6 |
| 4 | TGME49_236820-t26_1-p1 | hypothetical protein | 8 | 5 | 6 | 5 | 3 | 5 | 5 | 6 |
| 4 | TGME49_236940.P1609 | hypothetical protein | 7 | 6 | 6 | 5 | 4 | 3 | 4 | 6 |
| 5 | TGME49_237070-t26_1-p1 | hypothetical protein | 8 | 7 | 5 | 4 | 3 | 3 | 3 | 6 |
| 4 | TGME49_237180-t26_1-p1 | hypothetical protein | 8 | 7 | 6 | 5 | 3 | 3 | 4 | 6 |
| 4 | TGME49_237240-t26_1-p1 | pre-rRNA-processing protein TSR2, putative | 7 | 7 | 6 | 5 | 4 | 4 | 4 | 6 |
| 5 | TGME49_237450.P1628 | CAP domain-containing protein | 8 | 7 | 5 | 4 | 3 | 3 | 3 | 6 |
| 10 | TGME49_237470-t26_1-p1 | glycine cleavage T-protein (aminomethyl transferase) domain-containing protein | 10 | 8 | 3 | 7 | 6 | 3 | 3 | 6 |
| 4 | TGME49_237490.P1630 | centrin family protein | 8 | 6 | 6 | 6 | 5 | 5 | 3 | 6 |
| 5 | TGME49_237840-t26_1-p1 | hypothetical protein | 9 | 6 | 4 | 4 | 4 | 3 | 3 | 6 |
| 5 | TGME49_238020.P1639 | hypothetical protein | 10 | 8 | 5 | 4 | 4 | 3 | 3 | 6 |

| | | | | | | | | | | |
|---|---|---|---|---|---|---|---|---|---|---|
| 4 | TGME49_238060-t26_1-p1 | EF hand domain-containing protein | 8 | 6 | 6 | 5 | 3 | 3 | 3 | 6 |
| 5 | TGME49_238870-t26_1-p1 | hypothetical protein | 9 | 7 | 6 | 3 | 3 | 3 | 3 | 6 |
| 5 | TGME49_238930.P1656 | hypothetical protein | 10 | 5 | 5 | 4 | 3 | 3 | 3 | 6 |
| 10 | TGME49_238970.P1659 | hypothetical protein | 10 | 9 | 5 | 5 | 3 | 3 | 4 | 6 |
| 5 | TGME49_239270.P1667 | hypothetical protein | 9 | 6 | 3 | 3 | 3 | 3 | 3 | 6 |
| 5 | TGME49_239290.P1668 | hypothetical protein | 10 | 8 | 4 | 3 | 3 | 3 | 3 | 6 |
| 10 | TGME49_239340-t26_1-p1 | GTP-binding protein, putative | 10 | 9 | 5 | 5 | 4 | 4 | 4 | 6 |
| 5 | TGME49_239400.P1673 | inner membrane complex protein IMC28 | 10 | 8 | 5 | 4 | 4 | 3 | 3 | 6 |
| 5 | TGME49_239580-t26_1-p1 | hypothetical protein | 10 | 10 | 4 | 3 | 3 | 3 | 3 | 6 |
| 5 | TGME49_239610-t26_1-p1 | hypothetical protein | 9 | 7 | 6 | 4 | 3 | 3 | 3 | 6 |
| 4 | TGME49_239710-t26_1-p1 | phosphomannomutase | 8 | 7 | 7 | 6 | 4 | 3 | 3 | 6 |
| 5 | TGME49_239780-t26_1-p1 | hypothetical protein | 9 | 7 | 6 | 3 | 3 | 3 | 3 | 6 |
| 5 | TGME49_239810.P1688 | hypothetical protein | 10 | 8 | 3 | 3 | 3 | 3 | 3 | 6 |
| 5 | TGME49_239880-t26_1-p1 | hypothetical protein | 10 | 8 | 5 | 5 | 3 | 4 | 3 | 6 |
| 4 | TGME49_239890-t26_1-p1 | SCP family extracellular subfamily protein | 7 | 6 | 6 | 5 | 3 | 3 | 3 | 6 |
| 5 | TGME49_240310-t26_1-p1 | Toxoplasma gondii family E protein | 9 | 7 | 3 | 3 | 3 | 3 | 3 | 6 |
| 5 | TGME49_240370-t26_1-p1 | Toxoplasma gondii family E protein | 9 | 7 | 3 | 3 | 3 | 3 | 3 | 6 |
| 4 | TGME49_240470-t26_1-p1 | hypothetical protein | 9 | 6 | 6 | 3 | 5 | 3 | 3 | 6 |
| 4 | TGME49_240530-t26_1-p1 | hypothetical protein | 8 | 7 | 7 | 6 | 4 | 5 | 3 | 6 |
| 10 | TGME49_240630.P1710 | inner membrane complex protein IMC35 | 9 | 5 | 3 | 8 | 4 | 3 | 3 | 6 |
| 5 | TGME49_240900-t26_1-p1 | AP2 domain transcription factor AP2VI-2 | 10 | 6 | 5 | 3 | 3 | 3 | 3 | 6 |
| 4 | TGME49_240960-t26_1-p1 | AIG2 family protein | 9 | 7 | 6 | 5 | 3 | 3 | 3 | 6 |
| 5 | TGME49_240990-t26_1-p1 | hypothetical protein | 10 | 6 | 4 | 3 | 3 | 3 | 3 | 6 |
| 4 | TGME49_241010.P1723 | K+ channel tetramerisation domain-containing protein | 8 | 6 | 6 | 5 | 3 | 4 | 3 | 6 |
| 4 | TGME49_242080.P1731 | Qb-SNARE VTI1, putative | 7 | 7 | 6 | 6 | 4 | 4 | 4 | 6 |
| 4 | TGME49_242270-t26_1-p1 | Nudix hydrolase, putative | 8 | 6 | 6 | 5 | 4 | 3 | 4 | 6 |
| 5 | TGME49_242590-t26_1-p1 | hypothetical protein | 9 | 7 | 6 | 3 | 3 | 3 | 3 | 6 |
| 5 | TGME49_242670.P1740 | hypothetical protein | 10 | 6 | 5 | 4 | 4 | 3 | 3 | 6 |
| 4 | TGME49_243290-t26_1-p1 | hypothetical protein | 9 | 6 | 5 | 4 | 4 | 4 | 4 | 6 |
| 4 | TGME49_243560.P1765 | coiled-coil domain-containing protein 81, putative | 7 | 7 | 7 | 7 | 3 | 5 | 3 | 6 |
| 5 | TGME49_243635-t26_1-p1 | hypothetical protein | 10 | 9 | 3 | 4 | 3 | 3 | 3 | 6 |
| 4 | TGME49_243720-t26_1-p1 | peroxisomal biogenesis factor PEX11 | 8 | 7 | 7 | 5 | 5 | 4 | 4 | 6 |
| 4 | TGME49_243790-t26_1-p1 | SAG-related sequence SRS33 | 7 | 6 | 6 | 5 | 3 | 3 | 3 | 6 |
| 5 | TGME49_243970-t26_1-p1 | hypothetical protein | 10 | 8 | 4 | 4 | 4 | 3 | 3 | 6 |
| 5 | TGME49_244030-t26_1-p1 | inner membrane complex protein IMC3 | 10 | 7 | 3 | 3 | 3 | 3 | 3 | 6 |
| 4 | TGME49_244080-t26_1-p1 | hypothetical protein | 7 | 6 | 6 | 6 | 3 | 5 | 4 | 6 |
| 5 | TGME49_244140-t26_1-p1 | hypothetical protein | 9 | 7 | 5 | 3 | 3 | 3 | 3 | 6 |
| 4 | TGME49_244420-t26_1-p1 | rRNA biogenesis protein RRP36, putative | 8 | 7 | 6 | 5 | 4 | 3 | 4 | 6 |
| 10 | TGME49_244450-t26_1-p1 | protein phosphatase PPM3E, putative | 6 | 8 | 3 | 7 | 3 | 3 | 3 | 6 |
| 4 | TGME49_244570-t26_1-p1 | hypothetical protein | 9 | 6 | 7 | 5 | 3 | 4 | 3 | 6 |
| 4 | TGME49_244600-t26_1-p1 | hypothetical protein | 8 | 5 | 6 | 5 | 5 | 4 | 3 | 6 |
| 4 | TGME49_244630.P1799 | tRNA pseudouridine synthase D, putative | 9 | 6 | 6 | 5 | 4 | 3 | 3 | 6 |
| 5 | TGME49_244670-t26_1-p1 | hypothetical protein | 9 | 6 | 4 | 4 | 3 | 3 | 3 | 6 |
| 4 | TGME49_244690-t26_1-p1 | hypothetical protein | 9 | 7 | 7 | 5 | 4 | 3 | 4 | 6 |
| 4 | TGME49_244890.P1802 | cyclin, putative | 7 | 6 | 6 | 5 | 5 | 4 | 4 | 6 |
| 4 | TGME49_245520-t26_1-p1 | hypothetical protein | 7 | 6 | 6 | 5 | 4 | 3 | 4 | 6 |
| 4 | TGME49_245550.P1809 | hypothetical protein | 7 | 7 | 6 | 6 | 3 | 3 | 3 | 6 |
| 4 | TGME49_245600.P1812 | hypothetical protein | 8 | 7 | 6 | 4 | 3 | 3 | 3 | 6 |
| 4 | TGME49_245990.P1817 | hypothetical protein | 9 | 6 | 6 | 5 | 4 | 3 | 4 | 6 |

| | | | | | | | | | | |
|---|---|---|---|---|---|---|---|---|---|---|
| 5 | TGME49_246010.P1819 | hypothetical protein | 10 | 8 | 3 | 3 | 3 | 4 | 4 | 6 |
| 5 | TGME49_246030-t26_1-p1 | mediator complex subunit MED17 | 10 | 9 | 4 | 3 | 5 | 3 | 5 | 6 |
| 5 | TGME49_246090.P1823 | hypothetical protein | 10 | 8 | 3 | 3 | 7 | 3 | 3 | 6 |
| 10 | TGME49_246140-t26_1-p1 | hypothetical protein | 10 | 8 | 5 | 6 | 4 | 3 | 3 | 6 |
| 4 | TGME49_246190.P1828 | centrosome protein CEP530 | 9 | 7 | 6 | 5 | 5 | 4 | 4 | 6 |
| 4 | TGME49_246450-t26_1-p1 | hypothetical protein | 8 | 7 | 6 | 5 | 3 | 3 | 3 | 6 |
| 5 | TGME49_246590-t26_1-p1 | hypothetical protein | 9 | 7 | 5 | 3 | 3 | 3 | 3 | 6 |
| 5 | TGME49_246650.P1838 | palmitoyltransferase DHHC18 | 9 | 8 | 3 | 3 | 3 | 3 | 3 | 6 |
| 4 | TGME49_246670-t26_1-p1 | hypothetical protein | 8 | 7 | 6 | 5 | 5 | 4 | 3 | 6 |
| 4 | TGME49_246710-t26_1-p1 | hypothetical protein | 9 | 7 | 5 | 5 | 5 | 4 | 4 | 6 |
| 5 | TGME49_246750-t26_1-p1 | hypothetical protein | 9 | 6 | 5 | 4 | 3 | 3 | 4 | 6 |
| 4 | TGME49_246930-t26_1-p1 | calmodulin CAM1 | 7 | 6 | 6 | 5 | 3 | 3 | 4 | 6 |
| 4 | TGME49_246950.P1843 | apical cap protein AC9 | 9 | 7 | 6 | 4 | 4 | 4 | 4 | 6 |
| 10 | TGME49_247280-t26_1-p1 | hypothetical protein | 10 | 8 | 5 | 5 | 5 | 4 | 4 | 6 |
| 4 | TGME49_247310-t26_1-p1 | cpw-wpc domain-containing protein | 9 | 6 | 8 | 4 | 4 | 5 | 3 | 6 |
| 10 | TGME49_247320-t26_1-p1 | histone RNA hairpin-binding protein, putative | 10 | 8 | 4 | 6 | 5 | 4 | 3 | 6 |
| 4 | TGME49_247580-t26_1-p1 | glutaredoxin 3 | 8 | 5 | 6 | 5 | 4 | 4 | 4 | 6 |
| 10 | TGME49_247740-t26_1-p1 | methyltransferase, putative | 10 | 9 | 3 | 8 | 5 | 5 | 5 | 6 |
| 4 | TGME49_247750.P1864 | hypothetical protein | 8 | 6 | 5 | 5 | 3 | 3 | 3 | 6 |
| 4 | TGME49_247770.P1866 | cytochrome c oxidase subunit ApiCOX19 | 7 | 7 | 6 | 3 | 3 | 3 | 5 | 6 |
| 10 | TGME49_247790-t26_1-p1 | VRR-NUC domain-containing protein | 10 | 10 | 5 | 5 | 4 | 4 | 4 | 6 |
| 4 | TGME49_247940-t26_1-p1 | hypothetical protein | 8 | 8 | 6 | 5 | 3 | 3 | 3 | 6 |
| 5 | TGME49_247950.P1870 | DNA-directed RNA polymerase II largest subunit | 9 | 7 | 5 | 3 | 3 | 3 | 3 | 6 |
| 5 | TGME49_248140-t26_1-p1 | hypothetical protein | 9 | 6 | 5 | 3 | 3 | 3 | 3 | 6 |
| 4 | TGME49_248190-t26_1-p1 | hypothetical protein | 10 | 6 | 6 | 5 | 3 | 4 | 4 | 6 |
| 4 | TGME49_248210.P1878 | hypothetical protein | 8 | 7 | 6 | 3 | 5 | 3 | 3 | 6 |
| 5 | TGME49_248430-t26_1-p1 | hypothetical protein | 8 | 7 | 6 | 3 | 3 | 3 | 3 | 6 |
| 4 | TGME49_248520-t26_1-p1 | hypothetical protein | 8 | 7 | 6 | 5 | 4 | 3 | 3 | 6 |
| 5 | TGME49_248570.P1894 | hypothetical protein | 10 | 8 | 5 | 3 | 3 | 4 | 3 | 6 |
| 4 | TGME49_248690.P1901 | hypothetical protein | 8 | 7 | 6 | 6 | 3 | 3 | 4 | 6 |
| 4 | TGME49_248700.P1902 | alveolin domain containing intermediate filament IMC12 | 8 | 7 | 6 | 5 | 4 | 4 | 3 | 6 |
| 4 | TGME49_248780-t26_1-p1 | SRR1 protein | 9 | 6 | 5 | 5 | 3 | 4 | 4 | 6 |
| 5 | TGME49_248800-t26_1-p1 | hypothetical protein | 10 | 6 | 4 | 3 | 3 | 3 | 3 | 6 |
| 4 | TGME49_248870-t26_1-p1 | SNARE associated Golgi protein | 8 | 6 | 6 | 4 | 4 | 3 | 5 | 6 |
| 4 | TGME49_248960-t26_1-p1 | methyltransferase domain-containing protein | 9 | 6 | 7 | 5 | 4 | 5 | 5 | 6 |
| 5 | TGME49_248990.P1913 | prenylcysteine oxidase, putative | 10 | 9 | 3 | 3 | 3 | 3 | 3 | 6 |
| 4 | TGME49_249190-t26_1-p1 | AP2 domain transcription factor AP2XII-6 | 8 | 6 | 6 | 4 | 3 | 3 | 3 | 6 |
| 4 | TGME49_249200.P1920 | Ctr copper transporter family protein | 7 | 7 | 6 | 4 | 4 | 3 | 5 | 6 |
| 4 | TGME49_249310-t26_1-p1 | ER membrane protein complex subunit 8, putative | 8 | 7 | 6 | 4 | 5 | 4 | 4 | 6 |
| 5 | TGME49_249410-t26_1-p1 | hypothetical protein | 9 | 7 | 6 | 3 | 3 | 3 | 3 | 6 |
| 10 | TGME49_249600-t26_1-p1 | hypothetical protein | 10 | 8 | 3 | 7 | 6 | 5 | 5 | 6 |
| 4 | TGME49_249630.P1938 | glutathione S-transferase 1 | 8 | 7 | 7 | 5 | 5 | 3 | 3 | 6 |
| 4 | TGME49_249780.P1945 | hypothetical protein | 8 | 7 | 6 | 4 | 4 | 3 | 3 | 6 |
| 5 | TGME49_249800.P1946 | hypothetical protein | 9 | 6 | 4 | 4 | 3 | 3 | 3 | 6 |
| 5 | TGME49_250050-t26_1-p1 | hypothetical protein | 10 | 6 | 4 | 4 | 3 | 3 | 3 | 6 |
| 5 | TGME49_250910.P1970 | hypothetical protein | 9 | 7 | 5 | 4 | 4 | 3 | 3 | 6 |
| 5 | TGME49_250940.P1972 | hypothetical protein | 10 | 9 | 3 | 3 | 3 | 3 | 3 | 6 |
| 10 | TGME49_251400-t26_1-p1 | hypothetical protein | 10 | 8 | 5 | 5 | 4 | 4 | 4 | 6 |

| | | | | | | | | | | |
|---|---|---|---|---|---|---|---|---|---|---|
| 10 | TGME49_251510-t26_1-p1 | Ulp1 protease family, C-terminal catalytic domain-containing protein | 10 | 9 | 5 | 5 | 5 | 4 | 4 | 6 |
| 4 | TGME49_251520-t26_1-p1 | hypothetical protein | 7 | 6 | 6 | 6 | 3 | 3 | 3 | 6 |
| 4 | TGME49_251540-t26_1-p1 | dense granule protein GRA9 | 8 | 7 | 6 | 6 | 4 | 4 | 3 | 6 |
| 10 | TGME49_251560.P1978 | hypothetical protein | 10 | 9 | 4 | 6 | 4 | 4 | 4 | 6 |
| 5 | TGME49_251590.P1979 | hypothetical protein | 9 | 7 | 6 | 4 | 3 | 3 | 3 | 6 |
| 4 | TGME49_251640-t26_1-p1 | ubiquitin-conjugating enzyme subfamily protein | 8 | 6 | 7 | 6 | 3 | 4 | 4 | 6 |
| 4 | TGME49_251710-t26_1-p1 | protein transport protein GOS1, putative | 8 | 7 | 6 | 6 | 4 | 3 | 3 | 6 |
| 10 | TGME49_251740-t26_1-p1 | AP2 domain transcription factor AP2XII-9 | 10 | 9 | 4 | 6 | 5 | 5 | 5 | 6 |
| 10 | TGME49_251770-t26_1-p1 | hypothetical protein | 10 | 10 | 3 | 7 | 3 | 3 | 3 | 6 |
| 4 | TGME49_251790.P1982 | zinc finger Ran-binding domain-containing protein | 8 | 6 | 6 | 4 | 3 | 3 | 4 | 6 |
| 4 | TGME49_252230-t26_1-p1 | hypothetical protein | 9 | 8 | 5 | 5 | 4 | 3 | 4 | 6 |
| 4 | TGME49_252320-t26_1-p1 | Sas10/Utp3/C1D family protein | 8 | 6 | 5 | 5 | 4 | 4 | 4 | 6 |
| 5 | TGME49_252360-t26_1-p1 | rhoptry kinase family protein ROP24 (incomplete catalytic triad) | 9 | 8 | 3 | 3 | 3 | 3 | 3 | 6 |
| 10 | TGME49_252400-t26_1-p1 | HIT zinc finger protein | 9 | 7 | 4 | 5 | 4 | 4 | 3 | 6 |
| 4 | TGME49_253020.P2006 | hypothetical protein | 9 | 7 | 7 | 5 | 4 | 4 | 3 | 6 |
| 10 | TGME49_253100-t26_1-p1 | hypothetical protein | 9 | 8 | 3 | 6 | 3 | 3 | 4 | 6 |
| 10 | TGME49_253130-t26_1-p1 | transporter, major facilitator family protein | 7 | 9 | 3 | 8 | 3 | 3 | 4 | 6 |
| 5 | TGME49_253140-t26_1-p1 | hypothetical protein | 10 | 8 | 5 | 4 | 3 | 3 | 3 | 6 |
| 4 | TGME49_253160-t26_1-p1 | hypothetical protein | 9 | 7 | 6 | 6 | 4 | 4 | 3 | 6 |
| 5 | TGME49_253310.P2016 | hypothetical protein | 10 | 7 | 5 | 4 | 4 | 3 | 3 | 6 |
| 10 | TGME49_253330.P2017 | bradyzoite pseudokinase BPK1 | 9 | 8 | 3 | 6 | 3 | 3 | 3 | 6 |
| 5 | TGME49_253410-t26_1-p1 | hypothetical protein | 9 | 7 | 5 | 5 | 3 | 3 | 3 | 6 |
| 4 | TGME49_253500.P2025 | intraflagellar transport 57-like protein | 8 | 6 | 7 | 3 | 3 | 3 | 3 | 6 |
| 10 | TGME49_253790-t26_1-p1 | zinc finger (CCCH type) motif-containing protein | 10 | 10 | 3 | 5 | 3 | 3 | 3 | 6 |
| 4 | TGME49_253830-t26_1-p1 | hypothetical protein | 8 | 6 | 5 | 5 | 3 | 3 | 3 | 6 |
| 5 | TGME49_253850-t26_1-p1 | zinc finger, CCHC-type domain-containing protein | 9 | 7 | 5 | 4 | 3 | 3 | 3 | 6 |
| 5 | TGME49_253990-t26_1-p1 | hypothetical protein | 10 | 7 | 6 | 4 | 3 | 3 | 3 | 6 |
| 5 | TGME49_254000.P2041 | dense granule protein GRA47 | 9 | 6 | 5 | 4 | 3 | 3 | 3 | 6 |
| 5 | TGME49_254135-t26_1-p1 | hypothetical protein | 10 | 5 | 5 | 4 | 4 | 3 | 3 | 6 |
| 4 | TGME49_254220-t26_1-p1 | RNA recognition motif-containing protein | 9 | 7 | 6 | 4 | 4 | 4 | 4 | 6 |
| 10 | TGME49_254230-t26_1-p1 | hypothetical protein | 9 | 9 | 3 | 7 | 7 | 5 | 4 | 6 |
| 4 | TGME49_254260.P2053 | COX19 cytochrome c oxidase assembly family protein | 7 | 6 | 6 | 4 | 4 | 4 | 4 | 6 |
| 4 | TGME49_254270-t26_1-p1 | hypothetical protein | 8 | 7 | 7 | 4 | 4 | 3 | 3 | 6 |
| 4 | TGME49_254280-t26_1-p1 | DNA-directed RNA polymerase III RPC9 | 9 | 7 | 6 | 5 | 4 | 4 | 4 | 6 |
| 5 | TGME49_254530-t26_1-p1 | hypothetical protein | 10 | 8 | 5 | 4 | 4 | 3 | 3 | 6 |
| 4 | TGME49_254540-t26_1-p1 | CDP-alcohol phosphatidyltransferase superfamily protein | 9 | 6 | 6 | 5 | 3 | 4 | 3 | 6 |
| 5 | TGME49_254700.P2069 | RING finger protein, putative | 9 | 6 | 5 | 3 | 4 | 4 | 3 | 6 |
| 4 | TGME49_254890-t26_1-p1 | CRAL/TRIO domain-containing protein | 8 | 7 | 6 | 5 | 3 | 3 | 4 | 6 |
| 4 | TGME49_254920.P2081 | hypothetical protein | 8 | 7 | 6 | 4 | 4 | 3 | 4 | 6 |
| 5 | TGME49_254960-t26_1-p1 | hypothetical protein | 10 | 8 | 3 | 3 | 3 | 3 | 3 | 6 |
| 4 | TGME49_255270-t26_1-p1 | UNC-50 protein UNC1 | 8 | 7 | 6 | 6 | 3 | 3 | 4 | 6 |
| 4 | TGME49_255410-t26_1-p1 | hypothetical protein | 7 | 6 | 5 | 5 | 4 | 3 | 4 | 6 |
| 5 | TGME49_255440.P2101 | hypothetical protein | 9 | 7 | 3 | 3 | 3 | 3 | 3 | 6 |
| 10 | TGME49_255470.P2104 | GTP-binding protein, putative | 10 | 9 | 5 | 7 | 5 | 4 | 4 | 6 |
| 4 | TGME49_255620.P2108 | B9-C2 domain-containing protein | 6 | 7 | 7 | 5 | 3 | 5 | 3 | 6 |
| 4 | TGME49_255670.P2112 | methyltransferase domain-containing protein | 8 | 7 | 7 | 5 | 3 | 6 | 3 | 6 |

| | | | | | | | | | | |
|---|---|---|---|---|---|---|---|---|---|---|
| 4 | TGME49_255690.P2114 | 2-C-methyl-D-erythritol 2,4-cyclodiphosphate synthase domain-containing protein | 8 | 6 | 6 | 3 | 4 | 3 | 4 | 6 |
| 4 | TGME49_255930-t26_1-p1 | hypothetical protein | 9 | 6 | 6 | 5 | 3 | 3 | 3 | 6 |
| 10 | TGME49_256830.P2141 | SacI domain-containing protein | 10 | 8 | 3 | 6 | 5 | 5 | 4 | 6 |
| 5 | TGME49_256880.P2145 | protein kinase domain-containing protein | 9 | 7 | 6 | 4 | 3 | 3 | 3 | 6 |
| 4 | TGME49_257490-t26_1-p1 | prefoldin subunit 3, putative | 7 | 7 | 7 | 6 | 3 | 5 | 4 | 6 |
| 5 | TGME49_257595-t26_1-p1 | SET domain-containing protein | 10 | 5 | 6 | 3 | 3 | 3 | 3 | 6 |
| 5 | TGME49_257610-t26_1-p1 | hypothetical protein | 10 | 6 | 6 | 3 | 3 | 3 | 3 | 6 |
| 5 | TGME49_257625-t26_1-p1 | tetrahydrofolate dehydrogenase/cyclohydrolase, NAD(P)-binding domain-containing protein | 9 | 6 | 5 | 3 | 3 | 3 | 3 | 6 |
| 4 | TGME49_257650.P2181 | hypothetical protein | 8 | 7 | 6 | 4 | 4 | 3 | 3 | 6 |
| 5 | TGME49_257670-t26_1-p1 | hypothetical protein | 10 | 6 | 5 | 5 | 3 | 3 | 3 | 6 |
| 5 | TGME49_257700.P2183 | hypothetical protein | 10 | 6 | 4 | 4 | 3 | 3 | 3 | 6 |
| 10 | TGME49_257760-t26_1-p1 | hypothetical protein | 10 | 8 | 5 | 5 | 5 | 4 | 4 | 6 |
| 4 | TGME49_257980.P2195 | ribosome recycling factor protein | 9 | 7 | 6 | 4 | 5 | 3 | 3 | 6 |
| 4 | TGME49_258470.P2222 | inner membrane complex protein IMC24 | 8 | 7 | 6 | 5 | 5 | 4 | 4 | 6 |
| 4 | TGME49_258550-t26_1-p1 | SAG-related sequence SRS28 | 8 | 7 | 6 | 4 | 4 | 3 | 3 | 6 |
| 4 | TGME49_258625.P2230 | peptidyl-prolyl cis-trans isomerase, FKBP-type domain-containing protein | 9 | 6 | 6 | 5 | 4 | 4 | 4 | 6 |
| 5 | TGME49_258660-t26_1-p1 | rhoptry protein ROP6 | 9 | 8 | 5 | 5 | 3 | 3 | 3 | 6 |
| 5 | TGME49_258670.P2233 | hypothetical protein | 10 | 7 | 3 | 4 | 3 | 3 | 3 | 6 |
| 4 | TGME49_258800.P2241 | rhoptry kinase family protein ROP31 | 6 | 7 | 7 | 3 | 3 | 4 | 3 | 6 |
| 5 | TGME49_258910.P2249 | hypothetical protein | 10 | 8 | 5 | 5 | 3 | 3 | 3 | 6 |
| 4 | TGME49_259000.P2255 | ER membrane protein complex subunit 4, putative | 7 | 6 | 6 | 5 | 4 | 3 | 3 | 6 |
| 4 | TGME49_259020-t26_1-p1 | bradyzoite antigen BAG1 | 8 | 7 | 6 | 3 | 4 | 3 | 3 | 6 |
| 4 | TGME49_259140.P2264 | hypothetical protein | 10 | 7 | 6 | 5 | 4 | 4 | 4 | 6 |
| 5 | TGME49_259155-t26_1-p1 | hypothetical protein | 10 | 7 | 4 | 3 | 3 | 3 | 3 | 6 |
| 5 | TGME49_259168.P2266 | hypothetical protein | 10 | 7 | 5 | 3 | 3 | 3 | 4 | 6 |
| 4 | TGME49_259210.P2272 | jmjC domain-containing protein C2orf60 | 10 | 7 | 6 | 4 | 5 | 3 | 4 | 6 |
| 5 | TGME49_259270-t26_1-p1 | SAG-related sequence SRS26E | 9 | 7 | 4 | 3 | 3 | 3 | 3 | 6 |
| 4 | TGME49_259610.P2281 | hypothetical protein | 8 | 5 | 5 | 5 | 4 | 3 | 3 | 6 |
| 4 | TGME49_259650.P2285 | zinc finger protein, putative | 8 | 6 | 5 | 5 | 4 | 4 | 3 | 6 |
| 4 | TGME49_259700-t26_1-p1 | hypothetical protein | 7 | 7 | 6 | 6 | 3 | 4 | 4 | 6 |
| 4 | TGME49_259860.P2292 | Myb domain-containing protein | 10 | 7 | 6 | 5 | 4 | 4 | 4 | 6 |
| 5 | TGME49_259970.P2300 | hypothetical protein | 9 | 7 | 4 | 3 | 3 | 3 | 3 | 6 |
| 10 | TGME49_260150.P2306 | tetratricopeptide repeat-containing protein | 10 | 10 | 5 | 7 | 4 | 4 | 3 | 6 |
| 10 | TGME49_260190-t26_1-p1 | microneme protein MIC13 | 9 | 8 | 5 | 8 | 5 | 4 | 4 | 6 |
| 10 | TGME49_260380.P2318 | hypothetical protein | 10 | 9 | 5 | 5 | 3 | 4 | 3 | 6 |
| 10 | TGME49_260390.P2319 | hypothetical protein | 10 | 8 | 5 | 5 | 5 | 4 | 3 | 6 |
| 5 | TGME49_260580-t26_1-p1 | hypothetical protein | 10 | 9 | 4 | 4 | 3 | 4 | 4 | 6 |
| 5 | TGME49_260650.P2336 | glycosyltransferase, group 2 family protein | 10 | 5 | 5 | 4 | 4 | 3 | 3 | 6 |
| 4 | TGME49_260800.P2341 | hypothetical protein | 10 | 5 | 6 | 5 | 4 | 3 | 3 | 6 |
| 4 | TGME49_261475.P2365 | hypothetical protein | 7 | 6 | 6 | 5 | 5 | 3 | 4 | 6 |
| 5 | TGME49_261490.P2367 | hypothetical protein | 10 | 8 | 3 | 3 | 4 | 3 | 3 | 6 |
| 5 | TGME49_261560-t26_1-p1 | Tat binding protein 1(TBP-1)-interacting protein TBPIP | 9 | 8 | 4 | 3 | 3 | 5 | 3 | 6 |
| 4 | TGME49_261590.P2371 | ankyrin, putative | 7 | 7 | 6 | 6 | 3 | 4 | 3 | 6 |
| 4 | TGME49_261620-t26_1-p1 | hypothetical protein | 8 | 6 | 4 | 5 | 4 | 4 | 4 | 6 |
| 4 | TGME49_261630-t26_1-p1 | gamma interferon inducible lysosomal thiol reductase (GILT) protein | 8 | 6 | 7 | 3 | 3 | 5 | 3 | 6 |

| | | | | | | | | | | |
|---|---|---|---|---|---|---|---|---|---|---|
| 4 | TGME49_261670.P2375 | ribonuclease H1/H2 small subunit protein | 8 | 7 | 6 | 5 | 4 | 3 | 4 | 6 |
| 4 | TGME49_261690.P2377 | hypothetical protein | 7 | 7 | 7 | 4 | 6 | 3 | 3 | 6 |
| 5 | TGME49_261730.P2379 | hypothetical protein | 10 | 10 | 4 | 4 | 3 | 3 | 3 | 6 |
| 4 | TGME49_262090-t26_1-p1 | hypothetical protein | 8 | 7 | 7 | 5 | 3 | 5 | 3 | 6 |
| 4 | TGME49_262130.P2395 | hypothetical protein | 8 | 6 | 5 | 4 | 3 | 4 | 4 | 6 |
| 5 | TGME49_262170.P2399 | hypothetical protein | 8 | 7 | 5 | 3 | 3 | 3 | 3 | 6 |
| 4 | TGME49_262470-t26_1-p1 | C protein immunoglobulin-A-binding beta antigen, putative | 7 | 6 | 5 | 5 | 4 | 3 | 3 | 6 |
| 5 | TGME49_262630.P2415 | hypothetical protein | 9 | 7 | 3 | 4 | 3 | 3 | 3 | 6 |
| 4 | TGME49_262710-t26_1-p1 | Ctr copper transporter family protein | 8 | 6 | 6 | 5 | 5 | 3 | 5 | 6 |
| 5 | TGME49_262730.P2424 | rhoptry protein ROP16 | 10 | 7 | 6 | 3 | 3 | 3 | 4 | 6 |
| 4 | TGME49_262980.P2435 | hypothetical protein | 8 | 7 | 6 | 5 | 3 | 4 | 4 | 6 |
| 10 | TGME49_263010.P2438 | POZ domain-containing protein | 8 | 9 | 3 | 5 | 7 | 4 | 3 | 6 |
| 5 | TGME49_263120.P2447 | hypothetical protein | 10 | 6 | 5 | 3 | 3 | 3 | 3 | 6 |
| 4 | TGME49_263170.P2450 | HSP20-like chaperone, putative | 7 | 6 | 7 | 4 | 5 | 3 | 3 | 6 |
| 5 | TGME49_263230.P2456 | apicomplexan amino acid transporter ApiAT7-1 | 10 | 9 | 3 | 3 | 3 | 3 | 3 | 6 |
| 4 | TGME49_263240-t26_1-p1 | MerC domain-containing protein | 7 | 6 | 5 | 5 | 4 | 4 | 4 | 6 |
| 4 | TGME49_263280.P2458 | hypothetical protein | 7 | 7 | 6 | 3 | 5 | 4 | 3 | 6 |
| 4 | TGME49_263330.P2461 | Adaptin ear-binding coat-associated protein 2 (NECAP-2) isoform 2 family protein | 8 | 6 | 6 | 5 | 3 | 3 | 3 | 6 |
| 5 | TGME49_263350.P2463 | hypothetical protein | 8 | 7 | 6 | 3 | 3 | 3 | 3 | 6 |
| 4 | TGME49_263400.P2466 | hypothetical protein | 9 | 6 | 7 | 5 | 4 | 3 | 4 | 6 |
| 5 | TGME49_263540.P2475 | hypothetical protein | 10 | 9 | 4 | 4 | 3 | 3 | 3 | 6 |
| 5 | TGME49_263750.P2491 | hypothetical protein | 10 | 6 | 4 | 3 | 3 | 3 | 3 | 6 |
| 10 | TGME49_263980.P2499 | pentatricopeptide repeat domain-containing protein | 10 | 8 | 5 | 6 | 4 | 3 | 3 | 6 |
| 4 | TGME49_264150-t26_1-p1 | hypothetical protein | 8 | 7 | 6 | 5 | 4 | 3 | 3 | 6 |
| 4 | TGME49_264485-t26_1-p1 | AP2 domain transcription factor AP2IX-3 | 10 | 6 | 6 | 5 | 3 | 5 | 5 | 6 |
| 4 | TGME49_264600-t26_1-p1 | hypothetical protein | 7 | 7 | 6 | 6 | 3 | 4 | 4 | 6 |
| 4 | TGME49_264710-t26_1-p1 | bucentaur or craniofacial development domain-containing protein | 8 | 8 | 7 | 5 | 4 | 5 | 3 | 6 |
| 5 | TGME49_264805-t26_1-p1 | hypothetical protein | 10 | 6 | 3 | 3 | 3 | 3 | 4 | 6 |
| 5 | TGME49_264980-t26_1-p1 | hypothetical protein | 10 | 8 | 5 | 4 | 3 | 3 | 3 | 6 |
| 4 | TGME49_264990.P2529 | inner membrane complex protein IMC16 | 8 | 7 | 6 | 4 | 4 | 4 | 4 | 6 |
| 5 | TGME49_265030.P2531 | hypothetical protein | 9 | 7 | 4 | 4 | 3 | 3 | 3 | 6 |
| 10 | TGME49_265050.P2532 | zinc finger protein, putative | 10 | 8 | 3 | 5 | 3 | 4 | 3 | 6 |
| 10 | TGME49_265080-t26_1-p1 | Tubulin-tyrosine ligase family protein | 10 | 10 | 3 | 8 | 4 | 3 | 3 | 6 |
| 5 | TGME49_265260-t26_1-p1 | hypothetical protein | 9 | 7 | 5 | 5 | 3 | 3 | 3 | 6 |
| 4 | TGME49_265430-t26_1-p1 | hypothetical protein | 9 | 5 | 6 | 5 | 3 | 3 | 3 | 6 |
| 5 | TGME49_265455-t26_1-p1 | hypothetical protein | 9 | 7 | 6 | 4 | 3 | 3 | 3 | 6 |
| 4 | TGME49_265810.P2546 | hypothetical protein | 9 | 7 | 6 | 4 | 4 | 3 | 3 | 6 |
| 5 | TGME49_265840.P2549 | nuclear mesh protein NMP1 | 10 | 6 | 6 | 4 | 4 | 3 | 3 | 6 |
| 5 | TGME49_266410-t26_1-p1 | hypothetical protein | 9 | 6 | 3 | 3 | 3 | 3 | 3 | 6 |
| 4 | TGME49_266500-t26_1-p1 | OTU domain-containing protein OTU9 | 8 | 7 | 7 | 4 | 3 | 3 | 3 | 6 |
| 5 | TGME49_266650-t26_1-p1 | hypothetical protein | 9 | 6 | 4 | 3 | 3 | 3 | 3 | 6 |
| 5 | TGME49_266670.P2576 | hypothetical protein | 10 | 7 | 4 | 3 | 3 | 3 | 3 | 6 |
| 4 | TGME49_266700-t26_1-p1 | hypothetical protein | 7 | 7 | 6 | 5 | 3 | 3 | 3 | 6 |
| 10 | TGME49_266910-t26_1-p1 | cell-cycle-associated protein kinase GSK, putative | 8 | 8 | 3 | 6 | 6 | 6 | 3 | 6 |
| 10 | TGME49_266950.P2589 | protein kinase, putative | 10 | 8 | 3 | 7 | 6 | 3 | 3 | 6 |
| 10 | TGME49_267030.P2593 | ribonuclease type III Dicer | 10 | 8 | 4 | 7 | 3 | 4 | 4 | 6 |

| | | | | | | | | | | |
|---|---|---|---|---|---|---|---|---|---|---|
| 4 | TGME49_267090-t26_1-p1 | hypothetical protein | 8 | 7 | 6 | 5 | 4 | 3 | 3 | 6 |
| 5 | TGME49_267140-t26_1-p1 | SAG-related sequence SRS38B | 8 | 7 | 6 | 3 | 3 | 3 | 3 | 6 |
| 4 | TGME49_267150-t26_1-p1 | SAG-related sequence SRS38C | 9 | 7 | 6 | 5 | 3 | 3 | 3 | 6 |
| 4 | TGME49_267300-t26_1-p1 | golgi apparatus membrane protein TVP15, putative | 8 | 7 | 7 | 4 | 6 | 4 | 3 | 6 |
| 4 | TGME49_267700.P2611 | hypothetical protein | 7 | 6 | 6 | 5 | 4 | 4 | 4 | 6 |
| 5 | TGME49_267855-t26_1-p1 | hypothetical protein | 10 | 6 | 6 | 4 | 4 | 3 | 3 | 6 |
| 5 | TGME49_268035.P2623 | hypothetical protein | 10 | 8 | 4 | 3 | 3 | 3 | 3 | 6 |
| 5 | TGME49_268182-t26_1-p1 | transporter, major facilitator family protein | 9 | 8 | 5 | 4 | 3 | 3 | 3 | 6 |
| 5 | TGME49_268340-t26_1-p1 | glycosyltransferase family 28 C-terminal domain-containing protein | 9 | 6 | 5 | 4 | 3 | 4 | 3 | 6 |
| 5 | TGME49_268680-t26_1-p1 | hypothetical protein | 10 | 8 | 3 | 4 | 3 | 3 | 3 | 6 |
| 4 | TGME49_268910.P2658 | signal peptidase I protein | 7 | 7 | 6 | 5 | 4 | 4 | 3 | 6 |
| 4 | TGME49_268970-t26_1-p1 | hypothetical protein | 8 | 7 | 5 | 6 | 3 | 3 | 3 | 6 |
| 5 | TGME49_269130.P2665 | hypothetical protein | 9 | 7 | 4 | 4 | 3 | 3 | 3 | 6 |
| 4 | TGME49_269175.P2668 | U6 snRNA phosphodiesterase, putative | 9 | 6 | 6 | 3 | 4 | 4 | 3 | 6 |
| 5 | TGME49_269210.P2672 | hypothetical protein | 10 | 6 | 5 | 4 | 3 | 3 | 3 | 6 |
| 5 | TGME49_269270.P2676 | leucine rich repeat-containing protein | 9 | 7 | 6 | 3 | 3 | 3 | 3 | 6 |
| 5 | TGME49_269600.P2688 | biotin-requiring enzyme domain-containing protein | 10 | 6 | 5 | 3 | 4 | 3 | 3 | 6 |
| 5 | TGME49_269620.P2689 | hypothetical protein | 9 | 7 | 4 | 4 | 3 | 3 | 3 | 6 |
| 4 | TGME49_269640.P2691 | hypothetical protein | 7 | 6 | 6 | 5 | 5 | 4 | 3 | 6 |
| 10 | TGME49_269760-t26_1-p1 | 'chromo' (CHRromatin Organization MOdifier) domain-containing protein | 10 | 9 | 5 | 5 | 4 | 4 | 4 | 6 |
| 10 | TGME49_269910-t26_1-p1 | transcription factor C2H2, putative | 8 | 8 | 3 | 7 | 3 | 3 | 5 | 6 |
| 5 | TGME49_270050.P2715 | hypothetical protein | 9 | 6 | 4 | 3 | 3 | 3 | 3 | 6 |
| 5 | TGME49_270060.P2716 | hypothetical protein | 9 | 8 | 4 | 3 | 3 | 3 | 3 | 6 |
| 5 | TGME49_270160.P2723 | hypothetical protein | 9 | 6 | 3 | 3 | 3 | 3 | 3 | 6 |
| 5 | TGME49_270230.P2728 | hypothetical protein | 10 | 6 | 6 | 3 | 4 | 4 | 4 | 6 |
| 5 | TGME49_270240-t26_1-p1 | MAG1 | 9 | 7 | 6 | 4 | 3 | 3 | 3 | 6 |
| 10 | TGME49_270450.P2737 | MCM2/3/5 family protein | 10 | 9 | 3 | 7 | 6 | 5 | 4 | 6 |
| 5 | TGME49_270610-t26_1-p1 | hypothetical protein | 9 | 7 | 3 | 3 | 3 | 4 | 3 | 6 |
| 4 | TGME49_270640-t26_1-p1 | serine/arginine-rich splicing factor SR4 | 7 | 6 | 6 | 5 | 3 | 3 | 5 | 6 |
| 5 | TGME49_270720.P2780 | hypothetical protein | 10 | 10 | 3 | 3 | 4 | 3 | 3 | 6 |
| 5 | TGME49_270760-t26_1-p1 | asparagine synthase | 10 | 8 | 5 | 5 | 4 | 3 | 3 | 6 |
| 4 | TGME49_270790.P2783 | hypothetical protein | 7 | 6 | 5 | 5 | 4 | 3 | 3 | 6 |
| 4 | TGME49_270810-t26_1-p1 | hypothetical protein | 9 | 7 | 6 | 5 | 4 | 3 | 3 | 6 |
| 5 | TGME49_270890-t26_1-p1 | hypothetical protein | 9 | 6 | 4 | 4 | 4 | 3 | 3 | 6 |
| 4 | TGME49_270980.P2795 | phosphoglycerate mutase family protein | 10 | 6 | 7 | 5 | 4 | 5 | 4 | 6 |
| 10 | TGME49_271090.P2799 | hypothetical protein | 10 | 7 | 5 | 5 | 4 | 4 | 4 | 6 |
| 4 | TGME49_271220-t26_1-p1 | hypothetical protein | 7 | 6 | 6 | 5 | 3 | 3 | 3 | 6 |
| 5 | TGME49_271260-t26_1-p1 | hypothetical protein | 10 | 7 | 5 | 3 | 3 | 3 | 3 | 6 |
| 5 | TGME49_271600.P2818 | hypothetical protein | 9 | 7 | 5 | 4 | 3 | 4 | 4 | 6 |
| 4 | TGME49_271750-t26_1-p1 | EF hand domain-containing protein | 8 | 6 | 5 | 5 | 4 | 3 | 4 | 6 |
| 5 | TGME49_271770-t26_1-p1 | hypothetical protein | 10 | 7 | 4 | 3 | 3 | 4 | 4 | 6 |
| 4 | TGME49_271930.P2831 | inner membrane complex protein IMC20 | 7 | 6 | 6 | 5 | 3 | 4 | 4 | 6 |
| 10 | TGME49_271990.P2834 | hypothetical protein | 10 | 9 | 5 | 6 | 5 | 5 | 5 | 6 |
| 4 | TGME49_272340.P2852 | hypothetical protein | 9 | 7 | 7 | 4 | 4 | 4 | 4 | 6 |
| 5 | TGME49_272380.P2856 | hypothetical protein | 9 | 8 | 3 | 3 | 3 | 3 | 3 | 6 |
| 4 | TGME49_272383.P2857 | hypothetical protein | 9 | 5 | 7 | 4 | 4 | 3 | 3 | 6 |
| 5 | TGME49_272460.P2865 | dense granule protein GRA72 | 9 | 7 | 4 | 3 | 3 | 3 | 3 | 6 |
| 4 | TGME49_272520.P2869 | transverse sutures component TSC4 | 9 | 7 | 5 | 5 | 4 | 4 | 4 | 6 |

| | | | | | | | | | | |
|---|---|---|---|---|---|---|---|---|---|---|
| 4 | TGME49_272530-t26_1-p1 | hypothetical protein | 9 | 7 | 6 | 5 | 4 | 3 | 4 | 6 |
| 5 | TGME49_272620.P2874 | hypothetical protein | 9 | 7 | 4 | 3 | 3 | 3 | 3 | 6 |
| 5 | TGME49_272730.P2883 | hypothetical protein | 10 | 5 | 5 | 4 | 4 | 3 | 3 | 6 |
| 10 | TGME49_272770-t26_1-p1 | LTV1 domain-containing protein | 10 | 8 | 5 | 7 | 5 | 5 | 4 | 6 |
| 4 | TGME49_273030.P2889 | phosphoglycerate mutase PGM1 | 8 | 8 | 7 | 4 | 5 | 3 | 3 | 6 |
| 5 | TGME49_273050.P2890 | basal complex component BCC8 | 10 | 7 | 5 | 4 | 3 | 3 | 3 | 6 |
| 4 | TGME49_273110-t26_1-p1 | SAG-related sequence SRS30D | 9 | 7 | 6 | 5 | 3 | 3 | 3 | 6 |
| 4 | TGME49_273280.P2895 | hypothetical protein | 9 | 6 | 7 | 6 | 3 | 4 | 4 | 6 |
| 5 | TGME49_273595.P2911 | hypothetical protein | 10 | 6 | 6 | 3 | 4 | 4 | 3 | 6 |
| 4 | TGME49_273630.P2913 | DnaJ domain-containing protein | 7 | 7 | 6 | 6 | 4 | 4 | 4 | 6 |
| 4 | TGME49_273640-t26_1-p1 | hypothetical protein | 9 | 7 | 5 | 5 | 4 | 3 | 3 | 6 |
| 5 | TGME49_273670.P2915 | hypothetical protein | 10 | 7 | 6 | 3 | 3 | 3 | 3 | 6 |
| 5 | TGME49_273790.P2920 | hypothetical protein | 10 | 8 | 5 | 5 | 3 | 3 | 3 | 6 |
| 5 | TGME49_273815-t26_1-p1 | hypothetical protein | 9 | 8 | 5 | 3 | 3 | 3 | 3 | 6 |
| 4 | TGME49_273930-t26_1-p1 | DNA topoisomerase domain-containing protein | 8 | 6 | 5 | 5 | 4 | 3 | 4 | 6 |
| 10 | TGME49_274000.P2931 | centrin-binding protein SFI1 | 10 | 10 | 4 | 9 | 7 | 4 | 3 | 6 |
| 5 | TGME49_274180.P2945 | hypothetical protein | 10 | 6 | 3 | 4 | 3 | 3 | 3 | 6 |
| 10 | TGME49_275450.P2953 | hypothetical protein | 10 | 8 | 4 | 5 | 3 | 4 | 4 | 6 |
| 4 | TGME49_275770-t26_1-p1 | hypothetical protein | 7 | 6 | 5 | 5 | 4 | 4 | 4 | 6 |
| 10 | TGME49_275780-t26_1-p1 | F-box domain-containing protein | 10 | 8 | 4 | 5 | 4 | 4 | 3 | 6 |
| 5 | TGME49_276200.P2968 | hypothetical protein | 10 | 7 | 4 | 3 | 3 | 3 | 3 | 6 |
| 4 | TGME49_276930-t26_1-p1 | IMP1-like protein, putative | 7 | 6 | 6 | 3 | 3 | 5 | 5 | 6 |
| 4 | TGME49_276990-t26_1-p1 | membrane-associated progesterone-binding protein homolog MAPR | 7 | 6 | 6 | 6 | 5 | 3 | 4 | 6 |
| 5 | TGME49_277060.P2976 | hypothetical protein | 10 | 6 | 5 | 4 | 3 | 3 | 3 | 6 |
| 5 | TGME49_277150-t26_1-p1 | XPG N-terminal domain-containing protein | 10 | 7 | 6 | 3 | 3 | 4 | 3 | 6 |
| 4 | TGME49_277220.P2978 | hypothetical protein | 9 | 7 | 6 | 5 | 4 | 4 | 3 | 6 |
| 4 | TGME49_277680-t26_1-p1 | hypothetical protein | 7 | 6 | 6 | 5 | 5 | 4 | 3 | 6 |
| 4 | TGME49_278160-t26_1-p1 | vesicle transport v-snare protein | 8 | 7 | 6 | 6 | 4 | 4 | 3 | 6 |
| 5 | TGME49_278180-t26_1-p1 | hypothetical protein | 8 | 7 | 5 | 3 | 3 | 3 | 3 | 6 |
| 4 | TGME49_278450-t26_1-p1 | hypothetical protein | 9 | 7 | 6 | 4 | 5 | 4 | 3 | 6 |
| 5 | TGME49_278510.P3000 | protein phosphatase PPM3F | 9 | 7 | 6 | 3 | 3 | 3 | 3 | 6 |
| 5 | TGME49_278580.P3004 | hypothetical protein | 10 | 7 | 3 | 5 | 3 | 3 | 3 | 6 |
| 4 | TGME49_278680.P3008 | hypothetical protein | 7 | 6 | 6 | 5 | 4 | 3 | 3 | 6 |
| 5 | TGME49_278760.P3013 | hypothetical protein | 10 | 7 | 6 | 4 | 4 | 3 | 3 | 6 |
| 4 | TGME49_278900-t26_1-p1 | protein kinase, putative | 8 | 7 | 7 | 5 | 4 | 4 | 4 | 6 |
| 5 | TGME49_280450-t26_1-p1 | hypothetical protein | 10 | 8 | 4 | 3 | 3 | 3 | 3 | 6 |
| 5 | TGME49_280460-t26_1-p1 | AP2 domain transcription factor AP2VIIa-2 | 10 | 8 | 5 | 3 | 3 | 3 | 3 | 6 |
| 5 | TGME49_280522.P3042 | hypothetical protein | 10 | 7 | 4 | 4 | 3 | 3 | 3 | 6 |
| 4 | TGME49_280530-t26_1-p1 | hypothetical protein | 8 | 5 | 5 | 4 | 4 | 4 | 4 | 6 |
| 5 | TGME49_280720-t26_1-p1 | hypothetical protein | 9 | 6 | 4 | 4 | 4 | 3 | 3 | 6 |
| 5 | TGME49_281470.P3062 | hypothetical protein | 9 | 6 | 4 | 3 | 3 | 3 | 3 | 6 |
| 4 | TGME49_281500-t26_1-p1 | aminotransferase, putative | 7 | 7 | 5 | 4 | 4 | 4 | 4 | 6 |
| 5 | TGME49_281650-t26_1-p1 | hypothetical protein | 9 | 6 | 5 | 3 | 4 | 4 | 4 | 6 |
| 10 | TGME49_281675.P3072 | rhoptry kinase family protein ROP35 | 10 | 9 | 3 | 8 | 3 | 3 | 3 | 6 |
| 4 | TGME49_281910.P3073 | hypothetical protein | 9 | 7 | 6 | 5 | 3 | 3 | 3 | 6 |
| 5 | TGME49_282055-t26_1-p1 | protein phosphatase PP2C-hn | 9 | 7 | 3 | 3 | 3 | 3 | 3 | 6 |
| 4 | TGME49_282190-t26_1-p1 | hydrolase, NUDIX family protein | 9 | 7 | 6 | 5 | 3 | 3 | 3 | 6 |
| 4 | TGME49_283510.P3088 | IMC-associated protein IAP1 | 8 | 7 | 6 | 4 | 3 | 5 | 4 | 6 |
| 4 | TGME49_283530.P3090 | small rab-related GTPase | 7 | 5 | 7 | 6 | 5 | 3 | 3 | 6 |

| | | | | | | | | | | |
|---|---|---|---|---|---|---|---|---|---|---|
| 4 | TGME49_283540.P3091 | microneme protein MIC20 | 8 | 7 | 7 | 5 | 4 | 3 | 3 | 6 |
| 5 | TGME49_283580.P3093 | DnaJ domain-containing protein | 9 | 7 | 5 | 4 | 3 | 3 | 3 | 6 |
| 4 | TGME49_284010-t26_1-p1 | 5'-3' exonuclease, N-terminal resolvase family domain-containing protein | 8 | 8 | 6 | 5 | 5 | 4 | 4 | 6 |
| 5 | TGME49_284040-t26_1-p1 | hypothetical protein | 10 | 9 | 4 | 4 | 3 | 3 | 3 | 6 |
| 4 | TGME49_284590.P3110 | mediator complex subunit MED10 | 8 | 7 | 6 | 5 | 5 | 4 | 4 | 6 |
| 4 | TGME49_285445.P3131 | cullin, putative | 10 | 6 | 6 | 5 | 4 | 4 | 3 | 6 |
| 4 | TGME49_285530-t26_1-p1 | ribosomal protein RPL35 | 8 | 5 | 5 | 5 | 4 | 3 | 3 | 6 |
| 4 | TGME49_285680-t26_1-p1 | dihydrolipoamide acyltransferase, putative | 9 | 6 | 6 | 5 | 4 | 4 | 4 | 6 |
| 4 | TGME49_285710-t26_1-p1 | hypothetical protein | 8 | 7 | 6 | 5 | 4 | 4 | 3 | 6 |
| 5 | TGME49_286000-t26_1-p1 | hypothetical protein | 10 | 8 | 4 | 4 | 3 | 4 | 3 | 6 |
| 4 | TGME49_286010-t26_1-p1 | hypothetical protein | 9 | 6 | 5 | 7 | 4 | 3 | 4 | 6 |
| 10 | TGME49_286030.P3144 | micropore protein MPP2 | 10 | 8 | 5 | 5 | 4 | 3 | 4 | 6 |
| 5 | TGME49_286160.P3150 | non-specific serine/threonine protein kinase | 10 | 8 | 3 | 3 | 3 | 3 | 3 | 6 |
| 5 | TGME49_286190-t26_1-p1 | hypothetical protein | 10 | 7 | 4 | 3 | 3 | 3 | 3 | 6 |
| 4 | TGME49_286260-t26_1-p1 | tetratricopeptide repeat-containing protein | 9 | 6 | 7 | 6 | 4 | 3 | 3 | 6 |
| 5 | TGME49_286520.P3154 | hypothetical protein | 10 | 8 | 3 | 3 | 3 | 3 | 3 | 6 |
| 4 | TGME49_286540-t26_1-p1 | hypothetical protein | 9 | 6 | 5 | 5 | 3 | 3 | 3 | 6 |
| 4 | TGME49_286590-t26_1-p1 | microtubule associated protein SPM2 | 9 | 7 | 5 | 6 | 3 | 4 | 3 | 6 |
| 4 | TGME49_286670-t26_1-p1 | hypothetical protein | 9 | 7 | 6 | 5 | 3 | 3 | 3 | 6 |
| 5 | TGME49_287220-t26_1-p1 | hypothetical protein | 9 | 7 | 4 | 4 | 3 | 3 | 3 | 6 |
| 5 | TGME49_288190.P3178 | E3 ubiquitin-protein ligase, putative | 10 | 7 | 6 | 4 | 4 | 3 | 4 | 6 |
| 4 | TGME49_288220-t26_1-p1 | CAP domain-containing protein | 8 | 6 | 6 | 6 | 3 | 4 | 3 | 6 |
| 4 | TGME49_288270-t26_1-p1 | DUF4602 domain-containing protein | 9 | 7 | 6 | 5 | 4 | 3 | 4 | 6 |
| 4 | TGME49_288320-t26_1-p1 | adenylate kinase | 7 | 7 | 7 | 6 | 3 | 3 | 3 | 6 |
| 5 | TGME49_288370.P3181 | hypothetical protein | 9 | 7 | 5 | 4 | 4 | 3 | 4 | 6 |
| 4 | TGME49_288390-t26_1-p1 | B3GNTL1 protein | 8 | 7 | 7 | 5 | 3 | 3 | 3 | 6 |
| 10 | TGME49_288475-t26_1-p1 | hypothetical protein | 10 | 10 | 5 | 6 | 3 | 3 | 3 | 6 |
| 10 | TGME49_288520-t26_1-p1 | ATPase, AAA family protein | 9 | 9 | 3 | 3 | 5 | 6 | 5 | 6 |
| 4 | TGME49_288620.P3184 | Erv1 / Alr family protein | 7 | 6 | 5 | 5 | 4 | 4 | 3 | 6 |
| 10 | TGME49_288650-t26_1-p1 | dense granule protein GRA12 | 9 | 7 | 3 | 6 | 3 | 3 | 5 | 6 |
| 4 | TGME49_288685-t26_1-p1 | Fe-S protein assembly co-chaperone HscB protein | 8 | 6 | 6 | 5 | 5 | 4 | 4 | 6 |
| 5 | TGME49_288840.P3192 | dense granule protein GRA18 | 9 | 7 | 5 | 3 | 3 | 3 | 3 | 6 |
| 4 | TGME49_288880-t26_1-p1 | hypothetical protein | 9 | 7 | 6 | 4 | 5 | 4 | 4 | 6 |
| 5 | TGME49_289060-t26_1-p1 | hypothetical protein | 9 | 7 | 5 | 3 | 3 | 3 | 3 | 6 |
| 5 | TGME49_289110-t26_1-p1 | hypothetical protein | 10 | 8 | 5 | 3 | 3 | 3 | 3 | 6 |
| 10 | TGME49_289120-t26_1-p1 | Uso1-like protein ULP1 | 10 | 9 | 4 | 5 | 5 | 4 | 3 | 6 |
| 4 | TGME49_289180-t26_1-p1 | thioredoxin family redox-active protein, putative | 7 | 6 | 6 | 4 | 5 | 5 | 3 | 6 |
| 4 | TGME49_289230-t26_1-p1 | 50S ribosomal protein L12, putative | 9 | 6 | 5 | 5 | 5 | 4 | 4 | 6 |
| 4 | TGME49_289290.P3201 | hypothetical protein | 7 | 6 | 5 | 5 | 4 | 3 | 3 | 6 |
| 5 | TGME49_289740.P3213 | POZ domain-containing protein | 10 | 9 | 3 | 3 | 3 | 3 | 3 | 6 |
| 5 | TGME49_289790.P3215 | spindle and kinetochore-associated protein SKA3 | 10 | 7 | 5 | 4 | 3 | 3 | 3 | 6 |
| 4 | TGME49_290225.P3225 | protein kinase, putative | 10 | 7 | 6 | 5 | 4 | 4 | 4 | 6 |
| 4 | TGME49_290300.P3226 | hypothetical protein | 8 | 7 | 5 | 5 | 5 | 4 | 4 | 6 |
| 10 | TGME49_290880-t26_1-p1 | DnaJ domain-containing protein | 10 | 9 | 5 | 5 | 3 | 4 | 4 | 6 |
| 4 | TGME49_290930-t26_1-p1 | Mnd1 family protein | 7 | 6 | 6 | 5 | 4 | 3 | 4 | 6 |
| 10 | TGME49_290970.P3241 | serine palmitoyltransferase SPT2 | 9 | 9 | 3 | 9 | 3 | 3 | 3 | 6 |
| 4 | TGME49_291010-t26_1-p1 | hypothetical protein | 7 | 6 | 5 | 5 | 5 | 3 | 4 | 6 |
| 4 | TGME49_291890.P3254 | microneme protein MIC1 | 9 | 7 | 6 | 5 | 3 | 3 | 3 | 6 |
| 4 | TGME49_291910-t26_1-p1 | hypothetical protein | 8 | 7 | 5 | 5 | 4 | 3 | 3 | 6 |

| | | | | | | | | | | |
|---|---|---|---|---|---|---|---|---|---|---|
| 5 | TGME49_291960.P3255 | rhoptry kinase family protein ROP40 (incomplete catalytic triad) | 9 | 7 | 6 | 3 | 4 | 3 | 3 | 6 |
| 5 | TGME49_292260-t26_1-p1 | SAG-related sequence SRS36B | 8 | 7 | 5 | 3 | 3 | 3 | 3 | 6 |
| 10 | TGME49_292300.P3263 | queuine tRNA-ribosyltransferase, putative | 10 | 9 | 4 | 5 | 5 | 5 | 4 | 6 |
| 10 | TGME49_292320.P3265 | queuine tRNA-ribosyltransferase, putative | 10 | 8 | 4 | 5 | 4 | 4 | 4 | 6 |
| 5 | TGME49_293250.P3281 | hypothetical protein | 10 | 9 | 5 | 3 | 3 | 3 | 3 | 6 |
| 5 | TGME49_293280.P3285 | cyclin protein | 10 | 7 | 5 | 5 | 3 | 3 | 3 | 6 |
| 4 | TGME49_293300.P3286 | Yip1 domain-containing protein | 9 | 7 | 6 | 5 | 5 | 4 | 4 | 6 |
| 4 | TGME49_293580.P3310 | prefoldin subunit 2, putative | 7 | 6 | 6 | 5 | 4 | 4 | 4 | 6 |
| 4 | TGME49_293600.P3312 | ribosomal protein RPL27 | 8 | 6 | 6 | 5 | 5 | 4 | 4 | 6 |
| 10 | TGME49_293700.P3319 | WD domain, G-beta repeat-containing protein | 9 | 9 | 6 | 7 | 4 | 5 | 4 | 6 |
| 5 | TGME49_293840.P3334 | EF hand domain-containing protein | 10 | 5 | 5 | 3 | 3 | 3 | 3 | 6 |
| 4 | TGME49_293860.P3336 | CS domain-containing protein | 8 | 7 | 6 | 5 | 4 | 3 | 5 | 6 |
| 5 | TGME49_294020.P3341 | zinc finger, C3HC4 type (RING finger) domain-containing protein | 9 | 7 | 4 | 3 | 4 | 4 | 4 | 6 |
| 5 | TGME49_294050.P3343 | hypothetical protein | 9 | 7 | 6 | 3 | 4 | 3 | 3 | 6 |
| 5 | TGME49_294060.P3344 | hypothetical protein | 10 | 8 | 5 | 3 | 4 | 3 | 3 | 6 |
| 10 | TGME49_294200.P3346 | glucose-6-phosphate 1-dehydrogenase | 10 | 9 | 3 | 5 | 3 | 3 | 3 | 6 |
| 4 | TGME49_294310-t26_1-p1 | hypothetical protein | 9 | 6 | 5 | 5 | 3 | 3 | 4 | 6 |
| 4 | TGME49_294340.P3352 | transverse sutures component TSC6 | 8 | 5 | 6 | 5 | 4 | 3 | 3 | 6 |
| 4 | TGME49_294720-t26_1-p1 | RING zinc finger protein | 9 | 6 | 6 | 4 | 4 | 3 | 3 | 6 |
| 4 | TGME49_294790.P3368 | microneme protein MIC19 | 8 | 7 | 5 | 5 | 4 | 3 | 4 | 6 |
| 4 | TGME49_294880-t26_1-p1 | 8-oxoguanine DNA glycosylase, N-terminal domain-containing protein | 9 | 6 | 7 | 5 | 5 | 4 | 3 | 6 |
| 5 | TGME49_295110-t26_1-p1 | rhoptry protein ROP7 | 8 | 8 | 5 | 3 | 3 | 3 | 3 | 6 |
| 4 | TGME49_295400-t26_1-p1 | MINDY deubiquitinase, putative | 9 | 8 | 6 | 4 | 5 | 4 | 4 | 6 |
| 5 | TGME49_295430-t26_1-p1 | hypothetical protein | 8 | 7 | 5 | 3 | 3 | 3 | 3 | 6 |
| 5 | TGME49_295650-t26_1-p1 | hypothetical protein | 9 | 7 | 3 | 3 | 3 | 3 | 3 | 6 |
| 4 | TGME49_295910-t26_1-p1 | ATP-dependent Clp protease adaptor protein ClpS protein | 8 | 5 | 5 | 5 | 5 | 4 | 3 | 6 |
| 5 | TGME49_295935-t26_1-p1 | KRUF family protein | 9 | 7 | 5 | 3 | 3 | 3 | 3 | 6 |
| 5 | TGME49_295980-t26_1-p1 | hypothetical protein | 10 | 7 | 5 | 4 | 4 | 3 | 3 | 6 |
| 5 | TGME49_297340-t26_1-p1 | hypothetical protein | 10 | 5 | 5 | 4 | 4 | 3 | 3 | 6 |
| 4 | TGME49_297400.P3404 | hypothetical protein | 8 | 7 | 6 | 5 | 5 | 3 | 4 | 6 |
| 4 | TGME49_297510.P3411 | protein farnesyltransferase subunit alpha, putative | 9 | 7 | 7 | 4 | 4 | 4 | 4 | 6 |
| 5 | TGME49_297690-t26_1-p1 | hypothetical protein | 10 | 10 | 3 | 3 | 4 | 3 | 4 | 6 |
| 4 | TGME49_297760-t26_1-p1 | hypothetical protein | 7 | 6 | 5 | 5 | 4 | 3 | 4 | 6 |
| 5 | TGME49_297910-t26_1-p1 | hypothetical protein | 8 | 7 | 5 | 3 | 3 | 3 | 3 | 6 |
| 4 | TGME49_297920-t26_1-p1 | hypothetical protein | 8 | 7 | 5 | 5 | 3 | 3 | 3 | 6 |
| 4 | TGME49_297925.P3424 | HesB-like domain-containing protein | 9 | 6 | 6 | 3 | 5 | 4 | 4 | 6 |
| 4 | TGME49_298040-t26_1-p1 | hypothetical protein | 8 | 6 | 5 | 6 | 5 | 3 | 3 | 6 |
| 5 | TGME49_299100-t26_1-p1 | hypothetical protein | 10 | 8 | 3 | 3 | 3 | 3 | 3 | 6 |
| 5 | TGME49_299270.P3441 | hypothetical protein | 10 | 7 | 4 | 3 | 3 | 3 | 3 | 6 |
| 5 | TGME49_299780.P3443 | dense granule protein GRA76 | 9 | 8 | 5 | 5 | 3 | 3 | 3 | 6 |
| 10 | TGME49_299990.P3448 | archease family protein | 8 | 6 | 3 | 6 | 5 | 3 | 4 | 6 |
| 4 | TGME49_300010.P3450 | hypothetical protein | 8 | 6 | 6 | 5 | 5 | 4 | 3 | 6 |
| 5 | TGME49_300070.P3452 | hypothetical protein | 9 | 7 | 6 | 3 | 3 | 3 | 3 | 6 |
| 10 | TGME49_300230-t26_1-p1 | nuclease, putative | 9 | 8 | 5 | 5 | 3 | 4 | 4 | 6 |
| 5 | TGME49_301000-t26_1-p1 | methyltransferase | 9 | 6 | 3 | 3 | 3 | 3 | 3 | 6 |
| 5 | TGME49_301240.P3470 | hypothetical protein | 9 | 7 | 4 | 4 | 3 | 3 | 3 | 6 |

| | | | | | | | | | | |
|---|---|---|---|---|---|---|---|---|---|---|
| 5 | TGME49_301290.P3472 | hypothetical protein | 9 | 7 | 3 | 4 | 3 | 3 | 3 | 6 |
| 4 | TGME49_301460-t26_1-p1 | hypothetical protein | 9 | 5 | 6 | 4 | 4 | 3 | 4 | 6 |
| 4 | TGME49_304450-t26_1-p1 | cation-transporting ATPase | 8 | 5 | 5 | 6 | 3 | 4 | 3 | 6 |
| 4 | TGME49_304730.P3489 | MOB-like protein phocein | 8 | 7 | 7 | 4 | 3 | 4 | 5 | 6 |
| 4 | TGME49_305060-t26_1-p1 | CAP-Gly domain-containing protein | 8 | 7 | 7 | 6 | 3 | 5 | 4 | 6 |
| 4 | TGME49_305250-t26_1-p1 | hypothetical protein | 8 | 7 | 6 | 4 | 3 | 3 | 3 | 6 |
| 5 | TGME49_305530-t26_1-p1 | hypothetical protein | 8 | 7 | 6 | 3 | 3 | 3 | 3 | 6 |
| 5 | TGME49_305620-t26_1-p1 | RFT1 domain-containing protein | 10 | 8 | 5 | 4 | 4 | 4 | 4 | 6 |
| 5 | TGME49_306190-t26_1-p1 | hypothetical protein | 9 | 7 | 5 | 3 | 3 | 3 | 3 | 6 |
| 5 | TGME49_306230-t26_1-p1 | hypothetical protein | 10 | 6 | 3 | 3 | 3 | 3 | 3 | 6 |
| 5 | TGME49_306250-t26_1-p1 | hypothetical protein | 10 | 8 | 5 | 4 | 3 | 3 | 3 | 6 |
| 4 | TGME49_306280-t26_1-p1 | mediator complex subunit MED7 | 8 | 7 | 6 | 5 | 4 | 3 | 3 | 6 |
| 5 | TGME49_306300-t26_1-p1 | hypothetical protein | 10 | 9 | 5 | 4 | 3 | 4 | 3 | 6 |
| 5 | TGME49_306320-t26_1-p1 | Myb family DNA-binding domain-containing protein | 10 | 6 | 5 | 4 | 4 | 4 | 4 | 6 |
| 10 | TGME49_306450-t26_1-p1 | short chain dehydrogenase family protein, putative | 8 | 7 | 3 | 6 | 3 | 3 | 3 | 6 |
| 4 | TGME49_306470-t26_1-p1 | isoprenylcysteine carboxyl methyltransferase (icmt) family protein | 9 | 7 | 5 | 5 | 4 | 4 | 4 | 6 |
| 10 | TGME49_306490.P3524 | rhoptry apical surface protein RASP3 | 10 | 9 | 5 | 5 | 4 | 3 | 3 | 6 |
| 5 | TGME49_306510-t26_1-p1 | zinc finger (CCCH type) motif-containing protein | 10 | 8 | 5 | 5 | 3 | 3 | 3 | 6 |
| 4 | TGME49_306590-t26_1-p1 | MIP18 domain-containing protein | 9 | 6 | 5 | 5 | 5 | 4 | 4 | 6 |
| 5 | TGME49_306620-t26_1-p1 | AP2 domain transcription factor AP2IX-9 | 10 | 8 | 5 | 4 | 4 | 3 | 3 | 6 |
| 5 | TGME49_306890.P3529 | hypothetical protein | 10 | 8 | 3 | 5 | 3 | 3 | 3 | 6 |
| 4 | TGME49_307820-t26_1-p1 | hypothetical protein | 9 | 7 | 6 | 5 | 4 | 3 | 3 | 6 |
| 4 | TGME49_308020-t26_1-p1 | SAG-related sequence SRS57 | 8 | 7 | 6 | 5 | 4 | 3 | 3 | 6 |
| 5 | TGME49_308090-t26_1-p1 | rhoptry protein ROP5 | 8 | 8 | 3 | 3 | 3 | 3 | 3 | 6 |
| 4 | TGME49_308830.P3543 | dual specificity protein phosphatase | 7 | 7 | 6 | 6 | 4 | 3 | 3 | 6 |
| 4 | TGME49_309040.P3557 | hypothetical protein | 9 | 8 | 6 | 5 | 4 | 5 | 4 | 6 |
| 5 | TGME49_309050-t26_1-p1 | hypothetical protein | 9 | 7 | 5 | 4 | 4 | 3 | 3 | 6 |
| 4 | TGME49_309070-t26_1-p1 | hypothetical protein | 9 | 5 | 6 | 5 | 4 | 4 | 3 | 6 |
| 4 | TGME49_309280-t26_1-p1 | zinc finger, C3HC4 type (RING finger) domain-containing protein | 7 | 8 | 6 | 3 | 4 | 4 | 3 | 6 |
| 4 | TGME49_309290-t26_1-p1 | hypothetical protein | 8 | 8 | 7 | 4 | 5 | 3 | 4 | 6 |
| 4 | TGME49_309760.P3575 | dense granule protein GRA55 | 8 | 7 | 7 | 5 | 3 | 3 | 3 | 6 |
| 10 | TGME49_309860.P3579 | hypothetical protein | 9 | 8 | 3 | 7 | 3 | 5 | 5 | 6 |
| 5 | TGME49_309880.P3581 | hypothetical protein | 10 | 9 | 5 | 4 | 3 | 3 | 3 | 6 |
| 4 | TGME49_309990-t26_1-p1 | hypothetical protein | 7 | 6 | 6 | 6 | 3 | 3 | 5 | 6 |
| 4 | TGME49_310030.P3590 | cyclase-associated protein CAP | 7 | 7 | 6 | 6 | 3 | 3 | 5 | 6 |
| 10 | TGME49_310140.P3597 | LMBR1 domain-containing protein | 10 | 9 | 4 | 5 | 3 | 4 | 5 | 6 |
| 10 | TGME49_310160-t26_1-p1 | argonaute AGO | 10 | 10 | 3 | 9 | 3 | 3 | 3 | 6 |
| 5 | TGME49_310240-t26_1-p1 | hypothetical protein | 9 | 6 | 5 | 4 | 3 | 3 | 3 | 6 |
| 4 | TGME49_310330.P3605 | hypothetical protein | 9 | 7 | 6 | 5 | 4 | 4 | 4 | 6 |
| 4 | TGME49_310390-t26_1-p1 | hypothetical protein | 8 | 7 | 6 | 5 | 4 | 3 | 3 | 6 |
| 4 | TGME49_310570-t26_1-p1 | RNA recognition motif-containing protein | 8 | 6 | 6 | 4 | 4 | 4 | 4 | 6 |
| 4 | TGME49_310990-t26_1-p1 | hypothetical protein | 7 | 7 | 6 | 6 | 3 | 4 | 3 | 6 |
| 5 | TGME49_311040-t26_1-p1 | hypothetical protein | 8 | 7 | 5 | 4 | 3 | 3 | 3 | 6 |
| 5 | TGME49_311140-t26_1-p1 | hypothetical protein | 9 | 7 | 5 | 3 | 3 | 3 | 3 | 6 |
| 4 | TGME49_311320.P3641 | Qb-SNARE protein, putative | 7 | 6 | 6 | 4 | 5 | 4 | 3 | 6 |
| 5 | TGME49_311335.P3642 | dual specificity phosphatase, catalytic domain-containing protein | 10 | 7 | 6 | 4 | 3 | 3 | 3 | 6 |
| 5 | TGME49_311490-t26_1-p1 | hypothetical protein | 10 | 7 | 3 | 3 | 3 | 3 | 3 | 6 |

| | | | | | | | | | | |
|---|---|---|---|---|---|---|---|---|---|---|
| 10 | TGME49_311510-t26_1-p1 | eIF2 kinase IF2K-B | 10 | 9 | 3 | 5 | 3 | 4 | 4 | 6 |
| 5 | TGME49_311650.P3653 | leucine rich repeat-containing protein | 10 | 9 | 4 | 3 | 3 | 3 | 3 | 6 |
| 5 | TGME49_311670-t26_1-p1 | transmembrane surface antigen TSA1 | 9 | 7 | 3 | 4 | 3 | 3 | 3 | 6 |
| 5 | TGME49_311730-t26_1-p1 | hypothetical protein | 10 | 8 | 5 | 5 | 3 | 3 | 3 | 6 |
| 5 | TGME49_311760-t26_1-p1 | hypothetical protein | 10 | 8 | 3 | 4 | 3 | 3 | 3 | 6 |
| 10 | TGME49_311765-t26_1-p1 | hypothetical protein | 10 | 10 | 3 | 8 | 3 | 3 | 3 | 6 |
| 5 | TGME49_311770.P3655 | basal complex component BCC3 | 10 | 9 | 5 | 4 | 3 | 3 | 3 | 6 |
| 5 | TGME49_311790-t26_1-p1 | hypothetical protein | 8 | 7 | 6 | 3 | 3 | 3 | 3 | 6 |
| 4 | TGME49_311905-t26_1-p1 | hypothetical protein | 8 | 7 | 6 | 4 | 4 | 4 | 4 | 6 |
| 5 | TGME49_312130-t26_1-p1 | hypothetical protein | 10 | 7 | 3 | 4 | 3 | 3 | 3 | 6 |
| 4 | TGME49_312410-t26_1-p1 | methyltransferase, putative | 9 | 7 | 7 | 4 | 4 | 4 | 4 | 6 |
| 4 | TGME49_312640.P3678 | selenoprotein S, putative | 8 | 7 | 6 | 5 | 5 | 3 | 3 | 6 |
| 4 | TGME49_312670-t26_1-p1 | hypothetical protein | 9 | 7 | 7 | 4 | 5 | 4 | 4 | 6 |
| 5 | TGME49_312830-t26_1-p1 | hypothetical protein | 10 | 7 | 5 | 3 | 4 | 3 | 3 | 6 |
| 10 | TGME49_312905-t26_1-p1 | polyribonucleotide 5-hydroxyl-kinase Clp1, putative | 10 | 9 | 5 | 6 | 5 | 4 | 4 | 6 |
| 5 | TGME49_312950-t26_1-p1 | RAP-1 family protein | 9 | 7 | 4 | 4 | 3 | 3 | 3 | 6 |
| 10 | TGME49_312970.P3685 | hypothetical protein | 10 | 8 | 5 | 5 | 4 | 4 | 3 | 6 |
| 5 | TGME49_313110-t26_1-p1 | hypothetical protein | 9 | 7 | 5 | 4 | 4 | 3 | 3 | 6 |
| 5 | TGME49_313360-t26_1-p1 | hypothetical protein | 10 | 8 | 4 | 4 | 3 | 3 | 3 | 6 |
| 5 | TGME49_313422-t26_1-p1 | hypothetical protein | 10 | 8 | 4 | 4 | 3 | 3 | 4 | 6 |
| 5 | TGME49_313445.P3707 | hypothetical protein | 9 | 6 | 4 | 3 | 3 | 3 | 3 | 6 |
| 4 | TGME49_313520.P3711 | hypothetical protein | 7 | 7 | 6 | 5 | 5 | 3 | 3 | 6 |
| 4 | TGME49_313580.P3713 | cytochrome b5 family heme/steroid binding domain-containing protein | 8 | 6 | 5 | 5 | 3 | 4 | 4 | 6 |
| 5 | TGME49_313660.P3717 | hypothetical protein | 9 | 6 | 3 | 3 | 3 | 3 | 3 | 6 |
| 4 | TGME49_313700-t26_1-p1 | hypothetical protein | 9 | 7 | 6 | 5 | 4 | 3 | 3 | 6 |
| 4 | TGME49_313750-t26_1-p1 | hypothetical protein | 8 | 7 | 6 | 3 | 5 | 4 | 3 | 6 |
| 5 | TGME49_313780-t26_1-p1 | hypothetical protein | 9 | 7 | 5 | 4 | 3 | 3 | 3 | 6 |
| 5 | TGME49_313900.P3725 | non-specific serine/threonine protein kinase | 10 | 8 | 5 | 4 | 4 | 3 | 3 | 6 |
| 4 | TGME49_314070.P3732 | hypothetical protein | 7 | 7 | 6 | 5 | 3 | 3 | 3 | 6 |
| 10 | TGME49_314265-t26_1-p1 | hypothetical protein | 10 | 9 | 5 | 4 | 4 | 5 | 4 | 6 |
| 4 | TGME49_314370.P3740 | hypothetical protein | 10 | 7 | 6 | 5 | 4 | 3 | 5 | 6 |
| 5 | TGME49_314390-t26_1-p1 | hypothetical protein | 9 | 6 | 4 | 3 | 3 | 3 | 3 | 6 |
| 4 | TGME49_314450-t26_1-p1 | hypothetical protein | 8 | 7 | 6 | 5 | 4 | 4 | 4 | 6 |
| 4 | TGME49_314460-t26_1-p1 | hypothetical protein | 8 | 6 | 5 | 5 | 3 | 3 | 4 | 6 |
| 4 | TGME49_314470.P3744 | hypothetical protein | 8 | 7 | 6 | 5 | 4 | 3 | 4 | 6 |
| 5 | TGME49_314515-t26_1-p1 | hypothetical protein | 10 | 9 | 5 | 4 | 3 | 3 | 3 | 6 |
| 5 | TGME49_314920.P3754 | hypothetical protein | 10 | 9 | 5 | 4 | 4 | 3 | 3 | 6 |
| 10 | TGME49_315080-t26_1-p1 | hypothetical protein | 10 | 9 | 4 | 5 | 5 | 4 | 5 | 6 |
| 4 | TGME49_315110-t26_1-p1 | prefoldin subunit 5, putative | 7 | 6 | 6 | 5 | 5 | 4 | 4 | 6 |
| 4 | TGME49_315540.P3778 | calcium binding egf domain-containing protein | 8 | 6 | 7 | 6 | 3 | 3 | 3 | 6 |
| 4 | TGME49_315700-t26_1-p1 | hypothetical protein | 10 | 6 | 5 | 5 | 3 | 5 | 3 | 6 |
| 5 | TGME49_315885.P3791 | glycosyltransferase, putative | 10 | 6 | 5 | 3 | 4 | 3 | 3 | 6 |
| 10 | TGME49_315950.P3796 | cytoskeleton salvaging RING family member 1 | 10 | 10 | 3 | 4 | 4 | 3 | 4 | 6 |
| 5 | TGME49_315970.P3798 | lipoxygenase-like enzyme LOXL1 | 9 | 6 | 4 | 4 | 3 | 3 | 3 | 6 |
| 4 | TGME49_315980.P3799 | EREBP-4 family protein | 8 | 7 | 6 | 5 | 4 | 4 | 5 | 6 |
| 4 | TGME49_316090-t26_1-p1 | RNAse P Rpr2/Rpp21 subunit domain-containing protein | 9 | 7 | 6 | 5 | 4 | 3 | 3 | 6 |
| 10 | TGME49_316470-t26_1-p1 | hypothetical protein | 10 | 10 | 5 | 6 | 6 | 3 | 4 | 6 |
| 5 | TGME49_316510-t26_1-p1 | zinc finger (CCCH type) motif-containing protein | 10 | 8 | 3 | 4 | 3 | 3 | 3 | 6 |

| | | | | | | | | | | |
|---|---|---|---|---|---|---|---|---|---|---|
| 10 | TGME49_316620.P3816 | cell division cycle protein CDC20, putative | 10 | 9 | 4 | 6 | 3 | 4 | 5 | 6 |
| 5 | TGME49_316650-t26_1-p1 | hypothetical protein | 10 | 9 | 3 | 3 | 5 | 3 | 4 | 6 |
| 5 | TGME49_316760-t26_1-p1 | hypothetical protein | 9 | 6 | 5 | 3 | 4 | 3 | 3 | 6 |
| 5 | TGME49_318130-t26_1-p1 | Toxoplasma gondii family E protein | 9 | 8 | 3 | 3 | 3 | 3 | 3 | 6 |
| 4 | TGME49_318170-t26_1-p1 | hypothetical protein | 9 | 6 | 6 | 5 | 4 | 3 | 4 | 6 |
| 5 | TGME49_318210-t26_1-p1 | hypothetical protein | 10 | 8 | 5 | 4 | 4 | 3 | 3 | 6 |
| 5 | TGME49_318220.P3824 | hypothetical protein | 10 | 5 | 5 | 4 | 3 | 3 | 3 | 6 |
| 4 | TGME49_318275-t26_1-p1 | peptidyl-prolyl cis-trans isomerase, FKBP-type domain-containing protein | 9 | 7 | 6 | 5 | 5 | 4 | 4 | 6 |
| 4 | TGME49_318320-t26_1-p1 | ATP-dependent Clp endopeptidase, proteolytic subunit ClpP domain-containing protein | 8 | 6 | 6 | 4 | 4 | 4 | 3 | 6 |
| 4 | TGME49_318370-t26_1-p1 | hypothetical protein | 8 | 6 | 5 | 4 | 3 | 4 | 4 | 6 |
| 5 | TGME49_318390-t26_1-p1 | hypothetical protein | 10 | 8 | 3 | 3 | 3 | 5 | 5 | 6 |
| 4 | TGME49_318600-t26_1-p1 | DNA /pantothenate metabolism flavoprotein | 9 | 6 | 7 | 6 | 4 | 4 | 5 | 6 |
| 5 | TGME49_319090.P3844 | IgA-specific serine endopeptidase | 9 | 7 | 5 | 4 | 4 | 3 | 3 | 6 |
| 5 | TGME49_319320-t26_1-p1 | hypothetical protein | 10 | 9 | 3 | 4 | 3 | 3 | 3 | 6 |
| 5 | TGME49_319360-t26_1-p1 | SAG-related sequence SRS17A | 9 | 7 | 5 | 4 | 3 | 3 | 3 | 6 |
| 5 | TGME49_319380.P3847 | hypothetical protein | 10 | 7 | 5 | 4 | 4 | 4 | 3 | 6 |
| 5 | TGME49_319520-t26_1-p1 | hypothetical protein | 10 | 8 | 3 | 3 | 3 | 3 | 3 | 6 |
| 4 | TGME49_319550.P3850 | membrane protein C17G8.08c, putative | 8 | 7 | 7 | 5 | 4 | 3 | 4 | 6 |
| 5 | TGME49_319560.P3851 | microneme protein MIC3 | 8 | 7 | 5 | 3 | 3 | 3 | 3 | 6 |
| 5 | TGME49_319662.P3854 | histone lysine methyltransferase, SET, putative | 10 | 7 | 6 | 3 | 4 | 3 | 3 | 6 |
| 4 | TGME49_319940.P3864 | synaptosomal-associated protein 29 | 8 | 7 | 6 | 3 | 5 | 4 | 3 | 6 |
| 4 | TGME49_320005-t26_1-p1 | hypothetical protein | 8 | 6 | 6 | 5 | 5 | 4 | 4 | 6 |
| 4 | TGME49_320040-t26_1-p1 | CHORD protein | 7 | 7 | 6 | 3 | 5 | 3 | 3 | 6 |
| 4 | TGME49_320060-t26_1-p1 | DnaJ domain-containing protein | 9 | 5 | 6 | 4 | 4 | 4 | 3 | 6 |
| 5 | TGME49_320170-t26_1-p1 | SAG-related sequence SRS16E | 8 | 7 | 6 | 3 | 3 | 3 | 3 | 6 |
| 4 | TGME49_320230-t26_1-p1 | SAG-related sequence SRS15C | 8 | 7 | 6 | 4 | 3 | 3 | 3 | 6 |
| 4 | TGME49_320440-t26_1-p1 | exoribonuclease, putative | 10 | 7 | 6 | 5 | 4 | 4 | 5 | 6 |
| 5 | TGME49_320515.P3880 | hypothetical protein | 10 | 7 | 4 | 4 | 3 | 3 | 3 | 6 |
| 4 | TGME49_320530-t26_1-p1 | hypothetical protein | 8 | 6 | 6 | 3 | 4 | 3 | 3 | 6 |
| 4 | TGME49_320650-t26_1-p1 | ankyrin repeat-containing protein | 8 | 7 | 7 | 4 | 3 | 4 | 4 | 6 |
| 10 | TGME49_320710.P3886 | hypothetical protein | 8 | 6 | 3 | 7 | 3 | 3 | 3 | 6 |
| 4 | TGME49_321300.P3890 | autophagy-related protein ATG12 | 9 | 6 | 4 | 5 | 5 | 5 | 3 | 6 |
| 5 | TGME49_321390-t26_1-p1 | hypothetical protein | 9 | 6 | 5 | 3 | 3 | 3 | 3 | 6 |
| 5 | TGME49_321540-t26_1-p1 | hypothetical protein | 10 | 8 | 4 | 4 | 5 | 4 | 3 | 6 |
| 4 | TGME49_321680-t26_1-p1 | RNA-binding protein, putative | 9 | 7 | 6 | 5 | 4 | 4 | 4 | 6 |
| 4 | TGME49_500004.P7 | peptidase, putative | 8 | 7 | 5 | 5 | 3 | 4 | 4 | 6 |
| 5 | TGME49_500009.P12 | kinesin-like protein | 9 | 8 | 4 | 4 | 3 | 4 | 3 | 6 |
| 4 | TGME49_500081.P382 | hypothetical protein, conserved | 6 | 8 | 7 | 5 | 3 | 3 | 3 | 6 |
| 4 | TGME49_500083.P384 | hypothetical protein, conserved | 8 | 8 | 5 | 5 | 3 | 3 | 3 | 6 |
| 4 | TGME49_500087.P388 | hypothetical protein, conserved | 8 | 6 | 7 | 3 | 3 | 4 | 4 | 6 |
| 4 | TGME49_500099.P400 | hypothetical protein, conserved | 10 | 5 | 7 | 3 | 4 | 3 | 3 | 6 |
| 5 | TGME49_500115.P417 | EGF-like domain-containing protein | 8 | 8 | 3 | 3 | 3 | 4 | 4 | 6 |
| 5 | TGME49_500135.P437 | Rab3-GAP domain-containing protein | 9 | 6 | 5 | 4 | 4 | 3 | 3 | 6 |
| 4 | TGME49_500190.P194 | selenoprotein T, putative | 7 | 6 | 6 | 5 | 4 | 3 | 3 | 6 |
| 5 | TGME49_500200.P205 | intraconoidal microtubule-associated protein ICMAP3 | 9 | 6 | 4 | 3 | 3 | 3 | 3 | 6 |
| 5 | TGME49_500220.P225 | hypothetical protein, conserved | 10 | 9 | 3 | 3 | 3 | 3 | 3 | 6 |
| 10 | TGME49_500258.P47 | EF hand domain-containing protein | 10 | 7 | 5 | 6 | 4 | 4 | 3 | 6 |

| | | | | | | | | | | |
|---|---|---|---|---|---|---|---|---|---|---|
| 5 | TGME49_500264.P53 | hypothetical protein, conserved | 10 | 9 | 4 | 4 | 3 | 3 | 3 | 6 |
| 5 | TGME49_500273.P62 | hypothetical protein, conserved | 9 | 8 | 5 | 3 | 3 | 3 | 3 | 6 |
| 10 | TGME49_500277.P66 | hypothetical protein, conserved | 10 | 9 | 3 | 6 | 3 | 3 | 3 | 6 |
| 4 | TGME49_500292.P81 | hypothetical protein, conserved | 9 | 5 | 6 | 4 | 3 | 4 | 3 | 6 |
| 5 | TGME49_500309.P99 | WW domain-containing protein | 10 | 6 | 4 | 4 | 3 | 3 | 3 | 6 |
| 10 | TGME49_500310.P100 | nucleotidyltransferase, putative | 9 | 9 | 3 | 7 | 3 | 3 | 4 | 6 |
| 5 | TGME49_500348.P138 | hypothetical protein | 10 | 10 | 3 | 3 | 3 | 3 | 3 | 6 |
| 4 | TGME49_500364.P263 | apicomplexan kinetochore protein AKIT9, putative | 8 | 7 | 6 | 5 | 4 | 3 | 3 | 6 |
| 10 | TGME49_500394.P293 | CAP domain-containing protein | 9 | 8 | 4 | 7 | 3 | 4 | 4 | 6 |
| 5 | TGME49_500400.P299 | hypothetical protein, conserved | 9 | 7 | 6 | 3 | 3 | 3 | 3 | 6 |
| 10 | TGME49_500422.P321 | RNA-dependent RNA polymerase, putative | 10 | 10 | 4 | 9 | 3 | 4 | 5 | 6 |
| 5 | TGME49_500429.P328 | hypothetical protein, conserved | 10 | 9 | 5 | 3 | 3 | 3 | 3 | 6 |

| Kmean_cluster | GeneID (all protein TG) | Product Description | Decile_bitHH | Decile_bitNcan | Decile_bitBB | Decilebit_CS | Decilebit_SN3 | Decile bit_Ef | Decile bit_CC | Meta_cluster |
|---|---|---|---|---|---|---|---|---|---|---|
| 20 | TGME49_200010.P447 | dense granule protein GRA20 | 8 | 4 | 3 | 3 | 3 | 3 | 3 | 7 |
| 24 | TGME49_200240-t26_1-p1 | microneme protein MIC17B | 5 | 7 | 3 | 3 | 3 | 3 | 3 | 7 |
| 16 | TGME49_200310.P448 | hypothetical protein | 5 | 5 | 4 | 4 | 5 | 3 | 4 | 7 |
| 20 | TGME49_200320.P451 | hypoxanthine-xanthine-guanine phosphoribosyl transferase HXGPRT | 8 | 3 | 3 | 3 | 7 | 3 | 3 | 7 |
| 26 | TGME49_200360-t26_1-p1 | hypothetical protein | 6 | 4 | 4 | 3 | 3 | 3 | 3 | 7 |
| 26 | TGME49_200590-t26_1-p1 | Toxoplasma gondii family C protein | 6 | 3 | 3 | 3 | 3 | 3 | 3 | 7 |
| 26 | TGME49_200600.P463 | hypothetical protein | 5 | 3 | 3 | 3 | 3 | 3 | 3 | 7 |
| 16 | TGME49_201170-t26_1-p1 | hypothetical protein | 6 | 5 | 5 | 5 | 4 | 4 | 3 | 7 |
| 20 | TGME49_201180.P465 | megakaryocyte stimulating factor MSF | 8 | 4 | 3 | 3 | 3 | 3 | 3 | 7 |
| 26 | TGME49_201730-t26_1-p1 | hypothetical protein | 5 | 3 | 3 | 3 | 3 | 3 | 3 | 7 |
| 16 | TGME49_201750-t26_1-p1 | Tctex-1 family protein | 6 | 6 | 6 | 5 | 5 | 3 | 4 | 7 |
| 20 | TGME49_201810-t26_1-p1 | hypothetical protein | 8 | 3 | 4 | 3 | 3 | 3 | 3 | 7 |
| 24 | TGME49_201860.P477 | hypothetical protein | 6 | 6 | 4 | 4 | 4 | 4 | 3 | 7 |
| 24 | TGME49_202090-t26_1-p1 | hypothetical protein | 7 | 6 | 5 | 4 | 3 | 3 | 3 | 7 |
| 16 | TGME49_202100.P482 | hypothetical protein | 6 | 5 | 5 | 4 | 4 | 3 | 3 | 7 |
| 24 | TGME49_202110-t26_1-p1 | hypothetical protein | 6 | 5 | 5 | 4 | 3 | 3 | 3 | 7 |
| 26 | TGME49_202205-t26_1-p1 | hypothetical protein | 6 | 3 | 5 | 3 | 3 | 3 | 3 | 7 |
| 9 | TGME49_202255.P487 | hypothetical protein | 8 | 3 | 7 | 4 | 3 | 4 | 3 | 7 |
| 9 | TGME49_202285-t26_1-p1 | EGF family domain-containing protein | 8 | 3 | 6 | 3 | 3 | 5 | 3 | 7 |
| 26 | TGME49_202450.P490 | hypothetical protein | 6 | 4 | 4 | 4 | 3 | 3 | 3 | 7 |
| 16 | TGME49_202568-t26_1-p1 | FAM32A domain-containing protein | 5 | 3 | 5 | 5 | 4 | 4 | 4 | 7 |
| 26 | TGME49_202600-t26_1-p1 | hypothetical protein | 5 | 3 | 3 | 3 | 3 | 3 | 3 | 7 |
| 20 | TGME49_202620.P497 | dense granule protein GRA64 | 8 | 4 | 4 | 3 | 3 | 3 | 3 | 7 |
| 20 | TGME49_202730-t26_1-p1 | hypothetical protein | 10 | 4 | 4 | 3 | 3 | 3 | 3 | 7 |
| 24 | TGME49_202760-t26_1-p1 | hypothetical protein | 8 | 6 | 5 | 3 | 3 | 3 | 3 | 7 |
| 16 | TGME49_202810-t26_1-p1 | mitochondrial import inner membrane translocase subunit TIM14 | 6 | 5 | 5 | 3 | 5 | 3 | 5 | 7 |
| 24 | TGME49_202890.P504 | hypothetical protein | 8 | 6 | 5 | 3 | 4 | 3 | 4 | 7 |
| 24 | TGME49_202930.P506 | IMC sutures component ISC5 | 8 | 6 | 5 | 3 | 4 | 3 | 4 | 7 |
| 24 | TGME49_202940-t26_1-p1 | hypothetical protein | 8 | 6 | 4 | 3 | 3 | 3 | 3 | 7 |
| 24 | TGME49_202960-t26_1-p1 | glycine cleavage H-protein | 8 | 6 | 5 | 3 | 4 | 3 | 4 | 7 |
| 20 | TGME49_202975.P508 | hypothetical protein | 8 | 3 | 3 | 3 | 3 | 3 | 3 | 7 |
| 26 | TGME49_203070-t26_1-p1 | hypothetical protein | 7 | 3 | 4 | 3 | 3 | 4 | 3 | 7 |
| 20 | TGME49_203250-t26_1-p1 | hypothetical protein | 10 | 4 | 3 | 3 | 3 | 3 | 3 | 7 |
| 26 | TGME49_203290.P516 | dense granule protein GRA34 | 7 | 4 | 3 | 3 | 3 | 3 | 3 | 7 |
| 26 | TGME49_203310.P517 | dense granule protein GRA7 | 6 | 4 | 4 | 3 | 3 | 3 | 3 | 7 |
| 9 | TGME49_203375-t26_1-p1 | cpw-wpc domain-containing protein | 8 | 3 | 7 | 3 | 3 | 5 | 3 | 7 |
| 20 | TGME49_203450.P524 | brain colonization protein BCP1 | 8 | 3 | 3 | 3 | 3 | 3 | 3 | 7 |
| 26 | TGME49_203460-t26_1-p1 | mitochondrial distribution and morphology protein, putative | 5 | 3 | 5 | 4 | 3 | 3 | 3 | 7 |
| 16 | TGME49_203505.P525 | hypothetical protein | 6 | 3 | 6 | 6 | 3 | 3 | 4 | 7 |
| 16 | TGME49_203630.P529 | ribosomal protein RPL44 | 5 | 5 | 5 | 3 | 5 | 5 | 5 | 7 |
| 26 | TGME49_203682-t26_1-p1 | hypothetical protein | 6 | 3 | 3 | 3 | 3 | 3 | 3 | 7 |
| 24 | TGME49_203720-t26_1-p1 | vitamin k epoxide reductase family protein | 7 | 6 | 6 | 3 | 3 | 3 | 3 | 7 |
| 26 | TGME49_203740-t26_1-p1 | hypothetical protein | 7 | 3 | 3 | 3 | 3 | 3 | 3 | 7 |
| 26 | TGME49_203770-t26_1-p1 | hypothetical protein | 7 | 3 | 4 | 4 | 3 | 3 | 3 | 7 |
| 20 | TGME49_203790.P532 | hypothetical protein | 8 | 5 | 3 | 3 | 3 | 3 | 3 | 7 |

| | | | | | | | | | | |
|---|---|---|---|---|---|---|---|---|---|---|
| 24 | TGME49_203930-t26_1-p1 | hypothetical protein | 8 | 7 | 3 | 4 | 4 | 3 | 3 | 7 |
| 24 | TGME49_203990-t26_1-p1 | rhoptry protein ROP12 | 7 | 6 | 4 | 4 | 3 | 3 | 3 | 7 |
| 26 | TGME49_204000-t26_1-p1 | hypothetical protein | 7 | 4 | 3 | 3 | 3 | 3 | 3 | 7 |
| 26 | TGME49_204010-t26_1-p1 | hypothetical protein | 5 | 3 | 3 | 3 | 3 | 3 | 3 | 7 |
| 20 | TGME49_204080-t26_1-p1 | histidine acid phosphatase superfamily protein | 10 | 3 | 3 | 5 | 3 | 4 | 4 | 7 |
| 20 | TGME49_204270.P545 | dense granule protein GRA60 | 9 | 5 | 3 | 3 | 3 | 3 | 3 | 7 |
| 24 | TGME49_204320-t26_1-p1 | hypothetical protein | 8 | 6 | 4 | 4 | 3 | 3 | 3 | 7 |
| 26 | TGME49_204330-t26_1-p1 | hypothetical protein | 7 | 3 | 3 | 3 | 3 | 3 | 3 | 7 |
| 9 | TGME49_204340.P548 | dense granule protein GRA54 | 10 | 3 | 7 | 4 | 3 | 3 | 3 | 7 |
| 16 | TGME49_204370.P549 | mitochondrial pyruvate carrier protein 2 | 6 | 6 | 6 | 4 | 4 | 3 | 3 | 7 |
| 26 | TGME49_204490.P555 | hypothetical protein | 5 | 4 | 4 | 3 | 3 | 3 | 3 | 7 |
| 20 | TGME49_204500-t26_1-p1 | hypothetical protein | 8 | 5 | 3 | 3 | 3 | 3 | 3 | 7 |
| 24 | TGME49_204520-t26_1-p1 | hypothetical protein | 7 | 6 | 5 | 4 | 3 | 3 | 3 | 7 |
| 24 | TGME49_205020-t26_1-p1 | hypothetical protein | 8 | 7 | 3 | 3 | 3 | 3 | 3 | 7 |
| 16 | TGME49_205030-t26_1-p1 | protein transport protein BET1, putative | 6 | 5 | 5 | 5 | 5 | 3 | 4 | 7 |
| 24 | TGME49_205070-t26_1-p1 | hypothetical protein | 6 | 5 | 4 | 3 | 3 | 3 | 3 | 7 |
| 26 | TGME49_205080.P559 | hypothetical protein | 7 | 3 | 3 | 3 | 3 | 3 | 4 | 7 |
| 26 | TGME49_205170.P562 | hypothetical protein | 6 | 5 | 3 | 3 | 3 | 3 | 3 | 7 |
| 26 | TGME49_205210-t26_1-p1 | hypothetical protein | 6 | 4 | 4 | 3 | 3 | 3 | 3 | 7 |
| 9 | TGME49_205250-t26_1-p1 | rhoptry protein ROP18 | 9 | 3 | 6 | 4 | 3 | 4 | 3 | 7 |
| 20 | TGME49_205398-t26_1-p1 | DNA repair and recombination protein RAD54, putative | 8 | 3 | 3 | 3 | 3 | 5 | 3 | 7 |
| 26 | TGME49_205450-t26_1-p1 | hypothetical protein | 6 | 4 | 3 | 3 | 3 | 3 | 3 | 7 |
| 9 | TGME49_205625-t26_1-p1 | transporter, major facilitator family protein | 9 | 3 | 5 | 5 | 4 | 4 | 4 | 7 |
| 20 | TGME49_205662-t26_1-p1 | LCCL domain-containing protein | 8 | 3 | 3 | 3 | 3 | 3 | 3 | 7 |
| 26 | TGME49_206290-t26_1-p1 | hypothetical protein | 5 | 3 | 3 | 3 | 3 | 3 | 3 | 7 |
| 9 | TGME49_206330-t26_1-p1 | PHD-finger domain-containing protein | 10 | 5 | 6 | 4 | 4 | 4 | 4 | 7 |
| 26 | TGME49_206340-t26_1-p1 | hypothetical protein | 6 | 5 | 3 | 3 | 3 | 3 | 3 | 7 |
| 9 | TGME49_206360.P581 | peptidyl-tRNA hydrolase domain-containing protein | 9 | 3 | 5 | 5 | 4 | 3 | 3 | 7 |
| 24 | TGME49_206440-t26_1-p1 | cpw-wpc domain-containing protein | 7 | 6 | 5 | 3 | 3 | 4 | 3 | 7 |
| 9 | TGME49_206480-t26_1-p1 | succinate dehydrogenase subunit SDH23 | 7 | 3 | 6 | 5 | 5 | 4 | 4 | 7 |
| 16 | TGME49_206680-t26_1-p1 | hypothetical protein | 7 | 3 | 5 | 5 | 4 | 4 | 4 | 7 |
| 26 | TGME49_207010.P599 | SAG-related sequence SRS48K | 7 | 3 | 3 | 3 | 3 | 3 | 3 | 7 |
| 24 | TGME49_207015.P600 | SRS domain-containing protein | 6 | 6 | 3 | 3 | 3 | 3 | 3 | 7 |
| 16 | TGME49_207020.P601 | hypothetical protein | 6 | 3 | 6 | 5 | 4 | 5 | 4 | 7 |
| 26 | TGME49_207050-t26_1-p1 | hypothetical protein | 7 | 4 | 3 | 3 | 3 | 3 | 3 | 7 |
| 20 | TGME49_207140.P607 | SAG-related sequence SRS49B | 8 | 3 | 5 | 3 | 3 | 3 | 3 | 7 |
| 24 | TGME49_207160-t26_1-p1 | SAG-related sequence SRS49D | 6 | 6 | 4 | 4 | 3 | 3 | 3 | 7 |
| 24 | TGME49_207170.P609 | cytochrome b-c1 complex subunit 12 | 6 | 5 | 5 | 3 | 4 | 3 | 3 | 7 |
| 26 | TGME49_207210-t26_1-p1 | hypothetical protein | 5 | 5 | 5 | 3 | 3 | 3 | 3 | 7 |
| 16 | TGME49_207240.P610 | hypothetical protein | 5 | 5 | 5 | 5 | 4 | 4 | 4 | 7 |
| 9 | TGME49_207250.P611 | electron-transfer-flavoprotein, alpha polypeptide | 9 | 3 | 7 | 3 | 3 | 6 | 5 | 7 |
| 26 | TGME49_207420.P615 | hypothetical protein | 6 | 3 | 4 | 4 | 4 | 4 | 4 | 7 |

| | | | | | | | | | | |
|---|---|---|---|---|---|---|---|---|---|---|
| 26 | TGME49_207650-t26_1-p1 | OTU domain-containing protein OTUD1A | 7 | 3 | 3 | 3 | 3 | 3 | 3 | 7 |
| 16 | TGME49_207690.P625 | programmed cell death 5 protein | 5 | 5 | 5 | 4 | 4 | 4 | 4 | 7 |
| 16 | TGME49_207790.P626 | enkurin, putative | 6 | 6 | 5 | 5 | 4 | 5 | 4 | 7 |
| 20 | TGME49_207920.P630 | hypothetical protein | 9 | 5 | 3 | 3 | 4 | 4 | 3 | 7 |
| 24 | TGME49_207950.P632 | hypothetical protein | 7 | 5 | 4 | 4 | 3 | 3 | 3 | 7 |
| 26 | TGME49_207955-t26_1-p1 | hypothetical protein | 6 | 3 | 3 | 4 | 3 | 3 | 3 | 7 |
| 16 | TGME49_208300-t26_1-p1 | hypothetical protein | 5 | 3 | 5 | 4 | 4 | 4 | 4 | 7 |
| 20 | TGME49_208330-t26_1-p1 | DUF2981 domain-containing protein | 8 | 3 | 5 | 3 | 3 | 3 | 3 | 7 |
| 20 | TGME49_208380-t26_1-p1 | hypothetical protein | 8 | 3 | 5 | 3 | 3 | 3 | 3 | 7 |
| 24 | TGME49_208430-t26_1-p1 | serine proteinase inhibitor PI-2, putative | 7 | 7 | 6 | 3 | 3 | 3 | 3 | 7 |
| 24 | TGME49_208450.P644 | protease inhibitor PI2 | 8 | 6 | 3 | 4 | 3 | 3 | 3 | 7 |
| 24 | TGME49_208550.P647 | hypothetical protein | 7 | 6 | 5 | 4 | 3 | 3 | 3 | 7 |
| 24 | TGME49_208770-t26_1-p1 | hypothetical protein | 6 | 6 | 5 | 4 | 3 | 3 | 3 | 7 |
| 26 | TGME49_208850-t26_1-p1 | SAG-related sequence SRS11 | 6 | 5 | 3 | 4 | 3 | 3 | 3 | 7 |
| 26 | TGME49_208980.P660 | hypothetical protein | 7 | 4 | 3 | 3 | 3 | 3 | 3 | 7 |
| 24 | TGME49_209070.P666 | hypothetical protein | 8 | 6 | 5 | 3 | 4 | 3 | 4 | 7 |
| 16 | TGME49_209120.P669 | PAT complex subunit Asterix, putative | 5 | 5 | 5 | 4 | 4 | 3 | 3 | 7 |
| 16 | TGME49_209290.P676 | ribosomal protein RPS28 | 5 | 5 | 5 | 3 | 4 | 3 | 4 | 7 |
| 16 | TGME49_209515.P682 | hypothetical protein | 6 | 3 | 3 | 8 | 6 | 3 | 3 | 7 |
| 26 | TGME49_209530-t26_1-p1 | hypothetical protein | 6 | 4 | 3 | 4 | 3 | 3 | 3 | 7 |
| 26 | TGME49_209583-t26_1-p1 | hypothetical protein | 6 | 3 | 3 | 4 | 3 | 3 | 3 | 7 |
| 20 | TGME49_209585-t26_1-p1 | hypothetical protein | 8 | 3 | 4 | 5 | 3 | 3 | 3 | 7 |
| 24 | TGME49_209640.P688 | hypothetical protein | 8 | 6 | 4 | 4 | 3 | 3 | 3 | 7 |
| 20 | TGME49_209705-t26_1-p1 | hypothetical protein | 8 | 3 | 3 | 3 | 3 | 3 | 3 | 7 |
| 24 | TGME49_209740-t26_1-p1 | hypothetical protein | 7 | 5 | 4 | 4 | 3 | 3 | 3 | 7 |
| 24 | TGME49_209755-t26_1-p1 | cyst matrix protein MAG2 | 8 | 6 | 3 | 3 | 3 | 3 | 3 | 7 |
| 16 | TGME49_209900-t26_1-p1 | hypothetical protein | 6 | 6 | 6 | 5 | 5 | 4 | 4 | 7 |
| 26 | TGME49_209930-t26_1-p1 | hypothetical protein | 6 | 3 | 5 | 5 | 3 | 3 | 3 | 7 |
| 20 | TGME49_210330-t26_1-p1 | SAG-related sequence SRS37B | 8 | 3 | 5 | 3 | 3 | 3 | 3 | 7 |
| 16 | TGME49_210408.P710 | HMG (high mobility group) box domain-containing protein | 5 | 5 | 5 | 5 | 5 | 4 | 4 | 7 |
| 9 | TGME49_210490-t26_1-p1 | hypothetical protein | 10 | 3 | 7 | 6 | 5 | 3 | 3 | 7 |
| 24 | TGME49_210600.P711 | KRUF family protein | 7 | 7 | 3 | 3 | 3 | 3 | 3 | 7 |
| 20 | TGME49_210678-t26_1-p1 | OTU domain-containing protein OTUD1D | 9 | 3 | 3 | 3 | 3 | 3 | 3 | 7 |
| 26 | TGME49_210740.P715 | hypothetical protein | 7 | 3 | 3 | 3 | 3 | 3 | 3 | 7 |
| 24 | TGME49_210770-t26_1-p1 | hypothetical protein | 8 | 7 | 4 | 3 | 3 | 3 | 3 | 7 |
| 16 | TGME49_210787-t26_1-p1 | Erv1 / Alr family protein | 6 | 3 | 5 | 5 | 4 | 3 | 3 | 7 |
| 16 | TGME49_210810.P719 | conoid associated protein CAP1 | 6 | 3 | 5 | 5 | 3 | 4 | 3 | 7 |
| 16 | TGME49_211040-t26_1-p1 | Sec61beta family protein | 5 | 5 | 5 | 4 | 4 | 4 | 4 | 7 |
| 16 | TGME49_211060.P725 | hypothetical protein | 6 | 5 | 5 | 5 | 3 | 3 | 4 | 7 |
| 16 | TGME49_211240-t26_1-p1 | peptidyl-tRNA hydrolase, putative | 7 | 6 | 5 | 5 | 5 | 4 | 4 | 7 |
| 24 | TGME49_211290-t26_1-p1 | rhoptry protein ROP15 | 8 | 6 | 5 | 4 | 3 | 3 | 3 | 7 |
| 24 | TGME49_211320-t26_1-p1 | hypothetical protein | 8 | 7 | 4 | 3 | 3 | 3 | 3 | 7 |
| 9 | TGME49_211385.P735 | hypothetical protein | 8 | 3 | 8 | 6 | 6 | 4 | 4 | 7 |
| 24 | TGME49_211420.P738 | serine/arginine-rich splicing factor SR3 | 6 | 5 | 3 | 3 | 5 | 3 | 3 | 7 |
| 24 | TGME49_211630-t26_1-p1 | hypothetical protein | 7 | 5 | 4 | 4 | 3 | 3 | 3 | 7 |
| 20 | TGME49_211650.P745 | hypothetical protein | 9 | 3 | 3 | 4 | 3 | 4 | 3 | 7 |

| | | | | | | | | | | |
|---|---|---|---|---|---|---|---|---|---|---|
| 26 | TGME49_211675-t26_1-p1 | hypothetical protein | 7 | 3 | 3 | 3 | 3 | 3 | 3 | 7 |
| 16 | TGME49_211710-t26_1-p1 | TB2/DP1, HVA22 family protein | 7 | 6 | 6 | 3 | 5 | 3 | 4 | 7 |
| 24 | TGME49_212160-t26_1-p1 | hypothetical protein | 7 | 7 | 6 | 3 | 3 | 3 | 3 | 7 |
| 24 | TGME49_212210.P757 | hypothetical protein | 7 | 6 | 4 | 4 | 3 | 3 | 3 | 7 |
| 26 | TGME49_212275.P762 | hypothetical protein | 7 | 3 | 3 | 5 | 3 | 3 | 5 | 7 |
| 26 | TGME49_212410-t26_1-p1 | dense granule protein GRA11A | 7 | 3 | 3 | 3 | 3 | 3 | 4 | 7 |
| 20 | TGME49_212710-t26_1-p1 | phenylalanine-4-hydroxylase | 9 | 3 | 3 | 3 | 3 | 3 | 3 | 7 |
| 16 | TGME49_212760-t26_1-p1 | hypothetical protein | 5 | 3 | 3 | 5 | 5 | 3 | 4 | 7 |
| 26 | TGME49_212830-t26_1-p1 | expressed protein JLP1 | 5 | 3 | 3 | 3 | 3 | 3 | 3 | 7 |
| 24 | TGME49_212860-t26_1-p1 | hypothetical protein | 8 | 5 | 3 | 5 | 4 | 4 | 4 | 7 |
| 24 | TGME49_212910-t26_1-p1 | rhomboid protease ROM3 | 7 | 7 | 5 | 4 | 3 | 4 | 3 | 7 |
| 24 | TGME49_212937.P770 | hypothetical protein | 8 | 6 | 5 | 4 | 4 | 3 | 3 | 7 |
| 26 | TGME49_213050.P778 | hypothetical protein | 6 | 3 | 3 | 3 | 3 | 3 | 3 | 7 |
| 20 | TGME49_213067.P780 | dense granule protein GRA36 | 8 | 3 | 4 | 3 | 3 | 3 | 3 | 7 |
| 24 | TGME49_213115.P783 | hypothetical protein | 8 | 6 | 5 | 4 | 4 | 3 | 3 | 7 |
| 24 | TGME49_213240-t26_1-p1 | hypothetical protein | 8 | 5 | 4 | 3 | 4 | 3 | 3 | 7 |
| 24 | TGME49_213280.P785 | SAG-related sequence SRS25 | 7 | 6 | 5 | 3 | 3 | 3 | 3 | 7 |
| 16 | TGME49_213410-t26_1-p1 | small nuclear ribonucleoprotein f (snrnp-f), putative | 5 | 5 | 3 | 3 | 5 | 5 | 5 | 7 |
| 9 | TGME49_213460.P792 | hypothetical protein | 10 | 3 | 7 | 3 | 3 | 3 | 3 | 7 |
| 20 | TGME49_213560-t26_1-p1 | hypothetical protein | 9 | 3 | 5 | 3 | 3 | 4 | 3 | 7 |
| 16 | TGME49_213590.P797 | hypothetical protein | 5 | 5 | 5 | 5 | 5 | 4 | 3 | 7 |
| 20 | TGME49_213610.P799 | hypothetical protein | 10 | 3 | 3 | 3 | 5 | 3 | 3 | 7 |
| 16 | TGME49_213690.P801 | E3 ubiquitin-protein ligase RBX1, putative | 6 | 6 | 5 | 5 | 4 | 5 | 3 | 7 |
| 24 | TGME49_213746.P804 | hypothetical protein | 7 | 6 | 3 | 4 | 4 | 3 | 3 | 7 |
| 16 | TGME49_213810-t26_1-p1 | iron-sulfur cluster protein ISCA | 6 | 6 | 5 | 5 | 3 | 3 | 4 | 7 |
| 20 | TGME49_213820-t26_1-p1 | hypothetical protein | 10 | 4 | 3 | 3 | 3 | 3 | 3 | 7 |
| 16 | TGME49_213850-t26_1-p1 | MORN repeat-containing protein | 8 | 4 | 5 | 3 | 6 | 5 | 3 | 7 |
| 9 | TGME49_213860.P810 | hypothetical protein | 8 | 4 | 7 | 7 | 3 | 3 | 3 | 7 |
| 24 | TGME49_213880.P811 | hypothetical protein | 7 | 5 | 4 | 3 | 3 | 3 | 3 | 7 |
| 16 | TGME49_213940-t26_1-p1 | CHCH domain-containing protein | 6 | 5 | 5 | 5 | 5 | 4 | 4 | 7 |
| 26 | TGME49_214080-t26_1-p1 | toxofilin | 6 | 4 | 3 | 3 | 3 | 3 | 3 | 7 |
| 26 | TGME49_214130-t26_1-p1 | hypothetical protein | 7 | 4 | 4 | 3 | 3 | 3 | 3 | 7 |
| 16 | TGME49_214220.P819 | hypothetical protein | 5 | 5 | 4 | 4 | 4 | 3 | 4 | 7 |
| 16 | TGME49_214250.P820 | cytochrome b-c1 complex subunit 11 | 5 | 5 | 5 | 3 | 3 | 3 | 4 | 7 |
| 24 | TGME49_214330-t26_1-p1 | DnaJ domain-containing protein | 8 | 6 | 5 | 3 | 3 | 3 | 3 | 7 |
| 26 | TGME49_214410-t26_1-p1 | hypothetical protein | 6 | 3 | 3 | 3 | 3 | 3 | 3 | 7 |
| 26 | TGME49_214420.P824 | hypothetical protein | 6 | 3 | 3 | 3 | 3 | 3 | 3 | 7 |
| 26 | TGME49_214430-t26_1-p1 | hypothetical protein | 7 | 3 | 3 | 3 | 3 | 3 | 3 | 7 |
| 16 | TGME49_214480.P826 | replication factor a protein 3 protein | 6 | 6 | 5 | 4 | 4 | 4 | 3 | 7 |
| 26 | TGME49_214575-t26_1-p1 | hypothetical protein | 5 | 3 | 3 | 3 | 3 | 3 | 3 | 7 |
| 16 | TGME49_214930.P837 | ATP synthase-associated protein | 6 | 6 | 5 | 4 | 5 | 3 | 3 | 7 |
| 26 | TGME49_214950.P839 | hypothetical protein | 6 | 3 | 3 | 3 | 3 | 3 | 3 | 7 |
| 20 | TGME49_215080-t26_1-p1 | hypothetical protein | 10 | 4 | 5 | 4 | 4 | 3 | 3 | 7 |
| 26 | TGME49_215165-t26_1-p1 | hypothetical protein | 5 | 3 | 3 | 3 | 3 | 4 | 3 | 7 |
| 24 | TGME49_215220.P849 | dense granule protein GRA22 | 8 | 7 | 3 | 5 | 3 | 3 | 3 | 7 |
| 9 | TGME49_215250-t26_1-p1 | thiamin pyrophosphokinase, catalytic domain-containing protein | 10 | 5 | 7 | 6 | 5 | 4 | 5 | 7 |
| 20 | TGME49_215300-t26_1-p1 | hypothetical protein | 9 | 5 | 5 | 4 | 3 | 3 | 3 | 7 |

| | | | | | | | | | | |
|---|---|---|---|---|---|---|---|---|---|---|
| 9 | TGME49_215328.P852 | transporter, putative | 9 | 3 | 7 | 7 | 5 | 3 | 3 | 7 |
| 26 | TGME49_215347-t26_1-p1 | hypothetical protein | 5 | 3 | 3 | 3 | 3 | 3 | 3 | 7 |
| 16 | TGME49_215350-t26_1-p1 | hypothetical protein | 6 | 5 | 5 | 5 | 5 | 4 | 3 | 7 |
| 9 | TGME49_215385-t26_1-p1 | transporter, major facilitator family protein | 9 | 3 | 8 | 8 | 3 | 5 | 5 | 7 |
| 16 | TGME49_215390-t26_1-p1 | TIM10 family protein, putative | 5 | 5 | 4 | 4 | 4 | 3 | 4 | 7 |
| 16 | TGME49_215530.P861 | hypothetical protein | 6 | 5 | 5 | 5 | 4 | 3 | 3 | 7 |
| 26 | TGME49_215540-t26_1-p1 | hypothetical protein | 6 | 4 | 3 | 4 | 3 | 3 | 3 | 7 |
| 26 | TGME49_215600-t26_1-p1 | hypothetical protein | 7 | 3 | 5 | 4 | 4 | 3 | 3 | 7 |
| 16 | TGME49_215610.P864 | ATP synthase-associated protein | 6 | 5 | 5 | 3 | 5 | 3 | 3 | 7 |
| 16 | TGME49_215930.P869 | mediator complex subunit MED21 | 6 | 6 | 5 | 5 | 4 | 4 | 4 | 7 |
| 26 | TGME49_215980-t26_1-p1 | hypothetical protein | 6 | 5 | 3 | 3 | 3 | 3 | 3 | 7 |
| 20 | TGME49_216053-t26_1-p1 | hypothetical protein | 8 | 3 | 4 | 5 | 4 | 4 | 3 | 7 |
| 24 | TGME49_216055-t26_1-p1 | hypothetical protein | 8 | 5 | 5 | 3 | 4 | 3 | 3 | 7 |
| 9 | TGME49_216090-t26_1-p1 | hypothetical protein | 8 | 3 | 7 | 3 | 3 | 4 | 4 | 7 |
| 26 | TGME49_216155-t26_1-p1 | hypothetical protein | 6 | 3 | 3 | 3 | 3 | 3 | 3 | 7 |
| 16 | TGME49_216180-t26_1-p1 | hypothetical protein | 6 | 6 | 6 | 5 | 3 | 3 | 4 | 7 |
| 24 | TGME49_216270-t26_1-p1 | hypothetical protein | 7 | 6 | 5 | 3 | 3 | 3 | 3 | 7 |
| 16 | TGME49_216275-t26_1-p1 | ACR, YggU family COG1872 domain-containing protein | 7 | 3 | 5 | 5 | 4 | 4 | 4 | 7 |
| 20 | TGME49_216290-t26_1-p1 | hypothetical protein | 10 | 3 | 3 | 6 | 4 | 3 | 3 | 7 |
| 24 | TGME49_216370.P882 | golgi localizing protein GLP2 | 8 | 6 | 4 | 3 | 3 | 3 | 3 | 7 |
| 20 | TGME49_216435-t26_1-p1 | hypothetical protein | 8 | 4 | 4 | 4 | 4 | 3 | 3 | 7 |
| 24 | TGME49_216700-t26_1-p1 | peptide deformylase, putative | 7 | 6 | 6 | 3 | 3 | 3 | 3 | 7 |
| 26 | TGME49_216720-t26_1-p1 | hypothetical protein | 6 | 3 | 3 | 3 | 3 | 3 | 3 | 7 |
| 26 | TGME49_216770-t26_1-p1 | hypothetical protein | 5 | 3 | 3 | 3 | 3 | 3 | 3 | 7 |
| 16 | TGME49_216910-t26_1-p1 | hypothetical protein | 6 | 6 | 6 | 5 | 3 | 4 | 4 | 7 |
| 9 | TGME49_216940-t26_1-p1 | hypothetical protein | 8 | 3 | 7 | 7 | 3 | 6 | 3 | 7 |
| 24 | TGME49_217030-t26_1-p1 | hypothetical protein | 7 | 5 | 4 | 4 | 3 | 4 | 4 | 7 |
| 24 | TGME49_217040-t26_1-p1 | hypothetical protein | 7 | 7 | 6 | 3 | 3 | 3 | 3 | 7 |
| 26 | TGME49_217370-t26_1-p1 | hypothetical protein | 6 | 4 | 3 | 4 | 3 | 3 | 3 | 7 |
| 16 | TGME49_217450.P904 | general transcription factor IIH polypeptide 5 GTF2H5 | 5 | 5 | 5 | 5 | 3 | 4 | 3 | 7 |
| 24 | TGME49_217490-t26_1-p1 | hypothetical protein | 6 | 7 | 4 | 5 | 3 | 4 | 3 | 7 |
| 26 | TGME49_217520-t26_1-p1 | hypothetical protein | 5 | 4 | 3 | 3 | 3 | 3 | 3 | 7 |
| 9 | TGME49_217530.P908 | dense granule protein GRA63 | 9 | 4 | 7 | 6 | 3 | 4 | 4 | 7 |
| 26 | TGME49_217555-t26_1-p1 | hypothetical protein | 5 | 4 | 3 | 4 | 3 | 3 | 3 | 7 |
| 16 | TGME49_217560-t26_1-p1 | DNA-directed RNA polymerase II RPB10 | 5 | 5 | 5 | 3 | 3 | 4 | 4 | 7 |
| 24 | TGME49_217640.P914 | hypothetical protein | 6 | 5 | 5 | 4 | 3 | 3 | 3 | 7 |
| 26 | TGME49_217665.P917 | hypothetical protein | 6 | 3 | 4 | 3 | 4 | 3 | 3 | 7 |
| 26 | TGME49_217720-t26_1-p1 | hypothetical protein | 6 | 4 | 3 | 3 | 3 | 3 | 3 | 7 |
| 20 | TGME49_217730-t26_1-p1 | WD repeat-containing protein | 9 | 3 | 5 | 3 | 3 | 3 | 3 | 7 |
| 20 | TGME49_217800.P926 | hypothetical protein | 10 | 3 | 3 | 4 | 3 | 3 | 3 | 7 |
| 24 | TGME49_218520.P943 | microneme protein MIC6 | 8 | 6 | 4 | 4 | 4 | 3 | 3 | 7 |
| 24 | TGME49_218860-t26_1-p1 | hypothetical protein | 8 | 5 | 4 | 3 | 3 | 5 | 3 | 7 |
| 26 | TGME49_218955-t26_1-p1 | hypothetical protein | 6 | 3 | 3 | 3 | 3 | 3 | 3 | 7 |
| 20 | TGME49_219110-t26_1-p1 | hypothetical protein | 8 | 4 | 3 | 3 | 3 | 3 | 3 | 7 |
| 9 | TGME49_219160-t26_1-p1 | hypothetical protein | 8 | 3 | 7 | 3 | 3 | 3 | 3 | 7 |

| | | | | | | | | | | |
|---|---|---|---|---|---|---|---|---|---|---|
| 24 | TGME49_219240.P966 | Peptidyl-tRNA hydrolase PTH2 domain-containing protein | 8 | 5 | 3 | 5 | 5 | 3 | 3 | 7 |
| 26 | TGME49_219348-t26_1-p1 | SAG-related sequence SRS55M | 7 | 3 | 3 | 3 | 3 | 3 | 3 | 7 |
| 24 | TGME49_219610.P977 | hypothetical protein | 7 | 6 | 3 | 4 | 3 | 3 | 3 | 7 |
| 26 | TGME49_219690.P981 | hypothetical protein | 5 | 4 | 4 | 3 | 4 | 3 | 3 | 7 |
| 20 | TGME49_219730-t26_1-p1 | hypothetical protein | 9 | 3 | 3 | 3 | 3 | 3 | 3 | 7 |
| 16 | TGME49_219828.P985 | HMG (high mobility group) box domain-containing protein | 5 | 5 | 5 | 5 | 5 | 3 | 4 | 7 |
| 20 | TGME49_220025.P986 | hypothetical protein | 10 | 3 | 3 | 3 | 3 | 3 | 3 | 7 |
| 26 | TGME49_220080-t26_1-p1 | Toxoplasma gondii family C protein | 5 | 3 | 3 | 3 | 3 | 3 | 3 | 7 |
| 24 | TGME49_220140-t26_1-p1 | EF hand domain-containing protein | 6 | 6 | 5 | 4 | 4 | 3 | 3 | 7 |
| 20 | TGME49_220175-t26_1-p1 | hypothetical protein | 9 | 4 | 3 | 4 | 3 | 3 | 3 | 7 |
| 20 | TGME49_220240.P990 | dense granule protein GRA31 | 8 | 4 | 3 | 3 | 3 | 3 | 3 | 7 |
| 20 | TGME49_220330-t26_1-p1 | hypothetical protein | 9 | 5 | 4 | 3 | 3 | 3 | 3 | 7 |
| 9 | TGME49_220350-t26_1-p1 | tRNA ligases class II (D, K and N) domain-containing protein | 10 | 3 | 8 | 5 | 5 | 6 | 4 | 7 |
| 9 | TGME49_220450.P999 | ribonuclease HI protein | 9 | 4 | 6 | 5 | 4 | 4 | 4 | 7 |
| 20 | TGME49_220630-t26_1-p1 | hypothetical protein | 9 | 5 | 3 | 3 | 3 | 3 | 3 | 7 |
| 26 | TGME49_220650-t26_1-p1 | hypothetical protein | 6 | 3 | 3 | 3 | 3 | 3 | 3 | 7 |
| 24 | TGME49_220870-t26_1-p1 | hypothetical protein | 7 | 6 | 3 | 5 | 5 | 3 | 3 | 7 |
| 16 | TGME49_220950-t26_1-p1 | hypothetical protein | 7 | 3 | 6 | 3 | 5 | 3 | 4 | 7 |
| 9 | TGME49_221265.P1016 | hypothetical protein | 10 | 3 | 7 | 5 | 4 | 4 | 3 | 7 |
| 26 | TGME49_221295-t26_1-p1 | hypothetical protein | 7 | 3 | 5 | 4 | 3 | 3 | 3 | 7 |
| 24 | TGME49_221480.P1023 | hypothetical protein | 7 | 5 | 4 | 3 | 3 | 3 | 3 | 7 |
| 9 | TGME49_221550.P1027 | CMGC kinase, MAPK family, MEK kinase-related (incomplete catalytic triad) | 9 | 4 | 6 | 5 | 3 | 6 | 6 | 7 |
| 26 | TGME49_221585-t26_1-p1 | hypothetical protein | 5 | 3 | 3 | 3 | 3 | 3 | 3 | 7 |
| 20 | TGME49_221665-t26_1-p1 | hypothetical protein | 10 | 3 | 3 | 3 | 3 | 3 | 3 | 7 |
| 26 | TGME49_221675.P1035 | hypothetical protein | 7 | 3 | 3 | 3 | 3 | 3 | 3 | 7 |
| 20 | TGME49_221685-t26_1-p1 | hypothetical protein | 8 | 3 | 3 | 3 | 3 | 3 | 3 | 7 |
| 24 | TGME49_221818.P1037 | hypothetical protein | 7 | 6 | 6 | 4 | 3 | 3 | 3 | 7 |
| 26 | TGME49_221865-t26_1-p1 | hypothetical protein | 5 | 3 | 3 | 3 | 3 | 3 | 3 | 7 |
| 24 | TGME49_221930.P1042 | hypothetical protein | 7 | 6 | 6 | 3 | 3 | 3 | 3 | 7 |
| 26 | TGME49_222080-t26_1-p1 | hypothetical protein | 6 | 4 | 3 | 3 | 3 | 3 | 3 | 7 |
| 16 | TGME49_222090-t26_1-p1 | hypothetical protein | 6 | 6 | 5 | 5 | 4 | 3 | 3 | 7 |
| 20 | TGME49_222155.P1050 | hypothetical protein | 10 | 3 | 4 | 3 | 3 | 3 | 3 | 7 |
| 24 | TGME49_222180-t26_1-p1 | hypothetical protein | 7 | 5 | 3 | 3 | 3 | 3 | 3 | 7 |
| 16 | TGME49_222260-t26_1-p1 | hypothetical protein | 5 | 5 | 5 | 5 | 3 | 3 | 3 | 7 |
| 24 | TGME49_222280-t26_1-p1 | hypothetical protein | 7 | 5 | 5 | 4 | 4 | 3 | 3 | 7 |
| 16 | TGME49_222290-t26_1-p1 | U6 snRNA-associated Sm-like protein LSm1, putative | 7 | 6 | 5 | 3 | 5 | 3 | 5 | 7 |
| 20 | TGME49_222370-t26_1-p1 | SAG-related sequence SRS13 | 9 | 5 | 4 | 3 | 3 | 3 | 3 | 7 |
| 9 | TGME49_222385.P1058 | hypothetical protein | 8 | 3 | 6 | 3 | 4 | 4 | 4 | 7 |
| 24 | TGME49_222390-t26_1-p1 | hypothetical protein | 8 | 6 | 5 | 4 | 4 | 3 | 3 | 7 |
| 16 | TGME49_222850.P1064 | stress responsive a/b barrel domain-containing protein | 5 | 5 | 5 | 3 | 5 | 3 | 4 | 7 |
| 16 | TGME49_222870.P1066 | proteasome assembly chaperone 3, putative | 6 | 6 | 6 | 3 | 5 | 4 | 4 | 7 |
| 20 | TGME49_222935-t26_1-p1 | hypothetical protein | 8 | 3 | 3 | 3 | 3 | 3 | 3 | 7 |

| | | | | | | | | | | |
|---|---|---|---|---|---|---|---|---|---|---|
| 16 | TGME49_223000.P1070 | dynein light chain DLC8a | 5 | 5 | 5 | 5 | 5 | 3 | 5 | 7 |
| 16 | TGME49_223120-t26_1-p1 | cyclin-dependent kinase regulatory subunit protein | 6 | 5 | 5 | 5 | 3 | 5 | 4 | 7 |
| 26 | TGME49_223125.P1076 | plastid ubiquitin-like protein PUBL | 5 | 3 | 3 | 5 | 3 | 4 | 4 | 7 |
| 26 | TGME49_223400-t26_1-p1 | hypothetical protein | 5 | 5 | 3 | 3 | 3 | 3 | 3 | 7 |
| 9 | TGME49_223450.P1080 | ubiquitin carboxyl-terminal hydrolase | 10 | 3 | 6 | 5 | 6 | 4 | 6 | 7 |
| 16 | TGME49_223485.P1082 | protein MPODD, putative | 5 | 3 | 5 | 4 | 5 | 4 | 4 | 7 |
| 16 | TGME49_223530-t26_1-p1 | mitotic-spindle organizing protein 1, putative | 5 | 5 | 4 | 4 | 4 | 3 | 4 | 7 |
| 9 | TGME49_223610.P1086 | hypothetical protein | 10 | 4 | 7 | 5 | 4 | 3 | 3 | 7 |
| 9 | TGME49_223668.P1088 | LYAR-type C2HC zinc finger protein | 7 | 3 | 6 | 6 | 4 | 4 | 3 | 7 |
| 26 | TGME49_223685-t26_1-p1 | hypothetical protein | 6 | 3 | 3 | 4 | 3 | 3 | 3 | 7 |
| 24 | TGME49_223770.P1090 | hypothetical protein | 6 | 5 | 4 | 3 | 3 | 3 | 3 | 7 |
| 26 | TGME49_223915-t26_1-p1 | hypothetical protein | 7 | 3 | 3 | 3 | 3 | 3 | 3 | 7 |
| 24 | TGME49_223940.P1099 | gliding-associated protein GAP45 | 7 | 5 | 3 | 5 | 4 | 3 | 4 | 7 |
| 16 | TGME49_224020-t26_1-p1 | hypothetical protein | 6 | 5 | 5 | 3 | 4 | 4 | 3 | 7 |
| 16 | TGME49_224130-t26_1-p1 | RNA binding protein, putative | 6 | 6 | 5 | 5 | 5 | 4 | 4 | 7 |
| 9 | TGME49_224170-t26_1-p1 | SAG-related sequence SRS60A | 8 | 3 | 6 | 4 | 3 | 3 | 3 | 7 |
| 9 | TGME49_224180.P1109 | BTB domain-containing protein | 10 | 3 | 6 | 3 | 5 | 3 | 3 | 7 |
| 9 | TGME49_224235-t26_1-p1 | translation initiation factor IF-3 protein | 9 | 3 | 5 | 5 | 5 | 4 | 3 | 7 |
| 26 | TGME49_224275-t26_1-p1 | hypothetical protein | 5 | 3 | 3 | 3 | 3 | 3 | 3 | 7 |
| 16 | TGME49_224340-t26_1-p1 | hypothetical protein | 5 | 6 | 5 | 5 | 5 | 4 | 4 | 7 |
| 24 | TGME49_224820.P1130 | hypothetical protein | 7 | 5 | 5 | 4 | 4 | 3 | 3 | 7 |
| 16 | TGME49_224845.P1132 | hypothetical protein | 6 | 3 | 5 | 6 | 5 | 4 | 4 | 7 |
| 16 | TGME49_224905.P1136 | hypothetical protein | 6 | 3 | 5 | 5 | 5 | 4 | 3 | 7 |
| 16 | TGME49_224990.P1139 | hypothetical protein | 6 | 6 | 5 | 5 | 4 | 4 | 4 | 7 |
| 26 | TGME49_225120.P1152 | hypothetical protein | 6 | 4 | 4 | 3 | 3 | 3 | 3 | 7 |
| 24 | TGME49_225160.P1155 | hypothetical protein | 8 | 6 | 5 | 3 | 3 | 3 | 3 | 7 |
| 26 | TGME49_225170-t26_1-p1 | hypothetical protein | 6 | 3 | 5 | 5 | 3 | 3 | 3 | 7 |
| 9 | TGME49_225315-t26_1-p1 | hypothetical protein | 8 | 3 | 7 | 4 | 3 | 3 | 3 | 7 |
| 24 | TGME49_225350-t26_1-p1 | hypothetical protein | 8 | 6 | 4 | 3 | 3 | 3 | 3 | 7 |
| 20 | TGME49_225370.P1166 | hypothetical protein | 8 | 3 | 4 | 3 | 3 | 3 | 3 | 7 |
| 26 | TGME49_225413-t26_1-p1 | AMMECR1 domain-containing protein | 5 | 3 | 3 | 3 | 3 | 3 | 3 | 7 |
| 20 | TGME49_225440.P1167 | hypothetical protein | 10 | 5 | 4 | 4 | 3 | 3 | 3 | 7 |
| 16 | TGME49_225460.P1169 | hypothetical protein | 5 | 6 | 6 | 3 | 3 | 4 | 3 | 7 |
| 26 | TGME49_225555-t26_1-p1 | hypothetical protein | 5 | 3 | 4 | 4 | 4 | 3 | 3 | 7 |
| 26 | TGME49_225680-t26_1-p1 | hypothetical protein | 6 | 3 | 3 | 4 | 3 | 3 | 3 | 7 |
| 16 | TGME49_225720-t26_1-p1 | hypothetical protein | 7 | 5 | 5 | 5 | 4 | 5 | 3 | 7 |
| 16 | TGME49_225730.P1177 | ATP synthase-associated protein | 6 | 6 | 5 | 3 | 5 | 4 | 4 | 7 |
| 16 | TGME49_225790.P1181 | thioredoxin-like protein 2 | 7 | 6 | 5 | 5 | 5 | 5 | 3 | 7 |
| 26 | TGME49_225953.P1187 | hypothetical protein | 6 | 3 | 5 | 4 | 3 | 4 | 3 | 7 |
| 20 | TGME49_225955.P1189 | leucine rich repeat-containing protein | 8 | 3 | 3 | 6 | 3 | 5 | 4 | 7 |
| 24 | TGME49_225970-t26_1-p1 | mediator complex subunit MED31 | 7 | 6 | 6 | 3 | 3 | 3 | 4 | 7 |
| 16 | TGME49_226010.P1191 | pterin-4-alpha-carbinolamine dehydratase | 5 | 5 | 5 | 5 | 4 | 4 | 4 | 7 |
| 9 | TGME49_226072.P1195 | Ser/Thr phosphatase family protein | 9 | 3 | 8 | 4 | 4 | 5 | 5 | 7 |
| 16 | TGME49_226240.P1196 | bud site selection protein, putative | 5 | 5 | 5 | 5 | 5 | 4 | 4 | 7 |
| 9 | TGME49_226245.P1197 | hypothetical protein | 8 | 3 | 6 | 8 | 3 | 4 | 3 | 7 |
| 24 | TGME49_226260-t26_1-p1 | hypothetical protein | 7 | 5 | 3 | 3 | 3 | 3 | 3 | 7 |

| | | | | | | | | | | |
|---|---|---|---|---|---|---|---|---|---|---|
| 20 | TGME49_226330-t26_1-p1 | hypothetical protein | 10 | 3 | 5 | 3 | 3 | 3 | 3 | 7 |
| 24 | TGME49_226380.P1204 | dense granule protein GRA35 | 8 | 6 | 4 | 3 | 3 | 3 | 3 | 7 |
| 26 | TGME49_226490.P1211 | hypothetical protein | 6 | 5 | 3 | 4 | 3 | 3 | 3 | 7 |
| 16 | TGME49_226500-t26_1-p1 | succinate dehydrogenase subunit SDH11 | 5 | 5 | 5 | 5 | 4 | 4 | 4 | 7 |
| 20 | TGME49_226515.P1212 | hypothetical protein | 8 | 3 | 3 | 3 | 3 | 3 | 3 | 7 |
| 9 | TGME49_226532-t26_1-p1 | hypothetical protein | 8 | 3 | 7 | 3 | 5 | 4 | 3 | 7 |
| 24 | TGME49_226570.P1214 | hypothetical protein | 6 | 6 | 5 | 4 | 4 | 3 | 3 | 7 |
| 20 | TGME49_226670-t26_1-p1 | hypothetical protein | 8 | 4 | 4 | 3 | 3 | 3 | 3 | 7 |
| 16 | TGME49_226705-t26_1-p1 | hypothetical protein | 5 | 4 | 4 | 4 | 4 | 4 | 3 | 7 |
| 26 | TGME49_226825-t26_1-p1 | protein phosphatase inhibitor protein | 5 | 3 | 4 | 4 | 3 | 3 | 3 | 7 |
| 24 | TGME49_226860.P1226 | SAG-related sequence SRS67 | 7 | 6 | 5 | 4 | 3 | 3 | 3 | 7 |
| 24 | TGME49_226920.P1231 | hypothetical protein | 8 | 7 | 4 | 3 | 4 | 3 | 3 | 7 |
| 26 | TGME49_226925-t26_1-p1 | hypothetical protein | 5 | 3 | 3 | 3 | 3 | 3 | 3 | 7 |
| 26 | TGME49_226930-t26_1-p1 | hypothetical protein | 5 | 3 | 3 | 3 | 3 | 3 | 3 | 7 |
| 26 | TGME49_226985.P1233 | hypothetical protein | 7 | 3 | 6 | 4 | 3 | 3 | 3 | 7 |
| 9 | TGME49_227070-t26_1-p1 | hypothetical protein | 8 | 3 | 7 | 5 | 3 | 3 | 3 | 7 |
| 24 | TGME49_227100-t26_1-p1 | glutaredoxin 5 | 6 | 5 | 5 | 3 | 3 | 3 | 3 | 7 |
| 26 | TGME49_227280-t26_1-p1 | dense granule protein GRA3 | 6 | 4 | 3 | 3 | 3 | 3 | 3 | 7 |
| 26 | TGME49_227310.P1243 | hypothetical protein | 6 | 3 | 5 | 4 | 3 | 3 | 3 | 7 |
| 26 | TGME49_227330-t26_1-p1 | hypothetical protein | 6 | 4 | 3 | 3 | 3 | 3 | 3 | 7 |
| 9 | TGME49_227440.P1248 | WD domain, G-beta repeat-containing protein | 9 | 3 | 8 | 7 | 5 | 4 | 4 | 7 |
| 24 | TGME49_227590.P1252 | hypothetical protein | 7 | 6 | 5 | 4 | 3 | 3 | 3 | 7 |
| 16 | TGME49_227600.P1253 | ribosomal protein RPL34 | 6 | 6 | 6 | 6 | 6 | 3 | 3 | 7 |
| 26 | TGME49_227620-t26_1-p1 | dense granule protein GRA2 | 6 | 4 | 4 | 3 | 3 | 3 | 3 | 7 |
| 26 | TGME49_227630.P1255 | hypothetical protein | 5 | 3 | 3 | 3 | 3 | 3 | 3 | 7 |
| 16 | TGME49_227870.P1262 | Tim10/DDP family zinc finger superfamily protein | 5 | 5 | 5 | 5 | 4 | 3 | 3 | 7 |
| 16 | TGME49_227910.P1266 | cytochrome b-c1 complex subunit 8 | 6 | 6 | 6 | 5 | 5 | 5 | 3 | 7 |
| 16 | TGME49_227920-t26_1-p1 | succinate dehydrogenase subunit SDH10 | 5 | 5 | 4 | 4 | 4 | 3 | 3 | 7 |
| 16 | TGME49_227970.P1269 | histone family DNA-binding protein | 7 | 6 | 5 | 5 | 5 | 4 | 4 | 7 |
| 24 | TGME49_228010-t26_1-p1 | hypothetical protein | 8 | 5 | 5 | 4 | 3 | 3 | 3 | 7 |
| 24 | TGME49_228110.P1274 | transmembrane protein TMEM14C | 6 | 5 | 5 | 3 | 4 | 3 | 3 | 7 |
| 20 | TGME49_228175.P1280 | hypothetical protein | 9 | 3 | 5 | 4 | 3 | 3 | 3 | 7 |
| 20 | TGME49_228270-t26_1-p1 | hypothetical protein | 8 | 3 | 5 | 4 | 4 | 3 | 3 | 7 |
| 9 | TGME49_228280-t26_1-p1 | hypothetical protein | 8 | 3 | 5 | 4 | 4 | 4 | 4 | 7 |
| 16 | TGME49_228310.P1289 | hypothetical protein | 6 | 6 | 6 | 4 | 3 | 4 | 4 | 7 |
| 9 | TGME49_228390-t26_1-p1 | hypothetical protein | 10 | 3 | 9 | 7 | 3 | 3 | 4 | 7 |
| 24 | TGME49_228620-t26_1-p1 | phosphatidylethanolamine-binding protein, putative | 7 | 6 | 6 | 4 | 3 | 3 | 3 | 7 |
| 26 | TGME49_229220-t26_1-p1 | hypothetical protein | 7 | 3 | 3 | 4 | 4 | 3 | 3 | 7 |
| 16 | TGME49_229250-t26_1-p1 | ribosomal protein RPL28 | 6 | 6 | 6 | 5 | 5 | 4 | 4 | 7 |
| 16 | TGME49_229450-t26_1-p1 | hypothetical protein | 6 | 5 | 4 | 4 | 4 | 4 | 4 | 7 |
| 16 | TGME49_229680.P1333 | hypothetical protein | 6 | 6 | 6 | 3 | 5 | 3 | 5 | 7 |
| 16 | TGME49_230070-t26_1-p1 | BolA family protein | 5 | 5 | 5 | 5 | 4 | 3 | 4 | 7 |
| 24 | TGME49_230122.P1353 | hypothetical protein | 6 | 6 | 5 | 3 | 3 | 3 | 3 | 7 |
| 24 | TGME49_230130.P1354 | hypothetical protein | 6 | 5 | 4 | 4 | 3 | 3 | 3 | 7 |
| 26 | TGME49_230135.P1355 | Kinesin-associated protein | 6 | 3 | 5 | 5 | 3 | 3 | 3 | 7 |

| | | | | | | | | | | |
|---|---|---|---|---|---|---|---|---|---|---|
| 26 | TGME49_230160-t26_1-p1 | RNA-binding protein, putative | 6 | 5 | 3 | 4 | 3 | 3 | 3 | 7 |
| 26 | TGME49_230180.P1360 | dense granule protein GRA24 | 6 | 3 | 3 | 3 | 3 | 3 | 3 | 7 |
| 16 | TGME49_230380.P1366 | protein translocation complex, SEC61 gamma subunit | 5 | 5 | 5 | 3 | 5 | 5 | 5 | 7 |
| 24 | TGME49_230470.P1370 | rhoptry kinase family protein ROP46, putative | 7 | 7 | 3 | 3 | 3 | 3 | 3 | 7 |
| 16 | TGME49_230480.P1371 | hypothetical protein | 5 | 5 | 5 | 5 | 3 | 3 | 4 | 7 |
| 16 | TGME49_230610-t26_1-p1 | transmembrane protein TMEM222 | 7 | 5 | 6 | 3 | 5 | 3 | 3 | 7 |
| 24 | TGME49_230640-t26_1-p1 | hypothetical protein | 6 | 5 | 3 | 4 | 4 | 3 | 3 | 7 |
| 26 | TGME49_230705.P1383 | dense granule protein GRA51 | 7 | 3 | 3 | 3 | 3 | 3 | 3 | 7 |
| 26 | TGME49_230870-t26_1-p1 | hypothetical protein | 6 | 3 | 3 | 3 | 3 | 3 | 3 | 7 |
| 26 | TGME49_230970.P1393 | hypothetical protein | 5 | 3 | 4 | 3 | 3 | 3 | 3 | 7 |
| 24 | TGME49_231020-t26_1-p1 | hypothetical protein | 7 | 6 | 6 | 3 | 3 | 3 | 3 | 7 |
| 26 | TGME49_231050-t26_1-p1 | hypothetical protein | 6 | 3 | 3 | 3 | 3 | 3 | 3 | 7 |
| 26 | TGME49_231055-t26_1-p1 | hypothetical protein | 7 | 3 | 3 | 3 | 3 | 3 | 3 | 7 |
| 16 | TGME49_231080-t26_1-p1 | ribosomal protein RPL38 | 5 | 5 | 5 | 3 | 4 | 4 | 3 | 7 |
| 16 | TGME49_231110.P1402 | COPI associated protein | 6 | 6 | 5 | 3 | 5 | 3 | 3 | 7 |
| 24 | TGME49_231125.P1403 | hypothetical protein | 5 | 6 | 3 | 3 | 3 | 3 | 3 | 7 |
| 16 | TGME49_231140.P1405 | ribosomal protein RPS25 | 6 | 5 | 5 | 3 | 5 | 5 | 4 | 7 |
| 24 | TGME49_231160-t26_1-p1 | hypothetical protein | 6 | 6 | 5 | 4 | 4 | 3 | 3 | 7 |
| 20 | TGME49_231180-t26_1-p1 | hypothetical protein | 9 | 3 | 3 | 3 | 3 | 3 | 3 | 7 |
| 16 | TGME49_231380-t26_1-p1 | DNA-directed RNA polymerase II RPB4 | 6 | 6 | 6 | 3 | 5 | 5 | 4 | 7 |
| 20 | TGME49_231625-t26_1-p1 | hypothetical protein | 9 | 3 | 3 | 3 | 5 | 3 | 3 | 7 |
| 24 | TGME49_231860-t26_1-p1 | hypothetical protein | 8 | 5 | 4 | 3 | 3 | 3 | 3 | 7 |
| 26 | TGME49_231865-t26_1-p1 | hypothetical protein | 5 | 3 | 3 | 3 | 3 | 3 | 3 | 7 |
| 24 | TGME49_231880.P1428 | hypothetical protein | 8 | 6 | 3 | 4 | 3 | 3 | 3 | 7 |
| 24 | TGME49_231950-t26_1-p1 | hypothetical protein | 8 | 6 | 3 | 3 | 3 | 3 | 3 | 7 |
| 20 | TGME49_231960.P1433 | dense granule protein GRA28 | 9 | 3 | 3 | 3 | 4 | 3 | 3 | 7 |
| 26 | TGME49_231990-t26_1-p1 | hypothetical protein | 5 | 4 | 4 | 4 | 3 | 3 | 3 | 7 |
| 26 | TGME49_231991-t26_1-p1 | hypothetical protein | 5 | 3 | 3 | 3 | 3 | 3 | 3 | 7 |
| 26 | TGME49_231994-t26_1-p1 | hypothetical protein | 5 | 3 | 3 | 3 | 3 | 3 | 3 | 7 |
| 26 | TGME49_231996-t26_1-p1 | hypothetical protein | 5 | 3 | 3 | 3 | 3 | 3 | 3 | 7 |
| 26 | TGME49_231998.P1436 | hypothetical protein | 5 | 3 | 3 | 3 | 3 | 3 | 3 | 7 |
| 24 | TGME49_232030.P1440 | inner membrane complex protein IMC21 | 6 | 6 | 3 | 3 | 4 | 3 | 3 | 7 |
| 24 | TGME49_232200-t26_1-p1 | hypothetical protein | 6 | 6 | 5 | 3 | 3 | 3 | 3 | 7 |
| 16 | TGME49_232230-t26_1-p1 | ribosomal protein RPL30 | 6 | 6 | 6 | 5 | 5 | 3 | 3 | 7 |
| 20 | TGME49_232320.P1458 | hypothetical protein | 10 | 5 | 4 | 4 | 3 | 3 | 3 | 7 |
| 24 | TGME49_232400.P1464 | PAN domain-containing protein | 8 | 6 | 3 | 3 | 3 | 3 | 3 | 7 |
| 24 | TGME49_232630.P1477 | hypothetical protein | 6 | 6 | 5 | 4 | 4 | 3 | 3 | 7 |
| 9 | TGME49_232655-t26_1-p1 | hypothetical protein | 7 | 3 | 6 | 3 | 3 | 5 | 4 | 7 |
| 26 | TGME49_232700-t26_1-p1 | hypothetical protein | 7 | 3 | 6 | 3 | 3 | 4 | 3 | 7 |
| 16 | TGME49_232760-t26_1-p1 | protein phosphatase inhibitor IPP2 | 6 | 6 | 5 | 5 | 4 | 3 | 4 | 7 |
| 20 | TGME49_232950-t26_1-p1 | hypothetical protein | 8 | 5 | 3 | 3 | 3 | 3 | 3 | 7 |
| 24 | TGME49_232960.P1485 | prolyl hydroxylase PHYa | 8 | 7 | 5 | 3 | 4 | 3 | 3 | 7 |
| 9 | TGME49_232980.P1487 | hypothetical protein | 10 | 5 | 7 | 4 | 4 | 3 | 3 | 7 |
| 24 | TGME49_233030.P1490 | gliding-associated protein GAP70 | 6 | 5 | 4 | 4 | 3 | 3 | 3 | 7 |
| 16 | TGME49_233170.P1496 | hypothetical protein | 6 | 6 | 6 | 6 | 3 | 5 | 3 | 7 |
| 16 | TGME49_233200.P1499 | ubiquitin-like protein URM1 | 6 | 5 | 5 | 5 | 4 | 3 | 4 | 7 |
| 26 | TGME49_233290-t26_1-p1 | hypothetical protein | 7 | 3 | 3 | 3 | 3 | 3 | 3 | 7 |

| | | | | | | | | | | |
|---|---|---|---|---|---|---|---|---|---|---|
| 26 | TGME49_233315-t26_1-p1 | hypothetical protein | 5 | 3 | 3 | 3 | 3 | 3 | 3 | 7 |
| 16 | TGME49_233360-t26_1-p1 | hypothetical protein | 6 | 6 | 6 | 6 | 3 | 4 | 4 | 7 |
| 20 | TGME49_233405-t26_1-p1 | hypothetical protein | 8 | 3 | 3 | 3 | 3 | 4 | 3 | 7 |
| 20 | TGME49_233450-t26_1-p1 | SAG-related sequence SRS29A | 8 | 3 | 5 | 4 | 3 | 3 | 3 | 7 |
| 24 | TGME49_233460.P1508 | SAG-related sequence SRS29B | 8 | 7 | 3 | 4 | 3 | 3 | 3 | 7 |
| 24 | TGME49_233480-t26_1-p1 | SAG-related sequence SRS29C | 8 | 6 | 3 | 3 | 3 | 3 | 3 | 7 |
| 26 | TGME49_233610-t26_1-p1 | hypothetical protein | 7 | 3 | 3 | 3 | 3 | 3 | 3 | 7 |
| 26 | TGME49_233670-t26_1-p1 | hypothetical protein | 6 | 3 | 3 | 3 | 3 | 3 | 3 | 7 |
| 16 | TGME49_233695-t26_1-p1 | hypothetical protein | 6 | 4 | 5 | 4 | 4 | 3 | 3 | 7 |
| 24 | TGME49_233780-t26_1-p1 | hypothetical protein | 6 | 5 | 5 | 3 | 3 | 4 | 3 | 7 |
| 26 | TGME49_233792-t26_1-p1 | hypothetical protein | 6 | 3 | 4 | 3 | 3 | 3 | 3 | 7 |
| 26 | TGME49_234310.P1531 | hypothetical protein | 6 | 4 | 4 | 4 | 4 | 3 | 3 | 7 |
| 24 | TGME49_234400.P1534 | hypothetical protein | 7 | 6 | 6 | 4 | 3 | 3 | 3 | 7 |
| 9 | TGME49_234470.P1538 | hypothetical protein | 9 | 3 | 6 | 5 | 4 | 3 | 3 | 7 |
| 9 | TGME49_234507-t26_1-p1 | hypothetical protein | 8 | 3 | 7 | 3 | 3 | 3 | 3 | 7 |
| 9 | TGME49_234510.P1539 | acyl-CoA binding protein ACBP2 | 8 | 3 | 7 | 6 | 5 | 4 | 5 | 7 |
| 16 | TGME49_234530.P1540 | hypothetical protein | 5 | 3 | 7 | 3 | 4 | 4 | 4 | 7 |
| 9 | TGME49_234640.P1544 | hypothetical protein | 10 | 3 | 7 | 7 | 6 | 4 | 3 | 7 |
| 24 | TGME49_234930-t26_1-p1 | SAG-related sequence SRS43 | 8 | 7 | 3 | 5 | 3 | 3 | 3 | 7 |
| 9 | TGME49_234950.P1545 | protein kinase (incomplete catalytic triad) | 9 | 3 | 6 | 6 | 3 | 3 | 3 | 7 |
| 24 | TGME49_235010-t26_1-p1 | hypothetical protein | 7 | 5 | 4 | 3 | 3 | 3 | 3 | 7 |
| 26 | TGME49_235140.P1549 | secreted effector protein NSM | 7 | 3 | 3 | 3 | 3 | 3 | 3 | 7 |
| 24 | TGME49_235440.P1560 | LSM domain-containing protein | 6 | 5 | 4 | 3 | 3 | 3 | 3 | 7 |
| 26 | TGME49_235590-t26_1-p1 | hypothetical protein | 6 | 3 | 3 | 3 | 3 | 3 | 3 | 7 |
| 26 | TGME49_235630.P1567 | hypothetical protein | 5 | 3 | 3 | 3 | 3 | 3 | 3 | 7 |
| 16 | TGME49_235710-t26_1-p1 | hypothetical protein | 5 | 5 | 4 | 4 | 4 | 3 | 3 | 7 |
| 24 | TGME49_235720.P1572 | hypothetical protein | 7 | 5 | 4 | 4 | 4 | 3 | 4 | 7 |
| 24 | TGME49_235880.P1577 | mitochondrial pyruvate carrier protein 1 | 6 | 6 | 5 | 5 | 3 | 3 | 3 | 7 |
| 16 | TGME49_235890-t26_1-p1 | dual-specificity protein phosphatase | 6 | 5 | 5 | 3 | 4 | 4 | 3 | 7 |
| 20 | TGME49_236060-t26_1-p1 | hypothetical protein | 9 | 3 | 4 | 3 | 3 | 3 | 3 | 7 |
| 16 | TGME49_236130.P1588 | signal recognition particle (SRP9) domain-containing protein | 6 | 5 | 5 | 4 | 4 | 4 | 4 | 7 |
| 26 | TGME49_236190.P1590 | hypothetical protein | 7 | 3 | 3 | 4 | 3 | 3 | 3 | 7 |
| 26 | TGME49_236400-t26_1-p1 | hypothetical protein | 7 | 4 | 4 | 4 | 3 | 3 | 3 | 7 |
| 16 | TGME49_236550-t26_1-p1 | hypothetical protein | 6 | 6 | 5 | 5 | 5 | 3 | 3 | 7 |
| 24 | TGME49_236630.P1599 | hypothetical protein | 8 | 7 | 3 | 3 | 3 | 3 | 3 | 7 |
| 20 | TGME49_236670-t26_1-p1 | hypothetical protein | 9 | 3 | 4 | 3 | 3 | 3 | 3 | 7 |
| 24 | TGME49_236870.P1604 | dense granule protein GRA42 | 6 | 5 | 4 | 3 | 3 | 3 | 3 | 7 |
| 26 | TGME49_236890.P1605 | dense granule protein GRA37 | 6 | 4 | 3 | 3 | 3 | 3 | 3 | 7 |
| 24 | TGME49_236900-t26_1-p1 | hypothetical protein | 7 | 5 | 3 | 3 | 3 | 4 | 3 | 7 |
| 16 | TGME49_236920.P1607 | hypothetical protein | 6 | 5 | 5 | 4 | 4 | 3 | 4 | 7 |
| 16 | TGME49_236950-t26_1-p1 | hypothetical protein | 6 | 5 | 5 | 4 | 4 | 3 | 3 | 7 |
| 26 | TGME49_237015.P1614 | dense granule protein GRA43 | 7 | 3 | 4 | 3 | 3 | 3 | 3 | 7 |
| 24 | TGME49_237190.P1619 | hypothetical protein | 7 | 5 | 4 | 4 | 3 | 3 | 3 | 7 |
| 26 | TGME49_237230.P1623 | membrane protein MYR3 | 6 | 3 | 4 | 3 | 3 | 3 | 3 | 7 |
| 26 | TGME49_237810-t26_1-p1 | hypothetical protein | 5 | 3 | 3 | 3 | 3 | 3 | 3 | 7 |
| 26 | TGME49_237880.P1637 | dense granule protein GRA13 | 6 | 3 | 4 | 3 | 3 | 3 | 3 | 7 |

| | | | | | | | | | | |
|---|---|---|---|---|---|---|---|---|---|---|
| 24 | TGME49_238080-t26_1-p1 | zinc finger HIT domain-containing protein | 7 | 5 | 3 | 4 | 3 | 3 | 3 | 7 |
| 16 | TGME49_238110-t26_1-p1 | replication factor a protein 3 protein | 5 | 5 | 5 | 4 | 4 | 4 | 3 | 7 |
| 16 | TGME49_238250.P1646 | ribosomal protein RPL36 | 5 | 5 | 5 | 5 | 3 | 5 | 5 | 7 |
| 24 | TGME49_238260.P1647 | hypothetical protein | 8 | 7 | 4 | 3 | 3 | 3 | 3 | 7 |
| 26 | TGME49_238270.P1648 | hypothetical protein | 5 | 4 | 3 | 3 | 3 | 3 | 3 | 7 |
| 24 | TGME49_238410.P1650 | hypothetical protein | 7 | 7 | 5 | 4 | 3 | 3 | 3 | 7 |
| 20 | TGME49_238420-t26_1-p1 | hypothetical protein | 10 | 5 | 4 | 4 | 4 | 3 | 3 | 7 |
| 24 | TGME49_238440-t26_1-p1 | SAG-related sequence SRS22A | 6 | 5 | 4 | 4 | 3 | 3 | 3 | 7 |
| 26 | TGME49_238460.P1651 | SAG-related sequence SRS22B | 5 | 3 | 3 | 3 | 3 | 3 | 3 | 7 |
| 26 | TGME49_238470.P1652 | SAG-related sequence SRS22C | 6 | 3 | 4 | 3 | 3 | 3 | 3 | 7 |
| 26 | TGME49_238480-t26_1-p1 | SAG-related sequence SRS22D | 6 | 3 | 3 | 3 | 3 | 3 | 3 | 7 |
| 26 | TGME49_238490-t26_1-p1 | SAG-related sequence SRS22E | 5 | 3 | 3 | 3 | 3 | 3 | 3 | 7 |
| 26 | TGME49_238500.P1653 | SAG-related sequence SRS22F | 6 | 3 | 3 | 3 | 3 | 3 | 3 | 7 |
| 26 | TGME49_238520-t26_1-p1 | SAG-related sequence SRS22G | 5 | 3 | 4 | 3 | 3 | 3 | 3 | 7 |
| 26 | TGME49_238850.P1655 | SAG-related sequence SRS22I | 6 | 3 | 3 | 3 | 3 | 3 | 3 | 7 |
| 9 | TGME49_238960-t26_1-p1 | EF hand domain-containing protein | 8 | 3 | 6 | 3 | 3 | 4 | 4 | 7 |
| 20 | TGME49_238980.P1660 | hypothetical protein | 10 | 5 | 4 | 4 | 3 | 3 | 3 | 7 |
| 26 | TGME49_239010.P1662 | E2F4-associated EZH2-inducing gene regulator TEEGR | 7 | 3 | 3 | 3 | 3 | 3 | 3 | 7 |
| 24 | TGME49_239087-t26_1-p1 | hypothetical protein | 6 | 6 | 5 | 3 | 3 | 4 | 3 | 7 |
| 24 | TGME49_239090-t26_1-p1 | SAG-related sequence SRS23 | 8 | 6 | 4 | 3 | 3 | 3 | 3 | 7 |
| 16 | TGME49_239260-t26_1-p1 | histone H4 | 5 | 5 | 5 | 5 | 5 | 3 | 5 | 7 |
| 24 | TGME49_239320.P1670 | SufE-like protein 1 | 7 | 6 | 5 | 3 | 4 | 4 | 4 | 7 |
| 16 | TGME49_239330-t26_1-p1 | ribosomal protein RPL37 | 5 | 5 | 5 | 3 | 5 | 5 | 3 | 7 |
| 20 | TGME49_239365.P1672 | hypothetical protein | 8 | 3 | 3 | 3 | 3 | 3 | 3 | 7 |
| 16 | TGME49_239550-t26_1-p1 | dynein light chain 2B, cytoplasmic, putative | 6 | 6 | 6 | 3 | 3 | 5 | 5 | 7 |
| 26 | TGME49_239640-t26_1-p1 | hypothetical protein | 6 | 4 | 3 | 3 | 3 | 3 | 3 | 7 |
| 20 | TGME49_239650-t26_1-p1 | hypothetical protein | 8 | 5 | 3 | 3 | 3 | 3 | 3 | 7 |
| 20 | TGME49_239660-t26_1-p1 | hypothetical protein | 8 | 4 | 3 | 4 | 3 | 3 | 3 | 7 |
| 16 | TGME49_239690.P1681 | ER membrane protein complex subunit 6, putative | 6 | 5 | 5 | 3 | 4 | 4 | 4 | 7 |
| 24 | TGME49_239740-t26_1-p1 | dense granule protein GRA14 | 8 | 5 | 4 | 3 | 3 | 3 | 3 | 7 |
| 9 | TGME49_239748-t26_1-p1 | hypothetical protein | 9 | 3 | 6 | 5 | 4 | 3 | 4 | 7 |
| 16 | TGME49_239755-t26_1-p1 | hypothetical protein | 8 | 3 | 3 | 6 | 7 | 3 | 3 | 7 |
| 16 | TGME49_239770.P1685 | alveolin domain containing intermediate filament IMC11 | 5 | 6 | 5 | 5 | 3 | 3 | 3 | 7 |
| 26 | TGME49_239795.P1686 | hypothetical protein | 7 | 3 | 3 | 3 | 3 | 3 | 3 | 7 |
| 20 | TGME49_240060.P1692 | inhibitor of STAT1 transcription | 8 | 3 | 3 | 3 | 3 | 3 | 3 | 7 |
| 20 | TGME49_240200.P1696 | hypothetical protein | 8 | 3 | 3 | 5 | 3 | 3 | 3 | 7 |
| 24 | TGME49_240230-t26_1-p1 | hypothetical protein | 7 | 6 | 5 | 5 | 3 | 3 | 3 | 7 |
| 9 | TGME49_240243-t26_1-p1 | hypothetical protein | 8 | 3 | 7 | 6 | 4 | 3 | 3 | 7 |
| 26 | TGME49_240255-t26_1-p1 | hypothetical protein | 7 | 3 | 5 | 3 | 3 | 3 | 4 | 7 |
| 26 | TGME49_240270.P1699 | hypothetical protein | 5 | 5 | 5 | 3 | 3 | 3 | 3 | 7 |
| 20 | TGME49_240325-t26_1-p1 | Toxoplasma gondii family E protein | 9 | 3 | 3 | 3 | 3 | 3 | 3 | 7 |
| 20 | TGME49_240330-t26_1-p1 | Toxoplasma gondii family E protein | 9 | 3 | 3 | 3 | 3 | 3 | 3 | 7 |
| 20 | TGME49_240340-t26_1-p1 | Toxoplasma gondii family E protein | 9 | 3 | 3 | 3 | 3 | 3 | 3 | 7 |
| 24 | TGME49_240400-t26_1-p1 | hypothetical protein | 8 | 7 | 4 | 3 | 3 | 3 | 3 | 7 |
| 24 | TGME49_240440-t26_1-p1 | hypothetical protein | 8 | 6 | 5 | 3 | 3 | 3 | 3 | 7 |

| | | | | | | | | | | |
|---|---|---|---|---|---|---|---|---|---|---|
| 24 | TGME49_240480.P1704 | cpw-wpc domain-containing protein | 8 | 6 | 3 | 3 | 3 | 3 | 3 | 7 |
| 26 | TGME49_240490-t26_1-p1 | hypothetical protein | 6 | 4 | 3 | 3 | 3 | 3 | 3 | 7 |
| 20 | TGME49_240525-t26_1-p1 | hypothetical protein | 9 | 3 | 3 | 3 | 5 | 5 | 5 | 7 |
| 26 | TGME49_240550-t26_1-p1 | copper chaperone COX17-1, putative | 5 | 4 | 4 | 3 | 4 | 3 | 3 | 7 |
| 9 | TGME49_240575-t26_1-p1 | type iib dna topoisomerase | 8 | 3 | 7 | 4 | 3 | 5 | 4 | 7 |
| 24 | TGME49_240580.P1709 | hypothetical protein | 8 | 6 | 3 | 4 | 3 | 3 | 3 | 7 |
| 20 | TGME49_240680-t26_1-p1 | MOLO1 domain-containing protein | 8 | 3 | 3 | 3 | 4 | 6 | 3 | 7 |
| 16 | TGME49_240780-t26_1-p1 | hypothetical protein | 6 | 5 | 5 | 5 | 5 | 4 | 3 | 7 |
| 16 | TGME49_240790.P1715 | dynein light chain, putative | 5 | 5 | 5 | 5 | 3 | 3 | 3 | 7 |
| 24 | TGME49_240930.P1721 | MoaC family protein | 7 | 6 | 6 | 4 | 3 | 3 | 3 | 7 |
| 24 | TGME49_240950-t26_1-p1 | hypothetical protein | 8 | 6 | 5 | 3 | 4 | 4 | 4 | 7 |
| 26 | TGME49_241155-t26_1-p1 | hypothetical protein | 6 | 3 | 3 | 3 | 3 | 3 | 3 | 7 |
| 9 | TGME49_241165-t26_1-p1 | hypothetical protein | 8 | 3 | 7 | 4 | 3 | 4 | 3 | 7 |
| 26 | TGME49_241240.P1725 | dense granule protein GRA65 | 7 | 3 | 3 | 3 | 3 | 3 | 3 | 7 |
| 26 | TGME49_241305.P1726 | hypothetical protein | 6 | 3 | 3 | 3 | 3 | 3 | 3 | 7 |
| 26 | TGME49_241710-t26_1-p1 | hypothetical protein | 7 | 3 | 3 | 3 | 3 | 3 | 3 | 7 |
| 24 | TGME49_241890-t26_1-p1 | hypothetical protein | 6 | 6 | 5 | 5 | 3 | 3 | 3 | 7 |
| 20 | TGME49_242250-t26_1-p1 | Protein kinase domain-containing protein [Source:UniProtKB/TrEMBL;Acc:S8F3P2] | 8 | 3 | 3 | 3 | 3 | 3 | 3 | 7 |
| 26 | TGME49_242310-t26_1-p1 | hypothetical protein | 6 | 3 | 3 | 3 | 3 | 3 | 3 | 7 |
| 24 | TGME49_242650-t26_1-p1 | hypothetical protein | 8 | 7 | 4 | 3 | 3 | 3 | 3 | 7 |
| 26 | TGME49_242710.P1742 | hypothetical protein | 7 | 3 | 3 | 3 | 3 | 3 | 3 | 7 |
| 24 | TGME49_242740-t26_1-p1 | hypothetical protein | 7 | 7 | 6 | 3 | 3 | 3 | 3 | 7 |
| 26 | TGME49_242820.P1749 | hypothetical protein | 5 | 4 | 3 | 3 | 3 | 3 | 3 | 7 |
| 16 | TGME49_242840.P1750 | cytochrome c oxidase subunit ApiCOX14 | 6 | 6 | 5 | 5 | 4 | 3 | 3 | 7 |
| 26 | TGME49_242850.P1751 | hypothetical protein | 6 | 3 | 4 | 3 | 4 | 4 | 3 | 7 |
| 26 | TGME49_242900.P1753 | Toxoplasma gondii family A protein | 7 | 3 | 3 | 3 | 3 | 3 | 3 | 7 |
| 26 | TGME49_243100.P1754 | Toxoplasma gondii family A protein | 6 | 3 | 3 | 3 | 3 | 3 | 3 | 7 |
| 26 | TGME49_243110-t26_1-p1 | hypothetical protein | 7 | 3 | 3 | 3 | 3 | 3 | 3 | 7 |
| 24 | TGME49_243120-t26_1-p1 | Toxoplasma gondii family A protein | 8 | 6 | 3 | 4 | 3 | 3 | 3 | 7 |
| 24 | TGME49_243140-t26_1-p1 | Toxoplasma gondii family A protein | 8 | 6 | 3 | 3 | 3 | 3 | 3 | 7 |
| 26 | TGME49_243150-t26_1-p1 | Toxoplasma gondii family A protein | 6 | 3 | 5 | 3 | 3 | 3 | 3 | 7 |
| 24 | TGME49_243160-t26_1-p1 | Toxoplasma gondii family A protein | 8 | 6 | 3 | 3 | 3 | 3 | 3 | 7 |
| 20 | TGME49_243170-t26_1-p1 | Toxoplasma gondii family A protein | 8 | 3 | 3 | 3 | 3 | 3 | 3 | 7 |
| 20 | TGME49_243180-t26_1-p1 | Toxoplasma gondii family A protein | 8 | 3 | 4 | 3 | 3 | 3 | 3 | 7 |
| 24 | TGME49_243190-t26_1-p1 | Toxoplasma gondii family A protein | 8 | 6 | 3 | 3 | 3 | 3 | 3 | 7 |
| 24 | TGME49_243390-t26_1-p1 | ER membrane protein complex subunit 7, putative | 8 | 6 | 5 | 4 | 3 | 3 | 3 | 7 |
| 26 | TGME49_243545.P1763 | apical ring protein RNG1 | 5 | 3 | 4 | 4 | 3 | 3 | 3 | 7 |
| 9 | TGME49_243670.P1769 | ARM repeats containing protein | 9 | 4 | 6 | 5 | 3 | 4 | 3 | 7 |
| 26 | TGME49_243940.P1775 | dense granule protein GRA81 | 5 | 4 | 4 | 3 | 3 | 3 | 3 | 7 |
| 26 | TGME49_243980-t26_1-p1 | hypothetical protein | 6 | 4 | 4 | 4 | 4 | 3 | 3 | 7 |
| 24 | TGME49_244010-t26_1-p1 | HORMA domain-containing protein | 8 | 6 | 3 | 4 | 4 | 4 | 3 | 7 |
| 24 | TGME49_244020.P1778 | activator of hsp90 atpase 1 family protein | 6 | 6 | 3 | 3 | 5 | 3 | 3 | 7 |
| 26 | TGME49_244070-t26_1-p1 | hypothetical protein | 5 | 3 | 3 | 3 | 3 | 3 | 3 | 7 |
| 9 | TGME49_244130-t26_1-p1 | hypothetical protein | 10 | 5 | 8 | 4 | 4 | 4 | 4 | 7 |

| | | | | | | | | | | |
|---|---|---|---|---|---|---|---|---|---|---|
| 24 | TGME49_244160-t26_1-p1 | transcription initiation factor TFIID complex subunit TAF12 | 8 | 6 | 4 | 4 | 4 | 3 | 3 | 7 |
| 26 | TGME49_244170-t26_1-p1 | hypothetical protein | 6 | 3 | 3 | 3 | 3 | 3 | 3 | 7 |
| 20 | TGME49_244190-t26_1-p1 | hypothetical protein | 9 | 4 | 3 | 3 | 3 | 3 | 3 | 7 |
| 24 | TGME49_244260-t26_1-p1 | hypothetical protein | 7 | 6 | 4 | 4 | 3 | 3 | 3 | 7 |
| 26 | TGME49_244280-t26_1-p1 | hypothetical protein | 5 | 3 | 3 | 3 | 3 | 3 | 3 | 7 |
| 16 | TGME49_244480-t26_1-p1 | peptidase M16 inactive domain-containing protein | 3 | 3 | 3 | 8 | 6 | 3 | 3 | 7 |
| 20 | TGME49_244515-t26_1-p1 | hypothetical protein | 8 | 3 | 3 | 3 | 3 | 3 | 3 | 7 |
| 20 | TGME49_244530-t26_1-p1 | dense granule protein GRA49 | 9 | 4 | 3 | 3 | 3 | 3 | 3 | 7 |
| 24 | TGME49_244610.P1797 | zinc finger, C3HC4 type (RING finger) domain-containing protein | 8 | 6 | 5 | 4 | 3 | 3 | 3 | 7 |
| 20 | TGME49_244720.P1801 | hypothetical protein | 10 | 4 | 4 | 4 | 3 | 3 | 3 | 7 |
| 16 | TGME49_244900.P1803 | dynein light chain type 1, putative | 6 | 5 | 5 | 5 | 5 | 4 | 4 | 7 |
| 26 | TGME49_245428-t26_1-p1 | hypothetical protein | 6 | 3 | 3 | 3 | 3 | 3 | 3 | 7 |
| 26 | TGME49_245432-t26_1-p1 | dense granule protein GRA19 | 5 | 5 | 3 | 3 | 3 | 3 | 3 | 7 |
| 26 | TGME49_245475.P1806 | hypothetical protein | 6 | 5 | 3 | 3 | 3 | 3 | 3 | 7 |
| 20 | TGME49_245560.P1810 | hypothetical protein | 10 | 3 | 4 | 4 | 3 | 3 | 3 | 7 |
| 16 | TGME49_245630-t26_1-p1 | hypothetical protein | 5 | 5 | 5 | 5 | 4 | 4 | 3 | 7 |
| 16 | TGME49_245650-t26_1-p1 | proteasome assembly chaperone 4, putative | 7 | 5 | 5 | 5 | 4 | 3 | 3 | 7 |
| 26 | TGME49_245740-t26_1-p1 | hypothetical protein | 5 | 3 | 3 | 3 | 3 | 3 | 3 | 7 |
| 20 | TGME49_245980-t26_1-p1 | hypothetical protein | 9 | 5 | 3 | 3 | 3 | 3 | 3 | 7 |
| 16 | TGME49_246000.P1818 | large subunit ribosomal protein IMG2 | 7 | 5 | 5 | 5 | 5 | 5 | 4 | 7 |
| 24 | TGME49_246070-t26_1-p1 | SAG-related sequence SRS56 | 8 | 7 | 3 | 3 | 3 | 3 | 3 | 7 |
| 26 | TGME49_246535-t26_1-p1 | hypothetical protein | 5 | 3 | 3 | 3 | 3 | 3 | 3 | 7 |
| 26 | TGME49_246555.P1835 | hypothetical protein | 6 | 3 | 3 | 4 | 3 | 3 | 3 | 7 |
| 24 | TGME49_246610-t26_1-p1 | vesicle-associated membrane protein VAMP4 | 7 | 6 | 5 | 4 | 4 | 3 | 3 | 7 |
| 24 | TGME49_246940.P1842 | gliding-associated protein GAP80 | 8 | 6 | 4 | 5 | 4 | 3 | 3 | 7 |
| 9 | TGME49_246978-t26_1-p1 | hypothetical protein | 10 | 3 | 9 | 5 | 5 | 4 | 3 | 7 |
| 20 | TGME49_246990-t26_1-p1 | hypothetical protein | 9 | 5 | 4 | 4 | 3 | 3 | 3 | 7 |
| 9 | TGME49_246995-t26_1-p1 | hypothetical protein | 10 | 3 | 7 | 5 | 6 | 6 | 5 | 7 |
| 20 | TGME49_247015-t26_1-p1 | hypothetical protein | 9 | 3 | 4 | 4 | 3 | 3 | 3 | 7 |
| 9 | TGME49_247025-t26_1-p1 | PITH domain-containing protein | 8 | 3 | 7 | 6 | 3 | 3 | 3 | 7 |
| 24 | TGME49_247030-t26_1-p1 | hypothetical protein | 6 | 6 | 5 | 3 | 4 | 3 | 3 | 7 |
| 16 | TGME49_247340.P1848 | PIH1 domain-containing protein | 6 | 6 | 5 | 4 | 3 | 5 | 4 | 7 |
| 9 | TGME49_247390.P1853 | ATPase, AAA family protein | 10 | 3 | 9 | 5 | 6 | 3 | 3 | 7 |
| 24 | TGME49_247400-t26_1-p1 | hypothetical protein | 7 | 5 | 3 | 3 | 3 | 3 | 3 | 7 |
| 16 | TGME49_247410.P1854 | ATP synthase-associated protein | 6 | 6 | 5 | 5 | 5 | 3 | 4 | 7 |
| 26 | TGME49_247440.P1855 | dense granule protein GRA33 | 7 | 3 | 4 | 3 | 3 | 3 | 3 | 7 |
| 26 | TGME49_247530-t26_1-p1 | hypothetical protein | 6 | 3 | 3 | 3 | 3 | 3 | 3 | 7 |
| 24 | TGME49_247560.P1860 | dynein light chain protein, putative | 6 | 6 | 5 | 4 | 3 | 3 | 3 | 7 |
| 20 | TGME49_247570.P1861 | hypothetical protein | 9 | 5 | 5 | 3 | 3 | 4 | 4 | 7 |
| 16 | TGME49_247610-t26_1-p1 | U6 snRNA-associated Sm-like protein LSm5, putative | 6 | 6 | 5 | 3 | 5 | 3 | 4 | 7 |
| 20 | TGME49_247640-t26_1-p1 | hypothetical protein | 8 | 3 | 3 | 3 | 3 | 3 | 3 | 7 |
| 26 | TGME49_247648-t26_1-p1 | Sybindin family protein | 7 | 3 | 6 | 3 | 3 | 3 | 4 | 7 |
| 24 | TGME49_247960-t26_1-p1 | hypothetical protein | 8 | 6 | 3 | 5 | 3 | 3 | 3 | 7 |
| 26 | TGME49_247980-t26_1-p1 | hypothetical protein | 5 | 4 | 4 | 3 | 4 | 3 | 3 | 7 |

| | | | | | | | | | | |
|---|---|---|---|---|---|---|---|---|---|---|
| 20 | TGME49_248170.P1877 | peroxisomal protein PEX14, putative | 9 | 5 | 4 | 4 | 3 | 4 | 3 | 7 |
| 26 | TGME49_248180-t26_1-p1 | hypothetical protein | 5 | 4 | 3 | 3 | 3 | 3 | 3 | 7 |
| 16 | TGME49_248250-t26_1-p1 | translation initiation factor IF-2, putative | 6 | 6 | 6 | 3 | 6 | 6 | 3 | 7 |
| 24 | TGME49_248260-t26_1-p1 | hypothetical protein | 6 | 5 | 5 | 3 | 4 | 3 | 3 | 7 |
| 24 | TGME49_248330.P1882 | zinc finger, C3HC4 type (RING finger) domain-containing protein | 7 | 6 | 3 | 5 | 3 | 4 | 3 | 7 |
| 24 | TGME49_248360.P1884 | hypothetical protein | 8 | 7 | 3 | 4 | 3 | 3 | 3 | 7 |
| 26 | TGME49_248445-t26_1-p1 | hypothetical protein | 7 | 3 | 3 | 5 | 3 | 4 | 3 | 7 |
| 16 | TGME49_248490-t26_1-p1 | hypothetical protein | 6 | 6 | 5 | 3 | 5 | 4 | 3 | 7 |
| 16 | TGME49_248660.P1900 | cytochrome c oxidase assembly protein COX14, putative | 5 | 5 | 5 | 5 | 5 | 4 | 4 | 7 |
| 26 | TGME49_248673-t26_1-p1 | hypothetical protein | 5 | 3 | 3 | 5 | 3 | 3 | 3 | 7 |
| 26 | TGME49_248677-t26_1-p1 | nucleic acid binding protein, putative | 7 | 3 | 3 | 5 | 3 | 3 | 3 | 7 |
| 24 | TGME49_248820-t26_1-p1 | hypothetical protein | 8 | 7 | 4 | 3 | 3 | 3 | 3 | 7 |
| 16 | TGME49_248860-t26_1-p1 | hypothetical protein | 5 | 3 | 5 | 5 | 4 | 3 | 4 | 7 |
| 16 | TGME49_248940-t26_1-p1 | hypothetical protein | 5 | 5 | 5 | 3 | 5 | 3 | 4 | 7 |
| 16 | TGME49_249365.P1928 | intraflagellar transport protein 46, putative | 6 | 3 | 6 | 5 | 4 | 3 | 3 | 7 |
| 24 | TGME49_249370.P1929 | translation initiation factor SUI1, putative | 6 | 6 | 5 | 3 | 3 | 3 | 3 | 7 |
| 20 | TGME49_249425-t26_1-p1 | hypothetical protein | 8 | 3 | 3 | 3 | 3 | 3 | 3 | 7 |
| 26 | TGME49_249520-t26_1-p1 | hypothetical protein | 6 | 5 | 3 | 3 | 3 | 3 | 3 | 7 |
| 26 | TGME49_249570-t26_1-p1 | hypothetical protein | 6 | 4 | 3 | 3 | 3 | 3 | 3 | 7 |
| 26 | TGME49_249698.P1940 | hypothetical protein | 6 | 3 | 5 | 3 | 3 | 4 | 3 | 7 |
| 26 | TGME49_249750.P1943 | hypothetical protein | 6 | 5 | 3 | 3 | 3 | 3 | 3 | 7 |
| 24 | TGME49_249760-t26_1-p1 | vitamin k epoxide reductase family protein | 7 | 6 | 5 | 3 | 3 | 3 | 3 | 7 |
| 26 | TGME49_249930-t26_1-p1 | hypothetical protein | 7 | 3 | 3 | 4 | 3 | 3 | 3 | 7 |
| 26 | TGME49_249935.P1951 | hypothetical protein | 5 | 4 | 4 | 4 | 3 | 4 | 3 | 7 |
| 24 | TGME49_250030.P1956 | hypothetical protein | 7 | 5 | 4 | 4 | 3 | 3 | 3 | 7 |
| 26 | TGME49_250040.P1957 | hypothetical protein | 7 | 3 | 4 | 4 | 3 | 3 | 3 | 7 |
| 26 | TGME49_250115-t26_1-p1 | hypothetical protein | 7 | 3 | 3 | 3 | 3 | 3 | 3 | 7 |
| 16 | TGME49_250350.P1959 | CRIPT, putative | 5 | 5 | 5 | 3 | 3 | 4 | 3 | 7 |
| 26 | TGME49_250500-t26_1-p1 | hypothetical protein | 6 | 3 | 3 | 4 | 3 | 3 | 3 | 7 |
| 20 | TGME49_250670-t26_1-p1 | hypothetical protein | 8 | 3 | 3 | 3 | 3 | 3 | 3 | 7 |
| 24 | TGME49_250710.P1961 | microneme protein MIC10 | 6 | 5 | 4 | 3 | 4 | 3 | 3 | 7 |
| 16 | TGME49_250900.P1969 | hypothetical protein | 7 | 6 | 6 | 3 | 5 | 4 | 4 | 7 |
| 26 | TGME49_250950-t26_1-p1 | KRUF family protein | 7 | 3 | 3 | 3 | 3 | 3 | 3 | 7 |
| 16 | TGME49_251390.P1973 | rieske [2fe-2s] domain-containing protein | 6 | 6 | 5 | 5 | 4 | 3 | 3 | 7 |
| 16 | TGME49_251470.P1974 | V-type ATPase V0 subunit e, putative | 5 | 5 | 5 | 5 | 4 | 4 | 3 | 7 |
| 16 | TGME49_251550.P1977 | acyl-CoA binding protein ACBP1 | 5 | 5 | 5 | 5 | 4 | 3 | 4 | 7 |
| 26 | TGME49_251660-t26_1-p1 | hypothetical protein | 5 | 3 | 3 | 3 | 3 | 3 | 3 | 7 |
| 24 | TGME49_251800-t26_1-p1 | hypothetical protein | 8 | 5 | 3 | 4 | 4 | 3 | 3 | 7 |
| 26 | TGME49_251857.P1984 | hypothetical protein | 7 | 3 | 5 | 4 | 3 | 4 | 4 | 7 |
| 16 | TGME49_251870-t26_1-p1 | histone H2Bb | 6 | 3 | 6 | 3 | 3 | 3 | 5 | 7 |
| 9 | TGME49_251885-t26_1-p1 | hypothetical protein | 7 | 3 | 6 | 5 | 5 | 4 | 4 | 7 |
| 26 | TGME49_251900-t26_1-p1 | hypothetical protein | 6 | 4 | 4 | 4 | 3 | 3 | 3 | 7 |
| 24 | TGME49_251910-t26_1-p1 | hypothetical protein | 7 | 6 | 6 | 3 | 3 | 3 | 3 | 7 |

| | | | | | | | | | | |
|---|---|---|---|---|---|---|---|---|---|---|
| 20 | TGME49_251962-t26_1-p1 | SAG-related sequence SRS59K | 9 | 3 | 4 | 3 | 3 | 3 | 3 | 7 |
| 20 | TGME49_252070-t26_1-p1 | KRUF family protein | 10 | 4 | 3 | 3 | 3 | 3 | 3 | 7 |
| 20 | TGME49_252190-t26_1-p1 | KRUF family protein | 8 | 3 | 3 | 3 | 3 | 3 | 3 | 7 |
| 20 | TGME49_252255.P1988 | hypothetical protein | 8 | 3 | 3 | 3 | 3 | 3 | 3 | 7 |
| 20 | TGME49_252280-t26_1-p1 | hypothetical protein | 10 | 5 | 4 | 4 | 3 | 3 | 3 | 7 |
| 16 | TGME49_252310.P1992 | hypothetical protein | 7 | 4 | 5 | 4 | 4 | 4 | 3 | 7 |
| 9 | TGME49_252330-t26_1-p1 | hypothetical protein | 9 | 3 | 6 | 6 | 3 | 3 | 3 | 7 |
| 20 | TGME49_252350-t26_1-p1 | hypothetical protein | 10 | 4 | 3 | 3 | 3 | 3 | 3 | 7 |
| 26 | TGME49_252390.P1996 | hypothetical protein | 7 | 4 | 3 | 3 | 3 | 3 | 3 | 7 |
| 20 | TGME49_252395-t26_1-p1 | hypothetical protein | 10 | 3 | 4 | 4 | 3 | 3 | 3 | 7 |
| 9 | TGME49_252480-t26_1-p1 | hypothetical protein | 7 | 3 | 6 | 5 | 4 | 3 | 4 | 7 |
| 16 | TGME49_252630-t26_1-p1 | succinate dehydrogenase subunit SDH15 | 6 | 3 | 5 | 5 | 5 | 4 | 4 | 7 |
| 26 | TGME49_252860-t26_1-p1 | hypothetical protein | 7 | 3 | 4 | 3 | 3 | 3 | 3 | 7 |
| 9 | TGME49_252880-t26_1-p1 | hypothetical protein | 9 | 3 | 6 | 4 | 4 | 4 | 3 | 7 |
| 26 | TGME49_252890-t26_1-p1 | hypothetical protein | 5 | 3 | 3 | 3 | 3 | 3 | 3 | 7 |
| 24 | TGME49_252990-t26_1-p1 | hypothetical protein | 8 | 6 | 5 | 4 | 4 | 3 | 3 | 7 |
| 24 | TGME49_253460-t26_1-p1 | hypothetical protein | 7 | 5 | 3 | 3 | 3 | 3 | 3 | 7 |
| 16 | TGME49_253530-t26_1-p1 | ribonuclease P/MRP protein subunit POP5, putative | 6 | 6 | 5 | 5 | 4 | 4 | 4 | 7 |
| 26 | TGME49_253660.P2033 | RING finger protein, putative | 5 | 5 | 3 | 5 | 3 | 3 | 3 | 7 |
| 26 | TGME49_253690-t26_1-p1 | hypothetical protein | 5 | 4 | 4 | 4 | 4 | 3 | 3 | 7 |
| 26 | TGME49_253755-t26_1-p1 | hypothetical protein | 6 | 3 | 3 | 3 | 3 | 3 | 3 | 7 |
| 26 | TGME49_253770-t26_1-p1 | hypothetical protein | 5 | 3 | 3 | 3 | 3 | 3 | 3 | 7 |
| 24 | TGME49_253960.P2039 | oxidoreductase, short chain dehydrogenase/reductase family protein | 7 | 5 | 6 | 3 | 3 | 4 | 4 | 7 |
| 20 | TGME49_253970.P2040 | hypothetical protein | 9 | 5 | 3 | 3 | 3 | 3 | 3 | 7 |
| 16 | TGME49_254030.P2044 | cytochrome c oxidase subunit ApiCOX13 | 6 | 6 | 6 | 5 | 4 | 4 | 3 | 7 |
| 26 | TGME49_254040-t26_1-p1 | hypothetical protein | 5 | 3 | 3 | 3 | 3 | 3 | 3 | 7 |
| 24 | TGME49_254060-t26_1-p1 | SAG-related sequence SRS14A | 8 | 5 | 4 | 3 | 3 | 3 | 3 | 7 |
| 26 | TGME49_254140.P2047 | DNA-directed RNA polymerase II RPABC4 | 5 | 4 | 4 | 3 | 4 | 3 | 3 | 7 |
| 9 | TGME49_254160.P2048 | hypothetical protein | 10 | 5 | 6 | 5 | 3 | 4 | 4 | 7 |
| 16 | TGME49_254340.P2054 | SAYSvFN domain-containing protein | 6 | 6 | 5 | 5 | 4 | 4 | 3 | 7 |
| 16 | TGME49_254460.P2058 | hypothetical protein | 6 | 5 | 5 | 4 | 3 | 3 | 4 | 7 |
| 16 | TGME49_254520-t26_1-p1 | mediator complex subunit MED11 | 6 | 6 | 5 | 5 | 4 | 4 | 3 | 7 |
| 16 | TGME49_254590-t26_1-p1 | hypothetical protein | 6 | 6 | 5 | 5 | 5 | 3 | 3 | 7 |
| 16 | TGME49_254600-t26_1-p1 | ubiquitin family protein | 5 | 5 | 5 | 3 | 3 | 3 | 5 | 7 |
| 20 | TGME49_254603-t26_1-p1 | hypothetical protein | 8 | 3 | 5 | 3 | 3 | 3 | 3 | 7 |
| 16 | TGME49_254610-t26_1-p1 | Tim10/DDP family zinc finger superfamily protein | 5 | 5 | 5 | 5 | 5 | 3 | 3 | 7 |
| 26 | TGME49_254625.P2063 | hypothetical protein | 7 | 3 | 5 | 4 | 3 | 3 | 3 | 7 |
| 26 | TGME49_254635-t26_1-p1 | hypothetical protein | 7 | 3 | 3 | 3 | 3 | 3 | 3 | 7 |
| 24 | TGME49_254640.P2065 | DNA polymerase delta subunit 4, putative | 7 | 6 | 3 | 4 | 4 | 3 | 4 | 7 |
| 9 | TGME49_254660.P2067 | ankyrin repeat-containing protein | 8 | 3 | 7 | 3 | 3 | 5 | 4 | 7 |
| 26 | TGME49_254670-t26_1-p1 | hypothetical protein | 6 | 3 | 4 | 3 | 3 | 3 | 3 | 7 |
| 26 | TGME49_254680-t26_1-p1 | hypothetical protein | 6 | 5 | 3 | 4 | 3 | 3 | 3 | 7 |

| | | | | | | | | | | |
|---|---|---|---|---|---|---|---|---|---|---|
| 26 | TGME49_254720.P2070 | dense granule protein GRA8 | 6 | 4 | 3 | 3 | 3 | 3 | 3 | 7 |
| 26 | TGME49_254740-t26_1-p1 | endoribonuclease L-PSP protein | 5 | 3 | 3 | 4 | 3 | 3 | 3 | 7 |
| 24 | TGME49_254780.P2074 | hypothetical protein | 6 | 6 | 4 | 5 | 3 | 3 | 3 | 7 |
| 16 | TGME49_254800.P2075 | LYR domain-containing protein | 6 | 5 | 4 | 4 | 4 | 4 | 3 | 7 |
| 24 | TGME49_254820-t26_1-p1 | hypothetical protein | 8 | 6 | 4 | 3 | 3 | 3 | 3 | 7 |
| 9 | TGME49_255200.P2089 | Radial spoke head protein 9, putative | 8 | 5 | 7 | 7 | 3 | 5 | 4 | 7 |
| 26 | TGME49_255245.P2092 | mitochondrial import receptor subunit TOM22 | 5 | 4 | 4 | 3 | 4 | 3 | 3 | 7 |
| 20 | TGME49_255330-t26_1-p1 | hypothetical protein | 8 | 4 | 3 | 5 | 4 | 4 | 3 | 7 |
| 24 | TGME49_255360-t26_1-p1 | hypothetical protein | 7 | 7 | 3 | 4 | 3 | 4 | 3 | 7 |
| 24 | TGME49_255380-t26_1-p1 | hypothetical protein | 7 | 5 | 3 | 3 | 3 | 3 | 3 | 7 |
| 9 | TGME49_255400.P2098 | hypothetical protein | 8 | 3 | 5 | 6 | 5 | 3 | 3 | 7 |
| 24 | TGME49_255490.P2106 | hypothetical protein | 7 | 7 | 5 | 4 | 3 | 3 | 3 | 7 |
| 16 | TGME49_255660.P2111 | EF hand domain-containing protein EFP1 | 6 | 5 | 5 | 4 | 4 | 3 | 3 | 7 |
| 9 | TGME49_255700.P2115 | hypothetical protein | 10 | 3 | 7 | 5 | 4 | 3 | 3 | 7 |
| 26 | TGME49_255715-t26_1-p1 | hypothetical protein | 7 | 3 | 5 | 3 | 3 | 3 | 3 | 7 |
| 20 | TGME49_255720.P2116 | tRNA intron endonuclease, putative | 9 | 3 | 5 | 4 | 4 | 3 | 4 | 7 |
| 16 | TGME49_255910.P2122 | PfMNL-2 CISD1 family iron-sulfur protein, putative | 6 | 5 | 5 | 5 | 4 | 4 | 4 | 7 |
| 20 | TGME49_255940-t26_1-p1 | hypothetical protein | 8 | 3 | 4 | 3 | 4 | 4 | 4 | 7 |
| 9 | TGME49_256010.P2127 | hypothetical protein | 7 | 3 | 6 | 5 | 5 | 4 | 4 | 7 |
| 16 | TGME49_256025.P2129 | SPEF1 family protein | 6 | 3 | 6 | 3 | 5 | 5 | 4 | 7 |
| 26 | TGME49_256035-t26_1-p1 | hypothetical protein | 6 | 3 | 3 | 3 | 3 | 3 | 3 | 7 |
| 16 | TGME49_256050-t26_1-p1 | signal recognition particle 14kd protein | 6 | 6 | 6 | 5 | 3 | 3 | 3 | 7 |
| 9 | TGME49_256788.P2137 | intraflagellar transport 52 (Protein NGD5) family protein | 9 | 3 | 8 | 8 | 7 | 4 | 5 | 7 |
| 26 | TGME49_256792-t26_1-p1 | hypothetical protein | 7 | 3 | 5 | 3 | 3 | 3 | 3 | 7 |
| 24 | TGME49_257160-t26_1-p1 | hypothetical protein | 6 | 5 | 5 | 4 | 3 | 3 | 3 | 7 |
| 20 | TGME49_257330.P2166 | hypothetical protein | 10 | 4 | 4 | 3 | 3 | 3 | 3 | 7 |
| 16 | TGME49_257380.P2170 | inhibitor of cysteine protease 1 | 6 | 6 | 5 | 5 | 4 | 3 | 3 | 7 |
| 24 | TGME49_257420.P2172 | hypothetical protein | 7 | 5 | 5 | 3 | 3 | 3 | 3 | 7 |
| 24 | TGME49_257460-t26_1-p1 | hypothetical protein | 7 | 6 | 3 | 3 | 3 | 3 | 3 | 7 |
| 20 | TGME49_257572-t26_1-p1 | hypothetical protein | 8 | 4 | 3 | 3 | 3 | 3 | 3 | 7 |
| 24 | TGME49_257580-t26_1-p1 | hypothetical protein | 8 | 6 | 4 | 3 | 4 | 3 | 3 | 7 |
| 20 | TGME49_257925.P2192 | hypothetical protein | 9 | 3 | 3 | 3 | 3 | 3 | 3 | 7 |
| 24 | TGME49_257970-t26_1-p1 | hypothetical protein | 8 | 6 | 3 | 3 | 3 | 3 | 3 | 7 |
| 9 | TGME49_258000.P2197 | hypothetical protein | 10 | 5 | 8 | 6 | 4 | 3 | 3 | 7 |
| 16 | TGME49_258100.P2206 | succinate dehydrogenase assembly factor 2, putative | 6 | 5 | 5 | 4 | 4 | 4 | 4 | 7 |
| 9 | TGME49_258220-t26_1-p1 | hypothetical protein | 7 | 3 | 6 | 6 | 5 | 3 | 4 | 7 |
| 16 | TGME49_258720-t26_1-p1 | Ubiquitin family protein, putative | 5 | 5 | 5 | 5 | 5 | 5 | 3 | 7 |
| 24 | TGME49_258730-t26_1-p1 | hypothetical protein | 6 | 6 | 5 | 5 | 3 | 3 | 3 | 7 |
| 24 | TGME49_258810-t26_1-p1 | SAG-related sequence SRS27B | 8 | 6 | 3 | 4 | 3 | 3 | 3 | 7 |
| 20 | TGME49_258840-t26_1-p1 | hypothetical protein | 9 | 4 | 3 | 3 | 3 | 3 | 3 | 7 |
| 26 | TGME49_258890-t26_1-p1 | hypothetical protein | 6 | 5 | 3 | 3 | 3 | 3 | 3 | 7 |
| 24 | TGME49_258930-t26_1-p1 | peptidylprolyl isomerase | 6 | 6 | 5 | 5 | 3 | 3 | 3 | 7 |
| 26 | TGME49_258940-t26_1-p1 | acylphosphatase family protein | 6 | 3 | 5 | 3 | 3 | 4 | 3 | 7 |
| 24 | TGME49_259040.P2257 | hypothetical protein | 7 | 5 | 4 | 3 | 3 | 3 | 3 | 7 |

| | | | | | | | | | | |
|---|---|---|---|---|---|---|---|---|---|---|
| 24 | TGME49_259070.P2259 | acetyltransferase, GNAT family protein | 8 | 6 | 5 | 4 | 3 | 3 | 3 | 7 |
| 9 | TGME49_259080.P2260 | hypothetical protein | 10 | 3 | 4 | 5 | 4 | 4 | 4 | 7 |
| 20 | TGME49_259150-t26_1-p1 | hypothetical protein | 10 | 3 | 3 | 3 | 3 | 3 | 3 | 7 |
| 20 | TGME49_259205.P2271 | hypothetical protein | 8 | 3 | 4 | 3 | 3 | 3 | 3 | 7 |
| 16 | TGME49_259240-t26_1-p1 | ribosomal protein RPS21 | 5 | 5 | 5 | 3 | 5 | 5 | 5 | 7 |
| 26 | TGME49_259290-t26_1-p1 | SAG-related sequence SRS26C | 7 | 3 | 4 | 3 | 3 | 3 | 3 | 7 |
| 9 | TGME49_259580-t26_1-p1 | hypothetical protein | 7 | 3 | 6 | 5 | 4 | 3 | 4 | 7 |
| 26 | TGME49_259590-t26_1-p1 | membrane protein, putative | 6 | 3 | 5 | 3 | 3 | 3 | 3 | 7 |
| 26 | TGME49_259620.P2282 | hypothetical protein | 5 | 5 | 3 | 3 | 3 | 3 | 3 | 7 |
| 26 | TGME49_259895-t26_1-p1 | hypothetical protein | 5 | 3 | 3 | 3 | 3 | 3 | 3 | 7 |
| 24 | TGME49_260020.P2303 | hypothetical protein | 6 | 5 | 4 | 4 | 3 | 3 | 3 | 7 |
| 9 | TGME49_260030.P2304 | atypical MEK-related kinase (incomplete catalytic triad) | 10 | 3 | 6 | 4 | 4 | 4 | 4 | 7 |
| 16 | TGME49_260260.P2313 | ribosomal protein RPP1 | 6 | 5 | 5 | 4 | 4 | 4 | 4 | 7 |
| 26 | TGME49_260325-t26_1-p1 | hypothetical protein | 7 | 3 | 3 | 3 | 3 | 3 | 3 | 7 |
| 16 | TGME49_260340.P2315 | DNL zinc finger protein | 6 | 3 | 5 | 5 | 4 | 4 | 4 | 7 |
| 16 | TGME49_260370.P2317 | PH-containing protein PH1 | 6 | 6 | 6 | 3 | 6 | 5 | 3 | 7 |
| 26 | TGME49_260375-t26_1-p1 | hypothetical protein | 5 | 3 | 3 | 3 | 3 | 3 | 3 | 7 |
| 24 | TGME49_260400-t26_1-p1 | hypothetical protein | 7 | 6 | 4 | 4 | 3 | 3 | 3 | 7 |
| 16 | TGME49_260620-t26_1-p1 | hypothetical protein | 5 | 4 | 5 | 4 | 4 | 3 | 3 | 7 |
| 26 | TGME49_260825.P2343 | hypothetical protein | 5 | 3 | 4 | 4 | 3 | 3 | 3 | 7 |
| 16 | TGME49_260870-t26_1-p1 | zinc finger cdgsh type protein | 7 | 5 | 5 | 4 | 4 | 4 | 4 | 7 |
| 16 | TGME49_260990.P2345 | hypothetical protein | 6 | 6 | 6 | 4 | 5 | 3 | 4 | 7 |
| 24 | TGME49_261040.P2349 | hypothetical protein | 7 | 6 | 3 | 3 | 3 | 3 | 3 | 7 |
| 9 | TGME49_261450.P2363 | hypothetical protein | 10 | 3 | 8 | 6 | 6 | 5 | 6 | 7 |
| 16 | TGME49_261470.P2364 | U6 snRNA-associated Sm family protein LSm6, putative | 5 | 5 | 5 | 4 | 5 | 4 | 3 | 7 |
| 16 | TGME49_261540-t26_1-p1 | DNA-directed RNA polymerase I RPAC2 | 6 | 5 | 5 | 5 | 5 | 4 | 4 | 7 |
| 16 | TGME49_261580-t26_1-p1 | histone H2AX | 6 | 6 | 6 | 6 | 6 | 3 | 3 | 7 |
| 24 | TGME49_261700-t26_1-p1 | hypothetical protein | 7 | 5 | 3 | 3 | 3 | 3 | 3 | 7 |
| 24 | TGME49_261970-t26_1-p1 | hypothetical protein | 8 | 5 | 5 | 3 | 3 | 3 | 3 | 7 |
| 26 | TGME49_261990.P2389 | zinc finger, C3HC4 type (RING finger) domain-containing protein | 5 | 4 | 4 | 3 | 3 | 3 | 3 | 7 |
| 9 | TGME49_262050-t26_1-p1 | rhoptry kinase family protein ROP39 | 8 | 3 | 6 | 4 | 3 | 3 | 3 | 7 |
| 24 | TGME49_262160.P2398 | ubiquitin-like protein, putative | 7 | 6 | 5 | 5 | 3 | 3 | 3 | 7 |
| 24 | TGME49_262560-t26_1-p1 | hypothetical protein | 6 | 5 | 4 | 3 | 4 | 3 | 3 | 7 |
| 20 | TGME49_262740-t26_1-p1 | RING zinc finger protein | 9 | 3 | 5 | 3 | 4 | 4 | 4 | 7 |
| 16 | TGME49_262810-t26_1-p1 | iron donor protein CyaY protein | 7 | 4 | 3 | 5 | 5 | 4 | 4 | 7 |
| 9 | TGME49_262890-t26_1-p1 | hypothetical protein | 9 | 3 | 7 | 3 | 3 | 5 | 3 | 7 |
| 26 | TGME49_262930.P2433 | dolichol-phosphate mannosyltransferase subunit 3, putative | 6 | 5 | 3 | 4 | 3 | 3 | 3 | 7 |
| 9 | TGME49_262933-t26_1-p1 | hypothetical protein | 10 | 3 | 7 | 6 | 4 | 3 | 3 | 7 |
| 20 | TGME49_262935.P2434 | hypothetical protein | 9 | 3 | 3 | 4 | 4 | 3 | 4 | 7 |
| 16 | TGME49_263030-t26_1-p1 | dynein light chain 8 family E protein | 5 | 5 | 5 | 5 | 5 | 3 | 3 | 7 |
| 16 | TGME49_263080.P2442 | ATP synthase-associated protein | 6 | 6 | 5 | 5 | 4 | 3 | 4 | 7 |
| 9 | TGME49_263085.P2443 | zinc finger protein, putative | 10 | 3 | 7 | 5 | 4 | 3 | 3 | 7 |
| 20 | TGME49_263210.P2454 | ubiquitin interaction motif domain-containing protein | 10 | 4 | 5 | 4 | 4 | 3 | 3 | 7 |

| | | | | | | | | | | |
|---|---|---|---|---|---|---|---|---|---|---|
| 9 | TGME49_263245-t26_1-p1 | cpw-wpc domain-containing protein | 7 | 3 | 6 | 5 | 4 | 3 | 3 | 7 |
| 16 | TGME49_263250-t26_1-p1 | Pb-fam-5 protein | 6 | 3 | 5 | 5 | 3 | 5 | 3 | 7 |
| 16 | TGME49_263530.P2474 | chaperonin, putative | 5 | 5 | 5 | 5 | 5 | 4 | 4 | 7 |
| 16 | TGME49_263580.P2479 | bromodomain-containing protein BDP1 | 3 | 3 | 3 | 7 | 7 | 3 | 6 | 7 |
| 26 | TGME49_263590.P2480 | hypothetical protein | 7 | 4 | 4 | 3 | 3 | 3 | 3 | 7 |
| 16 | TGME49_263630.P2483 | hypothetical protein | 5 | 5 | 4 | 4 | 3 | 4 | 3 | 7 |
| 24 | TGME49_263660.P2485 | hypothetical protein | 8 | 6 | 5 | 4 | 4 | 3 | 3 | 7 |
| 16 | TGME49_263720.P2488 | high mobility group protein B1, putative | 5 | 5 | 5 | 3 | 5 | 3 | 3 | 7 |
| 9 | TGME49_263785.P2493 | phosphatidate cytidylyltransferase CDS2 | 9 | 5 | 7 | 5 | 3 | 4 | 4 | 7 |
| 16 | TGME49_263810-t26_1-p1 | hypothetical protein | 5 | 5 | 5 | 3 | 4 | 4 | 3 | 7 |
| 16 | TGME49_263990.P2500 | hypothetical protein | 6 | 6 | 5 | 5 | 4 | 3 | 3 | 7 |
| 26 | TGME49_264010-t26_1-p1 | U4/U6.U5 small nuclear ribonucleoprotein, putative | 7 | 4 | 4 | 3 | 4 | 3 | 3 | 7 |
| 26 | TGME49_264075-t26_1-p1 | hypothetical protein | 5 | 3 | 3 | 3 | 3 | 3 | 3 | 7 |
| 24 | TGME49_264110.P2505 | hypothetical protein | 8 | 5 | 3 | 5 | 3 | 3 | 4 | 7 |
| 26 | TGME49_264225-t26_1-p1 | Toxoplasma gondii family B protein | 5 | 3 | 3 | 3 | 3 | 3 | 3 | 7 |
| 24 | TGME49_264472.P2516 | hypothetical protein | 8 | 5 | 5 | 4 | 4 | 3 | 3 | 7 |
| 26 | TGME49_264620-t26_1-p1 | hypothetical protein | 6 | 4 | 4 | 4 | 4 | 3 | 3 | 7 |
| 24 | TGME49_264630-t26_1-p1 | hypothetical protein | 6 | 5 | 5 | 3 | 3 | 3 | 3 | 7 |
| 26 | TGME49_264700.P2521 | EF hand domain-containing protein | 5 | 3 | 4 | 3 | 3 | 3 | 3 | 7 |
| 16 | TGME49_264730-t26_1-p1 | hypothetical protein | 6 | 6 | 6 | 3 | 6 | 3 | 5 | 7 |
| 9 | TGME49_264900-t26_1-p1 | hypothetical protein | 9 | 4 | 6 | 5 | 5 | 4 | 4 | 7 |
| 9 | TGME49_264950.P2527 | tubulin binding cofactor c, putative | 9 | 5 | 7 | 4 | 4 | 5 | 4 | 7 |
| 24 | TGME49_264960.P2528 | spindle and kinetochore-associated protein SKA1 | 7 | 5 | 4 | 3 | 3 | 3 | 3 | 7 |
| 26 | TGME49_265060-t26_1-p1 | hypothetical protein | 5 | 5 | 3 | 3 | 3 | 3 | 3 | 7 |
| 26 | TGME49_265065-t26_1-p1 | hypothetical protein | 5 | 3 | 3 | 3 | 3 | 3 | 3 | 7 |
| 20 | TGME49_265270-t26_1-p1 | hypothetical protein | 8 | 5 | 3 | 3 | 3 | 3 | 3 | 7 |
| 24 | TGME49_265290-t26_1-p1 | hypothetical protein | 8 | 7 | 4 | 3 | 3 | 3 | 3 | 7 |
| 16 | TGME49_265370.P2537 | cytochrome c oxidase subunit ApiCOX16 | 6 | 6 | 5 | 5 | 5 | 4 | 3 | 7 |
| 26 | TGME49_265538-t26_1-p1 | hypothetical protein | 6 | 3 | 3 | 3 | 3 | 3 | 3 | 7 |
| 24 | TGME49_265990-t26_1-p1 | hypothetical protein | 7 | 6 | 5 | 4 | 4 | 4 | 3 | 7 |
| 26 | TGME49_266050-t26_1-p1 | hypothetical protein | 7 | 3 | 4 | 3 | 3 | 3 | 3 | 7 |
| 26 | TGME49_266065.P2555 | hypothetical protein | 7 | 3 | 4 | 3 | 3 | 3 | 3 | 7 |
| 16 | TGME49_266070.P2556 | ribosomal protein RPL31 | 5 | 6 | 6 | 3 | 5 | 5 | 5 | 7 |
| 9 | TGME49_266105.P2560 | EF hand family protein | 8 | 3 | 7 | 5 | 3 | 4 | 3 | 7 |
| 9 | TGME49_266130-t26_1-p1 | peroxiredoxin PRX2 | 8 | 3 | 7 | 6 | 6 | 3 | 3 | 7 |
| 20 | TGME49_266150-t26_1-p1 | hypothetical protein | 8 | 3 | 3 | 3 | 3 | 3 | 3 | 7 |
| 16 | TGME49_266300.P2561 | hypothetical protein | 7 | 5 | 5 | 4 | 4 | 4 | 4 | 7 |
| 20 | TGME49_266330-t26_1-p1 | Toxoplasma gondii family A protein | 8 | 3 | 3 | 3 | 3 | 3 | 3 | 7 |
| 24 | TGME49_266340-t26_1-p1 | Toxoplasma gondii family A protein | 6 | 6 | 5 | 4 | 3 | 3 | 3 | 7 |
| 20 | TGME49_266360.P2563 | hypothetical protein | 8 | 5 | 3 | 3 | 3 | 3 | 3 | 7 |
| 20 | TGME49_266380.P2565 | hypothetical protein | 9 | 4 | 4 | 3 | 4 | 3 | 3 | 7 |
| 16 | TGME49_266460.P2569 | small ubiquitin family modifier, putative | 5 | 5 | 5 | 5 | 5 | 5 | 4 | 7 |

| | | | | | | | | | | |
|---|---|---|---|---|---|---|---|---|---|---|
| 20 | TGME49_266610.P2571 | Kazal-type serine protease inhibitor domain-containing protein | 9 | 3 | 3 | 3 | 3 | 3 | 3 | 7 |
| 24 | TGME49_266720-t26_1-p1 | hypothetical protein | 6 | 6 | 3 | 3 | 3 | 3 | 3 | 7 |
| 16 | TGME49_266840-t26_1-p1 | hypothetical protein | 6 | 7 | 5 | 5 | 5 | 4 | 3 | 7 |
| 9 | TGME49_266870-t26_1-p1 | transporter, major facilitator family protein | 9 | 3 | 8 | 3 | 3 | 3 | 3 | 7 |
| 9 | TGME49_267045.P2594 | Ras family protein | 7 | 3 | 6 | 5 | 3 | 3 | 3 | 7 |
| 24 | TGME49_267120.P2597 | hypothetical protein | 6 | 5 | 4 | 4 | 3 | 4 | 3 | 7 |
| 20 | TGME49_267130-t26_1-p1 | SAG-related sequence SRS38A | 8 | 3 | 4 | 5 | 3 | 3 | 3 | 7 |
| 9 | TGME49_267160-t26_1-p1 | SAG-related sequence SRS38D | 8 | 3 | 6 | 5 | 3 | 3 | 3 | 7 |
| 16 | TGME49_267340-t26_1-p1 | hypothetical protein | 6 | 5 | 5 | 3 | 5 | 4 | 4 | 7 |
| 24 | TGME49_267350-t26_1-p1 | LSM domain-containing protein | 6 | 6 | 5 | 3 | 3 | 3 | 5 | 7 |
| 16 | TGME49_267400-t26_1-p1 | ribosomal protein RPL32 | 6 | 6 | 6 | 6 | 6 | 3 | 3 | 7 |
| 16 | TGME49_267470.P2602 | cold-shock DNA-binding domain-containing protein | 6 | 6 | 5 | 3 | 5 | 3 | 4 | 7 |
| 16 | TGME49_267520-t26_1-p1 | anaphase promoting complex subunit 11, putative | 6 | 5 | 5 | 5 | 3 | 3 | 4 | 7 |
| 20 | TGME49_267642-t26_1-p1 | hypothetical protein | 8 | 3 | 3 | 3 | 3 | 3 | 3 | 7 |
| 9 | TGME49_267710-t26_1-p1 | CPSF A subunit region protein | 10 | 3 | 9 | 5 | 6 | 6 | 4 | 7 |
| 20 | TGME49_267725-t26_1-p1 | hypothetical protein | 8 | 3 | 5 | 3 | 3 | 3 | 3 | 7 |
| 20 | TGME49_267740.P2614 | dense granule protein GRA48 | 8 | 4 | 3 | 3 | 3 | 3 | 3 | 7 |
| 9 | TGME49_267750.P2615 | hypothetical protein | 10 | 3 | 7 | 4 | 4 | 4 | 4 | 7 |
| 9 | TGME49_267790.P2616 | hypothetical protein | 10 | 3 | 6 | 6 | 7 | 5 | 4 | 7 |
| 20 | TGME49_267980-t26_1-p1 | hypothetical protein | 8 | 4 | 4 | 3 | 3 | 3 | 3 | 7 |
| 20 | TGME49_268176-t26_1-p1 | hypothetical protein | 9 | 3 | 3 | 3 | 3 | 3 | 3 | 7 |
| 24 | TGME49_268190.P2624 | hypothetical protein | 8 | 6 | 5 | 3 | 4 | 3 | 3 | 7 |
| 20 | TGME49_268220-t26_1-p1 | hypothetical protein | 10 | 4 | 4 | 3 | 3 | 3 | 3 | 7 |
| 24 | TGME49_268270.P2629 | hypothetical protein | 7 | 6 | 5 | 4 | 3 | 3 | 3 | 7 |
| 20 | TGME49_268360-t26_1-p1 | hypothetical protein | 9 | 3 | 5 | 3 | 3 | 3 | 3 | 7 |
| 24 | TGME49_268390-t26_1-p1 | hypothetical protein | 8 | 6 | 5 | 4 | 3 | 3 | 3 | 7 |
| 26 | TGME49_268610-t26_1-p1 | hypothetical protein | 7 | 4 | 3 | 4 | 3 | 3 | 3 | 7 |
| 16 | TGME49_268660-t26_1-p1 | hypothetical protein | 5 | 5 | 4 | 4 | 4 | 3 | 3 | 7 |
| 26 | TGME49_268670-t26_1-p1 | hypothetical protein | 7 | 3 | 3 | 3 | 3 | 4 | 3 | 7 |
| 16 | TGME49_268685.P2643 | mitochondrial large subunit ribosomal protein | 7 | 3 | 5 | 4 | 4 | 4 | 3 | 7 |
| 24 | TGME49_268700-t26_1-p1 | hypothetical protein | 8 | 6 | 4 | 3 | 3 | 3 | 3 | 7 |
| 16 | TGME49_268730-t26_1-p1 | glutaredoxin 1 | 7 | 5 | 5 | 3 | 5 | 4 | 3 | 7 |
| 24 | TGME49_268760-t26_1-p1 | hypothetical protein | 8 | 6 | 5 | 4 | 3 | 3 | 3 | 7 |
| 26 | TGME49_268790.P2648 | dense granule protein GRA58 | 7 | 4 | 3 | 3 | 3 | 3 | 3 | 7 |
| 24 | TGME49_268800-t26_1-p1 | hypothetical protein | 6 | 6 | 3 | 3 | 3 | 3 | 3 | 7 |
| 24 | TGME49_268980-t26_1-p1 | hypothetical protein | 7 | 5 | 3 | 4 | 4 | 3 | 3 | 7 |
| 26 | TGME49_268985-t26_1-p1 | hypothetical protein | 5 | 3 | 5 | 3 | 3 | 3 | 3 | 7 |
| 20 | TGME49_269170-t26_1-p1 | hypothetical protein | 8 | 3 | 4 | 3 | 3 | 4 | 3 | 7 |
| 26 | TGME49_269280-t26_1-p1 | hypothetical protein | 6 | 5 | 3 | 3 | 3 | 3 | 3 | 7 |
| 20 | TGME49_269315.P2679 | hypothetical protein | 8 | 3 | 3 | 4 | 3 | 3 | 3 | 7 |
| 20 | TGME49_269630.P2690 | hypothetical protein | 9 | 5 | 4 | 4 | 3 | 3 | 3 | 7 |
| 9 | TGME49_269710.P2697 | hypothetical protein | 10 | 3 | 9 | 7 | 5 | 3 | 3 | 7 |
| 26 | TGME49_269970-t26_1-p1 | hypothetical protein | 5 | 4 | 4 | 3 | 3 | 3 | 3 | 7 |
| 24 | TGME49_270120.P2720 | thioredoxin-like protein TLP1 | 6 | 6 | 5 | 4 | 4 | 3 | 3 | 7 |
| 26 | TGME49_270170-t26_1-p1 | hypothetical protein | 5 | 4 | 4 | 3 | 3 | 3 | 3 | 7 |

| | | | | | | | | | | |
|---|---|---|---|---|---|---|---|---|---|---|
| 24 | TGME49_270250-t26_1-p1 | dense granule protein GRA1 | 6 | 5 | 5 | 4 | 3 | 3 | 3 | 7 |
| 20 | TGME49_270280-t26_1-p1 | hypothetical protein | 10 | 4 | 4 | 3 | 3 | 3 | 3 | 7 |
| 16 | TGME49_270310.P2731 | acyl carrier protein, mitochondrial | 7 | 5 | 5 | 4 | 4 | 4 | 4 | 7 |
| 24 | TGME49_270350-t26_1-p1 | hypothetical protein | 8 | 6 | 5 | 4 | 3 | 3 | 4 | 7 |
| 16 | TGME49_270360.P2735 | ATP synthase-associated protein | 6 | 6 | 6 | 3 | 5 | 3 | 3 | 7 |
| 26 | TGME49_270520.P2746 | hypothetical protein | 6 | 3 | 3 | 3 | 3 | 3 | 3 | 7 |
| 20 | TGME49_270630.P2775 | hypothetical protein | 9 | 5 | 3 | 4 | 4 | 3 | 4 | 7 |
| 24 | TGME49_270660.P2776 | hypothetical protein | 8 | 7 | 4 | 3 | 3 | 3 | 3 | 7 |
| 24 | TGME49_270700.P2778 | membrane protein MYR2 | 8 | 5 | 4 | 3 | 3 | 3 | 3 | 7 |
| 26 | TGME49_270805-t26_1-p1 | hypothetical protein | 6 | 3 | 3 | 3 | 3 | 3 | 3 | 7 |
| 20 | TGME49_270975.P2794 | hypothetical protein | 10 | 4 | 5 | 4 | 3 | 3 | 3 | 7 |
| 20 | TGME49_271010.P2796 | hypothetical protein | 10 | 3 | 4 | 4 | 3 | 3 | 3 | 7 |
| 26 | TGME49_271015-t26_1-p1 | hypothetical protein | 7 | 3 | 3 | 3 | 3 | 3 | 3 | 7 |
| 24 | TGME49_271040-t26_1-p1 | hypothetical protein | 7 | 6 | 3 | 3 | 3 | 3 | 3 | 7 |
| 24 | TGME49_271050-t26_1-p1 | SAG-related sequence SRS34A | 6 | 5 | 4 | 3 | 3 | 3 | 3 | 7 |
| 20 | TGME49_271080-t26_1-p1 | hypothetical protein | 9 | 3 | 3 | 3 | 3 | 3 | 3 | 7 |
| 26 | TGME49_271174-t26_1-p1 | hypothetical protein | 7 | 3 | 5 | 3 | 3 | 4 | 3 | 7 |
| 9 | TGME49_271285-t26_1-p1 | leucine rich repeat-containing protein | 9 | 3 | 8 | 6 | 6 | 5 | 5 | 7 |
| 26 | TGME49_271320-t26_1-p1 | hypothetical protein | 7 | 3 | 3 | 3 | 3 | 3 | 3 | 7 |
| 9 | TGME49_271335.P2809 | hypothetical protein | 8 | 3 | 6 | 6 | 5 | 4 | 4 | 7 |
| 24 | TGME49_271360.P2811 | hypothetical protein | 8 | 6 | 4 | 4 | 3 | 4 | 3 | 7 |
| 24 | TGME49_271390-t26_1-p1 | U3 small nucleolar RNA-associated protein 11, putative | 8 | 7 | 4 | 3 | 5 | 3 | 4 | 7 |
| 24 | TGME49_271580-t26_1-p1 | Toxoplasma gondii family D protein | 6 | 7 | 3 | 3 | 3 | 3 | 3 | 7 |
| 26 | TGME49_271790-t26_1-p1 | hypothetical protein | 5 | 4 | 4 | 4 | 3 | 3 | 3 | 7 |
| 26 | TGME49_271850-t26_1-p1 | hypothetical protein | 6 | 3 | 3 | 3 | 3 | 4 | 4 | 7 |
| 9 | TGME49_271892.P2828 | hypothetical protein | 9 | 3 | 8 | 6 | 3 | 5 | 5 | 7 |
| 26 | TGME49_271935.P2832 | hypothetical protein | 5 | 3 | 3 | 3 | 3 | 3 | 3 | 7 |
| 24 | TGME49_272180.P2840 | 3-oxo-5-alpha-steroid 4-dehydrogenase | 8 | 5 | 5 | 3 | 4 | 4 | 4 | 7 |
| 20 | TGME49_272190.P2841 | hypothetical protein | 8 | 3 | 5 | 4 | 4 | 4 | 3 | 7 |
| 26 | TGME49_272265.P2846 | hypothetical protein | 7 | 3 | 5 | 5 | 3 | 3 | 3 | 7 |
| 20 | TGME49_272285.P2849 | tetratricopeptide repeat-containing protein | 10 | 3 | 5 | 3 | 3 | 3 | 3 | 7 |
| 26 | TGME49_272415.P2861 | hypothetical protein | 7 | 3 | 3 | 3 | 3 | 3 | 3 | 7 |
| 16 | TGME49_272440.P2864 | RNA recognition motif-containing protein | 5 | 4 | 6 | 5 | 5 | 3 | 4 | 7 |
| 16 | TGME49_272630.P2875 | U6 snRNA-associated Sm family protein LSm8, putative | 5 | 5 | 5 | 4 | 5 | 4 | 4 | 7 |
| 24 | TGME49_272660.P2878 | hypothetical protein | 7 | 6 | 5 | 4 | 3 | 3 | 3 | 7 |
| 24 | TGME49_272680-t26_1-p1 | hypothetical protein | 7 | 5 | 3 | 3 | 3 | 3 | 3 | 7 |
| 26 | TGME49_272905.P2888 | hypothetical protein | 7 | 3 | 4 | 3 | 3 | 3 | 3 | 7 |
| 26 | TGME49_273065-t26_1-p1 | hypothetical protein | 6 | 3 | 5 | 3 | 3 | 3 | 3 | 7 |
| 20 | TGME49_273120-t26_1-p1 | SAG-related sequence SRS30C | 8 | 3 | 3 | 5 | 3 | 3 | 3 | 7 |
| 26 | TGME49_273130.P2892 | SAG-related sequence SRS30A | 6 | 3 | 5 | 5 | 3 | 3 | 3 | 7 |
| 24 | TGME49_273320-t26_1-p1 | hypothetical protein | 7 | 5 | 5 | 3 | 3 | 3 | 4 | 7 |
| 26 | TGME49_273340-t26_1-p1 | hypothetical protein | 5 | 5 | 3 | 3 | 3 | 3 | 3 | 7 |
| 24 | TGME49_273350-t26_1-p1 | molybdopterin converting factor, subunit 2 protein | 6 | 6 | 5 | 4 | 4 | 3 | 3 | 7 |
| 16 | TGME49_273400.P2902 | hypothetical protein | 6 | 6 | 6 | 3 | 5 | 3 | 3 | 7 |

| | | | | | | | | | | |
|---|---|---|---|---|---|---|---|---|---|---|
| 24 | TGME49_273610.P2912 | hypothetical protein | 6 | 6 | 5 | 3 | 3 | 3 | 3 | 7 |
| 24 | TGME49_273720-t26_1-p1 | hypothetical protein | 8 | 5 | 3 | 5 | 3 | 3 | 3 | 7 |
| 24 | TGME49_273750-t26_1-p1 | hypothetical protein | 8 | 7 | 3 | 5 | 3 | 3 | 3 | 7 |
| 16 | TGME49_273830-t26_1-p1 | hypothetical protein | 6 | 6 | 6 | 3 | 5 | 3 | 4 | 7 |
| 24 | TGME49_273860-t26_1-p1 | hypothetical protein | 8 | 6 | 4 | 3 | 4 | 3 | 3 | 7 |
| 26 | TGME49_273980-t26_1-p1 | dense granule protein GRA80 | 5 | 4 | 3 | 3 | 3 | 3 | 3 | 7 |
| 26 | TGME49_273995.P2930 | hypothetical protein | 7 | 3 | 3 | 4 | 4 | 3 | 3 | 7 |
| 26 | TGME49_274090.P2937 | TIM10 family protein, putative | 5 | 5 | 4 | 3 | 3 | 4 | 3 | 7 |
| 20 | TGME49_274100.P2938 | hypothetical protein | 9 | 5 | 3 | 5 | 3 | 3 | 3 | 7 |
| 9 | TGME49_275330-t26_1-p1 | ribosomal protein RPL29 | 10 | 3 | 5 | 5 | 5 | 4 | 3 | 7 |
| 20 | TGME49_275380.P2951 | SAG-related sequence SRS47D | 8 | 5 | 3 | 3 | 3 | 3 | 3 | 7 |
| 26 | TGME49_275440-t26_1-p1 | dense granule protein GRA6 | 6 | 4 | 3 | 3 | 3 | 3 | 3 | 7 |
| 24 | TGME49_275460-t26_1-p1 | hypothetical protein | 8 | 6 | 4 | 3 | 3 | 3 | 3 | 7 |
| 20 | TGME49_275470-t26_1-p1 | dense granule protein GRA15 | 9 | 3 | 3 | 3 | 3 | 3 | 3 | 7 |
| 9 | TGME49_275600-t26_1-p1 | diacylglycerol acyltransferase | 9 | 3 | 8 | 7 | 7 | 3 | 3 | 7 |
| 24 | TGME49_275640.P2956 | hypothetical protein | 8 | 5 | 4 | 4 | 3 | 4 | 3 | 7 |
| 24 | TGME49_275700-t26_1-p1 | hypothetical protein | 7 | 5 | 5 | 3 | 4 | 3 | 3 | 7 |
| 16 | TGME49_275710.P2960 | hypothetical protein | 6 | 7 | 5 | 5 | 3 | 4 | 4 | 7 |
| 24 | TGME49_275760-t26_1-p1 | hypothetical protein | 8 | 5 | 5 | 4 | 3 | 3 | 3 | 7 |
| 9 | TGME49_275850-t26_1-p1 | dense granule protein GRA12C | 9 | 3 | 7 | 6 | 4 | 4 | 5 | 7 |
| 20 | TGME49_275990-t26_1-p1 | hypothetical protein | 10 | 3 | 4 | 3 | 3 | 3 | 3 | 7 |
| 16 | TGME49_276110.P2963 | cytochrome b5 family heme/steroid binding domain-containing protein | 6 | 5 | 5 | 5 | 4 | 4 | 4 | 7 |
| 20 | TGME49_276220-t26_1-p1 | hypothetical protein | 10 | 3 | 3 | 3 | 3 | 3 | 3 | 7 |
| 24 | TGME49_276840.P2970 | hypothetical protein | 6 | 5 | 3 | 5 | 3 | 3 | 4 | 7 |
| 16 | TGME49_276850-t26_1-p1 | LEA domain-containing protein LEA850 | 5 | 5 | 5 | 3 | 4 | 3 | 3 | 7 |
| 24 | TGME49_276870-t26_1-p1 | LEA domain-containing protein LEA870 | 6 | 5 | 5 | 4 | 3 | 3 | 3 | 7 |
| 26 | TGME49_276880.P2972 | LEA domain-containing protein LEA880 | 5 | 5 | 5 | 3 | 3 | 3 | 3 | 7 |
| 26 | TGME49_276940-t26_1-p1 | ribosome associated membrane protein RAMP4, putative | 5 | 4 | 4 | 3 | 3 | 4 | 3 | 7 |
| 20 | TGME49_276980-t26_1-p1 | hypothetical protein | 8 | 3 | 3 | 3 | 3 | 3 | 3 | 7 |
| 16 | TGME49_277010.P2975 | SufE-like protein 2 | 6 | 6 | 5 | 4 | 4 | 3 | 4 | 7 |
| 26 | TGME49_277080-t26_1-p1 | microneme protein MIC5 | 6 | 4 | 4 | 3 | 3 | 3 | 3 | 7 |
| 26 | TGME49_277230.P2979 | dense granule protein GRA82 | 6 | 4 | 3 | 3 | 3 | 3 | 3 | 7 |
| 20 | TGME49_277540-t26_1-p1 | hypothetical protein | 10 | 4 | 4 | 4 | 4 | 3 | 3 | 7 |
| 16 | TGME49_277880-t26_1-p1 | hypothetical protein | 5 | 5 | 5 | 4 | 3 | 3 | 4 | 7 |
| 16 | TGME49_277980-t26_1-p1 | dynein light chain | 5 | 5 | 5 | 5 | 5 | 3 | 4 | 7 |
| 24 | TGME49_278040-t26_1-p1 | G-patch domain-containing protein | 7 | 6 | 5 | 3 | 5 | 3 | 4 | 7 |
| 20 | TGME49_278090-t26_1-p1 | Toxoplasma gondii family A protein | 9 | 3 | 3 | 4 | 3 | 3 | 3 | 7 |
| 24 | TGME49_278100-t26_1-p1 | Toxoplasma gondii family A protein | 6 | 6 | 3 | 4 | 3 | 3 | 3 | 7 |
| 26 | TGME49_278245-t26_1-p1 | hypothetical protein | 5 | 3 | 3 | 3 | 3 | 3 | 3 | 7 |
| 20 | TGME49_278290-t26_1-p1 | Toxoplasma gondii family A protein | 8 | 3 | 3 | 3 | 3 | 3 | 3 | 7 |
| 24 | TGME49_278330-t26_1-p1 | Toxoplasma gondii family A protein | 8 | 6 | 3 | 3 | 3 | 3 | 3 | 7 |
| 20 | TGME49_278340-t26_1-p1 | Toxoplasma gondii family A protein | 9 | 3 | 4 | 3 | 3 | 3 | 3 | 7 |
| 20 | TGME49_278350-t26_1-p1 | Toxoplasma gondii family A protein | 9 | 3 | 3 | 3 | 3 | 3 | 3 | 7 |
| 20 | TGME49_278360-t26_1-p1 | Toxoplasma gondii family A protein | 9 | 3 | 4 | 3 | 3 | 3 | 3 | 7 |
| 26 | TGME49_278365-t26_1-p1 | Toxoplasma gondii family A protein | 6 | 3 | 3 | 3 | 3 | 3 | 3 | 7 |
| 24 | TGME49_278380-t26_1-p1 | Toxoplasma gondii family A protein | 6 | 6 | 5 | 3 | 3 | 3 | 3 | 7 |
| 24 | TGME49_278390-t26_1-p1 | Toxoplasma gondii family A protein | 7 | 6 | 5 | 3 | 3 | 3 | 3 | 7 |
| 24 | TGME49_278410-t26_1-p1 | Toxoplasma gondii family A protein | 8 | 6 | 5 | 3 | 3 | 3 | 3 | 7 |

| | | | | | | | | | | |
|---|---|---|---|---|---|---|---|---|---|---|
| 24 | TGME49_278420-t26_1-p1 | Toxoplasma gondii family A protein | 8 | 6 | 3 | 3 | 3 | 3 | 3 | 7 |
| 20 | TGME49_278430-t26_1-p1 | Toxoplasma gondii family A protein | 8 | 3 | 3 | 3 | 3 | 3 | 3 | 7 |
| 26 | TGME49_278460.P2999 | hypothetical protein | 6 | 3 | 5 | 3 | 3 | 3 | 3 | 7 |
| 26 | TGME49_278522.P3002 | hypothetical protein | 5 | 3 | 3 | 3 | 3 | 4 | 3 | 7 |
| 16 | TGME49_278540.P3003 | hypothetical protein | 6 | 5 | 4 | 4 | 4 | 4 | 3 | 7 |
| 24 | TGME49_278565-t26_1-p1 | hypothetical protein | 8 | 6 | 5 | 3 | 3 | 3 | 3 | 7 |
| 26 | TGME49_278620-t26_1-p1 | hypothetical protein | 7 | 4 | 3 | 3 | 3 | 3 | 3 | 7 |
| 24 | TGME49_278690-t26_1-p1 | cpw-wpc domain-containing protein | 8 | 6 | 3 | 3 | 3 | 4 | 3 | 7 |
| 20 | TGME49_278750.P3012 | AT hook motif domain-containing protein | 8 | 3 | 4 | 3 | 3 | 3 | 3 | 7 |
| 26 | TGME49_278753-t26_1-p1 | hypothetical protein | 5 | 3 | 3 | 3 | 3 | 3 | 3 | 7 |
| 16 | TGME49_278790-t26_1-p1 | Tctex2-related light chain | 6 | 3 | 5 | 5 | 3 | 3 | 4 | 7 |
| 20 | TGME49_278815.P3014 | hypothetical protein | 10 | 5 | 4 | 5 | 4 | 3 | 3 | 7 |
| 24 | TGME49_278860.P3021 | thioredoxin reductase-like selenoprotein, putative | 6 | 6 | 6 | 3 | 3 | 3 | 4 | 7 |
| 16 | TGME49_278950.P3024 | LSM domain-containing protein | 6 | 5 | 5 | 3 | 5 | 5 | 4 | 7 |
| 26 | TGME49_279100.P3028 | mitochondrial association factor 1a | 7 | 3 | 3 | 3 | 4 | 3 | 3 | 7 |
| 24 | TGME49_279340.P3030 | VMA21-like domain containing protein | 6 | 5 | 4 | 4 | 4 | 3 | 3 | 7 |
| 24 | TGME49_279350-t26_1-p1 | hypothetical protein | 6 | 6 | 5 | 3 | 3 | 3 | 3 | 7 |
| 16 | TGME49_279400.P3032 | glutaredoxin 2 | 6 | 5 | 5 | 4 | 5 | 4 | 4 | 7 |
| 24 | TGME49_279410-t26_1-p1 | hypothetical protein | 8 | 6 | 3 | 4 | 3 | 3 | 3 | 7 |
| 16 | TGME49_279430-t26_1-p1 | cwf18 pre-mRNA splicing factor protein | 6 | 6 | 5 | 5 | 5 | 3 | 5 | 7 |
| 26 | TGME49_279460-t26_1-p1 | Toxoplasma gondii family B protein | 5 | 3 | 3 | 3 | 3 | 3 | 3 | 7 |
| 26 | TGME49_279540-t26_1-p1 | hypothetical protein | 5 | 3 | 3 | 3 | 3 | 3 | 3 | 7 |
| 26 | TGME49_279550-t26_1-p1 | hypothetical protein | 6 | 3 | 3 | 3 | 3 | 3 | 3 | 7 |
| 26 | TGME49_280070-t26_1-p1 | hypothetical protein | 5 | 3 | 3 | 3 | 3 | 3 | 3 | 7 |
| 26 | TGME49_280375-t26_1-p1 | hypothetical protein | 5 | 3 | 3 | 3 | 3 | 3 | 3 | 7 |
| 9 | TGME49_280430.P3037 | ML domain-containing protein | 7 | 3 | 6 | 6 | 3 | 4 | 3 | 7 |
| 16 | TGME49_280510-t26_1-p1 | Dpy-30 motif protein | 6 | 6 | 5 | 5 | 4 | 3 | 4 | 7 |
| 20 | TGME49_280518-t26_1-p1 | hypothetical protein | 9 | 3 | 4 | 3 | 3 | 3 | 3 | 7 |
| 24 | TGME49_280570-t26_1-p1 | SAG-related sequence SRS35A | 6 | 6 | 3 | 3 | 3 | 3 | 3 | 7 |
| 24 | TGME49_280580.P3046 | SAG-related sequence SRS35B | 7 | 6 | 3 | 3 | 3 | 3 | 3 | 7 |
| 20 | TGME49_280605-t26_1-p1 | hypothetical protein | 10 | 3 | 5 | 3 | 3 | 3 | 3 | 7 |
| 26 | TGME49_280680-t26_1-p1 | hypothetical protein | 7 | 3 | 3 | 3 | 3 | 3 | 3 | 7 |
| 20 | TGME49_280725.P3052 | zinc finger, C2H2 type domain-containing protein | 8 | 3 | 4 | 4 | 3 | 4 | 3 | 7 |
| 16 | TGME49_280750.P3054 | rudimentary enhancer, putative | 5 | 5 | 5 | 3 | 5 | 5 | 5 | 7 |
| 26 | TGME49_280810-t26_1-p1 | hypothetical protein | 7 | 4 | 3 | 3 | 3 | 3 | 3 | 7 |
| 26 | TGME49_281350-t26_1-p1 | hypothetical protein | 7 | 3 | 3 | 3 | 3 | 3 | 3 | 7 |
| 26 | TGME49_281360-t26_1-p1 | hypothetical protein | 5 | 4 | 4 | 3 | 3 | 3 | 3 | 7 |
| 26 | TGME49_281460-t26_1-p1 | hypothetical protein | 6 | 4 | 4 | 3 | 3 | 3 | 3 | 7 |
| 26 | TGME49_281530-t26_1-p1 | hypothetical protein | 5 | 3 | 3 | 3 | 3 | 3 | 3 | 7 |
| 16 | TGME49_281560.P3066 | hypothetical protein | 7 | 3 | 5 | 4 | 4 | 4 | 3 | 7 |
| 26 | TGME49_281590.P3069 | hypothetical protein | 6 | 4 | 3 | 3 | 3 | 3 | 3 | 7 |
| 26 | TGME49_281600.P3070 | hypothetical protein | 5 | 3 | 4 | 3 | 3 | 3 | 3 | 7 |
| 26 | TGME49_281610.P3071 | hypothetical protein | 7 | 3 | 4 | 4 | 3 | 3 | 3 | 7 |
| 24 | TGME49_281930-t26_1-p1 | SAG-related sequence SRS39 | 8 | 6 | 3 | 3 | 3 | 3 | 3 | 7 |
| 9 | TGME49_282020-t26_1-p1 | hypothetical protein | 9 | 3 | 7 | 6 | 6 | 3 | 3 | 7 |
| 9 | TGME49_282140-t26_1-p1 | cwf21 protein | 9 | 3 | 3 | 6 | 4 | 5 | 5 | 7 |

| | | | | | | | | | | |
|---|---|---|---|---|---|---|---|---|---|---|
| 26 | TGME49_283450-t26_1-p1 | Toxoplasma gondii family B protein | 5 | 3 | 3 | 3 | 3 | 3 | 3 | 7 |
| 26 | TGME49_283460-t26_1-p1 | SAG-related sequence SRS21 | 6 | 3 | 3 | 3 | 4 | 3 | 3 | 7 |
| 26 | TGME49_283470.P3083 | Kazal-type serine protease inhibitor domain-containing protein | 5 | 4 | 4 | 3 | 3 | 3 | 3 | 7 |
| 16 | TGME49_283495.P3086 | hypothetical protein | 5 | 3 | 5 | 5 | 4 | 4 | 4 | 7 |
| 9 | TGME49_283500.P3087 | tetratricopeptide repeat-containing protein | 10 | 3 | 6 | 6 | 4 | 5 | 4 | 7 |
| 9 | TGME49_283765.P3100 | B9 domain-containing protein | 9 | 3 | 6 | 3 | 5 | 5 | 5 | 7 |
| 9 | TGME49_283810.P3102 | hypothetical protein | 10 | 3 | 9 | 8 | 3 | 3 | 3 | 7 |
| 20 | TGME49_283840-t26_1-p1 | GATA zinc finger domain-containing protein | 10 | 4 | 4 | 3 | 4 | 3 | 3 | 7 |
| 26 | TGME49_283882-t26_1-p1 | hypothetical protein | 7 | 3 | 4 | 3 | 3 | 3 | 3 | 7 |
| 24 | TGME49_284160.P3105 | histone lysine methyltransferase, SET, putative | 7 | 6 | 5 | 4 | 3 | 4 | 4 | 7 |
| 26 | TGME49_284420-t26_1-p1 | hypothetical protein | 5 | 3 | 3 | 3 | 3 | 3 | 3 | 7 |
| 16 | TGME49_284630.P3112 | hypothetical protein | 5 | 5 | 5 | 5 | 3 | 3 | 3 | 7 |
| 16 | TGME49_284660.P3114 | mitochondrial ribosomal protein s6-2, putative | 6 | 6 | 6 | 3 | 4 | 5 | 3 | 7 |
| 9 | TGME49_285260.P3127 | hypothetical protein | 10 | 5 | 7 | 4 | 4 | 4 | 4 | 7 |
| 24 | TGME49_285300-t26_1-p1 | hypothetical protein | 8 | 7 | 4 | 3 | 3 | 3 | 3 | 7 |
| 20 | TGME49_285425.P3130 | hypothetical protein | 10 | 5 | 4 | 4 | 3 | 3 | 3 | 7 |
| 24 | TGME49_285770.P3137 | coiled-coil domain-containing protein 189, putative | 8 | 6 | 5 | 3 | 3 | 3 | 4 | 7 |
| 24 | TGME49_285790-t26_1-p1 | hypothetical protein | 7 | 6 | 4 | 4 | 3 | 3 | 3 | 7 |
| 26 | TGME49_285820-t26_1-p1 | hypothetical protein | 5 | 3 | 3 | 3 | 3 | 3 | 3 | 7 |
| 24 | TGME49_285860-t26_1-p1 | SAG-related sequence SRS20C | 8 | 6 | 5 | 4 | 4 | 3 | 3 | 7 |
| 24 | TGME49_285930.P3141 | hypothetical protein | 7 | 6 | 4 | 5 | 4 | 3 | 3 | 7 |
| 26 | TGME49_286020-t26_1-p1 | hypothetical protein | 7 | 4 | 3 | 3 | 3 | 3 | 3 | 7 |
| 9 | TGME49_286040-t26_1-p1 | hypothetical protein | 9 | 3 | 5 | 6 | 3 | 3 | 3 | 7 |
| 16 | TGME49_286070.P3146 | cytochrome c oxidase assembly protein PET191, putative | 5 | 5 | 5 | 3 | 5 | 4 | 4 | 7 |
| 26 | TGME49_286460-t26_1-p1 | hypothetical protein | 5 | 3 | 3 | 3 | 3 | 3 | 3 | 7 |
| 26 | TGME49_286465-t26_1-p1 | hypothetical protein | 5 | 3 | 3 | 3 | 3 | 3 | 3 | 7 |
| 20 | TGME49_286510-t26_1-p1 | hypothetical protein | 9 | 3 | 4 | 3 | 3 | 4 | 4 | 7 |
| 16 | TGME49_286560-t26_1-p1 | U6 snRNA-associated Sm family protein | 5 | 5 | 5 | 3 | 5 | 5 | 5 | 7 |
| 26 | TGME49_286570-t26_1-p1 | hypothetical protein | 5 | 3 | 3 | 3 | 3 | 3 | 3 | 7 |
| 9 | TGME49_286610-t26_1-p1 | ribosomal protein RPS14, putative | 8 | 3 | 6 | 6 | 3 | 3 | 4 | 7 |
| 24 | TGME49_286650-t26_1-p1 | hypothetical protein | 7 | 6 | 3 | 5 | 3 | 3 | 3 | 7 |
| 24 | TGME49_286770-t26_1-p1 | hypothetical protein | 6 | 6 | 5 | 5 | 3 | 3 | 3 | 7 |
| 20 | TGME49_286910-t26_1-p1 | hypothetical protein | 9 | 4 | 4 | 3 | 3 | 3 | 3 | 7 |
| 26 | TGME49_287040-t26_1-p1 | hypothetical protein | 7 | 3 | 5 | 4 | 3 | 3 | 3 | 7 |
| 24 | TGME49_287170.P3167 | hypothetical protein | 8 | 6 | 5 | 4 | 3 | 3 | 3 | 7 |
| 16 | TGME49_287200.P3169 | TMEM234 domain-containing protein | 6 | 3 | 5 | 3 | 4 | 4 | 4 | 7 |
| 24 | TGME49_287250-t26_1-p1 | hypothetical protein | 7 | 6 | 5 | 4 | 3 | 3 | 3 | 7 |
| 24 | TGME49_288020-t26_1-p1 | hypothetical protein | 7 | 6 | 3 | 4 | 3 | 3 | 3 | 7 |
| 24 | TGME49_288030-t26_1-p1 | hypothetical protein | 8 | 6 | 5 | 4 | 4 | 3 | 3 | 7 |
| 20 | TGME49_288045-t26_1-p1 | hypothetical protein | 9 | 3 | 4 | 3 | 3 | 3 | 3 | 7 |
| 9 | TGME49_288230.P3180 | serine/threonine protein kinase STK19, putative | 10 | 3 | 7 | 6 | 4 | 3 | 3 | 7 |

| | | | | | | | | | | |
|---|---|---|---|---|---|---|---|---|---|---|
| 26 | TGME49_288265-t26_1-p1 | hypothetical protein | 6 | 3 | 3 | 4 | 3 | 3 | 3 | 7 |
| 24 | TGME49_288310-t26_1-p1 | hypothetical protein | 7 | 7 | 3 | 3 | 3 | 3 | 3 | 7 |
| 16 | TGME49_288580-t26_1-p1 | RNA methylase, putative | 7 | 5 | 6 | 3 | 4 | 5 | 5 | 7 |
| 26 | TGME49_288590-t26_1-p1 | hypothetical protein | 6 | 4 | 4 | 3 | 3 | 3 | 3 | 7 |
| 16 | TGME49_288850.P3193 | hypothetical protein | 5 | 6 | 5 | 5 | 4 | 4 | 3 | 7 |
| 16 | TGME49_288910-t26_1-p1 | DUF866 domain-containing protein | 6 | 5 | 6 | 6 | 6 | 3 | 3 | 7 |
| 9 | TGME49_288940.P3195 | hypothetical protein | 10 | 4 | 8 | 6 | 5 | 5 | 5 | 7 |
| 9 | TGME49_288945-t26_1-p1 | hypothetical protein | 10 | 3 | 7 | 6 | 3 | 5 | 3 | 7 |
| 24 | TGME49_289130-t26_1-p1 | hypothetical protein | 8 | 7 | 4 | 3 | 3 | 3 | 3 | 7 |
| 16 | TGME49_289210.P3199 | prefoldin subunit 4, putative | 6 | 5 | 5 | 5 | 5 | 3 | 4 | 7 |
| 24 | TGME49_289340-t26_1-p1 | hypothetical protein | 8 | 5 | 5 | 4 | 3 | 3 | 3 | 7 |
| 24 | TGME49_289590-t26_1-p1 | hypothetical protein | 7 | 7 | 4 | 3 | 3 | 4 | 3 | 7 |
| 24 | TGME49_289610-t26_1-p1 | hypothetical protein | 7 | 6 | 3 | 5 | 4 | 4 | 4 | 7 |
| 26 | TGME49_289615-t26_1-p1 | hypothetical protein | 7 | 3 | 4 | 5 | 3 | 3 | 3 | 7 |
| 26 | TGME49_289660-t26_1-p1 | hypothetical protein | 6 | 5 | 3 | 3 | 3 | 4 | 3 | 7 |
| 16 | TGME49_289860-t26_1-p1 | hypothetical protein | 6 | 5 | 5 | 5 | 3 | 4 | 3 | 7 |
| 26 | TGME49_289870-t26_1-p1 | hypothetical protein | 6 | 4 | 3 | 3 | 3 | 3 | 3 | 7 |
| 24 | TGME49_289920.P3217 | hypothetical protein | 8 | 6 | 3 | 4 | 3 | 3 | 3 | 7 |
| 24 | TGME49_289970.P3220 | hypothetical protein | 6 | 5 | 5 | 3 | 3 | 3 | 3 | 7 |
| 16 | TGME49_290000.P3221 | hypothetical protein | 7 | 5 | 5 | 4 | 5 | 3 | 4 | 7 |
| 16 | TGME49_290040-t26_1-p1 | macrophage migration inhibitory factor | 6 | 5 | 5 | 4 | 4 | 4 | 4 | 7 |
| 26 | TGME49_290250-t26_1-p1 | hypothetical protein | 6 | 3 | 5 | 3 | 3 | 3 | 3 | 7 |
| 24 | TGME49_290570-t26_1-p1 | hypothetical protein | 6 | 5 | 5 | 4 | 3 | 3 | 3 | 7 |
| 9 | TGME49_290605-t26_1-p1 | hypothetical protein | 9 | 3 | 5 | 6 | 3 | 4 | 4 | 7 |
| 24 | TGME49_290610-t26_1-p1 | hypothetical protein | 7 | 7 | 4 | 3 | 4 | 3 | 3 | 7 |
| 24 | TGME49_290700.P3231 | dense granule protein GRA25 | 7 | 5 | 4 | 3 | 3 | 3 | 3 | 7 |
| 16 | TGME49_291005.P3243 | hypothetical protein | 5 | 3 | 6 | 5 | 3 | 5 | 5 | 7 |
| 16 | TGME49_291160.P3245 | dynein light polypeptide 4, axonemal, putative | 5 | 5 | 5 | 5 | 5 | 4 | 4 | 7 |
| 9 | TGME49_291170.P3246 | vacuolar protein sorting-associated protein IST1, putative | 9 | 3 | 3 | 6 | 3 | 6 | 4 | 7 |
| 24 | TGME49_291330-t26_1-p1 | RNA recognition motif-containing protein | 7 | 6 | 5 | 3 | 5 | 3 | 4 | 7 |
| 24 | TGME49_291630-t26_1-p1 | hypothetical protein | 7 | 6 | 4 | 5 | 3 | 3 | 3 | 7 |
| 24 | TGME49_291660-t26_1-p1 | hypothetical protein | 7 | 6 | 3 | 3 | 3 | 3 | 3 | 7 |
| 24 | TGME49_291840-t26_1-p1 | hypothetical protein | 7 | 6 | 3 | 4 | 3 | 4 | 3 | 7 |
| 9 | TGME49_291860-t26_1-p1 | HORMA domain-containing protein | 9 | 3 | 7 | 6 | 3 | 4 | 3 | 7 |
| 20 | TGME49_292150-t26_1-p1 | hypothetical protein | 9 | 5 | 3 | 3 | 4 | 3 | 3 | 7 |
| 26 | TGME49_292245-t26_1-p1 | hypothetical protein | 5 | 3 | 4 | 3 | 4 | 3 | 3 | 7 |
| 26 | TGME49_292275-t26_1-p1 | SAG-related sequence SRS36E | 6 | 3 | 3 | 3 | 3 | 3 | 3 | 7 |
| 24 | TGME49_292280-t26_1-p1 | SAG-related sequence SRS36D | 6 | 7 | 6 | 3 | 3 | 3 | 3 | 7 |
| 26 | TGME49_292335-t26_1-p1 | hypothetical protein | 6 | 3 | 4 | 3 | 3 | 3 | 3 | 7 |
| 24 | TGME49_292350-t26_1-p1 | hypothetical protein | 6 | 5 | 5 | 3 | 3 | 3 | 3 | 7 |
| 20 | TGME49_292375-t26_1-p1 | KRUF family protein | 10 | 3 | 3 | 3 | 3 | 3 | 3 | 7 |
| 24 | TGME49_292600-t26_1-p1 | hypothetical protein | 7 | 6 | 4 | 3 | 3 | 3 | 3 | 7 |
| 24 | TGME49_293040-t26_1-p1 | hypothetical protein | 7 | 6 | 4 | 4 | 3 | 3 | 3 | 7 |
| 16 | TGME49_293200.P3277 | hypothetical protein | 6 | 4 | 5 | 4 | 4 | 4 | 3 | 7 |
| 16 | TGME49_293310.P3287 | ribosomal protein L20, putative | 6 | 6 | 6 | 5 | 3 | 3 | 3 | 7 |

| | | | | | | | | | | |
|---|---|---|---|---|---|---|---|---|---|---|
| 24 | TGME49_293360.P3292 | inner membrane complex protein IMC38 | 7 | 6 | 5 | 3 | 4 | 3 | 3 | 7 |
| 26 | TGME49_293372-t26_1-p1 | TIP49 family protein | 5 | 3 | 3 | 3 | 3 | 3 | 3 | 7 |
| 24 | TGME49_293400-t26_1-p1 | hypothetical protein | 7 | 6 | 5 | 4 | 3 | 3 | 3 | 7 |
| 26 | TGME49_293410-t26_1-p1 | protein PET117, putative | 5 | 4 | 4 | 3 | 4 | 3 | 3 | 7 |
| 20 | TGME49_293490.P3303 | hypothetical protein | 9 | 5 | 5 | 4 | 3 | 4 | 3 | 7 |
| 26 | TGME49_293740.P3323 | hypothetical protein | 5 | 5 | 4 | 3 | 3 | 3 | 3 | 7 |
| 24 | TGME49_293800.P3329 | transmembrane protein 65, putative | 7 | 6 | 3 | 3 | 4 | 3 | 4 | 7 |
| 16 | TGME49_293870.P3337 | thioredoxin 1 | 5 | 5 | 5 | 5 | 5 | 4 | 4 | 7 |
| 16 | TGME49_294230-t26_1-p1 | hypothetical protein | 5 | 5 | 5 | 4 | 4 | 3 | 4 | 7 |
| 26 | TGME49_294300-t26_1-p1 | hypothetical protein | 6 | 3 | 3 | 3 | 3 | 3 | 3 | 7 |
| 26 | TGME49_294370.P3355 | hypothetical protein | 6 | 3 | 5 | 3 | 3 | 3 | 3 | 7 |
| 16 | TGME49_294390-t26_1-p1 | myosin light chain MLC4, putative | 7 | 4 | 5 | 5 | 3 | 4 | 3 | 7 |
| 20 | TGME49_294410-t26_1-p1 | hypothetical protein | 9 | 5 | 3 | 3 | 3 | 3 | 3 | 7 |
| 26 | TGME49_294595.P3360 | EB1 domain-containing protein | 7 | 3 | 4 | 4 | 3 | 3 | 3 | 7 |
| 9 | TGME49_294750-t26_1-p1 | hypothetical protein | 7 | 3 | 6 | 5 | 4 | 3 | 4 | 7 |
| 20 | TGME49_294785.P3367 | zinc finger (CCCH type) motif-containing protein | 8 | 3 | 3 | 3 | 3 | 3 | 3 | 7 |
| 26 | TGME49_294808-t26_1-p1 | hypothetical protein | 7 | 3 | 3 | 3 | 3 | 3 | 3 | 7 |
| 24 | TGME49_294812-t26_1-p1 | RNA recognition motif-containing protein | 7 | 6 | 3 | 4 | 5 | 4 | 3 | 7 |
| 16 | TGME49_294890.P3374 | hypothetical protein | 6 | 3 | 5 | 4 | 4 | 4 | 3 | 7 |
| 24 | TGME49_294940-t26_1-p1 | hypothetical protein | 8 | 5 | 4 | 4 | 4 | 3 | 3 | 7 |
| 20 | TGME49_294970.P3376 | hypothetical protein | 8 | 4 | 3 | 3 | 3 | 3 | 3 | 7 |
| 16 | TGME49_295035-t26_1-p1 | hypothetical protein | 6 | 5 | 4 | 4 | 4 | 4 | 3 | 7 |
| 9 | TGME49_295080.P3379 | TLC domain-containing protein | 9 | 3 | 7 | 7 | 3 | 3 | 4 | 7 |
| 26 | TGME49_295105-t26_1-p1 | rhoptry protein, putative | 5 | 3 | 3 | 3 | 3 | 3 | 3 | 7 |
| 26 | TGME49_295125-t26_1-p1 | rhoptry protein ROP4 | 5 | 3 | 3 | 3 | 3 | 3 | 3 | 7 |
| 26 | TGME49_295300-t26_1-p1 | Toxoplasma gondii family B protein | 6 | 4 | 3 | 3 | 3 | 3 | 3 | 7 |
| 16 | TGME49_295310-t26_1-p1 | tubulin binding cofactor A protein | 7 | 6 | 5 | 5 | 5 | 4 | 4 | 7 |
| 24 | TGME49_295360.P3382 | inner membrane complex protein IMC18 | 8 | 6 | 5 | 4 | 3 | 3 | 3 | 7 |
| 26 | TGME49_295370.P3383 | hypothetical protein | 7 | 3 | 4 | 4 | 4 | 4 | 4 | 7 |
| 16 | TGME49_295450.P3385 | DIP13 homolog | 6 | 6 | 6 | 5 | 4 | 5 | 4 | 7 |
| 16 | TGME49_295460-t26_1-p1 | Got1 family protein | 6 | 6 | 6 | 6 | 4 | 3 | 3 | 7 |
| 26 | TGME49_295600-t26_1-p1 | SAG-related sequence SRS10 | 6 | 5 | 3 | 3 | 3 | 3 | 3 | 7 |
| 24 | TGME49_295630-t26_1-p1 | hypothetical protein | 6 | 5 | 4 | 3 | 4 | 3 | 3 | 7 |
| 24 | TGME49_295662.P3386 | hypothetical protein | 7 | 6 | 5 | 3 | 4 | 3 | 3 | 7 |
| 20 | TGME49_295740.P3390 | hypothetical protein | 10 | 5 | 4 | 3 | 3 | 3 | 3 | 7 |
| 20 | TGME49_296121-t26_1-p1 | hypothetical protein | 8 | 3 | 3 | 3 | 3 | 3 | 3 | 7 |
| 20 | TGME49_296231-t26_1-p1 | hypothetical protein | 9 | 3 | 3 | 3 | 3 | 3 | 3 | 7 |
| 16 | TGME49_296950-t26_1-p1 | hypothetical protein | 3 | 3 | 9 | 3 | 4 | 3 | 3 | 7 |
| 26 | TGME49_297090-t26_1-p1 | hypothetical protein | 6 | 3 | 3 | 3 | 3 | 3 | 3 | 7 |
| 24 | TGME49_297100-t26_1-p1 | hypothetical protein | 7 | 6 | 5 | 5 | 3 | 3 | 3 | 7 |
| 16 | TGME49_297140.P3401 | U6 snRNA-associated sm family protein Lsm2, putative | 5 | 5 | 5 | 5 | 4 | 5 | 5 | 7 |
| 24 | TGME49_297280.P3402 | hypothetical protein | 7 | 5 | 3 | 3 | 3 | 3 | 3 | 7 |
| 26 | TGME49_297290-t26_1-p1 | hypothetical protein | 7 | 3 | 3 | 3 | 3 | 3 | 3 | 7 |
| 24 | TGME49_297350-t26_1-p1 | hypothetical protein | 7 | 6 | 5 | 5 | 3 | 3 | 3 | 7 |
| 24 | TGME49_297430.P3406 | FoP domain-containing protein | 7 | 6 | 4 | 3 | 4 | 3 | 3 | 7 |

| | | | | | | | | | | |
|---|---|---|---|---|---|---|---|---|---|---|
| 26 | TGME49_297647-t26_1-p1 | hypothetical protein | 6 | 5 | 3 | 3 | 3 | 3 | 3 | 7 |
| 26 | TGME49_297700-t26_1-p1 | hypothetical protein | 5 | 3 | 3 | 3 | 3 | 3 | 3 | 7 |
| 20 | TGME49_297770.P3416 | hypothetical protein | 8 | 4 | 4 | 3 | 4 | 4 | 4 | 7 |
| 9 | TGME49_297780.P3417 | DNA gyrase subunit B | 9 | 3 | 8 | 3 | 3 | 4 | 3 | 7 |
| 20 | TGME49_297790-t26_1-p1 | hypothetical protein | 8 | 4 | 4 | 4 | 3 | 3 | 3 | 7 |
| 9 | TGME49_297830-t26_1-p1 | hypothetical protein | 10 | 3 | 8 | 4 | 4 | 4 | 4 | 7 |
| 26 | TGME49_297845-t26_1-p1 | hypothetical protein | 7 | 3 | 5 | 3 | 3 | 3 | 3 | 7 |
| 24 | TGME49_297880.P3423 | dense granule protein GRA23 | 7 | 6 | 5 | 3 | 4 | 3 | 3 | 7 |
| 9 | TGME49_297940.P3425 | single-stranded DNA-binding protein SSB | 9 | 5 | 6 | 5 | 4 | 5 | 5 | 7 |
| 24 | TGME49_297980.P3427 | hypothetical protein | 6 | 5 | 4 | 3 | 3 | 3 | 3 | 7 |
| 26 | TGME49_298060-t26_1-p1 | Toxoplasma gondii family C protein | 6 | 3 | 3 | 3 | 3 | 3 | 3 | 7 |
| 26 | TGME49_298090-t26_1-p1 | Toxoplasma gondii family B protein | 5 | 3 | 3 | 3 | 3 | 3 | 3 | 7 |
| 24 | TGME49_298590-t26_1-p1 | hypothetical protein | 7 | 5 | 5 | 4 | 4 | 3 | 3 | 7 |
| 9 | TGME49_298600.P3429 | leucine zipper-like transcriptional regulator | 10 | 3 | 9 | 6 | 7 | 4 | 6 | 7 |
| 9 | TGME49_298610.P3430 | GYF domain-containing protein | 10 | 3 | 9 | 5 | 5 | 4 | 4 | 7 |
| 16 | TGME49_298970.P3432 | U6 snRNA-associated Sm-like protein LSm3, putative | 5 | 5 | 5 | 3 | 5 | 3 | 4 | 7 |
| 26 | TGME49_299172.P3437 | hypothetical protein | 7 | 3 | 4 | 4 | 3 | 3 | 3 | 7 |
| 24 | TGME49_299220-t26_1-p1 | hypothetical protein | 8 | 6 | 6 | 3 | 3 | 3 | 3 | 7 |
| 20 | TGME49_299260-t26_1-p1 | hypothetical protein | 8 | 4 | 3 | 3 | 3 | 3 | 3 | 7 |
| 26 | TGME49_299570-t26_1-p1 | Toxoplasma gondii family B protein | 5 | 3 | 3 | 3 | 3 | 3 | 3 | 7 |
| 24 | TGME49_299670.P3442 | hypothetical protein | 8 | 7 | 3 | 5 | 4 | 3 | 3 | 7 |
| 24 | TGME49_299790.P3444 | hypothetical protein | 7 | 6 | 3 | 3 | 3 | 3 | 3 | 7 |
| 24 | TGME49_299800-t26_1-p1 | hypothetical protein | 8 | 6 | 3 | 3 | 3 | 3 | 3 | 7 |
| 24 | TGME49_299980.P3447 | hypothetical protein | 6 | 5 | 4 | 4 | 3 | 3 | 3 | 7 |
| 26 | TGME49_300048-t26_1-p1 | hypothetical protein | 6 | 3 | 3 | 3 | 3 | 3 | 3 | 7 |
| 26 | TGME49_300055-t26_1-p1 | hypothetical protein | 6 | 3 | 3 | 3 | 3 | 3 | 3 | 7 |
| 24 | TGME49_300180.P3459 | hypothetical protein | 6 | 5 | 4 | 4 | 4 | 3 | 3 | 7 |
| 16 | TGME49_300190-t26_1-p1 | ribosomal protein RPL37A | 5 | 5 | 5 | 5 | 5 | 5 | 3 | 7 |
| 9 | TGME49_300240-t26_1-p1 | syntaxin 6 | 7 | 3 | 6 | 5 | 4 | 4 | 3 | 7 |
| 9 | TGME49_300250.P3461 | MtN3/saliva family protein | 9 | 3 | 7 | 7 | 6 | 6 | 3 | 7 |
| 16 | TGME49_300280.P3463 | LSM domain-containing protein | 6 | 6 | 6 | 4 | 5 | 5 | 4 | 7 |
| 16 | TGME49_300290.P3464 | syntaxin-10 | 6 | 6 | 5 | 5 | 4 | 4 | 4 | 7 |
| 16 | TGME49_300601-t26_1-p1 | ribosomal protein S11 | 6 | 3 | 5 | 4 | 5 | 3 | 3 | 7 |
| 9 | TGME49_300606-t26_1-p1 | ribosomal protein S5, putative | 8 | 3 | 6 | 6 | 5 | 3 | 3 | 7 |
| 16 | TGME49_300608-t26_1-p1 | ribosomal protein L6, putative | 6 | 3 | 6 | 5 | 5 | 3 | 3 | 7 |
| 16 | TGME49_300611-t26_1-p1 | ribosomal protein L14 | 6 | 3 | 5 | 5 | 5 | 3 | 3 | 7 |
| 26 | TGME49_300612-t26_1-p1 | ribosomal protein S17 | 5 | 3 | 4 | 4 | 4 | 3 | 3 | 7 |
| 16 | TGME49_300613-t26_1-p1 | ribosomal protein L16, putative | 6 | 3 | 5 | 5 | 4 | 3 | 3 | 7 |
| 9 | TGME49_300615-t26_1-p1 | ribosomal protein S3, putative | 7 | 3 | 6 | 6 | 5 | 3 | 3 | 7 |
| 26 | TGME49_300618-t26_1-p1 | ribosomal protein S19 | 5 | 3 | 4 | 3 | 4 | 3 | 3 | 7 |
| 9 | TGME49_300631-t26_1-p1 | ribosomal protein L4 | 7 | 3 | 7 | 6 | 6 | 3 | 3 | 7 |
| 16 | TGME49_300641-t26_1-p1 | ribosomal protein S4 | 3 | 3 | 6 | 5 | 6 | 3 | 3 | 7 |
| 9 | TGME49_300700-t26_1-p1 | ribosomal protein S2, putative | 7 | 3 | 7 | 6 | 6 | 3 | 3 | 7 |
| 16 | TGME49_301130.P3467 | hypothetical protein | 6 | 5 | 5 | 5 | 3 | 4 | 4 | 7 |
| 20 | TGME49_301140-t26_1-p1 | SAG-related sequence SRS19A | 8 | 3 | 3 | 3 | 3 | 3 | 3 | 7 |
| 20 | TGME49_301180-t26_1-p1 | SAG-related sequence SRS19F | 8 | 3 | 3 | 3 | 3 | 3 | 3 | 7 |
| 20 | TGME49_301250-t26_1-p1 | hypothetical protein | 9 | 4 | 4 | 4 | 3 | 3 | 3 | 7 |

| | | | | | | | | | | |
|---|---|---|---|---|---|---|---|---|---|---|
| 16 | TGME49_301300-t26_1-p1 | PHAX RNA-binding domain protein | 7 | 5 | 5 | 5 | 5 | 4 | 4 | 7 |
| 20 | TGME49_301370.P3476 | palmitoyltransferase DHHC10 | 8 | 3 | 3 | 3 | 4 | 3 | 3 | 7 |
| 26 | TGME49_301470-t26_1-p1 | hypothetical protein | 7 | 4 | 3 | 3 | 3 | 3 | 3 | 7 |
| 20 | TGME49_301690-t26_1-p1 | hypothetical protein | 8 | 5 | 3 | 3 | 3 | 3 | 3 | 7 |
| 26 | TGME49_302003-t26_1-p1 | ORF E | 5 | 3 | 4 | 4 | 3 | 3 | 3 | 7 |
| 16 | TGME49_302055-t26_1-p1 | ribosomal protein S12 | 4 | 3 | 6 | 4 | 5 | 3 | 3 | 7 |
| 16 | TGME49_302057-t26_1-p1 | ribosomal protein S7 | 5 | 3 | 6 | 6 | 5 | 3 | 3 | 7 |
| 20 | TGME49_304480.P3482 | 3-oxo-5-alpha-steroid 4-dehydrogenase | 8 | 3 | 3 | 3 | 3 | 4 | 3 | 7 |
| 9 | TGME49_304670.P3485 | inner membrane complex protein IMC23 | 8 | 3 | 7 | 7 | 3 | 4 | 3 | 7 |
| 26 | TGME49_304980-t26_1-p1 | hypothetical protein | 6 | 3 | 3 | 3 | 3 | 3 | 3 | 7 |
| 20 | TGME49_305000-t26_1-p1 | hypothetical protein | 8 | 3 | 3 | 3 | 3 | 3 | 3 | 7 |
| 24 | TGME49_305050.P3495 | essential light chain ELC2 | 6 | 5 | 5 | 4 | 3 | 3 | 3 | 7 |
| 24 | TGME49_305070-t26_1-p1 | hypothetical protein | 6 | 6 | 5 | 3 | 4 | 3 | 3 | 7 |
| 20 | TGME49_305080-t26_1-p1 | hypothetical protein | 8 | 3 | 4 | 3 | 3 | 3 | 3 | 7 |
| 9 | TGME49_305090-t26_1-p1 | kinase binding protein cgi-121 protein | 8 | 3 | 6 | 6 | 4 | 3 | 3 | 7 |
| 9 | TGME49_305100-t26_1-p1 | hypothetical protein | 10 | 3 | 7 | 3 | 3 | 3 | 3 | 7 |
| 26 | TGME49_305110-t26_1-p1 | hypothetical protein | 7 | 3 | 4 | 3 | 3 | 3 | 3 | 7 |
| 9 | TGME49_305120-t26_1-p1 | transporter, solute:sodium symporter (SSS) family protein | 9 | 3 | 8 | 3 | 7 | 6 | 3 | 7 |
| 16 | TGME49_305160.P3498 | histone H2Ba | 6 | 6 | 6 | 6 | 5 | 3 | 3 | 7 |
| 16 | TGME49_305220-t26_1-p1 | C-Myc-binding family protein | 5 | 5 | 5 | 3 | 3 | 4 | 3 | 7 |
| 20 | TGME49_305280-t26_1-p1 | hypothetical protein | 9 | 3 | 3 | 3 | 3 | 3 | 3 | 7 |
| 20 | TGME49_305498-t26_1-p1 | hypothetical protein | 8 | 3 | 3 | 3 | 3 | 3 | 3 | 7 |
| 26 | TGME49_305635-t26_1-p1 | hypothetical protein | 5 | 3 | 4 | 3 | 3 | 3 | 3 | 7 |
| 9 | TGME49_305760-t26_1-p1 | zinc finger MIZ domain-containing protein | 8 | 3 | 6 | 4 | 4 | 4 | 4 | 7 |
| 9 | TGME49_305800-t26_1-p1 | 6-pyruvoyl tetrahydrobiopterin synthase | 9 | 3 | 7 | 6 | 6 | 6 | 5 | 7 |
| 16 | TGME49_305830.P3509 | EF hand domain-containing protein | 6 | 5 | 5 | 6 | 3 | 4 | 4 | 7 |
| 9 | TGME49_305840-t26_1-p1 | Nicotinate-nucleotide adenylyltransferase | 8 | 3 | 7 | 6 | 6 | 4 | 3 | 7 |
| 26 | TGME49_305870-t26_1-p1 | DAD family protein | 7 | 4 | 4 | 5 | 3 | 4 | 3 | 7 |
| 16 | TGME49_305900-t26_1-p1 | hypothetical protein | 7 | 5 | 5 | 5 | 5 | 4 | 4 | 7 |
| 24 | TGME49_305930.P3511 | IMC sutures component ISC4 | 7 | 5 | 4 | 4 | 3 | 3 | 3 | 7 |
| 26 | TGME49_306195-t26_1-p1 | DUF1764 domain-containing protein | 7 | 3 | 5 | 4 | 3 | 4 | 3 | 7 |
| 24 | TGME49_306270-t26_1-p1 | hypothetical protein | 6 | 5 | 4 | 3 | 3 | 3 | 3 | 7 |
| 26 | TGME49_306334-t26_1-p1 | hypothetical protein | 7 | 3 | 3 | 3 | 3 | 3 | 3 | 7 |
| 16 | TGME49_306380-t26_1-p1 | U1 zinc finger protein | 7 | 5 | 5 | 4 | 4 | 4 | 4 | 7 |
| 16 | TGME49_306390.P3521 | cytochrome c oxidase subunit 6B, putative | 6 | 5 | 5 | 5 | 3 | 3 | 4 | 7 |
| 24 | TGME49_306530-t26_1-p1 | hypothetical protein | 8 | 5 | 4 | 4 | 4 | 3 | 3 | 7 |
| 26 | TGME49_306680-t26_1-p1 | hypothetical protein | 7 | 3 | 3 | 3 | 3 | 3 | 3 | 7 |
| 9 | TGME49_306895-t26_1-p1 | hypothetical protein | 7 | 3 | 6 | 5 | 3 | 3 | 4 | 7 |
| 16 | TGME49_307000-t26_1-p1 | hypothetical protein | 6 | 4 | 4 | 4 | 4 | 3 | 4 | 7 |
| 26 | TGME49_307070-t26_1-p1 | hypothetical protein | 5 | 3 | 3 | 3 | 3 | 3 | 3 | 7 |
| 9 | TGME49_307603.P3536 | adenylate kinase | 9 | 3 | 7 | 7 | 3 | 6 | 4 | 7 |
| 9 | TGME49_307605-t26_1-p1 | hypothetical protein | 7 | 3 | 6 | 6 | 4 | 3 | 3 | 7 |
| 24 | TGME49_307830-t26_1-p1 | hypothetical protein | 7 | 6 | 4 | 4 | 4 | 4 | 4 | 7 |

| | | | | | | | | | | |
|---|---|---|---|---|---|---|---|---|---|---|
| 9 | TGME49_308010.P3540 | hypothetical protein | 10 | 4 | 6 | 4 | 4 | 4 | 3 | 7 |
| 20 | TGME49_308096-t26_1-p1 | rhoptry kinase family protein (incomplete catalytic triad) | 9 | 3 | 4 | 3 | 3 | 3 | 3 | 7 |
| 24 | TGME49_308840.P3544 | SAG-related sequence SRS51 | 6 | 6 | 3 | 3 | 3 | 3 | 3 | 7 |
| 20 | TGME49_308965.P3552 | hypothetical protein | 10 | 3 | 3 | 5 | 3 | 3 | 3 | 7 |
| 20 | TGME49_309080-t26_1-p1 | hypothetical protein | 10 | 5 | 4 | 3 | 3 | 3 | 3 | 7 |
| 26 | TGME49_309160.P3561 | IgA-specific metalloendopeptidase | 6 | 4 | 3 | 3 | 3 | 3 | 3 | 7 |
| 20 | TGME49_309330-t26_1-p1 | SAG-related sequence SRS55F | 8 | 3 | 4 | 3 | 3 | 3 | 3 | 7 |
| 20 | TGME49_309352-t26_1-p1 | SAG-related sequence SRS55N | 9 | 3 | 3 | 3 | 3 | 3 | 3 | 7 |
| 26 | TGME49_309590.P3572 | rhoptry protein ROP1 | 7 | 4 | 3 | 3 | 3 | 3 | 3 | 7 |
| 16 | TGME49_309610-t26_1-p1 | hypothetical protein | 6 | 4 | 5 | 4 | 3 | 4 | 3 | 7 |
| 24 | TGME49_309780-t26_1-p1 | hypothetical protein | 7 | 6 | 5 | 4 | 4 | 3 | 3 | 7 |
| 16 | TGME49_309810-t26_1-p1 | ribosomal protein RPP2 | 6 | 5 | 4 | 3 | 4 | 4 | 4 | 7 |
| 24 | TGME49_309840.P3578 | hypothetical protein | 8 | 6 | 3 | 3 | 3 | 3 | 3 | 7 |
| 20 | TGME49_309885.P3582 | hypothetical protein | 9 | 5 | 4 | 3 | 3 | 3 | 3 | 7 |
| 26 | TGME49_310055-t26_1-p1 | hypothetical protein | 5 | 3 | 3 | 3 | 3 | 3 | 3 | 7 |
| 26 | TGME49_310177.P3599 | hypothetical protein | 6 | 3 | 3 | 3 | 3 | 3 | 3 | 7 |
| 16 | TGME49_310180.P3600 | ATP synthase-associated protein | 5 | 5 | 5 | 3 | 4 | 4 | 4 | 7 |
| 26 | TGME49_310215-t26_1-p1 | hypothetical protein | 5 | 3 | 3 | 3 | 3 | 3 | 3 | 7 |
| 20 | TGME49_310235-t26_1-p1 | hypothetical protein | 8 | 3 | 4 | 4 | 3 | 3 | 4 | 7 |
| 20 | TGME49_310260-t26_1-p1 | hypothetical protein | 8 | 5 | 3 | 3 | 3 | 3 | 3 | 7 |
| 20 | TGME49_310275-t26_1-p1 | hypothetical protein | 9 | 3 | 3 | 3 | 3 | 3 | 3 | 7 |
| 20 | TGME49_310340-t26_1-p1 | hypothetical protein | 8 | 3 | 3 | 3 | 3 | 3 | 3 | 7 |
| 16 | TGME49_310360-t26_1-p1 | ATP synthase-associated protein | 7 | 5 | 5 | 5 | 3 | 5 | 3 | 7 |
| 26 | TGME49_310420.P3608 | hypothetical protein | 5 | 4 | 3 | 3 | 3 | 3 | 3 | 7 |
| 26 | TGME49_310510.P3612 | hypothetical protein | 5 | 4 | 4 | 4 | 4 | 3 | 3 | 7 |
| 16 | TGME49_310600-t26_1-p1 | heat shock factor binding protein 1 protein | 6 | 4 | 4 | 4 | 4 | 4 | 4 | 7 |
| 16 | TGME49_310610-t26_1-p1 | hypothetical protein | 7 | 5 | 5 | 4 | 5 | 4 | 3 | 7 |
| 24 | TGME49_310690-t26_1-p1 | hypothetical protein | 7 | 5 | 4 | 3 | 3 | 4 | 3 | 7 |
| 26 | TGME49_310780-t26_1-p1 | dense granule protein GRA4 | 6 | 4 | 3 | 3 | 3 | 3 | 3 | 7 |
| 24 | TGME49_310790.P3623 | cyst wall protein CST9 | 7 | 6 | 5 | 4 | 3 | 3 | 3 | 7 |
| 26 | TGME49_310840.P3625 | hypothetical protein | 5 | 4 | 4 | 4 | 3 | 3 | 3 | 7 |
| 26 | TGME49_310940-t26_1-p1 | hypothetical protein | 6 | 3 | 3 | 3 | 3 | 3 | 3 | 7 |
| 16 | TGME49_311000.P3630 | hypothetical protein | 7 | 5 | 5 | 5 | 4 | 3 | 3 | 7 |
| 24 | TGME49_311010.P3631 | hypothetical protein | 7 | 6 | 4 | 3 | 3 | 3 | 3 | 7 |
| 24 | TGME49_311020.P3632 | hypothetical protein | 8 | 6 | 5 | 4 | 4 | 3 | 3 | 7 |
| 26 | TGME49_311075-t26_1-p1 | hypothetical protein | 5 | 3 | 3 | 3 | 3 | 3 | 3 | 7 |
| 16 | TGME49_311110-t26_1-p1 | ubiquitin-fold modifier 1 | 5 | 5 | 3 | 3 | 5 | 5 | 4 | 7 |
| 26 | TGME49_311155.P3635 | hypothetical protein | 5 | 3 | 4 | 4 | 3 | 3 | 3 | 7 |
| 24 | TGME49_311170-t26_1-p1 | hypothetical protein | 6 | 5 | 4 | 3 | 3 | 3 | 4 | 7 |
| 9 | TGME49_311220.P3638 | hypothetical protein | 10 | 3 | 7 | 5 | 5 | 4 | 4 | 7 |
| 16 | TGME49_311260-t26_1-p1 | myosin light chain MLC5 | 6 | 6 | 5 | 5 | 5 | 3 | 4 | 7 |
| 26 | TGME49_311375.P3646 | hypothetical protein | 6 | 3 | 3 | 3 | 3 | 5 | 3 | 7 |
| 24 | TGME49_311420-t26_1-p1 | hypothetical protein | 7 | 6 | 3 | 3 | 3 | 3 | 3 | 7 |
| 24 | TGME49_311440-t26_1-p1 | SAG-related sequence SRS50 | 7 | 6 | 3 | 3 | 3 | 3 | 3 | 7 |
| 26 | TGME49_311485-t26_1-p1 | hypothetical protein | 6 | 3 | 3 | 3 | 3 | 3 | 3 | 7 |
| 16 | TGME49_311680-t26_1-p1 | FUN14 family protein | 7 | 6 | 6 | 3 | 5 | 4 | 4 | 7 |
| 26 | TGME49_311815-t26_1-p1 | hypothetical protein | 5 | 3 | 3 | 3 | 3 | 3 | 3 | 7 |
| 16 | TGME49_311830-t26_1-p1 | hypothetical protein | 6 | 6 | 5 | 5 | 4 | 3 | 3 | 7 |

| | | | | | | | | | | |
|---|---|---|---|---|---|---|---|---|---|---|
| 24 | TGME49_311840-t26_1-p1 | hypothetical protein | 7 | 6 | 3 | 5 | 3 | 3 | 3 | 7 |
| 24 | TGME49_312160.P3664 | hypothetical protein | 6 | 6 | 5 | 4 | 4 | 3 | 3 | 7 |
| 20 | TGME49_312175-t26_1-p1 | hypothetical protein | 10 | 5 | 4 | 4 | 3 | 3 | 3 | 7 |
| 26 | TGME49_312210-t26_1-p1 | hypothetical protein | 6 | 3 | 3 | 3 | 3 | 3 | 3 | 7 |
| 20 | TGME49_312240.P3666 | hypothetical protein | 10 | 3 | 5 | 4 | 4 | 3 | 3 | 7 |
| 16 | TGME49_312250-t26_1-p1 | DNA-directed RNA polymerase III RPC11 | 6 | 6 | 5 | 5 | 5 | 5 | 3 | 7 |
| 24 | TGME49_312270-t26_1-p1 | rhoptry protein ROP13 | 8 | 7 | 3 | 3 | 3 | 3 | 3 | 7 |
| 24 | TGME49_312340-t26_1-p1 | hypothetical protein | 8 | 6 | 4 | 4 | 3 | 3 | 3 | 7 |
| 26 | TGME49_312460.P3672 | zinc finger, MYND-type containing 12 family protein | 6 | 3 | 3 | 6 | 3 | 3 | 3 | 7 |
| 16 | TGME49_312470-t26_1-p1 | hypothetical protein | 6 | 5 | 5 | 5 | 3 | 3 | 3 | 7 |
| 26 | TGME49_312540-t26_1-p1 | hypothetical protein | 7 | 3 | 3 | 5 | 3 | 5 | 4 | 7 |
| 24 | TGME49_312600-t26_1-p1 | heat shock protein HSP21 | 7 | 6 | 6 | 3 | 3 | 3 | 3 | 7 |
| 16 | TGME49_312680.P3681 | 60S ribosomal protein L27, putative | 5 | 5 | 5 | 3 | 4 | 4 | 3 | 7 |
| 26 | TGME49_312690-t26_1-p1 | hypothetical protein | 5 | 3 | 3 | 3 | 3 | 3 | 3 | 7 |
| 26 | TGME49_313025-t26_1-p1 | hypothetical protein | 5 | 3 | 3 | 3 | 3 | 3 | 3 | 7 |
| 24 | TGME49_313070-t26_1-p1 | hypothetical protein | 8 | 5 | 4 | 4 | 4 | 3 | 3 | 7 |
| 16 | TGME49_313160-t26_1-p1 | hypothetical protein | 6 | 6 | 6 | 6 | 5 | 3 | 3 | 7 |
| 20 | TGME49_313210.P3694 | hypothetical protein | 10 | 3 | 4 | 3 | 3 | 3 | 3 | 7 |
| 26 | TGME49_313235-t26_1-p1 | hypothetical protein | 7 | 3 | 6 | 3 | 3 | 3 | 3 | 7 |
| 20 | TGME49_313318.P3699 | hypothetical protein | 8 | 3 | 4 | 4 | 3 | 3 | 3 | 7 |
| 26 | TGME49_313385-t26_1-p1 | hypothetical protein | 5 | 3 | 4 | 3 | 4 | 4 | 3 | 7 |
| 24 | TGME49_313440.P3706 | dense granule protein GRA59 | 7 | 5 | 3 | 3 | 3 | 3 | 3 | 7 |
| 16 | TGME49_313450-t26_1-p1 | AP-2 complex subunit sigma-1, putative | 6 | 6 | 6 | 5 | 6 | 3 | 4 | 7 |
| 26 | TGME49_313677-t26_1-p1 | hypothetical protein | 5 | 3 | 3 | 3 | 3 | 3 | 3 | 7 |
| 9 | TGME49_313710-t26_1-p1 | hypothetical protein | 9 | 3 | 7 | 6 | 4 | 3 | 3 | 7 |
| 16 | TGME49_313740.P3720 | zinc finger (CCCH type) motif-containing protein | 5 | 5 | 6 | 3 | 3 | 4 | 3 | 7 |
| 26 | TGME49_313848-t26_1-p1 | hypothetical protein | 7 | 3 | 3 | 3 | 3 | 3 | 3 | 7 |
| 20 | TGME49_313852-t26_1-p1 | hypothetical protein | 10 | 3 | 4 | 4 | 3 | 3 | 3 | 7 |
| 20 | TGME49_313890.P3724 | hypothetical protein | 10 | 3 | 3 | 4 | 4 | 3 | 3 | 7 |
| 24 | TGME49_313940-t26_1-p1 | hypothetical protein | 8 | 7 | 4 | 4 | 3 | 3 | 3 | 7 |
| 16 | TGME49_314000-t26_1-p1 | peptide methionine sulfoxide reductase msrB, putative | 7 | 6 | 4 | 4 | 3 | 5 | 5 | 7 |
| 24 | TGME49_314080-t26_1-p1 | hypothetical protein | 7 | 6 | 5 | 5 | 3 | 3 | 3 | 7 |
| 26 | TGME49_314250-t26_1-p1 | bradyzoite rhoptry protein BRP1 | 5 | 3 | 4 | 3 | 3 | 3 | 3 | 7 |
| 26 | TGME49_314260-t26_1-p1 | hypothetical protein | 6 | 4 | 4 | 3 | 3 | 3 | 3 | 7 |
| 26 | TGME49_314310.P3737 | hypothetical protein | 5 | 3 | 4 | 4 | 3 | 3 | 3 | 7 |
| 24 | TGME49_314415-t26_1-p1 | hypothetical protein | 7 | 6 | 5 | 4 | 5 | 3 | 3 | 7 |
| 20 | TGME49_314422.P3742 | rhodanese family domain-containing protein | 8 | 3 | 5 | 5 | 3 | 3 | 3 | 7 |
| 26 | TGME49_314482-t26_1-p1 | hypothetical protein | 6 | 3 | 3 | 4 | 3 | 3 | 3 | 7 |
| 26 | TGME49_314485.P3745 | hypothetical protein | 7 | 3 | 3 | 3 | 3 | 3 | 3 | 7 |
| 24 | TGME49_314550-t26_1-p1 | hypothetical protein | 8 | 7 | 3 | 3 | 3 | 3 | 3 | 7 |
| 26 | TGME49_314650-t26_1-p1 | hypothetical protein | 5 | 3 | 3 | 3 | 3 | 3 | 3 | 7 |
| 20 | TGME49_314670-t26_1-p1 | hypothetical protein | 8 | 3 | 5 | 3 | 3 | 3 | 3 | 7 |
| 26 | TGME49_314715-t26_1-p1 | hypothetical protein | 5 | 3 | 3 | 3 | 3 | 3 | 3 | 7 |

| | | | | | | | | | | |
|---|---|---|---|---|---|---|---|---|---|---|
| 16 | TGME49_314790-t26_1-p1 | small nuclear ribonucleoprotein G, putative | 5 | 5 | 5 | 5 | 3 | 5 | 5 | 7 |
| 16 | TGME49_314800.P3748 | hypothetical protein | 5 | 5 | 5 | 5 | 3 | 4 | 4 | 7 |
| 16 | TGME49_314820-t26_1-p1 | atp synthase F1, epsilon subunit, putative | 5 | 5 | 5 | 4 | 4 | 4 | 4 | 7 |
| 16 | TGME49_314830-t26_1-p1 | pre-mRNA splicing factor subunit, putative | 6 | 5 | 5 | 5 | 5 | 4 | 4 | 7 |
| 16 | TGME49_314910.P3753 | starch binding domain-containing protein | 7 | 6 | 6 | 3 | 5 | 5 | 4 | 7 |
| 26 | TGME49_315090-t26_1-p1 | hypothetical protein | 7 | 4 | 3 | 3 | 3 | 3 | 3 | 7 |
| 16 | TGME49_315120-t26_1-p1 | hypothetical protein | 7 | 6 | 5 | 5 | 5 | 4 | 4 | 7 |
| 24 | TGME49_315320.P3763 | SAG-related sequence SRS52A | 8 | 6 | 4 | 3 | 3 | 3 | 3 | 7 |
| 26 | TGME49_315330-t26_1-p1 | SAG-related sequence SRS52D | 6 | 3 | 3 | 3 | 3 | 3 | 3 | 7 |
| 26 | TGME49_315340-t26_1-p1 | SAG-related sequence SRS52C | 7 | 3 | 4 | 4 | 3 | 3 | 3 | 7 |
| 26 | TGME49_315345-t26_1-p1 | SAG-related sequence SRS52F | 7 | 3 | 3 | 3 | 3 | 3 | 3 | 7 |
| 9 | TGME49_315350-t26_1-p1 | vacuolar protein sorting-associated protein 46, putative | 8 | 3 | 6 | 6 | 3 | 5 | 5 | 7 |
| 24 | TGME49_315370.P3765 | SAG-related sequence SRS53A | 8 | 6 | 4 | 4 | 3 | 3 | 3 | 7 |
| 24 | TGME49_315380.P3766 | SAG-related sequence SRS53B | 8 | 6 | 5 | 4 | 3 | 3 | 3 | 7 |
| 24 | TGME49_315390.P3767 | SAG-related sequence SRS53C | 8 | 6 | 4 | 4 | 4 | 3 | 3 | 7 |
| 24 | TGME49_315410-t26_1-p1 | SAG-related sequence SRS53F | 8 | 6 | 5 | 4 | 3 | 3 | 3 | 7 |
| 24 | TGME49_315470.P3771 | hypothetical protein | 7 | 5 | 3 | 4 | 4 | 3 | 3 | 7 |
| 20 | TGME49_315490.P3773 | rhoptry protein ROP10 | 9 | 5 | 4 | 4 | 3 | 4 | 3 | 7 |
| 26 | TGME49_315610-t26_1-p1 | hypothetical protein | 5 | 4 | 3 | 3 | 3 | 3 | 3 | 7 |
| 26 | TGME49_315630.P3783 | hypothetical protein | 6 | 3 | 5 | 4 | 3 | 3 | 3 | 7 |
| 20 | TGME49_315740-t26_1-p1 | SAG-related sequence SRS54 | 8 | 3 | 5 | 3 | 3 | 3 | 3 | 7 |
| 16 | TGME49_315750.P3787 | inner membrane complex protein IMC26 | 7 | 5 | 6 | 5 | 5 | 4 | 4 | 7 |
| 20 | TGME49_315820-t26_1-p1 | hypothetical protein | 9 | 4 | 4 | 3 | 3 | 3 | 3 | 7 |
| 24 | TGME49_315910.P3793 | hypothetical protein | 8 | 5 | 5 | 5 | 3 | 3 | 3 | 7 |
| 9 | TGME49_316110-t26_1-p1 | ribosomal protein RPS15 | 9 | 5 | 6 | 6 | 5 | 4 | 4 | 7 |
| 26 | TGME49_316235-t26_1-p1 | hypothetical protein | 6 | 3 | 5 | 3 | 3 | 3 | 3 | 7 |
| 26 | TGME49_316255.P3801 | hypothetical protein | 5 | 3 | 5 | 4 | 3 | 4 | 3 | 7 |
| 20 | TGME49_316290-t26_1-p1 | hypothetical protein | 8 | 3 | 3 | 3 | 3 | 3 | 3 | 7 |
| 16 | TGME49_316360-t26_1-p1 | hypothetical protein | 6 | 6 | 5 | 5 | 4 | 4 | 4 | 7 |
| 26 | TGME49_316410-t26_1-p1 | hypothetical protein | 6 | 3 | 3 | 3 | 3 | 3 | 3 | 7 |
| 24 | TGME49_316420-t26_1-p1 | leucine rich repeat-containing protein | 7 | 6 | 3 | 3 | 3 | 4 | 3 | 7 |
| 26 | TGME49_316580-t26_1-p1 | hypothetical protein | 5 | 3 | 3 | 3 | 3 | 3 | 3 | 7 |
| 26 | TGME49_316600.P3814 | selenoprotein K, putative | 5 | 4 | 4 | 3 | 4 | 3 | 3 | 7 |
| 26 | TGME49_316690-t26_1-p1 | hypothetical protein | 6 | 3 | 3 | 3 | 3 | 3 | 3 | 7 |
| 9 | TGME49_316700-t26_1-p1 | uridine kinase | 9 | 3 | 7 | 6 | 4 | 3 | 3 | 7 |
| 26 | TGME49_316710-t26_1-p1 | hypothetical protein | 5 | 3 | 3 | 3 | 3 | 3 | 3 | 7 |
| 20 | TGME49_316780-t26_1-p1 | hypothetical protein | 9 | 3 | 3 | 3 | 3 | 3 | 3 | 7 |
| 26 | TGME49_317820-t26_1-p1 | hypothetical protein | 5 | 3 | 3 | 3 | 3 | 3 | 3 | 7 |
| 26 | TGME49_318180.P3823 | peroxisomal membrane protein PEX16, putative | 6 | 3 | 3 | 4 | 3 | 3 | 4 | 7 |
| 26 | TGME49_318340.P3827 | hypothetical protein | 6 | 4 | 4 | 4 | 4 | 3 | 3 | 7 |
| 26 | TGME49_318380-t26_1-p1 | hypothetical protein | 5 | 3 | 3 | 3 | 3 | 3 | 3 | 7 |
| 24 | TGME49_318450-t26_1-p1 | zinc finger, C3HC4 type (RING finger) domain-containing protein | 8 | 7 | 3 | 3 | 3 | 3 | 3 | 7 |

| | | | | | | | | | | |
|---|---|---|---|---|---|---|---|---|---|---|
| 20 | TGME49_318500-t26_1-p1 | cpw-wpc domain-containing protein | 8 | 4 | 3 | 3 | 3 | 3 | 3 | 7 |
| 24 | TGME49_318550-t26_1-p1 | hypothetical protein | 7 | 6 | 5 | 3 | 3 | 3 | 3 | 7 |
| 20 | TGME49_319350-t26_1-p1 | SAG-related sequence SRS17B | 8 | 3 | 3 | 3 | 3 | 3 | 3 | 7 |
| 24 | TGME49_319510-t26_1-p1 | hypothetical protein | 7 | 6 | 4 | 4 | 3 | 3 | 3 | 7 |
| 9 | TGME49_319590.P3853 | helicase, putative | 10 | 3 | 7 | 6 | 4 | 5 | 4 | 7 |
| 24 | TGME49_319630-t26_1-p1 | hypothetical protein | 7 | 5 | 4 | 3 | 3 | 3 | 3 | 7 |
| 16 | TGME49_319670-t26_1-p1 | hypothetical protein | 8 | 5 | 5 | 3 | 5 | 4 | 4 | 7 |
| 24 | TGME49_319690-t26_1-p1 | hypothetical protein | 6 | 6 | 5 | 3 | 3 | 3 | 4 | 7 |
| 26 | TGME49_319695-t26_1-p1 | hypothetical protein | 5 | 3 | 5 | 5 | 3 | 3 | 3 | 7 |
| 16 | TGME49_319730-t26_1-p1 | YOU2 family C2C2 zinc finger protein | 6 | 6 | 6 | 5 | 4 | 4 | 4 | 7 |
| 24 | TGME49_319890.P3860 | tyrosine-rich oocyst wall protein | 6 | 5 | 4 | 3 | 3 | 3 | 3 | 7 |
| 16 | TGME49_319960.P3865 | hypothetical protein | 6 | 3 | 5 | 5 | 4 | 3 | 3 | 7 |
| 20 | TGME49_320015.P3867 | hypothetical protein | 8 | 3 | 5 | 3 | 3 | 3 | 3 | 7 |
| 26 | TGME49_320105-t26_1-p1 | hypothetical protein | 5 | 3 | 3 | 3 | 3 | 3 | 3 | 7 |
| 16 | TGME49_320130.P3870 | hypothetical protein | 6 | 5 | 5 | 4 | 4 | 4 | 4 | 7 |
| 16 | TGME49_320140.P3871 | cytochrome b-c1 complex subunit 6, putative | 5 | 5 | 5 | 3 | 4 | 5 | 4 | 7 |
| 24 | TGME49_320180-t26_1-p1 | SAG-related sequence SRS16C | 8 | 7 | 4 | 3 | 3 | 3 | 3 | 7 |
| 20 | TGME49_320200-t26_1-p1 | SAG-related sequence SRS16A | 8 | 3 | 4 | 3 | 3 | 3 | 3 | 7 |
| 26 | TGME49_320250-t26_1-p1 | SAG-related sequence SRS15A | 5 | 3 | 6 | 3 | 3 | 3 | 3 | 7 |
| 26 | TGME49_320470.P3877 | hypothetical protein | 5 | 4 | 4 | 3 | 4 | 3 | 4 | 7 |
| 26 | TGME49_320525.P3881 | DUF1308 domain-containing protein | 7 | 3 | 6 | 4 | 3 | 3 | 3 | 7 |
| 9 | TGME49_320580-t26_1-p1 | hypothetical protein | 8 | 3 | 7 | 3 | 3 | 5 | 3 | 7 |
| 16 | TGME49_320600.P3884 | cold-shock DNA-binding domain-containing protein | 6 | 6 | 6 | 5 | 3 | 5 | 3 | 7 |
| 9 | TGME49_320715-t26_1-p1 | hypothetical protein | 10 | 3 | 7 | 7 | 3 | 3 | 3 | 7 |
| 16 | TGME49_320720.P3887 | hypothetical protein | 6 | 6 | 6 | 5 | 4 | 3 | 3 | 7 |
| 16 | TGME49_321400-t26_1-p1 | microsomal signal peptidase (spc12) domain-containing protein | 6 | 5 | 5 | 5 | 4 | 3 | 4 | 7 |
| 24 | TGME49_321490-t26_1-p1 | SAG-related sequence SRS12A | 6 | 7 | 6 | 3 | 3 | 3 | 3 | 7 |
| 26 | TGME49_321550-t26_1-p1 | hypothetical protein | 7 | 4 | 4 | 3 | 3 | 3 | 3 | 7 |
| 9 | TGME49_321580-t26_1-p1 | membrane protein, putative | 8 | 3 | 7 | 3 | 3 | 6 | 4 | 7 |
| 16 | TGME49_322210-t26_1-p1 | apocytochrome b, putative | 3 | 3 | 8 | 4 | 3 | 3 | 3 | 7 |
| 26 | TGME49_322500-t26_1-p1 | hypothetical protein | 5 | 3 | 3 | 3 | 3 | 3 | 3 | 7 |
| 26 | TGME49_323010-t26_1-p1 | hypothetical protein | 6 | 3 | 3 | 3 | 3 | 3 | 3 | 7 |
| 26 | TGME49_323020-t26_1-p1 | KRUF family protein | 7 | 3 | 3 | 3 | 3 | 3 | 3 | 7 |
| 26 | TGME49_324600-t26_1-p1 | heat shock protein | 7 | 3 | 3 | 3 | 3 | 3 | 3 | 7 |
| 26 | TGME49_326200-t26_1-p1 | type I fatty acid synthase | 7 | 3 | 3 | 3 | 3 | 3 | 3 | 7 |
| 20 | TGME49_329710-t26_1-p1 | SAG-related sequence SRS26I | 9 | 3 | 5 | 4 | 3 | 3 | 3 | 7 |
| 20 | TGME49_329800-t26_1-p1 | hypothetical protein | 9 | 3 | 3 | 3 | 3 | 3 | 3 | 7 |
| 16 | TGME49_330000-t26_1-p1 | cytochrome b | 3 | 3 | 9 | 3 | 3 | 3 | 3 | 7 |
| 20 | TGME49_500001.P4 | hypothetical protein, conserved | 8 | 3 | 4 | 4 | 3 | 3 | 3 | 7 |
| 9 | TGME49_500005.P8 | apicoplast DER1 | 9 | 3 | 7 | 6 | 3 | 3 | 3 | 7 |
| 26 | TGME49_500015.P18 | hypothetical protein, conserved | 7 | 3 | 3 | 3 | 3 | 3 | 3 | 7 |
| 24 | TGME49_500031.P34 | hypothetical protein, conserved | 7 | 5 | 5 | 3 | 3 | 3 | 3 | 7 |
| 9 | TGME49_500032.P35 | DLEC1 domain-containing protein, putative | 9 | 3 | 9 | 3 | 3 | 4 | 3 | 7 |
| 26 | TGME49_500048.P349 | hypothetical protein, conserved | 7 | 3 | 4 | 4 | 3 | 4 | 4 | 7 |
| 24 | TGME49_500052.P353 | hypothetical protein, conserved | 6 | 6 | 4 | 3 | 3 | 3 | 3 | 7 |
| 16 | TGME49_500054.P355 | c-Myc-binding protein, putative | 5 | 7 | 6 | 5 | 3 | 5 | 4 | 7 |

| | | | | | | | | | | |
|---|---|---|---|---|---|---|---|---|---|---|
| 16 | TGME49_500055.P356 | arginine and glutamate-rich protein, putative | 6 | 7 | 7 | 3 | 5 | 3 | 3 | 7 |
| 16 | TGME49_500068.P369 | hypothetical protein, conserved | 3 | 3 | 9 | 7 | 3 | 5 | 4 | 7 |
| 9 | TGME49_500069.P370 | hypothetical protein, conserved | 7 | 3 | 5 | 5 | 3 | 4 | 4 | 7 |
| 20 | TGME49_500071.P372 | sugar transporter, putative | 9 | 3 | 4 | 4 | 3 | 3 | 3 | 7 |
| 26 | TGME49_500075.P376 | tetratricopeptide repeat-containing protein | 5 | 3 | 3 | 3 | 4 | 4 | 3 | 7 |
| 20 | TGME49_500086.P387 | hypothetical protein, conserved | 10 | 4 | 3 | 3 | 3 | 3 | 3 | 7 |
| 16 | TGME49_500093.P394 | EF hand domain-containing protein | 3 | 3 | 7 | 3 | 5 | 3 | 3 | 7 |
| 16 | TGME49_500094.P395 | transmembrane protein, putative | 3 | 3 | 7 | 6 | 4 | 4 | 3 | 7 |
| 20 | TGME49_500100.P401 | transmembrane protein TMEM42, putative | 8 | 3 | 5 | 3 | 3 | 3 | 3 | 7 |
| 20 | TGME49_500108.P409 | hypothetical protein, conserved | 9 | 3 | 4 | 3 | 3 | 3 | 3 | 7 |
| 26 | TGME49_500111.P412 | Toxoplasma gondii family B protein | 6 | 3 | 3 | 3 | 3 | 3 | 3 | 7 |
| 24 | TGME49_500117.P419 | hypothetical protein, conserved | 8 | 6 | 4 | 3 | 3 | 3 | 3 | 7 |
| 24 | TGME49_500118.P420 | hypothetical protein, conserved | 7 | 6 | 5 | 5 | 3 | 3 | 3 | 7 |
| 16 | TGME49_500119.P421 | CRAL/TRIO domain-containing protein, putative | 3 | 3 | 6 | 6 | 3 | 4 | 3 | 7 |
| 9 | TGME49_500132.P434 | hypothetical protein, conserved | 6 | 3 | 7 | 6 | 3 | 5 | 4 | 7 |
| 9 | TGME49_500134.P436 | RNA binding protein, putative | 8 | 3 | 6 | 4 | 3 | 5 | 5 | 7 |
| 9 | TGME49_500139.P441 | EF-hand domain-containing protein | 9 | 3 | 8 | 3 | 7 | 3 | 5 | 7 |
| 9 | TGME49_500147.P151 | nucleoside transporter, putative | 10 | 3 | 8 | 3 | 3 | 3 | 3 | 7 |
| 26 | TGME49_500150.P154 | hypothetical protein, conserved | 6 | 3 | 4 | 3 | 3 | 4 | 3 | 7 |
| 26 | TGME49_500151.P155 | hypothetical protein, conserved | 7 | 3 | 4 | 3 | 3 | 3 | 3 | 7 |
| 26 | TGME49_500152.P156 | SAG domain-containing protein | 7 | 3 | 5 | 4 | 3 | 3 | 3 | 7 |
| 26 | TGME49_500156.P160 | hypothetical protein | 7 | 3 | 6 | 3 | 3 | 3 | 3 | 7 |
| 26 | TGME49_500161.P165 | hypothetical protein, conserved | 5 | 3 | 4 | 3 | 4 | 3 | 4 | 7 |
| 26 | TGME49_500162.P166 | SSNA1 domain-containing protein | 6 | 3 | 5 | 5 | 3 | 3 | 3 | 7 |
| 20 | TGME49_500166.P170 | hypothetical protein, conserved | 9 | 3 | 3 | 3 | 3 | 3 | 3 | 7 |
| 26 | TGME49_500170.P174 | hypothetical protein, conserved | 5 | 3 | 3 | 3 | 3 | 3 | 3 | 7 |
| 24 | TGME49_500171.P175 | lipocalin, putative | 7 | 6 | 3 | 3 | 3 | 3 | 3 | 7 |
| 26 | TGME49_500172.P176 | hypothetical protein, conserved | 7 | 3 | 3 | 3 | 3 | 3 | 3 | 7 |
| 26 | TGME49_500174.P178 | KRUF family protein | 6 | 3 | 4 | 3 | 3 | 3 | 3 | 7 |
| 20 | TGME49_500176.P180 | hypothetical protein, conserved | 8 | 3 | 5 | 3 | 3 | 3 | 3 | 7 |
| 20 | TGME49_500177.P181 | hypothetical protein, conserved | 9 | 4 | 3 | 4 | 3 | 3 | 3 | 7 |
| 20 | TGME49_500187.P191 | hypothetical protein, conserved | 9 | 3 | 5 | 3 | 3 | 3 | 3 | 7 |
| 20 | TGME49_500192.P196 | hypothetical protein, conserved | 10 | 5 | 3 | 3 | 3 | 3 | 3 | 7 |
| 9 | TGME49_500193.P197 | chromo domain-containing protein | 9 | 3 | 6 | 4 | 4 | 4 | 4 | 7 |
| 9 | TGME49_500198.P202 | hypothetical protein, conserved | 9 | 3 | 6 | 4 | 4 | 3 | 3 | 7 |
| 24 | TGME49_500199.P203 | SAG-related sequence | 7 | 6 | 5 | 3 | 3 | 3 | 3 | 7 |
| 9 | TGME49_500216.P221 | hypothetical protein, conserved | 9 | 3 | 7 | 5 | 4 | 3 | 3 | 7 |
| 26 | TGME49_500230.P236 | hypothetical protein, conserved | 7 | 3 | 4 | 4 | 3 | 4 | 4 | 7 |
| 9 | TGME49_500235.P241 | hypothetical protein, conserved | 9 | 3 | 6 | 5 | 4 | 3 | 4 | 7 |
| 20 | TGME49_500248.P254 | hypothetical protein, conserved | 9 | 3 | 3 | 3 | 3 | 3 | 3 | 7 |
| 16 | TGME49_500282.P71 | thioredoxin, putative | 6 | 3 | 5 | 5 | 3 | 3 | 4 | 7 |
| 20 | TGME49_500290.P79 | hypothetical protein, conserved | 10 | 4 | 3 | 3 | 3 | 3 | 3 | 7 |
| 20 | TGME49_500296.P85 | KRUF family protein | 9 | 5 | 5 | 3 | 3 | 3 | 3 | 7 |
| 26 | TGME49_500297.P86 | hypothetical protein, conserved | 7 | 3 | 3 | 3 | 3 | 3 | 3 | 7 |
| 9 | TGME49_500298.P87 | hypothetical protein, conserved | 9 | 3 | 7 | 3 | 4 | 3 | 3 | 7 |
| 24 | TGME49_500300.P89 | hypothetical protein, conserved | 7 | 6 | 5 | 3 | 3 | 3 | 3 | 7 |

| | | | | | | | | | | |
|---|---|---|---|---|---|---|---|---|---|---|
| 16 | TGME49_500304.P94 | P25-alpha family protein, putative | 6 | 5 | 5 | 5 | 3 | 5 | 5 | 7 |
| 20 | TGME49_500305.P95 | hypothetical protein, conserved | 8 | 3 | 4 | 3 | 3 | 3 | 3 | 7 |
| 9 | TGME49_500308.P98 | phosphatidate cytidylyltransferase, putative | 9 | 3 | 7 | 6 | 4 | 4 | 3 | 7 |
| 20 | TGME49_500315.P105 | hypothetical protein, conserved | 9 | 3 | 5 | 4 | 3 | 3 | 3 | 7 |
| 16 | TGME49_500318.P108 | zinc finger (CCCH type) motif-containing protein | 3 | 3 | 7 | 7 | 3 | 5 | 5 | 7 |
| 26 | TGME49_500328.P118 | hypothetical protein, conserved | 7 | 3 | 3 | 3 | 3 | 3 | 3 | 7 |
| 9 | TGME49_500330.P120 | hypothetical protein, conserved | 10 | 5 | 6 | 6 | 4 | 3 | 3 | 7 |
| 26 | TGME49_500331.P121 | hypothetical protein, conserved | 7 | 3 | 4 | 3 | 3 | 3 | 3 | 7 |
| 20 | TGME49_500338.P128 | hypothetical protein, conserved | 8 | 4 | 3 | 3 | 3 | 3 | 3 | 7 |
| 16 | TGME49_500349.P139 | hypothetical protein | 3 | 3 | 6 | 4 | 4 | 4 | 4 | 7 |
| 24 | TGME49_500353.P143 | SAG-related sequence | 7 | 6 | 4 | 3 | 3 | 3 | 3 | 7 |
| 26 | TGME49_500354.P144 | hypothetical protein, conserved | 5 | 4 | 3 | 3 | 3 | 3 | 3 | 7 |
| 20 | TGME49_500356.P146 | hypothetical protein, conserved | 9 | 3 | 4 | 3 | 3 | 3 | 3 | 7 |
| 9 | TGME49_500367.P266 | hypothetical protein, conserved | 8 | 3 | 3 | 8 | 3 | 4 | 4 | 7 |
| 16 | TGME49_500369.P268 | hypothetical protein, conserved | 3 | 3 | 6 | 6 | 3 | 5 | 4 | 7 |
| 24 | TGME49_500373.P272 | MORN repeat-containing protein | 7 | 4 | 7 | 3 | 3 | 3 | 3 | 7 |
| 20 | TGME49_500377.P276 | hypothetical protein, conserved | 8 | 3 | 3 | 3 | 3 | 3 | 3 | 7 |
| 26 | TGME49_500378.P277 | hypothetical protein, conserved | 5 | 3 | 4 | 4 | 3 | 3 | 3 | 7 |
| 20 | TGME49_500390.P289 | hypothetical protein, conserved | 9 | 3 | 5 | 4 | 3 | 3 | 3 | 7 |
| 26 | TGME49_500406.P305 | succinate dehydrogenase assembly factor 3, putative | 7 | 3 | 5 | 4 | 3 | 3 | 4 | 7 |
| 20 | TGME49_500409.P308 | hypothetical protein, conserved | 8 | 3 | 3 | 3 | 3 | 3 | 3 | 7 |
| 16 | TGME49_500410.P309 | hypothetical protein, conserved | 3 | 3 | 6 | 6 | 5 | 5 | 5 | 7 |
| 26 | TGME49_500413.P312 | hypothetical protein, conserved | 5 | 3 | 4 | 3 | 3 | 3 | 3 | 7 |
| 20 | TGME49_500414.P313 | hypothetical protein, conserved | 10 | 3 | 4 | 3 | 3 | 3 | 3 | 7 |
| 26 | TGME49_500415.P314 | Myb domain-containing protein | 5 | 3 | 3 | 3 | 3 | 3 | 3 | 7 |
| 20 | TGME49_500418.P317 | hypothetical protein, conserved | 9 | 3 | 5 | 4 | 3 | 3 | 3 | 7 |
| 26 | TGME49_500419.P318 | hypothetical protein | 5 | 3 | 3 | 3 | 3 | 3 | 3 | 7 |
| 9 | TGME49_500424.P323 | radial spoke head protein 3, putative | 10 | 3 | 6 | 4 | 3 | 3 | 3 | 7 |
| 26 | TGME49_500426.P325 | hypothetical protein, conserved | 5 | 3 | 5 | 4 | 3 | 3 | 3 | 7 |
| 20 | TGME49_500427.P326 | hypothetical protein, conserved | 8 | 4 | 4 | 3 | 3 | 4 | 4 | 7 |
| 26 | TGME49_500428.P327 | hypothetical protein, conserved | 5 | 3 | 3 | 3 | 3 | 3 | 3 | 7 |
| 20 | TGME49_500434.P333 | hypothetical protein, conserved | 10 | 3 | 4 | 3 | 3 | 3 | 3 | 7 |
| 26 | TGME49_500440.P339 | hypothetical protein, conserved | 7 | 3 | 3 | 3 | 3 | 4 | 3 | 7 |

| Kmean_cluster | GeneID (all protein TG) | Product Description | Decile_bitHH | Decile_bitNcan | Decile_bitBB | Decilebit_CS | Decilebit_SN3 | Decile bit_Ef | Decile bit_CC | Meta_cluster |
|---|---|---|---|---|---|---|---|---|---|---|
| 12 | TGME49_200110-t26_1-p1 | hypothetical protein | 3 | 3 | 3 | 3 | 3 | 3 | 3 | 8 |
| 12 | TGME49_200130-t26_1-p1 | Toxoplasma gondii family C protein | 3 | 3 | 3 | 3 | 3 | 3 | 3 | 8 |
| 12 | TGME49_200230-t26_1-p1 | microneme protein MIC17C | 3 | 3 | 3 | 3 | 3 | 3 | 3 | 8 |
| 12 | TGME49_200250-t26_1-p1 | microneme protein MIC17A | 3 | 3 | 7 | 3 | 3 | 3 | 3 | 8 |
| 12 | TGME49_200260-t26_1-p1 | hypothetical protein | 3 | 3 | 3 | 3 | 3 | 3 | 3 | 8 |
| 12 | TGME49_200320.P449 | hypoxanthine-xanthine-guanine phosphoribosyl transferase HXGPRT | 3 | 3 | 3 | 3 | 3 | 3 | 3 | 8 |
| 12 | TGME49_200320.P450 | hypoxanthine-xanthine-guanine phosphoribosyl transferase HXGPRT | 3 | 3 | 3 | 3 | 3 | 3 | 3 | 8 |
| 12 | TGME49_200480-t26_1-p1 | Toxoplasma gondii family B protein | 3 | 4 | 3 | 3 | 3 | 3 | 3 | 8 |
| 12 | TGME49_200595-t26_1-p1 | hypothetical protein | 4 | 3 | 3 | 3 | 3 | 3 | 3 | 8 |
| 12 | TGME49_200700-t26_1-p1 | Toxoplasma gondii family C protein | 3 | 3 | 3 | 3 | 3 | 3 | 3 | 8 |
| 12 | TGME49_201100-t26_1-p1 | hypothetical protein | 3 | 5 | 4 | 3 | 3 | 3 | 3 | 8 |
| 12 | TGME49_201110.P464 | hypothetical protein | 3 | 5 | 4 | 3 | 4 | 3 | 4 | 8 |
| 12 | TGME49_201160-t26_1-p1 | hypothetical protein | 3 | 3 | 3 | 3 | 3 | 3 | 3 | 8 |
| 12 | TGME49_201390-t26_1-p1 | hypothetical protein | 4 | 3 | 3 | 3 | 3 | 3 | 3 | 8 |
| 22 | TGME49_201520-t26_1-p1 | protein phosphatase PPM3G, putative | 3 | 7 | 3 | 3 | 3 | 3 | 3 | 8 |
| 12 | TGME49_201630-t26_1-p1 | protein phosphatase PPM3H, putative | 3 | 3 | 6 | 3 | 3 | 3 | 3 | 8 |
| 12 | TGME49_201775-t26_1-p1 | hypothetical protein | 3 | 3 | 3 | 3 | 3 | 3 | 3 | 8 |
| 12 | TGME49_201785-t26_1-p1 | hypothetical protein | 3 | 3 | 3 | 3 | 3 | 3 | 3 | 8 |
| 12 | TGME49_202065-t26_1-p1 | hypothetical protein | 3 | 3 | 3 | 3 | 3 | 3 | 3 | 8 |
| 12 | TGME49_202330-t26_1-p1 | hypothetical protein | 3 | 3 | 3 | 3 | 3 | 3 | 3 | 8 |
| 12 | TGME49_202375-t26_1-p1 | hypothetical protein | 3 | 3 | 3 | 3 | 3 | 3 | 3 | 8 |
| 12 | TGME49_202445-t26_1-p1 | hypothetical protein | 3 | 3 | 3 | 3 | 3 | 3 | 3 | 8 |
| 12 | TGME49_202735-t26_1-p1 | hypothetical protein | 3 | 3 | 3 | 3 | 3 | 3 | 3 | 8 |
| 12 | TGME49_202860-t26_1-p1 | hypothetical protein | 3 | 3 | 3 | 3 | 3 | 3 | 3 | 8 |
| 12 | TGME49_203100-t26_1-p1 | hypothetical protein | 3 | 3 | 3 | 3 | 3 | 3 | 3 | 8 |
| 12 | TGME49_203270-t26_1-p1 | ATP-dependent Clp endopeptidase, proteolytic subunit ClpP domain-containing protein | 3 | 4 | 5 | 4 | 3 | 4 | 4 | 8 |
| 12 | TGME49_203470-t26_1-p1 | hypothetical protein | 3 | 3 | 3 | 3 | 3 | 3 | 3 | 8 |
| 12 | TGME49_203685-t26_1-p1 | hypothetical protein | 3 | 3 | 3 | 3 | 3 | 3 | 3 | 8 |
| 12 | TGME49_203688-t26_1-p1 | hypothetical protein | 3 | 3 | 3 | 3 | 3 | 3 | 3 | 8 |
| 12 | TGME49_203705-t26_1-p1 | hypothetical protein | 3 | 3 | 3 | 3 | 3 | 3 | 3 | 8 |
| 12 | TGME49_203815-t26_1-p1 | hypothetical protein | 3 | 3 | 3 | 3 | 3 | 3 | 3 | 8 |
| 12 | TGME49_203985-t26_1-p1 | hypothetical protein | 3 | 3 | 3 | 3 | 3 | 3 | 3 | 8 |
| 12 | TGME49_204045-t26_1-p1 | hypothetical protein | 3 | 3 | 3 | 3 | 3 | 3 | 3 | 8 |
| 12 | TGME49_204090-t26_1-p1 | eIF2 kinase IF2K-C | 3 | 3 | 3 | 3 | 3 | 3 | 3 | 8 |
| 12 | TGME49_204110-t26_1-p1 | eIF2 kinase IF2K-C | 3 | 3 | 3 | 3 | 3 | 3 | 3 | 8 |
| 12 | TGME49_204325-t26_1-p1 | hypothetical protein | 3 | 3 | 4 | 3 | 3 | 3 | 3 | 8 |
| 12 | TGME49_204395-t26_1-p1 | hypothetical protein | 3 | 3 | 3 | 3 | 3 | 3 | 3 | 8 |
| 12 | TGME49_204505-t26_1-p1 | hypothetical protein | 3 | 3 | 3 | 3 | 3 | 3 | 3 | 8 |
| 12 | TGME49_205230-t26_1-p1 | hypothetical protein | 3 | 3 | 3 | 3 | 3 | 3 | 3 | 8 |
| 22 | TGME49_205402-t26_1-p1 | SNF2 family N-terminal domain-containing protein | 3 | 9 | 8 | 7 | 5 | 3 | 4 | 8 |
| 12 | TGME49_205425-t26_1-p1 | hypothetical protein | 3 | 3 | 3 | 3 | 3 | 3 | 3 | 8 |
| 12 | TGME49_205530-t26_1-p1 | hypothetical protein | 3 | 3 | 3 | 3 | 3 | 3 | 3 | 8 |
| 12 | TGME49_205630-t26_1-p1 | hypothetical protein | 3 | 3 | 3 | 3 | 3 | 3 | 3 | 8 |
| 12 | TGME49_205770-t26_1-p1 | hypothetical protein | 3 | 3 | 3 | 3 | 3 | 3 | 3 | 8 |
| 12 | TGME49_206460-t26_1-p1 | hypothetical protein | 3 | 3 | 3 | 3 | 3 | 3 | 3 | 8 |
| 12 | TGME49_206520-t26_1-p1 | hypothetical protein | 3 | 3 | 3 | 3 | 3 | 3 | 3 | 8 |

| | | | | | | | | | | |
|---|---|---|---|---|---|---|---|---|---|---|
| 22 | TGME49_206660-t26_1-p1 | hypothetical protein | 3 | 7 | 3 | 3 | 3 | 3 | 3 | 8 |
| 12 | TGME49_206910-t26_1-p1 | hypothetical protein | 3 | 3 | 3 | 3 | 3 | 3 | 3 | 8 |
| 12 | TGME49_207005.P598 | SAG-related sequence SRS48Q | 3 | 3 | 4 | 3 | 3 | 3 | 3 | 8 |
| 12 | TGME49_207065-t26_1-p1 | hypothetical protein | 3 | 3 | 3 | 3 | 3 | 3 | 3 | 8 |
| 12 | TGME49_207130.P606 | SAG-related sequence SRS49A | 3 | 3 | 3 | 4 | 3 | 3 | 3 | 8 |
| 12 | TGME49_207150.P608 | SAG-related sequence SRS49C | 3 | 3 | 3 | 3 | 3 | 3 | 3 | 8 |
| 22 | TGME49_207350.P612 | hypothetical protein | 5 | 7 | 7 | 3 | 3 | 3 | 3 | 8 |
| 12 | TGME49_207403-t26_1-p1 | hypothetical protein | 4 | 3 | 3 | 3 | 3 | 3 | 3 | 8 |
| 22 | TGME49_207780-t26_1-p1 | hypothetical protein | 4 | 6 | 3 | 3 | 3 | 3 | 3 | 8 |
| 12 | TGME49_208450.P645 | protease inhibitor PI2 | 3 | 3 | 5 | 4 | 3 | 3 | 3 | 8 |
| 12 | TGME49_208722-t26_1-p1 | hypothetical protein | 3 | 3 | 3 | 3 | 3 | 3 | 3 | 8 |
| 12 | TGME49_209005-t26_1-p1 | hypothetical protein | 3 | 3 | 3 | 3 | 3 | 3 | 3 | 8 |
| 12 | TGME49_209095-t26_1-p1 | hypothetical protein | 3 | 3 | 3 | 3 | 3 | 3 | 3 | 8 |
| 12 | TGME49_209780-t26_1-p1 | hypothetical protein | 3 | 3 | 3 | 3 | 3 | 3 | 3 | 8 |
| 12 | TGME49_209945.P700 | hypothetical protein | 4 | 3 | 4 | 4 | 3 | 3 | 3 | 8 |
| 22 | TGME49_209985-t26_1-p1 | rhoptry protein ROP42 | 3 | 8 | 7 | 3 | 3 | 3 | 3 | 8 |
| 12 | TGME49_210095.P703 | rhoptry protein ROP43 | 3 | 3 | 3 | 3 | 3 | 3 | 3 | 8 |
| 12 | TGME49_210100-t26_1-p1 | hypothetical protein | 3 | 3 | 3 | 3 | 3 | 3 | 3 | 8 |
| 12 | TGME49_210255.P706 | mitochondrial import receptor subunit TOM7 | 3 | 3 | 3 | 3 | 4 | 3 | 3 | 8 |
| 22 | TGME49_210320-t26_1-p1 | SAG-related sequence SRS37A | 3 | 6 | 5 | 4 | 3 | 3 | 3 | 8 |
| 12 | TGME49_210682-t26_1-p1 | hypothetical protein | 4 | 3 | 3 | 3 | 3 | 3 | 3 | 8 |
| 12 | TGME49_210715-t26_1-p1 | hypothetical protein | 3 | 3 | 3 | 3 | 3 | 3 | 3 | 8 |
| 12 | TGME49_210783-t26_1-p1 | hypothetical protein | 3 | 3 | 3 | 3 | 3 | 3 | 3 | 8 |
| 12 | TGME49_210815-t26_1-p1 | hypothetical protein | 3 | 3 | 3 | 3 | 3 | 3 | 3 | 8 |
| 12 | TGME49_210970-t26_1-p1 | hypothetical protein | 3 | 3 | 3 | 3 | 3 | 3 | 3 | 8 |
| 12 | TGME49_210975-t26_1-p1 | hypothetical protein | 3 | 3 | 3 | 3 | 3 | 3 | 3 | 8 |
| 12 | TGME49_211080-t26_1-p1 | hypothetical protein | 3 | 3 | 3 | 3 | 3 | 3 | 3 | 8 |
| 12 | TGME49_211420.P739 | serine/arginine-rich splicing factor SR3 | 3 | 3 | 5 | 3 | 3 | 3 | 3 | 8 |
| 12 | TGME49_211695-t26_1-p1 | hypothetical protein | 4 | 3 | 4 | 3 | 3 | 3 | 3 | 8 |
| 12 | TGME49_211920-t26_1-p1 | hypothetical protein | 3 | 3 | 3 | 3 | 3 | 3 | 3 | 8 |
| 12 | TGME49_212280-t26_1-p1 | hypothetical protein | 3 | 3 | 3 | 3 | 3 | 3 | 3 | 8 |
| 12 | TGME49_212420-t26_1-p1 | hypothetical protein | 3 | 3 | 3 | 3 | 3 | 3 | 3 | 8 |
| 12 | TGME49_212775-t26_1-p1 | hypothetical protein | 3 | 3 | 3 | 3 | 3 | 3 | 3 | 8 |
| 12 | TGME49_213065-t26_1-p1 | hypothetical protein | 3 | 3 | 3 | 3 | 3 | 3 | 3 | 8 |
| 12 | TGME49_213290.P786 | hypothetical protein | 3 | 5 | 3 | 3 | 3 | 3 | 3 | 8 |
| 12 | TGME49_213380-t26_1-p1 | hypothetical protein | 3 | 3 | 3 | 3 | 3 | 3 | 3 | 8 |
| 12 | TGME49_213415-t26_1-p1 | hypothetical protein | 3 | 3 | 3 | 3 | 3 | 3 | 3 | 8 |
| 12 | TGME49_213580.P796 | ribosomal protein L41 | 3 | 4 | 3 | 3 | 3 | 3 | 3 | 8 |
| 22 | TGME49_213610.P800 | hypothetical protein | 3 | 9 | 6 | 6 | 3 | 3 | 3 | 8 |
| 12 | TGME49_213635-t26_1-p1 | hypothetical protein | 3 | 3 | 3 | 3 | 3 | 3 | 3 | 8 |
| 12 | TGME49_213748-t26_1-p1 | hypothetical protein | 3 | 3 | 3 | 3 | 3 | 3 | 3 | 8 |
| 12 | TGME49_214070-t26_1-p1 | hypothetical protein | 4 | 3 | 3 | 3 | 3 | 3 | 3 | 8 |
| 12 | TGME49_214100-t26_1-p1 | hypothetical protein | 3 | 3 | 3 | 3 | 3 | 3 | 3 | 8 |
| 12 | TGME49_214115-t26_1-p1 | hypothetical protein | 3 | 3 | 3 | 3 | 3 | 3 | 3 | 8 |
| 12 | TGME49_214160-t26_1-p1 | hypothetical protein | 3 | 3 | 3 | 3 | 3 | 3 | 3 | 8 |
| 12 | TGME49_214295-t26_1-p1 | hypothetical protein | 3 | 3 | 3 | 3 | 3 | 3 | 3 | 8 |
| 12 | TGME49_214385.P823 | hypothetical protein | 4 | 4 | 4 | 4 | 4 | 4 | 3 | 8 |
| 12 | TGME49_214400-t26_1-p1 | hypothetical protein | 3 | 3 | 3 | 3 | 3 | 3 | 3 | 8 |
| 12 | TGME49_214783-t26_1-p1 | hypothetical protein | 3 | 3 | 3 | 3 | 3 | 3 | 3 | 8 |
| 12 | TGME49_214787-t26_1-p1 | hypothetical protein | 3 | 3 | 3 | 3 | 3 | 3 | 3 | 8 |

| | | | | | | | | | | |
|---|---|---|---|---|---|---|---|---|---|---|
| 12 | TGME49_214835-t26_1-p1 | hypothetical protein | 4 | 3 | 3 | 3 | 3 | 3 | 3 | 8 |
| 12 | TGME49_214860-t26_1-p1 | hypothetical protein | 3 | 3 | 3 | 3 | 3 | 3 | 3 | 8 |
| 12 | TGME49_215055-t26_1-p1 | hypothetical protein | 3 | 3 | 3 | 3 | 3 | 3 | 3 | 8 |
| 12 | TGME49_215270-t26_1-p1 | hypothetical protein | 3 | 3 | 3 | 3 | 3 | 3 | 3 | 8 |
| 12 | TGME49_215650-t26_1-p1 | hypothetical protein | 3 | 3 | 3 | 3 | 3 | 3 | 3 | 8 |
| 12 | TGME49_215680.P865 | SAG-related sequence SRS18 | 3 | 5 | 3 | 3 | 3 | 3 | 3 | 8 |
| 12 | TGME49_215785-t26_1-p1 | rhoptry protein ROP2A | 3 | 3 | 3 | 3 | 3 | 3 | 3 | 8 |
| 12 | TGME49_215885-t26_1-p1 | hypothetical protein | 3 | 3 | 3 | 3 | 3 | 3 | 3 | 8 |
| 12 | TGME49_215955-t26_1-p1 | hypothetical protein | 3 | 3 | 3 | 3 | 3 | 3 | 3 | 8 |
| 12 | TGME49_215960-t26_1-p1 | hypothetical protein | 3 | 3 | 3 | 3 | 3 | 3 | 3 | 8 |
| 12 | TGME49_215970-t26_1-p1 | hypothetical protein | 3 | 3 | 3 | 3 | 3 | 3 | 3 | 8 |
| 12 | TGME49_216200-t26_1-p1 | hypothetical protein | 3 | 3 | 3 | 3 | 3 | 3 | 3 | 8 |
| 12 | TGME49_216375-t26_1-p1 | hypothetical protein | 3 | 3 | 5 | 4 | 3 | 3 | 3 | 8 |
| 12 | TGME49_216415-t26_1-p1 | hypothetical protein | 3 | 3 | 3 | 3 | 3 | 3 | 3 | 8 |
| 12 | TGME49_216420-t26_1-p1 | hypothetical protein | 4 | 3 | 3 | 3 | 3 | 3 | 3 | 8 |
| 12 | TGME49_216585-t26_1-p1 | hypothetical protein | 3 | 3 | 3 | 3 | 3 | 3 | 3 | 8 |
| 12 | TGME49_216785-t26_1-p1 | hypothetical protein | 3 | 3 | 3 | 3 | 3 | 3 | 3 | 8 |
| 12 | TGME49_216875-t26_1-p1 | cpw-wpc domain-containing protein | 3 | 3 | 3 | 3 | 3 | 4 | 4 | 8 |
| 22 | TGME49_217660.P916 | Tctex-1 family protein | 3 | 6 | 5 | 3 | 3 | 4 | 3 | 8 |
| 12 | TGME49_217722-t26_1-p1 | hypothetical protein | 3 | 3 | 3 | 3 | 3 | 3 | 3 | 8 |
| 12 | TGME49_217724-t26_1-p1 | hypothetical protein | 3 | 3 | 3 | 3 | 3 | 3 | 3 | 8 |
| 12 | TGME49_217726-t26_1-p1 | hypothetical protein | 3 | 3 | 3 | 3 | 3 | 3 | 3 | 8 |
| 12 | TGME49_217728-t26_1-p1 | hypothetical protein | 3 | 3 | 3 | 3 | 3 | 3 | 3 | 8 |
| 12 | TGME49_217810-t26_1-p1 | hypothetical protein | 3 | 3 | 3 | 3 | 3 | 3 | 3 | 8 |
| 12 | TGME49_217855-t26_1-p1 | hypothetical protein | 4 | 3 | 3 | 3 | 3 | 3 | 3 | 8 |
| 12 | TGME49_217875-t26_1-p1 | hypothetical protein | 3 | 3 | 3 | 3 | 3 | 3 | 3 | 8 |
| 12 | TGME49_217940.P929 | hypothetical protein | 3 | 3 | 7 | 3 | 3 | 3 | 3 | 8 |
| 12 | TGME49_217951-t26_1-p1 | hypothetical protein | 3 | 3 | 3 | 3 | 3 | 3 | 3 | 8 |
| 22 | TGME49_217961-t26_1-p1 | hypothetical protein | 4 | 8 | 3 | 6 | 3 | 3 | 3 | 8 |
| 12 | TGME49_218130-t26_1-p1 | hypothetical protein | 3 | 3 | 3 | 3 | 3 | 3 | 3 | 8 |
| 12 | TGME49_218350-t26_1-p1 | hypothetical protein | 3 | 3 | 3 | 3 | 3 | 3 | 3 | 8 |
| 12 | TGME49_218460-t26_1-p1 | hypothetical protein | 3 | 3 | 6 | 3 | 5 | 3 | 3 | 8 |
| 12 | TGME49_219090-t26_1-p1 | hypothetical protein | 3 | 3 | 3 | 3 | 3 | 3 | 3 | 8 |
| 12 | TGME49_219570-t26_1-p1 | hypothetical protein | 3 | 3 | 3 | 3 | 3 | 3 | 3 | 8 |
| 12 | TGME49_219780-t26_1-p1 | hypothetical protein | 3 | 3 | 3 | 3 | 3 | 3 | 3 | 8 |
| 12 | TGME49_219840-t26_1-p1 | hypothetical protein | 3 | 3 | 3 | 3 | 3 | 3 | 3 | 8 |
| 12 | TGME49_220060-t26_1-p1 | hypothetical protein | 3 | 3 | 3 | 3 | 3 | 3 | 3 | 8 |
| 12 | TGME49_220070-t26_1-p1 | hypothetical protein | 3 | 3 | 3 | 3 | 3 | 3 | 3 | 8 |
| 12 | TGME49_220145-t26_1-p1 | hypothetical protein | 3 | 3 | 3 | 3 | 3 | 3 | 3 | 8 |
| 12 | TGME49_220180-t26_1-p1 | hypothetical protein | 3 | 3 | 3 | 3 | 3 | 3 | 3 | 8 |
| 12 | TGME49_220310-t26_1-p1 | hypothetical protein | 3 | 3 | 3 | 3 | 3 | 3 | 3 | 8 |
| 12 | TGME49_220525-t26_1-p1 | hypothetical protein | 3 | 3 | 3 | 3 | 3 | 3 | 3 | 8 |
| 12 | TGME49_220530-t26_1-p1 | AP2 domain transcription factor AP2V-1 | 3 | 3 | 3 | 3 | 3 | 3 | 3 | 8 |
| 12 | TGME49_220540-t26_1-p1 | hypothetical protein | 3 | 3 | 3 | 3 | 3 | 3 | 3 | 8 |
| 12 | TGME49_220640-t26_1-p1 | hypothetical protein | 3 | 3 | 3 | 3 | 3 | 3 | 3 | 8 |
| 12 | TGME49_220720-t26_1-p1 | hypothetical protein | 3 | 3 | 3 | 3 | 3 | 3 | 3 | 8 |
| 12 | TGME49_220730-t26_1-p1 | hypothetical protein | 3 | 3 | 3 | 3 | 3 | 3 | 3 | 8 |
| 12 | TGME49_220740-t26_1-p1 | hypothetical protein | 3 | 3 | 3 | 3 | 3 | 3 | 3 | 8 |
| 12 | TGME49_220850-t26_1-p1 | hypothetical protein | 3 | 3 | 3 | 3 | 3 | 3 | 3 | 8 |
| 12 | TGME49_221240-t26_1-p1 | hypothetical protein | 3 | 3 | 3 | 3 | 3 | 3 | 3 | 8 |

| | | | | | | | | | | |
|---|---|---|---|---|---|---|---|---|---|---|
| 12 | TGME49_221275-t26_1-p1 | hypothetical protein | 3 | 3 | 3 | 3 | 3 | 3 | 3 | 8 |
| 12 | TGME49_221415-t26_1-p1 | hypothetical protein | 3 | 3 | 3 | 3 | 3 | 3 | 3 | 8 |
| 12 | TGME49_221455-t26_1-p1 | hypothetical protein | 3 | 3 | 3 | 3 | 3 | 3 | 3 | 8 |
| 12 | TGME49_221855-t26_1-p1 | hypothetical protein | 3 | 3 | 3 | 3 | 3 | 3 | 3 | 8 |
| 12 | TGME49_221860-t26_1-p1 | hypothetical protein | 3 | 3 | 3 | 3 | 3 | 3 | 3 | 8 |
| 12 | TGME49_221910.P1040 | AN1 family Zinc finger domain-containing protein | 3 | 3 | 3 | 3 | 3 | 3 | 3 | 8 |
| 12 | TGME49_222075-t26_1-p1 | hypothetical protein | 3 | 3 | 3 | 3 | 3 | 3 | 3 | 8 |
| 12 | TGME49_222305-t26_1-p1 | hypothetical protein | 3 | 3 | 3 | 3 | 3 | 3 | 3 | 8 |
| 12 | TGME49_222948-t26_1-p1 | hypothetical protein | 3 | 3 | 3 | 3 | 3 | 3 | 3 | 8 |
| 12 | TGME49_223025-t26_1-p1 | hypothetical protein | 3 | 3 | 3 | 3 | 3 | 3 | 3 | 8 |
| 12 | TGME49_223045-t26_1-p1 | hypothetical protein | 3 | 3 | 3 | 3 | 3 | 3 | 3 | 8 |
| 12 | TGME49_223470-t26_1-p1 | hypothetical protein | 3 | 3 | 3 | 3 | 3 | 3 | 3 | 8 |
| 22 | TGME49_223845.P1094 | hypothetical protein | 3 | 8 | 6 | 5 | 3 | 3 | 3 | 8 |
| 12 | TGME49_224100-t26_1-p1 | hypothetical protein | 3 | 3 | 3 | 3 | 3 | 3 | 3 | 8 |
| 12 | TGME49_224560-t26_1-p1 | hypothetical protein | 3 | 3 | 3 | 3 | 3 | 3 | 3 | 8 |
| 12 | TGME49_224575-t26_1-p1 | hypothetical protein | 3 | 3 | 3 | 3 | 3 | 3 | 3 | 8 |
| 12 | TGME49_224780-t26_1-p1 | SAG-related sequence SRS40B [Source:UniProtKB/TrEMBL;Acc:S8EU84] | 3 | 3 | 3 | 3 | 3 | 3 | 3 | 8 |
| 12 | TGME49_225005-t26_1-p1 | hypothetical protein | 3 | 3 | 3 | 3 | 3 | 3 | 3 | 8 |
| 12 | TGME49_225435-t26_1-p1 | hypothetical protein | 3 | 3 | 3 | 3 | 3 | 3 | 3 | 8 |
| 12 | TGME49_225953.P1188 | hypothetical protein | 3 | 3 | 3 | 3 | 3 | 3 | 3 | 8 |
| 12 | TGME49_225958-t26_1-p1 | hypothetical protein | 3 | 3 | 3 | 3 | 3 | 3 | 3 | 8 |
| 12 | TGME49_226565-t26_1-p1 | hypothetical protein | 3 | 3 | 3 | 3 | 3 | 3 | 3 | 8 |
| 12 | TGME49_226820.P1224 | hypothetical protein | 3 | 3 | 3 | 3 | 3 | 3 | 3 | 8 |
| 12 | TGME49_227040-t26_1-p1 | hypothetical protein | 3 | 3 | 3 | 3 | 3 | 3 | 3 | 8 |
| 12 | TGME49_227130-t26_1-p1 | hypothetical protein | 3 | 3 | 3 | 3 | 3 | 3 | 3 | 8 |
| 12 | TGME49_227335-t26_1-p1 | hypothetical protein | 3 | 3 | 3 | 3 | 3 | 3 | 3 | 8 |
| 12 | TGME49_227615-t26_1-p1 | hypothetical protein | 3 | 3 | 3 | 3 | 3 | 3 | 3 | 8 |
| 12 | TGME49_227940-t26_1-p1 | hypothetical protein | 3 | 3 | 3 | 3 | 3 | 3 | 3 | 8 |
| 12 | TGME49_228020-t26_1-p1 | hypothetical protein | 3 | 3 | 3 | 3 | 3 | 3 | 3 | 8 |
| 12 | TGME49_228065-t26_1-p1 | hypothetical protein | 3 | 3 | 3 | 3 | 3 | 3 | 3 | 8 |
| 12 | TGME49_228145.P1276 | hypothetical protein | 3 | 3 | 3 | 3 | 3 | 3 | 3 | 8 |
| 12 | TGME49_228370.P1291 | hypothetical protein | 3 | 3 | 3 | 5 | 3 | 3 | 3 | 8 |
| 22 | TGME49_228395-t26_1-p1 | hypothetical protein | 3 | 10 | 3 | 3 | 3 | 3 | 3 | 8 |
| 12 | TGME49_228480-t26_1-p1 | hypothetical protein | 3 | 3 | 3 | 3 | 3 | 3 | 3 | 8 |
| 12 | TGME49_228640-t26_1-p1 | hypothetical protein | 3 | 3 | 3 | 3 | 3 | 3 | 3 | 8 |
| 12 | TGME49_228780-t26_1-p1 | Toxoplasma gondii family C protein | 3 | 3 | 3 | 3 | 3 | 3 | 3 | 8 |
| 12 | TGME49_228980-t26_1-p1 | hypothetical protein | 3 | 3 | 3 | 3 | 3 | 3 | 3 | 8 |
| 12 | TGME49_229005-t26_1-p1 | hypothetical protein | 3 | 3 | 3 | 3 | 3 | 3 | 3 | 8 |
| 12 | TGME49_229190-t26_1-p1 | hypothetical protein | 3 | 3 | 3 | 3 | 3 | 3 | 3 | 8 |
| 12 | TGME49_229300-t26_1-p1 | hypothetical protein | 3 | 3 | 3 | 3 | 3 | 3 | 3 | 8 |
| 12 | TGME49_229660-t26_1-p1 | hypothetical protein | 3 | 3 | 3 | 3 | 3 | 3 | 3 | 8 |
| 12 | TGME49_230180.P1359 | dense granule protein GRA24 | 3 | 3 | 4 | 3 | 3 | 3 | 3 | 8 |
| 12 | TGME49_230205-t26_1-p1 | hypothetical protein | 3 | 3 | 3 | 3 | 3 | 3 | 3 | 8 |
| 12 | TGME49_230900-t26_1-p1 | hypothetical protein | 3 | 3 | 3 | 3 | 3 | 3 | 3 | 8 |
| 12 | TGME49_231175-t26_1-p1 | hypothetical protein | 3 | 3 | 3 | 3 | 3 | 3 | 3 | 8 |
| 12 | TGME49_231230-t26_1-p1 | hypothetical protein | 3 | 3 | 3 | 3 | 3 | 3 | 3 | 8 |
| 12 | TGME49_231450-t26_1-p1 | hypothetical protein | 3 | 3 | 3 | 3 | 3 | 3 | 3 | 8 |
| 12 | TGME49_231992-t26_1-p1 | hypothetical protein | 4 | 3 | 3 | 3 | 3 | 3 | 3 | 8 |
| 12 | TGME49_232085-t26_1-p1 | hypothetical protein | 3 | 3 | 3 | 3 | 3 | 3 | 3 | 8 |

| | | | | | | | | | | |
|---|---|---|---|---|---|---|---|---|---|---|
| 12 | TGME49_232110-t26_1-p1 | hypothetical protein | 3 | 3 | 3 | 3 | 3 | 3 | 3 | 8 |
| 12 | TGME49_232770-t26_1-p1 | hypothetical protein | 3 | 4 | 3 | 3 | 3 | 3 | 3 | 8 |
| 12 | TGME49_232955-t26_1-p1 | hypothetical protein | 3 | 3 | 3 | 3 | 3 | 3 | 3 | 8 |
| 12 | TGME49_232990-t26_1-p1 | hypothetical protein | 3 | 3 | 3 | 3 | 3 | 3 | 3 | 8 |
| 12 | TGME49_233925-t26_1-p1 | hypothetical protein | 4 | 3 | 3 | 3 | 3 | 3 | 3 | 8 |
| 22 | TGME49_233950-t26_1-p1 | hypothetical protein | 3 | 8 | 3 | 3 | 3 | 3 | 3 | 8 |
| 12 | TGME49_234060-t26_1-p1 | hypothetical protein | 3 | 5 | 3 | 3 | 3 | 3 | 3 | 8 |
| 12 | TGME49_234340-t26_1-p1 | hypothetical protein | 3 | 3 | 3 | 3 | 3 | 3 | 3 | 8 |
| 12 | TGME49_234375-t26_1-p1 | hypothetical protein | 3 | 3 | 3 | 3 | 3 | 3 | 3 | 8 |
| 12 | TGME49_234450.P1535 | ribosomal protein RPS15A | 3 | 3 | 3 | 3 | 3 | 3 | 3 | 8 |
| 12 | TGME49_234550-t26_1-p1 | ribosomal protein RPL29 | 4 | 4 | 4 | 3 | 3 | 3 | 3 | 8 |
| 12 | TGME49_234590-t26_1-p1 | hypothetical protein | 3 | 3 | 3 | 3 | 3 | 3 | 3 | 8 |
| 12 | TGME49_235430-t26_1-p1 | hypothetical protein | 3 | 3 | 3 | 3 | 3 | 3 | 3 | 8 |
| 12 | TGME49_235600.P1565 | SAG-related sequence SRS45 | 4 | 4 | 3 | 3 | 3 | 3 | 3 | 8 |
| 12 | TGME49_235635-t26_1-p1 | hypothetical protein | 3 | 3 | 3 | 3 | 3 | 3 | 3 | 8 |
| 22 | TGME49_235730.P1573 | hypothetical protein | 3 | 6 | 6 | 5 | 5 | 3 | 3 | 8 |
| 12 | TGME49_236150-t26_1-p1 | hypothetical protein | 3 | 3 | 3 | 3 | 3 | 3 | 3 | 8 |
| 12 | TGME49_236230-t26_1-p1 | hypothetical protein | 3 | 3 | 3 | 3 | 3 | 3 | 3 | 8 |
| 22 | TGME49_236280-t26_1-p1 | hypothetical protein | 3 | 7 | 7 | 6 | 4 | 5 | 4 | 8 |
| 12 | TGME49_236290-t26_1-p1 | hypothetical protein | 3 | 3 | 3 | 3 | 3 | 3 | 3 | 8 |
| 12 | TGME49_236635-t26_1-p1 | hypothetical protein | 3 | 3 | 3 | 3 | 3 | 3 | 3 | 8 |
| 12 | TGME49_236830-t26_1-p1 | hypothetical protein | 3 | 3 | 3 | 3 | 3 | 3 | 3 | 8 |
| 12 | TGME49_236880-t26_1-p1 | hypothetical protein | 3 | 3 | 3 | 3 | 3 | 3 | 3 | 8 |
| 12 | TGME49_237120-t26_1-p1 | hypothetical protein | 3 | 3 | 6 | 4 | 3 | 3 | 3 | 8 |
| 12 | TGME49_237130-t26_1-p1 | cytochrome b, putative | 3 | 3 | 5 | 3 | 3 | 3 | 3 | 8 |
| 12 | TGME49_237270-t26_1-p1 | hypothetical protein | 4 | 3 | 3 | 3 | 3 | 3 | 3 | 8 |
| 12 | TGME49_237595-t26_1-p1 | hypothetical protein | 3 | 3 | 3 | 3 | 3 | 3 | 3 | 8 |
| 12 | TGME49_237700-t26_1-p1 | ADP/ATP carrier protein, putative | 3 | 3 | 3 | 3 | 3 | 3 | 3 | 8 |
| 12 | TGME49_237800.P1634 | dense granule protein GRA11B | 3 | 3 | 3 | 3 | 3 | 3 | 3 | 8 |
| 12 | TGME49_237835-t26_1-p1 | hypothetical protein | 3 | 3 | 3 | 3 | 3 | 3 | 3 | 8 |
| 12 | TGME49_237893-t26_1-p1 | hypothetical protein | 3 | 3 | 3 | 3 | 3 | 3 | 3 | 8 |
| 12 | TGME49_237894-t26_1-p1 | OTU domain-containing protein OTUD1B | 3 | 3 | 5 | 3 | 3 | 3 | 3 | 8 |
| 12 | TGME49_237900-t26_1-p1 | OTU domain-containing protein OTUD1E | 3 | 3 | 3 | 3 | 3 | 3 | 3 | 8 |
| 12 | TGME49_238165-t26_1-p1 | hypothetical protein | 3 | 3 | 3 | 3 | 3 | 3 | 3 | 8 |
| 12 | TGME49_238450-t26_1-p1 | hypothetical protein | 3 | 3 | 3 | 3 | 3 | 3 | 3 | 8 |
| 12 | TGME49_238505-t26_1-p1 | hypothetical protein | 3 | 3 | 3 | 3 | 3 | 3 | 3 | 8 |
| 12 | TGME49_238530-t26_1-p1 | SRS domain-containing protein | 3 | 3 | 3 | 3 | 3 | 3 | 3 | 8 |
| 12 | TGME49_238860-t26_1-p1 | hypothetical protein | 3 | 3 | 3 | 3 | 3 | 3 | 3 | 8 |
| 12 | TGME49_239083-t26_1-p1 | hypothetical protein | 3 | 3 | 3 | 3 | 3 | 3 | 3 | 8 |
| 22 | TGME49_239730.P1683 | MORN repeat-containing protein | 3 | 6 | 5 | 4 | 3 | 3 | 3 | 8 |
| 12 | TGME49_240260-t26_1-p1 | hypothetical protein | 4 | 3 | 4 | 4 | 3 | 3 | 3 | 8 |
| 22 | TGME49_240350-t26_1-p1 | Toxoplasma gondii family E protein | 3 | 7 | 3 | 3 | 3 | 3 | 3 | 8 |
| 12 | TGME49_240360-t26_1-p1 | Toxoplasma gondii family E protein | 3 | 3 | 3 | 3 | 3 | 3 | 3 | 8 |
| 12 | TGME49_240365-t26_1-p1 | hypothetical protein | 3 | 3 | 3 | 3 | 3 | 3 | 3 | 8 |
| 12 | TGME49_240620-t26_1-p1 | hypothetical protein | 4 | 5 | 3 | 3 | 3 | 3 | 3 | 8 |
| 12 | TGME49_240755-t26_1-p1 | hypothetical protein | 3 | 3 | 3 | 3 | 3 | 3 | 3 | 8 |
| 12 | TGME49_240920-t26_1-p1 | hypothetical protein | 3 | 3 | 3 | 3 | 3 | 3 | 3 | 8 |
| 12 | TGME49_240940-t26_1-p1 | hypothetical protein | 4 | 3 | 3 | 3 | 3 | 3 | 3 | 8 |
| 12 | TGME49_241175-t26_1-p1 | hypothetical protein | 4 | 3 | 3 | 3 | 3 | 3 | 3 | 8 |
| 12 | TGME49_241300-t26_1-p1 | hypothetical protein | 3 | 5 | 3 | 3 | 3 | 3 | 3 | 8 |

| | | | | | | | | | | |
|---|---|---|---|---|---|---|---|---|---|---|
| 12 | TGME49_241310-t26_1-p1 | hypothetical protein | 3 | 3 | 3 | 3 | 3 | 3 | 3 | 8 |
| 12 | TGME49_241610-t26_1-p1 | dense granule protein GRA21 | 3 | 3 | 4 | 3 | 3 | 3 | 3 | 8 |
| 12 | TGME49_242100-t26_1-p1 | rhoptry kinase family protein | 3 | 3 | 3 | 3 | 3 | 3 | 3 | 8 |
| 22 | TGME49_242110-t26_1-p1 | rhoptry kinase family protein ROP38 | 3 | 8 | 3 | 3 | 3 | 4 | 3 | 8 |
| 12 | TGME49_242118-t26_1-p1 | myosin-light-chain kinase | 3 | 3 | 3 | 3 | 3 | 3 | 3 | 8 |
| 12 | TGME49_242230-t26_1-p1 | rhoptry kinase family protein ROP29 | 3 | 3 | 3 | 3 | 3 | 3 | 3 | 8 |
| 12 | TGME49_242340.P1736 | ribosomal protein RPS29 | 4 | 4 | 4 | 4 | 4 | 3 | 3 | 8 |
| 22 | TGME49_243130-t26_1-p1 | Toxoplasma gondii family A protein | 3 | 6 | 3 | 3 | 3 | 3 | 3 | 8 |
| 12 | TGME49_243220-t26_1-p1 | hypothetical protein | 3 | 3 | 3 | 3 | 3 | 3 | 3 | 8 |
| 12 | TGME49_243302-t26_1-p1 | hypothetical protein | 3 | 3 | 3 | 3 | 3 | 3 | 3 | 8 |
| 12 | TGME49_243420-t26_1-p1 | hypothetical protein | 3 | 3 | 3 | 3 | 3 | 3 | 3 | 8 |
| 12 | TGME49_243520-t26_1-p1 | hypothetical protein | 3 | 3 | 3 | 3 | 3 | 3 | 3 | 8 |
| 12 | TGME49_243700-t26_1-p1 | hypothetical protein | 3 | 3 | 3 | 3 | 3 | 3 | 3 | 8 |
| 12 | TGME49_243770-t26_1-p1 | hypothetical protein | 3 | 3 | 3 | 3 | 3 | 3 | 3 | 8 |
| 12 | TGME49_243900-t26_1-p1 | hypothetical protein | 3 | 3 | 3 | 3 | 3 | 3 | 3 | 8 |
| 12 | TGME49_244015-t26_1-p1 | hypothetical protein | 3 | 3 | 3 | 3 | 3 | 3 | 3 | 8 |
| 12 | TGME49_244165-t26_1-p1 | hypothetical protein | 3 | 3 | 3 | 3 | 3 | 3 | 3 | 8 |
| 12 | TGME49_244920-t26_1-p1 | Toxoplasma gondii family B protein | 3 | 3 | 3 | 4 | 3 | 3 | 3 | 8 |
| 12 | TGME49_244930-t26_1-p1 | hypothetical protein | 3 | 3 | 3 | 3 | 3 | 3 | 3 | 8 |
| 12 | TGME49_244940-t26_1-p1 | hypothetical protein | 3 | 3 | 3 | 3 | 3 | 3 | 3 | 8 |
| 12 | TGME49_244950-t26_1-p1 | hypothetical protein | 3 | 3 | 3 | 3 | 3 | 3 | 3 | 8 |
| 12 | TGME49_245748-t26_1-p1 | hypothetical protein | 3 | 3 | 3 | 3 | 3 | 3 | 3 | 8 |
| 12 | TGME49_245760-t26_1-p1 | hypothetical protein | 3 | 3 | 3 | 3 | 3 | 3 | 3 | 8 |
| 12 | TGME49_245770-t26_1-p1 | hypothetical protein | 3 | 3 | 3 | 3 | 3 | 3 | 3 | 8 |
| 12 | TGME49_246210-t26_1-p1 | hypothetical protein | 3 | 3 | 3 | 3 | 3 | 3 | 3 | 8 |
| 22 | TGME49_246530.P1833 | phospholipase D active site domain-containing protein | 3 | 7 | 7 | 5 | 4 | 4 | 4 | 8 |
| 12 | TGME49_246700-t26_1-p1 | hypothetical protein | 3 | 3 | 3 | 3 | 3 | 3 | 3 | 8 |
| 12 | TGME49_246763-t26_1-p1 | hypothetical protein | 3 | 3 | 3 | 3 | 3 | 3 | 3 | 8 |
| 12 | TGME49_247260-t26_1-p1 | hypothetical protein | 3 | 3 | 3 | 3 | 3 | 3 | 3 | 8 |
| 12 | TGME49_247520.P1857 | WAVE complex interacting protein WIP | 3 | 3 | 3 | 3 | 3 | 3 | 3 | 8 |
| 12 | TGME49_247620-t26_1-p1 | hypothetical protein | 3 | 3 | 3 | 3 | 3 | 3 | 3 | 8 |
| 12 | TGME49_247720-t26_1-p1 | hypothetical protein | 3 | 3 | 3 | 3 | 3 | 3 | 3 | 8 |
| 12 | TGME49_248280-t26_1-p1 | hypothetical protein | 4 | 3 | 3 | 3 | 3 | 3 | 3 | 8 |
| 12 | TGME49_248350-t26_1-p1 | hypothetical protein | 3 | 5 | 4 | 3 | 3 | 3 | 3 | 8 |
| 12 | TGME49_248425-t26_1-p1 | hypothetical protein | 3 | 3 | 3 | 3 | 3 | 3 | 3 | 8 |
| 12 | TGME49_248560-t26_1-p1 | hypothetical protein | 3 | 3 | 3 | 3 | 3 | 3 | 3 | 8 |
| 12 | TGME49_248620-t26_1-p1 | hypothetical protein | 3 | 3 | 3 | 3 | 3 | 3 | 3 | 8 |
| 12 | TGME49_249400-t26_1-p1 | hypothetical protein | 3 | 3 | 3 | 3 | 3 | 3 | 3 | 8 |
| 12 | TGME49_249575-t26_1-p1 | hypothetical protein | 3 | 3 | 3 | 3 | 3 | 3 | 3 | 8 |
| 12 | TGME49_249665-t26_1-p1 | hypothetical protein | 3 | 3 | 3 | 3 | 3 | 3 | 3 | 8 |
| 12 | TGME49_249685-t26_1-p1 | hypothetical protein | 3 | 3 | 3 | 3 | 3 | 3 | 3 | 8 |
| 12 | TGME49_249725-t26_1-p1 | hypothetical protein | 4 | 3 | 3 | 3 | 3 | 3 | 3 | 8 |
| 12 | TGME49_249740.P1942 | translation machinery associated tma7 protein | 4 | 4 | 4 | 4 | 3 | 3 | 4 | 8 |
| 12 | TGME49_250000-t26_1-p1 | hypothetical protein | 3 | 3 | 3 | 3 | 3 | 3 | 3 | 8 |
| 12 | TGME49_250020-t26_1-p1 | hypothetical protein | 3 | 3 | 3 | 3 | 3 | 3 | 3 | 8 |
| 12 | TGME49_250100-t26_1-p1 | hypothetical protein | 3 | 3 | 3 | 3 | 3 | 3 | 3 | 8 |
| 12 | TGME49_250220-t26_1-p1 | hypothetical protein | 3 | 3 | 3 | 3 | 3 | 3 | 3 | 8 |
| 12 | TGME49_250685-t26_1-p1 | hypothetical protein | 3 | 3 | 3 | 3 | 3 | 3 | 3 | 8 |
| 12 | TGME49_250905-t26_1-p1 | hypothetical protein | 3 | 3 | 3 | 3 | 3 | 3 | 3 | 8 |

| | | | | | | | | | | |
|---|---|---|---|---|---|---|---|---|---|---|
| 12 | TGME49_250930-t26_1-p1 | hypothetical protein | 3 | 3 | 3 | 3 | 3 | 3 | 3 | 8 |
| 12 | TGME49_250955-t26_1-p1 | KRUF family protein | 3 | 3 | 3 | 3 | 3 | 3 | 3 | 8 |
| 12 | TGME49_251530-t26_1-p1 | hypothetical protein | 3 | 3 | 3 | 3 | 3 | 3 | 3 | 8 |
| 12 | TGME49_251580-t26_1-p1 | hypothetical protein | 3 | 3 | 3 | 3 | 3 | 3 | 3 | 8 |
| 12 | TGME49_251650-t26_1-p1 | hypothetical protein | 4 | 3 | 3 | 3 | 3 | 3 | 3 | 8 |
| 12 | TGME49_251665-t26_1-p1 | hypothetical protein | 3 | 3 | 3 | 3 | 3 | 3 | 3 | 8 |
| 12 | TGME49_251860-t26_1-p1 | hypothetical protein | 3 | 3 | 3 | 3 | 3 | 3 | 3 | 8 |
| 12 | TGME49_251958-t26_1-p1 | SAG-related sequence SRS59B | 3 | 3 | 4 | 3 | 3 | 3 | 3 | 8 |
| 12 | TGME49_252065-t26_1-p1 | KRUF family protein | 3 | 3 | 3 | 3 | 3 | 3 | 3 | 8 |
| 12 | TGME49_252080-t26_1-p1 | hypothetical protein | 3 | 3 | 3 | 3 | 3 | 3 | 3 | 8 |
| 12 | TGME49_252090-t26_1-p1 | hypothetical protein | 3 | 3 | 3 | 3 | 3 | 3 | 3 | 8 |
| 12 | TGME49_252450-t26_1-p1 | hypothetical protein | 3 | 3 | 3 | 3 | 3 | 3 | 3 | 8 |
| 12 | TGME49_252620.P2002 | initiation factor, subunit 2 family protein | 3 | 3 | 3 | 3 | 3 | 3 | 3 | 8 |
| 12 | TGME49_253180-t26_1-p1 | hypothetical protein | 3 | 3 | 3 | 3 | 3 | 3 | 3 | 8 |
| 22 | TGME49_253390.P2020 | hypothetical protein | 5 | 9 | 5 | 7 | 3 | 3 | 3 | 8 |
| 12 | TGME49_253760-t26_1-p1 | hypothetical protein | 3 | 3 | 3 | 3 | 3 | 3 | 3 | 8 |
| 12 | TGME49_254485-t26_1-p1 | hypothetical protein | 3 | 3 | 3 | 3 | 3 | 3 | 3 | 8 |
| 12 | TGME49_254606-t26_1-p1 | hypothetical protein | 3 | 3 | 3 | 3 | 3 | 3 | 3 | 8 |
| 12 | TGME49_254620.P2062 | ribosomal protein RPL39 | 4 | 4 | 4 | 4 | 4 | 3 | 3 | 8 |
| 22 | TGME49_255060-t26_1-p1 | cytochrome b(N-terminal)/b6/petB subfamily protein | 3 | 7 | 7 | 4 | 3 | 3 | 3 | 8 |
| 22 | TGME49_255160.P2083 | hypothetical protein | 3 | 9 | 4 | 4 | 3 | 3 | 3 | 8 |
| 12 | TGME49_255175-t26_1-p1 | hypothetical protein | 3 | 3 | 3 | 3 | 3 | 3 | 3 | 8 |
| 12 | TGME49_255190.P2087 | myosin C, myosin B | 3 | 3 | 3 | 3 | 3 | 3 | 3 | 8 |
| 12 | TGME49_255195.P2088 | hypothetical protein | 3 | 3 | 8 | 3 | 3 | 3 | 3 | 8 |
| 12 | TGME49_256220-t26_1-p1 | hypothetical protein | 3 | 3 | 3 | 3 | 3 | 3 | 3 | 8 |
| 12 | TGME49_256650-t26_1-p1 | hypothetical protein | 3 | 3 | 3 | 3 | 3 | 3 | 3 | 8 |
| 12 | TGME49_256700.P2135 | hypothetical protein | 3 | 3 | 3 | 3 | 3 | 3 | 3 | 8 |
| 12 | TGME49_256965-t26_1-p1 | hypothetical protein | 3 | 3 | 3 | 3 | 3 | 3 | 3 | 8 |
| 12 | TGME49_257568-t26_1-p1 | hypothetical protein | 3 | 3 | 3 | 3 | 3 | 3 | 3 | 8 |
| 12 | TGME49_257810-t26_1-p1 | hypothetical protein | 3 | 3 | 3 | 3 | 3 | 3 | 3 | 8 |
| 12 | TGME49_257910.P2191 | hypothetical protein | 3 | 3 | 3 | 3 | 3 | 3 | 3 | 8 |
| 12 | TGME49_258040-t26_1-p1 | hypothetical protein | 3 | 3 | 3 | 3 | 3 | 3 | 3 | 8 |
| 12 | TGME49_258085-t26_1-p1 | hypothetical protein | 3 | 3 | 3 | 3 | 3 | 3 | 3 | 8 |
| 12 | TGME49_258225-t26_1-p1 | hypothetical protein | 3 | 3 | 3 | 3 | 3 | 3 | 3 | 8 |
| 22 | TGME49_258230-t26_1-p1 | rhoptry kinase family protein ROP20 | 3 | 8 | 6 | 6 | 4 | 3 | 3 | 8 |
| 12 | TGME49_259030-t26_1-p1 | hypothetical protein | 4 | 5 | 4 | 3 | 3 | 3 | 3 | 8 |
| 12 | TGME49_259100-t26_1-p1 | hypothetical protein | 3 | 3 | 3 | 3 | 3 | 3 | 3 | 8 |
| 12 | TGME49_259280-t26_1-p1 | hypothetical protein | 3 | 3 | 3 | 3 | 3 | 3 | 3 | 8 |
| 12 | TGME49_259300-t26_1-p1 | SAG-related sequence SRS26B | 3 | 3 | 3 | 4 | 3 | 3 | 3 | 8 |
| 12 | TGME49_259410-t26_1-p1 | SAG-related sequence SRS26A | 3 | 3 | 5 | 4 | 3 | 3 | 3 | 8 |
| 22 | TGME49_259570-t26_1-p1 | CMP/dCMP deaminase, zinc-binding, putative | 3 | 7 | 6 | 3 | 3 | 3 | 4 | 8 |
| 12 | TGME49_259900-t26_1-p1 | hypothetical protein | 3 | 5 | 3 | 3 | 3 | 3 | 3 | 8 |
| 12 | TGME49_260330-t26_1-p1 | hypothetical protein | 3 | 3 | 3 | 3 | 3 | 3 | 3 | 8 |
| 12 | TGME49_260850-t26_1-p1 | mitochondrial import inner membrane translocase subunit TIM10, putative | 3 | 4 | 4 | 4 | 3 | 3 | 4 | 8 |
| 12 | TGME49_260860-t26_1-p1 | hypothetical protein | 3 | 3 | 3 | 3 | 3 | 3 | 3 | 8 |
| 12 | TGME49_261300-t26_1-p1 | hypothetical protein | 3 | 3 | 3 | 3 | 3 | 3 | 3 | 8 |
| 12 | TGME49_261390-t26_1-p1 | hypothetical protein | 3 | 3 | 3 | 3 | 3 | 3 | 3 | 8 |
| 12 | TGME49_261740.P2380 | rhoptry protein ROP47 | 3 | 4 | 3 | 3 | 3 | 3 | 3 | 8 |

| | | | | | | | | | | |
|---|---|---|---|---|---|---|---|---|---|---|
| 12 | TGME49_261770-t26_1-p1 | hypothetical protein | 3 | 3 | 3 | 3 | 3 | 3 | 3 | 8 |
| 12 | TGME49_262020-t26_1-p1 | hypothetical protein | 3 | 3 | 3 | 3 | 3 | 3 | 3 | 8 |
| 12 | TGME49_262690.P2421 | ribosomal protein RPL27 | 3 | 3 | 3 | 3 | 3 | 3 | 3 | 8 |
| 12 | TGME49_262715-t26_1-p1 | hypothetical protein | 3 | 3 | 3 | 3 | 3 | 3 | 3 | 8 |
| 12 | TGME49_263020-t26_1-p1 | 50S ribosomal protein L33, putative | 4 | 4 | 4 | 4 | 3 | 3 | 3 | 8 |
| 12 | TGME49_263135-t26_1-p1 | hypothetical protein | 4 | 3 | 3 | 3 | 3 | 3 | 3 | 8 |
| 12 | TGME49_264100-t26_1-p1 | hypothetical protein | 3 | 3 | 3 | 3 | 3 | 3 | 3 | 8 |
| 12 | TGME49_264230-t26_1-p1 | hypothetical protein | 3 | 3 | 3 | 3 | 3 | 3 | 3 | 8 |
| 12 | TGME49_264240-t26_1-p1 | hypothetical protein | 3 | 3 | 3 | 3 | 3 | 3 | 3 | 8 |
| 12 | TGME49_264250-t26_1-p1 | hypothetical protein | 3 | 3 | 3 | 3 | 3 | 3 | 3 | 8 |
| 12 | TGME49_264260-t26_1-p1 | hypothetical protein | 3 | 3 | 3 | 3 | 3 | 3 | 3 | 8 |
| 12 | TGME49_264875-t26_1-p1 | hypothetical protein | 3 | 3 | 3 | 3 | 3 | 3 | 3 | 8 |
| 22 | TGME49_265140-t26_1-p1 | hypothetical protein | 3 | 7 | 5 | 6 | 5 | 3 | 3 | 8 |
| 12 | TGME49_265160-t26_1-p1 | hypothetical protein | 3 | 3 | 3 | 3 | 3 | 3 | 3 | 8 |
| 12 | TGME49_265175-t26_1-p1 | hypothetical protein | 3 | 3 | 3 | 3 | 3 | 3 | 3 | 8 |
| 22 | TGME49_265340-t26_1-p1 | hypothetical protein | 3 | 7 | 5 | 3 | 3 | 3 | 3 | 8 |
| 12 | TGME49_265542-t26_1-p1 | protein phosphatase 2C domain-containing protein | 3 | 3 | 3 | 4 | 3 | 3 | 3 | 8 |
| 12 | TGME49_266290-t26_1-p1 | hypothetical protein | 3 | 3 | 3 | 3 | 3 | 3 | 3 | 8 |
| 12 | TGME49_266335-t26_1-p1 | Toxoplasma gondii family A protein | 3 | 3 | 3 | 3 | 3 | 3 | 3 | 8 |
| 12 | TGME49_266368-t26_1-p1 | hypothetical protein | 4 | 3 | 3 | 3 | 3 | 3 | 3 | 8 |
| 12 | TGME49_266480-t26_1-p1 | hypothetical protein | 3 | 4 | 3 | 3 | 3 | 3 | 3 | 8 |
| 12 | TGME49_266600-t26_1-p1 | Kazal-type serine protease inhibitor domain-containing protein | 3 | 5 | 3 | 4 | 3 | 3 | 3 | 8 |
| 12 | TGME49_266610.P2570 | Kazal-type serine protease inhibitor domain-containing protein | 3 | 3 | 3 | 3 | 3 | 3 | 3 | 8 |
| 12 | TGME49_266675-t26_1-p1 | hypothetical protein | 3 | 3 | 3 | 3 | 3 | 3 | 3 | 8 |
| 12 | TGME49_266980-t26_1-p1 | hypothetical protein | 3 | 3 | 3 | 3 | 3 | 3 | 3 | 8 |
| 12 | TGME49_267000-t26_1-p1 | hypothetical protein | 3 | 3 | 3 | 3 | 3 | 3 | 3 | 8 |
| 12 | TGME49_267010-t26_1-p1 | hypothetical protein | 3 | 3 | 3 | 3 | 3 | 3 | 3 | 8 |
| 12 | TGME49_267290-t26_1-p1 | hypothetical protein | 3 | 3 | 3 | 3 | 3 | 3 | 3 | 8 |
| 12 | TGME49_267435-t26_1-p1 | hypothetical protein | 3 | 3 | 3 | 3 | 3 | 3 | 3 | 8 |
| 12 | TGME49_267585-t26_1-p1 | hypothetical protein | 3 | 3 | 3 | 3 | 3 | 3 | 3 | 8 |
| 12 | TGME49_267610-t26_1-p1 | hypothetical protein | 3 | 3 | 3 | 3 | 3 | 3 | 3 | 8 |
| 12 | TGME49_267990-t26_1-p1 | hypothetical protein | 3 | 3 | 3 | 3 | 3 | 3 | 3 | 8 |
| 12 | TGME49_268170-t26_1-p1 | hypothetical protein | 4 | 3 | 3 | 3 | 3 | 3 | 3 | 8 |
| 12 | TGME49_268178-t26_1-p1 | hypothetical protein | 3 | 3 | 3 | 3 | 3 | 3 | 3 | 8 |
| 12 | TGME49_268300.P2631 | hypothetical protein | 3 | 3 | 3 | 3 | 3 | 3 | 3 | 8 |
| 12 | TGME49_268765-t26_1-p1 | hypothetical protein | 3 | 3 | 3 | 3 | 3 | 3 | 3 | 8 |
| 12 | TGME49_268835-t26_1-p1 | hypothetical protein | 4 | 3 | 3 | 3 | 3 | 3 | 3 | 8 |
| 12 | TGME49_269085-t26_1-p1 | hypothetical protein | 3 | 3 | 3 | 3 | 3 | 3 | 3 | 8 |
| 12 | TGME49_269417-t26_1-p1 | hypothetical protein | 4 | 3 | 3 | 3 | 3 | 3 | 3 | 8 |
| 12 | TGME49_269425-t26_1-p1 | hypothetical protein | 3 | 3 | 3 | 3 | 3 | 3 | 3 | 8 |
| 12 | TGME49_269470-t26_1-p1 | hypothetical protein | 3 | 3 | 3 | 3 | 3 | 3 | 3 | 8 |
| 12 | TGME49_269775-t26_1-p1 | hypothetical protein | 3 | 3 | 3 | 3 | 3 | 3 | 3 | 8 |
| 12 | TGME49_270020-t26_1-p1 | hypothetical protein | 3 | 3 | 3 | 3 | 3 | 3 | 3 | 8 |
| 12 | TGME49_270030-t26_1-p1 | hypothetical protein | 3 | 3 | 3 | 3 | 3 | 3 | 3 | 8 |
| 12 | TGME49_270040-t26_1-p1 | hypothetical protein | 3 | 3 | 3 | 3 | 3 | 3 | 3 | 8 |
| 12 | TGME49_270277-t26_1-p1 | hypothetical protein | 3 | 3 | 5 | 3 | 3 | 3 | 3 | 8 |
| 12 | TGME49_270290-t26_1-p1 | hypothetical protein | 3 | 3 | 3 | 3 | 3 | 3 | 3 | 8 |
| 12 | TGME49_270520.P2738 | hypothetical protein | 3 | 3 | 3 | 3 | 3 | 3 | 3 | 8 |

| | | | | | | | | | | |
|---|---|---|---|---|---|---|---|---|---|---|
| 12 | TGME49_270520.P2739 | hypothetical protein | 3 | 3 | 3 | 3 | 3 | 3 | 3 | 8 |
| 12 | TGME49_270520.P2740 | hypothetical protein | 3 | 3 | 3 | 3 | 3 | 3 | 3 | 8 |
| 12 | TGME49_270520.P2741 | hypothetical protein | 3 | 3 | 3 | 3 | 3 | 3 | 3 | 8 |
| 12 | TGME49_270520.P2742 | hypothetical protein | 3 | 3 | 3 | 3 | 3 | 3 | 3 | 8 |
| 12 | TGME49_270520.P2743 | hypothetical protein | 3 | 3 | 3 | 3 | 3 | 3 | 3 | 8 |
| 12 | TGME49_270520.P2744 | hypothetical protein | 3 | 3 | 3 | 3 | 3 | 3 | 3 | 8 |
| 12 | TGME49_270520.P2745 | hypothetical protein | 3 | 3 | 3 | 3 | 5 | 3 | 3 | 8 |
| 12 | TGME49_270520.P2747 | hypothetical protein | 3 | 3 | 3 | 4 | 3 | 3 | 3 | 8 |
| 12 | TGME49_270520.P2748 | hypothetical protein | 3 | 3 | 3 | 3 | 3 | 3 | 3 | 8 |
| 12 | TGME49_270520.P2749 | hypothetical protein | 3 | 3 | 3 | 3 | 3 | 3 | 3 | 8 |
| 12 | TGME49_270520.P2750 | hypothetical protein | 3 | 3 | 3 | 3 | 3 | 3 | 3 | 8 |
| 12 | TGME49_270520.P2751 | hypothetical protein | 3 | 3 | 3 | 3 | 3 | 3 | 3 | 8 |
| 12 | TGME49_270520.P2752 | hypothetical protein | 3 | 3 | 3 | 3 | 3 | 3 | 3 | 8 |
| 12 | TGME49_270520.P2753 | hypothetical protein | 3 | 3 | 3 | 3 | 3 | 3 | 3 | 8 |
| 12 | TGME49_270520.P2754 | hypothetical protein | 3 | 3 | 3 | 3 | 3 | 3 | 3 | 8 |
| 12 | TGME49_270520.P2755 | hypothetical protein | 3 | 3 | 3 | 3 | 3 | 3 | 3 | 8 |
| 12 | TGME49_270520.P2756 | hypothetical protein | 3 | 3 | 3 | 3 | 3 | 3 | 3 | 8 |
| 12 | TGME49_270520.P2757 | hypothetical protein | 3 | 3 | 3 | 3 | 3 | 3 | 3 | 8 |
| 12 | TGME49_270520.P2758 | hypothetical protein | 3 | 3 | 3 | 3 | 3 | 4 | 3 | 8 |
| 12 | TGME49_270520.P2759 | hypothetical protein | 3 | 3 | 3 | 3 | 3 | 3 | 3 | 8 |
| 12 | TGME49_270520.P2760 | hypothetical protein | 3 | 3 | 6 | 3 | 3 | 3 | 3 | 8 |
| 12 | TGME49_270520.P2761 | hypothetical protein | 3 | 3 | 3 | 3 | 3 | 3 | 3 | 8 |
| 12 | TGME49_270520.P2762 | hypothetical protein | 3 | 3 | 3 | 3 | 3 | 3 | 3 | 8 |
| 12 | TGME49_270520.P2763 | hypothetical protein | 3 | 3 | 3 | 3 | 3 | 3 | 3 | 8 |
| 12 | TGME49_270520.P2764 | hypothetical protein | 3 | 3 | 3 | 3 | 3 | 3 | 3 | 8 |
| 22 | TGME49_270520.P2765 | hypothetical protein | 3 | 6 | 3 | 3 | 3 | 3 | 3 | 8 |
| 12 | TGME49_270520.P2766 | hypothetical protein | 3 | 3 | 3 | 3 | 3 | 3 | 3 | 8 |
| 12 | TGME49_270520.P2767 | hypothetical protein | 3 | 3 | 3 | 3 | 3 | 3 | 3 | 8 |
| 12 | TGME49_270520.P2768 | hypothetical protein | 3 | 3 | 3 | 3 | 3 | 3 | 3 | 8 |
| 12 | TGME49_270520.P2769 | hypothetical protein | 3 | 3 | 3 | 3 | 3 | 3 | 3 | 8 |
| 12 | TGME49_270655-t26_1-p1 | hypothetical protein | 3 | 3 | 3 | 3 | 3 | 3 | 3 | 8 |
| 12 | TGME49_270680-t26_1-p1 | hypothetical protein | 3 | 3 | 3 | 3 | 3 | 3 | 3 | 8 |
| 12 | TGME49_271025-t26_1-p1 | hypothetical protein | 3 | 3 | 3 | 3 | 3 | 3 | 3 | 8 |
| 12 | TGME49_271115-t26_1-p1 | hypothetical protein | 3 | 3 | 3 | 3 | 3 | 3 | 3 | 8 |
| 12 | TGME49_271375-t26_1-p1 | hypothetical protein | 3 | 3 | 3 | 3 | 3 | 3 | 3 | 8 |
| 12 | TGME49_271400-t26_1-p1 | hypothetical protein | 3 | 3 | 3 | 4 | 3 | 3 | 3 | 8 |
| 12 | TGME49_271410-t26_1-p1 | hypothetical protein | 3 | 3 | 3 | 3 | 3 | 3 | 3 | 8 |
| 12 | TGME49_271570-t26_1-p1 | Toxoplasma gondii family D protein | 3 | 3 | 3 | 3 | 3 | 3 | 3 | 8 |
| 22 | TGME49_271590-t26_1-p1 | Toxoplasma gondii family D protein | 3 | 7 | 3 | 7 | 3 | 3 | 3 | 8 |
| 12 | TGME49_271985-t26_1-p1 | hypothetical protein | 3 | 3 | 3 | 3 | 3 | 3 | 3 | 8 |
| 12 | TGME49_272020.P2837 | hypothetical protein | 3 | 3 | 3 | 3 | 3 | 3 | 3 | 8 |
| 12 | TGME49_272300-t26_1-p1 | hypothetical protein | 3 | 3 | 3 | 3 | 3 | 3 | 3 | 8 |
| 12 | TGME49_272450-t26_1-p1 | hypothetical protein | 3 | 3 | 3 | 3 | 3 | 3 | 3 | 8 |
| 12 | TGME49_272560-t26_1-p1 | SAG-related sequence SRS31B | 3 | 3 | 3 | 3 | 3 | 3 | 3 | 8 |
| 12 | TGME49_272593-t26_1-p1 | hypothetical protein | 3 | 3 | 3 | 3 | 3 | 3 | 3 | 8 |
| 12 | TGME49_272595-t26_1-p1 | hypothetical protein | 3 | 3 | 3 | 3 | 3 | 3 | 3 | 8 |
| 12 | TGME49_273020-t26_1-p1 | hypothetical protein | 3 | 3 | 3 | 3 | 3 | 3 | 3 | 8 |
| 12 | TGME49_273075-t26_1-p1 | hypothetical protein | 3 | 3 | 3 | 3 | 3 | 3 | 3 | 8 |
| 12 | TGME49_273080-t26_1-p1 | hypothetical protein | 3 | 3 | 3 | 3 | 3 | 3 | 3 | 8 |
| 12 | TGME49_273300-t26_1-p1 | hypothetical protein | 3 | 3 | 3 | 3 | 3 | 3 | 3 | 8 |

| | | | | | | | | | | |
|---|---|---|---|---|---|---|---|---|---|---|
| 12 | TGME49_273310.P2897 | hypothetical protein | 3 | 4 | 4 | 3 | 3 | 3 | 3 | 8 |
| 12 | TGME49_273650-t26_1-p1 | kinesin B | 3 | 3 | 3 | 3 | 3 | 3 | 3 | 8 |
| 12 | TGME49_273770.P2918 | selenoprotein Q, putative | 4 | 4 | 4 | 4 | 3 | 3 | 3 | 8 |
| 12 | TGME49_273915-t26_1-p1 | hypothetical protein | 3 | 3 | 3 | 3 | 3 | 3 | 3 | 8 |
| 12 | TGME49_273985-t26_1-p1 | hypothetical protein | 3 | 3 | 3 | 3 | 3 | 3 | 3 | 8 |
| 12 | TGME49_274280-t26_1-p1 | hypothetical protein | 3 | 3 | 3 | 3 | 3 | 3 | 3 | 8 |
| 12 | TGME49_274290-t26_1-p1 | Toxoplasma gondii family B protein | 3 | 4 | 3 | 3 | 3 | 3 | 3 | 8 |
| 12 | TGME49_274700-t26_1-p1 | hypothetical protein | 3 | 3 | 3 | 3 | 3 | 3 | 3 | 8 |
| 12 | TGME49_275355-t26_1-p1 | SAG-related sequence SRS47A | 3 | 3 | 3 | 3 | 3 | 3 | 3 | 8 |
| 22 | TGME49_275360.P2950 | SAG-related sequence SRS47B | 3 | 6 | 3 | 3 | 3 | 3 | 3 | 8 |
| 12 | TGME49_275370-t26_1-p1 | SAG-related sequence SRS47C | 3 | 3 | 3 | 3 | 3 | 3 | 3 | 8 |
| 12 | TGME49_275390-t26_1-p1 | hypothetical protein | 3 | 3 | 3 | 3 | 3 | 3 | 3 | 8 |
| 12 | TGME49_275795-t26_1-p1 | hypothetical protein | 3 | 3 | 3 | 3 | 3 | 3 | 3 | 8 |
| 12 | TGME49_276230-t26_1-p1 | Toxoplasma gondii family B protein | 3 | 3 | 3 | 3 | 3 | 3 | 3 | 8 |
| 12 | TGME49_276810-t26_1-p1 | hypothetical protein | 3 | 3 | 3 | 3 | 3 | 3 | 3 | 8 |
| 12 | TGME49_276820-t26_1-p1 | hypothetical protein | 3 | 3 | 3 | 3 | 3 | 3 | 3 | 8 |
| 12 | TGME49_276830-t26_1-p1 | hypothetical protein | 3 | 3 | 3 | 3 | 3 | 3 | 3 | 8 |
| 12 | TGME49_277020-t26_1-p1 | hypothetical protein | 4 | 3 | 3 | 3 | 3 | 3 | 3 | 8 |
| 12 | TGME49_277055-t26_1-p1 | hypothetical protein | 3 | 3 | 3 | 3 | 3 | 3 | 3 | 8 |
| 12 | TGME49_277240-t26_1-p1 | NTPase I | 3 | 3 | 3 | 3 | 3 | 3 | 3 | 8 |
| 12 | TGME49_277270-t26_1-p1 | NTPase II | 3 | 3 | 5 | 3 | 3 | 3 | 3 | 8 |
| 12 | TGME49_277685-t26_1-p1 | hypothetical protein | 3 | 3 | 3 | 3 | 3 | 3 | 3 | 8 |
| 12 | TGME49_277705-t26_1-p1 | hypothetical protein | 3 | 3 | 3 | 3 | 3 | 3 | 3 | 8 |
| 12 | TGME49_277730-t26_1-p1 | hypothetical protein | 3 | 3 | 3 | 3 | 3 | 3 | 3 | 8 |
| 12 | TGME49_277830-t26_1-p1 | hypothetical protein | 3 | 3 | 3 | 3 | 3 | 3 | 3 | 8 |
| 12 | TGME49_277920.P2989 | hypothetical protein | 4 | 4 | 4 | 3 | 3 | 3 | 3 | 8 |
| 12 | TGME49_278005-t26_1-p1 | hypothetical protein | 3 | 3 | 3 | 3 | 3 | 3 | 3 | 8 |
| 12 | TGME49_278070-t26_1-p1 | Toxoplasma gondii family A protein | 3 | 3 | 3 | 3 | 3 | 3 | 3 | 8 |
| 12 | TGME49_278080-t26_1-p1 | Toxoplasma gondii family A protein | 3 | 3 | 3 | 3 | 3 | 3 | 3 | 8 |
| 12 | TGME49_278270.P2998 | nucleolar protein, structural component of H/ACA snoRNPs, putative | 4 | 4 | 4 | 3 | 4 | 3 | 3 | 8 |
| 12 | TGME49_278300-t26_1-p1 | Toxoplasma gondii family A protein | 3 | 3 | 3 | 3 | 3 | 3 | 3 | 8 |
| 12 | TGME49_278310-t26_1-p1 | hypothetical protein | 3 | 3 | 3 | 3 | 3 | 3 | 3 | 8 |
| 22 | TGME49_278320-t26_1-p1 | Toxoplasma gondii family A protein | 3 | 6 | 5 | 3 | 3 | 3 | 3 | 8 |
| 12 | TGME49_278370-t26_1-p1 | Toxoplasma gondii family A protein | 3 | 3 | 3 | 3 | 3 | 3 | 3 | 8 |
| 12 | TGME49_278400-t26_1-p1 | Toxoplasma gondii family A protein | 3 | 3 | 3 | 3 | 3 | 3 | 3 | 8 |
| 12 | TGME49_278757-t26_1-p1 | hypothetical protein | 3 | 3 | 3 | 3 | 3 | 3 | 3 | 8 |
| 12 | TGME49_278830.P3015 | glucose-6-phosphate 1-dehydrogenase | 3 | 3 | 3 | 3 | 3 | 3 | 3 | 8 |
| 12 | TGME49_278830.P3017 | glucose-6-phosphate 1-dehydrogenase | 3 | 3 | 3 | 3 | 3 | 3 | 3 | 8 |
| 22 | TGME49_278830.P3018 | glucose-6-phosphate 1-dehydrogenase | 3 | 10 | 3 | 3 | 3 | 3 | 3 | 8 |
| 12 | TGME49_278878-t26_1-p1 | GDA1/CD39 (nucleoside phosphatase) family protein | 3 | 3 | 3 | 3 | 3 | 3 | 3 | 8 |
| 12 | TGME49_279000-t26_1-p1 | SAG-related sequence SRS59J | 3 | 3 | 3 | 3 | 3 | 3 | 3 | 8 |
| 12 | TGME49_279310-t26_1-p1 | hypothetical protein | 3 | 3 | 3 | 3 | 3 | 3 | 3 | 8 |
| 12 | TGME49_279355-t26_1-p1 | hypothetical protein | 3 | 3 | 3 | 3 | 3 | 3 | 3 | 8 |
| 12 | TGME49_279405-t26_1-p1 | hypothetical protein | 3 | 3 | 3 | 3 | 3 | 3 | 3 | 8 |
| 12 | TGME49_279560-t26_1-p1 | hypothetical protein | 3 | 3 | 3 | 3 | 3 | 3 | 3 | 8 |
| 12 | TGME49_280465-t26_1-p1 | hypothetical protein | 3 | 3 | 3 | 3 | 3 | 3 | 3 | 8 |
| 12 | TGME49_281030-t26_1-p1 | hypothetical protein | 3 | 3 | 3 | 3 | 3 | 3 | 3 | 8 |
| 12 | TGME49_281040-t26_1-p1 | hypothetical protein | 3 | 3 | 3 | 3 | 3 | 3 | 3 | 8 |

| | | | | | | | | | | |
|---|---|---|---|---|---|---|---|---|---|---|
| 12 | TGME49_282160-t26_1-p1 | hypothetical protein | 3 | 3 | 3 | 3 | 3 | 3 | 3 | 8 |
| 12 | TGME49_283770-t26_1-p1 | hypothetical protein | 3 | 3 | 3 | 3 | 3 | 3 | 3 | 8 |
| 12 | TGME49_283800-t26_1-p1 | hypothetical protein | 3 | 3 | 3 | 3 | 3 | 3 | 3 | 8 |
| 12 | TGME49_283855-t26_1-p1 | hypothetical protein | 3 | 3 | 3 | 3 | 3 | 3 | 3 | 8 |
| 12 | TGME49_283870-t26_1-p1 | hypothetical protein | 3 | 3 | 3 | 3 | 3 | 3 | 3 | 8 |
| 12 | TGME49_284310-t26_1-p1 | hypothetical protein | 3 | 3 | 3 | 3 | 3 | 3 | 3 | 8 |
| 12 | TGME49_284680-t26_1-p1 | hypothetical protein | 3 | 3 | 3 | 3 | 3 | 3 | 3 | 8 |
| 12 | TGME49_285665-t26_1-p1 | hypothetical protein | 3 | 3 | 3 | 3 | 3 | 3 | 3 | 8 |
| 12 | TGME49_285740-t26_1-p1 | hypothetical protein | 3 | 3 | 3 | 3 | 3 | 3 | 3 | 8 |
| 12 | TGME49_285825-t26_1-p1 | hypothetical protein | 3 | 3 | 3 | 3 | 3 | 3 | 3 | 8 |
| 12 | TGME49_285865-t26_1-p1 | hypothetical protein | 3 | 3 | 3 | 3 | 3 | 3 | 3 | 8 |
| 22 | TGME49_285870-t26_1-p1 | SAG-related sequence SRS20A | 3 | 6 | 4 | 4 | 3 | 3 | 3 | 8 |
| 12 | TGME49_286170-t26_1-p1 | hypothetical protein | 3 | 3 | 3 | 3 | 3 | 3 | 3 | 8 |
| 12 | TGME49_286200-t26_1-p1 | hypothetical protein | 3 | 3 | 3 | 3 | 3 | 3 | 3 | 8 |
| 12 | TGME49_286450-t26_1-p1 | dense granule protein GRA5 | 4 | 3 | 3 | 3 | 3 | 3 | 3 | 8 |
| 12 | TGME49_286480-t26_1-p1 | hypothetical protein | 3 | 3 | 3 | 3 | 3 | 3 | 3 | 8 |
| 12 | TGME49_286485-t26_1-p1 | hypothetical protein | 3 | 3 | 3 | 3 | 3 | 3 | 3 | 8 |
| 12 | TGME49_286525-t26_1-p1 | hypothetical protein | 3 | 3 | 3 | 3 | 3 | 3 | 3 | 8 |
| 12 | TGME49_286595-t26_1-p1 | hypothetical protein | 3 | 3 | 3 | 3 | 3 | 3 | 3 | 8 |
| 12 | TGME49_286928-t26_1-p1 | hypothetical protein | 3 | 3 | 3 | 3 | 3 | 3 | 3 | 8 |
| 12 | TGME49_287235-t26_1-p1 | hypothetical protein | 4 | 3 | 3 | 3 | 3 | 3 | 3 | 8 |
| 12 | TGME49_287240-t26_1-p1 | hypothetical protein | 3 | 3 | 3 | 3 | 3 | 3 | 3 | 8 |
| 12 | TGME49_287280.P3173 | Toxoplasma gondii family C protein | 3 | 3 | 3 | 3 | 3 | 3 | 3 | 8 |
| 12 | TGME49_287430-t26_1-p1 | hypothetical protein | 3 | 3 | 3 | 3 | 3 | 3 | 3 | 8 |
| 12 | TGME49_287440-t26_1-p1 | hypothetical protein | 3 | 3 | 3 | 3 | 3 | 3 | 3 | 8 |
| 12 | TGME49_287450-t26_1-p1 | hypothetical protein | 4 | 3 | 3 | 3 | 3 | 3 | 3 | 8 |
| 12 | TGME49_287460-t26_1-p1 | hypothetical protein | 3 | 3 | 3 | 3 | 3 | 3 | 3 | 8 |
| 12 | TGME49_287470-t26_1-p1 | hypothetical protein | 3 | 4 | 3 | 3 | 3 | 3 | 3 | 8 |
| 12 | TGME49_287510-t26_1-p1 | aromatic amino acid hydrolase AAH1 | 3 | 3 | 3 | 3 | 8 | 3 | 3 | 8 |
| 12 | TGME49_287515-t26_1-p1 | hypothetical protein | 3 | 3 | 3 | 3 | 3 | 3 | 3 | 8 |
| 12 | TGME49_287990-t26_1-p1 | hypothetical protein | 3 | 3 | 3 | 3 | 3 | 3 | 3 | 8 |
| 12 | TGME49_288010-t26_1-p1 | hypothetical protein | 3 | 3 | 3 | 3 | 3 | 3 | 3 | 8 |
| 12 | TGME49_288245-t26_1-p1 | hypothetical protein | 4 | 3 | 4 | 3 | 4 | 3 | 3 | 8 |
| 12 | TGME49_288300-t26_1-p1 | hypothetical protein | 3 | 3 | 3 | 3 | 3 | 3 | 3 | 8 |
| 12 | TGME49_288455-t26_1-p1 | hypothetical protein | 4 | 3 | 3 | 3 | 3 | 3 | 3 | 8 |
| 12 | TGME49_288465-t26_1-p1 | hypothetical protein | 3 | 3 | 3 | 3 | 3 | 3 | 3 | 8 |
| 12 | TGME49_288900-t26_1-p1 | Yos1 family protein | 3 | 3 | 4 | 3 | 3 | 3 | 3 | 8 |
| 12 | TGME49_289030-t26_1-p1 | hypothetical protein | 3 | 3 | 3 | 3 | 3 | 3 | 3 | 8 |
| 12 | TGME49_289160-t26_1-p1 | hypothetical protein | 4 | 3 | 3 | 3 | 3 | 3 | 3 | 8 |
| 12 | TGME49_289370-t26_1-p1 | hypothetical protein | 4 | 3 | 3 | 3 | 3 | 3 | 3 | 8 |
| 12 | TGME49_289650.P3207 | PEP-carboxykinase I | 3 | 3 | 3 | 3 | 3 | 3 | 3 | 8 |
| 12 | TGME49_289745-t26_1-p1 | hypothetical protein | 3 | 3 | 3 | 3 | 3 | 3 | 3 | 8 |
| 12 | TGME49_289810-t26_1-p1 | hypothetical protein | 3 | 3 | 3 | 3 | 3 | 3 | 3 | 8 |
| 12 | TGME49_290010-t26_1-p1 | hypothetical protein | 3 | 3 | 3 | 3 | 3 | 3 | 3 | 8 |
| 12 | TGME49_290280-t26_1-p1 | hypothetical protein | 3 | 3 | 3 | 3 | 3 | 3 | 3 | 8 |
| 12 | TGME49_290645-t26_1-p1 | hypothetical protein | 3 | 3 | 3 | 3 | 3 | 3 | 3 | 8 |
| 12 | TGME49_290923.P3240 | hypothetical protein | 4 | 3 | 3 | 5 | 3 | 4 | 4 | 8 |
| 12 | TGME49_291580-t26_1-p1 | hypothetical protein | 3 | 3 | 3 | 3 | 3 | 3 | 3 | 8 |
| 22 | TGME49_291600-t26_1-p1 | gamma interferon inducible lysosomal thiol reductase (GILT) protein | 3 | 7 | 5 | 5 | 3 | 4 | 4 | 8 |

| | | | | | | | | | | |
|---|---|---|---|---|---|---|---|---|---|---|
| 22 | TGME49_291620-t26_1-p1 | hypothetical protein | 4 | 8 | 6 | 4 | 4 | 3 | 3 | 8 |
| 12 | TGME49_291850.P3252 | ribosomal protein RPS30 | 4 | 4 | 3 | 4 | 4 | 3 | 3 | 8 |
| 12 | TGME49_291920-t26_1-p1 | hypothetical protein | 3 | 3 | 3 | 3 | 3 | 3 | 3 | 8 |
| 12 | TGME49_292250-t26_1-p1 | SAG-related sequence SRS36A | 3 | 3 | 3 | 3 | 3 | 3 | 3 | 8 |
| 22 | TGME49_292270-t26_1-p1 | SAG-related sequence SRS36C | 3 | 7 | 3 | 3 | 3 | 3 | 3 | 8 |
| 12 | TGME49_292325-t26_1-p1 | hypothetical protein | 3 | 3 | 3 | 3 | 3 | 3 | 3 | 8 |
| 12 | TGME49_292390.P3267 | KRUF family protein | 3 | 3 | 6 | 3 | 3 | 3 | 3 | 8 |
| 12 | TGME49_292610-t26_1-p1 | Toxoplasma gondii family C protein | 3 | 3 | 3 | 3 | 3 | 3 | 3 | 8 |
| 12 | TGME49_293010.P3271 | hypothetical protein | 4 | 4 | 3 | 4 | 3 | 3 | 3 | 8 |
| 22 | TGME49_293252.P3282 | hypothetical protein | 3 | 9 | 4 | 3 | 3 | 3 | 3 | 8 |
| 12 | TGME49_293520-t26_1-p1 | hypothetical protein | 3 | 3 | 3 | 3 | 3 | 3 | 3 | 8 |
| 12 | TGME49_293740.P3324 | hypothetical protein | 3 | 3 | 3 | 3 | 3 | 3 | 3 | 8 |
| 12 | TGME49_293830.P3333 | methyltransferase domain-containing protein | 3 | 3 | 3 | 3 | 3 | 3 | 4 | 8 |
| 12 | TGME49_294030-t26_1-p1 | hypothetical protein | 3 | 3 | 3 | 3 | 3 | 3 | 3 | 8 |
| 12 | TGME49_294225-t26_1-p1 | hypothetical protein | 3 | 3 | 3 | 3 | 3 | 3 | 3 | 8 |
| 12 | TGME49_294285-t26_1-p1 | hypothetical protein | 3 | 3 | 3 | 3 | 3 | 3 | 3 | 8 |
| 12 | TGME49_294680-t26_1-p1 | hypothetical protein | 3 | 3 | 3 | 3 | 3 | 3 | 3 | 8 |
| 12 | TGME49_294760-t26_1-p1 | hypothetical protein | 3 | 3 | 3 | 3 | 3 | 3 | 3 | 8 |
| 12 | TGME49_294800-t26_1-p1 | elongation factor 1-alpha (EF-1-ALPHA), putative | 3 | 3 | 3 | 3 | 3 | 3 | 3 | 8 |
| 12 | TGME49_294805-t26_1-p1 | hypothetical protein | 3 | 3 | 3 | 3 | 3 | 3 | 3 | 8 |
| 12 | TGME49_294980-t26_1-p1 | hypothetical protein | 3 | 3 | 3 | 3 | 3 | 3 | 3 | 8 |
| 12 | TGME49_294990-t26_1-p1 | hypothetical protein | 3 | 3 | 3 | 3 | 3 | 3 | 3 | 8 |
| 12 | TGME49_295130-t26_1-p1 | hypothetical protein | 3 | 3 | 3 | 3 | 3 | 3 | 3 | 8 |
| 12 | TGME49_295140-t26_1-p1 | 1-phosphatidylinositol 4-kinase | 3 | 3 | 3 | 3 | 3 | 3 | 3 | 8 |
| 12 | TGME49_295685-t26_1-p1 | hypothetical protein | 3 | 3 | 3 | 3 | 3 | 3 | 3 | 8 |
| 12 | TGME49_295945-t26_1-p1 | hypothetical protein | 3 | 3 | 3 | 3 | 3 | 3 | 3 | 8 |
| 12 | TGME49_295970-t26_1-p1 | hypothetical protein | 3 | 5 | 3 | 3 | 3 | 3 | 3 | 8 |
| 12 | TGME49_296000.P3395 | rhoptry kinase family protein ROP2L12 | 3 | 3 | 3 | 3 | 3 | 3 | 3 | 8 |
| 12 | TGME49_296015-t26_1-p1 | rhoptry kinase family protein | 3 | 3 | 3 | 3 | 3 | 3 | 3 | 8 |
| 12 | TGME49_296020-t26_1-p1 | hypothetical protein | 3 | 3 | 3 | 3 | 3 | 3 | 3 | 8 |
| 12 | TGME49_296340-t26_1-p1 | hypothetical protein | 3 | 3 | 3 | 3 | 3 | 3 | 3 | 8 |
| 12 | TGME49_296640.P3397 | SAG-related sequence SRS48E | 3 | 3 | 3 | 3 | 3 | 3 | 3 | 8 |
| 12 | TGME49_297065-t26_1-p1 | hypothetical protein | 3 | 3 | 3 | 3 | 3 | 3 | 3 | 8 |
| 12 | TGME49_297190-t26_1-p1 | hypothetical protein | 3 | 3 | 3 | 3 | 3 | 3 | 3 | 8 |
| 12 | TGME49_297465.P3408 | hypothetical protein | 3 | 3 | 4 | 3 | 3 | 4 | 3 | 8 |
| 12 | TGME49_297643-t26_1-p1 | hypothetical protein | 3 | 3 | 3 | 3 | 3 | 3 | 3 | 8 |
| 12 | TGME49_297725-t26_1-p1 | hypothetical protein | 3 | 3 | 3 | 3 | 3 | 3 | 3 | 8 |
| 22 | TGME49_297780.P3418 | DNA gyrase subunit B | 3 | 8 | 3 | 7 | 5 | 3 | 5 | 8 |
| 12 | TGME49_298070-t26_1-p1 | hypothetical protein | 3 | 3 | 3 | 3 | 3 | 3 | 3 | 8 |
| 12 | TGME49_298080-t26_1-p1 | hypothetical protein | 3 | 3 | 3 | 3 | 3 | 3 | 3 | 8 |
| 12 | TGME49_298820-t26_1-p1 | hypothetical protein | 3 | 3 | 3 | 3 | 3 | 3 | 3 | 8 |
| 12 | TGME49_298830-t26_1-p1 | hypothetical protein | 3 | 3 | 3 | 3 | 3 | 3 | 3 | 8 |
| 12 | TGME49_298840-t26_1-p1 | hypothetical protein | 3 | 3 | 3 | 3 | 3 | 3 | 3 | 8 |
| 12 | TGME49_298960.P3431 | Toxoplasma gondii family C protein | 3 | 3 | 3 | 3 | 3 | 3 | 3 | 8 |
| 12 | TGME49_299000-t26_1-p1 | hypothetical protein | 3 | 4 | 4 | 3 | 3 | 3 | 3 | 8 |
| 12 | TGME49_299010-t26_1-p1 | hypothetical protein | 4 | 3 | 3 | 3 | 3 | 3 | 3 | 8 |
| 12 | TGME49_299015-t26_1-p1 | hypothetical protein | 3 | 3 | 3 | 3 | 3 | 3 | 3 | 8 |
| 12 | TGME49_299160-t26_1-p1 | hypothetical protein | 3 | 3 | 3 | 3 | 3 | 3 | 3 | 8 |
| 12 | TGME49_299995-t26_1-p1 | hypothetical protein | 3 | 3 | 3 | 3 | 3 | 3 | 3 | 8 |
| 12 | TGME49_300052-t26_1-p1 | hypothetical protein | 3 | 3 | 3 | 3 | 3 | 3 | 3 | 8 |

| | | | | | | | | | | |
|---|---|---|---|---|---|---|---|---|---|---|
| 12 | TGME49_300090-t26_1-p1 | hypothetical protein | 3 | 3 | 3 | 3 | 3 | 3 | 3 | 8 |
| 12 | TGME49_300160.P3458 | hypothetical protein | 3 | 3 | 4 | 3 | 3 | 3 | 3 | 8 |
| 12 | TGME49_300300-t26_1-p1 | hypothetical protein | 3 | 3 | 3 | 3 | 3 | 3 | 3 | 8 |
| 12 | TGME49_300370-t26_1-p1 | hypothetical protein | 3 | 3 | 3 | 3 | 3 | 3 | 3 | 8 |
| 12 | TGME49_300604-t26_1-p1 | ribosomal protein L36 | 4 | 3 | 4 | 3 | 3 | 3 | 3 | 8 |
| 12 | TGME49_300780-t26_1-p1 | hypothetical protein | 4 | 3 | 3 | 3 | 3 | 3 | 3 | 8 |
| 12 | TGME49_300790-t26_1-p1 | hypothetical protein | 3 | 3 | 3 | 3 | 3 | 3 | 3 | 8 |
| 12 | TGME49_300980-t26_1-p1 | hypothetical protein | 3 | 3 | 3 | 3 | 3 | 3 | 3 | 8 |
| 12 | TGME49_300990-t26_1-p1 | Toxoplasma gondii family C protein | 3 | 3 | 3 | 3 | 3 | 3 | 3 | 8 |
| 22 | TGME49_301150-t26_1-p1 | SAG-related sequence SRS19B | 3 | 8 | 3 | 5 | 3 | 3 | 3 | 8 |
| 22 | TGME49_301160-t26_1-p1 | SAG-related sequence SRS19C | 3 | 7 | 3 | 3 | 3 | 3 | 3 | 8 |
| 12 | TGME49_301170-t26_1-p1 | SAG-related sequence SRS19D | 3 | 3 | 3 | 3 | 3 | 3 | 3 | 8 |
| 12 | TGME49_301260-t26_1-p1 | hypothetical protein | 3 | 3 | 3 | 3 | 3 | 3 | 3 | 8 |
| 12 | TGME49_301310-t26_1-p1 | hypothetical protein | 3 | 3 | 4 | 3 | 4 | 3 | 3 | 8 |
| 12 | TGME49_301330-t26_1-p1 | hypothetical protein | 3 | 3 | 3 | 3 | 3 | 3 | 3 | 8 |
| 12 | TGME49_301415-t26_1-p1 | hypothetical protein | 3 | 3 | 3 | 3 | 3 | 3 | 3 | 8 |
| 12 | TGME49_301480-t26_1-p1 | hypothetical protein | 3 | 5 | 3 | 3 | 3 | 3 | 3 | 8 |
| 12 | TGME49_301890-t26_1-p1 | Toxoplasma gondii family B protein | 3 | 3 | 3 | 3 | 3 | 3 | 3 | 8 |
| 12 | TGME49_301996-t26_1-p1 | ORF D | 4 | 3 | 4 | 4 | 4 | 3 | 3 | 8 |
| 12 | TGME49_301998-t26_1-p1 | ORF C, putative | 4 | 3 | 4 | 4 | 4 | 3 | 3 | 8 |
| 12 | TGME49_302005-t26_1-p1 | ORF F | 4 | 3 | 4 | 4 | 4 | 3 | 3 | 8 |
| 12 | TGME49_302007-t26_1-p1 | ribosomal protein L11, putative | 3 | 3 | 5 | 5 | 5 | 3 | 3 | 8 |
| 12 | TGME49_302009-t26_1-p1 | ORF B | 3 | 3 | 3 | 3 | 3 | 3 | 3 | 8 |
| 12 | TGME49_302053-t26_1-p1 | ribosomal protein S7 | 3 | 3 | 3 | 3 | 3 | 3 | 3 | 8 |
| 12 | TGME49_302060-t26_1-p1 | elongation factor tu, apicoplast, putative | 3 | 3 | 3 | 3 | 3 | 3 | 3 | 8 |
| 12 | TGME49_302150-t26_1-p1 | ATPase, AAA family protein | 3 | 3 | 3 | 3 | 3 | 3 | 3 | 8 |
| 12 | TGME49_302450-t26_1-p1 | TFIIH basal transcription factor complex helicase XPB subunit, putative | 3 | 3 | 3 | 3 | 3 | 5 | 3 | 8 |
| 12 | TGME49_303050-t26_1-p1 | zinc finger, c2h2 type family protein | 3 | 3 | 3 | 3 | 3 | 3 | 3 | 8 |
| 12 | TGME49_303450-t26_1-p1 | coatomer gamma subunit | 3 | 3 | 3 | 3 | 3 | 3 | 3 | 8 |
| 12 | TGME49_304690-t26_1-p1 | hypothetical protein | 3 | 3 | 3 | 3 | 3 | 3 | 3 | 8 |
| 12 | TGME49_304755-t26_1-p1 | hypothetical protein | 3 | 3 | 3 | 3 | 3 | 3 | 3 | 8 |
| 12 | TGME49_304870-t26_1-p1 | hypothetical protein | 3 | 3 | 3 | 3 | 3 | 3 | 3 | 8 |
| 22 | TGME49_305230-t26_1-p1 | hypothetical protein | 3 | 9 | 7 | 4 | 4 | 3 | 4 | 8 |
| 12 | TGME49_305485-t26_1-p1 | hypothetical protein | 3 | 3 | 3 | 3 | 3 | 3 | 3 | 8 |
| 12 | TGME49_306010-t26_1-p1 | hypothetical protein | 3 | 3 | 3 | 3 | 3 | 3 | 3 | 8 |
| 12 | TGME49_306035-t26_1-p1 | hypothetical protein | 3 | 3 | 3 | 3 | 3 | 3 | 3 | 8 |
| 12 | TGME49_306455-t26_1-p1 | hypothetical protein | 4 | 3 | 3 | 3 | 3 | 3 | 3 | 8 |
| 12 | TGME49_306688-t26_1-p1 | hypothetical protein | 3 | 3 | 3 | 3 | 3 | 3 | 3 | 8 |
| 12 | TGME49_306692-t26_1-p1 | hypothetical protein | 3 | 3 | 3 | 3 | 3 | 3 | 3 | 8 |
| 12 | TGME49_306700-t26_1-p1 | hypothetical protein | 3 | 3 | 3 | 3 | 3 | 3 | 3 | 8 |
| 12 | TGME49_306710-t26_1-p1 | hypothetical protein | 3 | 3 | 3 | 3 | 3 | 3 | 3 | 8 |
| 12 | TGME49_306720-t26_1-p1 | hypothetical protein | 3 | 3 | 3 | 3 | 3 | 3 | 3 | 8 |
| 12 | TGME49_306730-t26_1-p1 | hypothetical protein | 3 | 3 | 3 | 3 | 3 | 3 | 3 | 8 |
| 12 | TGME49_306860-t26_1-p1 | hypothetical protein | 3 | 3 | 3 | 3 | 3 | 3 | 3 | 8 |
| 12 | TGME49_306870-t26_1-p1 | hypothetical protein | 3 | 3 | 3 | 3 | 3 | 3 | 3 | 8 |
| 12 | TGME49_306880-t26_1-p1 | hypothetical protein | 3 | 3 | 3 | 3 | 3 | 3 | 3 | 8 |
| 12 | TGME49_307015-t26_1-p1 | hypothetical protein | 3 | 3 | 3 | 3 | 3 | 3 | 3 | 8 |
| 12 | TGME49_307045-t26_1-p1 | Toxoplasma gondii family B protein | 3 | 3 | 3 | 3 | 3 | 3 | 3 | 8 |
| 12 | TGME49_307050-t26_1-p1 | hypothetical protein | 3 | 3 | 3 | 3 | 3 | 3 | 3 | 8 |

| | | | | | | | | | | |
|---|---|---|---|---|---|---|---|---|---|---|
| 12 | TGME49_307060-t26_1-p1 | hypothetical protein | 3 | 3 | 3 | 3 | 3 | 3 | 3 | 8 |
| 12 | TGME49_307080-t26_1-p1 | hypothetical protein | 3 | 3 | 3 | 3 | 3 | 3 | 3 | 8 |
| 12 | TGME49_307090-t26_1-p1 | hypothetical protein | 3 | 3 | 3 | 3 | 3 | 3 | 3 | 8 |
| 12 | TGME49_307260-t26_1-p1 | Toxoplasma gondii family C protein | 4 | 3 | 3 | 3 | 3 | 3 | 3 | 8 |
| 12 | TGME49_307270-t26_1-p1 | hypothetical protein | 3 | 3 | 3 | 3 | 3 | 3 | 3 | 8 |
| 12 | TGME49_307450-t26_1-p1 | hypothetical protein | 3 | 3 | 3 | 3 | 3 | 3 | 3 | 8 |
| 12 | TGME49_307460-t26_1-p1 | Toxoplasma gondii family B protein | 3 | 3 | 3 | 3 | 3 | 3 | 3 | 8 |
| 12 | TGME49_307470-t26_1-p1 | hypothetical protein | 3 | 3 | 3 | 3 | 3 | 3 | 3 | 8 |
| 12 | TGME49_307480-t26_1-p1 | Toxoplasma gondii family B protein | 3 | 3 | 3 | 3 | 3 | 3 | 3 | 8 |
| 12 | TGME49_307575-t26_1-p1 | hypothetical protein | 3 | 3 | 3 | 3 | 3 | 3 | 3 | 8 |
| 12 | TGME49_307815-t26_1-p1 | hypothetical protein | 3 | 3 | 3 | 3 | 3 | 3 | 3 | 8 |
| 12 | TGME49_308075-t26_1-p1 | hypothetical protein | 3 | 3 | 3 | 3 | 3 | 3 | 3 | 8 |
| 12 | TGME49_308093-t26_1-p1 | rhoptry kinase family protein (incomplete catalytic triad) | 3 | 3 | 6 | 3 | 3 | 3 | 3 | 8 |
| 12 | TGME49_308820-t26_1-p1 | hypothetical protein | 3 | 3 | 3 | 3 | 3 | 3 | 3 | 8 |
| 12 | TGME49_308945-t26_1-p1 | hypothetical protein | 3 | 3 | 3 | 3 | 3 | 3 | 3 | 8 |
| 12 | TGME49_309025-t26_1-p1 | hypothetical protein | 4 | 3 | 3 | 3 | 3 | 3 | 3 | 8 |
| 12 | TGME49_309095-t26_1-p1 | hypothetical protein | 4 | 3 | 3 | 3 | 3 | 3 | 3 | 8 |
| 12 | TGME49_309130-t26_1-p1 | hypothetical protein | 3 | 3 | 3 | 3 | 3 | 3 | 3 | 8 |
| 12 | TGME49_309150-t26_1-p1 | hypothetical protein | 3 | 3 | 3 | 3 | 3 | 3 | 3 | 8 |
| 12 | TGME49_309185-t26_1-p1 | hypothetical protein | 4 | 3 | 3 | 3 | 3 | 3 | 3 | 8 |
| 12 | TGME49_309230-t26_1-p1 | hypothetical protein | 3 | 3 | 3 | 3 | 3 | 3 | 3 | 8 |
| 12 | TGME49_309300.P3567 | SAG-related sequence SRS55A | 3 | 4 | 4 | 4 | 3 | 3 | 3 | 8 |
| 12 | TGME49_309405-t26_1-p1 | hypothetical protein | 3 | 3 | 3 | 3 | 3 | 3 | 3 | 8 |
| 12 | TGME49_309830.P3577 | hypothetical protein | 3 | 3 | 3 | 3 | 3 | 3 | 3 | 8 |
| 12 | TGME49_310030.P3591 | cyclase-associated protein CAP | 3 | 3 | 3 | 3 | 6 | 5 | 3 | 8 |
| 12 | TGME49_310045-t26_1-p1 | hypothetical protein | 3 | 3 | 3 | 3 | 3 | 3 | 3 | 8 |
| 12 | TGME49_310090-t26_1-p1 | hypothetical protein | 3 | 3 | 3 | 3 | 3 | 3 | 3 | 8 |
| 12 | TGME49_310110-t26_1-p1 | hypothetical protein | 3 | 3 | 3 | 3 | 3 | 3 | 3 | 8 |
| 12 | TGME49_310173-t26_1-p1 | hypothetical protein | 4 | 3 | 3 | 3 | 3 | 3 | 3 | 8 |
| 12 | TGME49_310455-t26_1-p1 | hypothetical protein | 3 | 3 | 3 | 3 | 3 | 3 | 3 | 8 |
| 12 | TGME49_310515-t26_1-p1 | hypothetical protein | 3 | 3 | 3 | 3 | 3 | 3 | 3 | 8 |
| 12 | TGME49_310590-t26_1-p1 | hypothetical protein | 3 | 3 | 3 | 3 | 3 | 3 | 3 | 8 |
| 12 | TGME49_310650-t26_1-p1 | hypothetical protein | 3 | 3 | 3 | 3 | 3 | 3 | 3 | 8 |
| 12 | TGME49_311050-t26_1-p1 | hypothetical protein | 3 | 3 | 3 | 3 | 3 | 3 | 3 | 8 |
| 12 | TGME49_311360.P3644 | protein kinase G AGC kinase family member PKG | 3 | 3 | 3 | 3 | 3 | 3 | 8 | 8 |
| 12 | TGME49_311425-t26_1-p1 | hypothetical protein | 3 | 3 | 3 | 3 | 3 | 3 | 3 | 8 |
| 12 | TGME49_311755.P3654 | plasma membrane protein 62D16 | 3 | 3 | 3 | 4 | 3 | 3 | 3 | 8 |
| 12 | TGME49_311850-t26_1-p1 | hypothetical protein | 3 | 3 | 3 | 3 | 3 | 3 | 3 | 8 |
| 12 | TGME49_312120-t26_1-p1 | hypothetical protein | 3 | 3 | 3 | 3 | 3 | 3 | 3 | 8 |
| 12 | TGME49_312245-t26_1-p1 | hypothetical protein | 3 | 3 | 3 | 3 | 3 | 3 | 3 | 8 |
| 12 | TGME49_312635.P3677 | P-loop NTPase, putative | 3 | 3 | 3 | 3 | 3 | 4 | 3 | 8 |
| 12 | TGME49_312850-t26_1-p1 | hypothetical protein | 3 | 3 | 3 | 3 | 3 | 3 | 3 | 8 |
| 12 | TGME49_312940.P3684 | hypothetical protein | 3 | 4 | 4 | 4 | 4 | 3 | 3 | 8 |
| 12 | TGME49_313055-t26_1-p1 | hypothetical protein | 3 | 3 | 3 | 3 | 3 | 3 | 3 | 8 |
| 12 | TGME49_313250-t26_1-p1 | hypothetical protein | 3 | 3 | 3 | 3 | 3 | 3 | 3 | 8 |
| 12 | TGME49_313277-t26_1-p1 | hypothetical protein | 3 | 3 | 3 | 3 | 3 | 3 | 3 | 8 |
| 12 | TGME49_313305-t26_1-p1 | hypothetical protein | 3 | 3 | 3 | 3 | 3 | 3 | 3 | 8 |
| 12 | TGME49_313322-t26_1-p1 | hypothetical protein | 3 | 3 | 3 | 3 | 3 | 3 | 3 | 8 |
| 12 | TGME49_313418-t26_1-p1 | hypothetical protein | 3 | 3 | 3 | 3 | 3 | 3 | 3 | 8 |

| | | | | | | | | | | |
|---|---|---|---|---|---|---|---|---|---|---|
| 12 | TGME49_313460-t26_1-p1 | hypothetical protein | 3 | 3 | 3 | 3 | 3 | 3 | 3 | 8 |
| 12 | TGME49_313475-t26_1-p1 | hypothetical protein | 3 | 3 | 3 | 3 | 3 | 3 | 3 | 8 |
| 22 | TGME49_313530.P3712 | transmembrane protein 167, putative | 3 | 5 | 5 | 5 | 3 | 3 | 3 | 8 |
| 12 | TGME49_313650.P3716 | hypothetical protein | 4 | 5 | 3 | 4 | 3 | 3 | 3 | 8 |
| 12 | TGME49_313775-t26_1-p1 | hypothetical protein | 3 | 3 | 3 | 3 | 3 | 3 | 3 | 8 |
| 12 | TGME49_313840-t26_1-p1 | hypothetical protein | 3 | 3 | 3 | 3 | 3 | 3 | 3 | 8 |
| 12 | TGME49_314350-t26_1-p1 | hypothetical protein | 3 | 3 | 3 | 3 | 3 | 3 | 3 | 8 |
| 12 | TGME49_314440-t26_1-p1 | hypothetical protein | 3 | 3 | 3 | 3 | 3 | 3 | 3 | 8 |
| 12 | TGME49_314490-t26_1-p1 | hypothetical protein | 3 | 3 | 3 | 3 | 3 | 3 | 3 | 8 |
| 12 | TGME49_314560-t26_1-p1 | hypothetical protein | 3 | 3 | 3 | 3 | 3 | 3 | 3 | 8 |
| 12 | TGME49_314570-t26_1-p1 | hypothetical protein | 3 | 3 | 3 | 3 | 3 | 3 | 3 | 8 |
| 12 | TGME49_314660-t26_1-p1 | TPRX1 protein | 3 | 3 | 3 | 3 | 3 | 3 | 3 | 8 |
| 12 | TGME49_314885-t26_1-p1 | hypothetical protein | 3 | 3 | 3 | 3 | 3 | 3 | 3 | 8 |
| 22 | TGME49_315160.P3757 | rhoptry apical surface protein RASP2 | 3 | 9 | 3 | 3 | 3 | 3 | 3 | 8 |
| 12 | TGME49_315400-t26_1-p1 | SAG-related sequence SRS53D | 3 | 3 | 3 | 3 | 3 | 3 | 3 | 8 |
| 12 | TGME49_315495.P3774 | hypothetical protein | 4 | 3 | 3 | 3 | 3 | 3 | 3 | 8 |
| 12 | TGME49_315570.P3780 | histone H1-like protein | 3 | 3 | 3 | 3 | 3 | 3 | 3 | 8 |
| 12 | TGME49_315705-t26_1-p1 | hypothetical protein | 3 | 3 | 3 | 3 | 3 | 3 | 3 | 8 |
| 12 | TGME49_315715-t26_1-p1 | hypothetical protein | 3 | 3 | 3 | 3 | 3 | 3 | 3 | 8 |
| 12 | TGME49_315855-t26_1-p1 | hypothetical protein | 3 | 3 | 3 | 3 | 3 | 3 | 3 | 8 |
| 12 | TGME49_316210-t26_1-p1 | hypothetical protein | 3 | 3 | 3 | 3 | 3 | 3 | 3 | 8 |
| 12 | TGME49_316550-t26_1-p1 | hypothetical protein | 4 | 4 | 3 | 3 | 3 | 3 | 3 | 8 |
| 12 | TGME49_317830-t26_1-p1 | hypothetical protein | 3 | 3 | 3 | 3 | 3 | 3 | 3 | 8 |
| 12 | TGME49_318120.P3819 | hypothetical protein | 3 | 3 | 3 | 3 | 3 | 3 | 3 | 8 |
| 12 | TGME49_318300-t26_1-p1 | hypothetical protein | 3 | 3 | 3 | 3 | 3 | 3 | 3 | 8 |
| 12 | TGME49_318360-t26_1-p1 | hypothetical protein | 3 | 3 | 3 | 3 | 3 | 3 | 3 | 8 |
| 12 | TGME49_318632-t26_1-p1 | hypothetical protein | 3 | 3 | 3 | 3 | 3 | 3 | 3 | 8 |
| 12 | TGME49_318880-t26_1-p1 | hypothetical protein | 3 | 3 | 3 | 3 | 3 | 3 | 3 | 8 |
| 12 | TGME49_319935-t26_1-p1 | hypothetical protein | 3 | 3 | 3 | 3 | 3 | 3 | 3 | 8 |
| 12 | TGME49_320135-t26_1-p1 | hypothetical protein | 3 | 3 | 3 | 3 | 3 | 3 | 3 | 8 |
| 12 | TGME49_320190-t26_1-p1 | SAG-related sequence SRS16B | 3 | 3 | 3 | 3 | 3 | 3 | 3 | 8 |
| 22 | TGME49_320240-t26_1-p1 | SAG-related sequence SRS15B | 3 | 7 | 6 | 3 | 3 | 3 | 3 | 8 |
| 12 | TGME49_320410-t26_1-p1 | hypothetical protein | 3 | 3 | 3 | 3 | 3 | 3 | 3 | 8 |
| 12 | TGME49_320660-t26_1-p1 | hypothetical protein | 4 | 3 | 3 | 3 | 3 | 3 | 3 | 8 |
| 12 | TGME49_320770-t26_1-p1 | Toxoplasma gondii family B protein | 3 | 3 | 3 | 3 | 3 | 3 | 3 | 8 |
| 12 | TGME49_320780-t26_1-p1 | hypothetical protein | 3 | 3 | 3 | 3 | 3 | 3 | 3 | 8 |
| 12 | TGME49_321170-t26_1-p1 | Toxoplasma gondii family C protein | 3 | 3 | 3 | 3 | 3 | 3 | 3 | 8 |
| 22 | TGME49_321370.P3893 | hypothetical protein | 3 | 7 | 5 | 4 | 4 | 3 | 3 | 8 |
| 12 | TGME49_321460-t26_1-p1 | hypothetical protein | 4 | 3 | 3 | 3 | 3 | 3 | 3 | 8 |
| 22 | TGME49_321470-t26_1-p1 | SAG-related sequence SRS12D | 3 | 7 | 6 | 5 | 3 | 3 | 3 | 8 |
| 22 | TGME49_321480-t26_1-p1 | SAG-related sequence SRS12B | 3 | 7 | 6 | 3 | 3 | 3 | 3 | 8 |
| 12 | TGME49_321685-t26_1-p1 | hypothetical protein | 3 | 3 | 3 | 3 | 3 | 3 | 3 | 8 |
| 12 | TGME49_321700-t26_1-p1 | Kinase-like domain-containing protein [Source:UniProtKB/TrEMBL;Acc:S8F8L7] | 3 | 3 | 3 | 3 | 3 | 3 | 3 | 8 |
| 12 | TGME49_321710-t26_1-p1 | Kinase-like domain-containing protein [Source:UniProtKB/TrEMBL;Acc:S8FBM1] | 3 | 3 | 3 | 3 | 3 | 3 | 3 | 8 |
| 12 | TGME49_321800-t26_1-p1 | EGF family domain-containing protein | 3 | 3 | 3 | 3 | 3 | 3 | 3 | 8 |
| 12 | TGME49_321900-t26_1-p1 | fructose-1,6-bisphosphate aldolase | 3 | 3 | 3 | 3 | 3 | 3 | 3 | 8 |
| 12 | TGME49_322000-t26_1-p1 | myosin-light-chain kinase | 3 | 3 | 4 | 3 | 3 | 3 | 3 | 8 |
| 12 | TGME49_322010-t26_1-p1 | myosin-light-chain kinase | 3 | 3 | 3 | 3 | 3 | 3 | 4 | 8 |

| | | | | | | | | | | |
|---|---|---|---|---|---|---|---|---|---|---|
| 12 | TGME49_322020-t26_1-p1 | hypothetical protein | 3 | 3 | 3 | 3 | 3 | 3 | 3 | 8 |
| 12 | TGME49_322030-t26_1-p1 | hypothetical protein | 3 | 3 | 3 | 3 | 3 | 3 | 3 | 8 |
| 12 | TGME49_322100-t26_1-p1 | myosin-light-chain kinase | 3 | 3 | 3 | 3 | 3 | 4 | 4 | 8 |
| 12 | TGME49_322110-t26_1-p1 | hypothetical protein | 3 | 3 | 3 | 3 | 3 | 3 | 3 | 8 |
| 12 | TGME49_322120-t26_1-p1 | rhoptry kinase family protein | 3 | 3 | 3 | 3 | 3 | 3 | 3 | 8 |
| 12 | TGME49_322130-t26_1-p1 | hypothetical protein | 3 | 3 | 3 | 3 | 3 | 3 | 3 | 8 |
| 12 | TGME49_322200-t26_1-p1 | apocytochrome b, putative | 4 | 3 | 5 | 3 | 3 | 3 | 3 | 8 |
| 12 | TGME49_322300-t26_1-p1 | hypothetical protein | 3 | 3 | 3 | 3 | 3 | 3 | 3 | 8 |
| 12 | TGME49_322400-t26_1-p1 | hypothetical protein | 3 | 3 | 3 | 3 | 3 | 3 | 3 | 8 |
| 12 | TGME49_322600-t26_1-p1 | cytochrome c oxidase subunit iii subfamily protein | 3 | 3 | 5 | 3 | 3 | 3 | 3 | 8 |
| 12 | TGME49_322700-t26_1-p1 | collagen, putative | 3 | 3 | 3 | 3 | 3 | 3 | 3 | 8 |
| 12 | TGME49_322800-t26_1-p1 | hypothetical protein | 3 | 3 | 3 | 3 | 3 | 3 | 3 | 8 |
| 12 | TGME49_322900-t26_1-p1 | hypothetical protein | 3 | 3 | 3 | 3 | 3 | 3 | 3 | 8 |
| 12 | TGME49_323000-t26_1-p1 | KRUF family protein | 4 | 3 | 3 | 3 | 3 | 3 | 3 | 8 |
| 12 | TGME49_323100-t26_1-p1 | hypothetical protein | 3 | 3 | 3 | 3 | 3 | 3 | 3 | 8 |
| 22 | TGME49_323110-t26_1-p1 | hypothetical protein | 3 | 6 | 3 | 3 | 3 | 3 | 3 | 8 |
| 12 | TGME49_323200-t26_1-p1 | OTU family cysteine protease | 3 | 3 | 3 | 3 | 3 | 3 | 3 | 8 |
| 12 | TGME49_323300-t26_1-p1 | myosin-light-chain kinase | 3 | 3 | 3 | 3 | 3 | 3 | 3 | 8 |
| 12 | TGME49_323310-t26_1-p1 | hypothetical protein | 3 | 3 | 3 | 3 | 3 | 3 | 3 | 8 |
| 12 | TGME49_323320-t26_1-p1 | hypothetical protein | 3 | 3 | 3 | 3 | 3 | 3 | 3 | 8 |
| 12 | TGME49_323330-t26_1-p1 | hypothetical protein | 3 | 3 | 3 | 3 | 3 | 3 | 3 | 8 |
| 12 | TGME49_323400-t26_1-p1 | cytochrome c oxidase subunit iii subfamily protein | 3 | 3 | 6 | 3 | 3 | 3 | 3 | 8 |
| 12 | TGME49_323500-t26_1-p1 | hypothetical protein | 3 | 3 | 3 | 3 | 3 | 3 | 3 | 8 |
| 12 | TGME49_323600-t26_1-p1 | OTU family cysteine protease | 3 | 3 | 3 | 3 | 3 | 3 | 3 | 8 |
| 12 | TGME49_323700-t26_1-p1 | OTU family cysteine protease | 3 | 3 | 3 | 3 | 3 | 3 | 3 | 8 |
| 22 | TGME49_323800-t26_1-p1 | OTU family cysteine protease | 4 | 7 | 6 | 3 | 3 | 3 | 3 | 8 |
| 12 | TGME49_323900-t26_1-p1 | hypothetical protein | 3 | 3 | 3 | 3 | 3 | 3 | 3 | 8 |
| 12 | TGME49_324000-t26_1-p1 | dynein heavy chain, n-terminal region 2 protein | 3 | 3 | 3 | 3 | 3 | 3 | 3 | 8 |
| 12 | TGME49_324100-t26_1-p1 | hypothetical protein | 3 | 3 | 3 | 3 | 3 | 3 | 3 | 8 |
| 12 | TGME49_324200-t26_1-p1 | hypothetical protein | 3 | 3 | 3 | 3 | 3 | 3 | 3 | 8 |
| 12 | TGME49_324300-t26_1-p1 | transitional endoplasmic reticulum ATPase, putative | 3 | 3 | 3 | 3 | 3 | 3 | 3 | 8 |
| 12 | TGME49_324400-t26_1-p1 | hypothetical protein | 3 | 3 | 3 | 3 | 3 | 3 | 3 | 8 |
| 12 | TGME49_324500-t26_1-p1 | nuclear distribution protein C, putative | 3 | 3 | 3 | 3 | 3 | 3 | 3 | 8 |
| 12 | TGME49_324700-t26_1-p1 | hypothetical protein | 3 | 3 | 3 | 3 | 3 | 3 | 3 | 8 |
| 12 | TGME49_324800-t26_1-p1 | tryptophanyl-tRNA synthetase | 3 | 3 | 3 | 3 | 3 | 3 | 3 | 8 |
| 12 | TGME49_324900-t26_1-p1 | STAS domain-containing protein | 3 | 3 | 3 | 3 | 3 | 3 | 3 | 8 |
| 12 | TGME49_325000-t26_1-p1 | hypothetical protein | 3 | 3 | 3 | 3 | 3 | 3 | 3 | 8 |
| 12 | TGME49_325100-t26_1-p1 | hypothetical protein | 3 | 3 | 3 | 3 | 3 | 3 | 3 | 8 |
| 12 | TGME49_325200-t26_1-p1 | pyridoxal-phosphate dependent superfamily protein | 3 | 3 | 3 | 3 | 3 | 3 | 3 | 8 |
| 12 | TGME49_325300-t26_1-p1 | proteasome subunit alpha type, putative | 3 | 3 | 3 | 3 | 3 | 3 | 3 | 8 |
| 12 | TGME49_325400-t26_1-p1 | ribosomal L37ae family protein | 3 | 3 | 3 | 3 | 3 | 3 | 3 | 8 |
| 12 | TGME49_325500-t26_1-p1 | ATP-binding Mrp/Nbp35 family protein | 3 | 3 | 3 | 3 | 3 | 3 | 3 | 8 |
| 12 | TGME49_325600-t26_1-p1 | hypothetical protein | 3 | 3 | 3 | 3 | 3 | 3 | 3 | 8 |
| 12 | TGME49_325700-t26_1-p1 | ATP-binding protein | 3 | 3 | 3 | 3 | 3 | 3 | 3 | 8 |
| 12 | TGME49_325800-t26_1-p1 | EGF family domain-containing protein | 3 | 3 | 4 | 5 | 3 | 3 | 3 | 8 |
| 12 | TGME49_325900-t26_1-p1 | zinc finger (CCCH type) motif-containing protein | 3 | 3 | 3 | 3 | 3 | 3 | 3 | 8 |
| 12 | TGME49_326000-t26_1-p1 | RNA binding protein, putative | 3 | 3 | 3 | 3 | 3 | 3 | 3 | 8 |
| 12 | TGME49_326100-t26_1-p1 | dynamin gtpase | 3 | 3 | 3 | 3 | 3 | 3 | 3 | 8 |
| 12 | TGME49_326300-t26_1-p1 | hypothetical protein | 3 | 3 | 3 | 3 | 3 | 3 | 3 | 8 |
| 12 | TGME49_326400-t26_1-p1 | ribosomal protein L10-like, putative | 3 | 3 | 3 | 3 | 3 | 3 | 3 | 8 |

| | | | | | | | | | | |
|---|---|---|---|---|---|---|---|---|---|---|
| 12 | TGME49_326500-t26_1-p1 | hypothetical protein | 3 | 3 | 3 | 3 | 3 | 3 | 3 | 8 |
| 12 | TGME49_326600-t26_1-p1 | histone, putative | 3 | 3 | 3 | 3 | 3 | 3 | 3 | 8 |
| 12 | TGME49_326700-t26_1-p1 | DNA-directed RNA polymerase, putative | 3 | 3 | 3 | 3 | 3 | 3 | 3 | 8 |
| 12 | TGME49_326800-t26_1-p1 | S-Adenosyl homocysteine hydrolase | 3 | 3 | 3 | 3 | 3 | 3 | 3 | 8 |
| 12 | TGME49_326900-t26_1-p1 | hypothetical protein | 3 | 3 | 3 | 3 | 3 | 3 | 3 | 8 |
| 12 | TGME49_327000-t26_1-p1 | U5 small nuclear ribonucleoprotein component (U5 snRNP), putative | 3 | 3 | 3 | 3 | 3 | 3 | 3 | 8 |
| 12 | TGME49_327100-t26_1-p1 | U5 small nuclear ribonuclear protein, putative | 3 | 3 | 3 | 3 | 3 | 3 | 3 | 8 |
| 12 | TGME49_327200-t26_1-p1 | rhoptry neck protein, putative | 3 | 3 | 3 | 3 | 3 | 3 | 3 | 8 |
| 12 | TGME49_327300-t26_1-p1 | Toxoplasma gondii family A protein | 3 | 3 | 3 | 3 | 3 | 3 | 3 | 8 |
| 12 | TGME49_327400-t26_1-p1 | dynein heavy chain family protein | 3 | 3 | 3 | 3 | 3 | 3 | 3 | 8 |
| 12 | TGME49_327500-t26_1-p1 | hypothetical protein | 3 | 3 | 3 | 3 | 3 | 3 | 3 | 8 |
| 12 | TGME49_327600-t26_1-p1 | exoribonuclease, putative | 3 | 3 | 3 | 3 | 3 | 3 | 3 | 8 |
| 12 | TGME49_327700-t26_1-p1 | hypothetical protein | 3 | 3 | 3 | 3 | 3 | 3 | 3 | 8 |
| 12 | TGME49_327800-t26_1-p1 | dynein-1-alpha heavy chain, flagellar inner arm I1 complex, putative | 3 | 3 | 3 | 3 | 3 | 3 | 3 | 8 |
| 12 | TGME49_327900-t26_1-p1 | hypothetical protein | 3 | 3 | 3 | 3 | 3 | 3 | 3 | 8 |
| 12 | TGME49_328000-t26_1-p1 | hypothetical protein | 3 | 3 | 3 | 3 | 3 | 3 | 3 | 8 |
| 12 | TGME49_328100-t26_1-p1 | RNA recognition motif-containing protein | 3 | 3 | 3 | 3 | 3 | 3 | 3 | 8 |
| 12 | TGME49_328200-t26_1-p1 | phosphatidylinositol 4-kinase | 3 | 3 | 5 | 3 | 3 | 3 | 3 | 8 |
| 12 | TGME49_328300-t26_1-p1 | CCR4-NOT transcription complex subunit 4, putative | 3 | 3 | 3 | 3 | 3 | 3 | 3 | 8 |
| 12 | TGME49_328400-t26_1-p1 | aminotransferase, class V family protein | 3 | 3 | 3 | 3 | 3 | 3 | 3 | 8 |
| 12 | TGME49_328500-t26_1-p1 | hypothetical protein | 3 | 3 | 3 | 3 | 3 | 3 | 3 | 8 |
| 12 | TGME49_328600-t26_1-p1 | Isoleucine-tRNA ligase | 3 | 3 | 3 | 3 | 3 | 3 | 3 | 8 |
| 12 | TGME49_328700-t26_1-p1 | hypothetical protein | 3 | 3 | 3 | 3 | 3 | 3 | 3 | 8 |
| 12 | TGME49_328800-t26_1-p1 | hypothetical protein | 3 | 3 | 3 | 3 | 3 | 3 | 3 | 8 |
| 12 | TGME49_328900-t26_1-p1 | hypothetical protein | 3 | 3 | 3 | 3 | 3 | 3 | 3 | 8 |
| 12 | TGME49_329000-t26_1-p1 | hypothetical protein | 3 | 3 | 3 | 3 | 3 | 3 | 3 | 8 |
| 12 | TGME49_329100-t26_1-p1 | Toxoplasma gondii family C protein | 3 | 3 | 3 | 3 | 3 | 3 | 3 | 8 |
| 12 | TGME49_329200-t26_1-p1 | hypothetical protein | 3 | 3 | 3 | 3 | 3 | 3 | 3 | 8 |
| 12 | TGME49_329300-t26_1-p1 | hypothetical protein | 3 | 3 | 3 | 3 | 3 | 3 | 3 | 8 |
| 12 | TGME49_329400-t26_1-p1 | Toxoplasma gondii family C protein | 3 | 3 | 3 | 3 | 3 | 3 | 3 | 8 |
| 12 | TGME49_329500-t26_1-p1 | hypothetical protein | 3 | 3 | 3 | 3 | 3 | 3 | 3 | 8 |
| 12 | TGME49_329600-t26_1-p1 | hypothetical protein | 3 | 3 | 3 | 3 | 3 | 3 | 3 | 8 |
| 12 | TGME49_329620-t26_1-p1 | hypothetical protein | 3 | 3 | 3 | 3 | 3 | 3 | 3 | 8 |
| 12 | TGME49_329700-t26_1-p1 | SAG-related sequence SRS26J | 3 | 4 | 4 | 4 | 3 | 3 | 3 | 8 |
| 12 | TGME49_500000.P3 | hypothetical protein | 3 | 3 | 3 | 3 | 3 | 3 | 3 | 8 |
| 12 | TGME49_500002.P5 | hypothetical protein, conserved | 3 | 3 | 3 | 3 | 3 | 3 | 3 | 8 |
| 12 | TGME49_500003.P6 | hypothetical protein, conserved | 3 | 3 | 3 | 3 | 3 | 3 | 3 | 8 |
| 12 | TGME49_500006.P9 | hypothetical protein, conserved | 3 | 3 | 5 | 3 | 3 | 3 | 3 | 8 |
| 22 | TGME49_500010.P13 | hypothetical protein, conserved | 3 | 7 | 3 | 6 | 3 | 3 | 3 | 8 |
| 12 | TGME49_500011.P14 | hypothetical protein, conserved | 3 | 3 | 3 | 3 | 3 | 3 | 3 | 8 |
| 12 | TGME49_500014.P17 | hypothetical protein, conserved | 3 | 3 | 4 | 4 | 3 | 3 | 3 | 8 |
| 12 | TGME49_500016.P19 | hypothetical protein, conserved | 3 | 3 | 3 | 3 | 3 | 3 | 3 | 8 |
| 12 | TGME49_500018.P21 | hypothetical protein, conserved | 3 | 3 | 3 | 4 | 3 | 3 | 3 | 8 |
| 12 | TGME49_500019.P22 | hypothetical protein, conserved | 3 | 3 | 6 | 3 | 3 | 3 | 3 | 8 |
| 12 | TGME49_500020.P23 | SAM domain-containing protein | 3 | 3 | 3 | 3 | 3 | 3 | 3 | 8 |
| 12 | TGME49_500021.P24 | hypothetical protein, conserved | 3 | 3 | 6 | 3 | 3 | 3 | 3 | 8 |
| 12 | TGME49_500023.P26 | hypothetical protein, conserved | 3 | 3 | 4 | 3 | 3 | 3 | 3 | 8 |

| | | | | | | | | | | |
|---|---|---|---|---|---|---|---|---|---|---|
| 22 | TGME49_500024.P27 | mitochondrial import inner membrane translocase subunit TIM9, putative | 3 | 5 | 5 | 5 | 3 | 3 | 4 | 8 |
| 12 | TGME49_500026.P29 | Toxoplasma gondii family B protein | 3 | 3 | 3 | 3 | 3 | 3 | 3 | 8 |
| 12 | TGME49_500027.P30 | hypothetical protein, conserved | 3 | 3 | 3 | 3 | 3 | 3 | 3 | 8 |
| 12 | TGME49_500028.P31 | hypothetical protein, conserved | 3 | 3 | 3 | 3 | 3 | 3 | 3 | 8 |
| 12 | TGME49_500033.P36 | hypothetical protein, conserved | 3 | 3 | 6 | 3 | 3 | 3 | 3 | 8 |
| 12 | TGME49_500034.P37 | hypothetical protein, conserved | 3 | 3 | 3 | 3 | 3 | 3 | 3 | 8 |
| 12 | TGME49_500035.P38 | hypothetical protein, conserved | 4 | 3 | 5 | 4 | 3 | 3 | 3 | 8 |
| 22 | TGME49_500036.P39 | RNA binding protein, putative | 3 | 5 | 5 | 5 | 4 | 3 | 4 | 8 |
| 12 | TGME49_500038.P1 | hypothetical protein, conserved | 3 | 3 | 3 | 3 | 3 | 4 | 4 | 8 |
| 12 | TGME49_500043.P343 | hypothetical protein, conserved | 3 | 3 | 3 | 3 | 3 | 3 | 3 | 8 |
| 12 | TGME49_500044.P344 | 3'5'-cyclic nucleotide phosphodiesterase PDE16 | 3 | 3 | 3 | 3 | 6 | 3 | 3 | 8 |
| 12 | TGME49_500049.P350 | hypothetical protein, conserved | 3 | 3 | 3 | 4 | 3 | 3 | 4 | 8 |
| 12 | TGME49_500050.P351 | hypothetical protein, conserved | 3 | 3 | 3 | 3 | 3 | 3 | 3 | 8 |
| 12 | TGME49_500056.P357 | selenoprotein W, putative | 4 | 3 | 3 | 4 | 4 | 3 | 3 | 8 |
| 12 | TGME49_500060.P361 | dynein light chain, putative | 3 | 3 | 3 | 3 | 3 | 3 | 3 | 8 |
| 12 | TGME49_500063.P364 | hypothetical protein, conserved | 3 | 3 | 3 | 3 | 3 | 3 | 3 | 8 |
| 22 | TGME49_500066.P367 | DUF501 domain-containing protein | 3 | 7 | 7 | 5 | 3 | 3 | 4 | 8 |
| 22 | TGME49_500067.P368 | hypothetical protein, conserved | 3 | 6 | 6 | 5 | 4 | 3 | 3 | 8 |
| 12 | TGME49_500073.P374 | hypothetical protein, conserved | 3 | 3 | 3 | 3 | 3 | 3 | 3 | 8 |
| 22 | TGME49_500074.P375 | DUF4200 domain-containing protein | 3 | 9 | 8 | 3 | 3 | 5 | 4 | 8 |
| 12 | TGME49_500078.P379 | REJ domain-containing protein | 3 | 3 | 3 | 4 | 3 | 4 | 3 | 8 |
| 12 | TGME49_500084.P385 | hypothetical protein, conserved | 3 | 3 | 4 | 3 | 3 | 3 | 3 | 8 |
| 12 | TGME49_500090.P391 | hypothetical protein, conserved | 3 | 3 | 4 | 3 | 3 | 3 | 4 | 8 |
| 12 | TGME49_500091.P392 | leucine-rich repeat containing protein | 3 | 3 | 3 | 3 | 3 | 4 | 3 | 8 |
| 12 | TGME49_500095.P396 | hypothetical protein, conserved | 3 | 3 | 5 | 3 | 3 | 3 | 3 | 8 |
| 12 | TGME49_500096.P397 | hypothetical protein, conserved | 3 | 3 | 3 | 4 | 4 | 3 | 4 | 8 |
| 22 | TGME49_500097.P398 | hypothetical protein, conserved | 3 | 5 | 5 | 3 | 3 | 3 | 3 | 8 |
| 12 | TGME49_500102.P403 | hypothetical protein, conserved | 3 | 3 | 3 | 3 | 3 | 3 | 3 | 8 |
| 12 | TGME49_500103.P404 | hypothetical protein | 3 | 3 | 3 | 4 | 3 | 3 | 3 | 8 |
| 12 | TGME49_500104.P405 | hypothetical protein | 3 | 3 | 3 | 3 | 3 | 3 | 3 | 8 |
| 12 | TGME49_500105.P406 | hypothetical protein, conserved | 3 | 3 | 3 | 3 | 3 | 3 | 3 | 8 |
| 12 | TGME49_500107.P408 | hypothetical protein, conserved | 4 | 3 | 3 | 3 | 3 | 3 | 3 | 8 |
| 22 | TGME49_500110.P411 | sentrin-specific protease 8, putative | 3 | 8 | 3 | 3 | 3 | 3 | 3 | 8 |
| 12 | TGME49_500113.P414 | WD repeat-containing protein | 3 | 3 | 3 | 3 | 3 | 3 | 3 | 8 |
| 22 | TGME49_500120.P422 | hypothetical protein, conserved | 3 | 6 | 4 | 3 | 4 | 3 | 3 | 8 |
| 12 | TGME49_500121.P423 | dephospho-CoA kinase | 3 | 3 | 3 | 3 | 3 | 3 | 3 | 8 |
| 12 | TGME49_500122.P424 | hypothetical protein, conserved | 4 | 3 | 3 | 3 | 3 | 3 | 3 | 8 |
| 12 | TGME49_500126.P428 | hypothetical protein, conserved | 4 | 3 | 4 | 3 | 3 | 3 | 3 | 8 |
| 12 | TGME49_500129.P431 | ankyrin repeat-containing protein | 3 | 3 | 3 | 3 | 3 | 3 | 4 | 8 |
| 22 | TGME49_500130.P432 | SND2 domain-containing protein | 3 | 6 | 6 | 3 | 3 | 4 | 4 | 8 |
| 12 | TGME49_500131.P433 | hypothetical protein, conserved | 3 | 3 | 3 | 4 | 3 | 3 | 3 | 8 |
| 12 | TGME49_500138.P440 | hypothetical protein | 3 | 3 | 3 | 3 | 3 | 3 | 3 | 8 |
| 22 | TGME49_500140.P442 | hypothetical protein, conserved | 3 | 7 | 3 | 4 | 3 | 3 | 3 | 8 |
| 12 | TGME49_500141.P443 | hypothetical protein, conserved | 3 | 3 | 4 | 3 | 3 | 3 | 3 | 8 |
| 12 | TGME49_500149.P153 | hypothetical protein, conserved | 3 | 3 | 3 | 3 | 3 | 3 | 3 | 8 |
| 12 | TGME49_500157.P161 | prefoldin, putative | 4 | 3 | 3 | 3 | 3 | 3 | 3 | 8 |
| 12 | TGME49_500158.P162 | calmodulin-binding protein, putative | 3 | 3 | 3 | 3 | 4 | 4 | 3 | 8 |
| 12 | TGME49_500159.P163 | hypothetical protein | 3 | 3 | 3 | 3 | 3 | 3 | 3 | 8 |
| 12 | TGME49_500164.P168 | RIIa domain-containing protein | 3 | 3 | 3 | 3 | 3 | 4 | 3 | 8 |

| | | | | | | | | | | |
|---|---|---|---|---|---|---|---|---|---|---|
| 12 | TGME49_500173.P177 | hypothetical protein, conserved | 3 | 3 | 3 | 3 | 3 | 3 | 3 | 8 |
| 22 | TGME49_500175.P179 | ankyrin repeat-containing protein | 3 | 7 | 6 | 6 | 3 | 5 | 4 | 8 |
| 12 | TGME49_500178.P182 | hypothetical protein, conserved | 3 | 3 | 3 | 3 | 3 | 3 | 3 | 8 |
| 12 | TGME49_500180.P184 | dynein light chain, putative | 3 | 3 | 4 | 5 | 3 | 3 | 3 | 8 |
| 12 | TGME49_500184.P188 | Toxoplasma gondii family C protein | 3 | 3 | 3 | 3 | 3 | 3 | 3 | 8 |
| 22 | TGME49_500191.P195 | hypothetical protein, conserved | 3 | 7 | 3 | 3 | 3 | 3 | 3 | 8 |
| 12 | TGME49_500194.P198 | hypothetical protein, conserved | 3 | 3 | 3 | 3 | 3 | 3 | 3 | 8 |
| 12 | TGME49_500195.P199 | hypothetical protein, conserved | 3 | 3 | 3 | 3 | 3 | 3 | 3 | 8 |
| 12 | TGME49_500196.P200 | hypothetical protein, conserved | 4 | 3 | 4 | 3 | 3 | 3 | 3 | 8 |
| 12 | TGME49_500197.P201 | intraflagellar transport protein 20, putative | 3 | 3 | 3 | 3 | 3 | 3 | 3 | 8 |
| 12 | TGME49_500201.P206 | hypothetical protein, conserved | 3 | 3 | 4 | 3 | 3 | 3 | 3 | 8 |
| 12 | TGME49_500203.P208 | cytochrome c oxidase subunit 6B, putative | 3 | 3 | 3 | 3 | 3 | 3 | 3 | 8 |
| 12 | TGME49_500205.P210 | hypothetical protein, conserved | 3 | 4 | 3 | 3 | 4 | 4 | 3 | 8 |
| 12 | TGME49_500208.P213 | hypothetical protein, conserved | 3 | 3 | 3 | 3 | 3 | 3 | 3 | 8 |
| 12 | TGME49_500211.P216 | coiled-coil domain-containing protein 103, putative | 3 | 3 | 3 | 3 | 3 | 4 | 4 | 8 |
| 12 | TGME49_500212.P217 | hypothetical protein, conserved | 3 | 3 | 3 | 3 | 3 | 3 | 3 | 8 |
| 12 | TGME49_500213.P218 | LisH domain-containing protein, putative | 3 | 3 | 5 | 3 | 4 | 4 | 3 | 8 |
| 12 | TGME49_500214.P219 | hypothetical protein, conserved | 3 | 3 | 3 | 3 | 3 | 3 | 3 | 8 |
| 12 | TGME49_500215.P220 | hypothetical protein, conserved | 3 | 3 | 4 | 3 | 3 | 3 | 3 | 8 |
| 12 | TGME49_500217.P222 | hypothetical protein | 3 | 3 | 3 | 3 | 3 | 4 | 3 | 8 |
| 12 | TGME49_500218.P223 | Toxoplasma gondii family C protein | 3 | 3 | 3 | 3 | 3 | 3 | 3 | 8 |
| 12 | TGME49_500219.P224 | zinc finger protein, putative | 3 | 3 | 3 | 3 | 3 | 3 | 3 | 8 |
| 12 | TGME49_500224.P230 | GAF domain-containing protein | 3 | 3 | 3 | 3 | 3 | 3 | 3 | 8 |
| 12 | TGME49_500225.P231 | hypothetical protein, conserved | 3 | 3 | 3 | 6 | 4 | 3 | 3 | 8 |
| 22 | TGME49_500228.P234 | 30S ribosomal protein S10, putative | 3 | 8 | 8 | 6 | 5 | 4 | 4 | 8 |
| 12 | TGME49_500229.P235 | hypothetical protein, conserved | 3 | 4 | 3 | 3 | 3 | 3 | 3 | 8 |
| 12 | TGME49_500233.P239 | hypothetical protein, conserved | 3 | 3 | 3 | 4 | 4 | 4 | 4 | 8 |
| 12 | TGME49_500234.P240 | IQ calmodulin domain-containing protein | 3 | 3 | 3 | 3 | 3 | 3 | 3 | 8 |
| 22 | TGME49_500238.P244 | armadillo repeat-containing protein | 3 | 9 | 3 | 3 | 3 | 4 | 4 | 8 |
| 22 | TGME49_500239.P245 | hypothetical protein, conserved | 3 | 9 | 3 | 3 | 3 | 3 | 3 | 8 |
| 12 | TGME49_500240.P246 | hypothetical protein, conserved | 3 | 3 | 3 | 3 | 3 | 3 | 3 | 8 |
| 12 | TGME49_500242.P248 | hypothetical protein, conserved | 3 | 3 | 3 | 3 | 3 | 3 | 3 | 8 |
| 12 | TGME49_500243.P249 | hypothetical protein, conserved | 3 | 3 | 7 | 3 | 3 | 3 | 3 | 8 |
| 12 | TGME49_500244.P250 | RIB43A-like with coiled-coils protein, putative | 4 | 3 | 3 | 3 | 3 | 5 | 3 | 8 |
| 12 | TGME49_500249.P255 | cAMP-dependent protein kinase regulatory subunit, putative | 4 | 3 | 4 | 3 | 4 | 4 | 4 | 8 |
| 12 | TGME49_500250.P256 | hypothetical protein, conserved | 3 | 3 | 3 | 3 | 3 | 3 | 3 | 8 |
| 12 | TGME49_500251.P257 | hypothetical protein, conserved | 3 | 3 | 3 | 3 | 3 | 3 | 3 | 8 |
| 12 | TGME49_500255.P44 | hypothetical protein, conserved | 3 | 3 | 3 | 3 | 3 | 3 | 3 | 8 |
| 12 | TGME49_500257.P46 | EF hand domain-containing protein | 3 | 3 | 5 | 3 | 3 | 3 | 4 | 8 |
| 12 | TGME49_500259.P48 | hypothetical protein, conserved | 3 | 3 | 3 | 3 | 3 | 3 | 3 | 8 |
| 22 | TGME49_500262.P51 | hypothetical protein, conserved | 3 | 10 | 7 | 3 | 3 | 3 | 3 | 8 |
| 12 | TGME49_500265.P54 | hypothetical protein, conserved | 3 | 3 | 3 | 3 | 3 | 3 | 3 | 8 |
| 12 | TGME49_500266.P55 | hypothetical protein, conserved | 3 | 3 | 3 | 3 | 3 | 3 | 3 | 8 |
| 12 | TGME49_500268.P57 | hypothetical protein, conserved | 3 | 3 | 3 | 3 | 3 | 3 | 3 | 8 |
| 12 | TGME49_500274.P63 | hypothetical protein, conserved | 3 | 3 | 3 | 3 | 3 | 3 | 3 | 8 |
| 12 | TGME49_500278.P67 | hypothetical protein, conserved | 3 | 3 | 3 | 4 | 3 | 3 | 3 | 8 |
| 12 | TGME49_500280.P69 | hypothetical protein, conserved | 3 | 3 | 3 | 3 | 3 | 3 | 3 | 8 |
| 12 | TGME49_500281.P70 | hypothetical protein | 3 | 3 | 3 | 3 | 3 | 3 | 3 | 8 |
| 12 | TGME49_500285.P74 | CPBP intramembrane metalloprotease, putative | 3 | 3 | 5 | 5 | 3 | 3 | 3 | 8 |

| | | | | | | | | | | |
|---|---|---|---|---|---|---|---|---|---|---|
| 12 | TGME49_500286.P75 | IQ calmodulin-binding protein, putative | 3 | 3 | 3 | 3 | 3 | 4 | 3 | 8 |
| 12 | TGME49_500288.P77 | hypothetical protein, conserved | 3 | 3 | 5 | 4 | 3 | 3 | 4 | 8 |
| 12 | TGME49_500289.P78 | hypothetical protein, conserved | 3 | 3 | 3 | 3 | 3 | 3 | 3 | 8 |
| 12 | TGME49_500294.P83 | protein BRAWNIN, putative | 3 | 3 | 3 | 4 | 3 | 3 | 3 | 8 |
| 12 | TGME49_500295.P84 | Toxoplasma gondii family B protein | 4 | 3 | 3 | 3 | 3 | 3 | 3 | 8 |
| 12 | TGME49_500299.P88 | hypothetical protein | 3 | 3 | 3 | 3 | 3 | 3 | 3 | 8 |
| 12 | TGME49_500311.P101 | hypothetical protein, conserved | 3 | 3 | 3 | 3 | 3 | 3 | 3 | 8 |
| 12 | TGME49_500312.P102 | dense granule protein GRA41 | 3 | 3 | 3 | 3 | 3 | 3 | 3 | 8 |
| 12 | TGME49_500313.P103 | hypothetical protein, conserved | 3 | 3 | 3 | 3 | 3 | 3 | 3 | 8 |
| 12 | TGME49_500316.P106 | hypothetical protein, conserved | 3 | 3 | 3 | 3 | 3 | 3 | 3 | 8 |
| 12 | TGME49_500319.P109 | hypothetical protein, conserved | 3 | 3 | 3 | 3 | 3 | 3 | 3 | 8 |
| 12 | TGME49_500320.P110 | hypothetical protein, conserved | 3 | 3 | 3 | 3 | 3 | 3 | 3 | 8 |
| 12 | TGME49_500324.P114 | hypothetical protein | 3 | 3 | 3 | 3 | 3 | 3 | 3 | 8 |
| 12 | TGME49_500326.P116 | hypothetical protein, conserved | 3 | 3 | 7 | 4 | 3 | 3 | 4 | 8 |
| 12 | TGME49_500332.P122 | hypothetical protein, conserved | 3 | 3 | 6 | 3 | 3 | 3 | 3 | 8 |
| 12 | TGME49_500334.P124 | hypothetical protein, conserved | 3 | 3 | 3 | 3 | 3 | 3 | 3 | 8 |
| 12 | TGME49_500336.P126 | hypothetical protein, conserved | 3 | 5 | 3 | 4 | 3 | 3 | 3 | 8 |
| 22 | TGME49_500337.P127 | ubiquitin-like protein, putative | 3 | 6 | 5 | 3 | 4 | 4 | 3 | 8 |
| 12 | TGME49_500346.P136 | hypothetical protein, conserved | 3 | 3 | 4 | 3 | 3 | 3 | 3 | 8 |
| 22 | TGME49_500347.P137 | hypothetical protein, conserved | 3 | 6 | 3 | 4 | 3 | 3 | 3 | 8 |
| 12 | TGME49_500350.P140 | hypothetical protein, conserved | 3 | 3 | 3 | 3 | 3 | 3 | 3 | 8 |
| 12 | TGME49_500351.P141 | hypothetical protein | 3 | 3 | 3 | 3 | 3 | 3 | 3 | 8 |
| 12 | TGME49_500357.P147 | hypothetical protein, conserved | 3 | 3 | 5 | 3 | 3 | 3 | 3 | 8 |
| 12 | TGME49_500359.P258 | hypothetical protein, conserved | 3 | 3 | 3 | 3 | 3 | 3 | 3 | 8 |
| 12 | TGME49_500360.P259 | regulator of chromosome condensation, putative | 3 | 3 | 3 | 6 | 3 | 5 | 4 | 8 |
| 12 | TGME49_500368.P267 | hypothetical protein, conserved | 3 | 3 | 4 | 3 | 3 | 3 | 3 | 8 |
| 12 | TGME49_500370.P269 | histidine phosphatase, putative | 3 | 3 | 6 | 3 | 3 | 3 | 3 | 8 |
| 12 | TGME49_500371.P270 | hypothetical protein | 3 | 3 | 3 | 3 | 3 | 3 | 3 | 8 |
| 12 | TGME49_500375.P274 | rhoptry kinase family protein | 3 | 3 | 3 | 3 | 3 | 3 | 3 | 8 |
| 22 | TGME49_500379.P278 | WD repeat-containing protein | 3 | 5 | 7 | 3 | 4 | 3 | 3 | 8 |
| 22 | TGME49_500381.P280 | EF-hand domain-containing protein | 3 | 9 | 3 | 8 | 4 | 4 | 4 | 8 |
| 12 | TGME49_500382.P281 | hypothetical protein, conserved | 3 | 3 | 3 | 3 | 3 | 4 | 4 | 8 |
| 12 | TGME49_500386.P285 | hypothetical protein, conserved | 3 | 3 | 4 | 3 | 3 | 3 | 3 | 8 |
| 12 | TGME49_500388.P287 | hypothetical protein, conserved | 3 | 3 | 4 | 3 | 3 | 3 | 3 | 8 |
| 12 | TGME49_500395.P294 | hypothetical protein, conserved | 3 | 3 | 7 | 3 | 3 | 4 | 4 | 8 |
| 12 | TGME49_500396.P295 | hypothetical protein | 3 | 3 | 3 | 3 | 3 | 3 | 3 | 8 |
| 12 | TGME49_500397.P296 | MoaD/ThiS family protein | 3 | 3 | 4 | 3 | 3 | 3 | 3 | 8 |
| 12 | TGME49_500399.P298 | hypothetical protein, conserved | 3 | 3 | 4 | 4 | 3 | 3 | 3 | 8 |
| 22 | TGME49_500402.P301 | hypothetical protein, conserved | 3 | 6 | 3 | 3 | 3 | 3 | 3 | 8 |
| 12 | TGME49_500407.P306 | hypothetical protein | 3 | 3 | 3 | 3 | 3 | 3 | 3 | 8 |
| 12 | TGME49_500412.P311 | hypothetical protein, conserved | 3 | 3 | 4 | 3 | 3 | 3 | 3 | 8 |
| 22 | TGME49_500417.P316 | hypothetical protein, conserved | 3 | 9 | 4 | 3 | 3 | 3 | 3 | 8 |
| 12 | TGME49_500420.P319 | hypothetical protein, conserved | 3 | 3 | 7 | 3 | 3 | 3 | 3 | 8 |
| 12 | TGME49_500425.P324 | hypothetical protein, conserved | 3 | 3 | 3 | 3 | 3 | 3 | 3 | 8 |
| 12 | TGME49_500430.P329 | hypothetical protein, conserved | 3 | 3 | 3 | 3 | 3 | 4 | 3 | 8 |
| 12 | TGME49_500431.P330 | SHIPPO repeat domain-containing protein | 3 | 3 | 3 | 4 | 3 | 3 | 3 | 8 |
| 12 | TGME49_500432.P331 | hypothetical protein, conserved | 3 | 3 | 3 | 3 | 3 | 3 | 3 | 8 |
| 12 | TGME49_500436.P335 | EF-hand calcium-binding domain-containing protein 10, putative | 3 | 3 | 3 | 3 | 4 | 4 | 3 | 8 |
| 12 | TGME49_500437.P336 | CASC1 domain-containing protein | 3 | 3 | 6 | 3 | 3 | 3 | 3 | 8 |
